# Zika viral proteome analysis reveals an epitope cluster within NS3 helicase as a potential vaccine candidate: an *in silico* study

**DOI:** 10.1101/2020.04.24.059543

**Authors:** Md Tangigul Haque, Md Nur Ahad Shah, Shatabdi Paul, Md Kawsar Khan, Payal Barua

**Author notes:** Corresponding Author: Payal Barua.

## Abstract

Zika virus (ZIKV), a mosquito-borne flavivirus, is now an emerging global public health concern. Currently, the pathogenicity, genetic diversity and the consequences of ZIKV infection are little known but a protective vaccine against ZIKV is an urgency. In this study, we have taken an immunoinformatics based approach to predict epitope cluster region in the whole proteome (3423 amino acids) of ZIKV. We have operated a range of bioinformatics algorithms to determine the epitopes of CD8+ cytotoxic T-cell (CTL), CD4+ helper T-cell (THL) and B cell. We have predicted an epitope cluster of 23 contiguous amino acids (region 1989-2011, WLEARMLLDNIYLQDGLIASLYR) residing on the protein NS3 helicase in ZIKV proteome. This epitope cluster contains fourteen CD4+ (THL) epitopes and six CD8+ (CTL) epitopes. The cluster region predicted to provide 93.86% population coverage worldwide. Finally, we have validated the epitopes by analysing their binding efficiency (binding energy within −4.7 to −6.9 kcal/mol) with specific HLA alleles. Based on our immunoinformatics analysis, we propose the peptide WLEARMLLDNIYLQDGLIASLYR as a new peptide vaccine candidate against Zika virus for further validation.

## 1. Introduction

Zika virus (ZIKV), an arthropod-borne virus (arbovirus) of the family Flaviviridae, is one of the most alarming infectious agents of the present time. After the first identification of the virus in Uganda in 1947, outbreaks of ZIKV infection were recorded in Yap Island, Micronesia, French Polynesia, North Pacific Islands and Brazil till 2010 [1–3]. Currently, the virus is known to occur in 84 countries including parts of South America, some parts of North America and Asia. The vectors for transmission of the ZIKV encompass a number of hosts including arboreal mosquitos that transmits the virus from the non-human primates, mosquitoes of *Aedes* genus (e.g. *A. aegypti, A. africanus, A. albopictus, A. apicoargenteus, A. furcifer, A. luteocephalus, A. opok, and A. vittatus*), and human for primary amplification [4]. ZIKV can spark outbreaks of infection around the world, as it can also be transmitted via placental, perinatal, sexual, and blood-borne routes in humans [5]. The primary clinical features of ZIKV infection in adults range from asymptomatic or mild to acute febrile illness similar to Dengue infection [2, 6, 7] as well as Guillain-Barré syndrome [8]. On the other hand, congenital infections of ZIKV often cause brain abnormalities and microcephaly in developing fetus [9, 10]. Despite that, very little or no protective measures are available for this fatal virus infection worldwide.

ZIKV has a single-stranded positive sense RNA genome of approximately 10.8 kb (10,807-nt) length with a single open reading frame (ORF) of 10,272-nt flanked by 5′ noncoding region (NCR) of 107-nt and the 3′ NCR region of 428-nt [11]. The ORF encodes a single polyprotein of 3,423 amino acids. The genome polyprotein of ZIKV is the mother of other viral proteins. The polyprotein is cleaved by viral and cellular proteases into three structural proteins located at the N terminus: capsid (C – 122 aa), precursor of membrane (prM – 168 aa), and envelope (E – 504 aa) protein and seven non-structural (NS) proteins essential for viral genomic RNA replication located at the C terminus: NS1 (352 aa), NS2A (226 aa), NS2B (130 aa), NS3 (617 aa), NS4A (150 aa), NS4B (251 aa), NS5 (903 aa) [12]. The structural proteins of the virion aid its entry via cellular receptor as well as its encapsidation [13]. The NS proteins primarily mediate the replication of the viral genome, cleavage of the polyproteins and host immune evasion [14]. Nevertheless, immunogenic epitopes are present in all three types of ZIKV proteins, E, prM and some NS [4, 15]. Infection of ZIKV in human elicits inflammatory responses which later induces both humoral and cell-mediated immune response [4]. Antibody and cell-mediated immune responses are initiated following infection with the release of IgM antibodies in about 9.1 days followed by IgG antibodies after 2–3 days [4, 16]. Because of the dynamic evolution of host immuneevasion mechanism of the flaviviruses, viral clearance or prevention of disease progression is difficult [17].

Along with the African strains from the 1950s, all ZIKV strains are limited to a single serotype, unlike DENV, making the ZIKV vaccine development process highly effective that is capable of inducing an immune response against all strains of this virus [18]. A modified mRNA vaccine showed promising immune responses against ZIKV in both mice, and *Rhesus macaques* model [19] and a live attenuated ZIKV vaccine succeeded to stop maternal transmission of this virus in mice model [20]. These types of vaccines are in various stages of development; some are in human trials, and some of these have not yet been tested in pregnancy [21, 22]. The main limitation of those vaccine candidates is the obscureness of whether any candidates can protect the fetus or not [21]. As an alternative to live attenuated or modified mRNA vaccine, some epitope-based peptide vaccines were proposed using in silico methods [23–26]. Epitope-based vaccine design determines the most immunogenic epitopes of a protein with the aid of computational approach. The in silico methods can predict B and T cell-specific epitopes and estimate the population coverage, leading to a possible prediction of activity of the vaccine in vivo. Till date, most of the predicted epitope vaccines are still in the pipeline of in vitro and in vivo testing with the goal of long-term protective and robust immune response in humans [22].

The immune epitope database (IEDB) hosts a plethora of sophisticated tools to design B and T cell-specific epitopes. IEDB tools have been successfully used previously to predict suitable vaccine candidate against many viruses, such as influenza, dengue, respiratory syncytial and human immunodeficiency virus [27–30] as well as for bacteria, such as *Mycobacterium tuberculosis*, *Streptococcus pneumoniae*, *Haemophilus influenzae*, *Helicobacter pylori* and *Mycobacterium bovis* [31–34]. Efficacy of some of these vaccine candidates has also been tested successfully in vivo and in vitro [31, 35, 36]. Some of the studies were conducted on specific proteins (the envelope glycoprotein and a couple of non-structural proteins) of the virus rather than the whole proteome [23, 37], and in another case, the study was done to identify individual epitopes across the proteome rather than epitope rich cluster regions [38]. Although the whole genome polyprotein of the ZIKV has the potential for residing continuous and discontinuous epitope regions for both B and T cells, this protein remains to be investigated by the immunoinformatic method. In this study, we aimed to map the multiple epitope cluster regions by screening the whole proteome of ZIKV to construct a potential epitope vaccine candidate. For this, we scanned the genome polyprotein of ZIKV to predict the most immunogenic and epitope dense region in ZIKV. We applied a range of bioinformatics tools to predict a cluster of epitopes with high population coverage throughout the world and with no cross-reactivity with human self-peptides.

## 2. Materials and Methods

### 2.1 Retrieving the protein

The protein sequences of Zika Virus (strain Mr 766) (ZIKV) (TaxID: 64320) genome polyprotein were retrieved from UniProt database (www.uniprot.org). Only the whole genome polyprotein sequences consisting of 3423 amino acids were taken for this study from different geographical location submitted from the year 1947 to 2016.

### 2.2 Multiple sequence alignment

The retrieved protein sequences of ZIKV were analyzed for conserved sequences by generating multiple sequence alignment. The alignment was done by utilizing the BLOSUM matrix algorithm from MEGA (version 6.06) using ClustalW tools [39]. A weblogo was generated from the retrieved sequences using the Weblogo tool (version 3.0) (weblogo.berkeley.edu) [40].

### 2.3 Construction of Phylogenetic tree with protein

A phylogenetic tree was constructed to calculate the evolutionary distances by using MEGA software (version 7.0.26). The evolutionary history was inferred using the neighbor-joining method with the number of bootstrap replication at 1000 which was used to denote the confidence level for the phylogenetic tree [41].

### 2.4 Calculation of non-synonymous – synonymous substitution ratio

The nucleotide sequences, corresponding to the previously retrieved Zika virus genome polyproteins, were collected from The European Nucleotide Archive (ENA) (https://www.ebi.ac.uk/). An average value of non-synonymous and synonymous substitution was calculated using MEGA (version 6.06) software by applying Nei-Gojobori (Jukes-Cantor) method [42]. Moreover, the non-synonymous – synonymous substitution ratio (Ka/Ks) was calculated to infer the direction and magnitude of natural selection acting on protein encoding genes.

### 2.5 Identification of T cell epitope

Epitope prediction tools provided in The Immune Epitope Database (IEDB) (http://www.iedb.org) was used to infer the epitopes capable of inducing an antigenic response, namely helper T cell (THL) and cytotoxic T cell (CTL) specific epitopes. The NN-align method (Nielsen and Lund 2009) was applied to predict THL epitopes. This artificial neural network based method can predict the binding affinity of the putative epitope with the MHC class II binding core, represented by the inverse function of the IC50 value. An IC50 value of less than 500 nM indicates higher affinity between the epitope and HLA. The epitopes with an IC50 value lower than 500 nM were selected for further analysis. For the prediction of CTL epitopes, ‘Proteasomal cleavage/Tap transport/MHC class I combined predictor’ (http://tools.immuneepitope.org/processing/) tool was used. The Artificial Neural Network (ANN) prediction method [43] was applied to predict successful CTL epitopes from a given protein sequence based on their ability to bind with MHC class I molecules as well as to be processed by both the proteasomes and transporters associated with antigen presentation (TAP). A cut-off value for MHC score −2.7, equivalent to the IC50 value of 500 nM, was used [28, 31, 44, 45]. The threshold value 1.00 for proteasomal cleavage and 1.14 for TAP binding was selected, which corresponds to an expected specificity rate of 76% [46].

### 2.6 Epitope conservation analysis

The ‘Conservation Across Antigens’ tool from IEDB (http://tools.iedb.org/conservancy/) was employed to inspect the conservancy of the predicted epitopes. The epitopes with 100% conservancy across all the variants were selected for further investigation.

### 2.7 Position specific amino acid antigenicity and epitope clustering

The IC50 value which is the inhibitory concentration of the amino acids of the selected epitopes of ZIKV proteome were analysed using an in-house algorithm. The presence of multiple overlapping epitopes resulted in a few IC50 values for each amino acid. In this case, the lowest IC50 value was taken for further analysis. Similarly, to predict the number of binding alleles of each amino acid, only the frequency of the unique alleles was counted. The redundant alleles were ignored. K-means clustering method was applied to classify the ZIKV proteome into epitope dense, moderate and poor regions based on their IC50 values and unique allele count [47]. The location of the cluster region within the ZIKV proteome was further investigated using the NCBI BlastP program.

### 2.8 B cell antigenicity prediction of the epitope cluster

In the immune system, B cell epitope is the sections of a molecule that is recognized by antibodies. Till date, the hidden Markov model has proven to be one of the most reliable methods for B cell epitope prediction. The Bepipred linear epitope prediction method available at the IEDB was employed to calculate the B cell antigenicity of our predicted epitope cluster of ZIKV (http://tools.iedb.org/bcell/). This method is a combination of the hidden Markov model and the propensity scale method delivering significantly better performance [48].

### 2.9 Population coverage prediction

The population coverage of the selected epitopes was calculated using the Population Coverage Analysis tool from IEDB [49]. HLA allele genotypic frequencies obtained from the Allele Frequency Net Database calculates the fraction of individuals predicted to respond to a given set of epitopes with known HLA restrictions [50]. During the analysis HLAs unavailable in the database were ignored.

### 2.10 Determination of cross-reacting human self-peptides in the predicted epitope

The presence of human peptides, analogous to the predicted multi-epitope cluster, was determined using the NCBI BlastP program (www.ncbi.nlm.nih.gov/BLAST/). The predicted epitope sequences were compared with the non-redundant protein sequences database, restricted to human (TaxID: 9606). The following parameters were set for prediction: word size of two, expected threshold value of 30,000, “no adjustment” setting for composition-based statistics and PAM30 as the matrix of choice [51]. Hits involving fewer than four amino acids match were ignored considering the likelihood of such matches being random.

### 2.11 3D structure prediction of epitopes and HLA molecules for analyzing their interaction using molecular docking

The PEP-FOLD peptide structure prediction server (bioserv.rpbs.univ-paris-diderot.fr/services/PEP-FOLD/) was used to design the three-dimensional structures of the predicted epitopes. The 3D structure of HLA-DRB1*04:05, HLA-DRB4*01:01, HLA-DRB1*01:01, HLA-DRB1*03:01, HLA-DRB1*04:04, HLA-DRB1*09:01, HLA-DRB1*15:01, HLA-A*30:02, HLA-A*02:06 and HLA-A*29:02 allele was predicted using Phyre2 protein prediction server (Protein Homology/analogy Recognition Engine V 2.0). The coding DNA sequences of the respective HLA molecules were obtained from the NCBI database (GenBank ID: CCM44108.1, SAL89153.1, BAN59849.1, CBM42731.1, CZQ50726.1, ADC42109.1, AEJ87208.1, SFW97183.1, AEQ 26027 and CDH35265 respectively).

The AutoDOCK tool from the MGL software package (version 1.5.6) was employed for docking purpose [52, 53]. For visualization, PyMOL (Version 1.4.1) was used to observe the docking interactions of the epitope structures with their corresponding HLAs. The best output with the lowest binding energy was selected in this case.

## 3. Results

### 3.1 Evolutionary divergence in ZIKV genome polyprotein

A total of 48 sequences were retrieved (Supplementary file 1) from different geographical locations (Supplementary file 2). A weblogo was generated using the multiple sequence alignment (Supplementary file 2). A phylogenetic tree constructed from the aligned sequences showed evolutionary relationships and divergence of ZIKV polyproteins over the period and geographical location (figure 1).

**Figure 01.**
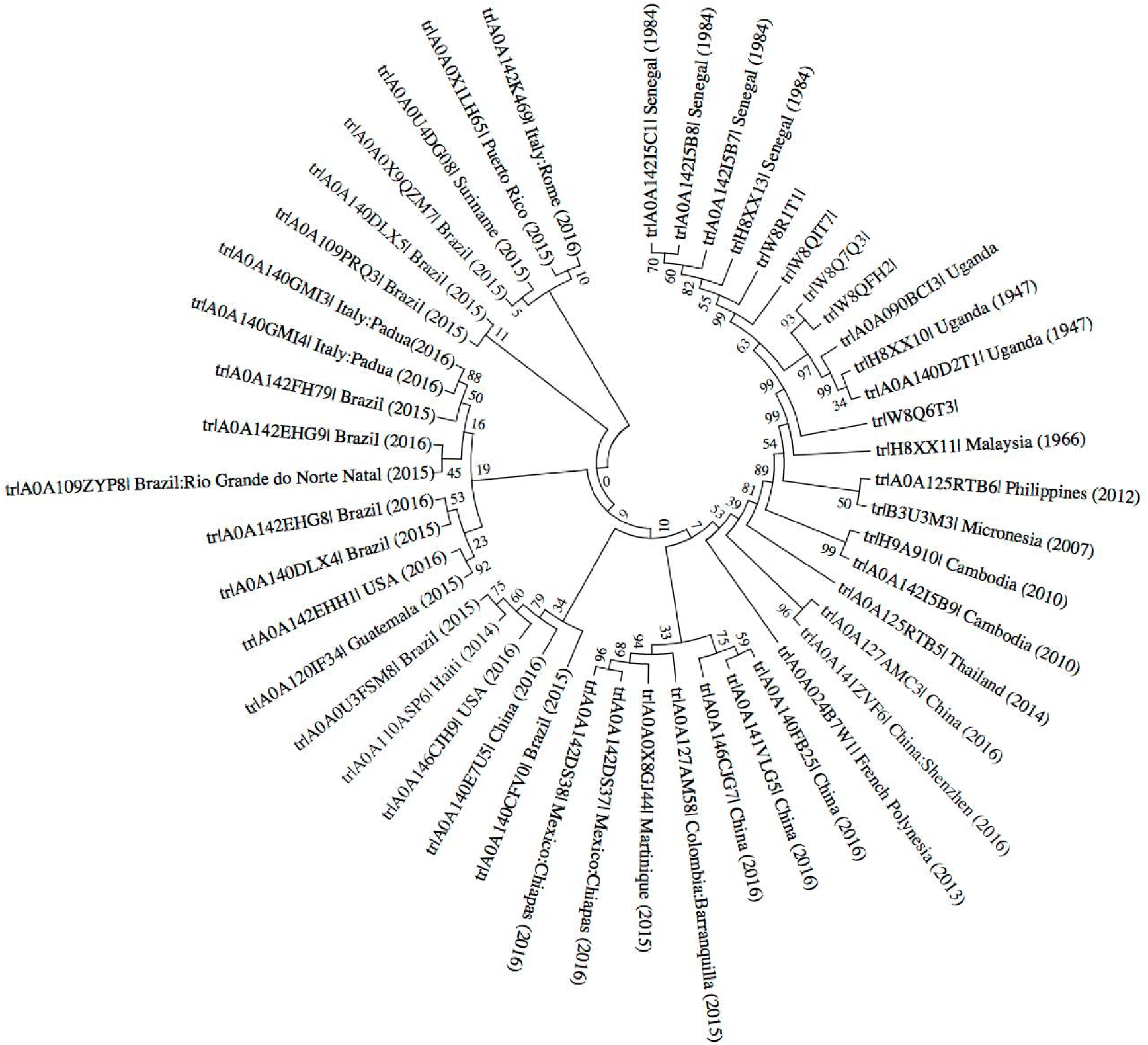
Phylogram of Zika virus genome polyprotein

### 3.2 Identification of the evolutionarily conserved region in ZIKV genome polyprotein

To infer the direction and force of natural selection acting on the ZIKV polyprotein, we calculated the non-synonymous (Ka) and synonymous (Ks) mutations rates. For this purpose, the DNA sequences of each of the corresponding polyprotein were retrieved from European Nucleotide Archive (ENA), and the mutation rates were calculated using MEGA. The average values for non-synonymous and synonymous mutations were 0.065 and 0.021 respectively for the polyprotein. The Ka/Ks ratio found for the polyprotein was 3.095, implying that the protein follows the positive or Darwinian selection. A Ka/Ks ratio below one indicates purifying selection, while a ratio of one means neutral selection. Here, the Ka/Ks ratio is significantly higher than one, which implies that the mutations in this protein are advantageous to the organism over the course of evolution.

### 3.3 Identification of CD8+ (CTL) cell-specific epitopes

Epitopes specific to CTL were identified using a few tools available at IEDB. In this case, the sequence with accession id A0A125RTB5_ZIKV was used. A total of 51 nine-mer peptide sequences (Supplementary file 3) were found using Artificial Neural Network (ANN) methods which satisfied all the desired criteria. These sequences were further inspected for their conservancy using the ‘Conservation Across Antigens’ tool from IEDB. The conservancy analysis identified 18 epitopes which are conserved across all the organisms (Table 01).

**Table 01.**
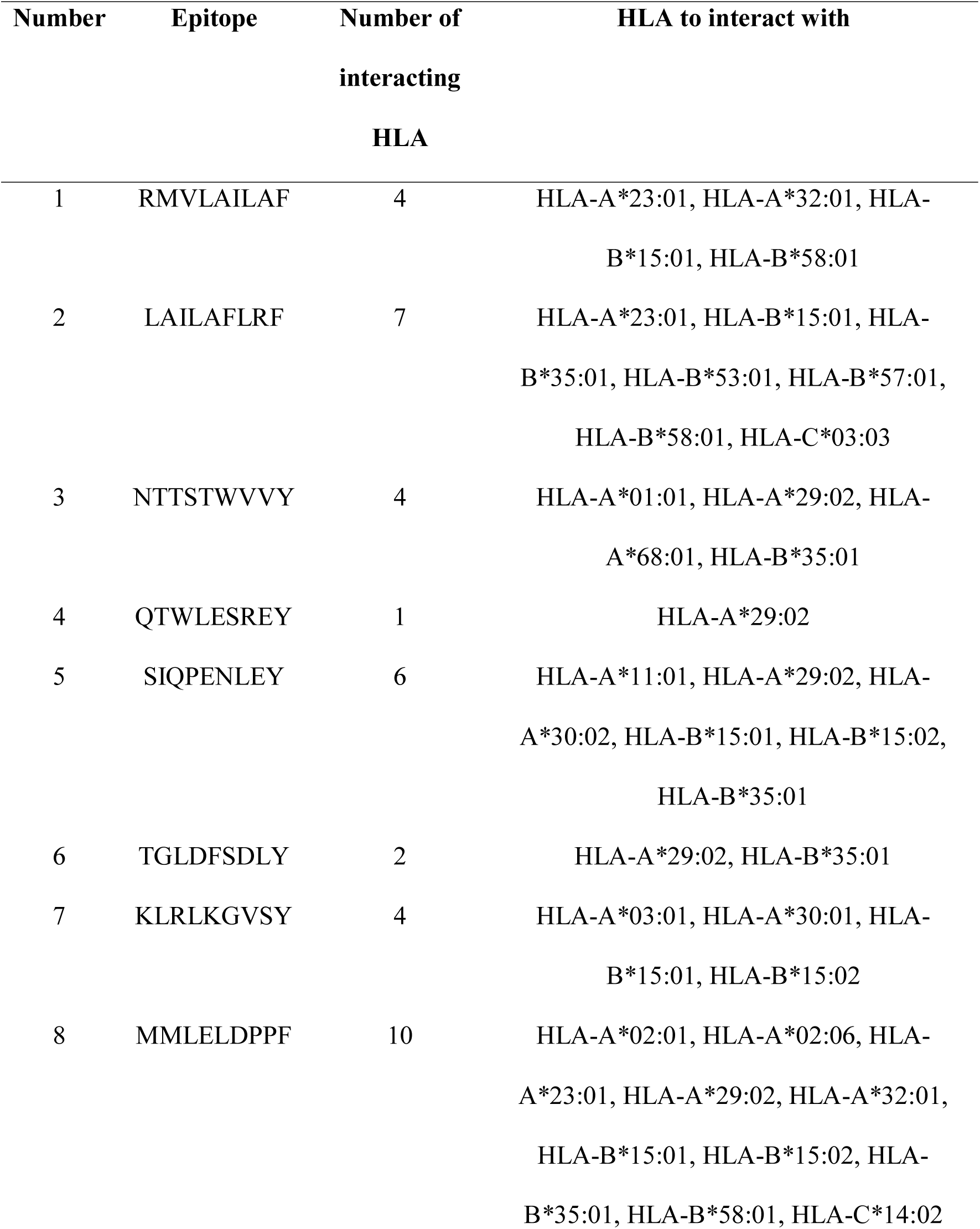
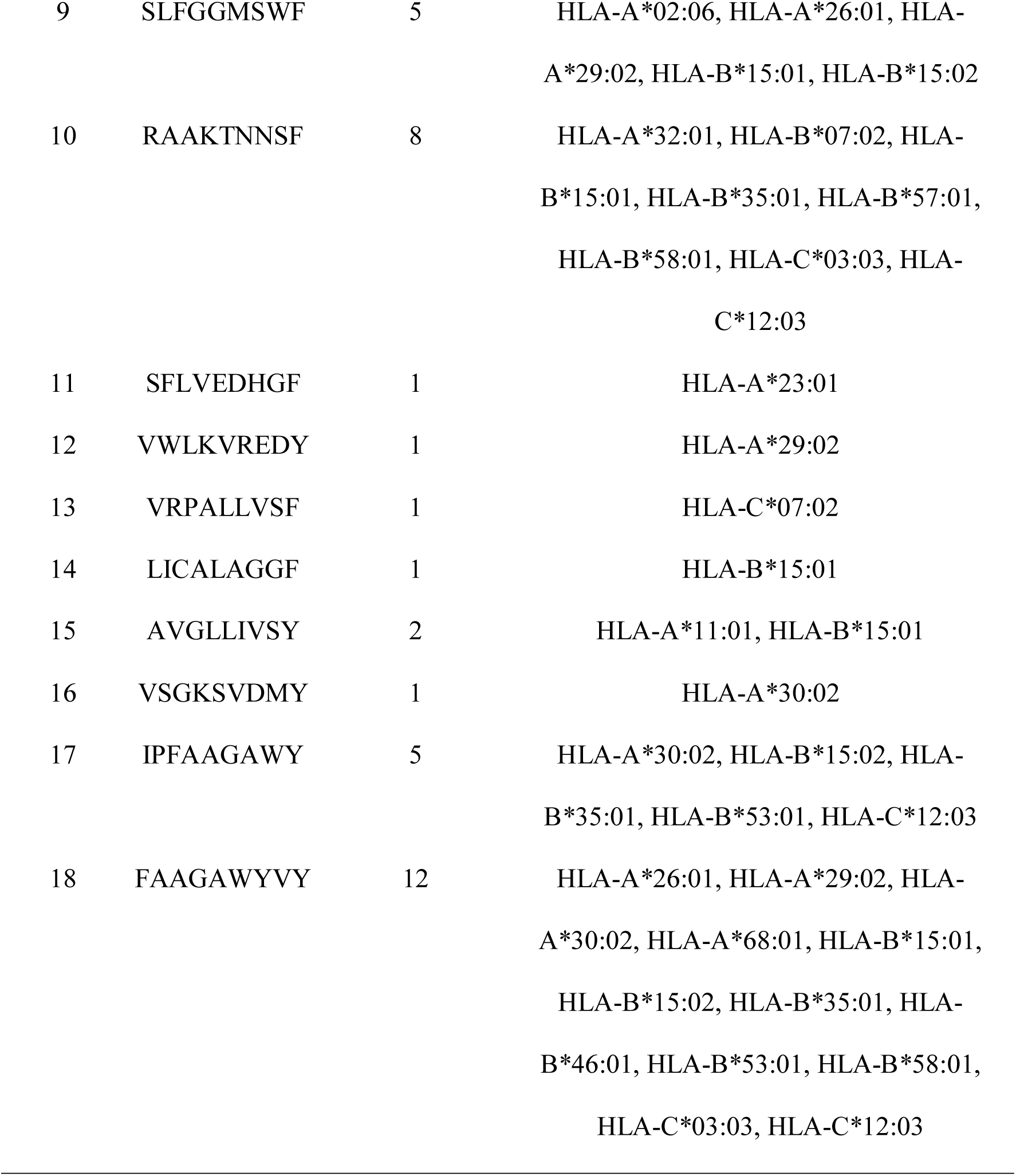
Selected CTL epitopes with their corresponding HLAs.

Among the predicted epitopes, the epitope FAAGAWYVY was found to have the highest number of binding HLAs which is twelve (HLA-B*35:01, HLA-C*03:03, HLA-A*29:02, HLA-C*12:03, HLA-B*15:01, HLA-A*30:02, HLA-A*26:01, HLA-B*15:02, HLA-B*53:01, HLA-A*68:01, HLA-B*46:01, HLA-B*58:01). On the other hand, six predicted epitopes (QTWLESREY, SFLVEDHGF, VWLKVREDY, VRPALLVSF, LICALAGGF, VSGKSVDMY) were found to be binding with only one HLA. In contrast, HLA-B*15:01 was found to be binding with the maximum number of predicted epitopes (10 epitopes), whereas, HLA-A*29:02 and HLA-B*35:01 were found to be binding with eight different epitopes each. Again, 15 different HLAs were found to be unique, binding with only one epitope each.

The positional distributions of HLA allele frequency and IC50 values for each amino acid position were obtained from the data using an in-house algorithm (Supplementary file 2) and were plotted accordingly (figure 02). In figure 2A, two peaks with the amino acid sequences 1490-AGAWYVYVK-1498 and 2541-ALEFYSY-2547 showed the highest allele counts of twelve. In figure 2B the amino acids were plotted against their respective IC50 values which correspond to their inhibitory concentrations. Here, IC50 values over 500 nM were ignored as amino acids with higher IC50 scores possess very low affinity towards the MHC molecules.

**Figure 02.**
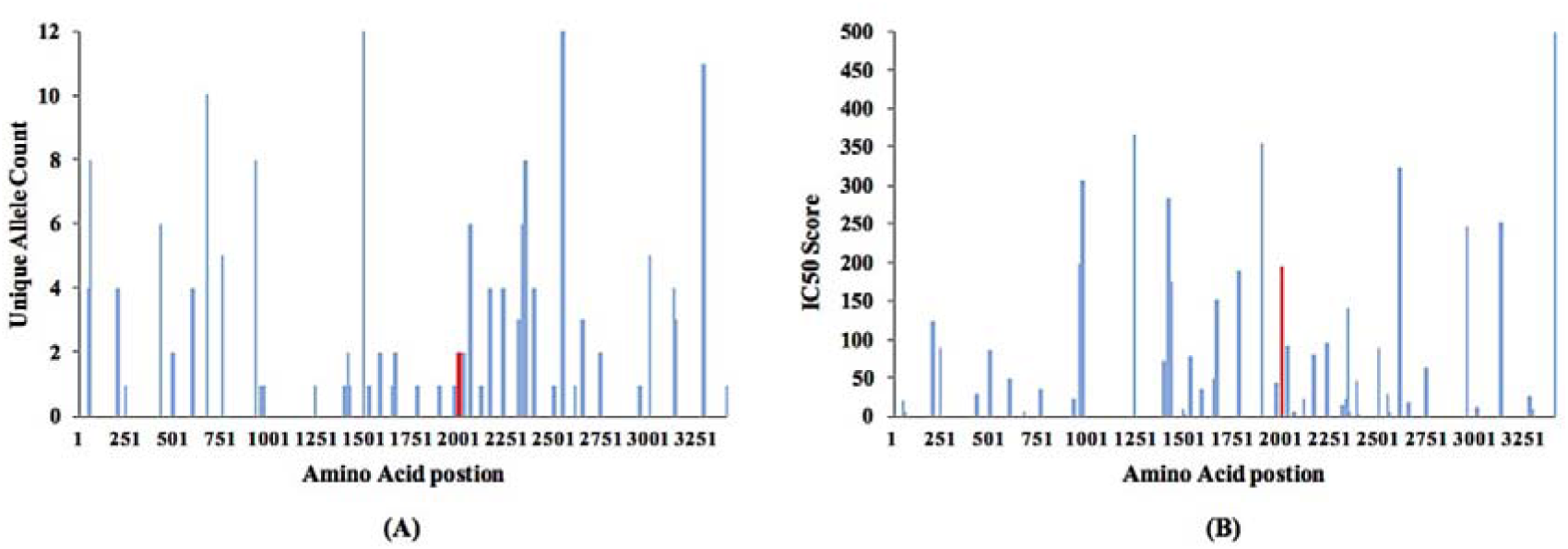
Position specific (A) Unique Allele Count (B) IC50 values of each amino acid for CTL cell epitopes. The lower the IC50 value, the higher the affinity towards MHC. The red region corresponds to the predicted epitope cluster. The predictions were derived from the algorithm ANN.

### 3.4 Identification of CD4+ (THL) cell-specific epitopes and determination of epitope cluster region

THL epitopes were predicted by the binding affinities between the putative epitope regions and HLA Class II molecules using the NN align method (Supplementary file 3). These epitopes were further tested for their conservancy. Using the predicted THL specific response data, positional distributions of the frequency of corresponding HLA alleles and IC50 values for each amino acid position were obtained and plotted respectively (figure 3A and 3B). IC50 values over 500 nM were ignored.

**Figure 03.**
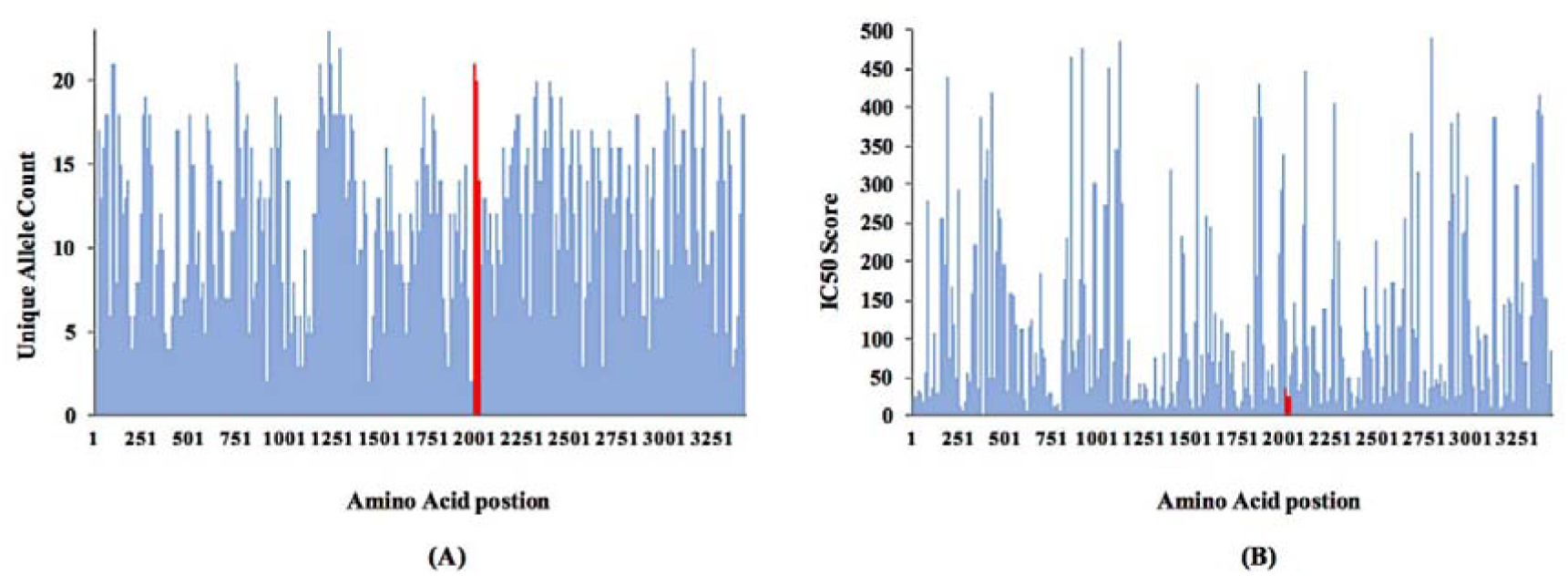
Position specific (A) Unique Allele Count (B) IC50 values of each amino acid for THL cell epitopes. The lower the IC50 value, the higher the affinity towards MHC. The red region corresponds to the predicted epitope cluster. The predictions were derived from the algorithm NN-align.

The CD4+ (THL) cell-specific epitope were further analyzed using the K-means clustering method to find the epitope cluster region on ZIKV polyprotein. Clustering allows amino acids to be assigned regarding their probability to become a constituent of an immunogenic epitope, based on two non-discrete variables: position-specific (1) unique allele counts and (2) IC50 scores (Supplementary file 3). A cluster of 23 conserved continuous amino acids was identified. The identified epitope cluster spanned from position 1989 to 2011 region with multiple overlapping THL and CTL epitopes that possess low IC50 values and high allele counts (Figure 4A). This sequence of the cluster region (1989-WLEARMLLDNIYLQDGLIASLYR-2011) was designated and used for all ensuing analyses. Using the NCBI BlastP program, we have identified the viral protein NS3 Helicase (PDB ID 5VI7), that contains the cluster sequence spanning over its region 313-335 (Figure 4B).

**Figure 04.**
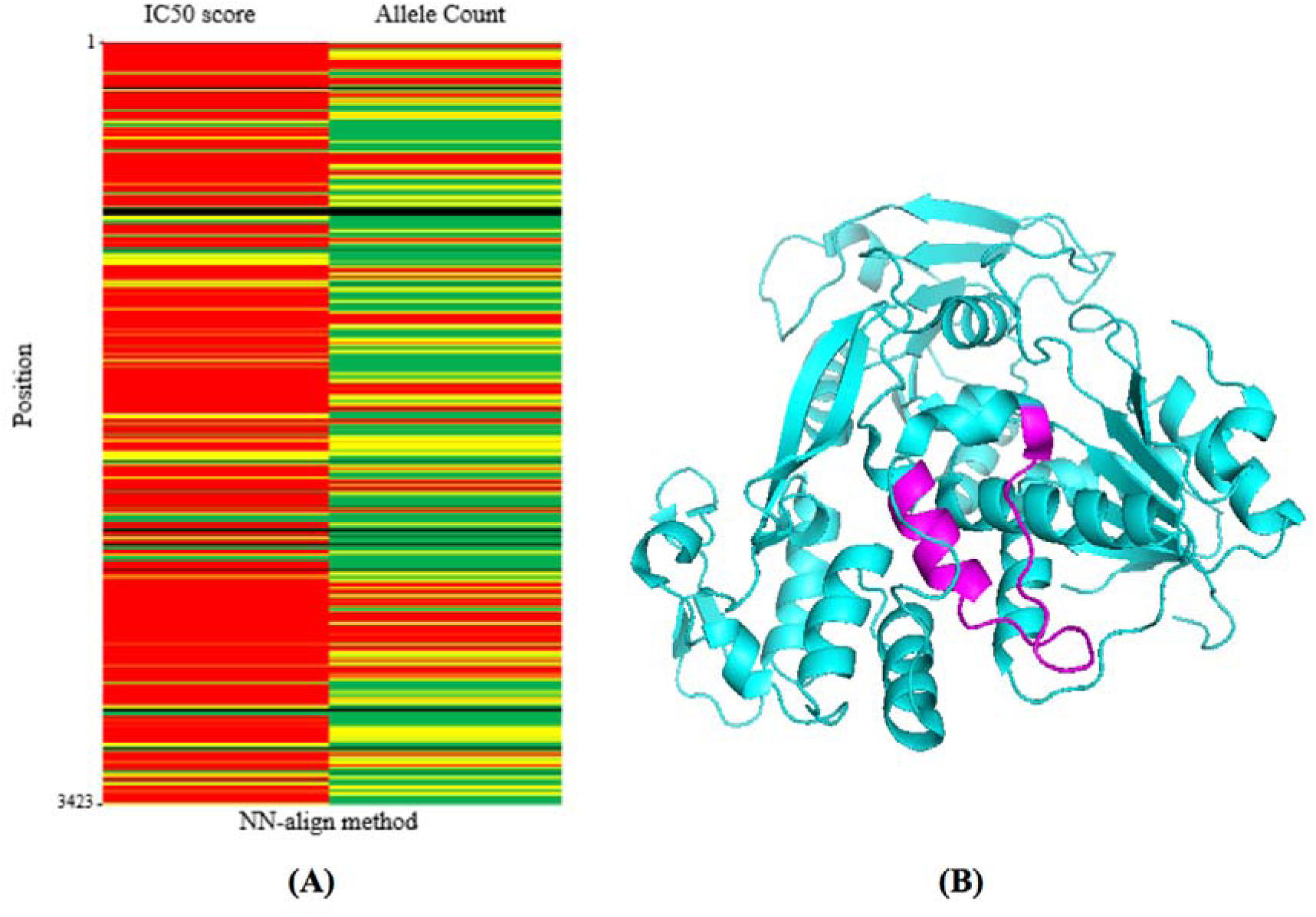
(A) Epitope cluster predicted by k-means cluster using NN-align method where each amino acid position represents as an epitope dense, average and poor region. The colour red denotes high allele count with low IC50 values, yellow denotes average both for allele count and IC50 value, green denotes low allele count with high IC50 values. (B) Crystal structure of the NS3 Helicase protein. The predicted epitope cluster region is colored in magenta.

### 3.5 T cell and B cell antigenicity of the predicted epitope cluster region

The conserved epitope cluster region was analysed for its CTL and THL specific response as well as B cell antigenicity. A total of six epitopes were found to elicit CTL mediated response (Table 02). Three epitopes – RMLLDNIYL, MLLDNIYLQ, and LQDGLIASL, were found to be the most immunogenic, binding with three different HLAs. Among the HLAs, HLA-A*02:01 and HLA-A*02:06 were the most abundant, each of them showing the affinity towards four different epitopes.

**Table 02.** Selected CD8+ (CTL) epitope within the cluster region with their corresponding HLAs.

| Epitope | Number of<br>interacting HLA | Interact with corresponding Allele |
| --- | --- | --- |
| ARMLLDNIY | 2 | HLA-A*30:02, HLA-B*27:05 |
| RMLLDNIYL | 3 | HLA-A*02:01, HLA-A*02:06, HLA-C*14:02 |
| MLLDNIYLQ | 3 | HLA-A*02:01, HLA-A*02:06, HLA-A*29:02 |
| YLQDGLIAS | 2 | HLA-A*02:01, HLA-A*02:06 |
| LQDGLIASL | 3 | HLA-A*02:01, HLA-A*02:06, HLA-B*39:01 |
| QDGLIASLY | 1 | HLA-A*29:02 |

In contrast to CTL specific response, the THL specific response of the cluster region was more evident resulting in a total of fourteen epitopes (Table 03). Among them, the epitope WLEARMLLD was the most effective one binding with 13 different HLA molecules. The epitopes MLLDNIYLQ and IYLQDGLIA showed the affinity towards nine different HLAs each. On the other hand, LDNIYLQDG and QDGLIASLY epitopes were the least immunogenic binding with only one HLA. HLA-DPA1*02:01 and HLA-DRB1*04:05 were the most evident HLAs that bind to five different epitopes.

**Table 03.**
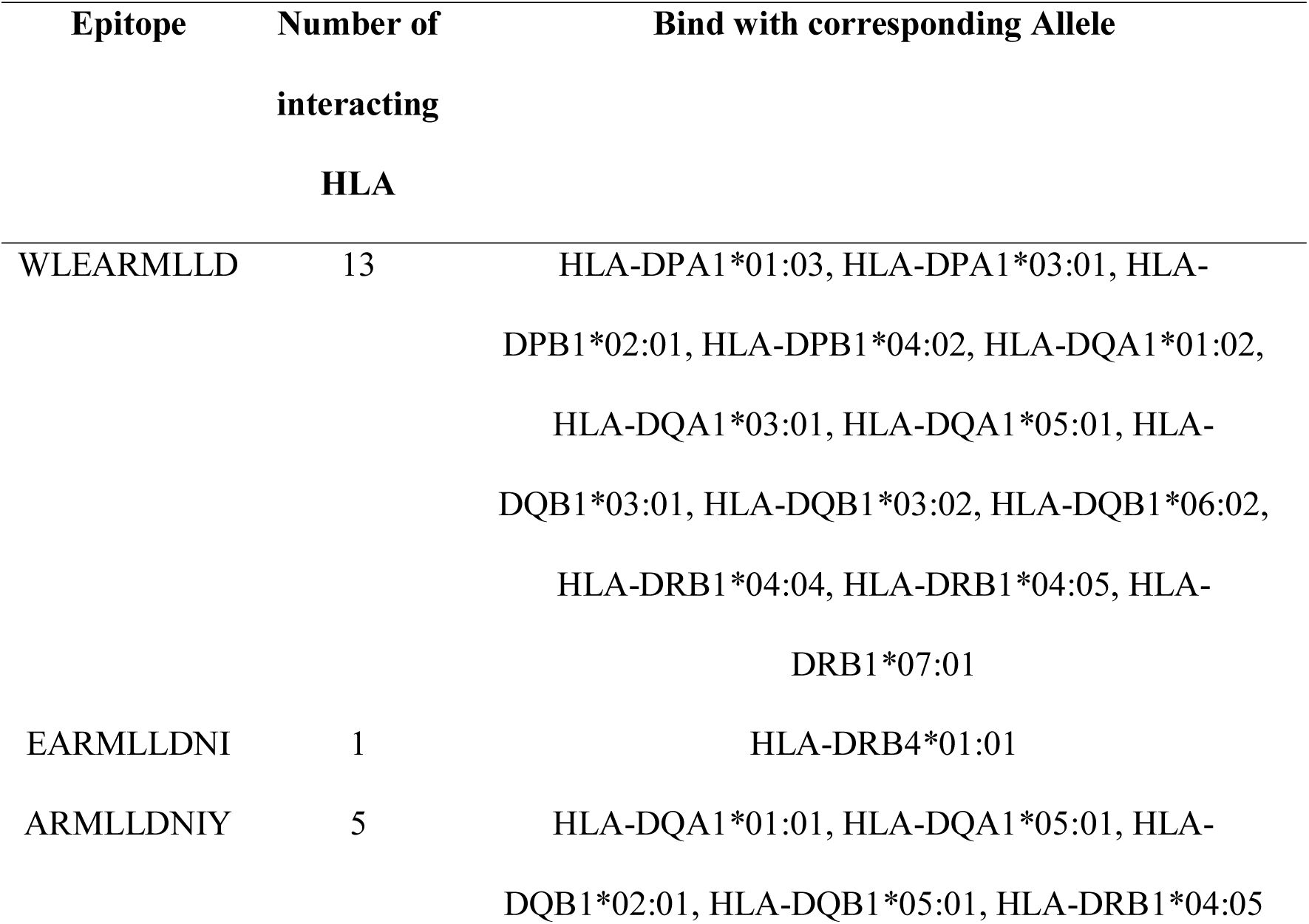
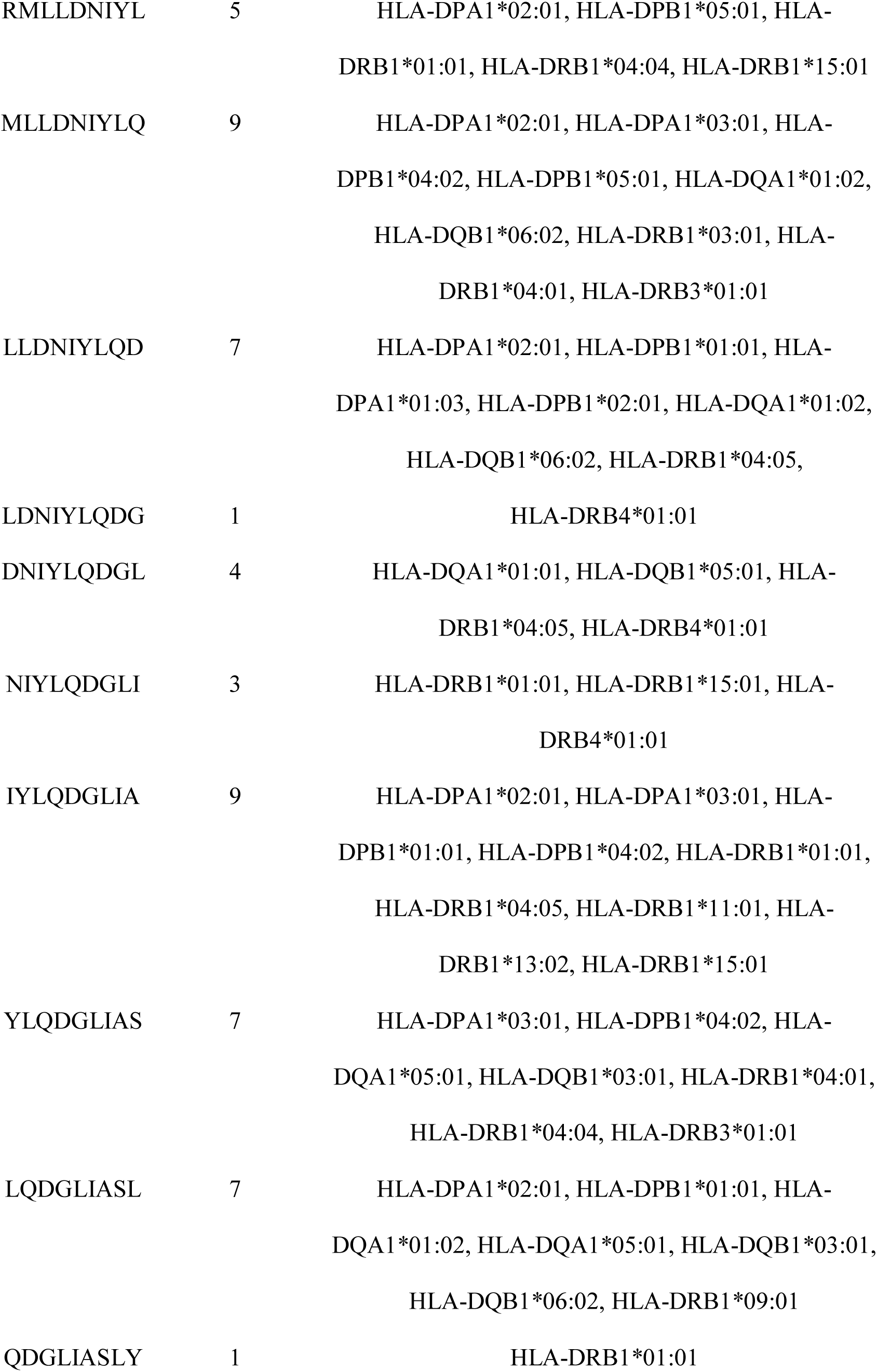
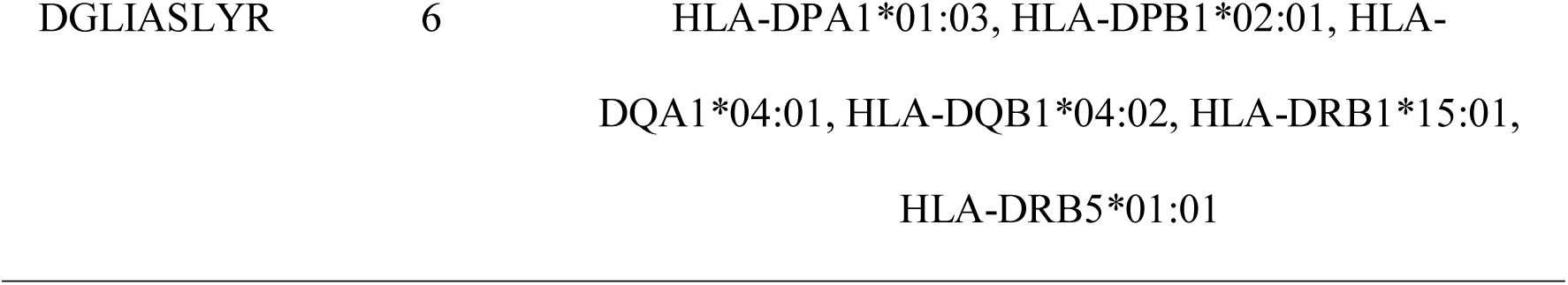
Selected CD4+ (THL) epitopes within the cluster region with their corresponding HLAs.

For B cell antigenicity of the epitope cluster, The Bepipred linear epitope prediction method was employed. Within the cluster, the region 1999-IYLQDGLIAS-2008 was predicted to induce a humoral immune response (Figure 05). In this case, the average predicted residue score, which is −1.053, was selected as the threshold score with the minimum score of −1.637 and the maximum score of −0.490 (Supplementary file 3).

**Figure 05.**
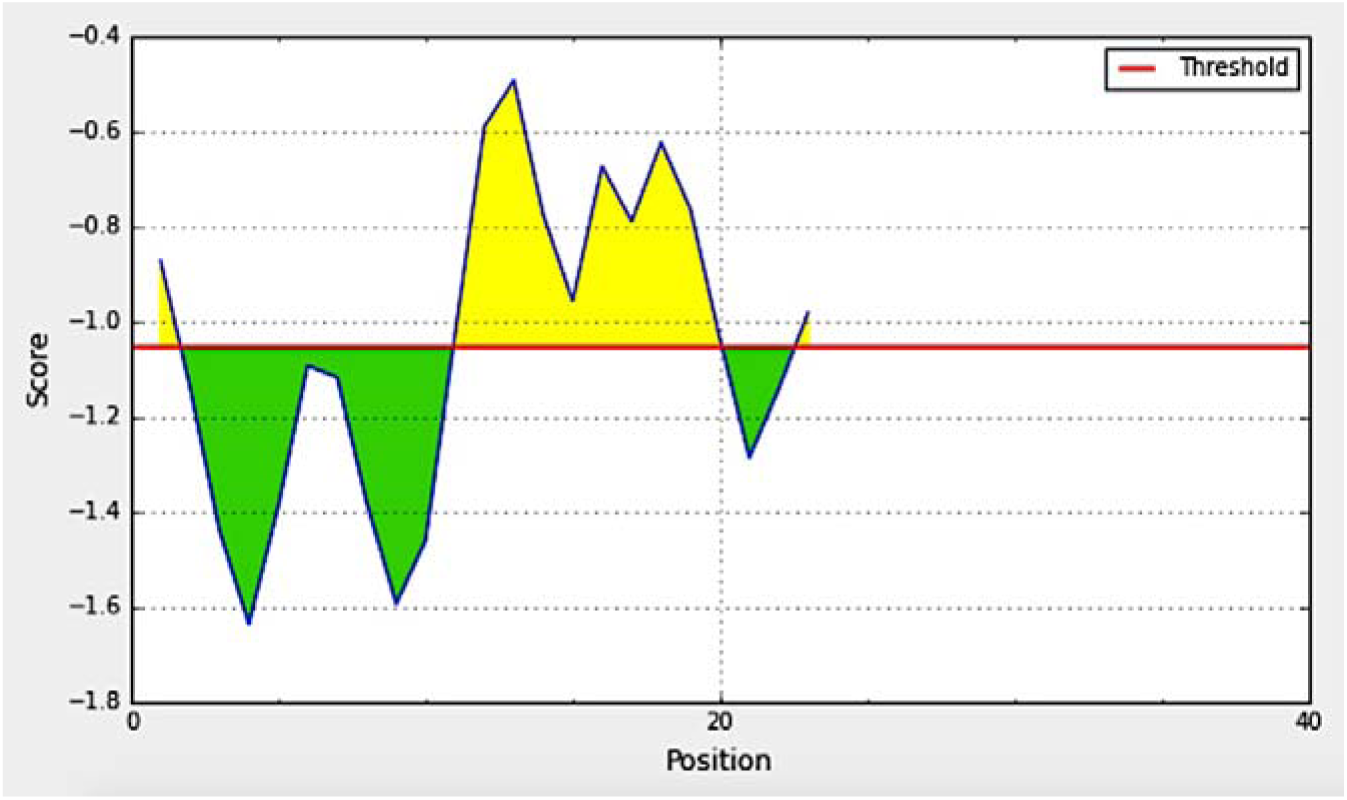
B cell epitope prediction using The Bepipred Linear Epitope Prediction method. The residues with scores above the threshold value are predicted to be part of an epitope and are colored in yellow.

### 3.6 Human peptide cross-reactivity analysis

Some whole cell and subunit vaccines include epitopes that occasionally mimic the native human peptides. We assessed the occurrence of contiguous sequences in human identical to the epitopes using the human non-redundant protein database. Our analysis resulted in a total of fourteen incidences where the epitopes matched with thirteen unique human protein sequences (Table 04). The maximum match of seven amino acids was found with proteins, germ cell-less protein-like 1 isoform X2, emopamil-binding protein-like isoform 1 and emopamil-binding protein-like isoform 2.

**Table 04.**
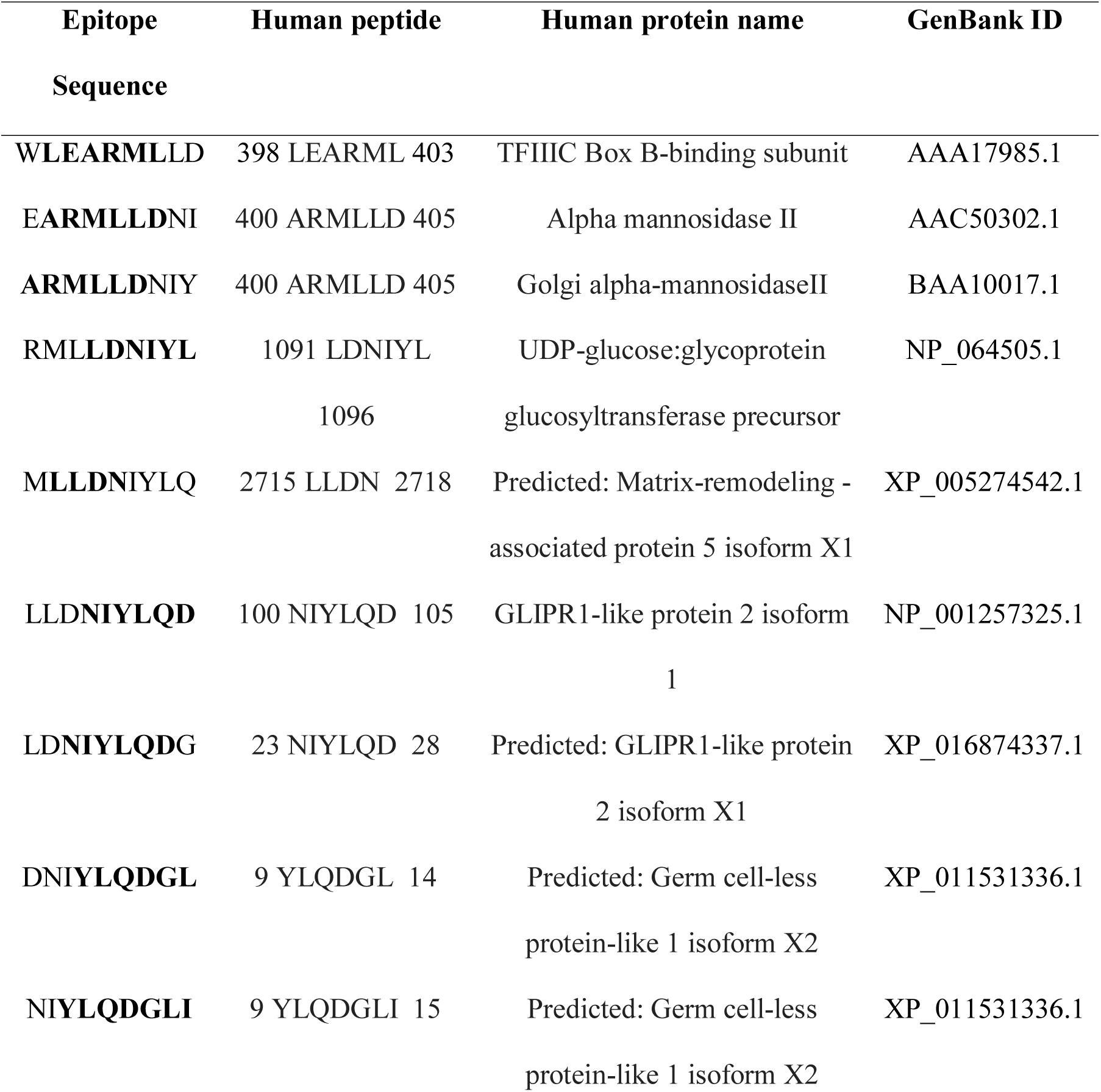
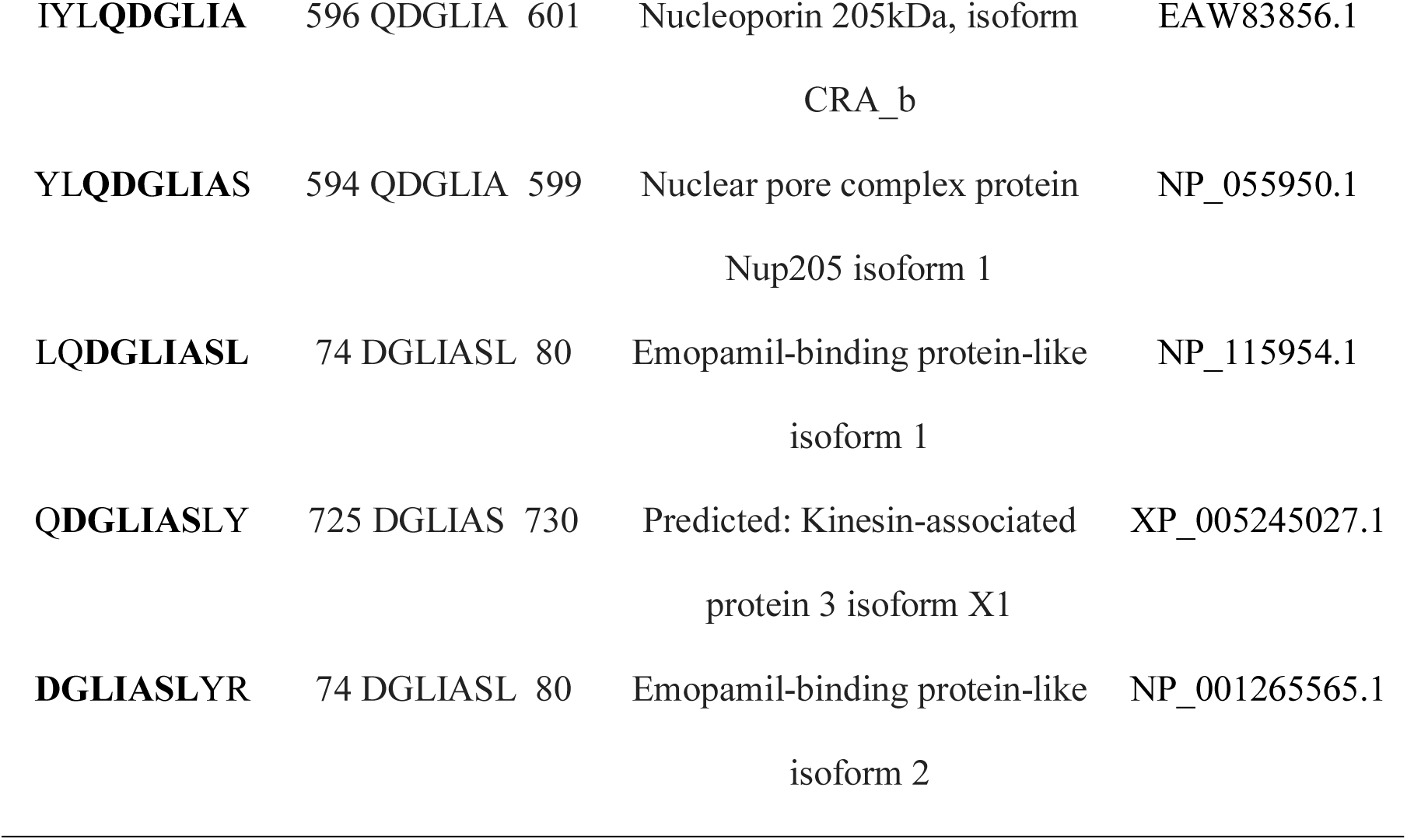
Human peptides analogous to the predicted epitopes. Amino acids in bold correspond to matched human peptide sequences. Positions of matched sequences within the human protein are denoted by numbers on either side of the peptide.

| Epitope<br>Sequence | Human peptide | Human protein name | GenBank ID |
| --- | --- | --- | --- |
| **WLEARMLLD** | 398 LEARML 403 | TFIIIC Box B-binding subunit | AAA17985.1 |
| **EARMLLDNI** | 400 ARMLLD 405 | Alpha mannosidase II | AAC50302.1 |
| **ARMLLDNIY** | 400 ARMLLD 405 | Golgi alpha-mannosidaseII | BAA10017.1 |
| **RMLLDNIYL** | 1091 LDNIYL<br>1096 | UDP-glucose:glycoprotein<br>glucosyltransferase precursor | NP_064505.1 |
| **MLLDNIYLQ** | 2715 LLDN 2718 | Predicted: Matrix-remodeling -<br>associated protein 5 isoform X1 | XP_005274542.1 |
| **LLDNIYLQD** | 100 NIYLQD 105 | GLIPR1-like protein 2 isoform<br>1 | NP_001257325.1 |
| **LDNIYLQDG** | 23 NIYLQD 28 | Predicted: GLIPR1-like protein<br>2 isoform X1 | XP_016874337.1 |
| **DNIYLQDGL** | 9 YLQDGL 14 | Predicted: Germ cell-less<br>protein-like 1 isoform X2 | XP_011531336.1 |
| **NIYLQDGLI** | 9 YLQDGLI 15 | Predicted: Germ cell-less<br>protein-like 1 isoform X2 | XP_011531336.1 |
| **IYLQDGLIA** | 596 QDGLIA 601 | Nucleoporin 205kDa, isoform<br>CRA_b | EAW83856.1 |
| **YLQDGLIAS** | 594 QDGLIA 599 | Nuclear pore complex protein<br>Nup205 isoform 1 | NP_055950.1 |
| **LQDGLIASL** | 74 DGLIASL 80 | Emopamil-binding protein-like<br>isoform 1 | NP_115954.1 |
| **QDGLIASLY** | 725 DGLIAS 730 | Predicted: Kinesin-associated<br>protein 3 isoform X1 | XP_005245027.1 |
| **DGLIASLYR** | 74 DGLIASL 80 | Emopamil-binding protein-like<br>isoform 2 | NP_001265565.1 |

### 3.7 Population coverage of the epitopes

Due to a smaller size, the usage of epitope-based vaccines remains restricted as they generate only a limited number of epitopes with low HLA allele coverage, even though epitopes with short peptides are HLA permissive [54, 55]. Due to the prevalence of human HLA polymorphism in different ethnic groups, the immunogenic responses elicited by epitope based vaccines may vary from one population to another. For this purpose, we calculated the population coverage in two ways – (1) Class I and II combined (2) Class II separate. Our results indicate that the predicted epitopes can generate a broad range of population coverage which is about 93.86% worldwide, with the maximum in North America (99.96%) and Europe (99.72%) for class I and II combined (Table 05). For class II, a similar population coverage was predicted that covers 91.80% of the population worldwide with the maximum in North America (99.91%) and Europe (99.46%) (Table 06).

**Table 05.** Class I and II combined percentage of population coverage of the selected epitopes in different geographical locations.

| Population/Area | Class I and II combined |  |  |
| --- | --- | --- | --- |
|  | Coverage <sup>a</sup> | Average hit <sup>b</sup> | PC90 <sup>c</sup> |
| Central Africa | 91.44 | 3.22 | 1.09 |
| Central America | 2.72 | 0.03 | 0.1 |
| East Africa | 92.09 | 3.48 | 1.13 |
| East Asia | 58.20 | 1.64 | 0.24 |
| Europe | 99.72 | 5.36 | 3.3 |
| North Africa | 50.32 | 1.37 | 0.2 |
| North America | 99.96 | 5.16 | 3.29 |
| Northeast Asia | 85.87 | 3.05 | 0.71 |
| Oceania | 83.87 | 2.99 | 0.62 |
| South America | 92.32 | 3.54 | 1.18 |
| South Asia | 97.47 | 3.57 | 1.8 |
| Southeast Asia | 43.94 | 0.95 | 0.18 |
| Southwest Asia | 49.64 | 1.39 | 0.2 |
| West Africa | 95.07 | 3.07 | 1.18 |
| West Indies | 63.19 | 1.8 | 0.27 |
| World | 93.86 | 4.31 | 1.44 |

<sup>a</sup> Projected population coverage.
<sup>b</sup> Average number of epitope hits/HLA combinations recognized by the population.
<sup>c</sup> Minimum number of epitope hits/HLA combinations recognized by 90% of the population

**Table 06.** Class II specific percentage of population coverage of the selected epitopes in different geographical locations.

| Population/Area | Class II |  |  |
| --- | --- | --- | --- |
|  | Coverage <sup>a</sup> | Average hit <sup>b</sup> | PC90 <sup>c</sup> |
| Central Africa | 97.87% | 4.21 | 1.74 |
| Central America | 1.34% | 0.03 | 0.2 |
| East Africa | 96.65% | 4.31 | 1.66 |
| East Asia | 46.18% | 1.59 | 0.37 |
| Europe | 99.46% | 4.41 | 3.17 |
| North Africa | 19.3% | 0.39 | 0.25 |
| North America | 99.91% | 3.92 | 3.15 |
| Northeast Asia | 84.23% | 3.29 | 0.63 |
| Oceania | 80.69% | 2.98 | 1.04 |
| South America | 95.93% | 4.12 | 1.9 |
| South Asia | 97.08% | 4.51 | 3.14 |
| Southeast Asia | 33.96% | 0.68 | 0.3 |
| Southwest Asia | 26.55% | 0.54 | 0.27 |
| West Africa | 96.68% | 4.73 | 3.12 |
| West Indies | 28.51% | 0.61 | 0.28 |
| World | 91.80% | 3.98 | 1.66 |

<sup>a</sup> Projected population coverage.
<sup>b</sup> Average number of epitope hits/HLA combinations recognized by the population.
<sup>c</sup> Minimum number of epitope hits/HLA combinations recognized by 90% of the population

### 3.8 Molecular docking of epitope and HLA interaction

The three-dimensional structures of epitopes were observed via PyMOL (Figure 06). The binding energy of predicted epitopes to their corresponding HLA molecules for both class I and class II were generated using AutoDock Vina tool (Figure 07 and 08). The grid box orientations and the sizes of the ligands and receptors across X, Y, and Z axes were provided separately to allow the epitopes to bind to the binding groove of the HLAs (supplementary file 2). The docking study showed strong epitope-HLA interaction for the epitopes within the cluster region. In case of MHC class I, the epitope ‘QDGLIASLY’ showed the most stable binding to the groove of HLA-A*29:02, with the energy of −6.9 kcal/mol (Table 07). Rest of the epitopes exhibited their binding energy below −6.0 kcal/mol with their corresponding HLAs except for ‘ARMLLDNIY’ (−4.9 kcal/mol) and ‘LQDGLIASL’ (−5.5 kcal/mol). With the binding energy of −6.3 kcal/mol, epitope ‘WLEARMLLD’ exhibited the most stable affinity towards the MHC class II molecule HLA-DRB1*04:05 (Table 08). In this case the average binding affinity was around −5.0 kcal/mol. The most potential epitopes were selected based on the higher binding energy.

**Table 07.** Docking results of class I epitopes with their corresponding allele interaction

| HLA Allele | Binding Epitope | Binding Energy (kcal/mol) |
| --- | --- | --- |
| HLA-A*30:02 | ARMLLDNIY | -4.9 |
| HLA-A*02:06 | RMLLDNIYL | -6.3 |
| HLA-A*02:06 | MLLDNIYLQ | -6.4 |
| HLA-A*02:06 | YLQDGLIAS | -6.4 |
| HLA-A*02:06 | LQDGLIASL | -5.5 |
| HLA-A*29:02 | QDGLIASLY | -6.9 |

**Table 08.** Docking results of class II epitopes with their corresponding alleles interaction.

| HLA Allele | Binding Epitope | Binding Energy (kcal/mol) |
| --- | --- | --- |
| HLA-DRB1*04:05 | WLEARMLLD | -6.3 |
| HLA-DRB4*01:01 | EARMLLDNI | -4.8 |
| HLA-DRB1*04:05 | ARMLLDNIY | -4.7 |
| HLA-DRB1*01:01 | RMLLDNIYL | -5.5 |
| HLA-DRB1*03:01 | MLLDNIYLQ | -5.7 |
| HLA-DRB1*04:05 | LLDNIYLQD | -5.7 |
| HLA-DRB4*01:01 | LDNIYLQDG | -5.0 |
| HLA-DRB4*01:01 | DNIYLQDGL | -5.8 |
| HLA-DRB1*01:01 | NIYLQDGLI | -5.0 |
| HLA-DRB1*04:05 | IYLQDGLIA | -5.2 |
| HLA-DRB1*04:04 | YLQDGLIAS | -5.9 |
| HLA-DRB1*09:01 | LQDGLIASL | -4.9 |
| HLA-DRB1*01:01 | QDGLIASLY | -4.8 |
| HLA-DRB1*15:01 | DGLIASLYR | -5.1 |

**Figure 06.**
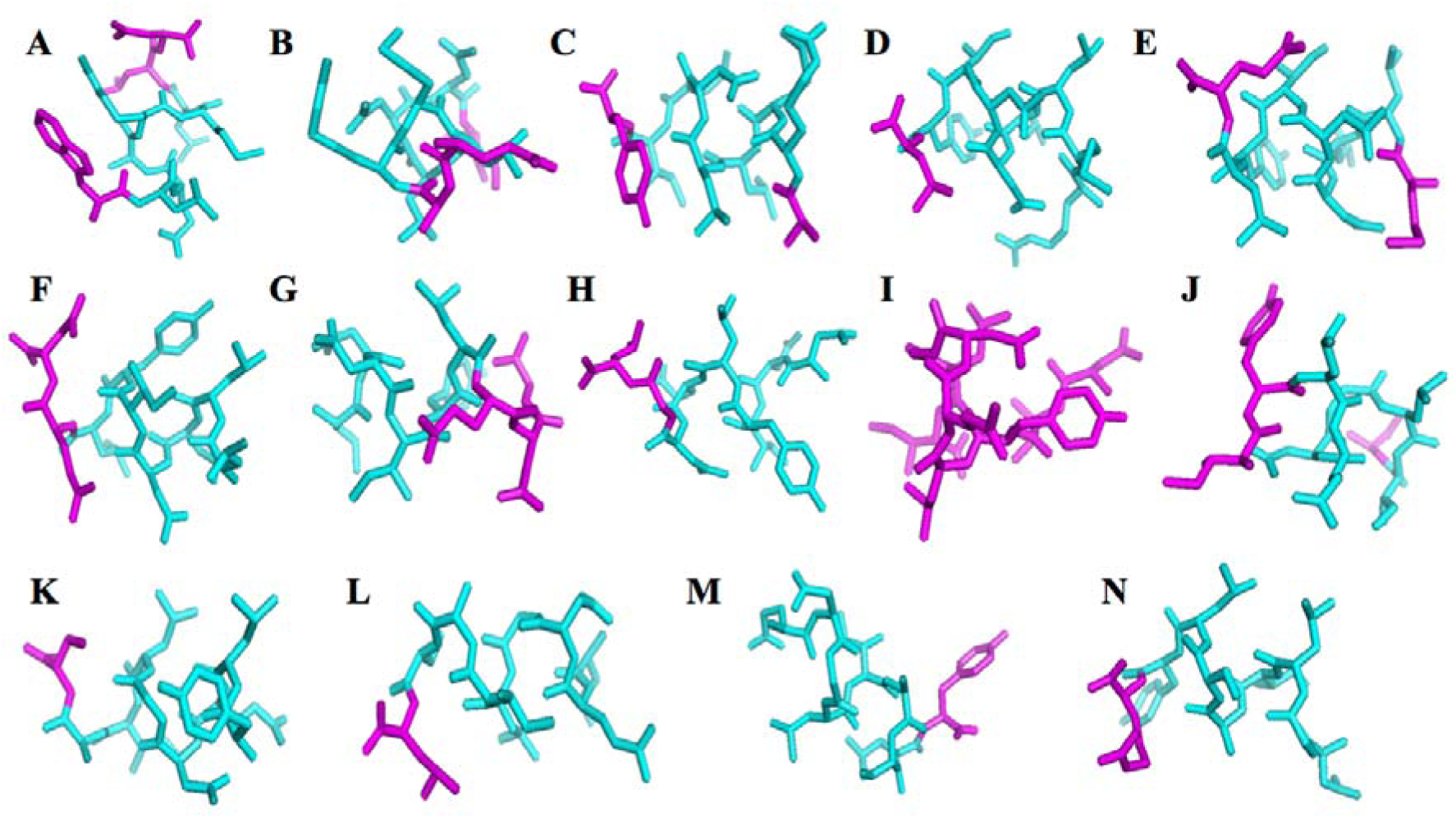
Three Dimensional structures of predicted epitopes. (A) WLEARMLLD (B) EARMLLDNI (C) ARMLLDNIY (D) RMLLDNIYL (E) MLLDNIYLQ (F) LLDNIYLQD (G) LDNIYLQDG (H) DNIYLQDGL (I) NIYLQDGL (J) IYLQDGLIA (K) YLQDGLIAS (L) LQDGLIASL (M) QDGLIASLY (N) DGLIASLYR. Cyan color indicates the helices and magenta color indicates the coiled portions.

**Figure 07.**
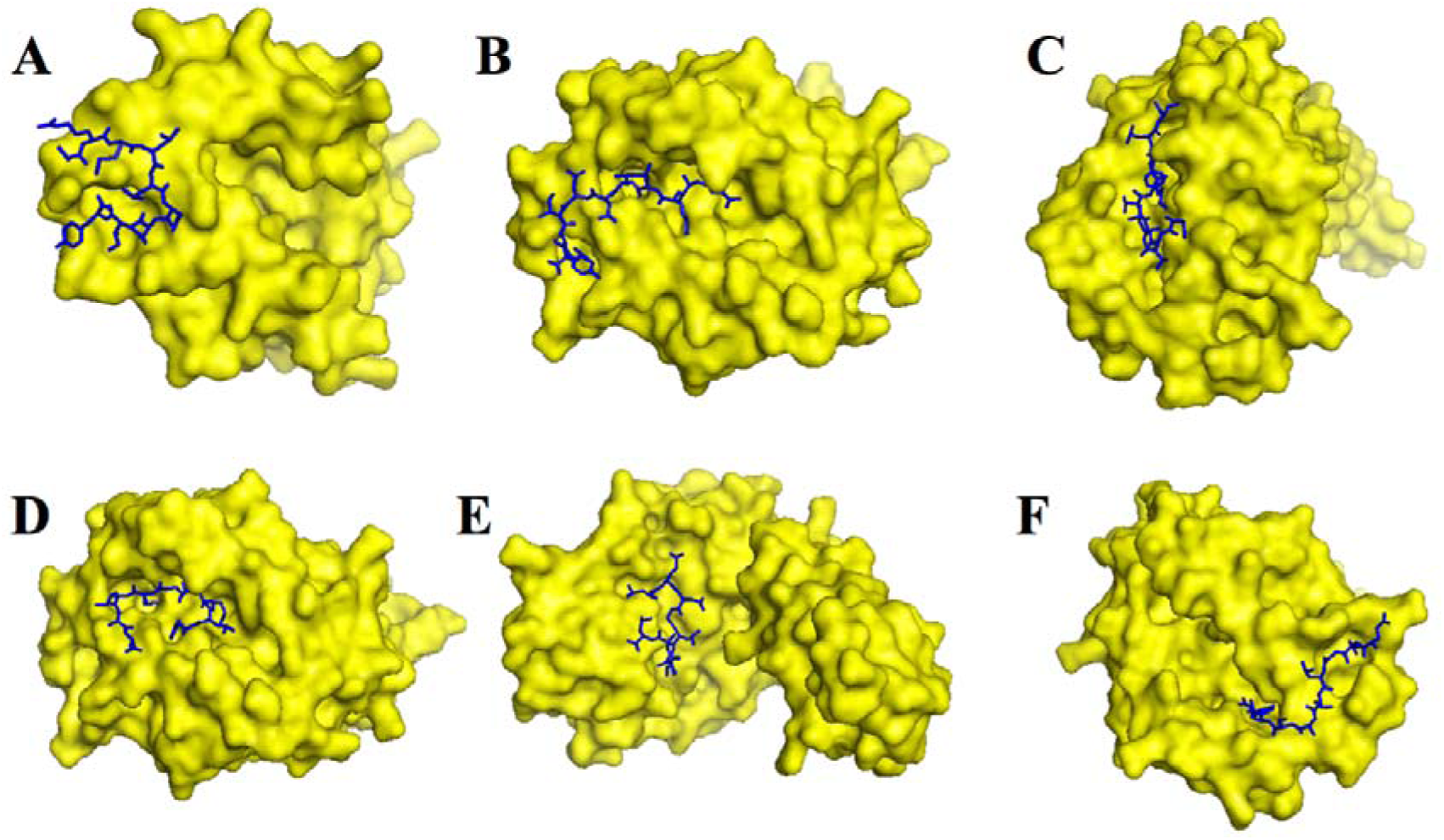
Molecular docking of class I (CTL) epitopes with their corresponding HLA. (A) ARMLLDNIY with HLA-A*30:02 (B) RMLLDNIYL with HLA-A*02:06 (C) MLLDNIYLQ with HLA-A*02:06 (D) YLQDGLIAS with HLA-A*02:06 (E) LQDGLIASL with HLA-A*02:06 (F) QDGLIASLY with HLA-A*29:02.

**Figure 08.**
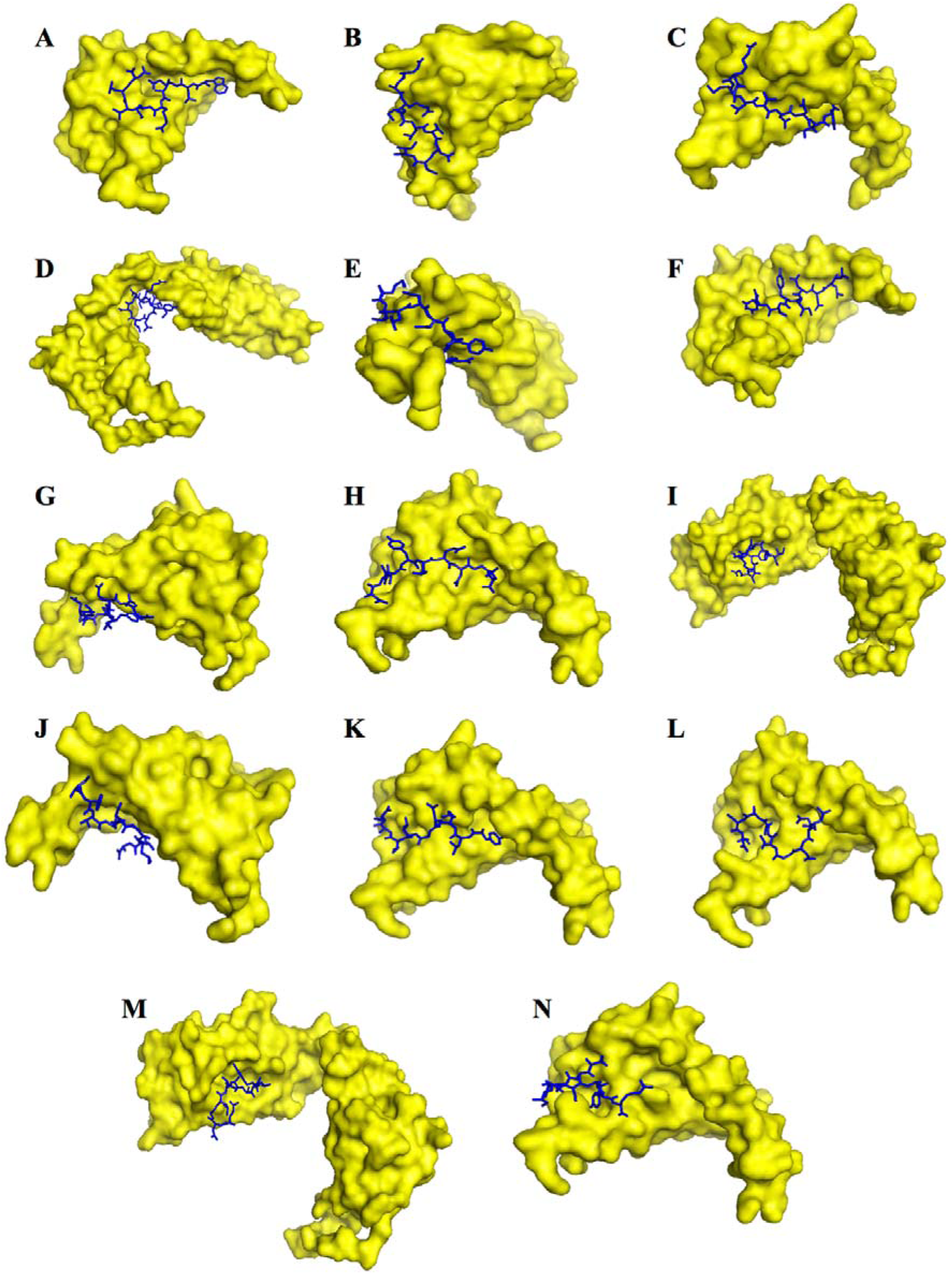
Molecular docking of class II (THL) epitopes with their corresponding HLA. (A) WLEARMLLD with HLA-DRB1*04:05 (B) EARMLLDNI with HLA-DRB4*01:01 (C) ARMLLDNIY with HLA-DRB1*04:05 (D) RMLLDNIYL with HLA-DRB1*01:01 (E) MLLDNIYLQ with HLA-DRB1*03:01 (F) LLDNIYLQD with HLA-DRB1*04:05 (G) LDNIYLQDG with HLA-DRB4*01:01 (H) DNIYLQDGL with HLA-DRB4*01:01 (I) NIYLQDGLI with HLA-DRB1*01:01 (J) IYLQDGLIA with HLA-DRB1*04:05 (K) YLQDGLIAS with HLA-DRB1*04:04 (L) LQDGLIASL with HLA-DRB1*09:01 (M) QDGLIASLY with HLA-DRB1*01:01 (N) DGLIASLYR with HLA-DRB1*15:01.

## 4. Discussion

In this study, we have focused on the prediction of an epitope cluster within Zika Virus proteome using a range of epitope predicting tools from immune epitope database. To be considered as a potential vaccine candidate for infection, an epitope is required to fulfil some criteria that comply with the host genetic makeup, defence mechanism, and autoimmune response. The present in silico study helped us predict an epitope cluster of 23 amino acids in ZIKV genome polyprotein that is capable of eliciting cell-mediated and humoral immune response. The predicted epitope cluster was then analysed for autoimmune response, population coverage worldwide as well as for HLA binding capacity by molecular docking analysis.

For CTL epitope prediction, we have employed “Proteasomal cleavage/ Tap transport/ MHC class I combined predictor” algorithm, which has also been applied in previous studies for successful epitope identification in *M. tuberculosis* and Respiratory Syncytial Virus proteome [28, 31]. THL mediated response study was conducted using the ANN method. K-means cluster helped in the identification of the epitope cluster in ZIKV proteome, where red color indicates the potential epitope dense region that has a high allele count with low IC50 value. The most potential epitope cluster region was identified in 1989-2011 region where 23 contiguous amino acids were found and visible by red color in both side -IC50 score and unique allele count (Figure 4A). Furthermore, the Bepipred linear epitope prediction method aided in the B cell antigenicity of the cluster region. The combined results of these tools along with sequence conservation study assisted us in the successful prediction of an epitope cluster capable of evoking humoral and cell-mediated immunity. Further investigation within the proteome helped us identify the protein NS3 helicase containing the cluster region (Figure 4B). The protein NS3 RNA helicase plays a crucial role in unwinding the dsRNA in 3’ to 5’ direction upon its binding with RNA and has previously been suggested to be an attractive target in the discovery of antiviral drugs [56, 57]. Being a member of superfamily 2 (SF2) of helicases, NS3 helicase renders the virus incapable of replicating upon its inactivation [58, 59]. Residing in this protein, our predicted cluster region can generate six different epitopes capable of binding with seven different MHC class I alleles (Table 02) and 14 different epitopes capable of binding with 31 different MHC class II alleles (Table 03). Previously, the C and E proteins of ZIKV have been observed to elicit CD4+ and CD8+ T cell responses in monkeys [60, 61] and NS1 and E proteins in ZIKV-infected humans [62]. Although unclear, the role of CD8+ T cells has been suggested to be associated with the reduction of viral loads, mortality and ZIKV induced weight loss in both wild-type and immune-compromised mouse models [63]. For B cell-mediated response, this cluster is predicted to generate a ten amino acids long continuous peptide sequence and a total of 12 residues out of 23 possess significantly higher B cell antigenicity than the rest of the cluster region (Figure 05). These results are evidential to the predicted epitope cluster of being a broad-spectrum vaccine candidate upon its development.

In vaccine development, subjugating the autoimmune or allergic response is considered to be a substantial hurdle. We compared our cluster sequence to the human non-redundant protein sequence database for analyzing the possible mimicry of the epitopes to the native human peptides. This scrutiny resulted in three human proteins which are analogous to the cluster sequence with the maximum of seven amino acids (Table 04). The absence of the identical sequences of eight or more amino acids is incapable of binding to the grooves of HLA molecules. This dramatically reduces the likelihood of priming autoreactive T cells and thus minimizing the probability of autoimmune diseases [51]. In conjunction with an autoimmune response, polymorphism across diverse human population also presents to be an imposing task in the development of a vaccine with high effectiveness. A highly polymorphic group of cell-surface receptors located on all the cells of the body known as the MHC plays a critical role in the immune system. The two major categories namely MHC class I, found on all nucleated cells, and MHC class II, expressed by the antigen presenting complexes (APCs), are associated with the function of differentiating between self-proteins and non-self-proteins in the body. Variation within the MHC alleles in human population yields different types of immune response for the same vaccine. Also, because the epitope vaccines are small and they bind with a smaller number of HLA, epitope vaccines are often suspected of providing low population coverage. Thus, ensuring high population coverage is another major challenge of epitope vaccine. On that account, our epitope cluster was subjected to the population coverage study (Tables 5–6). Our selected epitope cluster is predicted to provide high population coverage worldwide (93.86%), with the maximum in North America and Europe which might be due to these regions having a higher number of travellers [64].

Furthermore, we undertook the molecular docking analysis of each of these epitopes with one of their corresponding HLAs to calculate their affinity towards each other using the AutoDock Vina tool. In this study, the HLAs for each epitope was selected based on their prevalence to assess the potency of the epitope cluster. In all the cases, the binding energy was found to be below or around −5.0 kcal/mol, with the highest of −4.9 and −4.7 kcal/mol for MHC class I and II molecules respectively, which demonstrates the competence of the epitope cluster as a potential vaccine candidate in vivo.

## 5. Conclusion

ZIKV is a significant public health emergency at present. Fast-track investigations on vaccine development are desired to control the spread of ZIKV infection. Immunoinformatics approaches are the emerging field nowadays to predict epitope vaccine candidate agent against virus and bacteria. Here, we report a cluster of 23 contiguous amino acids after screening the whole proteome of ZIKV that includes THL, CTL and B cell epitopes overlapped within it. These epitopes are predicted to provide 93.86% population coverage worldwide as well as potent HLA binding capability. The in silico data about the reported epitope cluster will pave the way to design precisely targeted synthetic epitope vaccine against ZIKV. We do hope, our work would join some loose ends of in silico and clinical analysis of ZIKV immunity and vaccine development.

## Acknowledgements

All the authors acknowledge support of Department of Biochemistry and Molecular Biology, Shahjalal University of Science and Technology, Sylhet, Bangladesh.

## Declarations

### Ethics approval and consent to participate

Not applicable

