## Supplementary file 1 for "Zika viral proteome analysis reveals an epitope cluster within NS3 helicase as a potential vaccine candidate: an *in silico* study"

**Protein sequences (Aligned)**

>tr|A0A125RTB5|A0A125RTB5_ZIKV Genome polyprotein OSZika virus (strain Mr 766) PE4 SV1

MKNPKKKSGGFRIVNMLKRGVARVSPFGGLKRLPAGLLLGHGPIRMVLAILAFLRFTAIKPSLGLINRWGSVGKKEAMEIIKKFKKDLAAMLRIINARKEKKRRGTDTSVGIVGLLLTTAMAAEVTRRGSAYYMYLDRSDAGEAISFPTTLGMNKCYIQIMDLGHMCDATMSYECPMLDEGVEPDDVDCWCNTTSTWVVYGTCHHKKGEARRSRRAVTLPSHSTRKLQTRSQTWLESREYTKHLIRVENWIFRNPGFALAAAAIAWLLGSSTSQKVIYLVMILLIAPAYSIRCIGVSNRDFVEGMSGGTWVDVVLEHGGCVTVMAQDKPTVDIELVTTTVSNMAEVRSYCYEASISDMASDSRCPTQGEAYLDKQSDTQYVCKRTLVDRGWGNGCGLFGKGSLVTCAKFACSKKMTGKSIQPENLEYRIMLSVHGSQHSGMIVNDTGHETDENRAKVEITPNSPRAEATLGGFGSLGLDCEPRTGLDFSDLYYLTMNNKHWLVHKEWFHDIPLPWHTGADTGTPHWNNKEALVEFKDAHAKRQTVVVLGSQEGAVHTALAGALEAEMDGAKGRLSSGHLKCRLKMDKLRLKGVSYSLCTAAFTFTKIPAETLHGTVTVEVQYAGTDGPCKVPAQMAVDMQTLTPVGRLITANPVITEGTENSKMMLELDPPFGDSYIVIGVGEKKITHHWHRSGSTIGKAFEATVRGAKRMAVLGDTAWDFGSVGGVLNSLGKGIHQIFGAAFKSLFGGMSWFSQILIGTLLMWLGLNTKNGSISLMCLALGGVLIFLSTAVSADVGCSVDFSKKETRCGTGVFVYNDVEAWRDRYKYHPDSPRRLAAVVKQAWEDGICGISSVSRMENIMWRSVEGELNAILEENGVQLTVVVGSVKNPMWRGPQRLPVPVNELPHGWKAWGKSYFVRAAKTNNSFVVDGDTLKECPLKHRAWNSFLVEDHGFGVFHTSVWLKVREDYSLECDPAVIGTAVKGKEAVHSDLGYWIESEKNDTWRLRRAHLIEMKTCEWPKSHTLWTDGIEESDLIIPKSLAGPLSHHNTREGYRTQMKGPWHSEELEIRFEECPGTKVHVEETCGTRGPSLRSTTASGRVIEEWCCRECTMPPLSFRAKDGCWYGMEIRPRKEPESNLVRSMVTAGSTDHMDHFSLGVLVILLMVQEGLKKRMTTKIIISTSMAVLVAMILGGFSMSDLAKLAILMGATFAEMNTGGDVAHLALVAAFKVRPALLVSFIFRANWTPRESMLLALASCLLQTAISALEGDLMVLINGFALAWLAIRAMVVPRTDNITLAILAALTPLARGTLLVAWRAGLATCGGFMLLSLKGKGSVKKNLPFVMALGLTAVRLVDPINVVGLLLLTRSGKRSWPPSEVLTAVGLICALAGGFAKADIEMAGPMAAVGLLIVSYVVSGKSVDMYIERAGDITWEKDAEVTGNSPRLDVALDESGDFSLVEDDGPPMREIILKVVLMTICGMNPIAIPFAAGAWYVYVKTGKRSGALWDVPAPKEVKKGETTDGVYRVMTRRLLGSTQVGVGVMQEGVFHTMWHVTKGSALRSGEGRLDPYWGDVKQDLVSYCGPWKLDAAWDGHSEVQLLAVPPGERARNIQTLPGIFKTKDGDIGAVALDYPAGTSGSPILDKCGRVIGLYGNGVVIKNGSYVSAITQGRREEETPVECFEPSMLKKKQLTVLDLHPGAGKTRRVLPEIVREAIKTRLRTVILAPTRVVAAEMEEALRGLPVRYMTTAVNVTHSGTEIVDLMCHATFTSRLLQPIRVPNYNLYIMDEAHFTDPSSIAARGYISTRVEMGEAAAIFMTATPPGTRDAFPDSNSPIMDTEVEVPERAWSSGFDWVTDHSGKTVWFVPSVRNGNEIAACLTKAGKRVIQLSRKTFETEFQKTKHQEWDFVVTTDISEMGANFKADRVIDSRRCLKPVILDGERVILAGPMPVTHASAAQRRGRIGRNPNKPGDEYLYGGGCAETDEDHAHWLEARMLLDNIYLQDGLIASLYRPEADKVAAIEGEFKLRTEQRKTFVELMKRGDLPVWLAYQVASAGITYTDRRWCFDGMTNNTIMEDSVPAEVWTRHGEKRVLKPRWMDARVCSDHAALKSFKEFAAGKRGAAFGVMEALGTLPGHMTERFQEAIDNLAVLMRAETGSRPYKAAAAQLPETLETIMLLGLLGTVSLGIFFVLMRNKGIGKMGFGMVTLGASAWLMWLSEIEPARIACVLIVVFLLLVVLIPEPEKQRSPQDNQMAIIIMVAVGLLGLITANELGWLERTKSDLSHLMGRREEGATIGFSMDIDLRPASAWAIYAALTTFITPAVQHAVTTSYNNYSLMAMATQAGVLFGMGKGMPFYAWDFGVPLLMIGCYSQLTPLTLIVAIILLVAHYMYLIPGLQAAAARAAQKRTAAGIMKNPVVDGIVVTDIDTMTIDPQVEKKMGQVLLIAVAVSSAILSRTAWGWGEAGALITAATSTLWEGSPNKYWNSSTATSLCNIFRGSYLAGASLIYTVTRNAGLVKRRGGGTGETLGEKWKARLNQMSALEFYSYKKSGITEVCREEARRALKDGVATGGHAVSRGSAKLRWLVERGYLQPYGKVIDLGCGRGGWSYYAATIRKVQEVKGYTKGGPGHEEPMLVQSYGWNIVRLKSGVDVFHMAAEPCDTLLCDIGESSSSPEVEEARTLRVLSMVGDWLEKRPGAFCVKVLCPYTSTMMETLERLQRRYGGGLVRVPLSRNSTHEMYWVSGAKSNTIKSVSTTSQLLLGRMDGPRRPVKYEEDVNLGSGTRAVVSCAEAPNMKIIGNRIERIRSEHAETWFFDENHPYRTWAYHGSYEAPTQGSASSLINGVVRLLSKPWDVVTGVTGIAMTDTTPYGQQRVFKEKVDTRVPDPQEGTRQVMSMVSSWLWKELGKHKRPRVCTKEEFINKVRSNAALGAIFEEEKEWKTAVEAVNDPRFWALVDKEREHHLRGECQSCVYNMMGKREKKQGEFGKAKGSRAIWYMWLGARFLEFEALGFLNEDHWMGRENSGGGVEGLGLQRLGYVLEEMSRIPGGRMYADDTAGWDTRISRFDLENEALITNQMEKGHRALALAIIKYTYQNKVVKVLRPAEKGKTVMDIISRQDQRGSGQVVTYALNTFTNLVVQLIRNMEAEEVLEMQDLWLLRRSEKVTNWLQSNGWDRLKRMAVSGDDCVVKPIDDRFAHALRFLNDMGKVRKDTQEWKPSTGWDNWEEVPFCSHHFNKLHLKDGRSIVVPCRHQDELIGRARVSPGAGWSIRETACLAKSYAQMWQLLYFHRRDLRLMANAICSSVPVDWVPTGRTTWSIHGKGEWMTTEDMLVVWNRVWIEENDHMEDKTPVTKWTDIPYLGKREDLWCGSLIGHRPRTTWAENIKNTVNMVRRIIGDEEKYMDYLSTQVRYLGEEGSTPGVL

>tr|A0A125RTB6|A0A125RTB6_ZIKV Genome polyprotein OSZika virus (strain Mr 766) PE4 SV1

MKNPKKKSGGFRIVNMLKRGVARVSPFGGLKRLPAGLLLGHGPIRMVLAILAFLRFTAIKPSLGLINRWGSVGKKEAMEIIKKFKKDLAAMLRIINARKEKKRRGADTSVGIVGLLLTTAMAVEVTRRGSAYYMYLDRSDAGEAISFPTTLGMNKCYIQIMDLGHMCDATMSYECPMLDEGVEPDDVDCWCNTTSTWVVYGTCHHKKGEARRSRRAVTLPSHSTRKLQTRSQTWLESREYTKHLIRVENWIFRNPGFALAAAVIAWLLGSSTSQKVIYLVMILLIAPAYSIRCIGVSNRDFVEGMSGGTWVDVVLEHGGCVTVMAQDKPTVDIELVTTTVSNMAEVRSYCYEASISDMASDSRCPTQGEAYLDKQSDTQYVCKRTLVDRGWGNGCGLFGKGSLVTCAKFACSKKMTGKSIQPENLEYRIMLSVHGSQHSGMIVNDTGHETDENRAKVEITPNSPRAEATLGGFGSLGLDCEPRTGLDFSDLYYLTMNNKHWLVHKEWFHDIPLPWHAGADTGTPHWNNKEALVEFKDAHAKRQTVVVLGSQEGAVHTALAGALEAEMDGAKGRLSSGHLKCRLKMDKLRLKGVSYSLCTAAFTFTKIPAETLHGTVTVEVQYAGTDGPCKVPAQMAVDMQTLTPVGRLITANPVITESTENSKMMLELDPPFGDSYIVIGVGEKKITHHWHRSGSTIGKAFEATVRGAKRMAVLGDTAWDFGSVGGALNSLGKGIHQIFGAAFKSLFGGMSWFSQILIGTLLVWLGLNTKNGSISLTCLALGGVLIFLSTAVSADVGCSVDFSKKETRCGTGVFVYNDVEAWRDRYKYHPDSPRRLAAAVKQAWEDGICGISSVSRMENIMWRSVEGELNAILEENGVQLTVVVGSVKNPMWRGPQRLPVPVNELPHGWKAWGKSYFVRAAKTNNSFVVDGDTLKECPLKHRAWNSFLVEDHGFGVFHTSVWLKVREDYSLECDPAVIGTAAKGKEAVHSDLGYWIESEKNDTWRLKRAHLIEMKTCEWPKSHTLWTDGVEESDLIIPKSLAGPLSHHNTREGYRTQMKGPWHSEELEIRFEECPGTKVHVEETCGTRGPSLRSTTASGRVIEEWCCRECTMPPLSFRAKDGCWYGMEIRPRKEPESNLVRSMVTAGSTDHMDHFSLGVLVILLMVQEGLKKRMTTKIIISTSMAVLVAMILGGFSMSDLAKLAILMGATFAEMNTGGDVAHLALIAAFKVRPALLVSFIFRANWTPRESMLLALASCLLQTAISALEGDLMVLINGFALAWLAIRAMVVPRTDNITLAILAALTPLARGTLLVAWRAGLATCGGFMLLSLKGKGSVKKNLPFVMALGLTAVRLVDPINVVGLLLLTRSGKRSWPPSEVLTAVGLICALAGGFAKADIEMAGPMAAVGLLIVSYVVSGKSVDMYIERAGDITWEKDAEITGNSPRLDVALDESGDFSLVEDDGPPMREIILKVVLMTICGMNPIAIPFAAGAWYVYVKTGKRSGALWDVPAPKEVKKGETTDGVYRVMTRRLLGSTQVGVGVMQEGVFHTMWHVTKGSALRSGEGRLDPYWGDVKQDLVSYCGPWKLDAAWDGHSEVQLLAVPPGERARNIQTLPGTFKTKDGDIGAVALDYPAGTSGSPILDKCGRVIGLYGNGVVIKNGSYVSAITQGRREEETPVECFEPSMLKKKQLTVLDLHPGAGKTRRVLPEIVREAIKTRLRTVILAPTRVVAAEMEEALRGLPVRYMTTAVNVTHSGTEIVDLMCHATFTSRLLQPIRVPNYNLYIMDEAHFTDPSSIAARGYISTRVEMGEAAAIFMTATPPGTRDAFPDSNSPIMDTEVEVPERAWSTGFDWVTDHSGKTVWFVPSVRNGNEIAACLTKAGKRVIQLSRKTFETEFQKTKNQEWDFVVTTDISEMGANFKADRVIDSRRCLKPVILDGERVILAGPMPVTHASAAQRRGRIGRNPNKPGDEYLYGGGCAETDEDHAHWLEARMLLDNIYLQDGLIASLYRPEADKVAAIEGEFKLRTEQRKTFVELMKRGDLPVWLAYQVASAGITYTDRRWCFDGMTNNTIMEDSVPAEVWTRYGEKRVLKPRWMDARVCSDHAALKSFKEFAAGKRGAAFGVIEALGTLPGHMTERFQEAIDNLAVLMRAETGSRPYKAAAAQLPETLETIMLLGLLGTVSLGIFFVLMRNKGMGKMGFGMVTLGASAWLMWLSEIEPARIACVLIVVFLLLVVLIPEPEKQRSPQDNQMAIIIMVAVGLLGLITANELGWLERTKSDLSHLMGRREEGATTGFSMDIDLRPASAWAIYAALTTFITPAVQHAVTTSYNNYSLMAMATQAGVLFGMGKGMPFYAWDFGVPLLMMGCYSQLTPLTLIVAIILLVAHYMYLIPGLQAAAARAAQKRTAAGIMKNPVVDGIVVTDIDTMTIDPQVEKKMGQVLLIAVAVSSAILSRTAWGWGEAGALITAATSTLWEGSPNKYWNSSTATSLCNIFRGSYLAGASLIYTVTRNAGLVKRRGGGTGETLGEKWKARLNQMSALEFYSYKKSGITEVCREEARRALKDGVATGGHAVSRGSAKLRWLVERGYLQPYGKVIDLGCGRGGWSYYAATIRKVQEVKGYTKGGPGHEEPMLVQSYGWNIVRLKSGVDVFHMAAEPCDTLLCDIGESSSSPEVEEARTLRVLSMVGDWLEKRPGAFCIKVLCPYTSTMMETLERLQRRYGGGLVRVPLSRNSTHEMYWVSGAKSNTIKSVSTTSQLLLGRMDGPRRPVKYEEDVNLGSGTRAVVSCAEAPNMKIIGNRIERIRSEHAETWFFDENHPYRTWAYHGSYEAPTQGSASSLINGVVRLLSKPWDVVTGVTGIAMTDTTPYGQQRVFKEKVDTRVPDPQEGTRQVMSMVSSWLWKELGKHKRPRVCTKEEFINKVRSNAALGAIFEEEKEWKTAVEAVNDPRFWALVDKEREHHLRGECQSCVYNMMGKREKKQGEFGKAKGSRAIWYMWLGARFLEFEALGFLNEDHWMGRENSGGGVEGLGLQRLGYVLEEMSRIPGGRMYADDTAGWDTRISRFDLENEALITNQMEKGHRALALAIIKYTYQNKVVKVLRPAEKGKTVMDIISRQDQRGSGQVVTYALNTFTNLVVQLIRNMEAEEVLEMQDLWLLRRPEKVTNWLQSNGWDRLKRMAVSGDDCVVKPIDDRFAHALRFLNDMGKVRKDTQEWKPSTGWDNWEEVPFCSHHFNKLHLKDGRSIVVPCRHQDELIGRARVSPGAGWSIRETACLAKSYAQMWQLLYFHRRDLRLMANAICSSVPVDWVPTGRTTWSIHGKGEWMTTEDMLVVWNRVWIEENDHMEDKTPVTKWTDIPYLGKREDLWCGSLIGHRPRTTWAENIKNTVNMMRRIIGDEEKYMDYLSTQVRYLGEEGSTPGVL

>tr|A0A024B7W1|A0A024B7W1_ZIKV Genome polyprotein OSZika virus (strain Mr 766) PE1 SV1

MKNPKKKSGGFRIVNMLKRGVARVSPFGGLKRLPAGLLLGHGPIRMVLAILAFLRFTAIKPSLGLINRWGSVGKKEAMEIIKKFKKDLAAMLRIINARKEKKRRGADTSVGIVGLLLTTAMAAEVTRRGSAYYMYLDRNDAGEAISFPTTLGMNKCYIQIMDLGHMCDATMSYECPMLDEGVEPDDVDCWCNTTSTWVVYGTCHHKKGEARRSRRAVTLPSHSTRKLQTRSQTWLESREYTKHLIRVENWIFRNPGFALAAAAIAWLLGSSTSQKVIYLVMILLIAPAYSIRCIGVSNRDFVEGMSGGTWVDVVLEHGGCVTVMAQDKPTVDIELVTTTVSNMAEVRSYCYEASISDMASDSRCPTQGEAYLDKQSDTQYVCKRTLVDRGWGNGCGLFGKGSLVTCAKFACSKKMTGKSIQPENLEYRIMLSVHGSQHSGMIVNDTGHETDENRAKVEITPNSPRAEATLGGFGSLGLDCEPRTGLDFSDLYYLTMNNKHWLVHKEWFHDIPLPWHAGADTGTPHWNNKEALVEFKDAHAKRQTVVVLGSQEGAVHTALAGALEAEMDGAKGRLSSGHLKCRLKMDKLRLKGVSYSLCTAAFTFTKIPAETLHGTVTVEVQYAGTDGPCKVPAQMAVDMQTLTPVGRLITANPVITESTENSKMMLELDPPFGDSYIVIGVGEKKITHHWHRSGSTIGKAFEATVRGAKRMAVLGDTAWDFGSVGGALNSLGKGIHQIFGAAFKSLFGGMSWFSQILIGTLLMWLGLNTKNGSISLMCLALGGVLIFLSTAVSADVGCSVDFSKKETRCGTGVFVYNDVEAWRDRYKYHPDSPRRLAAAVKQAWEDGICGISSVSRMENIMWRSVEGELNAILEENGVQLTVVVGSVKNPMWRGPQRLPVPVNELPHGWKAWGKSYFVRAAKTNNSFVVDGDTLKECPLKHRAWNSFLVEDHGFGVFHTSVWLKVREDYSLECDPAVIGTAVKGKEAVHSDLGYWIESEKNDTWRLKRAHLIEMKTCEWPKSHTLWTDGIEESDLIIPKSLAGPLSHHNTREGYRTQMKGPWHSEELEIRFEECPGTKVHVEETCGTRGPSLRSTTASGRVIEEWCCRECTMPPLSFRAKDGCWYGMEIRPRKEPESNLVRSMVTAGSTDHMDHFSLGVLVILLMVQEGLKKRMTTKIIISTSMAVLVAMILGGFSMSDLAKLAILMGATFAEMNTGGDVAHLALIAAFKVRPALLVSFIFRANWTPRESMLLALASCLLQTAISALEGDLMVLINGFALAWLAIRAMVVPRTDNITLAILAALTPLARGTLLVAWRAGLATCGGFMLLSLKGKGSVKKNLPFVMALGLTAVRLVDPINVVGLLLLTRSGKRSWPPSEVLTAVGLICALAGGFAKADIEMAGPMAAVGLLIVSYVVSGKSVDMYIERAGDITWEKDAEVTGNSPRLDVALDESGDFSLVEDDGPPMREIILKVVLMTICGMNPIAIPFAAGAWYVYVKTGKRSGALWDVPAPKEVKKGETTDGVYRVMTRRLLGSTQVGVGVMQEGVFHTMWHVTKGSALRSGEGRLDPYWGDVKQDLVSYCGPWKLDAAWDGHSEVQLLAVPPGERARNIQTLPGIFKTKDGDIGAVALDYPAGTSGSPILDKCGRVIGLYGNGVVIKNGSYVSAITQGRREEETPVECFEPSMLKKKQLTVLDLHPGAGKTRRVLPEIVREAIKTRLRTVILAPTRVVAAEMEEALRGLPVRYMTTAVNVTHSGTEIVDLMCHATFTSRLLQPIRVPNYNLYIMDEAHFTDPSSIAARGYISTRVEMGEAAAIFMTATPPGTRDAFPDSNSPIMDTEVEVPERAWSSGFDWVTDHSGKTVWFVPSVRNGNEIAACLTKAGKRVIQLSRKTFETEFQKTKHQEWDFVVTTDISEMGANFKADRVIDSRRCLKPVILDGERVILAGPMPVTHASAAQRRGRIGRNPNKPGDEYLYGGGCAETDEDHAHWLEARMLLDNIYLQDGLIASLYRPEADKVAAIEGEFKLRTEQRKTFVELMKRGDLPVWLAYQVASAGITYTDRRWCFDGTTNNTIMEDSVPAEVWTRHGEKRVLKPRWMDARVCSDHAALKSFKEFAAGKRGAAFGVMEALGTLPGHMTERFQEAIDNLAVLMRAETGSRPYKAAAAQLPETLETIMLLGLLGTVSLGIFFVLMRNKGIGKMGFGMVTLGASAWLMWLSEIEPARIACVLIVVFLLLVVLIPEPEKQRSPQDNQMAIIIMVAVGLLGLITANELGWLERTKSDLSHLMGRREEGATIGFSMDIDLRPASAWAIYAALTTFITPAVQHAVTTSYNNYSLMAMATQAGVLFGMGKGMPFYAWDFGVPLLMIGCYSQLTPLTLIVAIILLVAHYMYLIPGLQAAAARAAQKRTAAGIMKNPVVDGIVVTDIDTMTIDPQVEKKMGQVLLIAVAVSSAILSRTAWGWGEAGALITAATSTLWEGSPNKYWNSSTATSLCNIFRGSYLAGASLIYTVTRNAGLVKRRGGGTGETLGEKWKARLNQMSALEFYSYKKSGITEVCREEARRALKDGVATGGHAVSRGSAKLRWLVERGYLQPYGKVIDLGCGRGGWSYYAATIRKVQEVKGYTKGGPGHEEPMLVQSYGWNIVRLKSGVDVFHMAAEPCDTLLCDIGESSSSPEVEEARTLRVLSMVGDWLEKRPGAFCIKVLCPYTSTMMETLERLQRRYGGGLVRVPLSRNSTHEMYWVSGAKSNTIKSVSTTSQLLLGRMDGPRRPVKYEEDVNLGSGTRAVVSCAEAPNMKIIGNRIERIRSEHAETWFFDENHPYRTWAYHGSYEAPTQGSASSLINGVVRLLSKPWDVVTGVTGIAMTDTTPYGQQRVFKEKVDTRVPDPQEGTRQVMSMVSSWLWKELGKHKRPRVCTKEEFINKVRSNAALGAIFEEEKEWKTAVEAVNDPRFWALVDKEREHHLRGECQSCVYNMMGKREKKQGEFGKAKGSRAIWYMWLGARFLEFEALGFLNEDHWMGRENSGGGVEGLGLQRLGYVLEEMSRIPGGRMYADDTAGWDTRISRFDLENEALITNQMEKGHRALALAIIKYTYQNKVVKVLRPAEKGKTVMDIISRQDQRGSGQVVTYALNTFTNLVVQLIRNMEAEEVLEMQDLWLLRRSEKVTNWLQSNGWDRLKRMAVSGDDCVVKPIDDRFAHALRFLNDMGKVRKDTQEWKPSTGWDNWEEVPFCSHHFNKLHLKDGRSIVVPCRHQDELIGRARVSPGAGWSIRETACLAKSYAQMWQLLYFHRRDLRLMANAICSSVPVDWVPTGRTTWSIHGKGEWMTTEDMLVVWNRVWIEENDHMEDKTPVTKWTDIPYLGKREDLWCGSLIGHRPRTTWAENIKNTVNMVRRIIGDEEKYMDYLSTQVRYLGEEGSTPGVL

>tr|A0A109PRQ3|A0A109PRQ3_ZIKV Genome polyprotein OSZika virus (strain Mr 766) PE4 SV1

MKNPKKKSGGFRIVNMLKRGVARVSPFGGLKRLPAGLLLGHGPIRMVLAILAFLRFTAIKPSLGLINRWGSVGKKEAMEIIKKFKKDLAAMLRIINARKEKKRRGADTSVGIVGLLLTTAMAAEVTRRGSAYYMYLDRNDAGEAISFPTTLGMNKCYIQIMDLGHMCDATMSYECPMLDEGVEPDDVDCWCNTTSTWVVYGTCHHKKGEARRSRRAVTLPSHSTRKLQTRSQTWLESREYTKHLIRVENWIFRNPGFALAAAAIAWLLGSSTSQKVIYLVMILLIAPAYSIRCIGVSNRDFVEGMSGGTWVDVVLEHGGCVTVMAQDKPTVDIELVTTTVSNMAEVRSYCYEASISDMASDSRCPTQGEAYLDKQSDTQYVCKRTLVDRGWGNGCGLFGKGSLVTCAKFACSKKMTGKSIQPENLEYRIMLSVHGSQHSGMIVNDTGHETDENRAKVEITPNSPRAEATLGGFGSLGLDCEPRTGLDFSDLYYLTMNNKHWLVHKEWFHDIPLPWHAGADTGTPHWNNKEALVEFKDAHAKRQTVVVLGSQEGAVHTALAGALEAEMDGAKGRLSSGHLKCRLKMDKLRLKGVSYSLCTAAFTFTKIPAETLHGTVTVEVQYAGTDGPCKVPAQMAVDMQTLTPVGRLITANPVITESTENSKMMLELDPPFGDSYIVIGVGEKKITHHWHRSGSTIGKAFEATVRGAKRMAVLGDTAWDFGSVGGALNSLGKGIHQIFGAAFKSLFGGMSWFSQILIGTLLMWLGLNTKNGSISLMCLALGGVLIFLSTAVSADVGCSVDFSKKETRCGTGVFVYNDVEAWRDRYKYHPDSPRRLAAAVKQAWEDGICGISSVSRMENIMWRSVEGELNAILEENGVQLTVVVGSVKNPMWRGPQRLPVPVNELPHGWKAWGKSYFVRAAKTNNSFVVDGDTLKECPLKHRAWNSFLVEDHGFGVFHTSVWLKVREDYSLECDPAVIGTAVKGKEAVHSDLGYWIESEKNDTWRLKRAHLIEMKTCEWPKSHTLWTDGIEESDLIIPKSLAGPLSHHNTREGYRTQMKGPWHSEELEIRFEECPGTKVHVEETCGTRGPSLRSTTASGRVIEEWCCRECTMPPLSFRAKDGCWYGMEIRPRKEPESNLVRSMVTAGSTDHMDHFSLGVLVILLMVQEGLKKRMTTKIIISTSMAVLVAMILGGFSMSDLAKLAILMGATFAEMNTGGDVAHLALIAAFKVRPALLVSFIFRANWTPRESMLLALASCLLQTAISALEGDLMVLINGFALAWLAIRAMVVPRTDNITLAILAALTPLARGTLLVAWRAGLATCGGFMLLSLKGKGSVKKNLPFVMALGLTAVRLVDPINVVGLLLLTRSGKRSWPPSEVLTAVGLICALAGGFAKADIEMAGPMAAVGLLIVSYVVSGKSVDMYIERAGDITWEKDAEVTGNSPRLDVALDESGDFSLVEDDGPPMREIILKVVLMTICGMNPIAIPFAAGAWYVYVKTGKRSGALWDVPAPKEVKKGETTDGVYRVMTRRLLGSTQVGVGVMQEGVFHTMWHVTKGSALRSGEGRLDPYWGDVKQDLVSYCGPWKLDAAWDGHSEVQLLAVPPGERARNIQTLPGIFKTKDGDIGAVALDYPAGTSGSPILDKCGRVIGLYGNGVVIKNGSYVSAITQGRREEETPVECFEPSMLKKKQLTVLDLHPGAGKTRRVLPEIVREAIKTRLRTVILAPTRVVAAEMEEALRGLPVRYMTTAVNVTHSGTEIVDLMCHATFTSRLLQPIRVPNYNLYIMDEAHFTDPSSIAARGYISTRVEMGEAAAIFMTATPPGTRDAFPDSNSPIMDTEVEVPERAWSSGFDWVTDHSGKTVWFVPSVRNGNEIAACLTKAGKRVIQLSRKTFETEFQKTKHQEWDFVVTTDISEMGANFKADRVIDSRRCLKPVILDGERVILAGPMPVTHASAAQRRGRIGRNPNKPGDEYLYGGGCAETDEDHAHWLEARMLLDNIYLQDGLIASLYRPEADKVAAIEGEFKLRTEQRKTFVELMKRGDLPVWLAYQVASAGITYTDRRWCFDGTTNNTIMEDSVPAEVWTRHGEKRVLKPRWMDARVCSDHAALKSFKEFAAGKRGAAFGVMEALGTLPGHMTERFQEAIDNLAVLMRAETGSRPYKAAAAQLPETLETIMLLGLLGTVSLGIFFVLMRNKGIGKMGFGMVTLGASAWLMWLSEIEPARIACVLIVVFLLLVVLIPEPEKQRSPQDNQMAIIIMVAVGLLGLITANELGWLERTKSDLSHLMGRREEGATIGFSMDIDLRPASAWAIYAALTTFITPAVQHAVTTSYNNYSLMAMATQAGVLFGMGKGMPFYAWDFGVPLLMIGCYSQLTPLTLIVAIILLVAHYMYLIPGLQAAAARAAQKRTAAGIMKNPVVDGIVVTDIDTMTIDPQVEKKMGQVLLIAVAVSSAILSRTAWGWGEAGALITAATSTLWEGSPNKYWNSSTATSLCNIFRGSYLAGASLIYTVTRNAGLVKRRGGGTGETLGEKWKARLNQMSALEFYSYKKSGITEVCREEARRALKDGVATGGHAVSRGSAKLRWLVERGYLQPYGKVIDLGCGRGGWSYYAATIRKVQEVKGYTKGGPGHEEPVLVQSYGWNIVRLKSGVDVFHMAAEPCDTLLCDIGESSSSPEVEEARTLRVLSMVGDWLEKRPGAFCIKVLCPYTSTMMETLERLQRRYGGGLVRVPLSRNSTHEMYWVSGAKSNTIKSVSTTSQLLLGRMDGPRRPVKYEEDVNLGSGTRAVVSCAEAPNMKIIGNRIERIRSEHAETWFFDENHPYRTWAYHGSYEAPTQGSASSLINGVVRLLSKPWDVVTGVTGIAMTDTTPYGQQRVFKEKVDTRVPDPQEGTRQVMSMVSSWLWKELGKHKRPRVCTKEEFINKVRSNAALGAIFEEEKEWKTAVEAVNDPRFWALVDKEREHHLRGECQSCVYNMMGKREKKQGEFGKAKGSRAIWYMWLGARFLEFEALGFLNEDHWMGRENSGGGVEGLGLQRLGYVLEEMSRIPGGRMYADDTAGWDTRISRFDLENEALITNQMEKGHRALALAIIKYTYQNKVVKVLRPAEKGKTVMDIISRQDQRGSGQVVTYALNTFTNLVVQLIRNMEAEEVLEMQDLWLLRRSEKVTNWLQSNGWDRLKRMAVSGDDCVVKPIDDRFAHALRFLNDMGKVRKDTQEWKPSTGWDNWEEVPFCSHHFNKLHLKDGRSIVVPCRHQDELIGRARVSPGAGWSIRETACLAKSYAQMWQLLYFHRRDLRLMANAICSSVPVDWVPTGRTTWSIHGKGEWMTTEDMLVVWNRVWIEENDHMEDKTPVTKWTDIPYLGKREDLWCGSLIGHRPRTTWAENIKNTVNMVRRIIGDEEKYMDYLSTQVRYLGEEGSTPGVL

>tr|B3U3M3|B3U3M3_ZIKV Genome polyprotein OSZika virus (strain Mr 766) PE4 SV1

MKNPKEEIRRIRIVNMLKRGVARVSPFGGLKRLPAGLLLGHGPIRMVLAILAFLRFTAIKPSLGLINRWGSVGKKEAMEIIKKFKKDLAAMLRIINARKEKKRRGTDTSVGIVGLLLTTAMAVEVTRRGSAYYMYLDRSDAGEAISFPTTLGMNKCYIQIMDLGHMCDATMSYECPMLDEGVEPDDVDCWCNTTSTWVVYGTCHHKKGEARRSRRAVTLPSHSTRKLQTRSQTWLESREYTKHLIRVENWIFRNPGFALAAAAIAWLLGSSTSQKVIYLVMILLIAPAYSIRCIGVSNRDFVEGMSGGTWVDVVLEHGGCVTVMAQDKPAVDIELVTTTVSNMAEVRSYCYEASISDMASDSRCPTQGEAYLDKQSDTQYVCKRTLVDRGWGNGCGLFGKGSLVTCAKFACSKKMTGKSIQPENLEYRIMLSVHGSQHSGMIVNDTGHETDENRAKVEITPNSPRAEATLGGFGSLGLDCEPRTGLDFSDLYYLTMNNKHWLVHKEWFHDIPLPWHAGADTGTPHWNNKEALVEFKDAHAKRQTVVVLGSQEGAVHTALAGALEAEMDGAKGRLSSGHLKCRLKMDKLRLKGVSYSLCTAAFTFTKIPAETLHGTVTVEVQYAGTDGPCKVPAQMAVDMQTLTPVGRLITANPVITESTENSKMMLELDPPFGDSYIVIGVGEKKITHHWHRSGSTIGKAFEATVRGAKRMAVLGDTAWDFGSVGGALNSLGKGIHQIFGAAFKSLFGGMSWFSQILIGTLLVWLGLNTKNGSISLTCLALGGVLIFLSTAVSADVGCSVDFSKKETRCGTGVFVYNDVDAWRDRYKYHPDSPRRLAAAVKQAWXDGICGISSVSRMENIMWRSVEGELNAILEENGVQLTVVVGSVKNPMWRGPQRLPVPVNELPHGWKAWGKSYFVRAAKTNNSFVVDGDTLKECPLKHRAWNSFLVEDHGFGVFHTSVWLKVREDYSLECDPAVIGTAAKGKEAVHSDLGYWIESEKNDTWRLKRAHLIEMKTCEWPKSHTLWTDGIEESDLIIPKSLAGPLSHHNTREGYRTQMKGPWHSEELEIRFEECPGTKVHVEETCGTRGPSLRSTTASGRVIEEWCCRECTMPPLSFRAKDGCWYGMEIRPRKEPESNLVRSMVTAGSTDHMDHFSLGVLVILLMVQEGLKKRMTTKIIISTSMAVLVAMILGGFSMSDLAKLAILMGATFAEMNTGGDVAHLALIAAFKVRPALLVSFIFRANWTPRESMLLALASCLLQTAISALEGDLMVLINGFALAWLAIRAMVVPRTDNITLAILTALTPLARGTLLVAWRAGLATCGGFMLLSLKGKGSVKKNLPFVMALGLTAVRLVDPINVVGLLLLTRSGKRSWPPSEVLTAVGLICALAGGFAKADIEMAGPMAAVGLLIVSYVVSGKSVDMYIERAGDITWEKDAEVTGNSPRLDVALDESGDFSLVEDDGPPMREIILKVVLMTICGMNPIAIPFAAGAWYVYVKTGKRSGALWDVPAPKEVKKGETTDGVYRVMTRRLLGSTQVGVGVMQEGVFHTMWHVTKGSALRSGEGRLDPYWGDVKQDLVSYCGPWKLDAAWDGHSEVQLLAVPPGERARNIQTPPGIFKTKDGDIGAVALDYPAGTSGSPILDKCGRVIGLYGNGVVIKNGSYVSAITQGRREEETPVECFEPSMLKKKQLTVLDLHPGAGKTRRVLPEIVREAIKTRLRTVILAPTRVVAAEMEEALRGLPVRYMTTAVNVTHSGTEIVDLMCHATFTSRLLQPIRVPNYNLYIMDEAHFTDPSSIAARGYISTRVEMGEAAAIFMTATPPGTRDAFPDSNSPIMDTEVEVPERAWSSGFDWVTDHSGKTVWFVPSVRNGNEIAACLTKAGKRVIQLSRKTFETEFLKTKNQEWDFVVTTDISEMGANFKADRVIDSRRCLKPVILDGERVILAGPMPVTHASAAQRRGRIGRNPNKPGDEYLYGGGCAETDEDHAHWLEARMLLDNIYLQDGLIASLYRPEADKVAAIEGEFKLRTEQRKTFVELMKRGDLPVWLAYQVASAGITYTDRKWCFDGTTNNTIMEDSVPAEVWTRYGEKRVLKPRWMDARVCSDHAALKSFKEFAAGKRGXAFGVMEALGTLPGHMTERFQEAIDNLAVLMRAETGSRPYEAAAAQLPETLETIMLLGLLGTVSLGIFFVLMRNKGIGKMGFGMVTLGASAWLMWLSEIEPARIACVLIVVFLLLVVLIPEPEKQRSPQDNQMAIIIMIAVGLLGLITANELGWLERTKSDLSHLMGRREEGATIGFSMDIDLRPASAWAIYAALTTFITPAVQHAVTTSYNNYSLMAMATQAGVLFGMGKGMPFYAWDFGVPLLMMGCYSQLTPLTLIVAIILLVAHYMYLIPGLQAAAARAAQKRTAAGIMKNPVVDGIVVTDIDTMTIDHRVEKKMGQVLLIAVAVSSAILSRTAWGWGEAGALITAATSTLWEGSPNKYWNSSTATSLCNIFRGSYLAGASLIYTVTRNAGLVKRRGGGTGETLGEKWKARLNQMSALEFYSYKKSGITEVCREEARRALKDGVATGGHAVSRGSAKLRWLVERGYLQPYGKVIDLGCGRGGWSYYAATIRKVQEVKGYTKGGPGHEEPMLVQSYGWNIVRLKSGVDVFHMAAEPCDTLLCDIGESSSSPEVEEARTLRVLSMVGDWLEKRPGAFCIKVLCPYTSTMMETLERLQRRYGGGLVRVPLSRNSTHEMYWVSGAKSNTIKSVSTTSQLLLGRMDGPRRPVKYEEDVNLGSGTRAVVSCAEAPNMKIIGNRIERIRSEHAETWFFDENHPYRTWAYHGSYEAPTQGSASSLINGVVRLLSKPWDVVTGVTGIAMTDTTPYGQQRVFKEKVDTRVPDPQEGTRQVMSMVSSWLWKELGKHKRPRVCTKEEFINKVRSNAALGAIFEEEKEWKTAVEAVNDPRFWALVDKEREHHLRGECQSCVYNMMGKREKKQGEFGKAKGSRAIWYMWLGARFLEFEALGFLNEDHWMGRENSGGGVEGLGLQRLGYVLEEMSRIPGGRMYADDTAGWDTRISRFDLENEALITNQMEKGHRALALAIIKYTYQNKVVKVLRPAEKGKTVMDIISRQDQRGSGQVVTYALNTFTNLVVQLIRNMEAEEVLEMQDLWLLRRSEKVTNWLQSNGWDRLKRMAVSGDDCVVKPIDDRFAHALRFLNDMGKVRKDTQEWKPSTGWDNWEEVPFCSHHFNKLHLKDGRSIVVPCRHQDELIGRARVSPGAGWSIRETACLAKSYAQMWQLLYFHRRDLRLMANAICSSVPVDWVPTGRTTWSIHGKGEWMTTEDMLVVWNRVWIEENDHMEDKTPVTKWTDIPYLGKREDLWCGSLIGHRPRTTWAENIKNTVNMMRRIIGDEEKYMDYLSTQVRYLGEEGSTPGVL

>tr|A0A0X9QZM7|A0A0X9QZM7_ZIKV Genome polyprotein OSZika virus (strain Mr 766) PE4 SV1

MKNPKKKSGGFRIVNMLKRGVARVSPFGGLKRLPAGLLLGHGPIRMVLAILAFLRFTAIKPSLGLINRWGSVGKKEAMEIIKKFKKDLAAMLRIINARKEKKRRGADTSVGIVGLLLTTAMAAEVTRRGSAYYMYLDRNDAGEAISFPTTLGMNKCYIQIMDLGHMCDATMSYECPMLDEGVEPDDVDCWCNTTSTWVVYGTCHHKKGEARRSRRAVTLPSHSTRKLQTRSQTWLESREYTKHLIRVENWIFRNPGFALAAAAIAWLLGSSTSQKVIYLVMILLIAPAYSIRCIGVSNRDFVEGMSGGTWVDVVLEHGGCVTVMAQDKPTVDIELVTTTVSNMAEVRSYCYEASISDMASDSRCPTQGEAYLDKQSDTQYVCKRTLVDRGWGNGCGLFGKGSLVTCAKFACSKKMTGKSIQPENLEYRIMLSVHGSQHSGMIVNDTGHETDENRAKVEITPNSPRAEATLGGFGSLGLDCEPRTGLDFSDLYYLTMNNKHWLVHKEWFHDIPLPWHAGADTGTPHWNNKEALVEFKDAHAKRQTVVVLGSQEGAVHTALAGALEAEMDGAKGRLSSGHLKCRLKMDKLRLKGVSYSLCTAAFTFTKIPAETLHGTVTVEVQYAGTDGPCKVPAQMAVDMQTLTPVGRLITANPVITESTENSKMMLELDPPFGDSYIVIGVGEKKITHHWHRSGSTIGKAFEATVRGAKRMAVLGDTAWDFGSVGGALNSLGKGIHQIFGAAFKSLFGGMSWFSQILIGTLLMWLGLNTKNGSISLMCLALGGVLIFLSTAVSADVGCSVDFSKKETRCGTGVFVYNDVEAWRDRYKYHPDSPRRLAAAVKQAWEDGICGISSVSRMENIMWRSVEGELNAILEENGVQLTVVVGSVKNPMWRGPQRLPVPVNELPHGWKAWGKSYFVRAAKTNNSFVVDGDTLKECPLKHRAWNSFLVEDHGFGVFHTSVWLKVREDYSLECDPAVIGTAVKGKEAVHSDLGYWIESEKNDTWRLKRAHLIEMKTCEWPKSHTLWTDGIEESDLIIPKSLAGPLSHHNTREGYRTQMKGPWHSEELEIRFEECPGTKVHVEETCGTRGPSLRSTTASGRVIEEWCCRECTMPPLSFRAKDGCWYGMEIRPRKEPESNLVRSMVTAGSTDHMDHFSLGVLVILLMVQEGLKKRMTTKIIISTSMAVLVAMILGGFSMSDLAKLAILMGATFAEMNTGGDVAHLALIAAFKVRPALLVSFIFRANWTPRESMLLALASCLLQTAISALEGDLMVLINGFALAWLAIRAMVVPRTDNITLAILAALTPLARGTLLVAWRAGLATCGGFMLLSLKGKGSVKKNLPFVMALGLTAVRLVDPINVVGLLLLTRSGKRSWPPSEVLTAVGLICALAGGFAKADIEMAGPMAAVGLLIVSYVVSGKSVDMYIERAGDITWEKDAEVTGNSPRLDVALDESGDFSLVEDDGPPMREIILKVVLMTICGMNPIAIPFAAGAWYVYVKTGKRSGALWDVPAPKEVKKGETTDGVYRVMTRRLLGSTQVGVGVMQEGVFHTMWHVTKGSALRSGEGRLDPYWGDVKQDLVSYCGPWKLDAAWDGHSEVQLLAVPPGERARNIQTLPGIFKTKDGDIGAVALDYPAGTSGSPILDKCGRVIGLYGNGVVIKNGSYVSAITQGRREEETPVECFEPSMLKKKQLTVLDLHPGAGKTRRVLPEIVREAIKTRLRTVILAPTRVVAAEMEEALRGLPVRYMTTAVNVTHSGTEIVDLMCHATFTSRLLQPIRVPNYNLYIMDEAHFTDPSSIAARGYISTRVEMGEAAAIFMTATPPGTRDAFPDSNSPIMDTEVEVPERAWSSGFDWVTDHSGKTVWFVPSVRNGNEIAACLTKAGKRVIQLSRKTFETEFQKTKHQEWDFVVTTDISEMGANFKADRVIDSRRCLKPVILDGERVILAGPMPVTHASAAQRRGRIGRNPNKPGDEYLYGGGCAETDEDHAHWLEARMLLDNIYLQDGLIASLYRPEADKVAAIEGEFKLRTEQRKTFVELMKRGDLPVWLAYQVASAGITYTDRRWCFDGTTNNTIMEDSVPAEVWTRHGEKRVLKPRWMDARVCSDHAALKSFKEFAAGKRGAAFGVMEALGTLPGHMTERFQEAIDNLAVLMRAETGSRPYKAAAAQLPETLETIMLLGLLGTVSLGIFFVLMRNKGIGKMGFGMVTLGASAWLMWLSEIEPARIACVLIVVFLLLVVLIPEPEKQRSPQDNQMAIIIMVAVGLLGLITANELGWLERTKSDLSHLMGRREEGATIGFSMDIDLRPASAWAIYAALTTFITPAVQHAVTTSYNNYSLMAMATQAGVLFGMGKGMPFYAWDFGVPLLMIGCYSQLTPLTLIVAIILLVAHYMYLIPGLQAAAARAAQKRTAAGIMKNPVVDGIVVTDIDTMTIDPQVEKKMGQVLLIAVAVSSAILSRTAWGWGEAGALITAATSTLWEGSPNKYWNSSTATSLCNIFRGSYLAGASLIYTVTRNAGLVKRRGGGTGETLGEKWKARLNQMSALEFYSYKKSGITEVCREEARRALKDGVATGGHAVSRGSAKLRWLVERGYLQPYGKVIDLGCGRGGWSYYAATIRKVQEVKGYTKGGPGHEEPVLVQSYGWNIVRLKSGVDVFHMAAEPCDTLLCDIGESSSSPEVEEARTLRVLSMVGDWLEKRPGAFCIKVLCPYTSTMMETLERLQRRYGGGLVRVPLSRNSTHEMYWVSGAKSNTIKSVSTTSQLLLGRMDGPRRPVKYEEDVDLGSGTRAVVSCAEAPNMKIIGNRIERIRSEHAETWFFDENHPYRTWAYHGSYEAPTQGSASSLINGVVRLLSKPWDVVTGVTGIAMTDTTPYGQQRVFKEKVDTRVPDPQEGTRQVMSMVSSWLWKELGKHKRPRVCTKEEFINKVRSNAALGAIFEEEKEWKTAVEAVNDPRFWALVDKEREHHLRGECQSCVYNMMGKREKKQGEFGKAKGSRAIWYMWLGARFLEFEALGFLNEDHWMGRENSGGGVEGLGLQRLGYVLEEMSRIPGGRMYADDTAGWDTRISRFDLENEALITNQMEKGHRALALAIIKYTYQNKVVKVLRPAEKGKTVMDIISRQDQRGSGQVVTYALNTFTNLVVQLIRNMEAEEVLEMQDLWLLRRSEKVTNWLQSNGWDRLKRMAVSGDDCVVKPIDDRFAHALRFLNDMGKVRKDTQEWKPSTGWDNWEEVPFCSHHFNKLHLKDGRSIVVPCRHQDELIGRARVSPGAGWSIRETACLAKSYAQMWQLLYFHRRDLRLMANAICSSVPVDWVPTGRTTWSIHGKGEWMTTEDMLVVWNRVWIEENDHMEDKTPVTKWTDIPYLGKREDLWCGSLIGHRPRTTWAENIKNTVNMVRRIIGDEEKYMDYLSTQVRYLGEEGSTPGVL

>tr|A0A120IF34|A0A120IF34_ZIKV Genome polyprotein OSZika virus (strain Mr 766) PE4 SV1

MKNPKKKSGGFRIVNMLKRGVARVSPFGGLKRLPAGLLLGHGPIRMVLAILAFLRFTAIKPSLGLINRWGSVGKKEAMEIIKKFKKDLAAMLRIINARKEKKRRGADTSVGIVGLLLTTAMAAEVTRRGSAYYMYLDRNDAGEAISFPTTLGMNKCYIQIMDLGHMCDATMSYECPMLDEGVEPDDVDCWCNTTSTWVVYGTCHHKKGEARRSRRAVTLPSHSTRKLQTRSQTWLESREYTKHLIRVENWIFRNPGFALAAAAIAWLLGSSTSQKVIYLVMILLIAPAYSIRCIGVSNRDFVEGMSGGTWVDVVLEHGGCVTVMAQDKPTVDIELVTTTVSNMAEIRSYCYEASISDMASDSRCPTQGEAYLDKQSDTQYVCKRTLVDRGWGNGCGLFGKGSLVTCAKFACSKKMTGKSIQPENLEYRIMLSVHGSQHSGMIVNDTGHETDENRAKVEITPNSPRAEATLGGFGSLGLDCEPRTGLDFSDLYYLTMNNKHWLVHKEWFHDIPLPWHAGADTGTPHWNNKEALVEFKDAHAKRQTVVVLGSQEGAVHTALAGALEAEMDGAKGRLSSGHLKCRLKMDKLRLKGVSYSLCTAAFTFTKIPAETLHGTVTVEVQYAGTDGPCKVPAQMAVDMQTLTPVGRLITANPVITESTENSKMMLELDPPFGDSYIVIGVGEKKITHHWHRSGSTIGKAFEATVRGAKRMAVLGDTAWDFGSVGGALNSLGKGIHQIFGAAFKSLFGGMSWFSQILIGTLLMWLGLNTKNGSISLMCLALGGVLIFLSTAVSADVGCSVDFSKKETRCGTGVFVYNDVEAWRDRYKYHPDSPRRLAAAVKQAWEDGICGISSVSRMENIMWRSVEGELNAILEENGVQLTVVVGSVKNPMWRAPQRLPVPVNELPHGWKAWGKSYFVRAAKTNNSFVVDGDTLKECPLKHRAWNSFLVEDHGFGVFHTSVWLKVREDYSLECDPAVIGTAVKGKEAVHSDLGYWIESEKNDTWRLKRAHLIEMKTCEWPKSHTLWTDGIEESDLIIPKSLAGPLSHHNTREGYRTQMKGPWHSEELEIRFEECPGTKVHVEETCGTRGPSLRSTTASGRVIEEWCCRECTMPPLSFRAKDGCWYGMEIRPRKEPESNLVRSMVTAGSTDHMDHFSLGVLVILLMVQEGLKKRMTTKIIISTSMAVLVAMILGGFSMSDLAKLAILMGATFAEMNTGGDVAHLALIAAFKVRPALLVSFIFRANWTPRESMLLALASCLLQTAISALEGDLMVLINGFALAWLAIRAMVVPRTDNITLAILAALTPLARGTLLVAWRAGLATCGGFMLLSLKGKGSVKKNLPFVMALGLTAVRLVDPINVVGLLLLTRSGKRSWPPSEVLTAVGLICALAGGFAKADIEMAGPMAAVGLLIVSYVVSGKSVDMYIERAGDITWEKDAEVTGNSPRLDVALDESGDFSLVEDDGPPMREIILKVVLMTICGMNPIAIPFAAGAWYVYVKTGKRSGALWDVPAPKEVKKGETTDGVYRVMTRRLLGSTQVGVGVMQEGVFHTMWHVTKGSALRSGEGRLDPYWGDVKQDLVSYCGPWKLDAAWDGHSEVQLLAVPPGERARNIQTLPGIFKTKDGDIGAVALDYPAGTSGSPILDKCGRVIGLYGNGVVIKNGSYVSAITQGRREEETPVECFEPSMLKKKQLTVLDLHPGAGKTRRVLPEIVREAIKTRLRTVILAPTRVVAAEMEEALRGLPVRYMTTAVNVTHSGTEIVDLMCHATFTSRLLQPIRVPNYNLYIMDEAHFTDPSSIAARGYISTRVEMGEAAAIFMTATPPGTRDAFPDSNSPIMDTEVEVPERAWSSGFDWVTDHSGKTVWFVPSVRNGNEIAACLTKAGKRVIQLSRKTFETEFQKTKHQEWDFVVTTDISEMGANFKADRVIDSRRCLKPVILDGERVILAGPMPVTHASAAQRRGRIGRNPNKPGDEYLYGGGCAETDEDHAHWLEARMLLDNIYLQDGLIASLYRPEADKVAAIEGEFKLRTEQRKTFVELMKRGDLPVWLAYQVASAGITYTDRRWCFDGTTNNTILEDSVPAEVWTRHGEKRVLKPRWMDARVCSDHAALKSFKEFAAGKRGAAFGVMEALGTLPGHMTERFQEAIDNLAVLMRAETGSRPYKAAAAQLPETLETIMLLGLLGTVSLGIFFVLMRNKGIGKMGFGMVTLGASAWLMWLSEIEPARIACVLIVVFLLLVVLIPEPEKQRSPQDNQMAIIIMVAVGLLGLITANELGWLERTKSDLSHLMGRREEGATIGFSMDIDLRPASAWAIYAALTTFITPAVQHAVTTSYNNYSLMAMATQAGVLFGMGKGMPFYAWDFGVPLLMIGCYSQLTPLTLIVAIILLVAHYMYLIPGLQAAAARAAQKRTAAGIMKNPVVDGIVVTDIDTMTIDPQVEKKMGQVLLIAVAVSSAILSRTAWGWGEAGALITAATSTLWEGSPNKYWNSSTATSLCNIFRGSYLAGASLIYTVTRNAGLVKRRGGGTGETLGEKWKARLNQMSALEFYSYKKSGITEVCREEARRALKDGVATGGHAVSRGSAKLRWLVERGYLQPYGKVIDLGCGRGGWSYYAATIRKVQEVKGYTKGGPGHEEPVLVQSYGWNIVRLKSGVDVFHMAAEPCDTLLCDIGESSSSPEVEEARTLRVLSMVGDWLERRPGAFCIKVLCPYTSTMMETLERLQRRYGGGLVRVPLSRNSTHEMYWVSGAKSNTIKSVSTTSQLLLGRMDGPRRPVKYEEDVNLGSGTRAVVSCAEAPNMKIIGNRIERIRSEHAETWFFDENHPYRTWAYHGSYEAPTQGSASSLINGVVRLLSKPWDVVTGVTGIAMTDTTPYGQQRVFKEKVDTRVPDPQEGTRQVMSMVSSWLWKELGKHKRPRVCTKEEFINKVRSNAALGAIFEEEKEWKTAVEAVNDPRFWALVDKEREHHLRGECQSCVYNMMGKREKKQGEFGKAKGSRAIWYMWLGARFLEFEALGFLNEDHWMGRENSGGGVEGLGLQRLGYVLEEMSCIPGGRMYADDTAGWDTRISRFDLENEALITNQMEKGHRALALAIIKYTYQNKVVKVLRPAEKGKTVMDIISRQDQRGSGQVVTYALNTFTNLVVQLIRNMEAEEVLEMQDLWLLRRSEKVTNWLQSNGWDRLKRMAVSGDDCVVKPIDDRFAHALRFLNDMGKVRKDTQEWKPSTGWDNWEEVPFCSHHFNKLHLKDGRSIVVPCRHQDELIGRARVSPGAGWSIRETACLAKSYAQMWQLLYFHRRDLRLMANAICSSVPVDWVPTGRTTWSIHGKGEWMTTEDMLVVWNRVWIEENDHMEDKTPVTKWTDIPYLGKREDLWCGSLIGHRPRTTWAENIKNTVNMVRRIIGDEEKYMDYLSTQVRYLGEEGSTPGVL

>tr|A0A110ASP6|A0A110ASP6_ZIKV Genome polyprotein OSZika virus (strain Mr 766) PE4 SV1

MKNPKKKSGGFRIVNMLKRGVARVSPFGGLKRLPAGLLLGHGPIRMVLAILAFLRFTAIKPSLGLINRWGSVGKKEAMEIIKKFKKDLAAMLRIINARKEKKRRGADTSVGIVGLLLTTAMAAEVTRRGSAYYMYLDRNDAGEAISFPTTLGMNKCYIQIMDLGHMCDATMSYECPMLDEGVEPDDVDCWCNTTSTWVVYGTCHHKKGEARRSRRAVTLPSHSTRKLQTRSQTWLESREYTKHLIRVENWIFRNPGFALAAAAIAWLLGSSTSQKVIYLVMILLIAPAYSIRCIGVSNRDFVEGMSGGTWVDVVLEHGGCVTVMAQDKPTVDIELVTTTVSNMAEVRSYCYEASISDMASDSRCPTQGEAYLDKQSDTQYVCKRTLVDRGWGNGCGLFGKGSLVTCAKFACSKKMTGKSIQPENLEYRIMLSVHGSQHSGMIVNDTGHETDENRAKVEITPNSPRAEATLGGFGSLGLDCEPRTGLDFSDLYYLTMNNKHWLVHKEWFHDIPLPWHAGADTGTPHWNNKEALVEFKDAHAKRQTVVVLGSQEGAVHTALAGALEAEMDGAKGRLSSGHLKCRLKMDKLRLKGVSYSLCTAAFTFTKIPAETLHGTVTVEVQYAGTDGPCKVPAQMAVDMQTLTPVGRLITANPVITESTENSKMMLELDPPFGDSYIVIGVGEKKITHHWHRSGSTIGKAFEATVRGAKRMAVLGDTAWDFGSVGGALNSLGKGIHQIFGAAFKSLFGGMSWFSQILIGTLLMWLGLNTKNGSISLMCLALGGVLIFLSTAVSADVGCSVDFSKKETRCGTGVFVYNDVEAWRDRYKYHPDSPRRLAAAVKQAWEDGICGISSVSRMENIMWRSVEGELNAILEENGVQLTVVVGSVKNPMWRGPQRLPVPVNELPHGWKAWGKSHFVRAAKTNNSFVVDGDTLKECPLKHRAWNSFLVEDHGFGVFHTSVWLKVREDYSLECDPAVIGTAVKGKEAVHSDLGYWIESEKNDTWRLKRAHLIEMKTCEWPKSHTLWTDGIEESDLIIPKSLAGPLSHHNTREGYRTQMKGPWHSEELEIRFEECPGTKVHVEETCGTRGPSLRSTTASGRVIEEWCCRECTMPPLSFRAKDGCWYGMEIRPRKEPESNLVRSMVTAGSTDHMDHFSLGVLVILLMVQEGLKKRMTTKIIISTSMAVLVAMILGGFSMSDLAKLAILMGATFAEMNTGGDVAHLALIAAFKVRPALLVSFIFRANWTPRESMLLALASCLLQTAISALEGDLMVLINGFALAWLAIRAMVVPRTDNITLAILAALTPLARGTLLVAWRAGLATCGGFMLLSLKGKGSVKKNLPFVMALGLTAVRLVDPINVVGLLLLTRSGKRSWPPSEVLTAVGLICALAGGFAKADIEMAGPMAAVGLLIVSYVVSGKSVDMYIERAGDITWEKDAEVTGNSPRLDVALDESGDFSLVEDDGPPMREIILKVVLMTICGMNPIAIPFAAGAWYVYVKTGKRSGALWDVPAPKEVKKGETTDGVYRVMTRRLLGSTQVGVGVMQEGVFHTMWHVTKGSALRSGEGRLDPYWGDVKQDLVSYCGPWKLDAAWDGHSEVQLLAVPPGERARNIQTLPGIFKTKDGDIGAVALDYPAGTSGSPILDKCGRVIGLYGNGVVIKNGSYVSAITQGRREEETPVECFEPSMLKKKQLTVLDLHPGAGKTRRVLPEIVREAIKTRLRTVILAPTRVVAAEMEEALRGLPVRYMTTAVNVTHSGTEIVDLMCHATFTSRLLQPIRVPNYNLYIMDEAHFTDPSSIAARGYISTRVEMGEAAAIFMTATPPGTRDAFPDSNSPIMDTEVEVPERAWSSGFDWVTDYSGKTVWFVPSVRNGNEIAACLTKAGKRVIQLSRKTFETEFQKTKHQEWDFVVTTDISEMGANFKADRVIDSRRCLKPVILDGERVILAGPMPVTHASAAQRRGRIGRNPNKPGDEYLYGGGCAETDEDHAHWLEARMLLDNIYLQDGLIASLYRPEADKVAAIEGEFKLRTEQRKTFVELMKRGDLPVWLAYQVASAGITYTDRRWCFDGTTNNTIMEDSVPAEVWTRHGEKRVLKPRWMDARVCSDHAALKSFKEFAAGKRGAAFGVMEALGTLPGHMTERFQEAIDNLAVLMRAETGSRPYKAAAAQLPETLETIMLLGLLGTVSLGIFFVLMRNKGIGKMGFGMVTLGASAWLMWLSEIEPARIACVLIVVFLLLVVLIPEPEKQRSPQDNQMAIIIMVAVGLLGLITANELGWLERTKSDLSHLMGRREEGATMGFSMDIDLRPASAWAIYAALTTFITPAVQHAVTTSYNNYSLMAMATQAGVLFGMGKGMPFYAWDFGVPLLMIGCYSQLTPLTLIVAIILLVAHYMYLIPGLQAAAARAAQKRTAAGIMKNPVVDGIVVTDIDTMTIDPQVEKKMGQVLLMAVAVSSAILSRTAWGWGEAGALITAATSTLWEGSPNKYWNSSTATSLCNIFRGSYLAGASLIYTVTRNAGLVKRRGGGTGETLGEKWKARLNQMSALEFYSYKKSGITEVCREEARRALKDGVATGGHAVSRGSAKLRWLVERGYLQPYGKVIDLGCGRGGWSYYAATIRKVQEVKGYTKGGPGHEEPVLVQSYGWNIVRLKSGVDVFHMAAEPCDTLLCDIGESSSSPEVEEARTLRVLSMVGDWLEKRPGAFCIKVLCPYTSTMMETLERLQRRYGGGLVRVPLSRNSTHEMYWVSGAKSNTIKSVSTTSQLLLGRMDGPRRPVKYEEDVNLGSGTRAVVSCAEAPNMKIIGNRIERIRSEHAETWFFDENHPYRTWAYHGSYEAPTQGSASSLINGVVRLLSKPWDVVTGVTGIAMTDTTPYGQQRVFKEKVDTRVPDPQEGTRQVMSMVSSWLWKELGKHKRPRVCTKEEFINKVRSNAALGAIFEEEKEWKTAVEAVNDPRFWALVDKEREHHLRGECQSCVYNMMGKREKKQGEFGKAKGSRAIWYMWLGARFLEFEALGFLNEDHWMGRENSGGGVEGLGLQRLGYVLEEMSRIPGGRMYADDTAGWDTRISRFDLENEALITNQMEKGHRALALAIIKYTYQNKVVKVLRPAEKGKTVMDIISRQDQRGSGQVVTYALNTFTNLVVQLIRNMEAEEVLEMQDLWLLRRSEKVTNWLQSNGWDRLKRMAVSGDDCVVKPIDDRFAHALRFLNDMGKVRKDTQEWKPSTGWDNWEEVPFCSHHFNKLHLKDGRSIVVPCRHQDELIGRARVSPGAGWSIRETACLAKSYAQMWQLLYFHRRDLRLMANAICSSVPVDWVPTGRTTWSIHGKGEWMTTEDMLVVWNRVWIEENDHMEDKTPVTKWTDIPYLGKREDLWCGSLIGHRPRTTWAENIKNTVNMVRRIIGDEEKYMDYLSTQVRYLGEEGSTPGVL

>tr|A0A0X8GJ44|A0A0X8GJ44_ZIKV Genome polyprotein OSZika virus (strain Mr 766) PE4 SV1

MKNPKKKSGGFRIVNMLKRGVARVSPFGGLKRLPAGLLLGHGPIRMVLAILAFLRFTAIKPSLGLINRWGSVGKKEAMEIIKKFKKDLAAMLRIINARKEKKRRGAETSVGIVGLLLTTAMAAEVTRRGSAYYMYLDRNDAGEAISFPTTLGMNKCYIQIMDLGHMCDATMSYECPMLDEGVEPDDVDCWCNTTSTWVVYGTCHHKKGEARRSRRAVTLPSHSTRKLQTRSQTWLESREYTKHLIRVENWIFRNPGFALAAAAIAWLLGSSTSQKVIYLVMILLIAPAYSIRCIGVSNRDFVEGMSGGTWVDVVLEHGGCVTVMAQDKPTVDIELVTTTVSNMAEVRSYCYEASISDMASDSRCPTQGEAYLDKQSDTQYVCKRTLVDRGWGNGCGLFGKGSLVTCAKFACSKKMTGKSIQPENLEYRIMLSVHGSQHSGMIVNDTGHETDENRAKVEITPNSPRAEATLGGFGSLGLDCEPRTGLDFSDLYYLTMNNKHWLVHKEWFHDIPLPWHAGADTGTPHWNNKEALVEFKDAHAKRQTVVVLGSQEGAVHTALAGALEAEMDGAKGRLSSGHLKCRLKMDKLRLKGVSYSLCTAAFTFTKIPAETLHGTVTVEVQYAGTDGPCKVPAQMAVDMQTLTPVGRLITANPVITESTENSKMMLELDPPFGDSYIVIGVGEKKITHHWHRSGSTIGKAFEATVRGAKRMAVLGDTAWDFGSVGGALNSLGKGIHQIFGAAFKSLFGGMSWFSQILIGTLLMWLGLNTKNGSISLMCLALGGVLIFLSTAVSADVGCSVDFSKKETRCGTGVFVYNDVEAWRDRYKYHPDSPRRLAAAVKQAWEDGICGISSVSRMENIMWRSVEGELNAILEENGVQLTVVVGSVKNPMWRGPQRLPVPVNELPHGWKAWGKSYFVRAAKTNNSFVVDGDTLKECPLKHRAWNSFLVEDHGFGVFHTSVWLKVREDYSLECDPAVIGTAVKGKEAVHSDLGYWIESEKNDTWRLKRAHLIEMKTCEWPKSHTLWTDGIEESDLIIPKSLAGPLSHHNTREGYRTQMKGPWHSEELEIRFEECPGTKVHVEETCGTRGPSLRSTTASGRVIEEWCCRECTMPPLSFWAKDGCWYGMEIRPRKEPESNLVRSMVTAGSTDHMDHFSLGVLVILLMVQEGLKKRMTTKIIISTSMAVLVAMILGGFSMSDLAKLAILMGATFAEMNTGGDVAHLALTAAFKVRPALLVSFIFRANWTPRESMLLALASCLLQTAISALEGDLMVLINGFALAWLAIRAMVVPRTDNITLAILAALTPLARGTLLVAWRAGLATCGGFMLLSLKGKGSVKKNLPFVMALGLTAVRLVDPINVVGLLLLTRSGKRSWPPSEVLTAVGLICALAGGFAKADIEMAGPMAAVGLLIVSYVVSGKSVDMYIERAGDITWEKDAEVTGNSPRLDVALDESGDFSLVEDDGPPMREIILKVVLMTICGMNPIAIPFAAGAWYVYVKTGKRSGALWDVPAPKEVKKGETTDGVYRVMTRRLLGSTQVGVGVMQEGVFHTMWHVTKGSALRSGEGRLDPYWGDVKQDLVSYCGPWKLDAAWDGHSEVQLLAVPPGERARNIQTLPGIFKTKDGDIGAVALDYPAGTSGSPILDKCGRVIGLYGNGVVIKNGSYVSAITQGRREEETPVECFEPSMLKKKQLTVLDLHPGAGKTRRVLPEIVREAIKTRLRTVILAPTRVVAAEMEEALRGLPVRYMTTAVNVTHSGTEIVDLMCHATFTSRLLQPIRVPNYNLYIMDEAHFTDPSSIAARGYISTRVEMGEAAAIFMTATPPGTRDAFPDSNSPIMDTEVEVPERAWSSGFDWVTDHSGKTVWFVPSVRNGNEIAACLTKAGKRVIQLSRKTFETEFQKTKHQEWDFVVTTDISEMGANFKADRVIDSRRCLKPVILDGERVILAGPMPVTHASAAQRRGRIGRNPNKPGDEYLYGGGCAETDEDHAHWLEARMLLDNIYLQDGLIASLYRPEADKVAAIEGEFKLRTEQRKTFVELMKRGDLPVWLAYQVASAGITYTDRRWCFDGTTNNTIMEDSVPAEVWTRHGEKRVLKPRWMDARVCSDHAALKSFKEFAAGKRGAAFGVMEALGTLPGHMTERFQEAIDNLAVLMRAETGSRPYKAAAAQLPETLETIMLLGLLGTVSLGIFFVLMRNKGIGKMGFGMVTLGASAWLMWLSEIEPARIACVLIVVFLLLVVLIPEPEKQRSPQDNQMAIIIMVAVGLLGLITANELGWLERTKSDLSHLMGRREEGATIGFSMDIDLRPASAWAIYAALTTFITPAVQHAVTTSYNNYSLMAMATQAGVLFGMGKGMPFYAWDFGVPLLMIGCYSQLTPLTLIVAIILLVAHYMYLIPGLQAAAARAAQKRTAAGIMKNPVVDGIVVTDIDTMTIDPQVEKKMGQVLLIAVAVSSAILSRTAWGWGEAGALITAATSTLWEGSPNKYWNSSTATSLCNIFRGSYLAGASLIYTVTRNAGLVKRRGGGTGETLGEKWKARLNQMSALEFYSYKKSGITEVCREEARRALKDGVATGGHAVSRGSAKLRWLVERGYLQPYGKVIDLGCGRGGWSYYAATIRKVQEVKGYTKGGPGHEEPVLVQSYGWNIVRLKSGVDVFHMAAEPCDTLLCDIGESSSSPEVEEARTLRVLSMVGDWLEKRPGAFCIKVLCPYTSTMMETLERLQRRYGGGLVRVPLSRNSTHEMYWVSGAKSNTIKSVSTTSQLLLGRMDGPRRPVKYEEDVNLGSGTRAVVSCAEAPNMKIIGNRIERIRSEHAETWFFDENHPYRTWAYHGSYEAPTQGSASSLINGVVRLLSKPWDVVTGVTGIAMTDTTPYGQQRVFKEKVDTRVPDPQEGTRQVMSMVSSWLWKELGKHKRPRVCTKEEFINKVRSNAALGAIFEEEKEWKTAVEAVNDPRFWALVDKEREHHLRGECQSCVYNMMGKREKKQGEFGKAKGSRAIWYMWLGARFLEFEALGFLNEDHWMGRENSGGGVEGLGLQRLGYVLEEMSRIPGGRMYADDTAGWDTRISRFDLENEALITNQMEKGHRALALAIIKYTYQNKVVKVLRPAEKGKTVMDIISRQDQRGSGQVVTYALNTFTNLVVQLIRNMEAEEVLEMQDLWLLRRSEKVTNWLQSNGWDRLKRMAVSGDDCVVKPIDDRFAHALRFLNDMGKVRKDTQEWKPSTGWDNWEEVPFCSHHFNKLHLKDGRSIVVPCRHQDELIGRARVSPGAGWSIRETACLAKSYAQMWQLLYFHRRDLRLMANAICSSVPVDWVPTGRTTWSIHGKGEWMTTEDMLVVWNRVWIEENDHMEDKTPVAKWTDIPYLGKREDLWCGSLIGHRPRTTWAENIKNTVNMVRRIIGDEEKYMDYLSTQVRYLGEEGSTPGVL

>tr|A0A0X1LH65|A0A0X1LH65_ZIKV Genome polyprotein OSZika virus (strain Mr 766) PE4 SV1

MKNPKKKSGGFRIVNMLKRGVARVSPFGGLKRLPAGLLLGHGPIRMVLAILAFLRFTAIKPSLGLINRWGSVGKKEAMETIKKFKKDLAAMLRIINARKEKKRRGADTSVGIVGLLLTTAMAAEVTRRGSAYYMYLDRNDAGEAISFPTTLGMNKCYIQIMDLGHMCDATMSYECPMLDEGVEPDDVDCWCNTTSTWVVYGTCHHKKGEARRSRRAVTLPSHSTRKLQTRSQTWLESREYTKHLIRVENWIFRNPGFALAAAAIAWLLGSSTSQKVIYLVMILLIAPAYSIRCIGVSNRDFVEGMSGGTWVDVVLEHGGCVTVMAQDKPTVDIELVTTTVSNMAEVRSYCYEASISDMASDSRCPTQGEAYLDKQSDTQYVCKRTLVDRGWGNGCGLFGKGSLVTCAKFACSKKMTGKSIQPENLEYRIMLSVHGSQHSGMIVNDTGHETDENRAKVEITPNSPRAEATLGGFGSLGLDCEPRTGLDFSDLYYLTMNNKHWLVHKEWFHDIPLPWHAGADTGTPHWNNKEALVEFKDAHAKRQTVVVLGSQEGAVHTALAGALEAEMDGAKGRLSSGHLKCRLKMDKLRLKGVSYSLCTAAFTFTKIPAETLHGTVTVEVQYAGTDGPCKVPAQMAVDMQTLTPVGRLITANPVITESTENSKMMLELDPPFGDSYIVIGVGEKKITHHWHRSGSTIGKAFEATVRGAKRMAVLGDTAWDFGSVGGALNSLGKGIHQIFGAAFKSLFGGMSWFSQILIGTLLMWLGLNTKNGSISLMCLALGGVLIFLSTAVSADVGCSVDFSKKETRCGTGVFVYNDVEAWRDRYKYHPDSPRRLAAAVKQAWEDGICGISSVSRMENIMWRSVEGELNAILEENGVQLTVVVGSVKNPMWRGPQRLPVPVNELPHGWKAWGKSYFVRAAKTNNSFVVDGDTLKECPLKHRAWNSFLVEDHGFGVFHTSVWLKVREDYSLECDPAVIGTAVKGKEAVHSDLGYWIESEKNDTWRLKRAHLIEMKTCEWPKSHTLWTDGIEESDLIIPKSLAGPLSHHNTREGYRTQMKGPWHSEELEIRFEECPGTKVHVEETCGTRGPSLRSTTASGRVIEEWCCRECTMPPLSFRAKDGCWYGMEIRPRKEPESNLVRSMVTAGSTDHMDHFSLGVLVILLMVQEGLKKRMTTKIIISTSMAVLVAMILGGFSMSDLAKLAILMGATFAEMNTGGDVAHLALIAAFKVRPALLVSFIFRANWTPRESMLLALASCLLQTAISALEGDLMVLINGFALAWLAIRAMVVPRTDNITLAILAALTPLARGTLLVAWRAGLATCGGFMLLSLKGKGSVKKNLPFVMALGLTAVRLVDPINVVGLLLLTRSGKRSWPPSEVLTAVGLICALAGGFAKADIEMAGPMAAVGLLIVSYVVSGKSVDMYIERAGDITWEKDAEVTGNSPRLDVALDESGDFSLVEDDGPPMREIILKVVLMTICGMNPIAIPFAAGAWYVYVKTGKRSGALWDVPAPKEVKKGETTDGVYRVMTRRLLGSTQVGVGVMQEGVFHTMWHVTKGSALRSGEGRLDPYWGDVKQDLVSYCGPWKLDAAWDGHSEVQLLAVPPGERARNIQTLPGIFKTKDGDIGAVALDYPAGTSGSPILDKCGRVIGLYGNGVVIKNGSYVSAITQGRREEETPVECFEPSMLKKKQLTVLDLHPGAGKTRRVLPEIVREAIKTRLRTVILAPTRVVAAEMEEALRGLPVRYMTTAVNVTHSGTEIVDLMCHATFTSRLLQPIRVPNYNLYIMDEAHFTDPSSIAARGYISTRVEMGEAAAIFMTATPPGTRDAFPDSNSPIMDTEVEVPERAWSSGFDWVTDHSGKTVWFVPSVRNGNEIAACLTKAGKRVIQLSRKTFETEFQKTKHQEWDFVVTTDISEMGANFKADRVIDSRRCLKPVILDGERVILAGPMPVTHASAAQRRGRIGRNPNKPGDEYLYGGGCAETDEDHAHWLEARMLLDNIYLQDGLIASLYRPEADKVAAIEGEFKLRTEQRKTFVELMKRGDLPVWLAYQVASAGITYTDRRWCFDGTTNNTIMEDSVPAEVWTRHGEKRVLKPRWMDARVCSDHAALKSFKEFAAGKRGAAFGVMEALGTLPGHMTERFQEAIDNLAVLMRAETGSRPYKAAAAQLPETLETIMLLGLLGTVSLGIFFVLMRNKGIGKMGFGMVTLGASAWLMWLSEIEPARIACVLIVVFLLLVVLIPEPEKQRSPQDNQMAIIIMVAVGLLGLITANELGWLERTKSDLSHLMGRREEGATIGFSMDIDLRPASAWAIYAALTTFITPAVQHAVTTSYNNYSLMAMATQAGVLFGMGKGMPFYAWDFGVPLLMIGCYSQLTPLTLIVAIILLVAHYMYLIPGLQAAAARAAQKRTAAGIMKNPVVDGIVVTDIDTMTIDPQVEKKMGQVLLIAVAVSSAILSRTAWGWGEAGALITAATSTLWEGSPNKYWNSSTATSLCNIFRGSYLAGASLIYTVTRNAGLVKRRGGGTGETLGEKWKARLNQMSALEFYSYKKSGITEVCREEARRALKDGVATGGHAVSRGSAKLRWLVERGYLQPYGKVIDLGCGRGGWSYYVATIRKVQEVKGYTKGGPGHEEPVLVQSYGWNIVRLKSGVDVFHMAAEPCDTLLCDIGESSSSPEVEEARTLRVLSMVGDWLEKRPGAFCIKVLCPYTSTMMETLERLQRRYGGGLVRVPLSRNSTHEMYWVSGAKSNTIKSVSTTSQLLLGRMDGPRRPVKYEEDVNLGSGTRAVVSCAEAPNMKIIGNRIERIRSEHAETWFFDENHPYRTWAYHGSYEAPTQGSASSLINGVVRLLSKPWDVVTGVTGIAMTDTTPYGQQRVFKEKVDTRVPDPQEGTRQVMSMVSSWLWKELGKHKRPRVCTKEEFINKVRSNAALGAIFEEEKEWKTAVEAVNDPRFWALVDKEREHHLRGECQSCVYNMMGKREKKQGEFGKAKGSRAIWYMWLGARFLEFEALGFLNEDHWMGRENSGGGVEGLGLQRLGYVLEEMSRIPGGRMYADDTAGWDTRISRFDLENEALITNQMEKGHRALALAIIKYTYQNKVVKVLRPAEKGKTVMDIISRQDQRGSGQVVTYALNTFTNLVVQLIRNMEAEEVLEMQDLWLLRRSEKVTNWLQSNGWDRLKRMAVSGDDCVVKPIDDRFAHALRFLNDMGKVRKDTQEWKPSTGWDNWEEVPFCSHHFNKLHLKDGRSIVVPCRHQDELIGRARVSPGAGWSIRETACLAKSYAQMWQLLYFHRRDLRLMANAICSSVPVDWVPTGRTTWSIHGKGEWMTTEDMLVVWNRVWIEENDHMEDKTPVTKWTDIPYLGKREDLWCGSLIGHRPRTTWAENIKNTVNMVRRIIGDEEKYMDYLSTQVRYLGEEGSTPGVL

>tr|W8Q6T3|W8Q6T3_ZIKV Genome polyprotein OSZika virus (strain Mr 766) PE4 SV1

MKNPKKKSGGFRIVNMLKRGVARVNPLGGLKRLPAGLLLGHGPIRMVLAILAFLRFTAIKPSLGLINRWGSVGKKEAMEIIKKFKKDLAAMLRIINARKERKRRGADTSIGIIGLLLTTAMAAEITRRGSAYYMYLDRSDAGKAISFATTLGVNKCHVQIMDLGHMCDATMSYECPMLDEGVEPDDVDCWCNTTSTWVVYGTCHHKKGEARRSRRAVTLPSHSTRKLQTRSQTWLESREYTKHLIKVENWIFRNPGFALAAVAIAWLLGSSTSQKVIYLIMILLIAPAYSIRCIGVSNRDFVEGMSGGTWVDVVLEHGGCVTVMAQDKPTVDIELVTTTVSNMAEVRSYCYEASISDMASDSRCPTQGEAYLDKQSDTQYVCKRTLVDRGWGNGCGLFGKGSLVTCAKFTCSKKMTGKSIQPENLEYRIMLSVHGSQHSGMIVNDXXXXXXXNRAEVEVTPNSPRAEATLGGFGSLGLDCEPRTGLDFSDLYYLTMNNKHWLVHKEWFHDIPLPWHAGADTGTPHWNNKEALVEFKDAHAKRQTVVVLGSQEGAVHTALAGALEAEMDGAKGRLFSGHLKCRLKMDKLRLKGVSYSLCTAAFTFTKVPAETLHGTVTVEVQYAGTDGPCKVPAQMAVDMQTLTPVGRLITANPVITESTENSKMMLELDPPFGDSYIVIGVGDKKITHHWHRSGSTIGKAFEATVRGAKRMAVLGDTAWDFGSVGGVFNSLGKGIHQIFGAAFKSLFGGMSWFSQILIGTLLVWLGLNTKNGSISLTCLALGGVMIFLSTAVSADVGCSVDFSKRETRCGTGVFVYNDVEAWRDRYKYHPDSPRRLAAAVKQAWEEGICGISSVSRMENIMWKSVEGELNAILEENGVQLTVVVGSVKNPMWRGPQRLPVPVNELPHGWKAWGKSYFVRAAKTNNSFVVDGDTLKECPLKHRAWNSFLVEDHGFGVFHTSVWLKVREDYSLECDPAVIGTAVKGKEAAHSDLGYWIESEKNDTWRLKRAHLIEMKTCEWPKSHTLWTDGVEESDLIIPKSLAGPLSHHNTREGYRTQAKGPWHSEELEIRFEECPGTKVHVEETCGTRGPSLRSTTASGRVIEEWCCRECTMPPLSFRAKDGCWYGMEIRPRKEPESNLVRSMVTAGSTDHMDHFSLGVLVILLMVQEGLKKRMTTKIIVSTSMAVLVAMILGGFSMSDLAKLVILMGATFAEMNTGGDVAHLALVAAFKVRPALLVSFIFRANWTPRESMLLALASCLLQTAISALEGDLMVLVNGFALAWLAIRAMAVPRTDNIALAILAALTPLARGTLLVAWRAGLATCGGFMLLSLKGKGSVKKNLPFVMALGMTAVRIVDPINVVGLLLLTRSGKRSWPPSEVLTAVGLICALAGGFAKADIEMAGPMAAVGLLIVSYVVSGKSVDMYIERAGDITWEKDAEVTGNSPRLDVALDESGDFSLVEEDGPPMREIILKVVLMAICGMNPIAIPFAAGAWYVYVKTGKRSGALWDVPAPKEVKKGETTDGVYRVMTRRLLGSTQVGVGVMQEGVFHTMWHVTKGAALRSGEGRLDPYWGDVKQDLVSYCGPWKLDAAWDGLSEVQLLAVPPGERARNIQTLPGIFKTKDGDIGAVALDYPAGTSGSPILDKCGRVIGLYGNGVVIKNGSYVSAITQGKREEETPVECFEPSMLKKKQLTVLDLHPGAGKTRRVLPEIVREAIKKRLRTVILAPTRVVAAEMEEALRGLXVRYMTTAVNVTHSGTEIVDLMCHATFTSRLLQPIRVPNYNLNIMDEAHFTDPSSIAARGYISTRVEMGEAAAIFMTATPPGTRDAFPDSNSPIMDTEVEVPERAWSSGFDWVTDHSGKTVWFVPSVRNGNEITACLTKAGKRVIQLSRKTFETEFQKTKNQEWDFVITTDISEMGANFKADRVIDSRRCLKPVILDGERVILAGPMPVTHASAAQRRGRIGRNPNKPGDEYMYGGGCAETDEDHAHWLEARMLLDNIYLQDGLIASLYRPEADKVAAIEGEFKLRTEQRKTFVELMKRGDLPVWLAYQVASAGITYTDRRWCFDGTTNNTIMEDSVPAEVWTKYGEKRVLKPRWMDARVCSDHAALKSFKEFAAGKRGAALGVMEALGTLPGHMTERFQEAIDNLAVLMRAETGSRPYKAAAAPLPETLETIMLLGLLGTVSLGIFFVLMRNKGIGKMGFGMVTLGASAWLMWLSEIEPARIACVLIVVFLLLVVLIPEPEKQRSPQDNQMAIIIMVAVGLLGLITANELGWLERTKNDIAHLMGRREEGVTMGFSMDIDLRPASAWAIYAALTTLITPAVQHAVTTSYNNYSLMAMATQAGVLFGMGKGMPFYAWDFGVPLLMMGCYSQLTPLTLIVAIILLVAHYMYLIPGLQAAAARAAQKRTAAGIMKNPVVDGIVVTDIDTMTIDPQVEKKMGQVLLIAVAISSAVLLRTAWGWGEAGALITAATSTLWEGSPNKYWNSSTATSLCNIFRGSYLAGASLIYTVTRNAGLVKRRGGGTGETLGEKWKARLNQMSALEFYSYKKSGITEVCREEARRALKDGVATGGHAVSRGSAKLRWLVERGYLQPHGKVVDLGCGRGGWSYYAATIRKVQEVKGYTKGGPGHEEPMLVQSYGWNIVRLKSGVDVFHMAAEPCDTLLCDIGESSSSPEVEETRTLRVLSMVGDWLEKRPGAFCIKVLCPYTSTMMETMERLQRRYGGGLVRVPLSRNSTHEMYWVSGAKSNIIKSVSTTSQLLLGRMDGPRRPVKYEEDVNLGSGTRAVASCAEAPNMKIIGKRIERIRSEHAETWFFDENHPYRTWAYHGSYEAPTQGSASSLVNGVVRLLSKPWDVVTGVTGIAMTDTTPYGQQRVFKEKVDTRVPDPQEGTRQTMNMVSSWLWKELGKRKRPRVCTKEEFINKVRSNAALGAIFEEEKEWKTAVEAVNDPRFWALVDKEREHHLRGECHSCVYNMMGKREKKXGEFGKAKGSRAIWYMWLGARFLEFESLGFLNEDHWMGRENSGGGVEGLGLQRLGYILEEMNRAPGGKMYADDTAGWDTRISKFDLENEALITNQMEEGHRALALAVIKYTYQNKVVKVLRPAEGGKTVMDIISRQDQRGSGQVVTYALNTFTNLVVQLIRNMEAEEVLEMQDLWLLRKPEKVTRWLQSNGWDRLKRMAVSGDDCVVKPIDDRFACCLRFLNDMGKVRKDTQEWKPSTGWSNWEEVPFCSHHFNKLHLKDGRSIVVPCRHQDELIGRARVSPGAGWSIRETACLAKSYAQMWQLLYFHRRDLRLMANAICSAVPVDWVPTGRTTWSIHGKGEWMTTEDMLMVWNRVWIEENDHMEDKTPVTKWTDIPYLGKREDLWCGSLIGHRPRTTWAENIKDTVNMVRRIIGDEEKYMDYLSTQVRYLGEEGSTPGVL

>tr|A0A0U4DG08|A0A0U4DG08_ZIKV Genome polyprotein OSZika virus (strain Mr 766) PE4 SV1

MKNPKKKSGGFRIVNMLKRGVARVSPFGGLKRLPAGLLLGHGPIRMVLAILAFLRFTAIKPSLGLINRWGSVGKKEAMEIIKKFKKDLAAMLRIINARKEKKRRGADTSVGIVGLLLTTAMAAEVTRRGSAYYMYLDRNDAGEAISFPTTLGMNKCYIQIMDLGHTCDATMSYECPMLDEGVEPDDVDCWCNTTSTWVVYGTCHHKKGEARRSRRAVTLPSHSTRKLQTRSQTWLESREYTKHLIRVENWIFRNPGFALAAAAIAWLLGSSTSQKVIYLVMILLIAPAYSIRCIGVSNRDFVEGMSGGTWVDVVLEHGGCVTVMAQDKPTVDIELVTTTVSNMAEVRSYCYEASISDMASDSRCPTQGEAYLDKQSDTQYVCKRTLVDRGWGNGCGLFGKGSLVTCAKFACSKKMTGKSIQPENLEYRIMLSVHGSQHSGMIVNDTGHETDENRAKVEITPNSPRAEATLGGFGSLGLDCEPRTGLDFSDLYYLTMNNKHWLVHKEWFHDIPLPWHAGADTGTPHWNNKEALVEFKDAHAKRQTVVVLGSQEGAVHTALAGALEAEMDGAKGRLSSGHLKCRLKMDKLRLKGVSYSLCTAAFTFTKIPAETLHGTVTVEVQYAGTDGPCKVPAQMAVDMQTLTPVGRLITANPVITESTENSKMMLELDPPFGDSYIVIGVGEKKITHHWHRSGSTIGKAFEATVRGAKRMAVLGDTAWDFGSVGGALNSLGKGIHQIFGAAFKSLFGGMSWFSQILIGTLLMWLGLNAKNGSISLMCLALGGVLIFLSTAVSADVGCSVDFSKKETRCGTGVFVYNDVEAWRDRYKYHPDSPRRLAAAVKQAWEDGICGISSVSRMENIMWRSVEGELNAILEENGVQLTVVVGSVKNPMWRGPQRLPVPVNELPHGWKAWGKSYFVRAAKTNNSFVVDGDTLKECPLKHRAWNSFLVEDHGFGVFHTSVWLKVREDYSLECDPAVIGTAVKGKEAVHSDLGYWIESEKNDTWRLKRAHLIEMKTCEWPKSHTLWTDGIEESDLIIPKSLAGPLSHHNTREGYRTQMKGPWHSEELEIRFEECPGTKVHVEETCGTRGPSLRSTTASGRVIEEWCCRECTMPPLSFRAKDGCWYGMEIRPRKEPESNLVRSMVTAGSTDHMDHFSLGVLVILLMVQEGLKKRMTTKIIISTSMAVLVAMILGGFSMSDLAKLAILMGATFAEMNTGGDVAHLALIAAFKVRPALLVSFIFRANWTPRESMLLALASCLLQTAISALEGDLMVLINGFALAWLAIRAMVVPRTDNITLAILAALTPLARGTLLVAWRAGLATCGGFMLLSLKGKGSVKKNLPFVMALGLTAVRLVDPINVVGLLLLTRSGKRSWPPSEVLTAVGLICALAGGFAKADIEMAGPMAAVGLLIVSYVVSGKSVDMYIERAGDITWEKDAEVTGNSPRLDVALDESGDFSLVEDDGPPMREIILKVVLMTICGMNPIAIPFAAGAWYVYVKTGKRSGALWDVPAPKEVKKGETTDGVYRVMTRRLLGSTQVGVGVMQEGVFHTMWHVTKGSALRSGEGRLDPYWGDVKQDLVSYCGPWKLDAAWDGHSEVQLLAVPPGERARNIQTLPGIFKTKDGDIGAVALDYPAGTSGSPILDKCGRVIGLYGNGVVIKNGSYVSAITQGRREEETPVECFEPSMLKKKQLTVLDLHPGAGKTRRVLPEIVREAIKTRLRTVILAPTRVVAAEMEEALRGLPVRYMTTAVNVTHSGTEIVDLMCHATFTSRLLQPIRVPNYNLYIMDEAHFTDPSSIAARGYISTRVEMGEAAAIFMTATPPGTRDAFPDSNSPIMDTEVEVPERAWSSGFDWVTDHSGKTVWFVPSVRNGNEIAACLTKAGKRVIQLSRKTFETEFQKTKHQEWDFVVTTDISEMGANFKADRVIDSRRCLKPVILDGERVILAGPMPVTHASAAQRRGRIGRNPNKPGDEYLYGGGCAETDEDHAHWLEARMLLDNIYLQDGLIASLYRPEADKVAAIEGEFKLRTEQRKTFVELMKRGDLPVWLAYQVASAGITYTDRRWCFDGTTNNTIMEDSVPAEVWTRHGEKRVLKPRWMDARVCSDHAALKSFKEFAAGKRGAAFGVMEALGTLPGHMTERFQEAIDNLAVLMRAETGSRPYKAAAAQLPETLETIMLLGLLGTVSLGIFFVLMRNKGIGKMGFGMVTLGASAWLMWLSEIEPARIACVLIVVFLLLVVLIPEPEKQRSPQDNQMAIIIMVAVGLLGLITANELGWLERTKSDLSHLMGRREEGATIGFSMDIDLRPASAWAIYAALTTFITPAVQHAVTTSYNNYSLMAMATQAGVLFGMGKGMPFYAWDFGVPLLMIGCYSQLTPLTLIVAIILLVAHYMYLIPGLQAAAARAAQKRTAAGIMKNPVVDGIVVTDIDTMTIDPQVEKKMGQVLLIAVAVSSAILSRTAWGWGEAGALITAATSTLWEGSPNKYWNSSTATSLCNIFRGSYLAGASLIYTVTRNAGLVKRRGGGTGETLGEKWKARLNQMSALEFYSYKKSGITEVCREEARRALKDGVATGGHAVSRGSAKLRWLVERGYLQPYGKVIDLGCGRGGWSYYAATIRKVQEVKGYTKGGPGHEEPVLVQSYGWNIVRLKSGVDVFHMAAEPCDTLLCDIGESSSSPEVEEARTLRVLSMVGDWLEKRPGAFCIKVLCPYTSTMMETLERLQRRYGGGLVRVPLSRNSTHEMYWVSGAKSNTIKSVSTTSQLLLGRMDGPRRPVKYEEDVNLGSGTRAVVSCAEAPNMKIIGNRIERIRSEHAETWFFDENHPYRTWAYHGSYEAPTQGSASSLINGVVRLLSKPWDVVTGVTGIAMTDTTPYGQQRVFKEKVDTRVPDPQEGTRQVMSMVSSWLWKELGKHKRPRVCTKEEFINKVRSNAALGAIFEEEKEWKTAVEAVNDPRFWALVDKEREHHLRGECQSCVYNMMGKREKKQGEFGKAKGSRAIWYMWLGARFLEFEALGFLNEDHWMGRENSGGGVEGLGLQRLGYVLEEMSRIPGGRMYADDTAGWDTRISRFDLENEALITNQMEKGHRALALAIIKYTYQNKVVKVLRPAEKGKTVMDIISRQDQRGSGQVVTYALNTFTNLVVQLIRNMEAEEVLEMQDLWLLRRSEKVTNWLQSNGWDRLKRMAVSGDDCVVKPIDDRFAHALRFLNDMGKVRKDTQEWKPSTGWDNWEEVPFCSHHFNKLHLKDGRSIVVPCRHQDELIGRARVSPGAGWSIRETACLAKSYAQMWQLLYFHRRDLRLMANAICSSVPVDWVPTGRTTWSIHGKGEWMTTEDMLVVWNRVWIEENDHMEDKTPVTKWTDIPYLGKREDLWCGSLIGHRPRTTWAENIKNTVNMVRRIIGDEEKYMDYLSTQVRYLGEEGSTPGVL

>tr|W8Q7Q3|W8Q7Q3_ZIKV Genome polyprotein OSZika virus (strain Mr 766) PE4 SV1

MKNPKKKSGGFRIVNMLKRGVARVNPLGGLKRLPAGLLLGHGPIRMVLAILAFLRFTAIKPSLGLINRWGSVGKKEAMEIIKKFKKDLAAMLRIINARKERKRRGADTSIGIIGLLLTTAMAAEITRRGSAYYMYLDRSDAGKAISFATTLGVNKCHVQIMDLGHMCDATMSYECPMLDEGVEPDDVDCWCNTTSTWVVYGTCHHKKGEARRSRRAVTLPSHSTRKLQTRSQTWLESREYTKHLIKVENWIFRNPGFALVAVAIAWLLGSSTSQKVIYLVMILLIAPAYSIRCIGVSNRDFVEGMSGGTWVDVVLEHGGCVTVMAQDKPTVDIELVTTTVSNMAEVRSYCYEASISDMASDSRCPTQGEAYLDKQSDTQYVCKRTLVDRGWGNGCGLFGKGSLVTCAKFTCSKKMTGKSIQPENLEYRIMLSVHGSQHSGMIVNDIGHETDENRAKVEVTPNSPRAEATLGGFGSLGLDCEPRTGLDFSDLYYLTMNNKHWLVHKEWFHDIPLPWHAGADTGTPHWNNKEALVEFKDAHAKRQTVVVLGSQEGAVHTALAGALEAEMDGAKGRLFSGHLKCRLKMDKLRLKGVSYSLCTAAFTFTKVPAETLHGTVTVEVQYAGTDGPCKVPAQMAVDMQTLTPVGRLITANPVITESTENSKMMLELDPPFGDSYIVIGVGDKKITHHWHRSGSTIGKAFEATVRGAKRMAVLGDTAWDFGSVGGVFNSLGKGIHQIFGAAFKSLFGGMSWFSQILIGTLLVWLGLNTKNGSISLTCLALGGVMIFLSTAVSADVGCSVDFSKRETRCGTGVFIYNDVEAWRDRYKYHPDSPRRLAAAVKQAWEEGICGISSVSRMENIMWKSVEGELNAILEENGVQLTVVVGSVKNPMWRGPQRLPVPVNGLPHGWKAWGKSYFVRAAKTNNSFVVDGDTLKECPLKHRAWNSFLVEDHGFGVFHTSVWLKVREDYSLECDPAVIGTAVKGREAAHSDLGYWIESEKNDTWRLKRAHLIEMKTCEWPKSHTLWTDGVEESDLIIPKSLAGPLSHHNTREGYRTQVKGPWHSEELEIRFEECPGTKVHVEETCGTRGPSLRSTTASGRVIEEWCCRECTMPPLSFRAKDGCWYGMEIRPRKEPESNLVRSMVTAGSTDHMDHFSLGVLVILLMVQEGLKKRMTTKIIMSTSMAVLVVMILGGFSMSDLAKLVILMGATFAEMNTGGDVAHLALVAAFKVRPALLVSFILRANWTPRESMLLALASCLLQTAISALEGDLMVLVNGFALAWLAIRAMAVPRTDNIALAILAALTPLARGTLLVAWRAGLATCGGFMLLSLKGKGSVKKNLPFVMALGLTAVRVVDPINVVGLLLLTRSGKRSWPPSEVLTAVGLICALAGGFAKADIEMAGPMAAVGLLIVSYVVSGKSVDMYIERAGDITWEKDAEVTGNSPRLDVALDESGDFSLVEEDGPPMREIILKVVLMAICGMNPIAIPFAAGAWYVYVKTGKRSGALWDVPAPKEVKKGETTDGVYRVMTRRLLGSTQVGVGVMQEGVFHTMWHVTKGAALRSGEGRLDPYWGDVKQDLVSYCGPWKLDAAWDGLSEVQLLAVPPGERARNIQTLPGIFKTKDGDIGAVALDYPAGTSGSPILDKCGRVIGLYGNGVVIKNGSYVSAITQGKREEETPVECFEPSMLKKKQLTVLDLHPGAGKTRRVLPEIVREAIKKRLRTVILAPTRVVAAEMEEALRGLPVRYMTTAVNVTHSGTEIVDLMCHATFTSRLLQPIRVPNYNLNIMDEAHFTDPSSIAARGYISTRVEMGEAAAIFMTATPPGTRDAFPDSNSPIMDTEVEVPERAWSSGFDWVTDHSGKTVWFVPSVRNGNEIAACLTKAGKRVIQLSRKTFETEFQKTKNQEWDFVITTDISEMGANFKADRVIDSRRCLKPVILDGERVILAGPMPVTHASAAQRRGRIGRNPNKPGDEYMYGGGCAETDEGHAHWLEARMLLDNIYLQDGLIASLYRPEADKVAAIEGEFKLRTEQRKTFVELMKRGDLPVWLAYQVASAGITYTDRRWCFDGTTNNTIMEDSVPAEVWTKYGEKRVLKPRWMDARVCSDHAALKSFKEFAAGKRGAALGVMEALGTLPGHMTERFQEAIDNLAVLMRAETGSRPYKAAAAQLPETLETIMLLGLLGTVSLGIFFVLMRNKGIGKMGFGMVTLGASAWLMWLSEIEPARIACVLIVVFLLLVVLIPEPEKQRSPQDNQMAIIIMVAVGLLGLITANELGWLERTKNDIAHLMGRREEGATMGFSMDIDLRPASAWAIYAALTTLITPAVQHAVTTSYNNYSLMAMATQAGVLFGMGKGMPFYAWDFGVPLLMMGCYSQLTPLTLIVAIILLVAHYMYLIPGLQAAAARAAQKRTAAGIMKNPVVDGIVVTDIDTMTIDPQVEKKMGQVLLIAVAISSAVLLRTAWGWGEAGALITAATSTLWEGSPNKYWNSSTATSLCNIFRGSYLAGASLIYTVTRNAGLVKRRGGGTGETLGEKWKARLNQMSALEFYSYKKSGITEVCREEARRALKDGVATGGHAVSRGSAKLRWLVERGYLQPHGKVVDLGCGRGGWSYYAATIRKVQEVRGYTKGGPGHEEPMLVQSYGWNIVRLKSGVDVFHMAAEPCDTLLCDIGESSSSPEVEETRTLRVLSMVGDWLEKRPGAFCIKVLCPYTSTMMETMERLQRRHGGGLVRVPLSRNSTHEMYWVSGAKSNIIKSVSTTSQLLLGRMDGPRRPVKYEEDVNLGSGTRAVASCAEAPNMKIIGRRIERIRNEHAETWFLDENHPYRTWAYHGSYEAPTQGSASSLVNGVVRLLSKPWDVVTGVTGIAMTDTTPYGQQRVFKEKVDTRVPDPQEGTRQVMNMVSSWLWKELGKRKRPRVCTKEEFINKVRSNAALGAIFEEEKEWKTAVEAVNDPRFWALVDREREHHLRGECHSCVYNMMGKREKKQGEFGKAKGSRAIWYMWLGARFLEFEALGFLNEDHWMGRENSGGGVEGLGLQRLGYVLEEMSQAPGGKMYADDTAGWDTRISKFDLENEALITNQMEEGHRTLALAVIKYTYQNKVVKVLRPAEGGKTVMDIISRQDQRGSGQVVTYALNTFTNLVVQLIRNMEAEEVLEMQDLWLLRKPEKVTRWLQSNGWDGLKRMAVSGDDCVVKPIDDRFAHALRFLNDMGKVRKDTQEWKPSTGWSNWEEVPFCSHHFNKLYLKDGRSIVVPCRHQDELIGRARVSPGAGWSIRETACLAKSYAQMWQLLYFHRRDLRLMANAICSAVPVDWVPTGRTTWSIHGKGEWMTTEDMLMVWNRVWIEENDHMEDKTPVTKWTDIPYLGKREDLWCGSLIGHRPRTTWAENIKDTVNMVRRIIGDEEKYMDYLSTQVRYLGEEGSTPGVL

>tr|A0A109ZYP8|A0A109ZYP8_ZIKV Genome polyprotein OSZika virus (strain Mr 766) PE4 SV1

MKNPKKKSGGFRIVNMLKRGVARVSPFGGLKRLPAGLLLGHGPIRMVLAILAFLRFTAIKPSLGLINRWGSVGKKEAMEIIKKFKKDLAAMLRIINARKEKKRRGADTSVGIVGLLLTTAMAAEVTRRGSAYYMYLDRNDAGEAISFPTTLGMNKCYIQIMDLGHMCDATMSYECPMLDEGVEPDDVDCWCNTTSTWVVYGTCHHKKGEARRSRRAVTLPSHSTRKLQTRSQTWLESREYTKHLIRVENWIFRNPGFALAAAAIAWLLGSSTSQKVIYLVMILLIAPAYSIRCIGVSNRDFVEGMSGGTWVDVVLEHGGCVTVMAQDKPTVDIELVTTTVSNMAEVRSYCYEASISDMASDSRCPTQGEAYLDKQSDTQYVCKRTLVDRGWGNGCGLFGKGSLVTCAKFACSKKMTGKSIQPENLEYRIMLSVHGSQHSGMIVNDTGHETDENRAKVEITPNSPRAEATLGGFGSLGLDCEPRTGLDFSDLYYLTMNNKHWLVHKEWFHDIPLPWHAGADTGTPHWNNKEALVEFKDAHAKRQTVVVLGSQEGAVHTALAGALEAEMDGAKGRLSSGHLKCRLKMDKLRLKGVSYSLCTAAFTFTKIPAETLHGTVTVEVQYAGTDGPCKVPAQMAVDMQTLTPVGRLITANPVITESTENSKMMLELDPPFGDSYIVIGVGEKKITHHWHRSGSTIGKAFEATVRGAKRMAVLGDTAWDFGSVGGALNSLGKGIHQIFGAAFKSLFGGMSWFSQILIGTLLMWLGLNTKNGSISLMCLALGGVLIFLSTAVSADVGCSVDFSKKETRCGTGVFVYNDVEAWRDRYKYHPDSPRRLAAAVKQAWEDGICGISSVSRMENIMWRSVEGELNAILEENGVQLTVVVGSVKNPMWRGPQRLPVPVNELPHGWKAWGKSYFVRAAKTNNSFVVDGDTLKECPLEHRAWNSFLVEDHGFGVFHTSVWLKVREDYSLECDPAVIGTAVKGKEAVHSDLGYWIESEKNDTWRLKRAHLIEMKTCEWPKSHTLWADGIEESDLIIPKSLAGPLSHHNTREGYRTQMKGPWHSEELEIRFEECPGTKVHVEETCGTRGPSLRSTTASGRVIEEWCCRECTMPPLSFRAKDGCWYGMEIRPRKEPESNLVRSVVTAGSTDHMDHFSLGVLVILLMVQEGLKKRMTTKIIISTSMAVLVAMILGGFSMSDLAKLAILMGATFAEMNTGGDVAHLALIAAFKVRPALLVSFIFRANWTPRESMLLALASCLLQTAISALEGDLMVLINGFALAWLAIRAMVVPRTDNITLAILAALTPLARGTLLVAWRAGLATCGGFMLLSLKGKGSVKKNLPFVMALGLTAVRLVDPINVVGLLLLTRSGKRSWPPSEVLTAVGLICALAGGFAKADIEMAGPMAAVGLLIVSYVVSGKSVDMYIERAGDITWEKDAEVTGNSPRLDVALDESGDFSLVEDDGPPMREIILKVVLMTICGMNPIAIPFAAGAWYVYVKTGKRSGALWDVPAPKEVKKGETTDGVYRVMTRRLLGSTQVGVGVMQEGVFHTMWHVTKGSALRSGEGRLDPYWGDVKQDLVSYCGPWKLDAAWDGHSEVQLLAVPPGERARNIQTLPGIFKTKDGDIGAVALDYPAGTSGSPILDKCGRVIGLYGNGVVIKNGSYVSAITQGRREEETPVECFEPSMLKKKQLTVLDLHPGAGKTRRVLPEIVREAIKTRLRTVILAPTRVVAAEMEEALRGLPVRYMTTAVNVTHSGTEIVDLMCHATFTSRLLQPIRVPNYNLYIMDEAHFTDPSSIAARGYISTRVEMGEAAAIFMTATPPGTRDAFPDSNSPIMDTEVEVPERAWSSGFDWVTDHSGKTVWFVPSVRNGNEIAACLTKAGKRVIQLSRKTFETEFQKTKHQEWDFVVTTDISEMGANFKADRVIDSRRCLKPVILDGERVILAGPMPVTHASAAQRRGRIGRNPNKPGDEYLYGGGCAETDEDHAHWLEARMLLDNIYLQDGLIASLYRPEADKVAAIEGEFKLRTEQRKTFVELMKRGDLPVWLAYQVASAGITYTDRRWCFDGTTNNTIMEDSVPAEVWTRHGEKRVLKPRWMDARVCSDHAALKSFKEFAAGKRGAAFGVMEALGTLPGHMTERFQEAIDNLAVLMRAETGSRPYKAAAAQLPETLETIMLLGLLGTVSLGIFFVLMRNKGIGKMGFGMVTLGASAWLMWLSEIEPARIACVLIVVFLLLVVLIPEPEKQRSPQDNQMAIIIMVAVGLLGLITANELGWLERTKSDLSHLMGRREEGATIGFSMDIDLRPASAWAIYAALTTFITPAVQHAVTTSYNNYSLMAMATQAGVLFGMGKGMPFYAWDFGVPLLMIGCYSQLTPLTLIVAIILLVAHYMYLIPGLQAAAARAAQKRTAAGIMKNPVVDGIVVTDIDTMTIDPQVEKKMGQVLLIAVAVSSAILSRTAWGWGEAGALITAATSTLWEGSPNKYWNSSTATSLCNIFRGSYLAGASLIYIVTRNAGLVKRRGGGTGETLGEKWKARLNQMSALEFYSYKKSGITEVCREEARRALKDGVATGGHAVSRGSAKLRWLVERGYLQPYGKVIDLGCGRGGWSYYAATIRKVQEVKGYTKGGPGHEEPVLVQSYGWNIVRLKSGVDVFHMAAEPCDTLLCDIGESSSSPEVEEARTLRVLSMVGDWLEKRPGAFCIKVLCPYTSTMMETLERLQRRYGGGLVRVPLSRNSTHEMYWVSGAKSNTIKSVSTTSQLLLGRMDGPRRPVKYEEDVNLGSGTRAVVSCAEAPNMKIIGNRIERIRSEHAETWFFDENHPYRTWAYHGSYEAPTQGSASSLINGVVRLLSKPWDVVTGVTGIAMTDTTPYGQQRVFKEKVDTRVPDPQEGTRQVMSMVSSWLWKELGKHKRPRVCTKEEFINKVRSNAALGAIFEEEKEWKTAVEAVNDPRFWALVDKEREHHLRGECQSCVYNMMGKREKKQGEFGKAKGSRAIWYMWLGARFLEFEALGFLNEDHWMGRENSGGGVEGLGLQRLGYVLEEMSRIPGGRMYADDTAGWDTRISRFDLENEALITNQMEKGHRALALAIIKYTYQNKVVKVLRPAEKGKTVMDIISRQDQRGSGQVVTYALNTFTNLVVQLIRNMEAEEVLEMQDLWLLRRSEKVTNWLQSNGWDRLKRMAVSGDDCVVKPIDDRFAHALRFLNDMGKVRKDTQEWKPSTGWDNWEEVPFCSHHFNKLHLKDGRSIVVPCRHQDELIGRARVSPGAGWSIRETACLAKSYAQMWQLLYFHRRDLRLMANAICSSVPVDWVPTGRTTWSIHGKGEWMTTEDMLVVWNRVWIEENDHMEDKTPVTKWTDIPYLGKREDLWCGSLIGHRPRTTWAENIKNTVNMVRRIIGDEEKYMDYLSTQVRYLGEEGSTPGVL

>tr|W8QIT7|W8QIT7_ZIKV Genome polyprotein OSZika virus (strain Mr 766) PE4 SV1

MKNPKRKSGGFRIVNMLKRGVARVNPLGGLKRLPAGLLLGHGPIRMVLAILAFLRFTAIKPSLGLINRWGSVGKKEAMEIIKKFKKDLAAMLRIINARKERKRRGADTSIGIIGLLLTTAMAAEITRRGSAYYMYLDRSDAGKAISFATTLGVNKCHVQIMDLGHMCDATMSYECPMLDEGVEPDDVDCWCNTTSTWVVYGTCHHKKGEARRSRRAVTLPSHSTRKLQTRSQTWLESREYTKHLIKVENWIFRNPGFALAAVAIAWLLGSSTSQKVIYLVMILLIAPAYSIRCIGVSNRDFVEGMSGGTWVDVVLEHGGCVTVMAQDKPTVDIELVTTTVSNMAEVRSYCYEASISDMASDSRCPTQGEAYLDKQSDTQYVCKRTLVDRGWGNGCGLFGKGSLVTCAKFTCSKKMTGKSIQPENLEYRIMLSVHGSQHSGMXXXXXGHETDENRAKVEVTPNSPRAEATLGGFGSLGLDCEPRTGLDFSDLYYLTMNNKHRLVRKEWFHDIPLPWHAGADTGTPHWNNKEALVEFKDAHAKRQTVVVLGSQEGAVHTALAGALEAEMDGAKGRLFSGHLKCRLKMDKLRLKGVSYSLCTAAFTFTKVPAETLHGTVTVEVQYAGTDGPCKVPAQMAVDMQTLTPVGRLITANPVITESTENSKMMLELDPPFGDSYIVIGVGDKKITHHWLKKGSSIGKAFEATVRGAKRMAVLGDTAWDFGSVGGVFNSLGKGVHQIFGAAFKSLFGGMSWFSQILIGTLLVWLGLNTKNGSISLTCLALGGVMIFLSTAVSADVGCSVDFSKKETRCGTGVFVYNDVEAWRDRYKYHPDSPRRLAAAVKQAWEEGICGISSVSRMENIMWKSVEGELNAILEENGVQLTVVVGSVKNPMWRGPQRLPVPVNELPHGWKAWGKSYFVRAAKTNNSFVVDGDTLKECPLKHRAWNSFLVEDHGFGIFHTSVWLKVREDYSLECDPAVIGTAVKGKEAAHSDLCYWIESEKNDTWRLRRAHLIEMKTCEWPKSHTLWTDGVEESDLIIPKSLAGPLSHHNTREGYRTQVKGPWHSEELEIRFEECPGTKVYVEETCGTRGPSLRSTTASGRVIEEWCCRECTMPPLSFRAKDGCWYGMEIRPRKEPESNLVRSMVTAGSTDHMDHFSLGVLVILLMVQEGLKKRMTTKIIMSTSMAVLVAMVLGGFSMSDLAKLVILMGATFAEMNTGGDVAHLALVAAFKVRPALLVSFIFRANWTPRESMLLALASCLLQTAISALEGELMVLVNGFALAWLAIRAMAVPRTDNIALATLAALTPLARGTLLVAWRAGLATCGGFMLLSLKGKGSVKKNLPFVMALGLAAVRIVDPINVVGLLLLTRSGKRSWPPSEVLTAVGLICALAGGFAKADIEMAGPMAAVGLLIVSYVVSGKSVDMYIERAGDITWEKDAEVTGNSPRLDVALDESGDFSLVEEDGPPMREIILKVVLMAICGMNPIAIPFAAGAWYVYVKTGKRSGALWDVPAPKEVKKGETTDGVYRVMTRRLLGSTQVGVGVMQEGVFHTMWHVTKGAALRSGEGRLDPYWGDVKQDLVSYCGPWKLDATWDGLSEVQLLAVPPGERARNIQTLPGIFKTKDGDIGAVALDYPAGTSGSPILDKCGRVIGLYGNGVVIKNGSYVSAITQGKREEEAPVECFEPSMLRKKQLTVLDLHPGAGKTRRVLPEIVREAIKKRLRTVILAPTRVVAAEMEEALRGLPVRYMTTAVNVTHSGTEIVDLMCHATFTSRLLQPIRVPNYNLNIMDEAHFTDPSSIAARGYISTRVEMGEAAAIFMTATPPGTRDAFPDSNSPIMDTEVEVPERAWSSGFDWVTDHSGKTIWFVPSVRNGNEIAACLTKAGKRVIQLSRKTFETEFQKTKNQEWDFVITTDISEMGANFKADRVIDSRRCLKPVILDGERVILAGPMPVTHASAAQRRGRIGRNPNKPGDEYMYGGGCAETDEDHAHWLEARMLLDNIYLQDGLIASLYRPEADKVAAIEGEFKLRTEQRKTFVELMKRGDLPVWLAYQVASAGITYTDRRWCFDGTTNNTIMEDSVPAEVWTKYGEKRVLKPRWMDARVCSDHAALKSFKEFAAGKRGAALGVMDALGTLPGHMTERFQEAIDNLAVLMRAETGSRPYKAAAAQLPETLETIMLLGLLGTVSLGIFFVLMRNKGIGKMGFGMVTLGASAWLMWLSEIEPARIACVLIVVFLLLVVLIPEPEKQRSPQDNQMAIIIMVAVGLLGLITANELGWLERTKSDIAYLMGRKEEGTTIGFSMDIDLRPASAWAIYAALTTLITPAVQHAVTTSYNNYSLMAMATQAGVLFGMGKGMPFYAWDFGVPLLMIGCYSQLTPLTLIVAIILLVAHYMYLIPGLQAAAARAAQKRTAAGIMKNPVVDGIVVTDIDTMTIDPQVEKKMGQVLLIAVAVSSAVLLRTAGGWGEAGALITAATSTLWEGSPNKYWNSSTATSLCNIFRGSYLAGASLIYTVTRNAGLVKRRGGGTGETLGEKWKARLNQMSALEFYSYKKSGITEVCREEARRALKDGVATGGHAVSRGSAKLRWLVERGYLQPHGKVVDLGCGRGGWSYYAATIRKVQEVRGYTKGGPGHEEPMLVQSYGWNIVRLKSGVDVFHMAAEPCDTLLCDIGESSSSPEVEETRTLRVLSMVEDWLEKRPGAFCIKVLCPYTSTMMETMERLQRRHGGGLVRVPLSRNSTHEMYWVSGAKSNIIKSVSTTSQLLLGRMDGPRRPVKYEEDVNLGSGTRAVASCAEAPNMKIIGRRIERIRNEHAETWFFDENHPYRTWAYHGSYEAPTQGSASSLVNGVVRLLSKPWDVVTGVTGIAMTDTTPYGQQRVFKEKVDTRVPDPQEGTRQVMNIVSSWLWKELGKRKRPRVCTKEEFINKVRSNAALGAIFEEEKEWKTAVEAVNDPRFWALVDKEREHHLRGECHSCVYNMMGKREKKQGEFGKAKGSRAIWYMWLGARFLEFEALGFLNEDHWMGRENSGGGVEGLGLQRLGYILEEMNRAPGGKMYADDTAGWDTRISKFDLENEALITNQMEEGHRALALAVIKYTYQNKVVKVLRPAEGGKTVMDIISRQDQRGSGQVVTYALNTFTNLVVQLIRNMEAEEVLEMHDLWLLRKPEKVTRWLQSNGWDRLKRMAVSGDDCVVKPIDDRFAHALRFLNDMGKVRKDTQEWKPSTGWSNWEEVPFCSHHFNKLHLKDGRSIVVPCRHQDELIGRARVSPGAGWSIRETACLAKSYAQMWQLLYFHRRDLRLMANAICSAVPVDWVPTGRTTWSIHGKGEWMTTEDMLMVWNRVWIEENDHMEDKTPVPKWTEIPYLGKRESLWCGYIIGHRPRTTWAENIKDTVNMVLRLIGDEEKYMDYLSTQVRYLGEEGSTPGVL

>tr|A0A090BCI3|A0A090BCI3_ZIKV Genome polyprotein OSZika virus (strain Mr 766) PE4 SV1

MKNPKKKSGGFRIVNMLKRGVARVNPLGGLKRLPAGLLLGHGPIRMVLAILAFLRFTAIKPSLGLINRWGSVGKKEAMEIIKKFKKDLAAMLRIINARKERKRRGADTSIGIIGLLLTTAMAAEITRRGSAYYMYLDRSDAGKAISFATTLGVNKCHVQIMDLGHMCDATMSYECPMLDEGVEPDDVDCWCNTTSTWVVYGTCHHKKGEARRSRRAVTLPSHSTRKLQTRSQTWLESREYTKHLIKVENWIFRNPGFALVAVAIAWLLGSSTSQKVIYLVMILLIAPAYSIRCIGVSNRDFVEGMSGGTWVDVVLEHGGCVTVMAQDKPTVDIELVTTTVSNMAEVRSYCYEASISDMASDSRCPTQGEAYLDKQSDTQYVCKRTLVDRGWGNGCGLFGKGSLVTCAKFTCSKKMTGKSIQPENLEYRIMLSVHGSQHSGMTVNDIGYETDENRAKVEVTPNSPRAEATLGGFGSLGLDCEPRTGLDFSDLYYLTMNNKHWLVHKEWFHDIPLPWHAGADTGTPHWNNKEALVEFKDAHAKRQTVVVLGSQEGAVHTALAGALEAEMDGAKGKLFSGHLKCRLKMDKLRLKGVSYSLCTAAFTFTKVPAETLHGTVTVEVQYAGTDGPCKIPVQMAVDMQTLTPVGRLITANPVITESTENSKMMLELDPPFGDSYIVIGVGDKKITHHWHRSGSTIGKAFEATVRGAKRMAVLGDTAWDFGSVGGVFNSLGKGIHQIFGAAFKSLFGGMSWFSQILIGTLLVWLGLNTKNGSISLTCLALGGVMIFLSTAVSADVGCSVDFSKKETRCGTGVFIYNDVEAWRDRYKYHPDSPRRLAAAVKQAWEEGICGISSVSRMENIMWKSVEGELNAILEENGVQLTVVVGSVKNPMWRGPQRLPVPVNELPHGWKAWGKSYFVRAAKTNNSFVVDGDTLKECPLEHRAWNSFLVEDHGFGVFHTSVWLKVREDYSLECDPAVIGTAVKGREAAHSDLGYWIESEKNDTWRLKRAHLIEMKTCEWPKSHTLWTDGVEESDLIIPKSLAGPLSHHNTREGYRTQVKGPWHSEELEIRFEECPGTKVYVEETCGTRGPSLRSTTASGRVIEEWCCRECTMPPLSFRAKDGCWYGMEIRPRKEPESNLVRSMVTAGSTDHMDHFSLGVLVILLMVQEGLKKRMTTKIIMSTSMAVLVVMILGGFSMSDLAKLVILMGATFAEMNTGGDVAHLALVAAFKVRPALLVSFIFRANWTPRESMLLALASCLLQTAISALEGDLMVLINGFALAWLAIRAMAVPRTDNIALPILAALTPLARGTLLVAWRAGLATCGGIMLLSLKGKGSVKKNLPFVMALGLTAVRVVDPINVVGLLLLTRSGKRSWPPSEVLTAVGLICALAGGFAKADIEMAGPMAAVGLLIVSYVVSGKSVDMYIERAGDITWEKDAEVTGNSPRLDVALDESGDFSLVEEDGPPMREIILKVVLMAICGMNPIAIPFAAGAWYVYVKTGKRSGALWDVPAPKEVKKGETTDGVYRVMTRRLLGSTQVGVGVMQEGVFHTMWHVTKGAALRSGEGRLDPYWGDVKQDLVSYCGPWKLDAAWDGLSEVQLLAVPPGERARNIQTLPGIFKTKDGDIGAVALDYPAGTSGSPILDKCGRVIGLYGNGVVIKNGSYVSAITQGKREEETPVECFEPSMLKKKQLTVLDLHPGAGKTRRVLPEIVREAIKKRLRTVILAPTRVVAAEMEEALRGLPVRYMTTAVNVTHSGTEIVDLMCHATFTSRLLQPIRVPNYNLYIMDEAHFTDPSSIAARGYISTRVEMGEAAAIFMTATPPGTRDAFPDSNSPIMDTEVEVPERAWSSGFDWVTDHSGKTVWFVPSVRNGNEIAACLTKAGKRVIQLSRKTFETEFQKTKNQEWDFVITTDISEMGANFKADRVIDSRRCLKPVILDGERVILAGPMPVTHASAAQRRGRIGRNPNKPGDEYMYGGGCAETDEGHAHWLEARMLLDNIYLQDGLIASLYRPEADKVAAIEGEFKLRTEQRKTFVELMKRGDLPVWLAYQVASAGITYTDRRWCFDGTTNNTIMEDSVPAEVWTKYGEKRVLKPRWMDARVCSDHAALKSFKEFAAGKRGAALGVMEALGTLPGHMTERFQEAIDNLAVLMRAETGSRPYKAAAAQLPETLETIMLLGLLGTVSLGIFFVLMRNKGIGKMGFGMVTLGASAWLMWLSEIEPARIACVLIVVFLLLVVLIPEPEKQRSPQDNQMAIIIMVAVGLLGLITANELGWLERTKNDIAHLMGRREEGATMGFSMDIDLRPASAWAIYAALTTLITPAVQHAVTTSYNNYSLMAMATQAGVLFGMGKGMPFYAWDLGVPLLMMGCYSQLTPLTLIVAIILLVAHYMYLIPGLQAAAARAAQKRTAAGIMKNPVVDGIVVTDIDTMTIDPQVEKKMGQVLLIAVAISSAVLLRTAWGWGEAGALITAATSTLWEGSPNKYWNSSTATSLCNIFRGSYLAGASLIYTVTRNAGLVKRRGGGTGETLGEKWKARLNQMSALEFYSYKKSGITEVCREEARRALKDGVATGGHAVSRGSAKLRWLVERGYLQPYGKVVDLGCGRGGWSYYAATIRKVQEVRGYTKGGPGHEEPMLVQSYGWNIVRLKSGVDVFHMAAEPCDTLLCDIGESSSSPEVEETRTLRVLSMVGDWLEKRPGAFCIKVLCPYTSTMMETMERLQRRHGGGLVRVPLSRNSTHEMYWVSGAKSNIIKSVSTTSQLLLGRMDGPRRPVKYEEDVNLGSGTRAVASCAEAPNMKIIGRRIERIRNEHAETWFLDENHPYRTWAYHGSYEAPTQGSASSLVNGVVRLLSKPWDVVTGVTGIAMTDTTPYGQQRVFKEKVDTRVPDPQEGTRQVMNIVSSWLWKELGKRKRPRVCTKEEFINKVRSNAALGAIFEEEKEWKTAVEAVNDPRFWALVDREREHHLRGECHSCVYNMMGKREKKQGEFGKAKGSRAIWYMWLGARFLEFEALGFLNEDHWMGRENSGGGVEGLGLQRLGYILEEMNRAPGGKMYADDTAGWDTRISKFDLENEALITNQMEEGHRTLALAVIKYTYQNKVVKVLRPAEGGKTVMDIISRQDQRGSGQVVTYALNTFTNLVVQLIRNMEAEEVLEMQDLWLLRKPEKVTRWLQSNGWDRLKRMAVSGDDCVVKPIDDRFAHALRFLNDMGKVRKDTQEWKPSTGWSNWEEVPFCSHHFNKLYLKDGRSIVVPCRHQDELIGRARVSPGAGWSIRETACLAKSYAQMWQLLYFHRRDLRLMANAICSAVPVDWVPTGRTTWSIHGKGEWMTTEDMLMVWNRVWIEENDHMEDKTPVTKWTDIPYLGKREDLWCGSLIGHRPRTTWAENIKDTVNMVRRIIGDEEKYMDYLSTQVRYLGEEGSTPGVL

>tr|W8R1T1|W8R1T1_ZIKV Genome polyprotein OSZika virus (strain Mr 766) PE4 SV1

MKNPKKRSGGFRIVNMLKRGVARVNPLGGLKRLPAGLLLGHGPIRMVLAILAFLRFTAIKPSLGLINRWGSVGKKEAMEIIKKFKKDLAAMLRIINARKERKRRGADTSIGIVGLLLTTAMAAEITRRGSAYYMYLDRSDAGKAISFATTLGVNKCHVQIMDLGHMCDATMSYECPMLDEGVEPDDVDCWCNTTSTWVVYGTCQHKKGEARRSRRAVTLPSHSTRKLQTRSQTWLESREYTKHLIKVENWIFRNPGFALVAVAIAWLLGSSTSQKVIYLVMILLIAPAYSIRCIGVSNRDFVEGMSGGTWVDVVLEHGGCVTVMAQDKPTVDIELVTTTVSNMAEVRSYCYEASISDMASDSRCPTQGEAYLDKQSDTQYVCKRTLVDRGWGNGCGLFGKGSLVTCAKFTCSKKMTGKSIQPENLEYRIMLSVHGSQHSGMIVNDIGHETDENRAKVEVTPNSPRAEATLGGFGSLGLDCEPRTGLDFSDLYYLTMNNKHWLVHKEWFHDIPLPWHAGADTGTPHWNNKEALVEFKDAHAKRQTVVVLGSQEGAVHTALAGALEAEMDGAKGRLFSGHLKCRLKMDKLRLKGVSYSLCTAVCTAAKVPAETLHGTVTVEVQYAGTDGPCKVPAQMAVDMQTLTPVGRLITANPVITESTENSKMMLELDPPFGDSYIVIGVGDKKITHHWHRSGSTIGKAFEATVRGAKRMAVLGDTAWDFGSVGGVFNSLGKGIHQIFGAAFKSLFGGMSWFSQILIGTLLVWLGLNTKNGSISLTCLALGGVMIFLSTAVSADVGCSVDFSKKETRCGTGVFVYNDVEAWRDRYKYHPDSPRRLAAAVKQAWEEGICGISSVSRMENIMWKSVEGELNAILEENGVQLTVVVGSVKNPMWRGPQRLPVPVNELPHGWKAWGKSYFVRAAKTNNSFVVDGDTLKECPLKHRAWNSFLVEDHGFGIFHTSVWLKVREDYSLECDPAVIGTAVKGKEAAHSDLGYWIESEKNDTWRLRRAHLIEMKTCEWPKSHTLWTDGVEESDLIIPKSLAGPLSHHNTREGYRTQVKGPWHSEELEIRFEECPGTKVHVEETCGTRGPSLRSTTASGRVIEEWCCRECTMPPLSFRAKDGCWYGMEIRPRKEPESNLVRSMVTAGSTDHMDHFSLGVLVILLMVQEGLKKRMTTKIIMSTSMAVLVAMVLGGFSMSDLAKLVILMGATFAEMNTGGDVAHLALVAAFKVRPALLVSFIFRANWTPRESMLLALASCLLQTAISALEGELMVLVNGFALAWLAIRAMAVPRTDNIALAILAALTPLARGTLLVAWRAGLATCGGFMLLSLKGKGSVKKNLPFVMALGLTAVRIVDPINVVGLLLLTRSGKRSWPPSEVLTAVGLICALAGGFAKADIEMAGPMAAVGLLIVSYVVSGKSVDMYIERAGDITWEKDXEVTGNSPRLDVALDESGDFSLVEEDGPPMREIILKVVLMAICGMNPIAIPFAAGAWYVYVKTGKRSGALWDVPAPKEVKKGETTDGVYRVMTRRLLGSTQVGVGVMQEGVFHTMWHVTKGAALRSGEGRLDPYWGDVKQDLVSYCGPWKLDAAWDGLSEVQLLAVPPGERARNIQTLPGIFKTKDGDIGAVALDYPAGTSGSPILDKCGRVIGLYGNGVVIKNGSYVSAITQGKREEEAPVECFEPSMLRKKQLTVLDLHPGAGKTRRVLPEIVREAIKKRLRTLILAPTRVVAAEMEEALRGLXVRYMTTAVNVTHSGTEIVDLMCHATFTSRLLQPIRVPNYNLNIMDEAHFTDPSSIAARGYISTRVEMGEAAAIFMTATPPGTRDAFPDSNSPIMDTEVEVPERAWSSGFDWVTDHSGKTIWFVPSVRSGEESAACLTKAGKRVIQLSRKTFETEFQKTKNQEWDFVITTDISEMGANFKADRVIDSRRCLKPVILDGERVILAGPMPVTHASAAQRRGRIGRNPNKPGDEYMYGGGCAETDEDHAHWLEARMLLDNIYLQDGLIASLYRPEADKVAAIEGEFKLRTEQRKTFVELMKRGDLPVWLAYQVASAGITYTDRRWCFDGTTNNTIMEDSVPAEVWTKYGEKRVLKPRWMDARVCSDHAALKSFKEFAAGKRGAALGVMDALGTLPGHMTERFQEAIDNLAVLMRAETGSRPYKAAAAQLPETLETIMLLGLLGTVSLGIFFVLMRNKGIGKMGFGMVTLGASAWLMWLSEIEPARIACVLIVVFLLLVVLIPEPEKQRSPQDNQMAIIIMVAVGLLGLITANELGWLERTKSDIAHLMGRKEEGTTIGFSMDIDLRPASAWAIYAALTTLITPAVQHAVTTSYNNYSLMAMATQAGVLFGMGKGMPFYAWDFGVPLLMMGCYSQLTPLTLIVAIILLVAHYMYLIPGLQAAAARAAQKRTAAGIMKNPVVDGIVVTDIDTMTIDPQVEKKMGQVLLIAVAVSSAVLLRTAWGWGEAGALITAATSTLWEGSPNKYWNSSTATSLCNIFRGSYLAGASLIYTVTRNAGLVKRRGGGTGETLGEKWKARLNQMSALEFYSYKKSGITEVCREEARRALKDGVATGGHAVSRGSAKLRWLVERGYLQPHGKVVDLGCGRGGWSYYAATIRKVQEVRGYTKGGPGHEEPMLVQSYGWNIVRLKSGVDVFHMAAEPCDTLLCDIGESSSSPEVEETRTLRVLSMVGDWLEKRPGAFCIKVLCPYTSTMMETMERLQRRHGGGLVRVPLSRNSTHEMYWVSGAKSNIIKSVSTTSQLLLGRMDGPRRPVKYEEDVNLGSGTRAVASCAEAPNMKIIGRRIERIRNEHAETWFFDENHPYRTWAYHGSYEAPTQGSASSLVNGVVRLLSKPWDVVTGVTGIAMTDTTPYGQQRVFKEKVDTRVPDPQEGTRQVMNMVSSWLWKELGKRKRPRVCTKEEFINKVRSNAALGAIFEEEKEWKTAVEAVNDPRFWALVDKEREHHLRGECHSCVYNMMGKREKKQGEFGKAKGSRAIWYMWLGARFLEFEALGFLNEDHWMGRENSGGGVEGLGLQRLGYILEEMNRAPGGKMYADDTAGWDTRISKFDLENEALITNQMEEGHRALALAVIKYTYQNKVVKVLRPAEGGKTVMDIISRQDQRGSGQVVTYALNTFTNLVVQLIRNMEAEEVLEMHDLWLLRKPEKVTRWLQSNGWDRLKRMAVSGDDCVVKPIDDRFAHALRFLNDMGKVRKDTQEWKPSTGWSNWEEVPFCSHHFNKLHLKDGRSIVVPCRHQDELIGRARVSPGAGWSIRETACLAKSYAQMWQLLYFHKRDLRLMANAICSAVPVDWVPTGRTTWSIHGKGEWMTTEDMLMVWNRVWIEENDHMEDKTPVTKWTDIPYLGKREDLWCGSLIGHRPRTTWAENIKDTVNMVRRIIGDEEKYMDYLSTQVRYLGEEGSTPGVL

>tr|A0A0U3FSM8|A0A0U3FSM8_ZIKV Genome polyprotein OSZika virus (strain Mr 766) PE4 SV1

MKNPKKKSGGFRIVNMLKRGVARVSPFGGLKRLPAGLLLGHGPIRMVLAILAFLRFTAIKPSLGLINRWGSVGKKEAMEIIKKFKKDLAAMLRIINARKEKKRRGADTSVGIVGLLLTTAMAAEVTRRGSAYYMYLDRNDAGEAISFPTTLGMNKCYIQIMDLGHMCDATMSYECPMLDEGVEPDDVDCWCNTTSTWVVYGTCHHKKGEARRSRRAVTLPSHSTRKLQTRSQTWLESREYTKHLIRVENWIFRNPGFALAAAAIAWLLGSSTSQKVIYLVMILLIAPAYSIRCIGVSNRDFVEGMSGGTWVDIVLEHGGCVTVMAQDKPTVDIELVTTTVSNMAEVRSYCYEASISDMASDSRCPTQGEAYLDKQSDTQYVCKRTLVDRGWGNGCGLFGKGSLVTCAKFACSKKMTGKSIQPENLEYRIMLSVHGSQHSGMIVNDTGHETDENRAKVEITPNSPRAEATLGGFGSLGLDCEPRTGLDFSDLYYLTMNNKHWLVHKEWFHDIPLPWHAGADTGTPHWNNKEALVEFKDAHAKRQTVVVLGSQEGAVHTALAGALEAEMDGAKGRLSSGHLKCRLKMDKLRLKGVSYSLCTAAFTFTKIPAETLHGTVTVEVQYAGTDGPCKVPAQMAVDMQTLTPVGRLITANPVITESTENSKMMLELDPPFGDSYIVIGVGEKKITHHWHRSGSTIGKAFEATVRGAKRMAVLGDTAWDFGSVGGALNSLGKGIHQIFGAAFKSLFGGMSWFSQILIGTLLMWLGLNTKNGSISLMCLALGGVLIFLSTAVSADVGCSVDFSKKETRCGTGVFVYNDVEAWRDRYKYHPDSPRRLAAAVKQAWEDGICGISSVSRMENIMWRSVEGELNAILEENGVQLTVVVGSVKNPMWRGPQRLPVPVNELPHGWKAWGKSHFVRAAKTNNSFVVDGDTLKECPLKHRAWNSFLVEDHGFGVFHTSVWLKVREDYSLECDPAVIGTAVKGKEAVHSDLGYWIESEKNDTWRLKRAHLIEMKTCEWPKSHTLWTDGIEESDLIIPKSLAGPLSHHNTREGYRTQMKGPWHSEELEIRFEECPGTKVHVEETCGTRGPSLRSTTASGRVIEEWCCRECTMPPLSFRAKDGCWYGMEIRPRKEPESNLVRSMVTAGSTDHMDHFSLGVLVILLMVQEGLKKRMTTKIIISTSMAVLVAMILGGFSMSDLAKLAILMGATFAEMNTGGDVAHLALIAAFKVRPALLVSFIFRANWTPRESMLLALASCLLQTAISALEGDLMVLINGFALAWLAIRAMVVPRTDNITLAILAALTPLARGTLLVAWRAGLATCGGFMLLSLKGKGSVKKNLPFVMALGLTAVRLVDPINVVGLLLLTRSGKRSWPPSEVLTAVGLICALAGGFAKADIEMAGPMAAVGLLIVSYVVSGKSVDMYIERAGDITWEKDAEVTGNSPRLDVALDESGDFSLVEDDGPPMREIILKVVLMTICGMNPIAIPFAAGAWYVYVKTGKRSGALWDVPAPKEVKKGETTDGVYRVMTRRLLGSTQVGVGVMQEGVFHTMWHVTKGSALRSGEGRLDPYWGDVKQDLVSYCGPWKLDAAWDGHSEVQLLAVPPGERARNIQTLPGIFKTKDGDIGAVALDYPAGTSGSPILDKCGRVIGLYGNGVVIKNGSYVSAITQGRREEETPVECFEPSMLKKKQLTVLDLHPGAGKTRRVLPEIVREAIKTRLRTVILAPTRVVAAEMEEALRGLPVRYMTTAVNVTHSGTEIVDLMCHATFTSRLLQPIRVPNYNLYIMDEAHFTDPSSIAARGYISTRVEMGEAAAIFMTATPPGTRDAFPDSNSPIMDTEVEVPERAWSSGFDWVTDYSGKTVWFVPSVRNGNEIAACLTKAGKRVIQLSRKTFETEFQKTKHQEWDFVVTTDISEMGANFKADRVIDSRRCLKPVILDGERVILAGPMPVTHASAAQRRGRIGRNPNKPGDEYLYGGGCAETDEDHAHWLEARMLLDNIYLQDGLIASLYRPEADKVAAIEGEFKLRTEQRKTFVELMKRGDLPVWLAYQVASAGITYTDRRWCFDGTTNNTIMEDSVPAEVWTRHGEKRVLKPRWMDARVCSDHAALKSFKEFAAGKRGAAFGVMEALGTLPGHMTERFQEAIDNLAVLMRAETGSRPYKAAAAQLPETLETIMLLGLLGTVSLGIFFVLMRNKGIGKMGFGMVTLGASAWLMWLSEIEPARIACVLIVVFLLLVVLIPEPEKQRSPQDNQMAIIIMVAVGLLGLITANELGWLERTKSDLSHLMGRREEGATMGFSMDIDLRPASAWAIYAALTTFITPAVQHAVTTSYNNYSLMAMATQAGVLFGMGKGMPFYAWDFGVPLLMIGCYSQLTPLTLIVAIILLVAHYMYLIPGLQAAAARAAQKRTAAGIMKNPVVDGIVVTDIDTMTIDPQVEKKMGQVLLMAVAVSSAILSRTAWGWGEAGALITAATSTLWEGSPNKYWNSSTATSLCNIFRGSYLAGASLIYTVTRNAGLVKRRGGGTGETLGEKWKARLNQMSALEFYSYKKSGITEVCREEARRALKDGVATGGHAVSRGSAKLRWLVERGYLQPYGKVIDLGCGRGGWSYYAATIRKVQEVKGYTKGGPGHEEPVLVQSYGWNIVRLKSGVDVFHMAAEPCDTLLCDIGESSSSPEVEEARTLRVLSMVGDWLEKRPGAFCIKVLCPYTSTMMETLERLQRRYGGGLVRVPLSRNSTHEMYWVSGAKSNTIKSVSTTSQLLLGRMDGPRRPVKYEEDVNLGSGTRAVVSCAEAPNMKIIGNRIERIRSEHAETWFFDENHPYRTWAYHGSYEAPTQGSASSLINGVVRLLSKPWDVVTGVTGIAMTDTTPYGQQRVFKEKVDTRVPDPQEGTRQVMSMVSSWLWKELGKHKRPRVCTKEEFINKVRSNAALGAIFEEEKEWKTAVEAVNDPRFWALVDKEREHHLRGECQSCVYNMMGKREKKQGEFGKAKGSRAIWYMWLGARFLEFEALGFLNEDHWMGRENSGGGVEGLGLQRLGYVLEEMSRIPGGRMYADDTAGWDTRISRFDLENEALITNQMEKGHRALALAIIKYTYQNKVVKVLRPAEKGKTVMDIISRQDQRGSGQVVTYALNTFTNLVVQLIRNMEAEEVLEMQDLWLLRRSEKVTNWLQSNGWDRLKRMAVSGDDCVVKPIDDRFAHALRFLNDMGKVRKDTQEWKPSTGWDNWEEVPFCSHHFNKLHLKDGRSIVVPCRHQDELIGRARVSPGAGWSIRETACLAKSYAQMWQLLYFHRRDLRLMANAICSSVPVDWVPTGRTTWSIHGKGEWMTTEDMLVVWNRVWIEENDHMEDKTPVTKWTDIPYLGKREDLWCGSLIGHRPRTTWAENIKNTVNMVRRIIGDEEKYMDYLSTQVRYLGEEGSTPGVL

>tr|H9A910|H9A910_ZIKV Genome polyprotein (Fragment) OSZika virus (strain Mr 766) PE4 SV1

MKNPKKKSGGFRIVNMLKRGVARVSPFGGLKRLPAGLLLGHGPIRMVLAILAFLRFTAIKPSLGLINRWGSVGKKEAMEIIKKFKKDLAAMLRIINARKEKKRRGTDTSVGIVGLLLTTAMAVEVTRRGNAYYMYLDRSDAGEAISFPTTMGMNKCYIQIMDLGHMCDATMSYECPMLDEGVEPDDVDCWCNTTSTWVVYGTCHHKKGEARRSRRAVTLPSHSTRKLQTRSQTWLESREYTKHLIRVENWIFRNPGFALAAAAIAWLLGSSTSQKVIYLVMILLIAPAYSIRCIGVSNRDFVEGMSGGTWVDVVLEHGGCVTVMAQDKPTVDIELVTTTVSNMAEVRSYCYEASISDMASDSRCPTQGEAYLDKQSDTQYVCKRTLVDRGWGNGCGLFGKGSLVTCAKFACSKKMTGKSIQPENLEYRIMLSVHGSQHSGMIVNDTGHETDENRAKVEITPNSPRAEATLGGFGSLGLDCEPRTGLDFSDLYYLTMNNKHWLVHKEWFHDIPLPWHAGADTGTPHWNNKEALVEFKDAHAKRQTVVVLGSQEGAVHTALAGALEAEMDGAKGRLSSGHLKCRLKMDKLRLKGVSYSLCTAAFTFTKIPAETLHGTVTVEVQYAGTDGPCKVPAQMAVDMQTLTPVGRLITANPVITESTENSKMMLELDPPFGDSYIVIGVGEKKITHHWHRSGSTIGKAFEATVRGAKRMAVLGDTAWDFGSVGGALNSLGKGIHQIFGAAFKSLFGGMSWFSQILIGTLLVWLGLNTKNGSISLMCLALGGVLIFLSTAVSADVGCSVDFSKKETRCGTGVFVYNDVEAWRDRYKYHPDSPRRLAAAVKQAWEDGICGISSVSRMENIMWRSVEGELNAILEENGVQLTVVVGSVKNPMWRGPQRLPVPVNELPHGWKAWGKSYFVRAAKTNNSFVVDGDTLKECPLKHRAWNSFLVEDHGFGVFHTSVWLKVREDYSLECDPAVIGTAAKGKEAVHSDLGYWIESEKNDTWRLKRAHLIEMKTCEWPKSHTLWTDGIEESDLIIPKSLAGPLSHHNTREGYRTQMKGPWHSEELEIRFEECPGTKVHVEETCGTRGPSLRSTTASGRVIEEWCCRECTMPPLSFRAKDGCWYGMEIRPRKEPESNLVRSMVTAGSTDHMDHFSLGVLVILLMVQEGLKKRMTTKIIISTSMAVLVAMILGGFSMSDLAKLAILMGATFAEMNTGGDVAHLALIAAFKVRPALLVSFIFRANWTPRESMLLALASCLLQTAISALEGDLMVPINGFALAWLAIRAMVVPRTDNITLAILAALTPLARGTLLVAWRAGLATCGGFMLLSLKGKGSVKKNLPFVMALGLTAVRLVDPINVVGLLLLTRSGKRSWPPSEVLTAVGLICALAGGFAKADIEMAGPMAAVGLLIVSYVVSGKSVDMYIERAGDITWEKDAEVTGNSPRLDVALDESGDFSLVEDDGPPMREIILKVVLMXICGMNPIAIPFAAGAWYVYVKTGKRSGALWDVPAPKEVKKGETTDGVYRVMTRRLLGSTQVGVGVMQEGVFHTMWHVTKGSALRSGEGRLDPYWGDVKQDLVSYCGPWKLDAAWDGHSEVQLLAVPPGERARNIQTLPGIFKTKDGDIGAVALDYPAGTSGSPILDKCGRVIGLYGNGVVIKNGSYVSAITQGRREEETPVECFEPSMLKKKQLTVLDLHPGAGKTRRVLPEIVREAIKTRLRTVILAPTRVVAAEMEEALRGLPVRYMTTAVNVTHSGTEIVDLMCHATFTSRLLQPIRVPNYNLYIMDEAHFTDPSSIAARGYISTRVEMGEAAAIFMTATPPGTRDAFPDSNSPIMDTEVEVPERAWSSGFDWVTDHSGKTVWFVPSVRNGNEIAACLTKAGKRVIQLSRKTFETEFQKTKHQEWDFVVTTDISEMGANFKADRVIDSRRCLKPVILDGERVILAGPMPVTHASAAQRRGRIGRNPNKPGDEYLYGGGCAETDEDHAHWLEARMLLDNIYLQDGLIASLYRPEADKVAAIEGEFKLRTEQRKTFVELMKRGDLPVWLAYQVASAGITYTDRRWCFDGTTNNTIMEDSVPAEVWTRYGEKRVLKPRWMDARVCSDHAALKSFKEFAAGKRGAAFGVMEALGTLPGHMTERFQEAIDNLAVLMRAETGSRPYKAAAAQLPETLETIMLLGLLGTVSLGIFFVLMRNKGIGKMGFGMVTLGASAWLMWLSEIEPARIACVLIVVFLLLVVLIPEPEKQRSPQDNQMAIIIMVAVGLLGLITANELGWLERTKSDLSHLMGRREEGATIGFSMDIDLRPASAWAIYAALTTFITPAVQHAVTTSYNNYSLMAMATQAGVLFGMGKGMPFYAWDFGVPLLMIGCYSQLTPLTLIVAIILLVAHYMYLIPGLQAAAARAAQKRTAAGIMKNPVVDGIVVTDIDTMTIDPQVEKKMGQVLLIAVAVSSAILSRTAWGWGEAGALITAATSTLWEGSPNKYWNSSTATSLCNIFRGSYLAGASLIYTVTRNAGLVKRRGGGTGETLGEKWKARLNQMSALEFYSYKKSGITEVCREEARRALKDGVATGGHAVSRGSAKLRWLVERGYLQPYGKVIDLGCGRGGWSYYAATIRKVQEVKGYTKGGPGHEEPMLVQSYGWNIVRLKSGVDVFHMAAEPCDTLLCDIGESSSSPEVEEARTLRVLSMVGDWLEKRPGAFCIKVLCPYTSTMMETLERLQRRYGGGLVRVPLSRNSTHEMYWVSGAKSNTIKSVSTTSQLLLGRMDGPRRPVKYEEDVNLGSGTRAVVSCAEAPNMKIIGNRIERIRSEHAETWFFDENHPYRTWAYHGSYEAPTQGSASSLINGVVRLLSKPWDVVTGVTGIAMTDTTPYGQQRVFKEKVDTRVPDPQEGTRQVMSMVSSWLWKELGKHKRPRVCTKEEFINKVRSNAALGAIFEEEKEWKTAVEAVNDPRFWALVDKEREHHLRGECQSCVYNMMGKREKKQGEFGKAKGSRAIWYMWLGARFLEFEALGFLNEDHWMGRENSGGGVEGLGLQRLGYVLEEMSRIPGGRMYADDTAGWDTRISRFDLENEALITNQMEKGHRALALAIIKYTYQNKVVKVLRPAEKGKTVMDIISRQDQRGSGQVVTYALNTFTNLVVQLIRNMEAEEVLEMQDLWLLRRSEKVTNWLQSNGWDRLKRMAVSGDDCVVKPIDDRFAHALRFLNDMGKVRKDTQEWKPSTGWDNWEEVPFCSHHFNKLHLKDGRSIVVPCRHQDELIGRARVSPGAGWSIRETACLAKSYAQMWQLLYFHRRDLRLMANAICSSVPVDWVPTGRTTWSIHGKGEWMTTEDMLVVWNRVWIEENDHMEDKTPVTKWTDIPYLGKREDLWCGSLIGHRPRTTWAENIKNTVNMMRRIIGDEEKYVDYLSTQVRYLGEEGSTPGVL

>tr|H8XX10|H8XX10_ZIKV Genome polyprotein (Fragment) OSZika virus (strain Mr 766) PE4 SV1

MKNPKKKSGGFRIVNMLKRGVARVNPLGGLKRLPAGLLLGHGPIRMVLAILAFLRFTAIKPSLGLINRWGSVGKKEAMEIIKKFKKDLAAMLRIINARKERKRRGADTSIGIIGLLLTTAMAAEITRRGSAYYMYLDRSDAGKAISFATTLGVNKCHVQIMDLGHMCDATMSYECPMLDEGVEPDDVDCWCNTTSTWVVYGTCHHKKGEARRSRRAVTLPSHSTRKLQTRSQTWLESREYTKHLIKVENWIFRNPGFALVAVAIAWLLGSSTSQKVIYLVMILLIAPAYSIRCIGVSNRDFVEGMSGGTWVDVVLEHGGCVTVMAQDKPTVDIELVTTTVSNMAEVRSYCYEASISDMASDSRCPTQGEAYLDKQSDTQYVCKRTLVDRGWGNGCGLFGKGSLVTCAKFTCSKKMTGKSIQPENLEYRIMLSVHGSQHSGMIVNDTGYETDENRAKVEVTPNSPRAEATLGGFGSLGLDCEPRTGLDFSDLYYLTMNNKHWLVHKEWFHDIPLPWHAGADTGTPHWNNKEALVEFKDAHAKRQTVVVLGSQEGAVHTALAGALEAEMDGAKGKLFSGHLKCRLKMDKLRLKGVSYSLCTAAFTFTKVPAETLHGTVTVEVQYAGTDGPCKIPVQMAVDMQTLTPVGRLITANPVITESTENSKMMLELDPPFGDSYIVIGVGDKKITHHWHRSGSTIGKAFEATVRGAKRMAVLGDTAWDFGSVGGVFNSLGKGIHQIFGAAFKSLFGGMSWFSQILIGTLLVWLGLNTKNGSISLTCLALGGVMIFLSTAVSADVGCSVDFSKKETRCGTGVFIYNDVEAWRDRYKYHPDSPRRLAAAVKQAWEEGICGISSVSRMENIMWKSVEGELNAILEENGVQLTVVVGSVKNPMWRGPQRLPVPVNELPHGWKAWGKSYFVRAAKTNNSFVVDGDTLKECPLEHRAWNSFLVEDHGFGVFHTSVWLKVREDYSLECDPAVIGTAVKGREAAHSDLGYWIESEKNDTWRLKRAHLIEMKTCEWPKSHTLWTDGVEESDLIIPKSLAGPLSHHNTREGYRTQVKGPWHSEELEIRFEECPGTKVYVEETCGTRGPSLRSTTASGRVIEEWCCRECTMPPLSFRAKDGCWYGMEIRPRKEPESNLVRSMVTAGSTDHMDHFSLGVLVILLMVQEGLKKRMTTKIIMSTSMAVLVVMILGGFSMSDLAKLVILMGATFAEMNTGGDVAHLALVAAFKVRPALLVSFIFRANWTPRESMLLALASCLLQTAISALEGDLMVLINGFALAWLAIRAMAVPRTDNIALPILAALTPLARGTLLVAWRAGLATCGGIMLLSLKGKGSVKKNLPFVMALGLTAVRVVDPINVVGLLLLTRSGKRSWPPSEVLTAVGLICALAGGFAKADIEMAGPMAAVGLLIVSYVVSGKSVDMYIERAGDITWEKDAEVTGNSPRLDVALDESGDFSLVEEDGPPMREIILKVVLMAICGMNPIAIPFAAGAWYVYVKTGKRSGALWDVPAPKEVKKGETTDGVYRVMTRRLLGSTQVGVGVMQEGVFHTMWHVTKGAALRSGEGRLDPYWGDVKQDLVSYCGPWKLDAAWDGLSEVQLLAVPPGERARNIQTLPGIFKTKDGDIGAVALDYPAGTSGSPILDKCGRVIGLYGNGVVIKNGSYVSAITQGKREEETPVECFEPSMLKKKQLTVLDLHPGAGKTRRVLPEIVREAIKKRLRTVILAPTRVVAAEMEEALRGLPVRYMTTAVNVTHSGTEIVDLMCHATFTSRLLQPIRVPNYNLYIMDEAHFTDPSSIAARGYISTRVEMGEAAAIFMTATPPGTRDAFPDSNSPIMDTEVEVPERAWSSGFDWVTDHSGKTVWFVPSVRNGNEIAACLTKAGKRVIQLSRKTFETEFQKTKNQEWDFVITTDISEMGANFKADRVIDSRRCLKPVILDGERVILAGPMPVTHASAAQRRGRIGRNPNKPGDEYMYGGGCAETDEGHAHWLEARMLLDNIYLQDGLIASLYRPEADKVAAIEGEFKLRTEQRKTFVELMKRGDLPVWLAYQVASAGITYTDRRWCFDGTTNNTIMEDSVPAEVWTKYGEKRVLKPRWMDARVCSDHAALKSFKEFAAGKRGAALGVMEALGTLPGHMTERFQEAIDNLAVLMRAETGSRPYKAAAAQLPETLETIMLLGLLGTVSLGIFFVLMRNKGIGKMGFGMVTLGASAWLMWLSEIEPARIACVLIVVFLLLVVLIPEPEKQRSPQDNQMAIIIMVAVGLLGLITANELGWLERTKNDIAHLMGRREEGATMGFSMDIDLRPASAWAIYAALTTLITPAVQHAVTTSYNNYSLMAMATQAGVLFGMGKGMPFYAWDLGVPLLMMGCYSQLTPLTLIVAIILLVAHYMYLIPGLQAAAARAAQKRTAAGIMKNPVVDGIVVTDIDTMTIDPQVEKKMGQVLLIAVAISSAVLLRTAWGWGEAGALITAATSTLWEGSPNKYWNSSTATSLCNIFRGSYLAGASLIYTVTRNAGLVKRRGGGTGETLGEKWKARLNQMSALEFYSYKKSGITEVCREEARRALKDGVATGGHAVSRGSAKLRWLVERGYLQPYGKVVDLGCGRGGWSYYAATIRKVQEVRGYTKGGPGHEEPMLVQSYGWNIVRLKSGVDVFHMAAEPCDTLLCDIGESSSSPEVEETRTLRVLSMVGDWLEKRPGAFCIKVLCPYTSTMMETMERLQRRHGGGLVRVPLSRNSTHEMYWVSGAKSNIIKSVSTTSQLLLGRMDGPRRPVKYEEDVNLGSGTRAVASCAEAPNMKIIGRRIERIRNEHAETWFLDENHPYRTWAYHGSYEAPTQGSASSLVNGVVRLLSKPWDVVTGVTGIAMTDTTPYGQQRVFKEKVDTRVPDPQEGTRQVMNIVSSWLWKELGKRKRPRVCTKEEFINKVRSNAALGAIFEEEKEWKTAVEAVNDPRFWALVDREREHHLRGECHSCVYNMMGKREKKQGEFGKAKGSRAIWYMWLGARFLEFEALGFLNEDHWMGRENSGGGVEGLGLQRLGYILEEMNRAPGGKMYADDTAGWDTRISKFDLENEALITNQMEEGHRTLALAVIKYTYQNKVVKVLRPAEGGKTVMDIISRQDQRGSGQVVTYALNTFTNLVVQLIRNMEAEEVLEMQDLWLLRKPEKVTRWLQSNGWDRLKRMAVSGDDCVVKPIDDRFAHALRFLNDMGKVRKDTQEWKPSTGWSNWEEVPFCSHHFNKLYLKDGRSIVVPCRHQDELIGRARVSPGAGWSIRETACLAKSYAQMWQLLYFHRRDLRLMANAICSAVPVDWVPTGRTTWSIHGKGEWMTTEDMLMVWNRVWIEENDHMEDKTPVTKWTDIPYLGKREDLWCGSLIGHRPRTTWAENIKDTVNMVRRIIGDEEKYMDYLSTQVRYLGEEGSTPGVL

>tr|H8XX13|H8XX13_ZIKV Genome polyprotein (Fragment) OSZika virus (strain Mr 766) PE4 SV1

MKNPKRKSGGFRIVNMLKRGVARVNPLGGLKRLPAGLLLGHGPIRMVLAILAFLRFTAIKPSLGLINRWGSVGKKEAMEIIKKFKKDLAAMLRIINARKERKRRGADTSIGIVGLLLTTAMAAEITRRGSAYYMYLDRSDAGKAISFATTLGVNKCHVQIMDLGHMCDATMSYECPMLDEGVEPDDVDCWCNTTSTWVVYGTCHHKKGEARRSRRAVTLPSHSTRKLQTRSQTWLESREYTKHLIKVENWIFRNPGFALVAVAIAWLLGSSTSQKVIYLVMILLIAPAYSIRCIGVSNRDFVEGMSGGTWVDVVLEHGGCVTVMAQDKPTVDIELVTTTVSNMAEVRSYCYEASISDMASDSRCPTQGEAYLDKQSDTQYVCKRTLVDRGWGNGCGLFGKGSLVTCAKFTCSKKMTGKSIQPENLEYRIMLSVHGSQHSGMIVNDTGHETDENRAKVEVTPNSPRAEATLGGFGSLGLDCEPRTGLDFSDLYYLTMNNKHWLVHKEWFHDIPLPWHAGADTGTPHWNNKEALVEFKDAHAKRQTVVVLGSQEGAVHTALAGALEAEMDGAKGRLFSGHLKCRLKMDKLRLKGVSYSLCTAAFTFTKVPAETLHGTVTVEVQYAGTDGPCKVPAQMAVDMQTLTPVGRLITANPVITESTENSKMMLELDPPFGDSYIVIGVGDKKITHHWHRSGSTIGKAFEATVRGAKRMAVLGDTAWDFGSVGGVFNSLGKGIHQIFGAAFKSLFGGMSWFSQILIGTLLVWLGLNTKNGSISLTCLALGGVMIFLSTAVSADVGCSVDFSKKETRCGTGVFVYNDVEAWRDRYKYHPDSPRRLAAAVKQAWEEGICGISSVSRMENIMWKSVEGELNAILEENGVQLTVVVGSVKNPMWRGPQRLPVPVNELPHGWKAWGKSYFVRAAKTNNSFVVDGDTLKECPLKHRAWNSFLVEDHGFGIFHTSVWLKVREDYSLECDPAVIGTAVKGKEAAHSDLGYWIESEKNDTWRLRRAHLIEMKTCEWPKSHTLWTDGVEESDLIIPKSLAGPLSHHNTREGYRTQLKGPWHSEELEIRFEECPGTKVHVEETCGTRGPSLRSTTASGRVIEEWCCRECTMPPLSFRAKDGCWYGMEIRPRKEPESNLVRSMVTAGSTDHMDHFSLGVLVILLMVQEGLKKRMTTKIIMSTSMAVLVAMVLGGFSMSDLAKLVILMGATFAEMNTGGDVAHLALVAAFKVRPALLVSFIFRANWTPRESMLLALASCLLQTAISALEGELMVLVNGFALAWLAIRAMAVPRTDNIALAILAALTPLARGTLLVAWRAGLATCGGFMLLSLKGKGSVKKNLPFVMALGLTAVRIVDPINVVGLLLLTRSGKRSWPPSEVLTAVGLICALAGGFAKADIEMAGPMAAVGLLIVSYVVSGKSVDMYIERAGDITWEKDAEVTGNSPRLDVALDESGDFSLVEEDGPPMREIILKVVLMAICGMNPIAIPFAAGAWYVYVKTGKRSGALWDVPAPKEVKKGETTDGVYRVMTRRLLGSTQVGVGVMQEGVFHTMWHVTKGAALRSGEGRLDPYWGDVKQDLVSYCGPWKLDAAWDGLSEVQLLAVPPGERARNIQTLPGIFKTKDGDIGAVALDYPAGTSGSPILDKCGRVIGLYGNGVVIKNGSYVSAITQGKREEEAPVECFEPSMLRKKQLTVLDLHPGAGKTRRVLPEIVREAIKKRLRTVILAPTRVVAAEMEEALRGLPVRYMTTAVNVTHSGTEIVDLMCHATFTSRLLQPIRVPNYNLYIMDEAHFTDPSSIAARGYISTRVEMGEAAAIFMTATPPGTRDAFPDSNSPIMDTEVEVPERAWSSGFDWVTDHSGKTIWFVPSVRNGNEIAACLTKAGKRVIQLSRKTFETEFQKTKNQEWDFVITTDISEMGANFKADRVIDSRRCLKPVILDGERVILAGPMPVTHASAAQRRGRVGRNPNKPGDEYMYGGGCAETDEDHAHWLEARMLLDNIYLQDGLIASLYRPEADKVAAIEGEFKLRTEQRKTFVELMKRGDLPVWLAYQVASAGITYTDRRWCFDGTTNNTIMEDSVPAEVWTKYGEKRVLKPRWMDARVCSDHAALKSFKEFAAGKRGAALGVMDALGTLPGHMTERFQEAIDNLAVLMRAETGSRPYKAAAAQLPETLETIMLLGLLGTVSLGIFFVLMRNKGIGKMGFGMVTLGASAWLMWLSEIEPARIACVLIVVFLLLVVLIPEPEKQRSPQDNQMAIIIMVAVGLLGLITANELGWLERTKSDIAHLMGRKEEGTTMGFSMDIDLRPASAWAIYAALTTLITPAVQHAVTTSYNNYSLMAMATQAGVLFGMGKGMPFYAWDFGVPLLMMGCYSQLTPLTLIVAIILLVAHYMYLIPGLQAAAARAAQKRTAAGIMKNPVVDGIVVTDIDTMTIDPQVEKKMGQVLLIAVAVSSAVLLRTAWGWGEAGALITAATSTLWEGSPNKYWNSSTATSLCNIFRGSYLAGASLIYTVTRNAGLVKRRGGGTGETLGEKWKARLNQMSALEFYSYKKSGITEVCREEARRALKDGVATGGHAVSRGSAKLRWLVERGYLQPHGKVVDLGCGRGGWSYYAATIRKVQEVRGYTKGGPGHEEPMLVQSYGWNIVRLKSGVDVFHMAAEPCDTLLCDIGESSSSPEVEETRTLRVLSMVGDWLEKRPGAFCIKVLCPYTSTMMETMERLQRRHGGGLVRVPLSRNSTHEMYWVSGAKSNIIKSVSTTSQLLLGRMEGPRRPVKYEEDVNLGSGTRAVASCAEAPNMKIIGRRIERIRNEHAETWFFDENHPYRTWAYHGSYEAPTQGSASSLVNGVVRLLSKPWDVVTGVTGIAMTDTTPYGQQRVFKEKVDTRVPDPQEGTRQVMNMVSSWLWKELGKRKRPRVCTKEEFINKVRSNAALGAIFEEEKEWKTAVEAVNDPRFWALVDKEREHHLRGECHSCVYNMMGKREKKQGEFGKAKGSRAIWYMWLGARFLEFEALGFLNEDHWMGRENSGGGVEGLGLQRLGYILEEMNRAPGGKMYADDTAGWDTRISKFDLENEALITNQMEEGHRALALAVIKYTYQNKVVKVLRPAEGGKTVMDIISRQDQRGSGQVVTYALNTFTNLVVQLIRNMEAEEVLEMHDLWLLRKPEKVTRWLQSNGWDRLKRMAVSGDDCVVKPIDDRFAHALRFLNDMGKVRKDTQEWKPSTGWSNWEEVPFCSHHFNKLHLKDGRSIVVPCRHQDELIGRARVSPGAGWSIRETACLAKSYAQMWQLLYFHRRDLRLMANAICSAVPVDWVPTGRTTWSIHGKGEWMTTEDMLMVWNRVWIEENDHMEDKTPVTKWTDIPYLGKREDLWCGSLIGHRPRTTWAENIKDTVNMVRRIIGDEEKXMDYLSTQVRYLGEEGSTPGVL

>tr|H8XX11|H8XX11_ZIKV Genome polyprotein (Fragment) OSZika virus (strain Mr 766) PE4 SV1

MKNPKKKSGGFRIVNMLKRGVARVSPFGGLKRLPAGLLLGHGPIRMVLAILAFLRFTAIKPSLGLINRWGSVGKKEAMEIIKKFKKDLAAMLRIINARKEKKRRGADTSVGIVGLLLTTAMAVEVTRRGSAYYMYLDRSDAGEAISFPTTLGVNKCYIQIMDLGHMCDATMSYECPMLDEGVEPDDVDCWCNTTSTWVVYGTCHHKKGEARRSRRAVTLPSHSTRKLQTRSQTWLESREYTKHLIRVENWIFRNPGFALAAAAIAWLLGSSTSQKVIYLVMILLIAPAYSIRCIGVSNRDFVEGMSGGTWVDVVLEHGGCVTVMAQDKPTVDIELVTTTVSNMAEVRSYCYEASISDMASDSRCPTQGEAYLDKQSDTQYVCKRTLVDRGWGNGCGLFGKGSLVTCAKFACSKKMTGKSIQPENLEYRIMLSVHGSQHSGMIVNDXGHETDENRAKVEITPNSPRAEATLGGFGSLGLDCEPRTGLDFSDLYYLTMNNKHWLVHKEWFHDIPLPWHAGADTGTPHWNNKEALVEFKDAHAKRQTVVVLGSQEGAVHTALAGALEAEMDGAKGRLSSGHLKCRLKMDKLRLKGVSYSLCTAAFTFTKIPAETLHGTVTVEVQYAGTDGPCKVPAQMAVDMQTLTPVGRLITANPVITESTENSKMMLELDPPFGDSYIVIGVGDKKITHHWXRSGSTIGKAFEATVRGAKRMAVLGDTAWDFGSVGGALNSLGKGIHQIFGAAFKSLFGGMSWFSQILIGTLLVWLGLNTKNGSISLTCLALGGVLIFLSTAVSADVGCSVDFSKKETRCGTGVFVYNDVEAWRDRYKYHPDSPRRLAAAVKQAWEDGICGISSVSRMENIMWRSVEGELNAILEENGVQLTVVVGSVKNPMWRGPQRLPVPVNELPHGWKAWGKSYFVRAAKTNNSFVVDGDTLKECPLKHRAWNSFLVEDHGFGVFHTSVWLKVREDYSLECDPAVIGTAAKGKEAVHSDLGYWIESEKNDTWRLKRAHLIEMKTCEWPKSHTLWTDGIEESDLIIPKSLAGPLSHHNTREGYRTQVKGPWHSEELEIRFEECPGTKVHVEETCGTRGPSLRSTTASGRVIEEWCCRECTMPPLSFRAKDGCWYGMEIRPRKEPESNLVRSMVTAGSTDHMDHFSLGVLVILLMVQEGLKKRMTTKIIISTSMAVLVAMILGGFSMSDLAKLAILMGATFAEMNTGGDVAHLALIAAFKVRPALLVSFIFRANWTPRESMLLALASCLLQTXISALEGDLMVLINGFALAWLAIRAMAVPRTDNITLAILAALTPLARGTLLVAWRAGLATCGGFMLLSLKGKGSVKKNLPFVMALGLTAVRLVDPINVVGLLLLTRSGKRSWPPSEVLTAVGLICALAGGFAKADIEMAGPMAAVGLLIVSYVVSGKSVDMYIERAGDITWEKDAEVTGNSPRLDVALDESGDFSLVEDDGPPMREIILKVVLMTICGMNPIAIPFAAGAWYVYVKTGKRSGALWDVPAPKEVKKGETTDGVYRVMTRRLLGSTQVGVGVMQEGVFHTMWHVTKGSALRSGEGRLDPYWGDVKQDLVSYCGPWKLDAAWDGHSEVQLLAVPPGERARNIQTLPGIFKTKDGDIGAVALDYPAGTSGSPILDKCGRVIGLYGNGVVIKNGSYVSAITQGRREEETPVECFEPSMLKKKQLTVLDLHPGAGKTRRVLPEIVREAIKTRLRTVILAPTRVVAAEMEEALRGLPVRYMTTAVNVTHSGTEIVDLMCHATFTSRLLQPIRVPNYNLYIMDEAHFTDPSSIAARGYISTRVEMGEAAAIFMTATPPGTRDAFPDSNSPIMDTEVEVPERAWSSGFDWVTDHSGKTVWFVPSVRNGNEIAACLTKAGKRVIQLSRKTFETEFQKTKNQEWDFVVTTDISEMGANFKADRVIDSRRCLKPVILDGERVILAGPMPVTHASAAQRRGRIGRNPNKPGDEYMYGGGCAETDEDHAHWLEARMLLDNIYLQDGLIASLYRPEADKVAAIEGEFKLRTEQRKTFVELMKRGDLPVWLAYQVASAGITYTDRRWCFDGTTNNTIMEDSVPAEVWTRYGEKRVLKPRWMDARVCSDHAALKSFKEFAAGKRGAAFGVMEALGTLPGHMTERFQEAIDNLAVLMRAETGSRPYKAAAAQLPETLETIMLLGLLGTVSLGIFFVLMRNKGIGKMGFGMVTLGASAWLMWLSEIEPARIACVLIVVFLLLVVLIPEPEKQRSPQDNQMAIIIMVAVGLLGLITANELGWLERTKSDLGHLMGRREEGATMGFSMDIDLRPASAWAIYAALTTLITPAVQHAVTTSYNNYSLMAMATQAGVLFGMGKGMPFYAWDFGVPLLMMGCYSQLTPLTLIVAIILLVAHYMYLIPGLQAAAARAAQKRTAAGIMKNPVVDGIVVTDIDTMTIDPQVEKKMGQVLLIAVAISSAVLLRTAWGWGEAGALITAATSTLWEGSPNKYWNSSTATSLCNIFRGSYLAGASLIYTVTRNAGLVKRRGGGTGETLGEKWKARLNQMSALEFYSYKKSGITEVCREEARRALKDGVATGGHAVSRGSAKLRWLVERGYLQPYGKVIDLGCGRGGWSYYAATIRKVQEVKGYTKGGPGHEEPTLVQSYGWNIVRLKSGVDVFHMAAESCDTLLCDIGESSSSPEVEEARTLRVLSMVGDWLEKRPGAFCIKVLCPYTSTMMETLERLQRRYGGGLVRVPLSRNSTHEMYWVSGAKSNIIKSVSTTSQLLLGRMDGPRRPVKYEEDVNLGSGTRAVASCAEAPNLKIIGNRVERIRSEHAETWFFDENHPYRTWAYHGSYEAPTQGSASSLINGVVRLLSKPWDVVTGVTGIAMTDTTPYGQQRVFKEKVDTRVPDPQEGTRQVMNMVSSWLWKELGKHKRPRVCTKEEFINKVRSNAALGAIFEEEKEWKTAVEAVNDPRFWALVDKEREHHLRGECQSCVYNMMGKREKKQGEFGKAKGSRAIWYMWLGARFLEFEALGFLNEDHWMGRENSGGGVEGLGLQRLGYVLEEMSRTPGGKMYADDTAGWDTRISRFDLENEALITNQMEKGHRALALAIIKYTYQNKVVKVLRPAERGKTVMDIISRQDQRGSGQVVTYALNTFTNLVVQLIRNMEAEEVLEMQDLWLLRRPEKVTSWLQSNGWDRLKRMAVSGDDCVVKPIDDRFAHALRFLNDMGKVRKDTQEWKPSTGWSNWEEVPFCSHHFNKLYLKDGRSIVVPCRHQDELIGRARVSPGAGWSIRETACLAKSYAQMWQLLYFHRRDLRLMANAICSSVPVDWVPTGRTTWSIHGKGEWMTTEDMLVVWNRVWIEENDHMEDKTPVTKWTDIPYLGKREDLWCGSLIGHRPRTTWAENIKDTVNMVRRIIGDEEKYMDYLSTQVRYLGEEGSTPGVL

>tr|W8QFH2|W8QFH2_ZIKV Polyprotein OSZika virus (strain Mr 766) PE4 SV1

MKNPKKKSGRFRIVNMLKRGVARVNPLGGLKRLPAGLLLGHGPIRMVLAILAFLRFTAIKPSLGLINRWGSVGKKEAMEIIKKFKKDLAAMLRIINARKERKRRGADTSIGIIGLLLTTAMAAEITRRGSAYYMYLDRSDAGKAISFATTLGVNKCHVQIMDLGHMCDATMSYECPMLDEGVEPDDVDCWCNTTSTWVVYGTCHHKKGETRRSRRSVSLRYHYTRKLQTRSQTWLESREYKKHLIMVENWIFRNPGFAIVSVAITWLMGSLTSQKVIYLVMIVLIVPAYSISCIGVSNRDLVEGMSGGTWVDVVLEHGGCVTEMAQDKPTVDIELVTMTVSNMAEVRSYCYEASLSDMASASRCPTQGEPSLDKQSDTQSVCKRTLGDRGWGNGCGIFGKGSLVTCSKFTCCKKMPGKSIQPENLEYRIMLPVHGSQHSGMIVNDIGHETDENRAKVEVTPNSPRAEATLGGFGSLGLDCEPRTGLDFSDLYYLTMNNKHWLVHKEWFHDIPLPWHAGADTGTPHWNNKEALVEFKDAHAKRQTVVVLGSQEGAVHTALAGALEAEMDGAKGRLFSGHLKCRLKMDKLRLKGVSYSLCTAAFTFTKVPAETLHGTVTVEVQSAGTDGPCKVPAQMAVDMQTLTPVGRLITANPVITESTENSKMMLELDPPFGDSYIVIGVGDKKITHHWHRSGSTIGKAFEATVRGAKRMAVLGDTAWDFGSVGGVFNSLGKGIHQIFGAAFKSLFGGMSWFSQILIGTLLVWLGLNTKNGSISLTCLALGGVMIFLSTAVSADVGCSVDFSKKETRCGTGVFIYNDVEAWRDRYKYHPDSPRRLAAAVKQAWEEGICGISSVSRMENIMWKSVEGELNAILEENGVQLTVVVGSVKNPMWRGPQRLPVPVNXLPHGWKAWGKSYFVRAAKTNNSFVVDGDTLKECPLKHRAWNSFLVEDHGFGVFHTSVWLKVREDYSLECDPAVIGTAVKGREAAHSDLGYWIESEKNDTWRLKRAHLIEMKTCEWPKSHTLWTDGVEESDLIIPKSLAGPLSHHNTREGYRTQVKGPWHSEELEIRFEECPGTKVHVEETCGTRGPSLRSTTASGRVIEEWCCRECTMPPLSFRAKDGCWYGMEIRPRKEPESNLVRSMVTAGSTDHMDHFSLGVLVILLMVQEGLKKRMTTKIIMSTSMAVLVVMILGGFSMSDLAKLVILMGATFAEMNTGGDVAHLALVAAFKVRPALLVSFILRANWTPRESMLLALASCLLQTAISALEGDLMVLVNGFALAWLAIRAMAVPRTDNIALAILAALTPLARGTLLVAWRAGLATCGGFMLLSLKGKGSVKKNLPFVMALGLTAVRVVDPINVVGLLLLTRSGKRSWPPSEVLTAVGLICALAGGFAKADIEMAGPMAAVGLLIVSYVVSGKSVDMYIERAGDITWEKDAEVTGNSPRLDVALDESGDFSLVEEDGPPMREIILKVVLMAICGMNPIAIPFAAGAWYVYVKTGKRSGALWDVPAPKEVKKGETTDGVYRVMTRRLLGSTQVGVGVMQEGVFHTMWHVTKGAALRSGEGRLDPYWGDVKQDLVSYCGPWKLDAAWDGLSEVQLLAVPPGERARNIQTLPGIFKTKDGDIGAVALDYPAGTSGSPILDKCGRVIGLYGNGVVIKNGSYVSAITQGKREEETPVECFEPSMLKKKQLTVLDLHPGAGKTRRVLPEIVREAIKKRLRTVILAPTPVERSYSRWSLWPSVARTQTTAVNVTHSGTEIVDLMCHATFTSRLLQPIRVPNYNLNIMDEAHFTDPSSIAARGYISTRVEMGEAAAIFMTATPPGTRDAFPDSNSPIMDTEVEVPERAWSSGFDWVTDHSGKTVWFVPSVRSGEESAACLTKAGKRVIQLSRKTFETEFQKTKNQEWDFVITTDISEMGANFKADRVIDSRRCLKPVILDGERVILAGPMPVTHASAAQRRGRIGRNPNKPGDEYMYGGGCAETDEDHAHWLEARMLLDNIYLQDGLIASLYRPEADKVAAIEGEFKLRTEQRKTFVELMKRGDLPVWLAYQVASAGITYTDRRWCFDGTTNNTIMEDSVPAEVWTKYGEKRVLKPRWMDARVCSDHAALKSFKEFAAGKRGAALGVMEALGTLPGHMTERFQEAIDNLAVLMRAETGSRPYKAAAAQLPETLETIMLLGLLGTVSLGIFFVLMRNKGIGKMGFGMVTLGASAWLMWLSEIEPARIACVLIVVFLLLVVLIPEPEKQRSPQDNQMAIIIMVAVGLLGLITANELGWLERTKNDIAHLMGRREEGATMGFSMDIDLRPASAWAIYAALTTLITPAVQHAVTTSYNNYSLMAMATQAGVLFGMGKGMPFMHGDLGVPLLMMGCYSQLTPLTLIVAIILLVAHYMYLIPGLQAAAARAAQKRTAAGIMKNPVVDGIVVTDIDTMTIDPQVEKKMGQVLLIAVAISSAVLLRTAWGWGEAGALITAATSTLWEGSPNKYWNSSTATSLCNIFRGSYLAGASLIYTVTRNAGLVKRRGGGTGETLGEKWKARLNQMSALEFYSYKKSGITEVCREEARRALKDGVATGGHAVSRGSAKLRWLVERGYLQPHGKVVDLGCGRGGWSYYAATIRKVQEVRGYTKGGPGHEEPMLVQSYGWNIVRLKSGVDVFHMAAEPCDTLLCDIGESSSSPEVEETRTLRVLSMVGDWLEKRPGAFCIKVLCPYTSTMMETMERLQRRHGGGLVRVPLSRNSTHEMYWVSGAKSNIIKSVSTTSQLLLGRMDGPRRPVKYEEDVNLGSGTRAVASCAEAPNMKIIGRRIERIRNEHAETWFLDENHPYRTWAYHGXYEAPTQGSASSLVNGVVRLLSKPWDVVTGVTGIAMTDTTPYGQQRVFKEKVDTRVPDPQEGTRQVMNMVSSWLWKELGKRKRPRVCTKEEFINKVRSNAALGAIFEEEKEWKTAVEAVNDPRFWALVDREREHHLRGECHSCVYNMMGKREKKQGEFGKAKGSRAIWYMWLGARFLEFESLGFLNEDHWMGRENSGGGVEGLGLQRLGYVLEEMSQAPGGKMYADDTAGWDTRISKFDLENEALITNQMEEGHRTLALAVIKYTYQNKVVKVLRPAEGGKTVMDIISRQDQRGSGQVVTYALNTFTNLVVQLIRNMEAEEVLEMQDLWLLRKPEKVTRWLQSNGWDRLKRMAVSGDDCVVKPIDDRFAHALRFLNDMGKVRKDTQEWKPSTGWSNWEEVPFCSHHFNKLYLKDGRSIVVPCRHQDELIGRARVSPGAGWSIRETACLAKSYAQMWQLLYFHRRDLRLMANAICSAVPVDWVPTGRTTWSIHGKGEWMTTEDMLMVWNRVWIEENDHMEDKTPVTKWTDIPYLGKREDLWCGSLIGHRPRTTWAENIKDTVNMVRRIIGDEEKYMDYLSTQVRYLGEEGSTPGVL

>tr|A0A140FB25|A0A140FB25_ZIKV Genome polyprotein OSZika virus (strain Mr 766) PE4 SV1

MKNPKKKSGGFRIVNMLKRGVARVSPFGGLKRLPAGLLLGHGPIRMVLAILAFLRFTAIKPSLGLINRWGSVGKKEAMEIIKKFKKDLAAMLRIINARKEKKRRGADTSVGIVGLLLTTAMAAEVTRRGSAYYMYLDRNDAGEAISFPTTLGMNKCYIQIMDLGHMCDATMSYECPMLDEGVEPDDVDCWCNTTSTWVVYGTCHHKKGEARRSRRAVTLPSHSTRKLQTRSQTWLESREYTKHLIRVENWIFRNPGFALAAAAIAWLLGSSTSQKVIYLVMILLIAPAYSIRCIGVSNRDFVEGMSGGTWVDVVLEHGGCVTVMAQDKPTVDIELVTTTVSNMAEVRSYCYEASISDMASDSRCPTQGEAYLDKQSDTQYVCKRTLVDRGWGNGCGLFGKGSLVTCAKFACSKKMTGKSIQPENLEYRIMLSVHGSQHSGMIVNDTGHETDENRAKVEITPNSPRAEATLGGFGSLGLDCEPRTGLDFSDLYYLTMNNKHWLVHKEWFHDIPLPWHAGADTGTPHWNNKEALVEFKDAHAKRQTVVVLGSQEGAVHTALAGALEAEMDGAKGRLSSGHLKCRLKMDKLRLKGVSYSLCTAAFTFTKIPAETLHGTVTVEVQYAGTDGPCKVPAQMAVDMQTLTPVGRLITANPVITESTENSKMMLELDPPFGDSYIVIGVGEKKITHHWHRSGSTIGKAFEATVRGAKRMAVLGDTAWDFGSVGGALNSLGKGIHQIFGAAFKSLFGGMSWFSQILIGTLLMWLGLNTKNGSISLMCLALGGVLIFLSTAVSADVGCSVDFSKKETRCGTGVFVYNDVEAWRDRYKYHPDSPRRLAAAVKQAWEDGICGISSVSRMENIMWRSVEGELNAILEENGVQLTVVVGSVKNPMWRGPQRLPVPVNELPHGWKAWGKSYFVRAAKTNNSFVVDGDTLKECPLKHRAWNSFLVEDHGFGVFHTSVWLKVREDYSLECDPAVIGTAVKGKEAVHSDLGYWIESEKNDTWRLKRAHLIEMKTCEWPKSHTLWTDGIEESDLIIPKSLAGPLSHHNTREGYRTQMKGPWHSEELEIRFEECPGTKVHVEETCGTRGPSLRSTTASGRVIEEWCCRECTMPPLSFRAKDGCWYGMEIRPRKEPESNLVRSMVTAGSTDHMDHFSLGVLVILLMVQEGLKKRMTTKIIISTSMAVLVAMILGGFSMSDLAKLAILMGATFAEMNTGGDVAHLALIAAFKVRPALLVSFIFRANWTPRESMLLALASCLLQTAISALEGDLMVLINGFALAWLAVRAMVVPRTDNITLAILAALTPLARGTLLVAWRAGLATCGGFMLLSLKGKGSVKKNLPFVMALGLTAVRLVDPINVVGLLLLTRSGKRSWPPSEVLTAVGLICALAGGFAKADIEMAGPMAAVGLLIVSYVVSGKSVDMYIERAGDITWEKDAEVTGNSPRLDVALDESGDFSLVEDDGPPMREIILKVVLMTICGMNPIAIPFAAGAWYVYVKTGKRSGALWDVPAPKEVKKGETTDGVYRVMTRRLLGSTQVGVGVMQEGVFHTMWHVTKGSALRSGEGRLDPYWGDVKQDLVSYCGPWKLDAAWDGHSEVQLLAVPPGERARNIQTLPGIFKTKDGDIGAVALDYPAGTSGSPILDKCGRVIGLYGNGVVIKNGSYVSAITQGRREEETPVECFEPSMLKKKQLTVLDLHPGAGKTRRVLPEIVREAIKTRLRTVILAPTRVVAAEMEEALRGLPVRYMTTAVNVTHSGTEIVDLMCHATFTSRLLQPIRVPNYNLYIMDEAHFTDPSSIAARGYISTRVEMGEAAAIFMTATPPGTRDAFPDSNSPIMDTEVEVPERAWSSGFDWVTDHSGKTVWFVPSVRNGNEIAACLTKAGKRVIQLSRKTFETEFQKTKHQEWDFVVTTDISEMGANFKADRVIDSRRCLKPVILDGERVILAGPMPVTHASAAQRRGRIGRNPNKPGDEYLYGGGCAETDEDHAHWLEARMLLDNIYLQDGLIASLYRPEADKVAAIEGEFKLRTEQRKTFVELMKRGDLPVWLAYQVASAGITYTDRRWCFDGTTNNTIMEDSVPAEVWTRHGEKRVLKPRWMDARVCSDHAALKSFKEFAAGKRGAAFGVMEALGTLPGHMTERFQEAIDNLAVLMRAETGSRPYKAAAAQLPETLETIMLLGLLGTVSLGIFFVLMRNKGIGKMGFGMVTLGASAWLMWLSEIEPARIACVLIVVFLLLVVLIPEPEKQRSPQDNQMAIIIMVAVGLLGLITANELGWLERTKSDLSHLMGRREEGATIGFSMDIDLRPASAWAIYAALTTFITPAVQHAVTTSYNNYSLMAMATQAGVLFGMGKGMPFYAWDFGVPLLMIGCYSQLTPLTLIVAIILLVAHYMYLIPGLQAAAARAAQKRTAAGIMKNPVVDGIVVTDIDTMTIDPQVEKKMGQVLLIAVAVSSAILSRTAWGWGEAGALITAATSTLWEGSPNKYWNSSTATSLCNIFRGSYLAGASLIYTVTRNAGLVKRRGGGTGETLGEKWKARLNQMSALEFYSYKKSGITEVCREEARRALKDGVATGGHAVSRGSAKLRWLVERGYLQPYGKVIDLGCGRGGWSYYAATIRKVQEVKGYTKGGPGHEEPVLVQSYGWNIVRLKSGVDVFHMAAEPCDTLLCDIGESSSSPEVEEARTLRVLSMVGDWLEKRPGAFCIKVLCPYTSTMMETLERLQRRYGGGLVRVPLSRNSTHEMYWVSGAKSNIIKSVSTTSQLLLGRMDGPRRPVKYEEDVNLGSGTRAVASCAEAPNMKIIGNRIERIRSEHAETWFFDENHPYRTWAYHGSYEAPTQGSASSLINGVVRLLSKPWDVVTGVTGIAMTDTTPYGQQRVFKEKVDTRVPDPQEGTRQVMSMVSSWLWKELGKHKRPRVCTKEEFINKVRSNAALGAIFEEEKEWKTAVEAVNDPRFWALVDKEREHHLRGECQSCVYNMMGKREKKQGEFGKAKGSRAIWYMWLGARFLEFEALGFLNEDHWMGRENSGGGVEGLGLQRLGYVLEEMSRIPGGRMYADDTAGWDTRISRFDLENEALITNQMEKGHRALALAIIKYTYQNKVVKVLRPAEKGKTVMDIISRQDQRGSGQVVTYALNTFTNLVVQLIRNMEAEEVLEMQDLWLLRRSEKVTNWLQSNGWDRLKRMAVSGDDCVVKPIDDRFAHALRFLNDMGKVRKDTQEWKPSTGWDNWEEVPFCSHHFNKLHLKDGRSIVVPCRHQDELIGRARVSPGAGWSIRETACLAKSYAQMWQLLYFHRRDLRLMANAICSSVPVDWVPTGRTTWSIHGKGEWMTTEDMLVVWNRVWIEENDHMEDKTPVTKWTDIPYLGKREDLWCGSLIGHRPRTTWAENIKNTVNMVRRIIGDEEKYMDYLSTQVRYLGEEGSTPGVL

>tr|A0A127AMC3|A0A127AMC3_ZIKV Genome polyprotein OSZika virus (strain Mr 766) PE4 SV1

MKNPKKKSGGFRIVNMLKRGVARVSPFGGLKRLPAGLLLGHGPIRMVLAILAFLRFTAIKPSLGLINRWGSVGKKEAMEIIKKFKKDLAAMLRIINARKEKKRRGADTNVGIVGLLLTTAMAAEVTRRGSAYYMYLDRNDAGEAISFPTTLGMNKCYIQIMDLGHMCDATMSYECPMLDEGVEPDDVDCWCNTTSTWVVYGTCHHKKGEARRSRRAVTLPSHSTRKLQTRSQTWLESREYTKHLIRVENWIFRNPGFALAAAAIAWLLGSSTSQKVIYLVMILLIAPAYSIRCIGVSNRDFVEGMSGGTWVDVVLEHGGCVTVMAQDKPTVDIELVTTTVSNMAEVRSYCYEASISDMASDSRCPTQGEAYLDKQSDTQYVCKRTLVDRGWGNGCGLFGKGSLVTCAKFACSKKMTGKSIQPENLEYRIMLSVHGSQHSGMIVNDTGHETDENRAKVEITPNSPRAEATLGGFGSLGLDCEPRTGLDFSDLYYLTMNNKHWLVHKEWFHDIPLPWHAGADTGTPHWNNKEALVEFKDAHAKRQTVVVLGSQEGAVHTALAGALEAEMDGAKGRLSSGHLKCRLKMDKLRLKGVSYSLCTAAFTFTKIPAETLHGTVTVEVQYAGTDGPCKVPAQMAVDMQTLTPVGRLITANPVITESTENSKMMLELDPPFGDSYIVIGVGEKKITHHWHRSGSTIGKAFEATVRGARRMAVLGDTAWDFGSVGGALNSLGKGIHQIFGAAFKSLFGGMSWFSQILIGTLLMWLGLNTKNGSISLMCLALGGVLIFLSTAVSADVGCSVDFSKKETRCGTGVFVYNDVEAWRDRYKYHPDSPRRLAAAVKQAWEDGICGISSVSRMENIMWRSVEGELNAILEENGVQLTVVVGSVKNPMWRGPQRLPVPVNELPHGWKAWGKSYFVRAAKTNNSFVVDGDTLKECPLKHRAWNSFLVEDHGFGVFHTSVWLKVREDYSLECDPAVIGTAVKGKEAVHSDLGYWIESEKNDTWRLKRAHLIEMKTCEWPKSHTLWTDGIEESDLIIPKSLAGPLSHHNTREGYRTQMKGPWHSEELEIRFEECPGTKVHVEETCGTRGPSLRSTTASGRVIEEWCCRECTMPPLSFQAKDGCWYGMEIRPRKEPESNLVRSMVTAGSTDHMDHFSLGVLVILLMVQEGLKKRMTTKIIISTSMAVLVAMILGGFSMSDLAKLAILMGATFAEMNTGGDVAHLALIAAFKVRPALLVSFIFRANWTPRESMLLALASCLLQTAISALEGDLMVLINGFALAWLAIRAMVVPRTDNITLAILAALTPLARGTLLVAWRAGLATCGGFMLLSLKGKGSVKKNLPFVMALGLTAVRLVDPINVVGLLLLTRSGKRSWPPSEVLTAVGLICALAGGFAKADIEMAGPMAAVGLLIVSYVVSGKSVDMYIERAGDITWEKDAEVTGNSPRLDVALDESGDFSLVEDDGPPMREIILKVVLMTICGMNPIAIPFAAGAWYVYVKTGKRSGALWDVPAPKEVKKGETTDGVYRVMTRRLLGSTQVGVGVMQEGVFHTMWHVTKGSALRSGEGRLDPYWGDVKQDLVSYCGPWKLDAAWDGHSEVQLLAVPPGERARNIQTLPGIFKTKDGDIGAVALDYPAGTSGSPILDKCGRVIGLYGNGVVIKNGSYVSAITQGRREEETPVECFEPSMLKKKQLTVLDLHPGAGKTRRVLPEIVREAIKTRLRTVILAPTRVVAAEMEEALRGLPVRYMTTAVNVTHSGTEIVDLMCHATFTSRLLQPIRVPNYNLYIMDEAHFTDPSSIAARGYISTRVEMGEAAAIFMTATPPGTRDAFPDSNSPIMDTEVEVPERAWSSGFDWVTDHSGKTVWFVPSVRNGNEIAACLTKAGKRVIQLSRKTFETEFQKTKHQEWDFVVTTDISEMGANFKADRVIDSRRCLKPVILDGERVILAGPMPVTHASAAQRRGRIGRNPNKPGDEYLYGGGCAETDEDHAHWLEARMLLDNIYLQDGLIASLYRPEADKVAAIEGEFKLRTEQRKTFVELMKRGDLPVWLAYQVASAGITYTDRRWCFDGTTNNTIMEDSVPAEVWTRHGEKRVLKPRWMDARVCSDHAALKSFKEFAAGKRGAAFGVMEALGTLPGHMTERFQEAIDNLAVLMRAETGSRPYKAAAAQLPETLETIMLLGLLGTVSLGIFFVLMRNKGIGKMGFGMVTLGASAWLMWLSEIEPARIACVLIVVFLLLVVLIPEPEKQRSPQDNQMAIIIMVAVGLLGLITANELGWLERTKSDLSHLMGRREEGATIGFSMDIDLRPASAWAIYAALTTFITPAVQHAVTTSYNNYSLMAMATQAGVLFGMGKGMPFYAWDFGVPLLMIGCYSQLTPLTLIVAIILLVAHYMYLIPGLQAAAARAAQKRTAAGIMKNPVVDGIVVTDIDTMTIDPQVEKKMGQVLLIAVAVSSAILSRTAWGWGEAGALITAATSTLWEGSPNKYWNSSTATSLCNIFRGSYLAGASLIYTVTRNAGLVKRRGGGTGETLGEKWKARLNQMSALEFYSYKKSGITEVCREEARRALKDGVATGGHAVSRGSAKLRWLVERGYLQPYGKVIDLGCGRGGWSYYAATIRKVQEVKGYTKGGPGHEEPMLVQSYGWNIVRLKSGVDVFHMAAEPCDTLLCDIGESSSSPEVEEARTLRVLSMVGDWLEKRPGAFCIKVLCPYTSTMMETLERLQRRYGGGLVRVPLSRNSTHEMYWVSGAKSNTIKSVSTTSQLLLGRMDGPRRPVKYEEDVNLGSGTRAVVSCAEAPNMKIIGNRIERIRSEHAETWFFDENHPYRTWAYHGSYEAPTQGSASSLINGVVRLLSKPWDVVTGVTGIAMTDTTPYGQQRVFKEKVDTRVPDPQEGTRQVMSMVSSWLWKELGKHKRPRVCTKEEFINKVRSNAALGAIFEEEKEWKTAVEAVNDPRFWALVDKEREHHLRGECQSCVYNMMGKREKKQGEFGKAKGSRAIWYMWLGARFLEFEALGFLNEDHWMGRENSGGGVEGLGLQRLGYVLEEMSRIPGGRMYADDTAGWDTRISRFDLENEALITNQMEKGHRALALAIIKYTYQNKVVKVLRPAEKGKTVMDIISRQDQRGSGQVVTYALNTFTNLVVQLIRSMEAEEVLEMQDLWLLRRSEKVTNWLQSNGWDRLKRMAVSGDDCVVRPIDDRFAHALRFLNDMGKVRKDTQEWKPSTGWDNWEEVPFCSHHFNKLHLKDGRSIVVPCRHQDELIGRARVSPGAGWSIRETACLAKSYAQMWQLLYFHRRDLRLMANAICSSVPVDWVPTGRTTWSIHGKGEWMTTEDMLVVWNRVWIEENDHMEDKTPVTKWTDIPYLGKREDLWCGSLIGHRPRTTWAENIKNTVNMVRRIIGDEEKYMDYLSTQVRYLGEEGSTPGVL

>tr|A0A140GMI4|A0A140GMI4_ZIKV Genome polyprotein OSZika virus (strain Mr 766) PE4 SV1

MKNPKKKSGGFRIVNMLKRGVARVSPFGGLKRLPAGLLLGHGPIRMVLAILAFLRFTAIKPSLGLINRWGSVGKKEAMEIIKKFKKDLAAMLRIINARKEKKRRGADTSVGIVGLLLTTAMAAEVTRRGSAYYMYLDRNDAGEAISFPTTLGMNKCYIQIMDLGHMCDATMSYECPMLDEGVEPDDVDCWCNTTSTWVVYGTCHHKKGEARRSRRAVTLPSHSTRKLQTRSQTWLESREYTKHLIRVENWIFRNPGFALAAAAIAWLLGSSTSQKVIYLVMILLIAPAYSIRCIGVSNRDFVEGMSGGTWVDVVLEHGGCVTVMAQDKPTVDIELVTTTVSNMAEVRSYCYEASISDMASDSRCPTQGEAYLDKQSDTQYVCKRTLVDRGWGNGCGLFGKGSLVTCAKFACSKKMTGKSIQPENLEYRIMLSVHGSQHSGMIVNDTGHETDENRAKVEITPNSPRAEATLGGFGSLGLDCEPRTGLDFSDLYYLTMNNKHWLVHKEWFHDIPLPWHAGADTGTPHWNNKEALVEFKDAHAKRQTVVVLGSQEGAVHTALAGALEAEMDGAKGRLSSGHLKCRLKMDKLRLKGVSYSLCTAAFTFTKIPAETLHGTVTVEVQYAGTDGPCKVPAQMAVDMQTLTPVGRLITANPVITESTENSKMMLELDPPFGDSYIVIGVGEKKITHHWHRSGSTIGKAFEATVRGAKRMAVLGDTAWDFGSVGGALNSLGKGIHQIFGAAFKSLFGGMSWFSQILIGTLLMWLGLNTKNGSISLMCLALGGVLIFLSTAVSADVGCSVDFSKKETRCGTGVFVYNDVEAWRDRYKYHPDSPRRLAAAVKQAWEDGICGISSVSRMENIMWRSVEGELNAILEENGVQLTVVVGSVKNPMWRGPQRLPVPVNELPHGWKAWGKSYFVRAAKTNNSFVVDGDTLKECPLKHRAWNSFLVEDHGFGVFHTSVWLKVREDYSLECDPAVIGTAVKGKEAVHSDLGYWIESEKNDTWRLKRAHLIEMKTCEWPKSHTLWTDGIEESDLIIPKSLAGPLSHHNTREGYRTQMKGPWHSEELEIRFEECPGTKVHVEETCGTRGPSLRSTTASGRVIEEWCCRECTMPPLSFRAKDGCWYGMEIRPRKEPESNLVRSVVTAGSTDHMDHFSLGVLVILLMVQEGLKKRMTTKIIISTSMAVLVAMILGGFSMSDLAKLAILMGATFAEMNTGGDVAHLALIAAFKVRPALLVSFIFRANWTPRESMLLALASCLLQTAISALEGDLMVLINGFALAWLAIRAMVVPRTDNITLAILAALTPLARGTLLVAWRAGLATCGGFMLLSLKGKGSVKKNLPFVMALGLTAVRLVDPINVVGLLLLTRSGKRSWPPSEVLTAVGLICALAGGFAKADIEMAGPMAAVGLLIVSYVVSGKSVDMYIERAGDITWEKDAEVTGNSPRLDVALDESGDFSLVEDDGPPMREIILKVVLMTICGMNPIAIPFAAGAWYVYVKTGKRSGALWDVPAPKEVKKGETTDGVYRVMTRRLLGSTQVGVGVMQEGVFHTMWHVTKGSALRSGEGRLDPYWGDVKQDLVSYCGPWKLDAAWDGHSEVQLLAVPPGERARNIQTLPGIFKTKDGDIGAVALDYPAGTSGSPILDKCGRVIGLYGNGVVIKNGSYVSAITQGRREEETPVECFEPSMLKKKQLTVLDLHPGAGKTRRVLPEIVREAIKTRLRTVILAPTRVVAAEMEEALRGLPVRYMTTAVNVTHSGTEIVDLMCHATFTSRLLQPIRVPNYNLYIMDEAHFTDPSSIAARGYISTRVEMGEAAAIFMTATPPGTRDAFPDSNSPIMDTEVEVPERAWSSGFDWVTDHSGKTVWFVPSVRNGNEIAACLTKAGKRVIQLSRKTFETEFQKTKHQEWDFVVTTDISEMGANFKADRVIDSRRCLKPVILDGERVILAGPMPVTHASAAQRRGRIGRNPNKPGDEYLYGGGCAETDEDHAHWLEARMLLDNIYLQDGLIASLYRPEADKVAAIEGEFKLRTEQRKTFVELMKRGDLPVWLAYQVASAGITYTDRRWCFDGTTNNTIMEDSVPAEVWTRHGEKRVLKPRWMDARVCSDHAALKSFKEFAAGKRGAAFGVMEALGTLPGHMTERFQEAIDNLAVLMRAETGSRPYKAAAAQLPETLETIMLLGLLGTVSLGIFFVLMRNKGIGKMGFGMVTLGASAWLMWLSEIEPARIACVLIVVFLLLVVLIPEPEKQRSPQDNQMAIIIMVAVGLLGLITANELGWLERTKSDLSHLMGRREEGATIGFSMDIDLRPASAWAIYAALTTFITPAVQHAVTTSYNNYSLMAMATQAGVLFGMGKGMPFYAWDFGVPLLMIGCYSQLTPLTLIVAIILLVAHYMYLIPGLQAAAARAAQKRTAAGIMKNPVVDGIVVTDIDTMTIDPQVEKKMGQVLLIAVAVSSAILSRTAWGWGEAGALITAATSTLWEGSPNKYWNSSTATSLCNIFRGSYLAGASLIYTVTRNAGLVKRRGGGTGETLGEKWKARLNQMSALEFYSYKKSGITEVCREEARRALKDGVATGGHAVSRGSAKLRWLVERGYLQPYGKVIDLGCGRGGWSYYAATIRKVQEVKGYTKGGPGHEEPVLVQSYGWNIVRLKSGVDVFHMAAEPCDTLLCDIGESSSSPEVEEARTLRVLSMVGDWLEKRPGAFCIKVLCPYTSTMMETLERLQRRYGGGLVRVPLSRNSTHEMYWVSGAKSNTIKSVSTTSQLLLGRMDGPRRPVKYEEDVNLGSGTRAVVSCAEAPNMKIIGNRIERIRSEHAETWFFDENHPYRTWAYHGSYEAPTQGSASSLVNGVVRLLSKPWDVVTGVTGIAMTDTTPYGQQRVFKEKVDTRVPDPQEGTRQVMSMVSSWLWKELGKHKRPRVCTKEEFINKVRSNAALGAIFEEEKEWKTAVEAVNDPRFWALVDKEREHHLRGECQSCVYNMMGKREKKQGEFGKAKGSRAIWYMWLGARFLEFEALGFLNEDHWMGRENSGGGVEGLGLQRLGYVLEEMSRIPGGRMYADDTAGWDTRISRFDLENEALITNQMEKGHRALALAIIKYTYQNKVVKVLRPAEKGKTVMDIISRQDQRGSGQVVTYALNTFTNLVVQLIRNMEAEEVLEMQDLWLLRRSEKVTNWLQSNGWDRLKRMAVSGDDCVVKPIDDRFAHALRFLNDMGKVRKDTQEWKPSTGWDNWEEVPFCSHHFNKLHLKDGRSIVVPCRHQDELIGRARVSPGAGWSIRETACLAKSYAQMWQLLYFHRRDLRLMANAICSSVPVDWVPTGRTTWSIHGKGEWMTTEDMLVVWNRVWIEENDHMEDKTPVTKWTDIPYLGKREDLWCGSLIGHRPRTTWAENIKNTVNMVRRIIGEEEKYMDYLSTQVRYLGEEGSTPGVL

>tr|A0A140CFV0|A0A140CFV0_ZIKV Genome polyprotein OSZika virus (strain Mr 766) PE4 SV1

MKNPKKKSGGFRIVNMLKRGVARVSPFGGLKRLPAGLLLGHGPIRMVLAILAFLRFTAIKPSLGLINRWGSVGKKEAMEIIKKFKKDLAAMLRIINARKEKKRRGADTSVGIVGLLLTTAMAAEVTRRGSAYYMYLDRNDAGEAISFPTTLGMNKCYIQIMDLGHMCDATMSYECPMLDEGVEPDDVDCWCNTTSTWVVYGTCHHKKGEARRSRRAVTLPSHSTRKLQTRSQTWLESREYTKHLIRVENWIFRNPGFALAAAAIAWLLGSSTSQKVIYLVMILLIAPAYSIRCIGVSNRDFVEGMSGGTWVDVVLEHGGCVTVMAQDKPTVDIELVTTTVSNMAEVRSYCYEASISDMASDSRCPTQGEAYLDKQSDTQYVCKRTLVDRGWGNGCGLFGKGSLVTCAKFACSKKMTGKSIQPENLEYRIMLSVHGSQHSGMIVNDTGHETDENRAKVEITPNSPRAEATLGGFGSLGLDCEPRTGLDFSDLYYLTMNNKHWLVHKEWFHDIPLPWHAGADTGTPHWNNKEALVEFKDAHAKRQTVVVLGTQEGAVHTALAGALEAEMDGAKGRLSSGHLKCRLKMDKLRLKGVSYSLCTAAFTFTKIPAETLHGTVTVEVQYAGTDGPCKVPAQMAVDMQTLTPVGRLITANPVITESTENSKMMLELDPPFGDSYIVIGVGEKKITHHWHRSGSTIGKAFEATVRGAKRMAVLGDTAWDFGSVGGALNSLGKGIHQIFGAAFKSLFGGMSWFSQILIGTLLMWLGLNTKNGSISLMCLALGGVLIFLSTAVSADVGCSVDFSKKETRCGTGVFVYNDVEAWRDRYKYHPDSPRRLAAAVKQAWEDGICGISSVSRMENIMWRSVEGELNAILEENGVQLTVVVGSVKNPMWRGPQRLPVPVNELPHGWKAWGKSYFVRAAKTNNSFVVDGDTLKECPLKHRAWNSFLVEDHGFGVFHTSVWLKVREDYSLECDPAVIGTAVKGKEAVHSDLGYWIESEKNDTWRLKRAHLIEMKTCEWPKSHTLWTDGIEESDLIIPKSLAGPLSHHNTREGYRTQMKGPWHSEELEIRFEECPGTKVHVEETCGTRGPSLRSTTASGRVIEEWCCRECTMPPLSFRAKDGCWYGMEIRPRKEPESNLVRSMVTAGSTDHMDHFSLGVLVILLMVQEGLKKRMTTKIIISTSMAVLVAMILGGFSMSDLAKLAILMGATFAEMNTGGDVAHLALIAAFKVRPALLVSFIFRANWTPRESMLLALASCFLQTAISALEGDLMVLINGFALAWLAIRAMVVPRTDNITLAILAALTPLARGTLLVAWRAGLATCGGFMLLSLKGKGSVKKNLPFVMALGLTAVRLVDPINVVGLLLLTRSGKRSWPPSEVLTAVGLICALAGGFAKADIEMAGPMAAVGLLIVSYVVSGKSVDMYIERAGDITWEKDAEVTGNSPRLDVALDESGDFSLVEDDGPPMREIILKVVLMTICGMNPIAIPFAAGAWYVYVKTGKRSGALWDVPAPKEVKKGETTDGVYRVMTRRLLGSTQVGVGVMQEGVFHTMWHVTKGSALRSGEGRLDPYWGDVKQDLVSYCGPWKLDAAWDGHSEVQLLAVPPGERARNIQTLPGIFKTKDGDIGAVALDYPAGTSGSPILDKCGRVIGLYGNGVVIKNGSYVSAITQGRREEETPVECFEPSMLKKKQLTVLDLHPGAGKTRRVLPEIVREAIKTRLRTVILAPTRVVAAEMEEALRGLPVRYMTTAVNVTHSGTEIVDLMCHATFTSRLLQPIRVPNYNLYIMDEAHFTDPSSIAARGYISTRVEMGEAAAIFMTATPPGTRDAFPDSNSPIMDTEVEVPERAWSSGFDWVTDHSGKTVWFVPSVRNGNEIAACLTKAGKRVIQLSRKTFETEFQKTKHQEWDFVVTTDISEMGANFKADRVIDSRRCLKPVILDGERVILAGPMPVTHASAAQRRGRIGRNPNKPGDEYLYGGGCAETDEDHAHWLEARMLLDNIYLQDGLIASLYRPEADKVAAIEGEFKLRTEQRKTFVELMKRGDLPVWLAYQVASAGITYTDRRWCFDGTTNNTIMEDSVPAEVWTRHGEKRVLKPRWMDARVCSDHAALKSFKEFAAGKRGAAFGVMEALGTLPGHMTERFQEAIDNLAVLMRAETGSRPYKAAAAQLPETLETIMLLGLLGTVSLGIFFVLMRNKGIGKMGFGMVTLGASAWLMWLSEIEPARIACVLIVVFLLLVVLIPEPEKQRSPQDNQMAIIIMVAVGLLGLITANELGWLERTKSDLSHLMGRREEGATIGFSMDIDLRPASAWAIYAALTTFITPAVQHAVTTSYNNYSLMAMATQAGVLFGMGKGMPFYAWDFGVPLLMIGCYSQLTPLTLIVAIILLVAHYMYLIPGLQAAAARAAQKRTAAGIMKNPVVDGIVVTDIDTMTIDPQVEKKMGQVLLIAVAVSSAILSRTAWGWGEAGALITAATSTLWEGSPNKYWNSSTATSLCNIFRGSYLAGASLIYTVTRNAGLVKRRGGGTGETLGEKWKARLNQMSALEFYSYKKSGITEVCREEARRALKDGVATGGHAVSRGSAKLRWLVERGYLQPYGKVIDLGCGRGGWSYYAATIRKVQEVKGYTKGGPGHEEPVLVQSYGWNIVRLKSGVDVFHMAAEPCDTLLCDIGESSSSPEVEEARTLRVLSMVGDWLEKRPGAFCIKVLCPYTSTMMETLERLQRRYGGGLVRVPLSRNSTHEMYWVSGAKSNTIKSVSTTSQLLLGRMDGPRRPVKYEEDVNLGSGTRAVVSCAEAPNMKIIGNRIERIRSEHAETWFFDENHPYRTWAYHGSYVAPTQGSASSLINGVVRLLSKPWDVVTGVTGIAMTDTTPYGQQRVFKEKVDTRVPDPQEGTRQVMSMVSSWLWKELGKHKRPRVCTKEEFINKVRSNAALGAIFEEEKEWKTAVEAVNDPRFWALVDKEREHHLRGECQSCVYNMMGKREKKQGEFGKAKGSRAIWYMWLGARFLEFEALGFLNEDHWMGRENSGGGVEGLGLQRLGYVLEEMSRIPGGRMYADDTAGWDTRISRFDLENEALITNQMEKGHRALALAIIKYTYQNKVVKVLRPAEKGKTVMDIISRQDQRGSGQVVTYALNTFTNLVVQLIRNMEAEEVLEMQDLWLLRRSEKVTNWLQSNGWDRLKRMAVSGDDCVVKPIDDRFAHALRFLNDMGKVRKDTQEWKPSTGWDNWEEVPFCSHHFNKLHLKDGRSIVVPCRHQDELIGRARVSPGAGWSIRETACLAKSYAQMWQLLYFHRRDLRLMANAICSSVPVDWVPTGRTTWSIHGKGEWMTTEDMLVVWNRVWIEENDHMEDKTPVTKWTDIPYLGKREDLWCGSLIGHRPRTTWAENIKNTVNMVRRIIGDEEKYMDYLSTQVRYLGEEGSTPGVL

>tr|A0A140DLX5|A0A140DLX5_ZIKV Genome polyprotein OSZika virus (strain Mr 766) PE4 SV1

MKNPKKKSGGFRIVNMLKRGVARVSPFGGLKRLPAGLLLGHGPIRMVLAILAFLRFTAIKPSLGLINRWGSVGKKEAMEIIKKFKKDLAAMLRIINARKEKKRRSADTSVGIVGLLLTTAMAAEVTRRGSAYYMYLDRNDAGEAISFPTTLGMNKCYIQIMDLGHMCDATMSYECPMLDEGVEPDDVDCWCNTTSTWVVYGTCHHKKGEARRSRRAVTLPSHSTRKLQTRSQTWLESREYTKHLIRVENWIFRNPGFALAAAAIAWLLGSSTSQKVIYLVMILLIAPAYSIRCIGVSNRDFVEGMSGGTWVDVVLEHGGCVTVMAQDKPTVDIELVTTTVSNMAEVRSYCYEASISDMASDSRCPTQGEAYLDKQSDTQYVCKRTLVDRGWGNGCGLFGKGSLVTCAKFACSKKMTGKSIQPENLEYRIMLSVHGSQHSGMIVNDTGHETDENRAKVEITPNSPRAEATLGGFGSLGLDCEPRTGLDFSDLYYLTMNNKHWLVHKEWFHDIPLPWHAGADTGTPHWNNKEALVEFKDAHAKRQTVVVLGSQEGAVHTALAGALEAEMDGAKGRLSSGHLKCRLKMDKLRLKGVSYSLCTAAFTFTKIPAETLHGTVTVEVQYAGTDGPCKVPAQMAVDTQTLTPVGRLITANPVITESTENSKMMLELDPPFGDSYIVIGVGEKKITHHWHRSGSTIGKAFEATVRGAKRMAVLGDTAWDFGSVGGALNSLGKGIHQIFGAAFKSLFGGMSWFSQILIGTLLMWLGLNTKNGSISLMCLALGGVLIFLSTAVSADVGCSVDFSKKETRCGTGVFVYNDVEAWRDRYKYHPDSPRRLAAAVKQAWEDGICGISSVSRMENIMWRSVEGELNAILEENGVQLTVVVGSVKNPMWRGPQRLPVPVNELPHGWKAWGKSYFVRAAKTNNSFVVDGDTLKECPLKHRAWNSFLVEDHGFGVFHTSVWLKVREDYSLECDPAVIGTAVKGKEAVHSDLGYWIESEKNDTWRLKRAHLIEMKTCEWPKSHTLWTDGIEESDLIIPKSLAGPLSHHNTREGYRTQMKGPWHSEELEIRFEECPGTKVHVEETCGTRGPSLRSTTASGRVIEEWCCRECTMPPLSFRAKDGCWYGMEIRPRKEPESNLVRSMVTAGSTDHMDHFSLGVLVILLMVQEGLKKRMTTKIIISTSMAVLVAMILGGFSMSDLAKLAILMGATFAEMNTGGDVAHLALIAAFKVRPALLVSFIFRANWTPRESMLLALASCLLQTAISALEGDLMVLINGFALAWLAIRAMVVPRTDNITLAILAALTPLARGTLLVAWRAGLATCGGFMLLSLKGKGSVKKNLPFVMALGLTAVRLVDPINVVGLLLLTRSGKRSWPPSEVLTAVGLICALAGGFAKADIEMAGPMAAVGLLIVSYVVSGKSVDMYIERAGDITWEKDAEVTGNSPRLDVALDESGDFSLVEDDGPPMREIILKVVLMTICGMNPIAIPFAAGAWYVYVKTGKRSGALWDVPAPKEVKKGETTDGVYRVMTRRLLGSTQVGVGVMQEGVFHTMWHVTKGSALRSGEGRLDPYWGDVKQDLVSYCGPWKLDAAWDGHSEVQLLAVPPGERARNIQTLPGIFKTKDGDIGAVALDYPAGTSGSPILDKCGRVIGLYGNGVVIKNGSYVSAITQGRREEETPVECFEPSMLKKKQLTVLDLHPGAGKTRRVLPEIVREAIKTRLRTVILAPTRVVAAEMEEALRGLPVRYMTTAVNVTHSGTEIVDLMCHATFTSRLLQPIRVPNYNLYIMDEAHFTDPSSIAARGYISTRVEMGEAAAIFMTATPPGTRDAFPDSNSPITDTEVEVPERAWSSGFDWVTDHSGKTVWFVPSVRNGNEIAACLTKAGKRVIQLSRKTFETEFQKTKHQEWDFVVTTDISEMGANFKADRVIDSRRCLKPVILDGERVILAGPMPVTHASAAQRRGRIGRNPNKPGDEYLYGGGCAETDEDHAHWLEARMLLDNIYLQDGLIASLYRPEADKVAAIEGEFKLRTEQRKTFVELMKRGDLPVWLAYQVASAGITYTDRRWCFDGTTNNTIMEDSVPAEVWTRHGEKRVLKPRWMDARVCSDHAALKSFKEFAAGKRGAAFGVMEALGTLPGHMTERFQEAIDNLAVLMRAETGSRPYKAAAAQLPETLETIMLLGLLGTVSLGIFFVLMRNKGIGKMGFGMVTLGASAWLMWLSEIEPARIACVLIVVFLLLVVLIPEPEKQRSPQDNQMAIIIMVAVGLLGLITANELGWLERTKSDLSHLMGRREEGATIGFSMDIDLRPASAWAIYAALTTFITPAVQHAVTTSYNNYSLMAMATQAGVLFGMGKGMPFYAWDFGVPLLMIGCYSQLTPLTLIVAIILLVAHYMYLIPGLQAAAARAAQKRTAAGIMKNPVVDGIVVTDIDTMTIDPQVEKKMGQVLLIAVAVSSAILSRTAWGWGEAGALITAATSTLWEGSPNKYWNSSTATSLCNIFRGSYLAGASLIYTVTRNAGLVKRRGGGTGETLGEKWKARLNQMSALEFYSYKKSGITEVCREEARRALKDGVATGGHAVSRGSAKLRWLVERGYLQPYGKVIDLGCGRGGWSYYAATIRKVQEVKGYTKGGPGHEEPVLVQSYGWNIVRLKSGVDVFHMAAEPCDTLLCDIGESSSSPEVEEARTLRVLSMVGDWLEKRPGAFCIKVLCPYTSTMMETLERLQRRYGGGLVRVPLSRNSTHEMYWVSGAKSNTIKSVSTTSQLLLGRMDGPRRPVKYEEDVNLGSGTRAVVSCAEAPNMKIIGNRIERIRSEHAETWFFDENHPYRTWAYHGSYEAPTQGSASSLINGVVRLLSKPWDVVTGVTGIAMTDTTPYGQQRVFKEKVDTRVPDPQEGTRQVMSMVSSWLWKELGKHKRPRVCTKEEFINKVRSNAALGAIFEEEKEWKTAVEAVNDPRFWALVDKEREHHLRGECQSCVYNMMGKREKKQGEFGKAKGSRAIWYMWLGARFLEFEALGFLNEDHWMGRENSGGGVEGLGLQRLGYVLEEMSRIPGGRMYADDTAGWDTRISRFDLENEALITNQMEKGHRALALAIIKYTYQNKVVKVLRPAEKGKTVMDIISRQDQRGSGQVVTYALNTFTNLVVQLIRNMEAEEVLEMQDLWLLRRSEKVTNWLQSNGWDRLKRMAVSGDDCVVKPIDDRFAHALRFLNDMGKVRKDTQEWKPSTGWDNWEEVPFCSHHFNKLHLKDGRSIVVPCRHQDELIGRARVSPGAGWSIRETACLAKSYAQMWQLLYFHRRDLRLMANAICSSVPVDWVPTGRTTWSIHGKGEWMTTEDMLVVWNRVWIEENDHMEDKTPVTKWTDIPYLGKREDLWCGSLIGHRPRTTWAENIKNTVNMVRRIIGDEEKYMDYLSTQVRYLGEEGSTPGVL

>tr|A0A140GMI3|A0A140GMI3_ZIKV Genome polyprotein OSZika virus (strain Mr 766) PE4 SV1

MKNPKKKSGGFRIVNMLKRGVARVSPFGGLKRLPAGLLLGHGPIRMVLAILAFLRFTAIKPSLGLINRWGSVGKKEAMEIIKKFKKDLAAMLRIINARKEKKRRGADTSVGIVGLLLTTAMAAEVTRRGSAYYMYLDRNDAGEAISFPTTLGMNKCYIQIMDLGHMCDATMSYECPMLDEGVEPDDVDCWCNTTSTWVVYGTCHHKKGEARRSRRAVTLPSHSTRKLQTRSQTWLESREYTKHLIRVENWIFRNPGFALAAAAIAWLLGSSTSQKVIYLVMILLIAPAYSIRCIGVSNRDFVEGMSGGTWVDVVLEHGGCVTVMAQDKPTVDIELVTTTVSNMAEVRSYCYEASISDMASDSRCPTQGEAYLDKQSDTQYVCKRTLVDRGWGNGCGLFGKGSLVTCAKFACSKKMTGKSIQPENLEYRIMLSVHGSQHSGMIVNDTGHETDENRAKVEITPNSPRAEATLGGFGSLGLDCEPRTGLDFSDLYYLTMNNKHWLVHKEWFHDIPLPWHAGADTGTPHWNNKEALVEFKDAHAKRQTVVVLGSQEGAVHTALAGALEAEMDGAKGRLSSGHLKCRLKMDKLRLKGVSYSLCTAAFTFTKIPAETLHGTVTVEVQYAGTDGPCKVPAQMAVDMQTLTPVGRLITANPVITESTENSKMMLELDPPFGDSYIVIGVGEKKITHHWHRSGSTIGKAFEATVRGAKRMAVLGDTAWDFGSVGGALNSLGKGIHQIFGAAFKSLFGGMSWFSQILIGTLLMWLGLNTKNGSISLMCLALGGVLIFLSTAVSADVGCSVDFSKKETRCGTGVFVYNDVEAWRDRYKYHPDSPRRLAAAVKQAWEDGICGISSVSRMENIMWRSVEGELNAILEENGVQLTVVVGSVKNPMWRGPQRLPVPVNELPHGWKAWGKSYFVRAAKTNNSFVVDGDTLKECPLKHRAWNSFLVEDHGFGVFHTSVWLKVREDYSLECDPAVIGTAVKGKEAVHSDLGYWIESEKNDTWRLKRAHLIEMKTCEWPKSHTLWTDGIEESDLIIPKSLAGPLSHHNTREGYRTQMKGPWHSEELEIRFEECPGTKVHVEETCGTRGPSLRSTTASGRVIEEWCCRECTMPPLSFRAKDGCWYGMEIRPRKEPESNLVRSVVTAGSTDHMDHFSLGVLVILLMVQEGLKKRMTTKIIISTSMAVLVAMILGGFSMSDLAKLAILMGATFAEMNTGGDVAHLALIAAFKVRPALLVSFIFRANWTPRESMLLALASCLLQTAISALEGDLMVLINGFALAWLAIRAMVVPRTDNITLAILAALTPLARGTLLVAWRAGLATCGGFMLLSLKGKGSVKKNLPFVMALGLTAVRLVDPINVVGLLLLTRSGKRSWPPSEVLTAVGLICALAGGFAKADIEMAGPMAAVGLLIVSYVVSGKSVDMYIERAGDITWEKDAEVTGNSPRLDVALDESGDFSLVEDDGPPMREIILKVVLMTICGMNPIAIPFAAGAWYVYVKTGKRSGALWDVPAPKEVKKGETTDGVYRVMTRRLLGSTQVGVGVMQEGVFHTMWHVTKGSALRSGEGRLDPYWGDVKQDLVSYCGPWKLDAAWDGHSEVQLLAVPPGERARNIQTLPGIFKTKDGDIGAVALDYPAGTSGSPILDKCGRVIGLYGNGVVIKNGSYVSAITQGRREEETPVECFEPSMLKKKQLTVLDLHPGAGKTRRVLPEIVREAIKTRLRTVILAPTRVVAAEMEEALRGLPVRYMTTAVNVTHSGTEIVDLMCHATFTSRLLQPIRVPNYNLYIMDEAHFTDPSSIAARGYISTRVEMGEAAAIFMTATPPGTRDAFPDSNSPIMDTEVEVPERAWSSGFDWVTDHSGKTVWFVPSVRNGNEIAACLTKAGKRVIQLSRKTFETEFQKTKHQEWDFVVTTDISEMGANFKADRVIDSRRCLKPVILDGERVILAGPMPVTHASAAQRRGRIGRNPNKPGDEYLYGGGCAETDEDHAHWLEARMLLDNIYLQDGLIASLYRPEADKVAAIEGEFKLRTEQRKTFVELMKRGDLPVWLAYQVASAGITYTDRRWCFDGTTNNTIMEDSVPAEVWTRHGEKRVLKPRWMDARVCSDHAALKSFKEFAAGKRGAAFGVMEALGTLPGHMTERFQEAIDNLAVLMRAETGSRPYKAAAAQLPETLETIMLLGLLGTVSLGIFFVLMRNKGIGKMGFGMVTLGASAWLMWLSEIEPARIACVLIVVFLLLVVLIPEPEKQRSPQDNQMAIIIMVAVGLLGLITANELGWLERTKSDLSHLMGRREEGATIGFSMDIDLRPASAWAIYAALTTFITPAVQHAVTTSYNNYSLMAMATQAGVLFGMGKGMPFYAWDFGVPLLMIGCYSQLTPLTLIVAIILLVAHYMYLIPGLQAAAARAAQKRTAAGIMKNPVVDGIVVTDIDTMTIDPQVEKKMGQVLLIAVAVSSAILSRTAWGWGEAGALITAATSTLWEGSPNKYWNSSTATSLCNIFRGSYLAGASLIYTVTRNAGLVKRRGGGTGETLGEKWKARLNQMSALEFYSYKKSGITEVCREEARRALKDGVATGGHAVSRGSAKLRWLVERGYLQPYGKVIDLGCGRGGWSYYAATIRKVQEVKGYTKGGPGHEEPVLVQSYGWNIVRLKSGVDVFHMAAEPCDTLLCDIGESSSSPEVEEARTLRVLSMVGDWLEKRPGAFCIKVLCPYTSTMMETLERLQRRYGGGLVRVPLSRNSTHEMYWVSGAKSNTIKSVSTTSQLLLGRMDGPRRPVKYEEDVNLGSGTRAVVSCAEAPNMKIIGNRIERIRSEHAETWFFDENHPYRTWAYHGSYEAPTQGSASSLVNGVVRLLSKPWDVVTGVTGIAMTDTTPYGQQRVFKEKVDTRVPDPQEGTRQVMSMVSSWLWKELGKHKRPRVCTKEEFINKVRSNAALGAIFEEEKEWKTAVEAVNDPRFWALVDKEREHHLRGECQSCVYNMMGKREKKQGEFGKAKGSRAIWYMWLGARFLEFEALGFLNEDHWMGRENSGGGVEGLGLQRLGYVLEEMSRIPGGRMYADDTAGWDTRISRFDLENEALITNQMEKGHRALALAIIKYTYQNKVVKVLRPAEKGKTVMDIISRQDQRGSGQVVTYALNTFTNLVVQLIRNMEAEEVLEMQDLWLLRRSEKVTNWLQSNGWDRLKRMAVSGDDCVVKPIDDRFAHALRFLNDMGKVRKDTQEWKPSTGWDNWEEVPFCSHHFNKLHLKDGRSIVVPCRHQDELIGRARVSPGAGWSIRETACLAKSYAQMWQLLYFHRRDLRLMANAICSSVPVDWVPTGRTTWSIHGKGEWMTTEDMLVVWNRVWIEENDHMEDKTPVTKWTDIPYLGKREDLWCGSLIGHRPRTTWAENIKNTVNMVRRIIGEEEKYMDYLSTQVRYLGEEGSTPGXL

>tr|A0A140D2T1|A0A140D2T1_ZIKV Genome polyprotein OSZika virus (strain Mr 766) PE4 SV1

MKNPKKKSGGFRIVNMLKRGVARVNPLGGLKRLPAGLLLGHGPIRMVLAILAFLRFTAIKPSLGLINRWGSVGKKEAMEIIKKFKKDLAAMLRIINARKERKRRGADTSIGIIGLLLTTAMAAEITRRGSAYYMYLDRSDAGKAISFATTLGVNKCHVQIMDLGHMCDATMSYECPMLDEGVEPDDVDCWCNTTSTWVVYGTCHHKKGEARRSRRAVTLPSHSTRKLQTRSQTWLESREYTKHLIKVENWIFRNPGFALVAVAIAWLLGSSTSQKVIYLVMILLIAPAYSIRCIGVSNRDFVEGMSGGTWVDVVLEHGGCVTVMAQDKPTVDIELVTTTVSNMAEVRSYCYEASISDMASDSRCPTQGEAYLDKQSDTQYVCKRTLVDRGWGNGCGLFGKGSLVTCAKFTCSKKMTGKSIQPENLEYRIMLSVHGSQHSGMIVNDTGYETDENRAKVEVTPNSPRAEATLGGFGSLGLDCEPRTGLDFSDLYYLTMNNKHWLVHKEWFHDIPLPWHAGADTGTPHWNNKEALVEFKDAHAKRQTVVVLGSQEGAVHTALAGALEAEMDGAKGKLFSGHLKCRLKMDKLRLKGVSYSLCTAAFTFTKVPAETLHGTVTVEVQYAGTDGPCKIPVQMAVDMQTLTPVGRLITANPVITESTENSKMMLELDPPFGDSYIVIGVGDKKITHHWHRSGSTIGKAFEATVRGAKRMAVLGDTAWDFGSVGGVFNSLGKGIHQIFGAAFKSLFGGMSWFSQILIGTLLVWLGLNTKNGSISLTCLALGGVMIFLSTAVSADVGCSVDFSKKETRCGTGVFIYNDVEAWRDRYKYHPDSPRRLAAAVKQAWEEGICGISSVSRMENIMWKSVEGELNAILEENGVQLTVVVGSVKNPMWRGPQRLPVPVNELPHGWKAWGKSYFVRAAKTNNSFVVDGDTLKECPLEHRAWNSFLVEDHGFGVFHTSVWLKVREDYSLECDPAVIGTAVKGREAAHSDLGYWIESEKNDTWRLKRAHLIEMKTCEWPKSHTLWTDGVEESDLIIPKSLAGPLSHHNTREGYRTQVKGPWHSEELEIRFEECPGTKVYVEETCGTRGPSLRSTTASGRVIEEWCCRECTMPPLSFRAKDGCWYGMEIRPRKEPESNLVRSMVTAGSTDHMDHFSLGVLVILLMVQEGLKKRMTTKIIMSTSMAVLVVMILGGFSMSDLAKLVILMGATFAEMNTGGDVAHLALVAAFKVRPALLVSFIFRANWTPRESMLLALASCLLQTAISALEGDLMVLINGFALAWLAIRAMAVPRTDNIALPILAALTPLARGTLLVAWRAGLATCGGIMLLSLKGKGSVKKNLPFVMALGLTAVRVVDPINVVGLLLLTRSGKRSWPPSEVLTAVGLICALAGGFAKADIEMAGPMAAVGLLIVSYVVSGKSVDMYIERAGDITWEKDAEVTGNSPRLDVALDESGDFSLVEEDGPPMREIILKVVLMAICGMNPIAIPFAAGAWYVYVKTGKRSGALWDVPAPKEVKKGETTDGVYRVMTRRLLGSTQVGVGVMQEGVFHTMWHVTKGAALRSGEGRLDPYWGDVKQDLVSYCGPWKLDAAWDGLSEVQLLAVPPGERARNIQTLPGIFKTKDGDIGAVALDYPAGTSGSPILDKCGRVIGLYGNGVVIKNGSYVSAITQGKREEETPVECFEPSMLKKKQLTVLDLHPGAGKTRRVLPEIVREAIKKRLRTVILAPTRVVAAEMEEALRGLPVRYMTTAVNVTHSGTEIVDLMCHATFTSRLLQPIRVPNYNLYIMDEAHFTDPSSIAARGYISTRVEMGEAAAIFMTATPPGTRDAFPDSNSPIMDTEVEVPERAWSSGFDWVTDHSGKTVWFVPSVRNGNEIAACLTKAGKRVIQLSRKTFETEFQKTKNQEWDFVITTDISEMGANFKADRVIDSRRCLKPVILDGERVILAGPMPVTHASAAQRRGRIGRNPNKPGDEYMYGGGCAETDEGHAHWLEARMLLDNIYLQDGLIASLYRPEADKVAAIEGEFKLRTEQRKTFVELMKRGDLPVWLAYQVASAGITYTDRRWCFDGTTNNTIMEDSVPAEVWTKYGEKRVLKPRWMDARVCSDHAALKSFKEFAAGKRGAALGVMEALGTLPGHMTERFQEAIDNLAVLMRAETGSRPYKAAAAQLPETLETIMLLGLLGTVSLGIFFVLMRNKGIGKMGFGMVTLGASAWLMWLSEIEPARIACVLIVVFLLLVVLIPEPEKQRSPQDNQMAIIIMVAVGLLGLITANELGWLERTKNDIAHLMGRREEGATMGFSMDIDLRPASAWAIYAALTTLITPAVQHAVTTSYNNYSLMAMATQAGVLFGMGKGMPFYAWDLGVPLLMMGCYSQLTPLTLIVAIILLVAHYMYLIPGLQAAAARAAQKRTAAGIMKNPVVDGIVVTDIDTMTIDPQVEKKMGQVLLIAVAISSAVLLRTAWGWGEAGALITAATSTLWEGSPNKYWNSSTATSLCNIFRGSYLAGASLIYTVTRNAGLVKRRGGGTGETLGEKWKARLNQMSALEFYSYKKSGITEVCREEARRALKDGVATGGHAVSRGSAKLRWLVERGYLQPYGKVVDLGCGRGGWSYYAATIRKVQEVRGYTKGGPGHEEPMLVQSYGWNIVRLKSGVDVFHMAAEPCDTLLCDIGESSSSPEVEETRTLRVLSMVGDWLEKRPGAFCIKVLCPYTSTMMETMERLQRRHGGGLVRVPLSRNSTHEMYWVSGAKSNIIKSVSTTSQLLLGRMDGPRRPVKYEEDVNLGSGTRAVASCAEAPNMKIIGRRIERIRNEHAETWFLDENHPYRTWAYHGSYEAPTQGSASSLVNGVVRLLSKPWDVVTGVTGIAMTDTTPYGQQRVFKEKVDTRVPDPQEGTRQVMNIVSSWLWKELGKRKRPRVCTKEEFINKVRSNAALGAIFEEEKEWKTAVEAVNDPRFWALVDREREHHLRGECHSCVYNMMGKREKKQGEFGKAKGSRAIWYMWLGARFLEFEALGFLNEDHWMGRENSGGGVEGLGLQRLGYILEEMNRAPGGKMYADDTAGWDTRISKFDLENEALITNQMEEGHRTLALAVIKYTYQNKVVKVLRPAEGGKTVMDIISRQDQRGSGQVVTYALNTFTNLVVQLIRNMEAEEVLEMQDLWLLRKPEKVTRWLQSNGWDRLKRMAVSGDDCVVKPIDDRFAHALRFLNDMGKVRKDTQEWKPSTGWSNWEEVPFCSHHFNKLYLKDGRSIVVPCRHQDELIGRARVSPGAGWSIRETACLAKSYAQMWQLLYFHRRDLRLMANAICSAVPVDWVPTGRTTWSIHGKGEWMTTEDMLMVWNRVWIEENDHMEDKTPVTKWTDIPYLGKREDLWCGSLIGHRPRTTWAENIKDTVNMVRRIIGDEEKYMDYLSTQVRYLGEEGSTPGVL

>tr|A0A140DLX4|A0A140DLX4_ZIKV Genome polyprotein OSZika virus (strain Mr 766) PE4 SV1

MKNPKKKSGGFRIVNMLKRGVARVSPFGGLKRLPAGLLLGHGPIRMVLAILAFLRFTAIKPSLGLINRWGSVGKKEAMEIIKKFKKDLAAMLRIINARKEKKRRGADTSVGIVGLLLTTAMAAEVTRRGSAYYMYLDRNDAGEAISFPTTLGMNKCYIQIMDLGHMCDATMSYECPMLDEGVEPDDVDCWCNTTSTWVVYGTCHHKKGEARRSRRAVTLPSHSTRKLQTRSQTWLESREYTKHLIRVENWIFRNPGFALAAAAIAWLLGSSTSQKVIYLVMILLIAPAYSIRCIGVSNRDFVEGMSGGTWVDVVLEHGGCVTVMAQDKPTVDIELVSTTVSNMAEVRSYCYEATISDIASDSRCPTQGEAYLDKQSDTQYVCKRTLVDRGWGNGCGLFGKGSLVTCAKFACSKKMTGKSIQPENLEYRIMLSVHGSQHSGMIVNDTGHETDENRAKVEITPNSPRAEATLGGFGSLGLDCEPRTGLDFSDLYYLTMNNKHWLVHKEWFHDIPLPWHAGADTGTPHWNNKEALVEFKDAHAKRQTAVVLGSQEGAVHTALAGALEAEMDGAKGRLSSGHLKCRLKMDKLRLKGVSYSLCTAAFTFTKIPAETLHGTVTVEVQYAGTDGPCKVPAQMAVDMQTLTPVGRLITANPVITESTENSKMMLELDPPFGDSYIVIGVGEKKITHHWHRSGSTIGKAFEATVRGAKRMAVLGDTAWDFGSVGGALNSLGKGIHQIFGAAFKSLFGGMSWFSQILIGTLLMWLGLNTKNGSISLMCLALGGVLIFLSTAVSADVGCSVDFSKKETRCGTGVFVYNDVEAWRDRYKYHPDSPRRLAAAVKQAWEDGICGISSVSRMENIMWRSVEGELNAILEENGVQLTVVVGSVKNPMWRGPQRLPVPVNELPHGWKAWGKSYFVRAAKTNNSFVVDGDTLKECPLKHRAWNSFLVEDHGFGVFHTSVWLKVREDYSLECDPAVIGTAVKEKEAVHSDLGYWIESEKNDTWRLKRAHLIEMKTCEWPKSHTLWTDGIEESDLIIPKSLAGPLSHHNTREGYRTQMKGPWHSEELEIRFEECPGTKVHVEETCGTRGPSLRSTTASGRVIEEWCCRECTMPPLSFRAKDGCWYGMEIRPRKEPESNLVRSVVTAGSTDHMDHFSLGVLVILLMVQEGLKKRMTTKIIISTSMAVLVAMILGGFSMSDLAKLAILMGATFAEMNTGGDVAHLALIAAFKVRPALLVSFIFRANWTPRESMLLALASCLLQTAISALEGDLMVLINGFALAWLAIRAMVVPRTDNITLAILAALTPLARGTLLVAWRAGLATCGGFMLLSLKGKGSVKKNLPFVMALGLTAVRLVDPINVVGLLLLTRSGKRSWPPSEVLTAVGLICALAGGFAKADIEMAGPIAAVGLLIVSYVVSGKSVDMYIERAGDITWEKDAEVTGNSPRLDVALDESGDFSLVEDDGPPMREIILKVVLMTICGMNPIAIPFAAGAWYVYVKTGKRSGALWDVPAPKEVKKGETTDGVYRVMTRRLLGSTQVGVGVMQEGVFHTMWHVTKGSALRSGEGRLDPYWGDVKQDLVSYCGPWKLDAAWDGHSEVQLLAVPPGERARNIQTLPGIFKTKDGDIGAVALDYPAGTSGSPILDKCGRVIGLYGNGVVIKNGSYVSAITQGRREEETPVECFEPSMLKKKQLTVLDLHPGAGKTRRVLPEIVREAIKTRLRTVILAPTRVVAAEMEEALRGLPVRYMTTAVNVTHSGTEIVDLMCHATFTSRLLQPIRVPNYNLYIMDEAHFTDPSSIAARGYISTRVEMGEAAAIFMTATPPGTRDAFPDSNSPIMDTEVEVPERAWSSGFDWVTDHSGKTVWFVPSVRNGNEIAACLTKAGKRVIQLSRKTFETEFQKTKHQEWDFVVTTDISEMGANFKADRVIDSRRCLKPVILDGERVILAGPMPVTHASAAQRRGRIGRNPNKPGDEYLYGGGCAETDEDHAHWLEARMLLDNIYLQDGLIASLYRPEADKVAAIEGEFKLRTEQRKTFVELMKRGDLPVWLAYQVASAGITYTDRRWCFDGTTNNTIMEDSVPAEVWTRHGEKRVLKPRWMDARVCSDHAALKSFKEFAAGKRGAAFGVMEALGTLPGHMTERFQEAIDNLAVLMRAETGSRPYKAAAAQLPETLETIMLLGLLGTVSLGIFFVLMRNKGIGKMGFGMVTLGASAWLMWLSEIEPARIACVLIVVFLLLVVLIPEPEKQRSPQDNQMAIIIMVAVGLLGLITANELGWLERTKSDLSHLMGRREEGATIGFSMDIDLRPASAWAIYAALTTFITPAVQHAVTTSYNNYSLMAMATQAGVLFGMGKGMPFYAWDFGVPLLMIGCYSQLTPLTLIVAIILLVAHYMYLIPGLQAAAARAAQKRTAAGIMKNPVVDGIVVTDIDTMTIDPQVEKKMGQVLLIAVAVSSAILSRTAWGWGEAGALITAATSTLWEGSPNKYWNSSTATSLCNIFRGSYLAGASLIYTVTRNAGLVKRRGGGTGETLGEKWKARLNQMSALEFYSYKKSGITEVCREEARRALKDGVATGGHAVSRGSAKLRWLVERGYLQPYGKVIDLGCGRGGWSYYAATIRKVQEVKGYTKGGPGHEEPVLVQSYGWNIVRLKSGVDVFHMAAEPCDTLLCDIGESSSSPEVEEARTLRVLSMVGDWLEKRPGAFCIKVLCPYTSTMMETLERLQRRYGGGLVRVPLSRNSTHEMYWVSGAKSNTIKSVSTTSQLLLGRMDGPRRPVKYEEDVNLGSGTRAVVSCAEAPNMKIIGDRIERIRSEHAETWFFDENHPYRTWAYHGSYEAPTQGSASSLINGVVRLLSKPWDVVTGVTGIAMTDTTPYGQQRVFKEKVDTRVPDPQEGTRQVMSMVSSWLWKELGKHKRPRVCTKEEFINKVRSNAALGAIFEEEKEWKTAVEAVNDPRFWALVDKEREHHLRGECQSCVYNMMGKREKKQGEFGKAKGSRAIWYMWLGARFLEFEALGFLNEDHWMGRENSGGGVEGLGLQRLGYVLEEMSRIPGGRMYADDTAGWDTRISRFDLENEALITNQMEKGHRALALAIIKYTYQNKVVKVLRPAEKGKTVMDIISRQDQRGSGQVVTYALNTFTNLVVQLIRNMEAEEVLEMQDLWLLRRSEKVTNWLQSNGWDRLKRMAVSGDDCVVKPIDDRFAHALRFLNDMGKVRKDTQEWKPSTGWDNWEEVPFCSHHFNKLHLKDGRSIVVPCRHQDELIGRARVSPGAGWSIRETACLAKSYAQMWQLLYFHRRDLRLMANAICSSVPVDWVPTGRTTWSIHGKGEWMTTEDMLVVWNRVWIEENDHMEDKTPVTKWTDIPYLGKREDLWCGSLIGHRPRTTWAENIKNTVNMVRRIIGDEEKYMDYLSTQVRYLGEEGSTPGVL

>tr|A0A140E7U5|A0A140E7U5_ZIKV Genome polyprotein OSZika virus (strain Mr 766) PE4 SV1

MKNPKKKSGGFRIVNMLKRGVARVSPFGGLKRLPAGLLLGHGPIRMVLAILAFLRFTAIKPSLGLINRWGSVGKKDAMEIIKKFKKDLAAMLRIINARKEKKRRGADTSVGIVGLLLTTAMAAEVTRRGSAYYMYLDRNDAGEAISFPTTLGMNKCYIQIMDLGHMCDATMSYECPMLDEGVEPDDVDCWCNTTSTWVVYGTCHHKKGEARRSRRAVTLPSHSTRKLQTRSQTWLESREYTKHLIRVENWIFRNPGFALAAAAIAWLLGSSTSQKVIYLVMILLIAPAYSIRCIGVSNRDFVEGMSGGTWVDVVLEHGGCVTAMAQDKPTVDIELVTTTVSNMAEVRSYCYEASISDMASDSRCPTQGEAYLDKQSDTQYVCKRTLVDRGWGNGCGLFGKGSLVTCAKFACSKKMTGKSIQPENLEYRIMLSVHGSQHSGMLVNDTGHETDENRAKVEITPNSPRAEATLGGFGSLGLDCEPRTGLDFSDLYYLTMNNKHWLAHKEWFHDIPLPWHAGAATGTPHWNNKEALVEFKDAHAKRQTVVVLGSQEGAVHTALAGALEAEMDGAKGRLSSGHLKCRLKMDKLRLKGVSYSLCTAAFTFTKIPAETVDGTVTVEGQYGGTDGPCKVPAQMAVDMQTLTPVGRLITANPVITESTENSKMMLELDPPFGDSYIVIGVGEKKITHHWHRSGSTIGKAFEATVRGAKRMAVLGDTAWDFGSVGGALNSLGKGIHQIIGAAFKSLFGGMSWFSQILIGTLLMWLGLNTKNGSISLMCLALGGVLIFLSTAVSGGVGCSVDFSKKETRCGTGVFVYNDVEAWRDRYKYHPDSPRRLAAAVKQAWEDGICGISSVSRMENIMWRSVEGELNAILEENGVQLTVVVGSVKNPMWRGPQRLPVPVNELPHGWKAWGKSYFVRAAKTNNSFVVDGDTLKECPLKHRAWNSFLVEDHGFGVFHTSVWLKVREDYWLECDPAVIGTAVKGKEAVHSDLGYWIESEKNDTWWLKRAHLIEMKTCEWPKSHTLWTDGIEESDLIIPKSLAGPLSHHNAREGYRTQMKGPWHSEELEIRFEECPGTKVHVEETCGTRGPSLRSTTASGRVIEEWCSRECTMPPLSFQAKDGCWYGMEIRPRKEPESNLVRSMVTAGSTDHMDHFSLGVLVILLMVQEGLKKRMTTKIIISTSMAVLVAMILGGFSMSDLAKLAILMGATFAEMNTGGDVAHLALIAAFKVRPALLVSFIFRANWTPRESMLLALASCLLQTAISALEGDLMVLINGFALAWLAIRAMVVPRTDNITLAILAALTPLARGTLLVAWRAGLATCGGFMLLSLKGKGSVKKNLPFVMALGLTAVRLVDPINVVGLLLLTRSGKRSWPPSEVLTAVGLICALAGGFAKADIEMAGPMAAVGLLIVSYVVSGKSVDMYIERAGDITWEKDAEVTGNSPRLDVALDESGDFSLVEDDGPPMREIILKVVLMTICGMNPIAIPFAAGAWYVYVKTGKRSGALWDVPAPKEVKKGETTDGVYRVMTRRLLGSTQVGVGVMQEGVFHTMWHVTKGSALRSGEGRLDPYWGDVKQDLVSYCGPWKLDAAWDGHSEVQLLAVPPGERARNIQTLPGIFKTKDGDIGAVALDYPAGTSGSPILDKCGRVIGLYGNGVVIKNGSYVSAITQGRREEETPVECFEPSMLKKKQLTVLDLHPGAGKTRRVLPEIVREAIKTRLRTVILAPTRVVAAEMEEALRGLPVRYMTTAVNVTHSGTEIVDLMCHATFTSRLLQPIRVPNYNLYIMDEAHFTDPSSIAARGYISTRVEMGEAAAIFMTATPPGTRDAFPDSNSPIMDTEVEVPERAWSSGFDWVTEYSGKTVWFVPRVRNGNEIAACLTKAGKRVIQLSRKTFETEFQKTKHQEWDFVVTTDISEMGANFKADRVIDSRRCLKPVILGGERVILAGPMPVTHASAAQRRGRIGRNPNKPGDEYLYGGGCAETDEDHAHWLEARMLLDNIYLQDGLIASLYRPEADKVAAIEGEFKLRTEQRKTFVELMKRGDLPVWLAYQVASAGITYTDRRWCFDGTTNNTIMEDSVPAEVWTRHGEKRVLKPRWMDARVCSDHAALKSFKEFAAGKRGAAFGVMEALGTLPGHMTERFQEAIDNLAVLMRAETGSRPYKAAAAQLPETLETIMLLGLLGTVSLGIFFVLMRNKGIGKMGFGMVTLGASAWLMWLSEIEPARIACVLIVVFLLLVVLIPEPEKQRSPQDNQMAIIIMVAVGLLGLITANELGWLERTKSDLSHLMGRREEGATMGFSMDIDLRPASAWAIYPALTSFITPAVQHAVTTSYNNYSLMAMATQAGVLFGMGKGMPFYAWDFGVPLLMIGCYSQLTPLTLIVAIILLVAHYMYLIPGLQAAAARAAQKRTAAGIMKNPVVEGIVVTDIDTMTIDPQVEKKMGQVLLMAVAVSSAILSRTAWGWGEAGALITAATSTLWEGSPNKYWNSSTATSLCNIFRGSYLAGASLIYTVTRNAGLVKRRGGGTGETLGEKWKARLNQMSALEFYSYKKSGITEVCREEARRALKDGVATGGHAVSRGSAKLRWLVERGYLQPYGKVIDLGCGRGGWSYYAATIRKVQEVKGYTKGGPGHEEPVLVQSYGWNIVRLKSGVDVFHMAAEPCDTLLCDIGESSSSPEVEEARTLRVLSMVGDWLEKRPGAFCIKVLCPYTSTMMETLERLQRRYGGGLVRVPLSRNSTHEMYWVSGAKSNTIKSVSTTSQLLLGRMDGPRRPVKYEEDVNLGSGTRAVVSCAEAPNMKIIGNRIERIRAEKAETWFFDENHPYRTWAYHGSYDAATQGSASSLINGVVRLLSKPWDVVTGVTGIAMTDTTPYGQQRVFKEKVDTRVPDPQEGTRQVMSMVSSWLWKELGKHKRPRVCTKEEFINKVRSNAALGAIFEEEKEWKTAVEAVNDPRFWALVDKEREHHLRGECQSCVYITMGKREKKQGEFGKAKGSRAIWYMWLGARFLEFEALGFLNEDHWMGRENSGGGVEGLGLQRLGYVLEEMSRIPGGRMYADDTAGWDTRISRFDLENEALITNQMEKGHRALALAIIKYTYQNKVVKVLRPAEKGKTVMDIISRQDQRGSGQVVTYALNTFTNLVVQLIRNMEAEEVLEMQDLWLLRRSEKVTNWLQSNGWDRLKRMAVSGDDCVVKPIDDRFAHALRFLNDMGKVRKDTQEWKPSTGWDNWEEVPFCSHHFNKLHLKDGRSIVVPCRHQDELIGRARVSPGAGWSIRETACLAKSYAQMWQLLYFHRRDLRLMANAICSSVPVDWVPTGRTTWSIHGKGEWMTTEDMLVAWNRVWIEENDHMEDKTPVTKWTDIPYLGKREDLWCGSLIGHRPRTTWAENIKNTVNMVRRIIGDEEKYMDYLSTQVRYLGEEGSTPGVL

>tr|A0A127AM58|A0A127AM58_ZIKV Genome polyprotein OSZika virus (strain Mr 766) PE4 SV1

MKNPKKKSGGFRIVNMLKRGVARVSPFGGLKRLPAGLLLGHGPIRMVLAILAFLRFTAIKPSLGLINRWGSVGKKEAMEIIKKFKKDLAAMLRIINARKEKKRRGAETSVGIVGLLLTTAMAAEVTRRGSAYYMYLDRNDAGEAISFPTTLGMNKCYIQIMDLGHMCDATMSYECPMLDEGVEPDDVDCWCNTTSTWVVYGTCHHKKGEARRSRRAVTLPSHSTRKLQTRSQTWLESREYTKHLIRVENWIFRNPGFALAAAAIAWLLGSSTSQKVIYLVMILLIAPAYSIRCIGVSNRDFVEGMSGGTWVDVVLEHGGCVTVMAQDKPTVDIELVTTTVSNMAEVRSYCYEASISDMASDSRCPTQGEAYLDKQSDTQYVCKRTLVDRGWGNGCGLFGKGSLVTCAKFACSKKMTGKSIQPENLEYRIMLSVHGSQHSGMIVNDTGHETDENRAKVEITPNSPRAEATLGGFGSLGLDCEPRTGLDFSDLYYLTMNNKHWLVHKEWFHDIPLPWHAGADTGTPHWNNKEALVEFKDAHAKRQTVVVLGSQEGAVHTALAGALEAEMDGAKGRLSSGHLKCRLKMDKLRLKGVSYSLCTAAFTFTKIPAETLHGTVTVEVQYAGTDGPCKVPAQMAVDMQTLTPVGRLITANPVITESTENSKMMLELDPPFGDSYIVIGVGEKKITHHWHRSGSTIGKAFEATVRGAKRMAVLGDTAWDFGSVGGALNSLGKGIHQIFGAAFKSLFGGMSWFSQILIGTLLMWLGLNTKNGSISLMCLALGGVLIFLSTAVSADVGCSVDFSKKETRCGTGVFVYNDVEAWRDRYKYHPDSPRRLAAAVKQAWEDGICGISSVSRMENIMWRSVEGELNAILEENGVQLTVVVGSVKNPMWRGPQRLPVPVNELPHGWKAWGKSYFVRAAKTNNSFVVDGDTLKECPLKHRAWNSFLVEDHGFGVFHTSVWLKVREDYSLECDPAVIGTAVKGKEAVHSDLGYWIESEKNDTWRLKRAHLIEMKTCEWPKSHTLWTDGIEESDLIIPKSLAGPLSHHNTREGYRTQMKGPWHSEELEIRFEECPGTKVHVEETCGTRGPSLRSTTASGRVIEEWCCRECTMPPLSFWAKDGCWYGMEIRPRKEPESNLVRSMVTAGSTDHMDHFSLGVLVILLMVQEGLKKRMTTKIIISTSMAVLVAMILGGFSMSDLAKLAILMGATFAEMNTGGDVAHLALIAAFKVRPALLVSFIFRANWTPRESMLLALASCLLQTAISALEGDLMVLINGFALAWLAIRAMVVPRTDNITLAILAALTPLARGTLLVAWRAGLATCGGFMLLSLKGKGSVKKNLPFVMALGLTAVRLVDPINVVGLLLLTRSGKRSWPPSEVLTAVGLICALAGGFAKADIEMAGPMAAVGLLIVSYVVSGKSVDMYIERAGDITWEKDAEVTGNSPRLDVALDESGDFSLVEDDGPPMREIILKVVLMTICGMNPIAIPFAAGAWYVYVKTGKRSGALWDVPAPKEVKKGETTDGVYRVMTRRLLGSTQVGVGVMQEGVFHTMWHVTKGSALRSGEGRLDPYWGDVKQDLVSYCGPWKLDAAWDGHSEVQLLAVPPGERARNIQTLPGIFKTKDGDIGAVALDYPAGTSGSPILDKCGRVIGLYGNGVVIKNGSYVSAITQGRREEETPVECFEPSMLKKKQLTVLDLHPGAGKTRRVLPEIVREAIKTRLRTVILAPTRVVAAEMEEALRGLPVRYMTTAVNVTHSGTEIVDLMCHATFTSRLLQPIRVPNYNLYIMDEAHFTDPSSIAARGYISTRVEMGEAAAIFMTATPPGTRDAFPDSNSPIMDTEVEVPERAWSSGFDWVTDHSGKTVWFVPSVRNGNEIAACLTKAGKRVIQLSRKTFETEFQKTKHQEWDFVVTTDISEMGANFKADRVIDSRRCLKPVILDGERVILAGPMPVTHASAAQRRGRIGRNPNKPGDEYLYGGGCAETDEDHAHWLEARMLLDNIYLQDGLIASLYRPEADKVAAIEGEFKLRTEQRKTFVELMKRGDLPVWLAYQVASAGITYTDRRWCFDGTTNNTIMEDSVPAEVWTRHGEKRVLKPRWMDARVCSDHAALKSFKEFAAGKRGAAFGVMEALGTLPGHMTERFQEAIDNLAVLMRAETGSRPYKAAAAQLPETLETIMLLGLLGTVSLGIFFVLMRNKGIGKMGFGMVTLGASAWLMWLSEIEPARIACVLIVVFLLLVVLIPEPEKQRSPQDNQMAIIIMVAVGLLGLITANELGWLERTKSDLSHLMGRREEGATIGFSMDIDLRPASAWAIYAALTTFITPAVQHAVTTSYNNYSLMAMATQAGVLFGMGKGMPFYAWDFGVPLLMIGCYSQLTPLTLIVAIILLVAHYMYLIPGLQAAAARAAQKRTAAGIMKNPVVDGIVVTDIDTMTIDPQVEKKMGQVLLIAVAVSSAILSRTAWGWGEAGALITAATSTLWEGSPNKYWNSSTATSLCNIFRGSYLAGASLIYTVTRNAGLVKRRGGGTGETLGEKWKARLNQMSALEFYSYKKSGITEVCREEARRALKDGVATGGHAVSRGSAKLRWLVERGYLQPYGKVIDLGCGRGGWSYYAATIRKVQEVKGYTKGGPGHEEPVLVQSYGWNIVRLKSGVDVFHMAAEPCDTLLCDIGESSSSPEVEEARTLRVLSMVGDWLEKRPGAFCIKVLCPYTSTMMETLERLQRRYGGGLVRVPLSRNSTHEMYWVSGAKSNTIKSVSTTSQLLLGRMDGPRRPVKYEEDVNLGSGTRAVVSCAEAPNMKIIGNRIERIRSEHAETWFFDENHPYRTWAYHGSYEAPTQGSASSLINGVVRLLSKPWDVVTGVTGIAMTDTTPYGQQRVFKEKVDTRVPDPQEGTRQVMSMVSSWLWKELGKHKRPRVCTKEEFINKVRSNAALGAIFEEEKEWKTAVEAVNDPRFWALVDKEREHHLRGECQSCVYNMMGKREKKQGEFGKAKGSRAIWYMWLGARFLEFEALGFLNEDHWMGRENSGGGVEGLGLQRLGYVLEEMSRIPGGRMYADDTAGWDTRISRFDLENEALITNQMEKGHRALALAIIKYTYQNKVVKVLRPAEKGKTVMDIISRQDQRGSGQVVTYALNTFTNLVVQLIRNMEAEEVLEMQDLWLLRRSEKVTNWLQSNGWDRLKRMAVSGDDCVVKPIDDRFAHALRFLNDMGKVRKDTQEWKPSTGWDNWEEVPFCSHHFNKLHLKDGRSIVVPCRHQDELIGRARVSPGAGWSIRETACLAKSYAQMWQLLYFHRRDLRLMANAICSSVPVDWVPTGRTTWSIHGKGEWMTTEDMLVVWNRVWIEENDHMEDKTPVAKWTDIPYLGKREDLWCGSLIGHRPRTTWAENIKNTVNMVRRIIGDEEKYMDYLSTQVRYLGEEGSTPGVL

>tr|A0A146CJG7|A0A146CJG7_ZIKV Polyprotein OSZika virus (strain Mr 766) PE4 SV1

MKNPKKKSGGFRIVNMLKRGVARVSPFGGLKRLPAGLLLGHGPIRMVLAILAFLRFTAIKPSLGLINRWGSVGKKEAMEIIKKFKKDLAAMLRIINARKEKKRRGADTSVGIVGLLLTTAMAAEVTRRGSAYYMYLDRNDAGEAISFPTTLGMNKCYIQIMDLGHMCDATMSYECPMLDEGVEPDDVDCWCNTTSTWVVYGTCHHKKGEARRSRRAVTLPSHSTRKLQTRSQTWLESREYTKHLIRVENWIFRNPGFALAAAAIAWLLGSSTSQKVIYLVMILLIAPAYSIRCIGVSNRDFVEGMSGGTWVDVVLEHGGCVTVMAQDKPTVDIELVTTTVSNMAEVRSYCYEASISDMASDSRCPTQGEAYLDKQSDTQYVCKRTLVDRGWGNGCGLFGKGSLVTCAKFACSKKMTGKSIQPENLEYRIMLSVHGSQHSGMIVNDTGHETDENRAKVEITPNSPRAEATLGGFGSLGLDCEPRTGLDFSDLYYLTMNNKHWLVHKEWFHDIPLPWHAGADTGTPHWNNKEALVEFKDAHAKRQTVVVLGSQEGAVHTALAGALEAEMDGAKGRLSSGHLKCRLKMDKLRLKGVSYSLCTAAFTFTKIPAETLHGTVTVEVQYAGTDGPCKVPAQMAVDMQTLTPVGRLITANPVITESTENSKMMLELDPPFGDSYIVIGVGEKKITHHWHRSGSTIGKAFEATVRGAKRMAVLGDTAWDFGSVGGALNSLGKGIHQIFGAAFKSLFGGMSWFSQILIGTLLMWLGLNTKNGSISLMCLALGGVLIFLSTAVSADVGCSVDFSKKETRCGTGVFVYNDVEAWRDRYKYHPDSPRRLAAAVKQAWEDGICGISSVSRMENIMWRSVEGELNAILEENGVQLTVVVGSVKNPMWRGPQRLPVPVNELPHGWKAWGKSYFVRAAKTNNSFVVDGDTLKECPLKHRAWNSFLVEDHGFGVFHTSVWLKVREDYSLECDPAVIGTAVKGKEAVHSDLGYWIESEKNDTWRLKRAHLIEMKTCEWPKSHTLWTDGIEESDLIIPKSLAGPLSHHNTREGYRTQMKGPWHSEELEIRFEECPGTKVHVEETCGTRGPSLRSTTASGRVIEEWCCRECTMPPLSFRAKDGCWYGMEIRPRKEPESNLVRSMVTAGSTDHMDHFSLGVLVILLMVQEGLKKRMTTKIIISTSMAVLVAMILGGFSMSDLAKLAILMGATFAEMNTGGDVAHLALIAAFKVRPALLVSFIFRANWTPRESMLLALASCLLQTAISALEGDLMVLINGFALAWLAVRAMVVPRTDNITLAILAALTPLARGTLLVAWRAGLATCGGFMLLSLKGKGSVKKNLPFVMALGLTAVRLVDPINVVGLLLLTRSGKRSWPPSEVLTAVGLICALAGGFAKADIEMAGPMAAVGLLIVSYVVSGKSVDMYIERAGDITWEKDAEVTGNSPRLDVALDESGDFSLVEDDGPPMREIILKVVLMTICGMNPIAIPFAAGAWYVYVKTGKRSGALWDVPAPKEVKKGETTDGVYRVMTRRLLGSTQVGVGVMQEGVFHTMWHVTKGSALRSGEGRLDPYWGDVKQDLVSYCGPWKLDAAWDGHSEVQLLAVPPGERARNIQTLPGIFKTKDGDIGAVALDYPAGTSGSPILDKCGRVIGLYGNGVVIKNGSYVSAITQGRREEETPVECFEPSMLKKKQLTVLDLHPGAGKTRRVLPEIVREAIKTRLRTVILAPTRVVAAEMEEALRGLPVRYMTTAVNVTHSGTEIVDLMCHATFTSRLLQPIRVPNYNLYIMDEAHFTDPSSIAARGYISTRVEMGEAAAIFMTATPPGTRDAFPDSNSPIMDTEVEVPERAWSSGFDWVTDHSGKTVWFVPSVRNGNEIAACLTKAGKRVIQLSRKTFETEFQKTKHQEWDFVVTTDISEMGANFKADRVIDSRRCLKPVILDGERVILAGPMPVTHASAAQRRGRIGRNPNKPGDEYLYGGGCAETDEDHAHWLEARMLLDNIYLQDGLIASLYRPEADKVAAIEGEFKLRTEQRKTFVELMKRGDLPVWLAYQVASAGITYTDRRWCFDGTTNNTIMEDSVPAEVWTRHGEKRVLKPRWMDARVCSDHAALKSFKEFAAGKRGAAFGVMEALGTLPGHMTERFQEAIDNLAVLMRAETGSRPYKAAAAQLPETLETIMLLGLLGTVSLGIFFVLMRNKGIGKMGFGMVTLGASAWLMWLSEIEPARIACVLIVVFLLLVVLIPEPEKQRSPQDNQMAIIIMVAVGLLGLITANELGWLERTKSDLSHLMGRREEGATIGFSMDIDLRPASAWAIYAALTTFITPAVQHAVTTSYNNYSLMAMATQAGVLFGMGKGMPFYAWDFGVPLLMIGCYSQLTPLTLIVAIILLVAHYMYLIPGLQAAAARAAQKRTAAGIMKNPVVDGIVVTDIDTMTIDPQVEKKMGQVLLIAVAVSSAILSRTAWGWGEAGALITAATSTLWEGSPNKYWNSSTATSLCNIFRGSYLAGASLIYTVTRNAGLVKRRGGGTGETLGEKWKARLNQMSALEFYSYKKSGITEVCREEARRALKDGVATGGHAVSRGSAKLRWLVDRGYLQPYGKVIDLGCGRGGWSYYAATIRKVQEVKGYTKGGPGHEEPVLVQSYGWNIVRLKSGVDVFHMAAEPCDTLLCDIGESSSSPEVEEARTLRVLSMVGDWLEKRPGAFCIKVLCPYTSTMMETLERLQRRYGGGLVRVPLSRNSTHEMYWVSGAKSNTIKSVSTTSQLLLGRMDGPRRPVKYEEDVNLGSGTRAVASCAEAPNMKIIGNRIERIRSEHAETWFFDENHPYRTWAYHGSYEAPTQGSASSLINGVVRLLSKPWDVVTGVTGIAMTDTTPYGQQRVFKEKVDTRVPDPQEGTRQVMSMVSSWLWKELGKHKRPRVCTKEEFINKVRSNAALGAIFEEEKEWKTAVEAVNDPRFWALVDKEREHHLRGECQSCVYNMMGKREKKQGEFGKAKGSRAIWYMWLGARFLEFEALGFLNEDHWMGRENSGGGVEGLGLQRLGYVLEEMSRIPGGRMYADDTAGWDTRISRFDLENEALITNQMEKGHRALALAIIKYTYQNKVVKVLRPAEKGKTVMDIISRQDQRGSGQVVTYALNTFTNLVVQLIRNMEAEEVLEMQDLWLLRRSEKVTNWLQSNGWDRLKRMAVSGDDCVVKPIDDRFAHALRFLNDMGKVRKDTQEWKPSTGWDNWEEVPFCSHHFNKLHLKDGRSIVVPCRHQDELIGRARVSPGAGWSIRETACLAKSYAQMWQLLYFHRRDLRLMANAICSSVPVDWVPTGRTTWSIHGKGEWMTTEDMLVVWNRVWIEENDHMEDKTPVTKWTDIPYLGKREDLWCGSLIGHRPRTTWAENIKNTVNMVRRIIGDEEKYMDYLSTQVRYLGEEGSTPGVL

>tr|A0A142I5B9|A0A142I5B9_ZIKV Polyprotein OSZika virus (strain Mr 766) PE4 SV1

MKNPKKKSGGFRIVNMLKRGVARVSPFGGLKRLPAGLLLGHGPIRMVLAILAFLRFTAIKPSLGLINRWGSVGKKEAMEIIKKFKKDLAAMLRIINARKEKKRRGTDTSVGIVGLLLTTAMAVEVTRRGNAYYMYLDRSDAGEAISFPTTMGMNKCYIQIMDLGHMCDATMSYECPMLDEGVEPDDVDCWCNTTSTWVVYGTCHHKKGEARRSRRAVTLPSHSTRKLQTRSQTWLESREYTKHLIRVENWIFRNPGFALAAAAIAWLLGSSTSQKVIYLVMILLIAPAYSIRCIGVSNRDFVEGMSGGTWVDVVLEHGGCVTVMAQDKPTVDIELVTTTVSNMAEVRSYCYEASISDMASDSRCPTQGEAYLDKQSDTQYVCKRTLVDRGWGNGCGLFGKGSLVTCAKFACSKKMTGKSIQPENLEYRIMLSVHGSQHSGMIVNDTGHETDENRAKVEITPNSPRAEATLGGFGSLGLDCEPRTGLDFSDLYYLTMNNKHWLVHKEWFHDIPLPWHAGADTGTPHWNNKEALVEFKDAHAKRQTVVVLGSQEGAVHTALAGALEAEMDGAKGRLSSGHLKCRLKMDKLRLKGVSYSLCTAAFTFTKIPAETLHGTVTVEVQYAGTDGPCKVPAQMAVDMQTLTPVGRLITANPVITESTENSKMMLELDPPFGDSYIVIGVGEKKITHHWHRSGSTIGKAFEATVRGAKRMAVLGDTAWDFGSVGGALNSLGKGIHQIFGAAFKSLFGGMSWFSQILIGTLLVWLGLNTKNGSISLMCLALGGVLIFLSTAVSADVGCSVDFSKKETRCGTGVFVYNDVEAWRDRYKYHPDSPRRLAAAVKQAWEDGICGISSVSRMENIMWRSVEGELNAILEENGVQLTVVVGSVKNPMWRGPQRLPVPVNELPHGWKAWGKSYFVRAAKTNNSFVVDGDTLKECPLKHRAWNSFLVEDHGFGVFHTSVWLKVREDYSLECDPAVIGTAAKGKEAVHSDLGYWIESEKNDTWRLKRAHLIEMKTCEWPKSHTLWTDGIEESDLIIPKSLAGPLSHHNTREGYRTQMKGPWHSEELEIRFEECPGTKVHVEETCGTRGPSLRSTTASGRVIEEWCCRECTMPPLSFRAKDGCWYGMEIRPRKEPESNLVRSMVTAGSTDHMDHFSLGVLVILLMVQEGLKKRMTTKIIISTSMAVLVAMILGGFSMSDLAKLAILMGATFAEMNTGGDVAHLALIAAFKVRPALLVSFIFRANWTPRESMLLALASCLLQTAISALEGDLMVPINGFALAWLAIRAMVVPRTDNITLAILAALTPLARGTLLVAWRAGLATCGGFMLLSLKGKGSVKKNLPFVMALGLTAVRLVDPINVVGLLLLTRSGKRSWPPSEVLTAVGLICALAGGFAKADIEMAGPMAAVGLLIVSYVVSGKSVDMYIERAGDITWEKDAEVTGNSPRLDVALDESGDFSLVEDDGPPMREIILKVVLMAICGMNPIAIPFAAGAWYVYVKTGKRSGALWDVPAPKEVKKGETTDGVYRVMTRRLLGSTQVGVGVMQEGVFHTMWHVTKGSALRSGEGRLDPYWGDVKQDLVSYCGPWKLDAAWDGHSEVQLLAVPPGERARNIQTLPGIFKTKDGDIGAVALDYPAGTSGSPILDKCGRVIGLYGNGVVIKNGSYVSAITQGRREEETPVECFEPSMLKKKQLTVLDLHPGAGKTRRVLPEIVREAIKTRLRTVILAPTRVVAAEMEEALRGLPVRYMTTAVNVTHSGTEIVDLMCHATFTSRLLQPIRVPNYNLYIMDEAHFTDPSSIAARGYISTRVEMGEAAAIFMTATPPGTRDAFPDSNSPIMDTEVEVPERAWSSGFDWVTDHSGKTVWFVPSVRNGNEIAACLTKAGKRVIQLSRKTFETEFQKTKHQEWDFVVTTDISEMGANFKADRVIDSRRCLKPVILDGERVILAGPMPVTHASAAQRRGRIGRNPNKPGDEYLYGGGCAETDEDHAHWLEARMLLDNIYLQDGLIASLYRPEADKVAAIEGEFKLRTEQRKTFVELMKRGDLPVWLAYQVASAGITYTDRRWCFDGTTNNTIMEDSVPAEVWTRYGEKRVLKPRWMDARVCSDHAALKSFKEFAAGKRGAAFGVMEALGTLPGHMTERFQEAIDNLAVLMRAETGSRPYKAAAAQLPETLETIMLLGLLGTVSLGIFFVLMRNKGIGKMGFGMVTLGASAWLMWLSEIEPARIACVLIVVFLLLVVLIPEPEKQRSPQDNQMAIIIMVAVGLLGLITANELGWLERTKSDLSHLMGRREEGATIGFSMDIDLRPASAWAIYAALTTFITPAVQHAVTTSYNNYSLMAMATQAGVLFGMGKGMPFYAWDFGVPLLMIGCYSQLTPLTLIVAIILLVAHYMYLIPGLQAAAARAAQKRTAAGIMKNPVVDGIVVTDIDTMTIDPQVEKKMGQVLLIAVAVSSAILSRTAWGWGEAGALITAATSTLWEGSPNKYWNSSTATSLCNIFRGSYLAGASLIYTVTRNAGLVKRRGGGTGETLGEKWKARLNQMSALEFYSYKKSGITEVCREEARRALKDGVATGGHAVSRGSAKLRWLVERGYLQPYGKVIDLGCGRGGWSYYAATIRKVQEVKGYTKGGPGHEEPMLVQSYGWNIVRLKSGVDVFHMAAEPCDTLLCDIGESSSSPEVEEARTLRVLSMVGDWLEKRPGAFCIKVLCPYTSTMMETLERLQRRYGGGLVRVPLSRNSTHEMYWVSGAKSNTIKSVSTTSQLLLGRMDGPRRPVKYEEDVNLGSGTRAVVSCAEAPNMKIIGNRIERIRSEHAETWFFDENHPYRTWAYHGSYEAPTQGSASSLINGVVRLLSKPWDVVTGVTGIAMTDTTPYGQQRVFKEKVDTRVPDPQEGTRQVMSMVSSWLWKELGKHKRPRVCTKEEFINKVRSNAALGAIFEEEKEWKTAVEAVNDPRFWALVDKEREHHLRGECQSCVYNMMGKREKKQGEFGKAKGSRAIWYMWLGARFLEFEALGFLNEDHWMGRENSGGGVEGLGLQRLGYVLEEMSRIPGGRMYADDTAGWDTRISRFDLENEALITNQMEKGHRALALAIIKYTYQNKVVKVLRPAEKGKTVMDIISRQDQRGSGQVVTYALNTFTNLVVQLIRNMEAEEVLEMQDLWLLRRSEKVTNWLQSNGWDRLKRMAVSGDDCVVKPIDDRFAHALRFLNDMGKVRKDTQEWKPSTGWDNWEEVPFCSHHFNKLHLKDGRSIVVPCRHQDELIGRARVSPGAGWSIRETACLAKSYAQMWQLLYFHRRDLRLMANAICSSVPVDWVPTGRTTWSIHGKGEWMTTEDMLVVWNRVWIEENDHMEDKTPVTKWTDIPYLGKREDLWCGSLIGHRPRTTWAENIKNTVNMMRRIIGDEEKYVDYLSTQVRYLGEEGSTPGVL

>tr|A0A142I5C1|A0A142I5C1_ZIKV Polyprotein OSZika virus (strain Mr 766) PE4 SV1

MKNPKKKSGGFRIVNMLKRGVARVNPLGGLKRLPAGLLLGHGPIRMVLAILAFLRFTAIKPSLGLINRWGSVGKKEAMEIIKKFKKDLAAMLRIINARKERKRRGADTSIGIVGLLLTTAMAAEITRRGSAYYMYLDRSDAGKAISFATTLGVNKCHVQIMDLGHMCDATMSYECPMLDEGVEPDDVDCWCNTTSTWVVYGTCHHKKGEARRSRRAVTLPSHSTRKLQTRSQTWLESREYTKHLIKVENWIFRNPGFALVAVAIAWLLGSSTSQKVIYLVMILLIAPAYSIRCIGVSNRDFVEGMSGGTWVDVVLEHGGCVTVMAQDKPTVDIELVTTTVSNMAEVRSYCYEASISDMASDSRCPTQGEAYLDKQSDTQYVCKRTLVDRGWGNGCGLFGKGSLVTCAKFTCSKKMTGKSIQPENLEYRIMLSVHGSQHSGMIVNDTGHETDENRAKVEVTPNSPRAEATLGGFGSLGLDCEPRTGLDFSDLYYLTMNNKHWLVHKEWFHDIPLPWHAGADTGTPHWNNKEALVEFKDAHAKRQTVVVLGSQEGAVHTALAGALEAEMDGAKGRLFSGHLKCRLKMDKLRLKGVSYSLCTAAFTFTKVPAETLHGTVTVEVQYAGTDGPCKVPAQMAVDMQTLTPVGRLITANPVITESTENSKMMLELDPPFGDSYIVIGVGDKKITHHWHRSGSTIGKAFEATVRGAKRMAVLGDTAWDFGSVGGVFNSLGKGIHQIFGAAFKSLFGGMSWFSQILIGTLLVWLGLNTKNGSISLTCLALGGVMIFLSTAVSADVGCSVDFSKKETRCGTGVFVYNDVEAWRDRYKYHPDSPRRLAAAVKQAWEEGICGISSVSRMENIMWKSVEGELNAILEENGVQLTVVVGSVKNPMWRGPQRLPVPVNELPHGWKAWGKSYFVRAAKTNNSFVVDGDTLKECPLKHRAWNSFLVEDHGFGIFHTSVWLKVREDYSLECDPAVIGTAVKGKEAAHSDLGYWIESEKNDTWRLRRAHLIEMKTCEWPKSHTLWTDGVEESDLIIPKSLAGPLSHHNTREGYRTQVKGPWHSEELEIRFEECPGTKVHVEETCGTRGPSLRSTTASGRVIEEWCCRECTMPPLSFRAKDGCWYGMEIRPRKEPESNLVRSMVTAGSTDHMDHFSLGVLVILLMVQEGLKKRMTTKIIMSTSMAVLVAMVLGGFSMSDLAKLVILMGATFAEMNTGGDVAHLALVAAFKVRPALLVSFIFRANWTPRESMLLALASCLLQTAISALEGELMVLVNGFALAWLAIRAMAVPRTDNIALAILAALTPLARGTLLVAWRAGLATCGGFMLLSLKGKGSVKKNLPFVMALGLTAVRIVDPINVVGLLLLTRSGKRSWPPSEVLTAVGLICALAGGFAKADIEMAGPMAAVGLLIVSYVVSGKSVDMYIERAGDITWEKDAEVTGNSPRLDVALDESGDFSLVEEDGPPMREIILKVVLMAICGMNPIAIPFAAGAWYVYVKTGKRSGALWDVPAPKEVKKGETTDGVYRVMTRRLLGSTQVGVGVMQEGVFHTMWHVTKGAALRSGEGRLDPYWGDVKQDLVSYCGPWKLDAAWDGLSEVQLLAVPPGERARNIQTLPGIFKTKDGDIGAVALDYPAGTSGSPILDKCGRVIGLYGNGVVIKNGSYVSAITQGKREEEAPVECFEPSMLRKKQLTVLDLHPGAGKTRRVLPEIVREAIKKRLRTVILAPTRVVAAEMEEALRGLPVRYMTTAVNVTHSGTEIVDLMCHATFTSRLLQPIRVPNYNLYIMDEAHFTDPSSIAARGYISTRVEMGEAAAIFMTATPPGTRDAFPDSNSPIMDTEVEVPERAWSSGFDWVTDHSGKTIWFVPSVRNGNEIAACLTKAGKRVIQLSRKTFETEFQKTKNQEWDFVITTDISEMGANFKADRVIDSRRCLKPVILDGERVILAGPMPVTHASAAQRRGRIGRNPNKPGDEYMYGGGCAETDEDHAHWLEARMLLDNIYLQDGLIASLYRPEADKVAAIEGEFKLRTEQRKTFVELMKRGDLPVWLAYQVASAGITYTDRRWCFDGTTNNTIMEDSVPAEVWTKYGEKRVLKPRWMDARVCSDHAALKSFKEFAAGKRGAALGVMDALGTLPGHMTERFQEAIDNLAVLMRAETGSRPYKAAAAQLPETLETIMLLGLLGTVSLGIFFVLMRNKGIGKMGFGMVTLGASAWLMWLSEIEPARIACVLIVVFLLLVVLIPEPEKQRSPQDNQMAIIIMVAVGLLGLITANELGWLERTKSDIAHLMGRKEEGTTMGFSMDIDLRPASAWAIYAALTTLITPAVQHAVTTSYNNYSLMAMATQAGVLFGMGKGMPFYAWDFGVPLLMMGCYSQLTPLTLIVAIILLVAHYMYLIPGLQAAAARAAQKRTAAGIMKNPVVDGIVVTDIDTMTIDPQVEKKMGQVLLIAVAVSSAVLLRTAWGWGEAGALITAATSTLWEGSPNKYWNSSTATSLCNIFRGSYLAGASLIHTVTRNAGLVKRRGGGTGETLGEKWKARLNQMSALEFYSYKKSGITEVCREEARRALKDGVATGGHAVSRGSAKLRWLVERGYLQPHGKVVDLGCGRGGWSYYAATIRKVQEVRGYTKGGPGHEEPMLVQSYGWNIVRLKSGVDVFHMAAEPCDTLLCDIGESSSSPEVEETRTLRVLSMVGDWLEKRPGAFCIKVLCPYTSTMMETMERLQRRHGGGLVRVPLSRNSTHEMYWVSGAKSNIIKSVSTTSQLLLGRMEGPRRPVKYEEDVNLGSGTRAVASCAEAPNMKIIGRRIERIRNEHAETWFFDENHPYRTWAYHGSYEAPTQGSASSLVNGVVRLLSKPWDVVTGVTGIAMTDTTPYGQQRVFKEKVDTRVPDPQEGTRQVMNMVSSWLWKELGKRKRPRVCTKEEFINKVRSNAALGAIFEEEKEWKTAVEAVNDPRFWALVDKEREHHLRGECHSCVYNMMGKREKKQGEFGKAKGSRAIWYMWLGARFLEFEALGFLNEDHWMGRENSGGGVEGLGLQRLGYILEEMNRAPGGKMYADDTAGWDTRISKFDLENEALITNQMEEGHRALALAVIKYTYQNKVVKVLRPAEGGKTVMDIISRQDQRGSGQVVTYALNTFTNLVVQLIRNMEAEEVLEMHDLWLLRKPEKVTRWLQSNGWDRLKRMAVSGDDCVVKPIDDRFAHALRFLNDMGKVRKDTQEWKPSTGWSNWEEVPFCSHHFNKLHLKDGRSIVVPCRHQDELIGRARVSPGAGWSIRETACLAKSYAQMWQLLYFHRRDLRLMANAICSAVPIDWVPTGRTTWSIHGKGEWMTTEDMLMVWNRVWIEENDHMEDKTPVTKWTDIPYLGKREDLWCGSLIGHRPRTTWAENIKDTVNMVRRIIGDEEKFMDYLSTQVRYLGEEGSTPGVL

>tr|A0A142I5B8|A0A142I5B8_ZIKV Polyprotein OSZika virus (strain Mr 766) PE4 SV1

MKNPKKKSGGFRIVNMLKRGVARVNPLGGLKRLPAGLLLGHGPIRMVLAILAFLRFTAIKPSLGLINRWGSVGKKEAMEIIKKFKKDLAAMLRIINARKERKRRGADTSIGIVGLLLTTAMAAEITRRGSAYYMYLDRSDAGKAISFATTLGVNKCHVQIMDLGHMCDATMSYECPMLDEGVEPDDVDCWCNTTSTWVVYGTCHHKKGEARRSRRAVTLPSHSTRKLQTRSQTWLESREYTKHLIKVENWIFRNPGFALVAVAIAWLLGSSTSQKVIYLVMILLIAPAYSIRCIGVSNRDFVEGMSGGTWVDVVLEHGGCVTVMAQDKPTVDIELVTTTVSNMAEVRSYCYEASISDMASDSRCPTQGEAYLDKQSDTQYVCKRTLVDRGWGNGCGLFGKGSLVTCAKFTCSKKMTGKSIQPENLEYRIMLSVHGSQHSGMIVNDTGHETDENRAKVEVTPNSPRAEATLGGFGSLGLDCEPRTGLDFSDLYYLTMNNKHWLVHKEWFHDIPLPWHAGADTGTPHWNNKEALVEFKDAHAKRQTVVVLGSQEGAVHTALAGALEAEMDGAKGRLFSGHLKCRLKMDKLRLKGVSYSLCTAAFTFTKVPAETLHGTVTVEVQYAGTDGPCKVPAQMAVDMQTLTPVGRLITANPVITESTENSKMMLELDPPFGDSYIVIGVGDKKITHHWHRSGSTIGKAFEATVRGAKRMAVLGDTAWDFGSVGGVFNSLGKGIHQIFGAAFKSLFGGMSWFSQILIGTLLVWLGLNTKNGSISLTCLALGGVMIFLSTAVSADVGCSVDFSKKETRCGTGVFVYNDVEAWRDRYKYHPDSPRRLAAAVKQAWEEGICGISSVSRMENIMWKSVEGELNAILEENGVQLTVVVGSVKNPMWRGPQRLPVPVNELPHGWKAWGKSYFVRAAKTNNSFVVDGDTLKECPLKHRAWNSFLVEDHGFGIFHTSVWLKVREDYSLECDPAVIGTAVKGKEAAHSDLGYWIESEKNDTWRLRRAHLIEMKTCEWPKSHTLWTDGVEESDLIIPKSLAGPLSHHNTREGYRTQVKGPWHSEELEIRFEECPGTKVHVEETCGTRGPSLRSTTASGRVIEEWCCRECTMPPLSFRAKDGCWYGMEIRPRKEPESNLVRSMVTAGSTDHMDHFSLGVLVILLMVQEGLKKRMTTKIIMSTSMAVLVAMVLGGFSMSDLAKLVILMGATFAEMNTGGDVAHLALVAAFKVRPALLVSFIFRANWTPRESMLLALASCLLQTAISALEGELMVLVNGFALAWLAIRAMAVPRTDNIALAILAALTPLARGTLLVAWRAGLATCGGFMLLSLKGKGSVKKNLPFVMALGLTAVRIVDPINVVGLLLLTRSGKRSWPPSEVLTAVGLICALAGGFAKADIEMAGPMAAVGLLIVSYVVSGKSVDMYIERAGDITWEKDAEVTGNSPRLDVALDESGDFSLVEEDGPPMREIILKVVLMAICGMNPIAIPFAAGAWYVYVKTGKRSGALWDVPAPKEVKKGETTDGVYRVMTRRLLGSTQVGVGVMQEGVFHTMWHVTKGAALRSGEGRLDPYWGDVKQDLVSYCGPWKLDAAWDGLSEVQLLAVPPGERARNIQTLPGIFKTKDGDIGAVALDYPAGTSGSPILDKCGRVIGLYGNGVVIKNGSYVSAITQGKREEEAPVECFEPSMLRKKQLTVLDLHPGAGKTRRVLPEIVREAIKKRLRTVILAPTRVVAAEMEEALRGLPVRYMTTAVNVTHSGTEIVDLMCHATFTSRLLQPIRVPNYNLYIMDEAHFTDPSSIAARGYISTRVEMGEAAAIFMTATPPGTRDAFPDSNSPIMDTEVEVPERAWSSGFDWVTDHSGKTIWFVPSVRNGNEIAACLTKAGKRVIQLSRKTFETEFQKTKNQEWDFVITTDISEMGANFKADRVIDSRRCLKPVILDGERVILAGPMPVTHASAAQRRGRIGRNPNKPGDEYMYGGGCAETDEDHAHWLEARMLLDNIYLQDGLIASLYRPEADKVAAIEGEFKLRTEQRKTFVELMKRGDLPVWLAYQVASAGITYTDRRWCFDGTTNNTIMEDSVPAEVWTKYGEKRVLKPRWMDARVCSDHAALKSFKEFAAGKRGAALGVMDALGTLPGHMTERFQEAIDNLAVLMRAETGSRPYKAAAAQLPETLETIMLLGLLGTVSLGIFFVLMRNKGIGKMGFGMVTLGASAWLMWLSEIEPARIACVLIVVFLLLVVLIPEPEKQRSPQDNQMAIIIMVAVGLLGLITANELGWLERTKSDIAHLMGRKEEGTTMGFSMDIDLRPASAWAIYAALTTLITPAVQHAVTTSYNNYSLMAMATQAGVLFGMGKGMPFYAWDFGVPLLMMGCYSQLTPLTLIVAIILLVAHYMYLIPGLQAAAARAAQKRTAAGIMKNPVVDGIVVTDIDTMTIDPQVEKKMGQVLLIAVAVSSAVLLRTAWGWGEAGALITAATSTLWEGSPNKYWNSSTATSLCNIFRGSYLAGASLIYTVTRNAGLVKRRGGGTGETLGEKWKARLNQMSALEFYSYKKSGITEVCREEARRALKDGVATGGHAVSRGSAKLRWLVERGYLQPHGKVVDLGCGRGGWSYYAATIRKVQEVRGYTKGGPGHEEPMLVQSYGWNIVRLKSGVDVFHMAAEPCDTLLCDIGESSSSPEVEETRTLRVLSMVGDWLEKRPGAFCIKVLCPYTSTMMETMERLQRRHGGGLVRVPLSRNSTHEMYWVSGAKSNIIKSVSTTSQLLLGRMEGPRRPVKYEEDVNLGSGTRAVASCAEAPNMKIIGRRIERIRNEHAETWFFDENHPYRTWAYHGSYEAPTQGSASSLVNGVVRLLSKPWDVVTGVTGIAMTDTTPYGQQRVFKEKVDTRVPDPQEGTRQVMNMVSSWLWKELGKRKRPRVCTKEEFINKVRSNAALGAIFEEEKEWKTAVEAVNDPRFWALVDKEREHHLRGECHSCVYNMMGKREKKQGEFGKAKGSRAIWYMWLGARFLEFEALGFLNEDHWMGRENSGGGVEGLGLQRLGYILEEMNRAPGGKMYADDTAGWDTRISKFDLENEALITNQMEEGHRALALAVIKYTYQNKVVKVLRPAEGGKTVMDIISRQDQRGSGQVVTYALNTFTNLVVQLIRNMEAEEVLEMHDLWLLRKPEKVTRWLQSNGWDRLKRMAVSGDDCVVKPIDDRFAHALRFLNDMGKVRKDTQEWKPSTGWSNWEEVPFCSHHFNKLHLKDGRSIVVPCRHQDELIGRARVSPGAGWSIRETACLAKSYAQMWQLLYFHRRDLRLMANAICSAVPIDWVPTGRTTWSIHGKGEWMTTEDMLMVWNRVWIEENDHMEDKTPVTKWTDIPYLGKREDLWCGSLIGHRPRTTWAENIKDTVNMVRRIIGDEEKFMDYLSTQVRYLGEEGSTPGVL

>tr|A0A142I5B7|A0A142I5B7_ZIKV Polyprotein OSZika virus (strain Mr 766) PE4 SV1

MKNPKKKSGGFRIVNMLKRGVARVNPLGGLKRLPAGLLLGHGPIRMVLAILAFLRFTAIKPSLGLINRWGSVGKKEAMEIIKKFKKDLAAMLRIINARKERKRRGADTSIGIVGLLLTTAMAAEITRRGSAYYMYLDRSDAGKAISFATTLGVNKCHVQIMDLGHMCDATMSYECPMLDEGVEPDDVDCWCNTTSTWVVYGTCHHKKGEARRSRRAVTLPSHSTRKLQTRSQTWLESREYTKHLIKVENWIFRNPGFALVAVAIAWLLGSSTSQKVIYLVMILLIAPAYSIRCIGVSNRDFVEGMSGGTWVDVVLEHGGCVTVMAQDKPTVDIELVTTTVSNMAEVRSYCYEASISDMASDSRCPTQGEAYLDKQSDTQYVCKRTLVDRGWGNGCGLFGKGSLVTCAKFTCSKKMTGKSIQPENLEYRIMLSVHGSQHSGMIVNDTGHETDENRAKVEVTPNSPRAEATLGGFGSLGLDCEPRTGLDFSDLYYLTMNNKHWLVHKEWFHDIPLPWHAGADTGTPHWNNKEALVEFKDAHAKRQTVVVLGSQEGAVHTALAGALEAEMDGAKGRLFSGHLKCRLKMDKLRLKGVSYSLCTAAFTFTKVPAETLHGTVTVEVQYAGTDGPCKVPAQMAVDMQTLTPVGRLITANPVITESTENSKMMLELDPPFGDSYIVIGVGDKKITHHWHRSGSTIGKAFEATVRGAKRMAVLGDTAWDFGSVGGVFNSLGKGIHQIFGAAFKSLFGGMSWFSQILIGTLLVWLGLNTKNGSISLTCLALGGVMIFLSTAVSADVGCSVDFSKKETRCGTGVFVYNDVEAWRDRYKYHPDSPRRLAAAVKQAWEEGICGISSVSRMENIMWKSVEGELNAILEENGVQLTVVVGSVKNPMWRGPQRLPVPVNELPHGWKAWGKSYFVRAAKTNNSFVVDGDTLKECPLKHRAWNSFLVEDHGFGIFHTSVWLKVREDYSLECDPAVIGTAVKGKEAAHSDLGYWIESEKNDTWRLRRAHLIEMKTCEWPKSHTLWTDGVEESDLIIPKSLAGPLSHHNTREGYRTQVKGPWHSEELEIRFEECPGTKVHVEETCGTRGPSLRSTTASGRVIEEWCCRECTMPPLSFRAKDGCWYGMEIRPRKEPESNLVRSMVTAGSTDHMDHFSLGVLVILLMVQEGLKKRMTTKIIMSTSMAVLVAMVLGGFSMSDLAKLVILMGATFAEMNTGGDVAHLALVAAFKVRPALLVSFIFRANWTPRESMLLALASCLLQTAISALEGELMVLVNGFALAWLAIRAMAVPRTDNIALAILAALTPLARGTLLVAWRAGLATCGGFMLLSLKGKGSVKKNLPFVMALGLTAVRIVDPINVVGLLLLTRSGKRSWPPSEVLTAVGLICALAGGFAKADIEMAGPMAAVGLLIVSYVVSGKSVDMYIERAGDITWEKDAEVTGNSPRLDVALDESGDFSLVEEDGPPMREIILKVVLMAICGMNPIAIPFAAGAWYVYVKTGKRSGALWDVPAPKEVKKGETTDGVYRVMTRRLLGSTQVGVGVMQEGVFHTMWHVTKGAALRSGEGRLDPYWGDVKQDLVSYCGPWKLDAAWDGLSEVQLLAVPPGERARNIQTLPGIFKTKDGDIGAVALDYPAGTSGSPILDKCGRVIGLYGNGVVIKNGSYVSAITQGKREEEAPVECFEPSMLRKKQLTVLDLHPGAGKTRRVLPEIVREAIKKRLRTVILAPTRVVAAEMEEALRGLPVRYMTTAVNVTHSGTEIVDLMCHATFTSRLLQPIRVPNYNLYIMDEAHFTDPSSIAARGYISTRVEMGEAAAIFMTATPPGTRDAFPDSNSPIMDTEVEVPERAWSSGFDWVTDHSGKTIWFVPSVRNGNEIAACLTKAGKRVIQLSRKTFETEFQKTKNQEWDFVITTDISEMGANFKADRVIDSRRCLKPVILDGERVILAGPMPVTHASAAQRRGRVGRNPNKPGDEYMYGGGCAETDEDHAHWLEARMLLDNIYLQDGLIASLYRPEADKVAAIEGEFKLRTEQRKTFVELMKRGDLPVWLAYQVASAGITYTDRRWCFDGTTNNTIMEDSVPAEVWTKYGEKRVLKPRWMDARVCSDHAALKSFKEFAAGKRGAALGVMDALGTLPGHMTERFQEAIDNLAVLMRAETGSRPYKAAAAQLPETLETIMLLGLLGTVSLGIFFVLMRNKGIGKMGFGMVTLGASAWLMWLSEIEPARIACVLIVVFLLLVVLIPEPEKQRSPQDNQMAIIIMVAVGLLGLITANELGWLERTKSDIAHLMGRKEEGTTMGFSMDIDLRPASAWAIYAALTTLITPAVQHAVTTSYNNYSLMAMATQAGVLFGMGKGMPFYAWDFGVPLLMMGCYSQLTPLTLIVAIILLVAHYMYLIPGLQAAAARAAQKRTAAGIMKNPVVDGIVVTDIDTMTIDPQVEKKMGQVLLIAVAVSSAVLLRTAWGWGEAGALITAATSTLWEGSPNKYWNSSTATSLCNIFRGSYLAGASLIYTVTRNAGLVKRRGGGTGETLGEKWKARLNQMSALEFYSYKKSGITEVCREEARRALKDGVATGGHAVSRGSAKLRWLVERGYLQPHGKVVDLGCGRGGWSYYAATIRKVQEVRGYTKGGPGHEEPMLVQSYGWNIVRLKSGVDVFHMAAEPCDTLLCDIGESSSSPEVEETRTLRVLSMVGDWLEKRPGAFCIKVLCPYTSTMMETMERLQRRHGGGLVRVPLSRNSTHEMYWVSGAKSNIIKSVSTTSQLLLGRMEGPRRPVKYEEDVNLGSGTRAVASCAEAPNMKIIGRRIERIRNEHAETWFFDENHPYRTWAYHGSYEAPTQGSASSLVNGVVRLLSKPWDVVTGVTGIAMTDTTPYGQQRVFKEKVDTRVPDPQEGTRQVMNMVSSWLWKELGKRKRPRVCTKEEFINKVRSNAALGAIFEEEKEWKTAVEAVNDPRFWALVDKEREHHLRGECHSCVYNMMGKREKKQGEFGKAKGSRAIWYMWLGARFLEFEALGFLNEDHWMGRENSGGGVEGLGLQRLGYILEEMNRAPGGKMYADDTAGWDTRISKFDLENEALITNQMEEGHRALALAVIKYTYQNKVVKVLRPAEGGKTVMDIISRQDQRGSGQVVTYALNTFTNLVVQLIRNMEAEEVLEMHDLWLLRKPEKVTRWLQSNGWDRLKRMAVSGDDCVVKPIDDRFAHALRFLNDMGKVRKDTQEWKPSTGWSNWEEVPFCSHHFNKLHLKDGRSIVVPCRHQDELIGRARVSPGAGWSIRETACLAKSYAQMWQLLYFHRRDLRLMANAICSAVPIDWVPTGRTTWSIHGKGEWMTTEDMLMVWNRVWIEENDHMEDKTPVTKWTDIPYLGKREDLWCGSLIGHRPRTTWAENIKDTVNMVRRIIGDEEKFMDYLSTQVRYLGEEGSTPGVL

>tr|A0A141ZVF6|A0A141ZVF6_ZIKV Polyprotein OSZika virus (strain Mr 766) PE4 SV1

MKNPKKKSGGFRIVNMLKRGVARVSPFGGLKRLPAGLLLGHGPIRMVLAILAFLRFTAIKPSLGLINRWGSVGKKEAMEIIKKFKKDLAAMLRIINARKEKKRRGADTNVGIVGLLLTTAMAAEVTRRGSAYYMYLDRNDAGEAISFPTTLGMNKCYIQIMDLGHMCDATMSYECPMLDEGVEPDDVDCWCNTTSTWVVYGTCHHKKGEARRSRRAVTLPSHSTRKLQTRSQTWLESREYTKHLIRVENWIFRNPGFALAAAAIAWLLGSSTSQKVIYLVMILLIAPAYSIRCIGVSNRDFVEGMSGGTWVDVVLEHGGCVTVMAQDKPTVDIELVTTTVSNMAEVRSYCYEASISDMASDSRCPTQGEAYLDKQSDTQYVCKRTLVDRGWGNGCGLFGKGSLVTCAKFACSKKMTGKSIQPENLEYRIMLSVHGSQHSGMIVNDTGHETDENRAKVEITPNSPRAEATLGGFGSLGLDCEPRTGLDFSDLYYLTMNNKHWLVHKEWFHDIPLPWHAGADTGTPHWNNKEALVEFKDAHAKRQTVVVLGSQEGAVHTALAGALEAEMDGAKGRLSSGHLKCRLKMDKLRLKGVSYSLCTAAFTFTKIPAETLHGTVTVEVQYAGTDGPCKVPAQMAVDMQTLTPVGRLITANPVITESTENSKMMLELDPPFGDSYIVIGVGEKKITHHWHRSGSTIGKAFEATVRGARRMAVLGDTAWDFGSVGGALNSLGKGIHQIFGAAFKSLFGGMSWFSQILIGTLLMWLGLNTKNGSISLMCLALGGVLIFLSTAVSADVGCSVDFSKKETRCGTGVFVYNDVEAWRDRYKYHPDSPRRLAAAVKQAWEDGICGISSVSRMENIMWRSVEGELNAILEENGVQLTVVVGSVKNPMWRGPQRLPVPVNELPHGWKAWGKSYFVRAAKTNNSFVVDGDTLKECPLKHRAWNSFLVEDHGFGVFHTSVWLKVREDYSLECDPAVIGTAVKGKEAVHSDLGYWIESEKNDTWRLKRAHLIEMKTCEWPKSHTLWTDGIEESDLIIPKSLAGPLSHHNTREGYRTQMKGPWHSEELEIRFEECPGTKVHVEETCGTRGPSLRSTTASGRVIEEWCCRECTMPPLSFQAKDGCWYGMEIRPRKEPESNLVRSMVTAGSTDHMDHFSLGVLVILLMVQEGLKKRMTTKIIISTSMAVLVAMILGGFSMSDLAKLAILMGATFAEMNTGGDVAHLALIAAFKVRPALLVSFIFRANWTPRESMLLALASCLLQTAISALEGDLMVLINGFALAWLAIRAMVVPRTDNITLAILAALTPLARGTLLVAWRAGLATCGGFMLLSLKGKGSVKKNLPFVMALGLTAVRLVDPINVVGLLLLTRSGKRSWPPSEVLTAVGLICALAGGFAKADIEMAGPMAAVGLLIVSYVVSGKSVDMYIERAGDITWEKDAEVTGNSPRLDVALDESGDFSLVEDDGPPMREIILKVVLMTICGMNPIAIPFAAGAWYVYVKTGKRSGALWDVPAPKEVKKGETTDGVYRVMTRRLLGSTQVGVGVMQEGVFHTMWHVTKGSALRSGEGRLDPYWGDVKQDLVSYCGPWKLDAAWDGHSEVQLLAVPPGERARNIQTLPGIFKTKDGDIGAVALDYPAGTSGSPILDKCGRVIGLYGNGVVIKNGSYVSAITQGRREEETPVECFEPSMLKKKQLTVLDLHPGAGKTRRVLPEIVREAIKTRLRTVILAPTRVVAAEMEEALRGLPVRYMTTAVNVTHSGTEIVDLMCHATFTSRLLQPIRVPNYNLYIMDEAHFTDPSSIAARGYISTRVEMGEAAAIFMTATPPGTRDAFPDSNSPIMDTEVEVPERAWSSGFDWVTDHSGKTVWFVPSVRNGNEIAACLTKAGKRVIQLSRKTFETEFQKTKHQEWDFVVTTDISEMGANFKADRVIDSRRCLKPVILDGERVILAGPMPVTHASAAQRRGRIGRNPNKPGDEYLYGGGCAETDEDHAHWLEARMLLDNIYLQDGLIASLYRPEADKVAAIEGEFKLKTEQRKTFVELMKRGDLPVWLAYQVASAGITYTDRRWCFDGTTNNTIMEDSVPAEVWTRHGEKRVLKPRWMDARVCSDHAALKSFKEFAAGKRGAAFGVMEALGTLPGHMTERFQEAIDNLAVLMRAETGSRPYKAAAAQLPETLETIMLLGLLGTVSLGIFFVLMRNKGIGKMGFGMVTLGASAWLMWLSEIEPARIACVLIVVFLLLVVLIPEPEKQRSPQDNQMAIIIMVAVGLLGLITANELGWLERTKSDLSHLMGRREEGATIGFSMDIDLRPASAWAIYAALTTFITPAVQHAVTTSYNNYSLMAMATQAGVLFGMGKGMPFYAWDFGVPLLMIGCYSQLTPLTLIVAIILLVAHYMYLIPGLQAAAARAAQKRTAAGIMKNPVVDGIVVTDIDTMTIDPQVEKKMGQVLLIAVAVSSAILSRTAWGWGEAGALITAATSTLWEGSPNKYWNSSTATSLCNIFRGSYLAGASLIYTVTRNAGLVKRRGGGTGETLGEKWKARLNQMSALEFYSYKKSGITEVCREEARRALKDGVATGGHAVSRGSAKLRWLVERGYLQPYGKVIDLGCGRGGWSYYAATIRKVQEVKGYTKGGPGHEEPMLVQSYGWNIIRLKSGVDVFHMAAEPCDTLLCDIGESSSSPEVEEARTLRVLSMVGDWLEKRPGAFCIKVLCPYTSTMMETLERLQRRYGGGLVRVPLSRNSTHEMYWVSGAKSNTIKSVSTTSQLLLGRMDGPRRPVKYEEDVNLGSGTRAVVSCAEAPNMKIIGNRIERIRSEHAETWFFDENHPYRTWAYHGSYEAPTQGSASSLINGVVRLLSKPWDVVTGVTGIAMTDTTPYGQQRVFKEKVDTRVPDPQEGTRQVMSMVSSWLWKELGKHKRPRVCTKEEFINKVRSNAALGAIFEEEKEWKTAVEAVNDPRFWALVDKEREHHLRGECQSCVYNMMGKREKKQGEFGKAKGSRAIWYMWLGARFLEFEALGFLNEDHWMGRENSGGGVEGLGLQRLGYVLEEMSRIPGGRMYADDTAGWDTRISRFDLENEALITNQMEKGHRALALAIIKYTYQNKVVKVLRPAEKGKTVMDIISRQDQRGSGQVVTYALNTFTNLVVQLIRSMEAEEVLEMQDLWLLRRSEKVTNWLQSNGWDRLKRMAVSGDDCVVRPIDDRFAHALRFLNDMGKVRKDTQEWKPSTGWDNWEEVPFCSHHFNKLHLKDGRSIVVPCRHQDELIGRARVSPGAGWSIRETACLAKSYAQMWQLLYFHRRDLRLMANAICSSVPVDWVPTGRTTWSIHGKGEWMTTEDMLVVWNRVWIEENDHMEDKTPVTKWTDIPYLGKREDLWCGSLIGHRPRTTWAENIKNTVNMMRRIIGDEEKYVDYLSTQVRYLGEEGSTPGVL

>tr|A0A142EHG8|A0A142EHG8_ZIKV Polyprotein OSZika virus (strain Mr 766) PE4 SV1

MKNPKKKSGGFRIVNMLKRGVARVSPFGGLKRLPAGLLLGHGPIRMVLAILAFLRFTAIKPSLGLINRWGSVGKKEAMEIIKKFKKDLAAMLRIINARKEKKRRGADTSVGIVGLLLTTAMAAEVTRRGSAYYMYLDRNDAGEAISFPTTLGMNKCYIQIMDLGHMCDATMSYECPMLDEGVEPDDVDCWCNTTSTWVVYGTCHHKKGEARRSRRAVTLPSHSTRKLQTRSQTWLESREYTKHLIRVENWIFRNPGFALAAAAIAWLLGSSTSQKVIYLVMILLIAPAYSIRCIGVSNRDFVEGMSGGTWVDVVLEHGGCVTVMAQDKPTVDIELVTTTVSNMAEVRSYCYEASISDMASDSRCPTQGEAYLDKQSDTQYVCKRTLVDRGWGNGCGLFGKGSLVTCAKFACSKKMTGKSIQPENLEYRIMLSVHGSQHSGMIVNDTGHETDENRAKVEITPNSPRAEATLGGFGSLGLDCEPRTGLDFSDLYYLTMNNKHWLVHKEWFHDIPLPWHAGADTGTPHWNNKEALVEFKDAHAKRQTVVVLGSQEGAVHTALAGALEAEMDGAKGRLSSGHLKCRLKMDKLRLKGVSYSLCTAAFTFTKIPAETLHGTVTVEVQYAGTDGPCKVPAQMAVDMQTLTPVGRLITANPVITESTENSKMMLELDPPFGDSYIVIGVGEKKITHHWHRSGSTIGKAFEATVRGAKRMAVLGDTAWDFGSVGGALNSLGKGIHQIFGAAFKSLFGGMSWFSQILIGTLLMWLGLNTKNGSISLMCLALGGVLIFLSTAVSADVGCSVDFSKKETRCGTGVFVYNDVEAWRDRYKYHPDSPRRLAAAVKQAWEDGICGISSVSRMENIMWRSVEGELNAILEENGVQLTVVVGSVKNPMWRGPQRLPVPVNELPHGWKAWGKSYFVRAAKTNNSFVVDGDTLKECPLKHRAWNSFLVEDHGFGVFHTSVWLKVREDYSLECDPAVIGTAVKGKEAVHSDLGYWIESEKNDTWRLKRAHLIEMKTCEWPKSHTLWTDGIEESDLIIPKSLAGPLSHHNTREGYRTQMKGPWHSEELEIRFEECPGTKVHVEETCGTRGPSLRSTTASGRVIEEWCCRECTMPPLSFRAKDGCWYGMEIRPRKEPESNLVRSMVTAGSTDHMDHFSLGVLVILLMVQEGLKKRMTTKIIISTSMAVLVAMILGGFSMSDLAKLAILMGATFAEMNTGGDVAHLALIAAFKVRPALLVSFIFRANWTPRESMLLALASCLLQTAISALEGDLMVLINGFALAWLAIRAMVVPRTDNITLAILAALTPLARGTLLVAWRAGLATCGGFMLLSLKGKGSVKKNLPFVMALGLTAVRLVDPINVVGLLLLTRSGKRSWPPSEVLTAVGLICALAGGFAKADIEMAGPIAAVGLLIVSYVVSGKSVDMYIERAGDITWEKDAEVTGNSPRLDVALDESGDFSLVEDDGPPMREIILKVVLMTICGMNPIAIPFAAGAWYVYVKTGKRSGALWDVPAPKEVKKGETTDGVYRVMTRRLLGSTQVGVGVMQEGVFHTMWHVTKGSALRSGEGRLDPYWGDVKQDLVSYCGPWKLDAAWDGHSEVQLLAVPPGERARNIQTLPGIFKTKDGDIGAVALDYPAGTSGSPILDKCGRVIGLYGNGVVIKNGSYVSAITQGRREEETPVECFEPSMLKKKQLTVLDLHPGAGKTRRVLPEIVREAIKTRLRTVILAPTRVVAAEMEEALRGLPVRYMTTAVNVTHSGTEIVDLMCHATFTSRLLQPIRVPNYNLYIMDEAHFTDPSSIAARGYISTRVEMGEAAAIFMTATPPGTRDAFPDSNSPIMDTEVEVPERAWSSGFDWVTDHSGKTVWFVPSVRNGNEIAACLTKAGKRVIQLSRKTFETEFQKTKHQEWDFVVTTDISEMGANFKADRVIDSRRCLKPVILDGERVILAGPMPVTHASAAQRRGRIGRNPNKPGDEYLYGGGCAETDEDHAHWLEARMLLDNIYLQDGLIASLYRPEADKVAAIEGEFKLRTEQRKTFVELMRRGDLPVWLAYQVASAGITYTDRRWCFDGTTNNTIMEDSVPAEVWTRHGEKRVLKPRWMDARVCSDHAALKSFKEFAAGKRGAAFGVMEALGTLPGHMTERFQEAIDNLAVLMRAETGSRPYKAAAAQLPETLETIMLLGLLGTVSLGIFFVLMRNKGIGKMGFGMVTLGASAWLMWLSEIEPARIACVLIVVFLLLVVLIPEPEKQRSPQDNQMAIIIMVAVGLLGLITANELGWLERTKSDLSHLMGRREEGATIGFSMDIDLRPASAWAIYAALTTFITPAVQHAVTTSYNNYSLMAMATQAGVLFGMGKGMPFYAWDFGVPLLMIGCYSQLTPLTLIVAIILLVAHYMYLIPGLQAAAARAAQKRTAAGIMKNPVVDGIVVTDIDTMTIDPQVEKKMGQVLLIAVAVSSAILSRTAWGWGEAGALITAATSTLWEGSPNKYWNSSTATSLCNIFRGSYLAGASLIYTVTRNAGLVKRRGGGTGETLGEKWKARLNQMSALEFYSYKKSGITEVCREEARRALKDGVATGGHAVSRGSAKLRWLVERGYLQPYGKVIDLGCGRGGWSYYAATIRKVQEVKGYTKGGPGHEEPVLVQSYGWNIVRLKSGVDVFHMAAEPCDTLLCDIGESSSSPEVEEARTLRVLSMVGDWLEKRPGAFCIKVLCPYTSTMMETLERLQRRYGGGLVRVPLSRNSTHEMYWVSGAKSNTIKSVSTTSQLLLGRMDGPRRPVKYEEDVNLGSGTRAVVSCAEAPNMKIIGNRIERIRSEHAETWFFDENHPYRTWAYHGSYEAPTQGSASSLINGVVRLLSKPWDVVTGVTGIAMTDTTPYGQQRVFKEKVDTRVPDPQEGTRQVMSMVSSWLWKELGKHKRPRVCTKEEFINKVRSNAALGAIFEEEKEWKTAVEAVNDPRFWALVDKEREHHLRGECQSCVYNMMGKREKKQGEFGKAKGSRAIWYMWLGARFLEFEALGFLNEDHWMGRENSGGGVEGLGLQRLGYVLEEMSRIPGGRMYADDTAGWDTRISRFDLENEALITNQMEKGHRALALAIIKYTYQNKVVKVLRPAEKGKTVMDIISRQDQRGSGQVVTYALNTFTNLVVQLIRNMEAEEVLEMQDLWLLRRSEKVTNWLQSNGWDRLKRMAVSGDDCVVKPIDDRFAHALRFLNDMGKVRKDTQEWKPSTGWDNWEEVPFCSHHFNKLHLKDGRSIVVPCRHQDELIGRARVSPGAGWSIRETACLAKSYAQMWQLLYFHRRDLRLMANAICSSVPVDWVPTGRTTWSIHGKGEWMTTEDMLVVWNRVWIEENDHMEDKTPVTKWTDIPYLGKREDLWCGSLIGHRPRTTWAENIKNTVNMVRRIIGDEEKYMDYLSTQVRYLGEEGSTPGVL

>tr|A0A142FH79|A0A142FH79_ZIKV Polyprotein OSZika virus (strain Mr 766) PE4 SV1

MKNPKKKSGGFRIVNMLKRGVARVSPFGGLKRLPAGLLLGHGPIRMVLAILAFLRFTAIKPSLGLINRWGSVGKKEAMEIIKKFKKDLAAMLRIINARKEKKRRGADTSVGIVGLLLTTAMAAEVTRRGSAYYMYLDRNDAGEAISFPTTLGMNKCYIQIMDLGHMCDATMSYECPMLDEGVEPDDVDCWCNTTSTWVVYGTCHHKKGEARRSRRAVTLPSHSTRKLQTRSQTWLESREYTKHLIRVENWIFRNPGFALAAAAIAWLLGSSTSQKVIYLVMILLIAPAYSIRCIGVSNRDFVEGMSGGTWVDVVLEHGGCVTVMAQDKPTVDIELVTTTVSNMAEVRSYCYEASISDMASDSRCPTQGEAYLDKQSDTQYVCKRTLVDRGWGNGCGLFGKGSLVTCAKFACSKKMTGKSIQPENLEYRIMLSVHGSQHSGMIVNDTGHETDENRAKVEITPNSPRAEATLGGFGSLGLDCEPRTGLDFSDLYYLTMNNKHWLVHKEWFHDIPLPWHAGADTGTPHWNNKEALVEFKDAHAKRQTVVVLGSQEGAVHTALAGALEAEMDGAKGRLSSGHLKCRLKMDKLRLKGVSYSLCTAAFTFTKIPAETLHGTVTVEVQYAGTDGPCKVPAQMAVDMQTLTPVGRLITANPVITESTENSKMMLELDPPFGDSYIVIGVGEKKITHHWHRSGSTIGKAFEATVRGAKRMAVLGDTAWDFGSVGGALNSLGKGIHQIFGAAFKSLFGGMSWFSQILIGTLLMWLGLNTKNGSISLMCLALGGVLIFLSTAVSADVGCSVDFSKKETRCGTGVFVYNDVEAWRDRYKYHPDSPRRLAAAVKQAWEDGICGISSVSRMENIMWRSVEGELNAILEENGVQLTVVVGSVKNPMWRGPQRLPVPVNELPHGWKAWGKSYFVRAAKTNNSFVVDGDTLKECPLKHRAWNSFLVEDHGFGVFHTSVWLKVREDYSLECDPAVIGTAVKGKEAVHSDLGYWIESEKNDTWRLKRAHLIEMKTCEWPKSHTLWTDGIEESDLIIPKSLAGPLSHHNTREGYRTQMKGPWHSEELEIRFEECPGTKVHVEETCGTRGPSLRSTTASGRVIEEWCCRECTMPPLSFRAKDGCWYGMEIRPRKEPESNLVRSMVTAGSTDHMDHFSLGVLVILLMVQEGLKKRMTTKIIISTSMAVLVAMILGGFSMSDLAKLAILMGATFAEMNTGGDVAHLALIAAFKVRPALLVSFIFRANWTPRESMLLALASCLLQTAISALEGDLMVLINGFALAWLAIRAMVVPRTDNITLAILAALTPLARGTLLVAWRAGLATCGGFMLLSLKGKGSVKKNLPFVMALGLTAVRLVDPINVVGLLLLTRSGKRSWPPSEVLTAVGLICALAGGFAKADIEMAGPMAAVGLLIVSYVVSGKSVDMYIERAGDITWEKDAEVTGNSPRLDVALDESGDFSLVEDDGPPMREIILKVVLMTICGMNPIAIPFAAGAWYVYVKTGKRSGALWDVPAPKEVKKGETTDGVYRVMTRRLLGSTQVGVGVMQEGVFHTMWHVTKGSALRSGEGRLDPYWGDVKQDLVSYCGPWKLDAAWDGHSEVQLLAVPPGERARNIQTLPGIFKTKGGDIGAVALDYPAGTSGSPILDKCGRVIGLYGNGVVIKNGSYVSAITQGRREEETPVECFEPSMLKKKQLTVLDLHPGAGKTRRVLPEIVREAIKTRLRTVILAPTRVVAAEMEEALRGLPVRYMTTAVNVTHSGTEIVDLMCHATFTSRLLQPIRVPNYNLYIMDEAHFTDPSSIAARGYISTRVEMGEAAAIFMTATPPGTRDAFPDSNSPIMDTEVEVPERAWSSGFDWVTDHSGKTVWFVPSVRNGNEIAACLTKAGKRVIQLSRKTFETEFQKTKHQEWDFVVTTDISEMGANFKADRVIDSRRCLKPVILDGERVILAGPMPVTHASAAQRRGRIGRNPNKPGDEYLYGGGCAETDEDHAHWLEARMLLDNIYLQDGLIASLYRPEADKVAAIEGEFKLRTEQRKTFVELMKRGDLPVWLAYQVASAGITYTDRRWCFDGTTNNTIMEDSVPAEVWTRHGEKRVLKPRWMDARVCSDHAALKSFKEFAAGKRGAALGVMEALGTLPGHMTERFQEAIDNLAVLMRAETGSRPYKAAAAQMPETLETIMLLGLLGTVSLGIFFVLMRNKGIGKMGFGMVTLGASAWLMWLSEIEPARIACVLIVVFLLLVVLIPEPEKQRSPQDNQMAIIIMVAVGLLGLITANELGWLERTKSDLSHLMGRREEGATIGFSMDIDLRPASAWAIYAALTTFITPAVQHAVTTSYNNYSLMAMATQAGVLFGMGKGMPFYAWDFGVPLLMIGCYSQLTPLTLIVAIILLVAHYMYLIPGLQAAAARAAQKRTAAGIMKNPVVDGIVVTDIDTMTIDPQVEKKMGQVLLIAVAVSSAILSRTAWGWGEAGALITAATSTLWEGSPNKYWNSSTATSLCNIFRGSYLAGASLIYTVTRNAGLVKRRGGGTGETLGEKWKARLNQMSALEFYSYKKSGITEVCREEARRALKDGVATGGHAVSRGSAKLRWLVERGYLQPHGKVIDLGCGRGGWSYYAATIRKVQEVKGYTKGGPGHEEPVLVQSYGWNIVRLKSGVDVFHMAAEPCDTLLCDIGESSSSPEVEEARTLRVLSMVGDWLEKRPGAFCIKVLCPYTSTMMETLERLQRRYGGGLVRVPLSRNSTHEMYWVSGAKSNTIKSVSTTSQLLLGRMDGPRRPVKYEEDVNLGSGTRAVVSCAEAPNMKIIGNRIERIRSEHAETWFFDENHPYRTWAYHGSYEAPTQGSASSLVNGVVRLLSKPWDVVTGVTGIAMTDTTPYGQQRVFKEKVDTRVPDPQEGTRQVMSMVSSWLWKELGKHKRPRVCTKEEFINKVRSNAALGAIFEEEKEWKTAVEAVNDPRFWALVDKEREHHLRGECQSCVYNMMGKREKKQGEFGKAKGSRAIWYMWLGARFLEFEALGFLNEDHWMGRENSGGGVEGLGLQRLGYVLEEMSRIPGGRMYADDTAGWDTRISRFDLENEALITNQMEKGHRALALAIIKYTYQNKVVKVLRPAEKGKTVMDIISRQDQRGSGQVVTYALNTFTNLVVQLIRNMEAEEVLEMQDLWLLRRPEKVTNWLQSNGWDRLKRMAVSGDDCVVKPIDDRFAHALRFLNDMGKVRKDTQEWKPSTGWDNWEEVPFCSHHFNKLHLKDGRSIVVPCRHQDELIGRARVSPGAGWSIRETACLAKSYAQMWQLLYFHRRDLRLMANAICSSVPVDWVPTGRTTWSIHGKGEWMTTEDMLVVWNRVWIEENDHMEDKTPVTKWTDIPYLGKREDLWCGSLIGHRPRTTWAENIKNTVNMVRRIIGDEEKYMDYLSTQVRYLGEEGSTPGVL

>tr|A0A142EHG9|A0A142EHG9_ZIKV Polyprotein OSZika virus (strain Mr 766) PE4 SV1

MKNPKKKSGGFRIVNMLKRGVARVSPFGGLKRLPAGLLLGHGPIRMVLAILAFLRFTAIKPSLGLINRWGSVGKKEAMEIIKKFKKDLAAMLRIINARKEKKRRGADTSVGIVGLLLTTAMAAEVTRRGSAYYMYLDRNDAGEAISFPTTLGMNKCYIQIMDLGHMCDATMSYECPMLDEGVEPDDVDCWCNTTSTWVVYGTCHHKKGEARRSRRAVTLPSHSTRKLQTRSQTWLESREYTKHLIRVENWIFRNPGFALAAAAIAWLLGSSTSQKVIYLVMILLIAPAYSIRCIGVSNRDFVEGMSGGTWVDVVLEHGGCVTVMAQDKPTVDIELVTTTVSNMAEVRSYCYEASISDMASDSRCPTQGEAYLDKQSDTQYVCKRTLVDRGWGNGCGLFGKGSLVTCAKFACSKKMTGKSIQPENLEYRIMLSVHGSQHSGMIVNDTGHETDENRAKVEITPNSPRAEATLGGFGSLGLDCEPRTGLDFSDLYYLTMNNKHWLVHKEWFHDIPLPWHAGADTGTPHWNNKEALVEFKDAHAKRQTVVVLGSQEGAVHTALAGALEAEMDGAKGRLSSGHLKCRLKMDKLRLKGVSYSLCTAAFTFTKIPAETLHGTVTVEVQYAGADGPCKVPAQMAVDMQTLTPVGRLITANPVITESTENSKMMLELDPPFGDSYIVIGVGEKKITHHWHRSGSTIGKAFEATVRGAKRMAVLGDTAWDFGSVGGALNSLGKGIHQIFGAAFKSLFGGMSWFSQILIGTLLMWLGLNTKNGSISLMCLALGGVLIFLSTAVSADVGCSVDFSKKETRCGTGVFVYNDVEAWRDRYKYHPDSPRRLAAAVKQAWEDGICGISSVSRMENIMWRSVEGELNAILEENGVQLTVVVGSVKNPMWRGPQRLPVPVNELPHGWKAWGKSYFVRAAKTNNSFVVDGDTLKECPLKHRAWNSFLVEDHGFGVFHTSVWLKVREDYSLECDPAVIGTAVKGKEAVHSDLGYWIESEKNDTWRLKRAHLIEMKTCEWPKSHTLWTDGIEESDLIIPKSLAGPLSHHNTREGYRTQMKGPWHSEELEIRFEECPGTKVHVEETCGTRGPSLRSTTASGRVIEEWCCRECTMPPLSFRAKDGCWYGMEIRPRKEPESNLVRSVVTAGSTDHMDHFSLGVLVILLMVQEGLKKRMTTKIIISTSMAVLVAMILGGFSMSDLAKLAILMGATFAEMNTGGDVAHLALIAAFKVRPALLVSFIFRANWTPRESMLLALASCLLQTAISALEGDLMVLINGFALAWLAIRAMVVPRTDNITLAILAALTPLARGTLLVAWRAGLATCGGFMLLSLKGKGSVKKNLPFVMALGLTAVRLVDPINVVGLLLLTRSGKRSWPPSEVLTAVGLICALAGGFAKADIEMAGPMAAVGLLIVSYVVSGKSVDMYIERAGDITWEKDAEVTGNSPRLDVALDESGDFSLVEDDGPPMREIILKVVLMTICGMNPIAIPFAAGAWYVYVKTGKRSGALWDVPAPKEVKKGETTDGVYRVMTRRLLGSTQVGVGVMQEGVFHTMWHVTKGSALRSGEGRLDPYWGDVKQDLVSYCGPWKLDAAWDGHSEVQLLAVPPGERARNIQTLPGIFKTKDGDIGAVALDYPAGTSGSPILDKCGRVIGLYGNGVVIKNGSYVSAITQGRREEETPVECFEPSMLKKKQLTVLDLHPGAGKTRRVLPEIVREAIKTRLRTVILAPTRVVAAEMEEALRGLPVRYMTTAVNVTHSGTEIVDLMCHATFTSRLLQPIRVPNYNLYIMDEAHFTDPSSIAARGYISTRVEMGEAAAIFMTATPPGTRDAFPDSNSPIMDTEVEVPERAWSSGFDWVTDHSGKTVWFVPSVRNGNEIAACLTKAGKRVIQLSRKTFETEFQKTKHQEWDFVVTTDISEMGANFKADRVIDSRRCLKPVILDGERVILAGPMPVTHASAAQRRGRIGRNPNKPGDEYLYGGGCAETDEDHAHWLEARMLLDNIYLQDGLIASLYRPEADKVAAIEGEFKLRTEQRKTFVELMKRGDLPVWLAYQVASAGITYTDRRWCFDGTTNNTIMEDSVPAEVWTRHGEKRVLKPRWMDARVCSDHAALKSFKEFAAGKRGATFGVMEALGTLPGHMTERFQEAIDNLAVLMRAETGSRPYKAAAAQLPETLETIMLLGLLGTVSLGIFFVLMRNKGIGKMGFGMVTLGASAWLMWLSEIEPARIACVLIVVFLLLVVLIPEPEKQRSPQDNQMAIIIMVAVGLLGLITANELGWLERTKSDLSHLMGRREEGATIGFSMDIDLRPASAWAIYAALTTFITPAVQHAVTTSYNNYSLMAMATQAGVLFGMGKGMPFYAWDFGVPLLMIGCYSQLTPLTLIVAIILLVAHYMYLIPGLQAAAARAAQKRTAAGIMKNPVVDGIVVTDIDTMTIDPQVEKKMGQVLLIAVAVSSAILSRTAWGWGEAGALITAATSTLWEGSPNKYWNSSTATSLCNIFRGSYLAGASLIYTVTRNAGLVKRRGGGTGETLGEKWKARLNQMSALEFYSYKKSGITEVCREEARRALKDGVATGGHAVSRGSAKLRWLVERGYLQPYGKVIDLGCGRGGWSYYAATIRKVQEVKGYTKGGPGHEEPVLVQSYGWNIVRLKSGVDVFHMAAEPCDTLLCDIGESSSSPEVEEARTLRVLSMAGDWLEKRPGAFCIKVLCPYTSTMMETLERLQRRYGGGLVRVPLSRNSTHEMYWVSGAKSNTIKSVSTTSQLLLGRMDGPRRPVKYEEDVNLGSGTRAVVSCAEAPNMKIIGNRIERIRSEHAETWFFDENHPYRTWAYHGSYEAPTQGSASSLINGVVRLLSKPWDVVTGVTGIAMTDTTPYGQQRVFKEKVDTRVPDPQEGTRQVMSMVSSWLWKELGKHKRPRVCTKEEFINKVRSNAALGAIFEEEKEWKTAVEAVNDPRFWALVDKEREHHLRGECQSCVYNMMGKREKKQGEFGKAKGSRAIWYMWLGARFLEFEALGFLNEDHWMGRENSGGGVEGLGLQRLGYVLEEMSRIPGGRMYADDTAGWDTRISRFDLENEALITNQMEKGHRALALAIIKYTYQNKVVKVLRPAEKGKTVMDIISRQDQRGSGQVVTYALNTFTNLVVQLIRNMEAEEVLEMQDLWLLRRSEKVTNWLQSNGWDRLKRMAVSGDDCVVKPIDDRFAHALRFLNDMGKVRKDTQEWKPSTGWDNWEEVPFCSHHFNKLHLKDGRSIVVPCRHQDELIGRARVSPGAGWSIRETACLAKSYAQMWQLLYFHRRDLRLMANAICSSVPVDWVPTGRTTWSIHGKGEWMTTEDMLVVWNRVWIEENDHMEDKTPVTKWTDIPYLGKREDLWCGSLIGHRPRTTWAENIKNTVNMVRRIIGDEEKYMDYLSTQVRYLGEEGSTPGVL

>tr|A0A142DS37|A0A142DS37_ZIKV Polyprotein OSZika virus (strain Mr 766) PE4 SV1

MKNPKKKSGGFRIVNMLKRGVARVSPFGGLKRLPAGLLLGHGPIRMVLAILAFLRFTAIKPSLGLINRWGSVGKKEAMEIIKKFKKDLAAMLRIINARKEKKRRGAETSVGIVGLLLTTAMAAEVTRRGSAYYMYLDRNDAGEAISFPTTLGMNKCYIQIMDLGHMCDATMSYECPMLDEGVEPDDVDCWCNTTSTWVVYGTCHHKKGEARRSRRAVTLPSHSTRKLQTRSQTWLESREYTKHLIRVENWIFRNPGFALAAAAIAWLLGSSTSQKVIYLVMILLIALAYSIRCIGVSNRDFVEGMSGGTWVDVVLEHGGCVTVMAQDKPTVDIELVTTTVSNMAEVRSYCYEASISDMASDSRCPTQGEAYLDKQSDTQYVCKRTLVDRGWGNGCGLFGKGSLVTCAKFACSKKMTGKSIQPENLEYRIMLSVHGSQHSGMIVNDTGHETDENRAKVEITPNSPRAEATLGGFGSLGLDCEPRTGLDFSDLYYLTMNNKHWLVHKEWFHDIPLPWHAGADTGTPHWNNKEALVEFKDAHAKRQTVVVLGSQEGAVHTALAGALEAEMDGAKGRLSSGHLKCRLKMDKLRLKGVSYSLCTAAFTFTKIPAETLHGTVTVEVQYAGTDGPCKVPAQMAVDMQTLTPVGRLITANPVITESTENSKMMLELDPPFGDSYIVIGVGEKKITHHWHRSGSTIGKAFEATVRGAKRMAVLGDTAWDFGSVGGALNSLGKGIHQIFGAAFKSLFGGMSWFSQILIGTLLMWLGLNTKNGSISLMCLALGGVLIFLSTAVSADVGCSVDFSKKETRCGTGVFVYNDVEAWRDRYKYHPDSPRRLAAAVKQAWEDGICGISSVSRMENIMWRSVEGELNAILEENGVQLTVVVGSVKNPMWRGPQRLPVPVNELPHGWKAWGKSYFVRAAKTNNSFVVDGDTLKECPLKHRAWNSFLVEDHGFGVFHTSVWLKVREDYSLECDPAVIGTAVKGKEAVHSDLGYWIESEKNDTWRLKRAHLIEMKTCEWPKSHTLWTDGIEESDLIIPKSLAGPLSHHNTREGYRTQMKGPWHSEELEIRFEECPGTKVHVEETCGTRGPSLRSTTASGRVIEEWCCRECTMPPLSFWAKDGCWYGMEIRPRKEPESNLVRSMVTAGSTDHMDHFSLGVLVILLMVQEGLKKRMTTKIIISTSMAVLVAMILGGFSMSDLAKLAILMGATFAEMNTGGDVAHLALTAAFKVRPALLVSFIFRANWTPRESMLLALASCLLQTAISALEGDLMVLINGFALAWLAIRAMVVPRTDNITLAILAALTPLARGTLLVAWRAGLATCGGFMLLSLKGKGSVKKNLPFVMALGLTAVRLVDPINVVGLLLLTRSGKRSWPPSEVLTAVGLICALAGGFAKADIEMAGPMAAVGLLIVSYVVSGKSVDMYIERAGDITWEKDAEVTGNSPRLDVALDESGDFSLVEDDGPPMREIILKVVLMTICGMNPIAIPFAAGAWYVYVKTGKRSGALWDVPAPKEVKKGETTDGVYRVMTRRLLGSTQVGVGVMQEGVFHTMWHVTKGSALRSGEGRLDPYWGDVKQDLVSYCGPWKLDAAWDGHSEVQLLAVPPGERERNIQTLPGIFKTKDGDIGAVALDYPAGTSGSPILDKCGRVIGLYGNGVVIKNGSYVSAITQGRREEETPVECFEPSMLKKKQLTVLDLHPGAGKTRRVLPEIVREAIKTRLRTVILAPTRVVAAEMEEALRGLPVRYMTTAVNVTHSGTEIVDLMCHATFTSRLLQPIRVPNYNLYIMDEAHFTDPSSIAARGYISTRVEMGEAAAIFMTATPPGTRDAFPDSNSPIMDTEVEVPERAWSSGFDWVTDHSGKTVWFVPSVRNGNEIAACLTKAGKRVIQLSRKTFETEFQKTKHQEWDFVVTTDISEMGANFKADRVIDSRRCLKPVILDGERVILAGPMPVTHASAAQRRGRIGRNPNKPGDEYLYGGGCAETDEDHAHWLEARMLLDNIYLQDGLIASLYRPEADKVAAIEGEFKLRTEQRKTFVELMKRGDLPVWLAYQVASAGITYTDRRWCFDGTTNNTIMEDSVPAEVWTRHGEKRVLKPRWMDARVCSDHAALKSFKEFAAGKRGAAFGVMEALGTLPGHMTERFQEAIDNLAVLMRAETGSRPYKAAAAQLPETLETIMLLGLLGTVSLGIFFVLMRNKGIGKMGFGMVTLGASAWLMWLSEIEPARIACVLIVVFLLLVVLIPEPEKQRSPQDNQMAIIIMVAVGLLGLITANELGWLERTKSDLSHLMGRREEGATIGFSMDIDLRPASAWAIYAALTTFITPAVQHAVTTSYNNYSLMAMATQAGVLFGMGKGMPFYAWDFGVPLLMIGCYSQLTPLTLIVAIILLVAHYMYLIPGLQAAAARAAQKRTAAGIMKNPVVDGIVVTDIDTMTIDPQVEKKMGQVLLIAVAVSSAILSRTAWGWGEAGALITAATSTLWEGSPNKYWNSSTATSLCNIFRGSYLAGASLIYTVTRNAGLVKRRGGGTGETLGEKWKARLNQMSALEFYSYKKSGITEVCREEARRALKDGVATGGHAVSRGSAKLRWLVERGYLQPYGKVIDLGCGRGGWSYYAATIRKVQEVKGYTKGGPGHEEPVLVQSYGWNIVRLKSGVDVFHMAAEPCDTLLCDIGESSSSPEVEEARTLRVLSMVGDWLEKRPGAFCIKVLCPYTSTMMETLERLQRRYGGGLVRVPLSRNSTHEMYWVSGAKSNTIKSVSTTSQLLLGRMDGPRRPVKYEEDVNLGSGTRAVVSCAEAPNMKIIGNRIERIRSEQAETWFFDENHPYRTWAYHGSYEAPTQGSASSLINGVVRLLSKPWDVVTGVTGIAMTDTTPYGQQRVFKEKVDTRVPDPQEGTRQVMSMVSSWLWKELGKHKRPRVCTKEEFINKVRSNAALGAIFEEEKEWKTAVEAVNDPRFWALVDKEREHHLRGECQSCVYNMMGKREKKQGEFGKAKGSRAIWYMWLGARFLEFEALGFLNEDHWMGRENSGGGVEVLGLQRLGYVLEEMSRIPGGRMYADDTAGWDTRISRFDLENEALITNQMEKGHRALALAIIKYTYQNKVVKVLRPAEKGKTVMDIISRQDQRGSGQVVTYALNTFTNLVVQLIRNMEAEEVLEMQDLWLLRRSEKVTNWLQSNGWDRLKRMAVSGDDCVVKPIDDRFAHALRFLNDMGKVRKDTQEWKPSTGWDNWEEVPFCSHHFNKLHLKDGRSIVVPCRHQDELIGRARVSPGAGWSIRETACLAKSYAQMWQLLYFHRRDLRLMANAICSSVPVDWVPTGRTTWSIHGKGEWMTTEDMLVVWNRVWIEENDHMEDKTPVAKWTDIPYLGKREDLWCGSLIGHRPRTTWAENIKNTVNMVRRIIGDEEKYMDYLSTQVRYLGEEGSTPGVL

>tr|A0A141VLG5|A0A141VLG5_ZIKV Polyprotein OSZika virus (strain Mr 766) PE4 SV1

MKNPKKKSGGFRIVNMLKRGVARVSPFGGLKRLPAGLLLGHGPIRMVLAILAFLRFTAIKPSLGLINRWGSVGKKEAMEIIKKFKKDLAAMLRIINARKEKKRRGAETSVGIVGLLLTTAMAAEVTRRGSAYYMYLDRNDAGEAISFPTTLGMNKCYIQIMDLGHMCDATMSYECPMLDEGVEPDDVDCWCNTTSTWVVYGTCHHKKGEARRSRRAVTLPSHSTRKLQTRSQTWLESREYTKHLIRVENWIFRNPGFALAAAAIAWLLGSSTSQKVIYLVMILLIAPAYSIRCIGVSNRDFVEGMSGGTWVDVVLEHGGCVTVMAQDKPTVDIELVTTTVSNMAEVRSYCYEASISDMASDSRCPTQGEAYLDKQSDTQYVCKRTLVDRGWGNGCGLFGKGSLVTCAKFACSKKMTGKSIQPENLEYRIMLSVHGSQHSGMIVNDTGHETDENRAKVEITPNSPRAEATLGGFGSLGLDCEPRTGLDFSDLYYLTMNNKHWLVHKEWFHDIPLPWHAGADTGTPHWNNKEALVEFKDAHAKRQTVVVLGSQEGAVHTALAGALEAEMDGAKGRLSSGHLKCRLKMDKLRLKGVSYSLCTAAFTFTKIPAETLHGTVTVEVQYAGTDGPCKVPAQMAVDMQTLTPVGRLITANPVITESTENSKMMLELDPPFGDSYIVIGVGEKKITHHWHRSGSTIGKAFEATVRGAKRMAVLGDTAWDFGSVGGALNSLGKGIHQIFGAAFKSLFGGMSWFSQILIGTLLMWLGLNTKNGSISLMCLALGGVLIFLSTAVSADVGCSVDFSKKETRCGTGVFVYNDVEAWRDRYKYHPDSPRRLAAAVKQAWEDGICGISSVSRMENIMWRSVEGELNAILEENGVQLTVVVGSVKNPMWRGPQRLPVPVNELPHGWKAWGKSYFVRAAKTNNSFVVDGDTLKECPLKHRAWNSFLVEDHGFGVFHTSVWLKVREDYSLECDPAVIGTAVKGKEAVHSDLGYWIESEKNDTWRLKRAHLIEMKTCEWPKSHTLWTDGIEESDLIIPKSLAGPLSHHNTREGYRTQMKGPWHSEELEIRFEECPGTKVHVEETCGTRGPSLRSTTASGRVIEEWCCRECTMPPLSFRAKDGCWYGMEIRPRKEPESNLVRSMVTAGSTDHMDHFSLGVLVILLMVQEGLKKRMTTKIIISTSMAVLVAMILGGFSMSDLAKLAILMGATFAEMNTGGDVAHLALIAAFKVRPALLVSFIFRANWTPRESMLLALASCLLQTAISALEGDLMVLINGFALAWLAVRAMVVPRTDNITLAILAALTPLARGTLLVAWRAGLATCGGFMLLSLKGKGSVKKNLPFVMALGLTAVRLVDPINVVGLLLLTRSGKRSWPPSEVLTAVGLICALAGGFAKADIEMAGPMAAVGLLIVSYVVSGKSVDMYIERAGDITWEKDAEVTGNSPRLDVALDESGDFSLVEDDGPPMREIILKVVLMTICGMNPIAIPFAAGAWYVYVKTGKRSGALWDVPAPKEVKKGETTDGVYRVMTRRLLGSTQVGVGVMQEGVFHTMWHVTKGSALRSGEGRLDPYWGDVKQDLVSYCGPWKLDAAWDGHSEVQLLAVPPGERARNIQTLPGIFKTKDGDIGAVALDYPAGTSGSPILDKCGRVIGLYGNGVVIKNGSYVSAITQGRREEETPVECFEPSMLKKKQLTVLDLHPGAGKTRRVLPEIVREAIKTRLRTVILAPTRVVAAEMEEALRGLPVRYMTTAVNVTHSGTEIVDLMCHATFTSRLLQPIRVPNYNLYIMDEAHFTDPSSIAARGYISTRVEMGEAAAIFMTATPPGTRDAFPDSNSPIMDTEVEVPERAWSSGFDWVTDHSGKTVWFVPSVRNGNEIAACLTKAGKRVIQLSRKTFETEFQKTKHQEWDFVVTTDISEMGANFKADRVIDSRRCLKPVILDGERVILAGPMPVTHASAAQRRGRIGRNPNKPGDEYLYGGGCAETDEDHAHWLEARMLLDNIYLQDGLIASLYRPEADKVAAIEGEFKLRTEQRKTFVELMKRGDLPVWLAYQVASAGITYTDRRWCFDGTTNNTIMEDSVPAEVWTRHGEKRVLKPRWMDARVCSDHAALKSFKEFAAGKRGAAFGVMEALGTLPGHMTERFQEAIDNLAVLMRAETGSRPYKAAAAQLPETLETIMLLGLLGTVSLGIFFVLMRNKGIGKMGFGMVTLGASAWLMWLSEIEPARIACVLIVVFLLLVVLIPEPEKQRSPQDNQMAIIIMVAVGLLGLITANELGWLERTKSDLSHLMGRREEGATIGFSMDIDLRPASAWAIYAALTTFITPAVQHAVTTSYNNYSLMAMATQAGVLFGMGKGMPFYAWDFGVPLLMIGCYSQLTPLTLIVAIILLVAHYMYLIPGLQAAAARAAQKRTAAGIMKNPVVDGIVVTDIDTMTIDPQVEKKMGQVLLIAVAVSSAILSRTAWGWGEAGALITAATSTLWEGSPNKYWNSSTATSLCNIFRGSYLAGASLIYTVTRNAGLVKRRGGGTGETLGEKWKARLNQMSALEFYSYKKSGITEVCREEARRALKDGVATGGHAVSRGSAKLRWLVERGYLQPYGKVIDLGCGRGGWSYYAATIRKVQEVKGYTKGGPGHEEPVLVQSYGWNIVRLKSGVDVFHMAAEPCDTLLCDIGESSSSPEVEEARTLRVLSMVGDWLEKRPGAFCIKVLCPYTSTMMETLERLQRRYGGGLVRVPLSRNSTHEMYWVSGAKSNIIKSVSTTSQLLLGRMDGPRRPVKYEEDVNLGSGTRAVASCAEAPNMKIIGNRIERIRSEHAETWFFDENHPYRTWAYHGSYEAPTQGSASSLINGVVRLLSKPWDVVTGVTGIAMTDTTPYGQQRVFKEKVDTRVPDPQEGTRQVMSMVSSWLWKELGKHKRPRVCTKEEFINKVRSNAALGAIFEEEKEWKTAVEAVNDPRFWALVDKEREHHLRGECQSCVYNMMGKREKKQGEFGKAKGSRAIWYMWLGARFLEFEALGFLNEDHWMGRENSGGGVEGLGLQRLGYVLEEMSRIPGGRMYADDTAGWDTRISRFDLENEALITNQMEKGHRALALAIIKYTYQNKVVKVLRPAEKGKTVMDIISRQDQRGSGQVVTYALNTFTNLVVQLIRNMEAEEVLEMQDLWLLRRSEKVTNWLQSNGWDRLKRMAVSGDDCVVKPIDDRFAHALRFLNDMGKVRKDTQEWKPSTGWDNWEEVPFCSHHFNKLHLKDGRSIVVPCRHQDELIGRARVSPGAGWSIRETACLAKSYAQMWQLLYFHRRDLRLMANAICSSVPVDWVPTGRTTWSIHGKGEWMTTEDMLVVWNRVWIEENDHMEDKTPVTKWTDIPYLGKREDLWCGSLIGHRPRTTWAENIKNTVNMVRRIIGDEEKYMDYLSTQVRYLGEEGSTPGVL

>tr|A0A142K469|A0A142K469_ZIKV Polyprotein OSZika virus (strain Mr 766) PE4 SV1

MKNPKKKSGGFRIVNMLKRGVARVSPFGGLKRLPAGLLLGHGPIRMVLAILAFLRFTAIKPSLGLINRWGSVGKKEAMEIIKKFKKDLAAMLRIINARKEKKRRGADTSVGIVGLLLTTAMAAEVTRRGSAYYMYLDRNDAGEAISFPTTLGMNKCYIQIMDLGHMCDATMSYECPMLDEGVEPDDVDCWCNTTSTWVVYGTCHHKKGEARRSRRAVTLPSHSTRKLQTRSQTWLESREYTKHLIRVENWIFRNPGFALAAAAIAWLLGSSTSQKVIYLVMILLIAPAYSIRCIGVSNRDFVEGMSGGTWVDVVLEHGGCVTVMAQDKPTVDIELVTTTVSNMAEVRSYCYEASISDMASDSRCPTQGEAYLDKQSDTQYVCKRTLVDRGWGNGCGLFGKGSLVTCAKFACSKKMTGKSIQPENLEYRIMLSVHGSQHSGMIVNDTGHETDENRAKVEITPNSPRAEATLGGFGSLGLDCEPRTGLDFSDLYYLTMNNKHWLVHKEWFHDIPLPWHAGADTGTPHWNNKEALVEFKDAHAKRQTVVVLGSQEGAVHTALAGALEAEMDGAKGRLSSGHLKCRLKMDKLRLKGVSYSLCTAAFTFTKIPAETLHGTVTVEVQYAGTDGPCKVPAQMAVDMQTLTPVGRLITANPVITESTENSKMMLELDPPFGDSYIVIGVGEKKITHHWHRSGSTIGKAFEATVRGAKRMAVLGDTAWDFGSVGGALNSLGKGIHQIFGAAFKSLFGGMSWFSQILIGTLLMWLGLNTKNGSISLMCLALGGVLIFLSTAVSADVGCSVDFSKKETRCGTGVFVYNDVEAWRDRYKYHPDSPRRLAAAVKQAWEDGICGISSVSRMENIMWRSVEGELNAILEENGVQLTVVVGSVKNPMWRGPQRLPVPVNELPHGWKAWGKSYFVRAAKTNNSFVVDGDTLKECPLKHRAWNSFLVEDHGFGVFHTSVWLKVREDYSLECDPAVIGTAVKGKEAVHSDLGYWIESEKNDTWRLKRAHLIEMKTCEWPKSHTLWTDGIEESDLIIPKSLAGPLSHHNTREGYRTQMEGPWHSEELEIRFEECPGTKVHVEETCGTRGPSLRSTTASGRVIEEWCCRECTMPPLSFRAKDGCWYGMEIRPRKEPESNLVRSMVTAGSTDHMDHFSLGVLVILLMVQEGLKKRMTTKIIISTSMAVLVAMILGGFSMSDLAKLAILMGATFAEMNTGGDVAHLALIAAFKVRPALLVSFIFRANWTPRESMLLALASCLLQTAISALEGDLMVLINGFALAWLAIRAMVVPRTDNITLAILAALTPLARGTLLVAWRAGLATCGGFMLLSLKGKGSVKKNLPFVMALGLTAVRLVDPINVVGLLLLTRSGKRSWPPSEVLTAVGLICALAGGFAKADIEMAGPMAAVGLLIVSYVVSGKSVDMYIERAGDITWEKDAEVTGNSPRLDVALDESGDFSLVEDDGPPMREIILKVVLMTICGMNPIAIPFAAGAWYVYVKTGKRSGALWDVPAPKEVKKGETTDGVYRVMTRRLLGSTQVGVGVMQEGVFHTMWHVTKGSALRSGEGRLDPYWGDVKQDLVSYCGPWKLDAAWDGHSEVQLLAVPPGERARNIQTLPGIFKTKDGDIGAVALDYPAGTSGSPILDKCGRVIGLYGNGVVIKNGSYVSAITQGRREEETPVDCFEPSMLKKKQLTVLDLHPGAGKTRRVLPEIVREAIKTRLRTVILAPTRVVAAEMEEALRGLPVRYMTTAVNVTHSGTEIVDLMCHATFTSRLLQPIRVPNYNLYIMDEAHFTDPSSIAARGYISTRVEMGEAAAIFMTATPPGTRDAFPDSNSPIMDTEVEVPERAWSSGFDWVTDHSGKTVWFVPSVRNGNEIAACLTKAGKRVIQLSRKTFETEFQKTKHQEWDFVVTTDISEMGANFKADRVIDSRRCLKPVILDGERVILAGPMPVTHASAAQRRGRIGRNPNKPGDEYLYGGGCAETDEDHAHWLEARMLLDNIYLQDGLIASLYRPEADKVAAIEGEFKLRTEQRKTFVELMKRGDLPVWLAYQVASAGITYTDRRWCFDGTTNNTIMEDSVPAEVWTRHGEKRVLKPRWMDARVCSDHAALKSFKEFAAGKRGAAFGVMEALGTLPGHMTERFQEAIDNLAVLMRAETGSRPYKAAAAQLPETLETIMLLGLLGTVSLGIFFVLMRNKGIGKMGFGMVTLGASAWLMWLSEIEPARIACVLIVVFLLLVVLIPEPEKQRSPQDNQMAIIIMVAVGLLGLITANELGWLERTKSDLSHLMGRREEGATIGFSMDIDLRPASAWAIYAALTTFITPAVQHAVTTSYNNYSLMAMATQAGVLFGMGKGMPFYAWDFGVPLLMIGCYSQLTPLTLIVAIILLVAHYMYLIPGLQAAAARAAQKRTAAGIMKNPVVDGIVVTDIDTMTIDPQVEKKMGQVLLIAVAVSSAILSRTAWGWGEAGALITAATSTLWEGSPNKYWNSSTATSLCNIFRGSYLAGASLIYTVTRNAGLVKRRGGGTGETLGEKWKARLNQMSALEFYSYKKSGITEVCREEARRALKDGVATGGHAVSRGSAKLRWLVERGYLQPYGKVIDLGCGRGGWSYYAATIRKVQEVKGYTKGGPGHEEPVLVQSYGWNIVRLKSGVDVFHMAAEPCDTLLCDIGESSSSPEVEEARTLRVLSMVGDWLEKRPGAFCIKVLCPYTSTMMETLERLQRRYGGGLVRVPLSRNSTHEMYWVSGAKSNTIKSVSTTSQLLLGRMDGPRRPVKYEEDVNLGSGTRAVVSCAEAPNMKIIGNRIERIRSEHAETWFFDENHPYRTWAYHGSYEAPTQGSASSLINGVVRLLSKPWDVVTGVTGIAMTDTTPYGQQRVFKEKVDTRVPDPQEGTRQVMSMVSSWLWKELGKHKRPRVCTKEEFINKVRSNAALGAIFEEEKEWKTAVEAVNDPRFWALVDKEREHHLRGECQSCVYNMMGKREKKQGEFGKAKGSRAIWYMWLGARFLEFEALGFLNEDHWMGRENSGGGVEGLGLQRLGYVLEEMSRIPGGRMYADDTAGWDTRISRFDLENEALITNQMEKGHRALALAIIKYTYQNKVVKVLRPAEKGKTVMDIISRQDQRGSGQVVTYALNTFTNLVVQLIRNMEAEEVLEMQDLWLLRRSEKVTNWLQSNGWDRLKRMAVSGDDCVVKPIDDRFAHALRFLNDMGKVRKDTQEWKPSTGWDNWEEVPFCSHHFNKLHLKDGRSIVVPCRHQDELIGRARVSPGAGWSIRETACLAKSYAQMWQLLYFHRRDLRLMANAICSSVPVDWVPTGRTTWSIHGKGEWMTTEDMLVVWNRVWIEENDHMEDKTPVTKWTDIPYLGKREDLWCGSLIGHRPRTTWAENIKNTVNMVRRIIGDEEKYMDYLSTQVRYLGEEGSTPGVL

>tr|A0A142DS38|A0A142DS38_ZIKV Polyprotein OSZika virus (strain Mr 766) PE4 SV1

MKNPKKKSGGFRIVNMLKRGVARVSPFGGLKRLPAGLLLGHGPIRMVLAILAFLRFTAIKPSLGLINRWGSVGKKEAMEIIKKFKKDLAAMLRIINARKEKKRRGAETSVGIVGLLLTTAMAAEVTRRGSAYYMYLDRNDAGEAISFPTTLGMNKCYIQIMDLGHMCDATMSYECPMLDEGVEPDDVDCWCNTTSTWVVYGTCHHKKGEARRSRRAVTLPSHSTRKLQTRSQTWLESREYTKHLIRVENWIFRNPGFALAAAAIAWLLGSSTSQKVIYLVMILLIAPAYSIRCIGVSNRDFVEGMSGGTWVDVVLEHGGCVTVMAQDKPTVDIELVTTTVSNMAEVRSYCYEASISDMASDSRCPTQGEAYLDKQSDTQYVCKRTLVDRGWGNGCGLFGKGSLVTCAKFACSKKMTGKSIQPENLEYRIMLSVHGSQHSGMIVNDTGHETDENRAKVEITPNSPRAEATLGGFGSLGLDCEPRTGLDFSDLYYLTMNNKHWLVHKEWFHDIPLPWHAGADTGTPHWNNKEALVEFKDAHAKRQTVVVLGSQEGAVHTALAGALEAEMDGAKGRLSSGHLKCRLKMDKLRLKGVSYSLCTAAFTFTKIPAETLHGTVTVEVQYAGTDGPCKVPAQMAVDMQTLTPVGRLITANPVITESTENSKMMLELDPPFGDSYIVIGVGEKKITHHWHRSGSTIGKAFEATVRGAKRMAVLGDTAWDFGSVGGALNSLGKGIHQIFGAAFKSLFGGMSWFSQILIGTLLMWLGLNTKNGSISLMCLALGGVLIFLSTAVSADVGCSVDFSKKETRCGTGVFVYNDVEAWRDRYKYHPDSPRRLAAAVKQAWEDGICGISSVSRMENIMWRSVEGELNAILEENGVQLTVVVGSVKNPMWRGPQRLPVPVNELPHGWKAWGKSYFVRAAKTNNSFVVDGDTLKECPLKHRAWNSFLVEDHGFGVFHTSVWLKVREDYSLECDPAVIGTAVKGKEAVHSDLGYWIESEKNDTWRLKRAHLIEMKTCEWPKSHTLWTDGIEESDLIIPKSLAGPLSHHNTREGYRTQMKGPWHSEELEIRFEECPGTKVHVEETCGTRGPSLRSTTASGRVIEEWCCRECTMPPLSFWAKDGCWYGMEIRPRKEPESNLVRSMVTAGSTDHMDHFSLGVLVILLMVQEGLKKRMTTKIIISTSMAVLVAMILGGFSMSDLAKLAILMGATFAEMNTGGDVAHLALTAAFKVRPALLVSFIFRANWTPRESMLLALASCLLQTAISALEGDLMVLINGFALAWLAIRAMVVPRTDNITLAILAALTPLARGTLLVAWRAGLATCGGFMLLSLKGKGSVKKNLPFVMALGLTAVRLVDPINVVGLLLLTRSGKRSWPPSEVLTAVGLICALAGGFAKADIEMAGPMAAVGLLIVSYVVSGKSVDMYIERAGDITWEKDAEVTGNSPRLDVALDESGDFSLVEDDGPPMREIILKVVLMTICGMNPIAIPFAAGAWYVYVKTGKRSGALWDVPAPKEVKKGETTDGVYRVMTRRLLGSTQVGVGVMQEGVFHTMWHVTKGSALRSGEGRLDPYWGDVKQDLVSYCGPWKLDAAWDGHSEVQLLAVPPGERERNIQTLPGIFKTKDGDIGAVALDYPAGTSGSPILDKCGRVIGLYGNGVVIKNGSYVSAITQGRREEETPVECFEPSMLKKKQLTVLDLHPGAGKTRRVLPEIVREAIKTRLRTVILAPTRVVAAEMEEALRGLPVRYMTTAVNVTHSGTEIVDLMCHATFTSRLLQPIRVPNYNLYIMDEAHFTDPSSIAARGYISTRVEMGEAAAIFMTATPPGTRDAFPDSNSPIMDTEVEVPERAWSSGFDWVTDHSGKTVWFVPSVRNGNEIAACLTKAGKRVIQLSRKTFETEFQKTKHQEWDFVVTTDISEMGANFKADRVIDSRRCLKPVILDGERVILAGPMPVTHASAAQRRGRIGRNPNKPGDEYLYGGGCAETDEDHAHWLEARMLLDNIYLQDGLIASLYRPEADKVAAIEGEFKLRTEQRKTFVELMKRGDLPVWLAYQVASAGITYTDRRWCFDGTTNNTIMEDSVPAEVWTRHGEKRVLKPRWMDARVCSDHAALKSFKEFAAGKRGAAFGVMEALGTLPGHMTERFQEAIDNLAVLMRAETGSRPYKAAAAQLPETLETIMLLGLLGTVSLGIFFVLMRNKGIGKMGFGMVTLGASAWLMWLSEIEPARIACVLIVVFLLLVVLIPEPEKQRSPQDNQMAIIIMVAVGLLGLITANELGWLERTKSDLSHLMGRREEGATIGFSMDIDLRPASAWAIYAALTTFITPAVQHAVTTSYNNYSLMAMATQAGVLFGMGKGMPFYAWDFGVPLLMIGCYSQLTPLTLIVAIILLVAHYMYLIPGLQAAAARAAQKRTAAGIMKNPVVDGIVVTDIDTMTIDPQVEKKMGQVLLIAVAVSSAILSRTAWGWGEAGALITAATSTLWEGSPNKYWNSSTATSLCNIFRGSYLAGASLIYTVTRNAGLVKRRGGGTGETLGEKWKARLNQMSALEFYSYKKSGITEVCREEARRALKDGVATGGHAVSRGSAKLRWLVERGYLQPYGKVIDLGCGRGGWSYYAATIRKVQEVKGYTKGGPGHEEPVLVQSYGWNIVRLKSGVDVFHMAAEPCDTLLCDIGESSSSPEVEEARTLRVLSMVGDWLEKRPGAFCIKVLCPYTSTMMETLERLQRRYGGGLVRVPLSRNSTHEMYWVSGAKSNTIKSVSTTSQLLLGRMDGPRRPVKYEEDVNLGSGTRAVVSCAEAPNMKIIGNRIERIRSEQAETWFFDENHPYRTWAYHGSYEAPTQGSASSLINGVVRLLSKPWDVVTGVTGIAMTDTTPYGQQRVFKEKVDTRVPDPQEGTRQVMSMVSSWLWKELGKHKRPRVCTKEEFINKVRSNAALGAIFEEEKEWKTAVEAVNDPRFWALVDKEREHHLRGECQSCVYNMMGKREKKQGEFGKAKGSRAIWYMWLGARFLEFEALGFLNEDHWMGRENSGGGVEVLGLQRLGYVLEEMSRIPGGRMYADDTAGWDTRISRFDLENEALITNQMEKGHRALALAIIKYTYQNKVVKVLRPAEKGKTVMDIISRQDQRGSGQVVTYALNTFTNLVVQLIRNMEAEEVLEMQDLWLLRRSEKVTNWLQSNGWDRLKRMAVSGDDCVVKPIDDRFAHALRFLNDMGKVRKDTQEWKPSTGWDNWEEVPFCSHHFNKLHLKDGRSIVVPCRHQDELIGRARVSPGAGWSIRETACLAKSYAQMWQLLYFHRRDLRLMANAICSSVPVDWVPTGRTTWSIHGKGEWMTTEDMLVVWNRVWIEENDHMEDKTPVAKWTDIPYLGKREDLWCGSLIGHRPRTTWAENIKNTVNMVRRIIGDEEKYMDYLSTQVRYLGEEGSTPGVL

>tr|A0A142EHH1|A0A142EHH1_ZIKV Polyprotein OSZika virus (strain Mr 766) PE4 SV1

MKNPKKKSGGFRIVNMLKRGVARVSPFGGLKRLPAGLLLGHGPIRMVLAILAFLRFTAIKPSLGLINRWGSVGKKEAMEIIKKFKKDLAAMLRIINARKEKKRRGADTSVGIVGLLLTTAMAAEVTRRGSAYYMYLDRNDAGEAISFPTTLGMNKCYIQIMDLGHMCDATMSYECPMLDEGVEPDDVDCWCNTTSTWVVYGTCHHKKGEARRSRRAVTLPSHSTRKLQTRSQTWLESREYTKHLIRVENWILRNPGFALAAAAIAWLLGSSTSQKVIYLVMILLIAPAYSIRCIGVSNRDFVEGMSGGTWVDVVLEHGGCVTVMAQDKPTVDIELVTTTVSNMAEVRSYCYEASISDMASDSRCPTQGEAYLDKQSDTQYVCKRTLVDRGWGNGCGLFGKGSLVTCAKFACSKKMTGKSIQPENLEYRIMLSVHGSQHSGMIVNDTGHETDENRAKVEITPNSPRAEATLGGFGSLGLDCEPRTGLDFSDLYYLTMNNKHWLVHKEWFHDIPLPWHAGADTGTPHWNNKEALVEFKDAHAKRQTVVVLGSQEGAVHTALAGALEAEMDGAKGRLSSGHLKCRLKMDKLRLKGVSYSLCTAAFTFTKIPAETLHGTVTVEVQYAGTDGPCKVPAQMAVDMQTLTPVGRLITANPVITESTENSKMMLELDPPFGDSYIVIGIGEKKITHHWHRSGSTIGKAFEATVRGAKRMAVLGDTAWDFGSVGGALNSLGKGIHQIFGAAFKSLFGGMSWFSQILIGTLLMWLGLNTKNGSISLMCLALGGVLIFLSTAVSADVGCSVDFSKKETRCGTGVFVYNDVEAWRDRYKYHPDSPRRLAAAVKQAWEDGICGISSVSRMENIMWRSVEGELNAILEENGVQLTVVVGSVKNPMWRAPQRLPVPVNELPHGWKAWGKSYFVRAAKTNNSFVVDGDTLKECPLKHRAWNSFLVEDHGFGVFHTSVWLKVREDYSLECDPAVIGTAVKGKEAVHSDLGYWIESEKNDTWRLKRAHLIEMKTCEWPQSHTLWTDGIEESDLIIPKSLAGPLSHHNTREGYRTQMKGPWHSEELEIRFEECPGTKVHVEETCGTRGPSLRSTTASGRVIEEWCCRECTMPPLSFRAKDGCWYGMEIRPRKEPESNLVRSMVTAGSTDHMDHFSLGVLVILLMVQEGLKKRMTTKIIISTSMAVLVAMILGGFSMSDLAKLAILMGATFAEMNTGGDVAHLALIAAFKVRPALLVSFIFRANWTPRESMLLALASCLLQTAISALEGDLMVLINGFALAWLAIRAMVVPRTDNITLAILAALTPLARGTLLVAWRAGLATCGGFMLLSLKGKGSVKKNLPFVMALGLTAVRLVDPINVVGLLLLTRSGKRSWPPSEVLTAVGLICALAGGFAKADIEMAGPVAAVGLLIVSYVVSGKSVDMYIERAGDITWEKDAEVTGNSPRLDVALDESGDFSLVEDDGPPMREIILKVVLMTICGMNPIAIPFAAGAWYVYVKTGKRSGALWDVPAPKEVKKGETTDGVYRVMTRRLLGSTQVGVGVMQEGVFHTMWHVTKGSALRSGEGRLDPYWGDVKQDLVSYCGPWKLDAAWDGHSEVQLLAVPPGERARNIQTLPGIFKTKDGDIGAVALDYPAGTSGSPILDKCGRVIGLYGNGVVIKNGSYVSAITQGRREEETPVECFEPSMLKKKQLTVLDLHPGAGKTRRVLPEIVREAIKTRLRTVILAPTRVVAAEMEEALRGLPVRYMTTAVNVTHSGTEIVDLMCHATFTSRLLQPIRVPNYNLYIMDEAHFTDPSSIAARGYISTRVEMGEAAAIFMTATPPGTRDAFPDSNSPIMDTEVEVPERAWSSGFDWVTDHSGKTVWFVPSVRNGNEIAACLTKAGKRVIQLSRKTFETEFQKTKHQEWDFVVTTDISEMGANFKADRVIDSRRCLKPVILDGERVILAGPMPVTHASAAQRRGRIGRNPNKPGDEYLYGGGCAETDEDHAHWLEARMLLDNIYLQDGLIASLYRPEADKVAAIEGEFKLRTEQRKTFVELMKRGDLPVWLAYQVASAGITYTDRRWCFDGATNNTILEDSVPAEVWTRHGEKRVLKPRWMDARVCSDHAALKSFKEFAAGKRGAAFGVMEALGTLPGHMTERFQEAIDNLAVLMRAETGSRPYKAAAAQLPETLETIMLLGLLGTVSLGIFFVLMRNKGIGKMGFGMVTLGASAWLMWLSEIEPARIACVLIVVFLLLVVLIPEPEKQRSPQDNQMAIIIMVAVGLLGLITANELGWLERTKSDLSHLMGRREEGATIGFSMDIDLRPASAWAIYAALTTFITPAVQHAVTTSYNNYSLMAMATQAGVLFGMGKGMPFYAWDFGVPLLMIGCYSQLTPLTLIVAIILLVAHYMYLIPGLQAAAARAAQKRTAAGIMKNPVVDGIVVTDIDTMTIDPQVEKKMGQVLLIAVAVSSAILSRTAWGWGEAGALITAATSTLWEGSPNKYWNSSTATSLCNIFRGSYLAGASLIYTVTRNAGLVKRRGGGTGETLGEKWKARLNQMSALEFYSYKKSGITEVCREEARRALKDGVATGGHAVSRGSAKLRWLVERGYLQPYGKVIDLGCGRGGWSYYAATIRKVQEVKGYTKGGPGHEEPVLVQSYGWNIVRLKSGVDVFHMAAEPCDTLLCDIGESSSSPEVEETRTLRVLSMVGDWLEKRPGAFCIKVLCPYTSTMMETLERLQRRYGGGLVRVPLSRNSTHEMYWVSGAKSNTIKSVSTTSQLLLGRMDGPRRPVKYEEDVNLGSGTRAVVSCAEAPNMKIIGNRIERIRSEHAETWFFDENHPYRTWAYHGSYEAPTQGSASSLINGVVRLLSKPWDVVTGVTGIAMTDTTPYGQQRVFKEKVDTRVPDPQEGTRQVMSMVSSWLWKELGKHKRPRVCTKEEFINKVRSNAALGAIFEEEKEWKTAVEAVNDPRFWALVDKEREHHLRGECQSCVYNMMGKREKKQGEFGKAKGSRAIWYMWLGARFLEFEALGFLNEDHWMGRENSGGGVEGLGLQRLGYVLEEMSCIPGGRMYADDTAGWDTRISRFDLENEALITNQMEKGHRALALAIIKYTYQNKVVKVLRPAEKGKTVMDIISRQDQRGSGQVVTYALNTFTNLVVQLIRNMEAEEVLEMQDLWLLRRSEKVTNWLQSNGWDRLKRMAVSGDDCVVKPIDDRFAHALRFLNDMGKVRKDTQEWKPSTGWDNWEEVPFCSHHFNKLHLKDGRSIVVPCRHQDELIGRARVSPGAGWSIRETACLAKSYAQMWQLLYFHRRDLRLMANAICSSVPVDWVPTGRTTWSIHGKGEWMTTEDMLVVWNRVWIEENDHMEDKTPVTKWTDIPYLGKREDLWCGSLIGHRPRTTWAENIKNTVNMVRRIIGDEEKYMDYLSTQVRYLGEEGSTPGVL

>tr|A0A146CJH9|A0A146CJH9_ZIKV Polyprotein OSZika virus (strain Mr 766) PE4 SV1

MKNPKKKSGGFRIVNMLKRGVARVSPFGGLKRLPAGLLLGHGPIRMVLAILAFLRFTAIKPSLGLINRWGSVGKKEAMEIIKKFKKDLAAMLRIINARKEKKRRGADTSVGIVGLLLTTAMAAEVTRRGSAYYMYLDRNDAGEAISFPTTLGMNKCYIQIMDLGHMCDATMSYECPMLDEGVEPDDVDCWCNTTSTWVVYGTCHHKKGEARRSRRAVTLPSHSTRKLQTRSQTWLESREYTKHLIRVENWIFRNPGFALAAAAIAWLLGSSTSQKVIYLVMILLIAPAYSIRCIGVSNRDFVEGMSGGTWVDVVLEHGGCVTVMAQDKPTVDIELVTTTVSNMAEVRSYCYEASISDMASDSRCPTQGEAYLDKQSDTQYVCKRTLVDRGWGNGCGLFGKGSLVTCAKFACSKKMTGKSIQPENLEYRIMLSVHGSQHSGMIVNDTGHETDENRAKVEITPNSPRAEATLGGFGSLGLDCEPRTGLDFSDLYYLTMNNKHWLVHKEWFHDIPLPWHAGADTGTPHWNNKEALVEFKDAHAKRQTVVVLGSQEGAVHTALAGALEAEMDGAKGRLSSGHLKCRLKMDKLRLKGVSYSLCTAAFTFTKIPAETLHGTVTVEVQYAGTDGPCKVPAQMAVDMQTLTPVGRLITANPVITESTENSKMMLELDPPFGDSYIVIGVGEKKITHHWHRSGSTIGKAFEATVRGAKRMAVLGDTAWDFGSVGGALNSLGKGIHQIFGAAFKSLFGGMSWFSQILIGTLLMWLGLNTKNGSISLMCLALGGVLIFLSTAVSADVGCSVDFSKKETRCGTGVFVYNDVEAWRDRYKYHPDSPRRLAAAVKQAWEDGICGISSVSRMENIMWRSVEGELNAILEENGVQLTVVVGSVKNPMWRGPQRLPVPVNELPHGWKAWGKSHFVRAAKTNNSFVVDGDTLKECPLKHRAWNSFLVEDHGFGVFHTSVWLKVREDYSLECDPAVIGTAVKGKEAVHSDLGYWIESEKNDTWRLKRAHLIEMKTCEWPKSHTLWTDGIEESDLIIPKSLAGPLSHHNTREGYRTQMKGPWHSEELEIRFEECPGTKVHVEETCGTRGPSLRSTTASGRVIEEWCCRECTMPPLSFRAKDGCWYGMEIRPRKEPESNLVRSMVTAGSTDHMDHFSLGVLVILLMVQEGLKKRMTTKIIISTSMAVLVAMILGGFSMSDLAKLAILMGATFAEMNTGGDVAHLALIAAFKVRPALLVSFIFRANWTPRESMLLALASCLLQTAISALEGDLMVLINGFALAWLAIRAMVVPRTDNITLAILAALTPLARGTLLVAWRAGLATCGGFMLLSLKGKGSVKKNLPFVMALGLTAVRLVDPINVVGLLLLTRSGKRSWPPSEVLTAVGLICALAGGFAKADIEMAGPMAAVGLLIVSYVVSGKSVDMYIERAGDITWEKDAEVTGNSPRLDVALDESGDFSLVEDDGPPMREIILKVVLMTICGMNPIAIPFAAGAWYVYVKTGKRSGALWDVPAPKEVKKGETTDGVYRVMTRRLLGSTQVGVGVMQEGVFHTMWHVTKGSALRSGEGRLDPYWGDVKQDLVSYCGPWKLDAAWDGHSEVQLLAVPPGERARNIQTLPGIFKTKDGDIGAVALDYPAGTSGSPILDKCGRVIGLYGNGVVIKNGSYVSAITQGRREEETPVECFEPSMLKKKQLTVLDLHPGAGKTRRVLPEIVREAIKTRLRTVILAPTRVVAAEMEEALRGLPVRYMTTAVNVTHSGTEIVDLMCHATFTSRLLQPIRVPNYNLYIMDEAHFTDPSSIAARGYISTRVEMGEAAAIFMTATPPGTRDAFPDSNSPIMDTEVEVPERAWSSGFDWVTDYSGKTVWFVPSVRNGNEIAACLTKAGKRVIQLSRKTFETEFQKTKHQEWDFVVTTDISEMGANFKADRVIDSRRCLKPVILDGERVILAGPMPVTHASAAQRRGRIGRNPNKPGDEYLYGGGCAETDEDHAHWLEARMLLDNIYLQDGLIASLYRPEADKVAAIEGEFKLRTEQRKTFVELMKRGDLPVWLAYQVASAGITYTDRRWCFDGTTNNTIMEDSVPAEVWTRHGEKRVLKPRWMDARVCSDHAALKSFKEFAAGKRGAAFGVMEALGTLPGHMTERFQEAIDNLAVLMRAETGSRPYKAAAAQLPETLETIMLLGLLGTVSLGIFFVLMRNKGIGKMGFGMVTLGASAWLMWLSEIEPARIACVLIVVFLLLVVLIPEPEKQRSPQDNQMAIIIMVAVGLLGLITANELGWLERTKSDLSHLMGRREEGATMGFSMDIDLRPASAWAIYAALTTFITPAVQHAVTTSYNNYSLMAMATQAGVLFGMGKGMPFYAWDFGVPLLMIGCYSQLTPLTLIVAIILLVAHYMYLIPGLQAAAARAAQKRTAAGIMKNPVVDGIVVTDIDTMTIDPQVEKKMGQVLLIAVAVSSAILSRTAWGWGEAGALITAATSTLWEGSPNKYWNSSTATSLCNIFRGSYLAGASLIYTVTRNAGLVKRRGGGTGETLGEKWKARLNQMSALEFYSYKKSGITEVCREEARRALKDGVATGGHAVSRGSAKLRWLVERGYLQPYGKVIDLGCGRGGWSYYAATIRKVQEVKGYTKGGPGHEEPVLVQSYGWNIVRLKSGVDVFHMAAEPCDTLLCDIGESSSSPEVEEARTLRVLSMVGDWLEKRPGAFCIKVLCPYTSTMMETLERLQRRYGGGLVRVPLSRNSTHEMYWVSGAKSNTIKSVSTTSQLLLGRMDGPRRPVKYEEDVNLGSGTRAVVSCAEAPNMKIIGNRIERIRSEHAETWFFDENHPYRTWAYHGSYEAPTQGSASSLINGVVRLLSKPWDVVTGVTGIAMTDTTPYGQQRVFKEKVDTRVPDPQEGTRQVMSMVSSWLWKELGKHKRPRVCTKEEFINKVRSNAALGAIFEEEKEWKTAVEAVNDPRFWALVDKEREHHLRGECQSCVYNMMGKREKKQGEFGKAKGSRAIWYMWLGARFLEFEALGFLNEDHWMGRENSGGGVEGLGLQRLGYVLEEMSRIPGGRMYADDTAGWDTRISRFDLENEALITNQMEKGHRALALAIIKYTYQNKVVKVLRPAEKGKTVMDIISRQDQRGSGQVVTYALNTFTNLVVQLIRNMEAEEVLEMQDLWLLRRSEKVTNWLQSNGWDRLKRMAVSGDDCVVKPIDDRFAHALRFLNDMGKVRKDTQEWKPSTGWDNWEEVPFCSHHFNKLHLKDGRSIVVPCRHQDELIGRARVSPGAGWSIRETACLAKSYAQMWQLLYFHRRDLRLMANAICSSVPVDWVPTGRTTWSIHGKGEWMTTEDMLVVWNRVWIEENDHMEDKTPVTKWTDIPYLGKREDLWCGSLIGHRPRTTWAENIKNTVNMVRRIIGDEEKYMDYLSTQVRYLGEEGSTPGVL

**DNA sequences (Aligned)**

>ENA|AMD61710|AMD61710.3 Zika virus polyprotein

ATGAAAAACCC-AAAAAAGAAATCCGGAGGATTCCGGATTGTCAATATGCTAAAACGCGGAGTAGCCCGTGTGAGCCCCTTTGGGGGCTTGAAGAGGCTGCCAGCCGGACTTCTGCTGGGTCATGGGCCCATCAGGATGGTCTTGGCGATTCTAGCCTTTTTGAGATTCACGGCAATCAAGCCATCACTGGGTCTCATCAATAGATGGGGTTCAGTGGGAAAAAAAGAGGCTATGGAAATAATAAAGAAGTTCAAGAAAGATCTGGCTGCCATGCTGAGAATAATCAATGCTAGGAAGGAGAAGAAGAGACGAGGCACAGATACTAGTGTCGGAATTGTTGGCCTCCTGCTGACCACAGCTATGGCAGCGGAGGTCACTAGACGTGGGAGTGCATACTATATGTACTTGGACAGAAGCGATGCTGGGGAGGCCATATCTTTTCCAACCACACTGGGGATGAATAAGTGTTATATACAGATCATGGATCTTGGACACATGTGTGATGCCACCATGAGCTATGAATGCCCTATGCTGGATGAGGGGGTAGAACCAGATGACGTCGATTGTTGGTGCAACACGACGTCAACTTGGGTTGTGTACGGAACCTGCCATCACAAAAAAGGTGAAGCACGGAGATCCAGAAGAGCTGTGACGCTCCCCTCCCATTCCACTAGGAAGCTGCAAACGCGGTCGCAGACCTGGTTGGAATCAAGAGAATACACAAAGCACTTGATTAGAGTCGAAAATTGGATATTCAGGAACCCTGGCTTCGCGTTAGCAGCAGCTGCCATCGCTTGGCTTTTGGGAAGCTCAACGAGCCAAAAAGTCATATACTTGGTCATGATACTGCTGATTGCCCCGGCATACAGCATCAGGTGCATAGGAGTCAGTAATAGGGACTTTGTGGAAGGTATGTCAGGTGGGACTTGGGTTGATGTTGTCTTGGAACATGGAGGTTGTGTCACCGTAATGGCACAGGACAAACCGACTGTCGACATAGAGCTGGTTACAACAACAGTCAGCAACATGGCGGAGGTAAGATCCTACTGCTATGAGGCATCAATATCGGACATGGCTTCGGACAGCCGCTGCCCAACACAAGGTGAAGCCTACCTTGACAAGCAATCAGACACTCAATATGTCTGCAAAAGAACGTTAGTGGACAGAGGCTGGGGAAATGGATGTGGACTTTTTGGCAAAGGGAGCCTGGTGACATGCGCTAAGTTTGCATGCTCCAAGAAAATGACCGGGAAGAGCATCCAGCCAGAGAATCTGGAGTACCGGATAATGCTGTCAGTTCATGGCTCCCAGCACAGTGGGATGATCGTTAATGACACAGGACATGAAACTGATGAGAATAGAGCGAAGGTTGAGATAACGCCCAATTCACCAAGAGCCGAAGCCACCCTGGGGGGTTTTGGAAGCCTAGGACTTGATTGTGAACCGAGGACAGGCCTTGACTTTTCAGATTTGTATTACTTGACTATGAACAACAAGCACTGGTTGGTTCACAAGGAGTGGTTCCACGACATTCCATTACCTTGGCACACTGGGGCAGACACCGGAACTCCACACTGGAACAACAAAGAAGCACTGGTAGAGTTCAAGGACGCACATGCCAAAAGGCAAACTGTCGTGGTTCTAGGGAGTCAAGAAGGAGCAGTTCACACGGCCCTTGCTGGAGCTCTGGAGGCTGAGATGGATGGTGCAAAGGGAAGGCTGTCCTCTGGCCACTTGAAATGTCGCCTGAAAATGGATAAACTTAGATTGAAGGGCGTGTCATACTCCTTGTGTACCGCAGCGTTCACATTCACCAAGATCCCGGCTGAAACACTGCACGGGACAGTCACAGTGGAGGTACAGTACGCAGGGACAGATGGACCTTGCAAGGTTCCAGCTCAGATGGCGGTGGACATGCAAACTCTGACCCCAGTTGGGAGGTTGATAACCGCTAACCCCGTAATCACTGAAGGCACTGAGAACTCTAAGATGATGCTGGAACTTGATCCACCATTTGGGGACTCTTACATTGTCATAGGAGTCGGGGAGAAGAAGATCACCCACCACTGGCACAGGAGTGGCAGCACCATTGGAAAAGCATTTGAAGCCACTGTGAGAGGTGCCAAGAGAATGGCAGTCTTGGGAGACACAGCCTGGGACTTTGGATCAGTTGGAGGCGTTCTTAACTCATTGGGCAAGGGCATCCATCAAATTTTTGGAGCAGCTTTCAAATCATTGTTTGGAGGAATGTCCTGGTTCTCACAAATTCTCATTGGAACGTTGCTGATGTGGTTGGGTCTGAATACAAAGAATGGATCTATTTCCCTTATGTGCTTGGCCTTAGGGGGAGTGTTGATCTTCTTATCCACAGCCGTCTCCGCTGATGTGGGGTGCTCGGTGGACTTCTCAAAGAAGGAAACGAGATGCGGTACAGGGGTGTTCGTCTATAACGACGTTGAAGCCTGGAGGGACAGGTACAAGTACCATCCTGACTCCCCTCGTAGATTGGCAGCAGTAGTCAAGCAAGCCTGGGAAGATGGTATCTGTGGGATCTCCTCTGTTTCAAGAATGGAAAACATCATGTGGAGATCAGTAGAAGGGGAGCTCAACGCAATCCTGGAAGAGAATGGAGTTCAACTGACGGTCGTTGTGGGATCTGTAAAAAACCCCATGTGGAGAGGTCCACAGAGATTGCCCGTGCCTGTGAACGAGCTGCCCCACGGCTGGAAGGCTTGGGGGAAATCGTACTTCGTCAGAGCAGCAAAGACAAATAACAGCTTTGTCGTGGATGGTGACACACTGAAGGAATGCCCACTCAAACATAGAGCATGGAACAGCTTTCTTGTGGAGGATCATGGGTTCGGGGTATTTCACACTAGTGTCTGGCTCAAGGTTAGAGAAGATTATTCACTAGAGTGTGATCCAGCCGTCATTGGAACAGCTGTTAAGGGAAAGGAGGCTGTACACAGTGATCTAGGCTACTGGATTGAGAGTGAGAAGAACGACACATGGAGGCTGAGGAGGGCCCACCTGATCGAGATGAAAACATGTGAATGGCCAAAGTCCCACACATTGTGGACAGATGGAATAGAAGAGAGTGATCTGATCATACCCAAGTCTTTAGCTGGGCCACTCAGCCATCACAACACCAGAGAGGGCTACAGGACCCAAATGAAAGGGCCATGGCACAGTGAAGAGCTTGAAATTCGGTTTGAGGAATGCCCAGGCACTAAGGTCCACGTGGAGGAAACATGTGGAACAAGAGGACCATCTCTGAGATCAACCACTGCAAGCGGAAGGGTGATCGAGGAATGGTGCTGCAGGGAGTGCACAATGCCCCCACTGTCGTTCCGGGCTAAAGATGGCTGTTGGTATGGAATGGAGATAAGGCCCAGGAAAGAACCAGAAAGTAACTTAGTAAGGTCAATGGTGACTGCAGGATCAACTGATCACATGGATCACTTTTCCCTTGGAGTGCTTGTGATTCTGCTCATGGTGCAGGAAGGGCTGAAGAAGAGAATGACCACAAAGATCATCATAAGCACATCAATGGCAGTGCTGGTAGCTATGATCCTGGGAGGATTTTCAATGAGTGATCTGGCTAAGCTTGCAATTTTGATGGGTGCCACCTTTGCGGAAATGAACACTGGAGGAGATGTAGCTCATCTGGCGCTGGTAGCGGCATTCAAAGTCAGACCAGCGTTGCTGGTATCTTTCATCTTCAGAGCTAATTGGACACCCCGTGAAAGCATGCTGCTGGCCTTGGCCTCGTGTCTTTTGCAAACTGCGATCTCCGCCTTGGAAGGCGACCTGATGGTTCTCATCAATGGTTTTGCTTTGGCCTGGTTGGCAATACGAGCGATGGTTGTTCCACGCACTGACAATATCACCTTGGCAATCCTGGCTGCTCTGACACCACTGGCCCGGGGCACACTGCTTGTGGCGTGGAGAGCAGGCCTTGCTACTTGCGGGGGGTTCATGCTCCTCTCTCTGAAGGGGAAAGGCAGTGTGAAGAAGAACTTACCATTTGTCATGGCCCTGGGACTAACCGCTGTGAGGCTGGTCGACCCCATCAACGTGGTGGGACTGCTGTTGCTCACAAGGAGTGGGAAGCGGAGCTGGCCCCCTAGCGAAGTACTCACAGCTGTTGGCCTGATATGCGCATTGGCTGGAGGGTTCGCCAAGGCAGATATAGAGATGGCTGGGCCCATGGCCGCGGTCGGTCTGCTAATTGTCAGTTACGTGGTCTCAGGAAAGAGTGTGGACATGTACATTGAAAGAGCAGGTGACATCACATGGGAAAAAGATGCGGAAGTTACTGGAAACAGTCCCCGGCTCGATGTGGCACTAGATGAGAGTGGTGATTTCTCCCTGGTGGAGGATGACGGTCCCCCCATGAGAGAGATCATACTCAAAGTGGTCCTGATGACCATCTGTGGCATGAACCCAATAGCCATACCCTTTGCAGCTGGAGCGTGGTACGTATACGTGAAAACTGGAAAAAGGAGTGGTGCTCTATGGGATGTGCCTGCTCCCAAGGAAGTAAAAAAGGGGGAGACCACAGATGGAGTGTACAGAGTAATGACTCGTAGACTGCTAGGTTCAACACAAGTTGGAGTGGGAGTTATGCAAGAGGGGGTCTTTCACACTATGTGGCATGTCACAAAAGGATCCGCGCTGAGAAGCGGTGAAGGGAGACTTGATCCATACTGGGGAGATGTCAAGCAGGATCTGGTGTCATACTGTGGTCCATGGAAGCTAGATGCCGCCTGGGACGGGCACAGCGAGGTGCAGCTCTTGGCCGTGCCCCCCGGAGAGAGAGCGAGGAACATCCAGACTCTGCCCGGAATATTTAAGACAAAGGATGGGGACATTGGAGCGGTTGCGCTGGACTATCCAGCAGGAACTTCAGGATCTCCAATCCTAGACAAGTGTGGGAGAGTGATAGGACTCTATGGCAATGGGGTCGTGATCAAGAATGGGAGTTATGTCAGTGCCATCACCCAAGGGAGGAGGGAGGAAGAGACTCCTGTTGAGTGCTTCGAGCCTTCGATGCTGAAGAAGAAGCAGCTAACTGTCTTAGACTTGCATCCTGGAGCTGGGAAAACCAGGAGAGTTCTTCCTGAAATAGTCCGTGAAGCCATAAAAACGAGACTCCGTACTGTGATCTTAGCTCCAACCAGGGTTGTCGCTGCTGAAATGGAGGAAGCCCTTAGAGGGCTTCCAGTGCGTTATATGACAACAGCAGTCAATGTCACCCATTCTGGGACAGAAATCGTTGACTTAATGTGCCATGCCACCTTCACTTCACGTCTACTACAGCCAATCAGAGTCCCCAACTATAATCTGTATATTATGGATGAGGCCCACTTCACAGATCCCTCAAGTATAGCAGCAAGAGGATACATTTCAACAAGGGTTGAGATGGGCGAGGCAGCTGCCATCTTCATGACCGCCACGCCACCAGGAACCCGTGACGCATTCCCGGACTCCAACTCACCAATTATGGACACCGAAGTGGAAGTCCCAGAGAGAGCCTGGAGCTCAGGCTTTGATTGGGTGACGGATCATTCTGGAAAAACAGTTTGGTTTGTCCCAAGCGTGAGGAACGGCAATGAGATCGCAGCTTGTCTGACAAAGGCTGGAAAACGGGTCATACAGCTCAGCAGAAAGACTTTTGAGACAGAGTTCCAGAAAACAAAACATCAAGAGTGGGACTTCGTCGTGACAACTGACATTTCAGAGATGGGCGCCAACTTTAAAGCTGACCGTGTCATAGATTCCAGGAGATGCCTAAAGCCGGTCATACTTGATGGCGAGAGAGTCATTCTGGCTGGACCCATGCCTGTCACACATGCCAGCGCTGCCCAGAGGAGGGGGCGCATAGGCAGGAATCCCAACAAACCTGGAGATGAGTATCTGTATGGAGGTGGGTGCGCAGAGACTGATGAAGACCATGCACACTGGCTTGAAGCAAGAATGCTCCTTGACAATATTTACCTCCAAGATGGCCTCATAGCCTCGCTCTATCGACCTGAGGCCGACAAAGTAGCAGCCATTGAGGGAGAGTTCAAGCTTAGGACGGAGCAAAGGAAGACCTTTGTGGAACTCATGAAAAGAGGAGATCTTCCTGTTTGGCTGGCCTATCAGGTTGCATCTGCCGGAATAACCTACACAGATAGAAGATGGTGCTTTGATGGCATGACCAACAACACCATAATGGAAGACAGTGTGCCGGCAGAGGTGTGGACCAGACACGGAGAGAAAAGAGTGCTCAAACCGAGGTGGATGGACGCCAGAGTTTGTTCAGATCATGCGGCCCTGAAGTCATTCAAGGAGTTTGCCGCTGGGAAAAGAGGAGCGGCTTTTGGAGTGATGGAAGCCCTGGGAACACTGCCAGGACACATGACGGAGAGATTCCAGGAAGCCATTGACAACCTCGCTGTGCTCATGCGGGCAGAGACTGGAAGCAGGCCTTACAAAGCCGCGGCGGCCCAATTGCCGGAGACCCTAGAGACCATTATGCTTTTGGGGTTGCTGGGAACAGTCTCGCTGGGAATCTTTTTCGTCTTGATGCGGAACAAGGGCATAGGGAAGATGGGCTTTGGAATGGTGACTCTTGGGGCCAGCGCATGGCTCATGTGGCTCTCGGAAATTGAGCCAGCCAGAATTGCATGCGTCCTCATTGTTGTGTTCCTATTGCTGGTGGTGCTCATACCTGAGCCAGAAAAGCAAAGATCCCCCCAGGACAACCAAATGGCAATCATCATCATGGTAGCAGTAGGTCTTCTGGGCTTGATTACCGCCAATGAACTCGGATGGTTGGAGAGAACAAAGAGTGACCTAAGCCATCTAATGGGAAGGAGAGAGGAGGGGGCAACCATAGGATTCTCAATGGACATTGACCTGCGGCCAGCCTCGGCCTGGGCCATCTATGCTGCCCTGACAACTTTCATTACCCCAGCCGTCCAACATGCAGTGACCACTTCATACAACAACTACTCCTTAATGGCGATGGCCACGCAAGCTGGAGTGTTGTTTGGTATGGGCAAAGGGATGCCATTCTACGCATGGG-ACTTTGGAGTCCCGCTGCTAATGATAGGTTGCTACTCACAATTAACACCCCTGACCCTAATAGTGGCTATCATTTTGCTCGTGGCGCACTACATGTACTTGATCCCAGGGCTGCAGGCAGCAGCTGCGCGTGCTGCCCAGAAGAGAACGGCAGCTGGCATCATGAAGAACCCTGTTGTGGATGGAATAGTGGTGACTGACATTGACACAATGACTATTGACCCCCAAGTGGAGAAAAAGATGGGACAGGTGCTACTCATAGCAGTAGCCGTCTCCAGCGCCATACTGTCGCGGACCGCCTGGGGGTGGGGGGAAGCTGGGGCCCTGATCACAGCTGCAACTTCCACTTTGTGGGAAGGCTCTCCGAACAAGTACTGGAACTCCTCTACAGCCACTTCACTGTGCAACATTTTTAGGGGAAGTTACTTGGCTGGAGCTTCTCTAATCTACACAGTAACAAGAAACGCTGGCTTGGTCAAGAGACGTGGGGGTGGAACAGGAGAGACCCTGGGAGAGAAATGGAAGGCCCGCTTGAACCAGATGTCGGCCCTGGAGTTCTACTCCTACAAAAAGTCAGGCATCACCGAGGTGTGCAGAGAAGAGGCCCGCCGCGCCCTCAAGGACGGTGTGGCAACGGGAGGCCATGCTGTGTCCCGAGGAAGTGCAAAGCTGAGATGGTTGGTGGAGCGGGGATACCTGCAGCCCTATGGAAAGGTCATTGATCTTGGATGTGGCAGAGGGGGCTGGAGTTACTACGCCGCCACCATCCGCAAAGTTCAAGAAGTGAAAGGATACACAAAAGGAGGCCCTGGTCATGAAGAACCCATGTTGGTGCAAAGCTATGGGTGGAACATAGTCCGTCTTAAGAGTGGGGTGGACGTCTTTCATATGGCGGCTGAGCCGTGTGACACGTTGCTGTGTGACATAGGTGAGTCATCATCTAGTCCTGAAGTGGAAGAAGCACGGACGCTCAGAGTCCTCTCCATGGTGGGGGATTGGCTTGAAAAAAGACCAGGAGCCTTTTGTGTAAAAGTGTTGTGCCCATACACCAGCACTATGATGGAAACCCTGGAGCGACTGCAGCGTAGGTATGGGGGAGGACTGGTCAGAGTGCCACTCTCCCGCAACTCTACACATGAGATGTACTGGGTCTCTGGAGCGAAAAGCAACACCATAAAAAGTGTGTCCACCACGAGCCAGCTCCTCTTGGGGCGCATGGACGGGCCCAGGAGGCCAGTGAAATATGAGGAGGATGTGAATCTCGGCTCTGGCACGCGGGCTGTGGTAAGCTGCGCTGAAGCTCCCAACATGAAGATCATTGGTAACCGCATTGAAAGGATCCGCAGTGAGCACGCGGAAACGTGGTTCTTTGACGAGAACCACCCATATAGGACATGGGCTTACCATGGAAGCTATGAGGCCCCTACACAAGGGTCAGCGTCCTCTCTAATAAACGGGGTTGTCAGGCTCCTGTCAAAACCCTGGGATGTGGTGACTGGAGTCACAGGAATAGCCATGACCGACACCACACCGTATGGTCAGCAAAGAGTTTTCAAGGAAAAAGTGGACACCAGGGTGCCAGACCCCCAAGAAGGCACTCGTCAGGTTATGAGCATGGTCTCTTCCTGGTTGTGGAAAGAGCTAGGCAAACACAAACGGCCACGAGTCTGTACCAAAGAAGAGTTCATCAACAAGGTTCGTAGCAATGCAGCATTAGGGGCAATATTTGAAGAGGAAAAAGAGTGGAAGACCGCAGTGGAAGCTGTGAACGATCCAAGGTTCTGGGCTCTAGTGGACAAGGAAAGAGAGCACCACCTGAGAGGAGAGTGCCAGAGCTGTGTGTACAACATGATGGGAAAAAGAGAAAAGAAACAAGGGGAATTTGGAAAGGCCAAGGGCAGCCGCGCCATCTGGTATATGTGGCTAGGGGCTAGATTTCTAGAGTTCGAAGCCCTTGGATTCTTAAATGAGGATCACTGGATGGGGAGAGAGAACTCAGGAGGTGGTGTTGAAGGGCTGGGATTACAAAGACTCGGATATGTCCTAGAAGAGATGAGTCGCATACCAGGAGGAAGGATGTATGCAGATGACACTGCTGGCTGGGACACCCGCATCAGCAGGTTTGATCTGGAGAATGAAGCTTTAATCACCAACCAAATGGAGAAAGGGCACAGGGCCTTAGCATTGGCCATAATCAAGTACACATACCAAAACAAAGTGGTAAAGGTCCTTAGACCAGCTGAAAAAGGGAAGACAGTTATGGACATTATTTCAAGACAAGACCAAAGGGGGAGCGGACAAGTTGTCACTTACGCTCTTAACACATTTACCAACCTAGTGGTGCAACTCATTCGGAATATGGAGGCTGAGGAAGTTCTAGAGATGCAAGACTTGTGGCTGCTGCGGAGGTCAGAGAAAGTGACCAACTGGTTGCAGAGCAACGGATGGGATAGGCTCAAACGAATGGCAGTCAGTGGAGATGATTGCGTTGTGAAGCCAATTGATGATAGGTTTGCACATGCCCTCAGGTTCTTGAATGATATGGGAAAAGTTAGGAAGGACACACAAGAGTGGAAACCCTCAACTGGATGGGACAACTGGGAAGAAGTTCCGTTTTGTTCCCACCACTTCAACAAGCTCCATCTCAAGGACGGGAGGTCCATTGTGGTTCCCTGCCGCCACCAAGATGAACTGATTGGCCGGGCCCGTGTCTCTCCAGGGGCGGGATGGAGCATCCGGGAGACTGCTTGCCTAGCAAAGTCATATGCGCAAATGTGGCAGCTCCTTTATTTCCACAGAAGGGACCTCCGACTGATGGCCAATGCCATCTGTTCATCTGTGCCAGTTGACTGGGTTCCAACTGGGAGAACTACCTGGTCAATCCATGGAAAGGGAGAATGGATGACCACTGAAGACATGCTTGTGGTGTGGAACAGAGTGTGGATTGAGGAGAACGACCACATGGAAGACAAGACCCCAGTTACGAAATGGACAGACATTCCCTATCTGGGAAAAAGGGAAGACTTGTGGTGTGGATCTCTCATAGGGCACAGACCGCGCACCACCTGGGCTGAGAACATTAAAAACACAGTCAACATGGTGCGCAGGATCATAGGTGATGAAGAAAAGTACATGGACTACCTATCCACCCAAGTTCGCTACTTGGGTGAAGAAGGGTCTACACCTGGAGTGCTATAA

>ENA|AMD61711|AMD61711.3 Zika virus polyprotein

ATGAAAAACCC-AAAAAAGAAATCCGGAGGATTCCGGATTGTCAATATGCTAAAACGCGGAGTAGCCCGTGTGAGCCCCTTTGGGGGCTTGAAGAGGCTGCCAGCCGGACTTCTGCTGGGCCATGGGCCCATCAGGATGGTCTTGGCGATACTAGCCTTTTTGAGATTCACGGCAATCAAGCCATCACTGGGTCTCATCAATAGATGGGGTTCAGTGGGGAAAAAAGAGGCTATGGAAATAATAAAGAAGTTCAAGAAAGATCTGGCTGCCATGCTGAGAATAATCAATGCTAGGAAGGAGAAGAAGAGACGAGGCGCAGATACTAGCGTCGGAATTGTTGGCCTCCTCCTGACCACAGCCATGGCAGTAGAGGTCACTAGACGTGGGAGTGCATACTATATGTACTTGGACAGAAGCGATGCTGGGGAGGCCATATCTTTTCCAACCACACTGGGGATGAATAAGTGTTACATACAAATCATGGATCTTGGACACATGTGTGATGCCACCATGAGCTATGAATGCCCTATGTTGGATGAGGGGGTAGAACCAGATGACGTCGATTGCTGGTGCAACACGACATCAACTTGGGTTGTGTATGGAACCTGCCACCACAAAAAAGGTGAAGCACGGAGATCTAGAAGAGCTGTGACGCTCCCCTCCCATTCCACTAGGAAGCTGCAAACGCGGTCGCAGACCTGGTTGGAATCAAGAGAATACACAAAGCACCTGATTAGAGTTGAAAATTGGATATTCAGGAACCCTGGCTTCGCGTTAGCAGCAGCTGTCATCGCTTGGCTTTTGGGAAGTTCAACGAGCCAAAAAGTCATATATCTGGTCATGATACTGCTGATTGCCCCGGCATACAGCATCAGGTGCATAGGAGTCAGCAATAGGGACTTTGTGGAAGGTATGTCAGGTGGGACTTGGGTTGATGTTGTCTTGGAACATGGAGGTTGTGTTACCGTAATGGCACAGGACAAACCGACTGTCGACATAGAGCTGGTTACAACAACAGTCAGCAACATGGCGGAGGTAAGATCCTACTGCTATGAGGCATCAATATCGGATATGGCTTCGGACAGCCGCTGCCCAACACAAGGTGAGGCCTACCTTGACAAGCAGTCAGACACTCAATATGTCTGCAAAAGAACGTTAGTGGACAGAGGCTGGGGAAATGGATGTGGACTTTTTGGCAAAGGGAGCCTGGTGACATGCGCTAAGTTTGCATGCTCCAAGAAAATGACCGGGAAGAGCATCCAGCCAGAGAATCTGGAGTACCGGATAATGCTGTCAGTTCATGGCTCCCAGCACAGTGGGATGATCGTTAATGACACAGGACATGAAACTGATGAGAATAGAGCGAAGGTTGAGATAACGCCCAATTCACCAAGAGCCGAAGCCACCCTGGGGGGTTTTGGGAGCCTAGGACTTGATTGTGAACCGAGGACAGGCCTTGACTTTTCAGATTTGTATTACCTGACTATGAATAACAAGCACTGGTTGGTTCACAAGGAGTGGTTCCACGACATTCCATTACCTTGGCATGCTGGGGCAGACACTGGAACTCCACATTGGAACAACAAAGAAGCACTGGTAGAGTTCAAGGACGCACATGCAAAAAGGCAAACTGTCGTGGTTCTAGGGAGTCAAGAAGGAGCAGTTCACACGGCCCTTGCTGGAGCTCTGGAGGCTGAGATGGATGGAGCCAAGGGAAGGCTGTCCTCTGGCCACTTGAAATGTCGCCTGAAAATGGATAAACTTAGATTGAAGGGCGTGTCATACTCCTTGTGCACTGCAGCGTTCACATTCACCAAGATCCCGGCTGAAACACTGCACGGGACAGTCACAGTGGAGGTACAGTACGCAGGGACAGATGGACCTTGCAAGGTTCCAGCTCAGATGGCGGTGGATATGCAAACTCTGACCCCAGTTGGGAGGTTGATAACCGCTAACCCTGTAATCACTGAAAGCACCGAGAACTCTAAGATGATGCTGGAACTTGATCCACCATTTGGGGACTCTTACATTGTCATAGGAGTCGGGGAGAAGAAGATCACCCATCACTGGCACAGGAGTGGCAGCACCATTGGAAAAGCATTTGAAGCCACTGTGAGAGGTGCCAAGAGAATGGCAGTCTTGGGAGACACAGCCTGGGACTTTGGATCAGTTGGGGGTGCTCTCAACTCATTGGGCAAGGGCATCCATCAAATTTTTGGAGCAGCTTTCAAATCATTGTTCGGAGGAATGTCCTGGTTCTCACAAATTCTCATTGGAACGTTGCTGGTGTGGTTGGGTCTGAATACAAAGAATGGATCTATTTCCCTTACGTGCTTGGCCTTAGGGGGAGTGTTGATCTTCTTATCCACAGCCGTTTCTGCTGATGTGGGGTGCTCGGTGGACTTCTCAAAGAAGGAAACGAGATGCGGTACAGGGGTGTTCGTCTATAACGACGTTGAAGCCTGGAGGGACAGGTACAAGTACCATCCTGACTCCCCTCGTAGATTGGCAGCAGCAGTCAAGCAAGCCTGGGAAGATGGGATCTGTGGGATCTCCTCTGTCTCAAGAATGGAAAACATCATGTGGAGATCAGTAGAAGGGGAGCTCAACGCAATCCTGGAAGAGAATGGAGTTCAACTGACGGTCGTTGTGGGATCTGTAAAAAACCCCATGTGGAGAGGTCCACAGAGATTGCCCGTGCCTGTGAACGAGCTGCCCCACGGCTGGAAGGCTTGGGGGAAATCGTACTTCGTCAGAGCAGCAAAGACAAATAACAGCTTTGTCGTGGATGGTGACACACTGAAGGAATGCCCACTCAAACATAGAGCATGGAACAGCTTTCTTGTGGAGGATCATGGGTTTGGGGTATTTCACACTAGTGTCTGGCTCAAGGTTAGAGAAGATTATTCATTAGAGTGTGATCCAGCCGTCATTGGAACAGCTGCTAAGGGAAAGGAGGCTGTGCACAGCGATCTAGGCTACTGGATTGAGAGTGAGAAGAACGACACATGGAGGCTGAAGAGGGCCCACCTGATCGAGATGAAAACATGTGAATGGCCAAAGTCCCACACATTGTGGACAGATGGAGTAGAAGAAAGTGATCTGATCATACCCAAGTCTTTAGCTGGGCCACTCAGCCATCACAACACCAGAGAGGGCTACAGGACTCAAATGAAAGGGCCATGGCACAGTGAAGAGCTTGAAATTCGGTTTGAGGAATGCCCAGGCACTAAGGTCCACGTGGAGGAAACATGTGGGACAAGAGGACCATCCCTGAGATCAACCACTGCAAGCGGAAGGGTGATCGAGGAATGGTGCTGCAGGGAATGCACAATGCCCCCACTGTCGTTCCGAGCTAAAGATGGCTGTTGGTATGGAATGGAGATAAGGCCCAGGAAAGAACCAGAAAGTAACTTAGTAAGGTCAATGGTGACTGCAGGATCAACTGATCACATGGATCACTTCTCTCTTGGAGTGCTTGTGATTTTGCTCATGGTGCAGGAAGGGCTGAAGAAGAGAATGACCACAAAGATCATCATAAGCACATCAATGGCAGTGCTGGTAGCCATGATCCTGGGAGGATTTTCAATGAGTGACCTGGCTAAGCTTGCAATTTTGATGGGTGCCACCTTCGCGGAAATGAACACTGGAGGAGATGTAGCTCATTTGGCGCTGATAGCGGCATTCAAAGTCAGACCTGCGTTGCTGGTATCTTTCATCTTCAGAGCTAATTGGACACCCCGTGAGAGCATGCTGCTGGCCTTGGCCTCGTGTCTTCTGCAAACTGCGATCTCCGCCTTGGAAGGCGACCTGATGGTTCTCATCAATGGTTTTGCTTTGGCCTGGTTGGCAATACGAGCGATGGTTGTTCCACGCACTGACAACATCACCTTGGCAATCCTGGCTGCTCTGACACCACTGGCCCGGGGCACACTGCTTGTGGCGTGGAGAGCAGGCCTTGCTACTTGCGGGGGGTTCATGCTCCTCTCTCTGAAGGGGAAAGGCAGTGTGAAGAAGAACCTACCATTTGTCATGGCCTTGGGACTAACTGCTGTGAGGCTGGTCGACCCCATCAACGTGGTGGGACTGCTGTTGCTCACAAGGAGTGGGAAGCGGAGCTGGCCCCCTAGTGAAGTACTCACAGCTGTTGGCCTGATATGCGCATTGGCTGGAGGGTTCGCCAAGGCGGATATAGAGATGGCTGGGCCCATGGCCGCGGTCGGTCTGCTAATTGTCAGTTACGTGGTCTCAGGAAAGAGTGTGGACATGTACATTGAAAGAGCAGGTGACATCACATGGGAAAAAGATGCGGAAATCACTGGAAACAGTCCCCGGCTCGATGTGGCACTAGATGAGAGTGGTGATTTCTCCCTAGTGGAGGATGATGGTCCACCCATGAGAGAGATCATACTCAAAGTGGTCCTGATGACCATCTGCGGCATGAACCCAATAGCCATACCCTTTGCAGCTGGAGCGTGGTACGTGTATGTGAAGACTGGAAAAAGGAGTGGTGCTCTATGGGATGTGCCTGCTCCCAAGGAAGTAAAAAAGGGGGAGACCACAGATGGAGTGTACAGAGTAATGACTCGTAGACTGCTTGGTTCAACACAAGTTGGAGTGGGAGTCATGCAAGAGGGGGTCTTCCACACTATGTGGCACGTCACAAAAGGATCCGCGCTGAGAAGCGGTGAAGGGAGACTTGATCCATACTGGGGAGATGTCAAGCAGGATCTGGTGTCATACTGTGGTCCGTGGAAGCTAGACGCCGCCTGGGACGGGCACAGCGAGGTGCAGCTCTTGGCCGTGCCCCCCGGAGAGAGAGCGAGGAACATCCAGACTCTGCCCGGAACATTTAAGACAAAGGATGGGGACATTGGAGCAGTTGCGCTGGACTACCCAGCAGGAACTTCAGGATCTCCAATCCTAGACAAGTGTGGGAGAGTGATAGGACTCTATGGTAATGGGGTCGTGATAAAAAATGGGAGTTATGTTAGTGCCATCACCCAAGGGAGGAGGGAGGAAGAGACTCCTGTTGAGTGCTTCGAGCCTTCGATGCTGAAGAAGAAGCAGCTAACTGTCTTAGACCTGCATCCTGGAGCCGGGAAAACCAGGAGAGTTCTTCCTGAAATAGTCCGTGAAGCCATAAAAACAAGACTCCGTACTGTGATCTTAGCTCCAACCAGGGTCGTCGCTGCTGAAATGGAGGAAGCCCTTAGAGGGCTTCCAGTTCGTTATATGACAACAGCAGTCAATGTCACCCATTCTGGGACAGAAATCGTTGACTTAATGTGCCATGCTACCTTCACTTCACGCCTACTACAACCAATCAGAGTCCCCAACTATAATTTGTATATTATGGATGAGGCCCACTTCACAGATCCCTCAAGTATAGCAGCAAGAGGATACATTTCAACAAGGGTTGAGATGGGCGAGGCGGCTGCCATCTTCATGACCGCCACGCCACCAGGAACCCGTGACGCATTCCCGGACTCCAACTCACCAATTATGGACACCGAGGTGGAAGTCCCAGAGAGAGCCTGGAGCACAGGCTTTGATTGGGTGACGGATCATTCTGGGAAAACAGTCTGGTTTGTTCCAAGCGTGAGGAACGGCAATGAGATCGCAGCTTGTCTGACAAAGGCTGGAAAACGGGTCATACAGCTCAGCAGAAAGACTTTTGAGACAGAGTTCCAGAAAACGAAAAATCAAGAGTGGGACTTCGTCGTGACAACCGACATTTCAGAGATGGGCGCCAACTTTAAAGCTGACCGTGTCATAGATTCCAGGAGATGCTTAAAGCCGGTCATACTTGATGGCGAGAGAGTCATTTTGGCTGGACCCATGCCTGTCACACATGCCAGCGCTGCTCAGAGGAGGGGGCGCATAGGCAGGAATCCCAACAAACCTGGAGATGAGTATCTGTATGGAGGTGGGTGCGCAGAGACTGATGAAGATCACGCACACTGGCTTGAAGCAAGAATGCTTCTTGACAACATTTACCTCCAAGATGGCCTCATAGCTTCGCTCTATCGACCTGAGGCCGACAAAGTAGCAGCTATTGAGGGAGAGTTCAAGCTTAGGACGGAGCAAAGGAAGACCTTTGTGGAACTCATGAAAAGAGGAGATCTTCCGGTTTGGTTGGCCTATCAGGTTGCATCTGCCGGAATAACCTACACAGATAGAAGATGGTGCTTTGATGGCATGACCAACAACACCATAATGGAAGACAGTGTGCCGGCAGAGGTGTGGACCAGATACGGAGAGAAAAGAGTGCTCAAACCGAGGTGGATGGACGCCAGAGTTTGTTCAGATCATGCGGCCCTGAAGTCATTCAAAGAGTTTGCCGCTGGGAAAAGAGGAGCGGCCTTTGGAGTGATAGAAGCCCTGGGAACACTGCCAGGACACATGACAGAGAGATTCCAGGAAGCCATTGACAACCTCGCTGTGCTCATGCGGGCAGAGACTGGAAGCAGGCCTTACAAAGCCGCGGCGGCCCAATTGCCGGAGACCCTAGAGACCATTATGCTTTTGGGGTTGCTGGGAACAGTCTCGCTGGGAATCTTTTTCGTCTTGATGCGGAACAAGGGCATGGGGAAGATGGGCTTTGGAATGGTGACTCTTGGGGCCAGCGCATGGCTTATGTGGCTCTCGGAAATTGAGCCAGCCAGAATTGCATGTGTCCTCATTGTCGTGTTCCTATTGCTGGTGGTGCTCATACCTGAGCCAGAAAAGCAAAGATCTCCTCAGGACAACCAAATGGCAATCATCATCATGGTAGCAGTGGGTCTTCTGGGCTTGATTACCGCCAATGAACTCGGATGGTTGGAGAGAACAAAAAGTGACCTAAGCCATCTAATGGGAAGGAGAGAGGAGGGGGCAACCACAGGATTCTCAATGGACATTGACCTGCGGCCAGCCTCAGCTTGGGCTATCTATGCTGCTCTGACAACTTTCATCACCCCAGCCGTCCAACATGCGGTGACCACTTCATACAACAACTACTCCTTAATGGCGATGGCCACGCAAGCTGGGGTGTTGTTTGGTATGGGCAAAGGGATGCCATTCTACGCATGGG-ACTTTGGAGTCCCGCTGCTAATGATGGGTTGCTACTCACAATTAACACCTCTGACCCTAATAGTGGCCATCATTTTGCTCGTGGCGCACTACATGTACTTGATCCCAGGGCTGCAGGCAGCAGCTGCGCGGGCTGCCCAGAAGAGAACGGCAGCTGGCATCATGAAGAACCCTGTTGTGGATGGAATAGTGGTGACTGACATTGACACAATGACAATTGACCCCCAAGTGGAAAAAAAGATGGGGCAGGTGCTACTCATAGCAGTAGCCGTCTCCAGCGCCATACTGTCGCGGACCGCCTGGGGGTGGGGGGAGGCTGGGGCCCTGATCACAGCTGCAACTTCCACCTTGTGGGAAGGCTCTCCGAACAAGTACTGGAACTCCTCCACAGCCACTTCACTGTGTAACATTTTTAGGGGAAGTTACTTGGCTGGAGCTTCTCTAATCTACACAGTAACAAGAAACGCTGGCTTGGTCAAGAGACGTGGGGGTGGAACGGGAGAGACCCTGGGAGAGAAATGGAAGGCCCGCCTGAACCAGATGTCGGCCCTGGAGTTCTACTCCTACAAAAAGTCAGGCATCACCGAGGTGTGCAGAGAAGAGGCCCGCCGTGCCCTCAAGGACGGTGTGGCAACAGGAGGCCATGCTGTGTCCCGAGGAAGTGCAAAGCTTAGATGGCTGGTGGAGAGAGGATACCTGCAGCCCTATGGAAAGGTCATTGATCTTGGATGTGGCAGAGGGGGCTGGAGTTACTATGCCGCCACCATCCGCAAAGTTCAGGAAGTGAAAGGATACACAAAAGGAGGCCCTGGTCATGAAGAACCCATGTTGGTGCAAAGCTATGGGTGGAACATAGTCCGTCTTAAGAGTGGGGTGGACGTCTTTCACATGGCGGCTGAGCCGTGTGACACTTTGCTGTGTGATATAGGTGAGTCATCATCTAGTCCTGAAGTGGAAGAAGCACGGACGCTCAGAGTCCTCTCCATGGTGGGGGATTGGCTTGAAAAAAGACCAGGAGCCTTTTGTATAAAAGTGTTGTGCCCATACACCAGCACTATGATGGAAACCCTGGAGCGACTGCAGCGTAGGTATGGGGGAGGACTGGTCAGGGTGCCACTCTCCCGCAACTCTACACATGAGATGTACTGGGTCTCTGGAGCGAAAAGCAACACCATAAAAAGTGTGTCCACCACGAGCCAGCTCCTCTTGGGGCGCATGGACGGGCCCAGGAGGCCAGTGAAATATGAGGAGGATGTGAATCTCGGCTCTGGCACGCGGGCTGTGGTAAGCTGCGCTGAAGCTCCCAACATGAAGATCATTGGTAACCGCATTGAGAGGATCCGCAGTGAGCACGCGGAAACGTGGTTCTTTGACGAGAACCACCCATATAGGACATGGGCTTACCATGGAAGCTATGAGGCCCCTACACAAGGGTCAGCGTCCTCTCTAATAAACGGGGTTGTCAGGCTCCTGTCAAAACCCTGGGATGTGGTGACTGGAGTCACAGGAATAGCCATGACTGACACCACACCGTATGGTCAGCAAAGAGTTTTCAAGGAAAAAGTGGACACTAGGGTGCCAGACCCCCAAGAAGGCACTCGTCAGGTTATGAGCATGGTCTCTTCCTGGTTATGGAAGGAGCTAGGCAAACACAAACGGCCACGAGTCTGTACCAAAGAAGAGTTCATCAACAAGGTTCGTAGCAATGCAGCATTAGGGGCAATATTTGAAGAGGAAAAAGAGTGGAAGACTGCAGTGGAAGCTGTGAATGATCCAAGGTTCTGGGCTCTAGTGGACAAGGAAAGAGAGCATCACCTGAGAGGAGAGTGTCAGAGCTGTGTGTACAACATGATGGGAAAAAGAGAAAAGAAACAAGGGGAATTTGGAAAGGCCAAGGGCAGCCGCGCCATCTGGTATATGTGGCTAGGGGCTAGATTCCTAGAGTTCGAAGCCCTTGGATTCTTGAATGAGGATCATTGGATGGGGAGAGAGAATTCAGGAGGTGGTGTTGAAGGACTGGGATTACAAAGACTCGGATATGTCCTAGAAGAGATGAGTCGCATACCAGGAGGAAGGATGTATGCAGATGATACTGCTGGCTGGGACACCCGCATCAGCAGGTTTGATCTGGAGAATGAAGCTCTAATCACCAACCAAATGGAGAAAGGGCACAGGGCCTTGGCATTGGCCATAATCAAGTACACATACCAAAACAAAGTGGTAAAGGTCCTTAGACCAGCTGAAAAAGGGAAGACAGTTATGGACATTATTTCAAGACAAGACCAAAGGGGGAGCGGACAAGTTGTCACTTACGCTCTTAATACATTCACCAACCTGGTGGTGCAGCTCATTCGGAATATGGAGGCTGAGGAAGTTCTAGAGATGCAAGACTTGTGGCTGCTGCGGAGGCCAGAGAAAGTGACCAACTGGTTGCAAAGCAACGGATGGGATAGGCTCAAAAGAATGGCAGTCAGTGGAGATGATTGCGTTGTGAAACCAATTGATGATAGGTTTGCACATGCCCTCAGGTTCTTGAATGATATGGGAAAAGTTAGGAAGGACACACAAGAGTGGAAACCCTCAACTGGATGGGACAACTGGGAAGAAGTTCCGTTTTGCTCCCACCACTTCAACAAACTCCATCTTAAGGACGGGAGGTCCATTGTGGTTCCCTGCCGCCACCAAGATGAACTGATTGGCCGAGCCCGCGTATCACCAGGGGCGGGATGGAGCATCCGGGAGACTGCTTGCCTAGCAAAATCATATGCGCAAATGTGGCAGCTCCTTTATTTCCACAGAAGGGACCTCCGACTGATGGCCAATGCCATTTGTTCATCTGTGCCAGTTGATTGGGTTCCAACTGGGAGAACTACCTGGTCAATCCATGGAAAGGGAGAATGGATGACCACTGAAGACATGCTTGTGGTATGGAACAGAGTGTGGATTGAGGAAAACGACCACATGGAAGACAAGACCCCAGTTACAAAATGGACAGACATTCCCTATTTGGGAAAAAGAGAAGACTTGTGGTGTGGATCTCTCATAGGGCACAGACCGCGTACTACCTGGGCTGAGAACATCAAAAATACAGTCAACATGATGCGCAGGATCATAGGTGATGAAGAAAAGTACATGGACTACCTATCCACCCAGGTTCGCTACTTGGGTGAAGAAGGGTCCACACCTGGAGTGCTGTAA

>ENA|AHZ13508|AHZ13508.1 Zika virus polyprotein

ATGAAAAACCC-AAAAAAGAAATCCGGAGGATTCCGGATTGTCAATATGCTAAAACGCGGAGTAGCCCGTGTGAGCCCCTTTGGGGGCTTGAAGAGGCTGCCAGCCGGACTTCTGCTGGGTCATGGGCCCATCAGGATGGTCTTGGCGATTCTAGCCTTTTTGAGATTCACGGCAATCAAGCCATCACTGGGTCTCATCAATAGATGGGGTTCAGTGGGGAAAAAAGAGGCTATGGAAATAATAAAGAAGTTCAAGAAAGATCTGGCTGCCATGCTGAGAATAATCAATGCTAGGAAGGAGAAGAAGAGACGAGGCGCAGATACTAGTGTCGGAATTGTTGGCCTCCTGCTGACCACAGCTATGGCAGCGGAGGTCACTAGACGTGGGAGTGCATACTATATGTACTTGGACAGAAACGACGCTGGGGAGGCCATATCTTTTCCAACCACATTGGGGATGAATAAGTGTTATATACAGATCATGGATCTTGGACACATGTGTGATGCCACCATGAGCTATGAATGCCCTATGCTGGATGAGGGGGTGGAACCAGATGACGTCGATTGTTGGTGCAACACGACGTCAACTTGGGTTGTGTACGGAACCTGCCATCACAAAAAAGGTGAAGCACGGAGATCTAGAAGAGCTGTGACGCTCCCCTCCCATTCCACTAGGAAGCTGCAAACGCGGTCGCAAACCTGGTTGGAATCAAGAGAATACACAAAGCACTTGATTAGAGTCGAAAATTGGATATTCAGGAACCCTGGCTTCGCGTTAGCAGCAGCTGCCATCGCTTGGCTTTTGGGAAGCTCAACGAGCCAAAAAGTCATATACTTGGTCATGATACTGCTGATTGCCCCGGCATACAGCATCAGGTGCATAGGAGTCAGCAATAGGGACTTTGTGGAAGGTATGTCAGGTGGGACTTGGGTTGATGTTGTCTTGGAACATGGAGGTTGTGTCACCGTAATGGCACAGGACAAACCGACTGTCGACATAGAGCTGGTTACAACAACAGTCAGCAACATGGCGGAGGTAAGATCCTACTGCTATGAGGCATCAATATCGGACATGGCTTCGGACAGCCGCTGCCCAACACAAGGTGAAGCCTACCTTGACAAGCAATCAGACACTCAATATGTCTGCAAAAGAACGTTAGTGGACAGAGGCTGGGGAAATGGATGTGGACTTTTTGGCAAAGGGAGCCTGGTGACATGCGCTAAGTTTGCATGCTCCAAGAAAATGACCGGGAAGAGCATCCAGCCAGAGAATCTGGAGTACCGGATAATGCTGTCAGTTCATGGCTCCCAGCACAGTGGGATGATCGTTAATGACACAGGACATGAAACTGATGAGAATAGAGCGAAGGTTGAGATAACGCCCAATTCACCAAGAGCCGAAGCCACCCTGGGGGGTTTTGGAAGCCTAGGACTTGATTGTGAACCGAGGACAGGCCTTGACTTTTCAGATTTGTATTACTTGACTATGAATAACAAGCACTGGTTGGTTCACAAGGAGTGGTTCCACGACATTCCATTACCTTGGCACGCTGGGGCAGACACCGGAACTCCACACTGGAACAACAAAGAAGCACTGGTAGAGTTCAAGGACGCACATGCCAAAAGGCAAACTGTCGTGGTTCTAGGGAGTCAAGAAGGAGCAGTTCACACGGCCCTTGCTGGAGCTCTGGAGGCTGAGATGGATGGTGCAAAGGGAAGGCTGTCCTCTGGCCACTTGAAATGTCGCCTGAAAATGGATAAACTTAGATTGAAGGGCGTGTCATACTCCTTGTGTACCGCAGCGTTCACATTCACCAAGATCCCGGCTGAAACACTGCACGGGACAGTCACAGTGGAGGTACAGTACGCAGGGACAGATGGACCTTGCAAGGTTCCAGCTCAGATGGCGGTGGACATGCAAACTCTGACCCCAGTTGGGAGGTTGATAACCGCTAACCCCGTAATCACTGAAAGCACTGAGAACTCTAAGATGATGCTGGAACTTGATCCACCATTTGGGGACTCTTACATTGTCATAGGAGTCGGGGAGAAGAAGATCACCCACCACTGGCACAGGAGTGGCAGCACCATTGGAAAAGCATTTGAAGCCACTGTGAGAGGTGCCAAGAGAATGGCAGTCTTGGGAGACACAGCCTGGGACTTTGGATCAGTTGGAGGCGCTCTCAACTCATTGGGCAAGGGCATCCATCAAATTTTTGGAGCAGCTTTCAAATCATTGTTTGGAGGAATGTCCTGGTTCTCACAAATTCTCATTGGAACGTTGCTGATGTGGTTGGGTCTGAACACAAAGAATGGATCTATTTCCCTTATGTGCTTGGCCTTAGGGGGAGTGTTGATCTTCTTATCCACAGCTGTCTCTGCTGATGTGGGGTGCTCGGTGGACTTCTCAAAGAAGGAGACGAGATGCGGTACAGGGGTGTTCGTCTATAACGACGTTGAAGCCTGGAGGGACAGGTACAAGTACCATCCTGACTCCCCCCGTAGATTGGCAGCAGCAGTCAAGCAAGCCTGGGAAGATGGTATCTGTGGGATCTCCTCTGTTTCAAGAATGGAAAACATCATGTGGAGATCAGTAGAAGGGGAGCTCAACGCAATCCTGGAAGAGAATGGAGTTCAACTGACGGTCGTTGTGGGATCTGTAAAAAACCCCATGTGGAGAGGTCCACAGAGATTGCCCGTGCCTGTGAACGAGCTGCCCCACGGCTGGAAGGCTTGGGGGAAATCGTACTTCGTCAGAGCAGCAAAGACAAATAACAGCTTTGTCGTGGATGGTGACACACTGAAGGAATGCCCACTCAAACATAGAGCATGGAACAGCTTTCTTGTGGAGGATCATGGGTTCGGGGTATTTCACACTAGTGTCTGGCTCAAGGTTAGAGAAGATTATTCATTAGAGTGTGATCCAGCCGTTATTGGAACAGCTGTTAAGGGAAAGGAGGCTGTACACAGTGATCTAGGCTACTGGATTGAGAGTGAGAAGAATGACACATGGAGGCTGAAGAGGGCCCATCTGATCGAGATGAAAACATGTGAATGGCCAAAGTCCCACACATTGTGGACAGATGGAATAGAAGAGAGTGATCTGATCATACCCAAGTCTTTAGCTGGGCCACTCAGCCATCACAATACCAGAGAGGGCTACAGGACCCAAATGAAAGGGCCATGGCACAGTGAAGAGCTTGAAATTCGGTTTGAGGAATGCCCAGGCACTAAGGTCCACGTGGAGGAAACATGTGGAACAAGAGGACCATCTCTGAGATCAACCACTGCAAGCGGAAGGGTGATCGAGGAATGGTGCTGCAGGGAGTGCACAATGCCCCCACTGTCGTTCCGGGCTAAAGATGGCTGTTGGTATGGAATGGAGATAAGGCCCAGGAAAGAACCAGAAAGTAACTTAGTAAGGTCAATGGTGACTGCAGGATCAACTGATCACATGGATCACTTCTCCCTTGGAGTGCTTGTGATTCTGCTCATGGTGCAGGAAGGGCTGAAGAAGAGAATGACCACAAAGATCATCATAAGCACATCGATGGCAGTGCTGGTAGCTATGATCCTGGGAGGATTTTCAATGAGTGACCTGGCTAAGCTTGCAATTTTGATGGGTGCCACCTTCGCGGAAATGAACACTGGAGGAGATGTAGCTCATCTGGCGCTGATAGCGGCATTCAAAGTCAGACCAGCGTTGCTGGTATCTTTCATCTTCAGAGCTAATTGGACACCCCGTGAAAGCATGCTGCTGGCCTTGGCCTCGTGTCTTTTGCAAACTGCGATCTCCGCCTTGGAAGGCGACCTGATGGTTCTCATCAATGGTTTTGCTTTGGCCTGGTTGGCAATACGAGCGATGGTTGTTCCACGCACTGATAACATCACCTTGGCAATCCTGGCTGCTCTGACACCACTGGCCCGGGGCACACTGCTTGTGGCGTGGAGAGCAGGCCTTGCTACTTGCGGGGGGTTTATGCTCCTCTCTCTGAAGGGAAAAGGCAGTGTGAAGAAGAACTTACCATTTGTCATGGCCCTGGGACTAACCGCTGTGAGGCTGGTCGACCCCATCAACGTGGTGGGACTGCTGTTGCTCACAAGGAGTGGGAAGCGGAGCTGGCCCCCTAGCGAAGTACTCACAGCTGTTGGCCTGATATGCGCATTGGCTGGAGGGTTCGCCAAGGCAGATATAGAGATGGCTGGGCCCATGGCCGCGGTCGGTCTGCTAATTGTCAGTTACGTGGTCTCAGGAAAGAGTGTGGACATGTACATTGAAAGAGCAGGTGACATCACATGGGAAAAAGATGCGGAAGTCACTGGAAACAGTCCCCGGCTCGATGTGGCGCTAGATGAGAGTGGTGATTTCTCCCTGGTGGAGGATGACGGTCCCCCCATGAGAGAGATCATACTCAAGGTGGTCCTGATGACCATCTGTGGCATGAACCCAATAGCCATACCCTTTGCAGCTGGAGCGTGGTACGTATACGTGAAGACTGGAAAAAGGAGTGGTGCTCTATGGGATGTGCCTGCTCCCAAGGAAGTAAAAAAGGGGGAGACCACAGATGGAGTGTACAGAGTAATGACTCGTAGACTGCTAGGTTCAACACAAGTTGGAGTGGGAGTTATGCAAGAGGGGGTCTTTCACACTATGTGGCACGTCACAAAAGGATCCGCGCTGAGAAGCGGTGAAGGGAGACTTGATCCATACTGGGGAGATGTCAAGCAGGATCTGGTGTCATACTGTGGTCCATGGAAGCTAGATGCCGCCTGGGACGGGCACAGCGAGGTGCAGCTCTTGGCCGTGCCCCCCGGAGAGAGAGCGAGGAACATCCAGACTCTGCCCGGAATATTTAAGACAAAGGATGGGGACATTGGAGCGGTTGCGCTGGATTACCCAGCAGGAACTTCAGGATCTCCAATCCTAGACAAGTGTGGGAGAGTGATAGGACTTTATGGCAATGGGGTCGTGATCAAAAATGGGAGTTATGTTAGTGCCATCACCCAAGGGAGGAGGGAGGAAGAGACTCCTGTTGAGTGCTTCGAGCCTTCGATGCTGAAGAAGAAGCAGCTAACTGTCTTAGACTTGCATCCTGGAGCTGGGAAAACCAGGAGAGTTCTTCCTGAAATAGTCCGTGAAGCCATAAAAACAAGACTCCGTACTGTGATCTTAGCTCCAACCAGGGTTGTCGCTGCTGAAATGGAGGAAGCCCTTAGAGGGCTTCCAGTGCGTTATATGACAACAGCAGTCAATGTCACCCACTCTGGAACAGAAATCGTCGACTTAATGTGCCATGCCACCTTCACTTCACGTCTACTACAGCCAATCAGAGTCCCCAACTATAATCTGTATATTATGGATGAGGCCCACTTCACAGATCCCTCAAGTATAGCAGCAAGAGGATACATTTCAACAAGGGTTGAGATGGGCGAGGCGGCTGCCATCTTCATGACCGCCACGCCACCAGGAACCCGTGACGCATTTCCGGACTCCAACTCACCAATTATGGACACCGAAGTGGAAGTCCCAGAGAGAGCCTGGAGCTCAGGCTTTGATTGGGTGACGGATCATTCTGGAAAAACAGTTTGGTTTGTTCCAAGCGTGAGGAACGGCAATGAGATCGCAGCTTGTCTGACAAAGGCTGGAAAACGGGTCATACAGCTCAGCAGAAAGACTTTTGAGACAGAGTTCCAGAAAACAAAACATCAAGAGTGGGACTTTGTCGTGACAACTGACATTTCAGAGATGGGCGCCAACTTTAAAGCTGACCGTGTCATAGATTCCAGGAGATGCCTAAAGCCGGTCATACTTGATGGCGAGAGAGTCATTCTGGCTGGACCCATGCCTGTCACACATGCCAGCGCTGCCCAGAGGAGGGGGCGCATAGGCAGGAATCCCAACAAACCTGGAGATGAGTATCTGTATGGAGGTGGGTGCGCAGAGACTGACGAAGACCATGCACACTGGCTTGAAGCAAGAATGCTCCTTGACAATATTTACCTCCAAGATGGCCTCATAGCCTCGCTCTATCGACCTGAGGCCGACAAAGTAGCAGCCATTGAGGGAGAGTTCAAGCTTAGGACGGAGCAAAGGAAGACCTTTGTGGAACTCATGAAAAGAGGAGATCTTCCTGTTTGGCTGGCCTATCAGGTTGCATCTGCCGGAATAACCTACACAGATAGAAGATGGTGCTTTGATGGCACGACCAACAACACCATAATGGAAGACAGTGTGCCGGCAGAGGTGTGGACCAGACACGGAGAGAAAAGAGTGCTCAAACCGAGGTGGATGGACGCCAGAGTTTGTTCAGATCATGCGGCCCTGAAGTCATTCAAGGAGTTTGCCGCTGGGAAAAGAGGAGCGGCTTTTGGAGTGATGGAAGCCCTGGGAACACTGCCAGGACACATGACAGAGAGATTCCAGGAAGCCATTGACAACCTCGCTGTGCTCATGCGGGCAGAGACTGGAAGCAGGCCTTACAAAGCCGCGGCGGCCCAATTGCCGGAGACCCTAGAGACCATTATGCTTTTGGGGTTGCTGGGAACAGTCTCGCTGGGAATCTTTTTCGTCTTGATGAGGAACAAGGGCATAGGGAAGATGGGCTTTGGAATGGTGACTCTTGGGGCCAGCGCATGGCTCATGTGGCTCTCGGAAATTGAGCCAGCCAGAATTGCATGTGTCCTCATTGTTGTGTTCCTATTGCTGGTGGTGCTCATACCTGAGCCAGAAAAGCAAAGATCTCCCCAGGACAACCAAATGGCAATCATCATCATGGTAGCAGTAGGTCTTCTGGGCTTGATTACCGCCAATGAACTCGGATGGTTGGAGAGAACAAAGAGTGACCTAAGCCATCTAATGGGAAGGAGAGAGGAGGGGGCAACCATAGGATTCTCAATGGACATTGACCTGCGGCCAGCCTCAGCTTGGGCCATCTATGCTGCCTTGACAACTTTCATTACCCCAGCCGTCCAACATGCAGTGACCACTTCATACAACAACTACTCCTTAATGGCGATGGCCACGCAAGCTGGAGTGTTGTTTGGTATGGGCAAAGGGATGCCATTCTACGCATGGG-ACTTTGGAGTCCCGCTGCTAATGATAGGTTGCTACTCACAATTAACACCCCTGACCCTAATAGTGGCCATCATTTTGCTCGTGGCGCACTACATGTACTTGATCCCAGGGCTGCAGGCAGCAGCTGCGCGTGCTGCCCAGAAGAGAACGGCAGCTGGCATCATGAAGAACCCTGTTGTGGATGGAATAGTGGTGACTGACATTGACACAATGACAATTGACCCCCAAGTGGAGAAAAAGATGGGACAGGTGCTACTCATAGCAGTAGCCGTCTCCAGCGCCATACTGTCGCGGACCGCCTGGGGGTGGGGGGAGGCTGGGGCCCTGATCACAGCGGCAACTTCCACTTTGTGGGAAGGCTCTCCGAACAAGTACTGGAACTCCTCTACAGCCACTTCACTGTGTAACATTTTTAGGGGAAGTTACTTGGCTGGAGCTTCTCTAATCTACACAGTAACAAGAAACGCTGGCTTGGTCAAGAGACGTGGGGGTGGAACAGGAGAGACCCTGGGAGAGAAATGGAAGGCCCGCTTGAACCAGATGTCGGCCCTGGAGTTCTACTCCTACAAAAAGTCAGGCATCACCGAGGTGTGCAGAGAAGAGGCCCGCCGCGCCCTCAAGGACGGTGTGGCAACGGGAGGCCATGCTGTGTCCCGAGGAAGTGCAAAGCTGAGATGGTTGGTGGAGCGGGGATACCTGCAGCCCTATGGAAAGGTCATTGATCTTGGATGTGGCAGAGGGGGCTGGAGTTACTACGCCGCCACCATCCGCAAAGTTCAAGAAGTGAAAGGATACACAAAAGGAGGCCCTGGTCATGAAGAACCCATGTTGGTGCAAAGCTATGGGTGGAACATAGTCCGTCTTAAGAGTGGGGTGGACGTCTTTCATATGGCGGCTGAGCCGTGTGACACGTTGCTGTGTGACATAGGTGAGTCATCATCTAGTCCTGAAGTGGAAGAAGCACGGACGCTCAGAGTCCTCTCCATGGTGGGGGATTGGCTTGAAAAAAGACCAGGAGCCTTTTGTATAAAAGTGTTGTGCCCATACACCAGCACTATGATGGAAACCCTGGAGCGACTGCAGCGTAGGTATGGGGGAGGACTGGTCAGAGTGCCACTCTCCCGCAACTCTACACATGAGATGTACTGGGTCTCTGGAGCGAAAAGCAACACCATAAAAAGTGTGTCCACCACGAGCCAGCTCCTCTTGGGGCGCATGGACGGGCCCAGGAGGCCAGTGAAATATGAGGAGGATGTGAATCTCGGCTCTGGCACGCGGGCTGTGGTAAGCTGCGCTGAAGCTCCCAACATGAAGATCATTGGTAACCGCATTGAAAGGATCCGCAGTGAGCACGCGGAAACGTGGTTCTTTGACGAGAACCACCCATATAGGACATGGGCTTACCATGGAAGCTATGAGGCCCCCACACAAGGGTCAGCGTCCTCTCTAATAAACGGGGTTGTCAGGCTCCTGTCAAAACCCTGGGATGTGGTGACTGGAGTCACAGGAATAGCCATGACCGACACCACACCGTATGGTCAGCAAAGAGTTTTCAAGGAAAAAGTGGACACTAGGGTGCCAGACCCCCAAGAAGGCACTCGTCAGGTTATGAGCATGGTCTCTTCCTGGTTGTGGAAAGAGCTAGGCAAACACAAACGGCCACGAGTCTGTACCAAAGAAGAGTTCATCAACAAGGTTCGTAGCAATGCAGCATTAGGGGCAATATTTGAAGAGGAAAAAGAGTGGAAGACTGCAGTGGAAGCTGTGAACGATCCAAGGTTCTGGGCTCTAGTGGACAAGGAAAGAGAGCACCACCTGAGAGGAGAGTGCCAGAGTTGTGTGTACAACATGATGGGAAAAAGAGAAAAGAAACAAGGGGAATTTGGAAAGGCCAAGGGCAGCCGCGCCATCTGGTATATGTGGCTAGGGGCTAGATTTCTAGAGTTCGAAGCCCTTGGATTCTTGAACGAGGATCACTGGATGGGGAGAGAGAACTCAGGAGGTGGTGTTGAAGGGCTGGGATTACAAAGACTCGGATATGTCCTAGAAGAGATGAGTCGCATACCAGGAGGAAGGATGTATGCAGATGACACTGCTGGCTGGGACACCCGCATCAGCAGGTTTGATCTGGAGAATGAAGCTCTAATCACCAACCAAATGGAGAAAGGGCACAGGGCCTTGGCATTGGCCATAATCAAGTACACATACCAAAACAAAGTGGTAAAGGTCCTTAGACCAGCTGAAAAAGGGAAGACAGTTATGGACATTATTTCGAGACAAGACCAAAGGGGGAGCGGACAAGTTGTCACTTACGCTCTTAACACATTTACCAACCTAGTGGTGCAACTCATTCGGAATATGGAGGCTGAGGAAGTTCTAGAGATGCAAGACTTGTGGCTGCTGCGGAGGTCAGAGAAAGTGACCAACTGGTTGCAGAGCAACGGATGGGATAGGCTCAAACGAATGGCAGTCAGTGGAGATGATTGCGTTGTGAAGCCAATTGATGATAGGTTTGCACATGCCCTCAGGTTCTTGAATGATATGGGAAAAGTTAGGAAGGACACACAAGAGTGGAAACCCTCAACTGGATGGGACAACTGGGAAGAAGTTCCGTTTTGCTCCCACCACTTCAACAAGCTCCATCTCAAGGACGGGAGGTCCATTGTGGTTCCCTGCCGCCACCAAGATGAACTGATTGGCCGGGCCCGCGTCTCTCCAGGGGCGGGATGGAGCATCCGGGAGACTGCTTGCCTAGCAAAATCATATGCGCAAATGTGGCAGCTCCTTTATTTCCACAGAAGGGACCTCCGACTGATGGCCAATGCCATTTGTTCATCTGTGCCAGTTGACTGGGTTCCAACTGGGAGAACTACCTGGTCAATCCATGGAAAGGGAGAATGGATGACCACTGAAGACATGCTTGTGGTGTGGAACAGAGTGTGGATTGAGGAGAACGACCACATGGAAGACAAGACCCCAGTTACGAAATGGACAGACATTCCCTATTTGGGAAAAAGGGAAGACTTGTGGTGTGGATCTCTCATAGGGCACAGACCGCGCACCACCTGGGCTGAGAACATTAAAAACACAGTCAACATGGTGCGCAGGATCATAGGTGATGAAGAAAAGTACATGGACTACCTATCCACCCAAGTTCGCTACTTGGGTGAAGAAGGGTCTACACCTGGAGTGCTGTAA

>ENA|AMA12085|AMA12085.1 Zika virus polyprotein

ATGAAAAACCC-AAAAAAGAAATCCGGAGGATTCCGGATTGTCAATATGCTAAAACGCGGAGTAGCCCGTGTGAGCCCCTTTGGGGGCTTGAAGAGGCTGCCAGCCGGACTTCTGCTGGGTCATGGGCCCATCAGGATGGTCTTGGCGATTCTAGCCTTTTTGAGATTCACGGCAATCAAGCCATCACTGGGTCTCATCAATAGATGGGGTTCAGTTGGGAAAAAAGAGGCTATGGAAATAATAAAGAAGTTCAAGAAAGATCTGGCTGCCATGCTGAGAATAATCAATGCTAGGAAGGAGAAGAAGAGACGAGGCGCAGATACTAGTGTCGGAATTGTTGGCCTCCTGCTGACCACAGCTATGGCAGCGGAGGTCACTAGACGTGGGAGTGCATACTATATGTACTTGGACAGAAACGATGCTGGGGAGGCCATATCTTTTCCAACCACATTGGGGATGAATAAGTGTTATATACAGATCATGGATCTTGGACACATGTGTGATGCCACCATGAGCTATGAATGCCCTATGCTGGATGAGGGGGTGGAACCAGATGACGTCGATTGTTGGTGCAACACGACGTCAACTTGGGTTGTGTACGGAACCTGCCATCACAAAAAAGGTGAAGCACGGAGATCTAGAAGAGCTGTGACGCTCCCCTCCCATTCCACTAGGAAGCTGCAAACGCGGTCGCAAACCTGGTTGGAATCAAGAGAATACACAAAGCACTTGATTAGAGTCGAAAATTGGATATTCAGGAACCCTGGCTTCGCGTTAGCAGCAGCTGCCATCGCTTGGCTTTTGGGAAGCTCAACGAGCCAAAAAGTCATATACTTGGTCATGATACTGCTGATTGCCCCGGCATACAGCATCAGGTGCATAGGAGTCAGCAATAGGGACTTTGTGGAAGGTATGTCAGGTGGGACTTGGGTTGATGTTGTCTTGGAACATGGAGGTTGTGTCACCGTAATGGCACAGGACAAACCGACTGTCGACATAGAGCTGGTTACAACAACAGTCAGCAACATGGCGGAGGTAAGATCCTACTGCTATGAGGCATCAATATCAGACATGGCTTCGGACAGCCGCTGCCCAACACAAGGTGAAGCCTACCTTGACAAGCAATCAGACACTCAATATGTCTGCAAAAGAACGTTAGTGGACAGAGGCTGGGGAAATGGATGTGGACTTTTTGGCAAAGGGAGCCTGGTGACATGCGCTAAGTTTGCATGCTCCAAGAAAATGACCGGGAAGAGCATCCAGCCAGAGAATCTGGAGTACCGGATAATGCTGTCAGTTCATGGCTCCCAGCACAGTGGGATGATCGTTAATGACACAGGACATGAAACTGATGAGAATAGAGCGAAAGTTGAGATAACGCCCAATTCACCAAGAGCCGAAGCCACCCTGGGGGGTTTTGGAAGCCTAGGACTTGATTGTGAACCGAGGACAGGCCTTGACTTTTCAGATTTGTATTACTTGACTATGAATAACAAGCACTGGTTGGTTCACAAGGAGTGGTTCCACGACATTCCATTACCTTGGCACGCTGGGGCAGACACCGGAACTCCACACTGGAACAACAAAGAAGCACTGGTAGAGTTCAAGGACGCACATGCCAAAAGGCAAACTGTCGTGGTTCTAGGGAGTCAAGAAGGAGCAGTTCACACGGCCCTTGCTGGAGCTCTGGAGGCTGAGATGGATGGTGCAAAGGGAAGGCTGTCCTCTGGCCACTTGAAATGTCGCCTGAAAATGGATAAACTTAGATTGAAGGGCGTGTCATACTCCTTGTGTACTGCAGCGTTCACATTCACCAAGATCCCGGCTGAAACACTGCACGGGACAGTCACAGTGGAGGTACAGTACGCAGGGACAGATGGACCTTGCAAGGTTCCAGCTCAGATGGCGGTGGACATGCAAACTCTGACCCCAGTTGGGAGGTTGATAACCGCTAACCCCGTAATCACTGAAAGCACTGAGAACTCTAAGATGATGCTGGAACTTGATCCACCATTTGGGGACTCTTACATTGTCATAGGAGTCGGGGAGAAGAAGATCACCCACCACTGGCACAGGAGTGGCAGCACCATTGGAAAAGCATTTGAAGCCACTGTGAGAGGTGCCAAGAGAATGGCAGTCTTGGGAGACACAGCCTGGGACTTTGGATCAGTTGGAGGCGCTCTCAACTCATTGGGCAAGGGCATCCATCAAATTTTTGGAGCAGCTTTCAAATCATTGTTTGGAGGAATGTCCTGGTTCTCACAAATTCTCATTGGAACGTTGCTGATGTGGTTGGGTCTGAACACAAAGAATGGATCTATTTCCCTTATGTGCTTGGCCTTAGGGGGGGTGTTGATCTTCTTATCCACAGCCGTCTCTGCTGATGTGGGGTGCTCGGTGGACTTCTCAAAGAAGGAGACGAGATGCGGTACAGGGGTGTTCGTCTATAACGACGTTGAAGCCTGGAGGGACAGGTACAAGTACCATCCTGACTCCCCCCGTAGATTGGCAGCAGCAGTCAAGCAAGCCTGGGAAGATGGTATCTGCGGGATCTCCTCTGTTTCAAGAATGGAAAACATCATGTGGAGATCAGTAGAAGGGGAGCTCAACGCAATCCTGGAAGAGAATGGAGTTCAACTGACGGTCGTTGTGGGATCTGTAAAAAACCCCATGTGGAGAGGTCCACAGAGATTGCCCGTGCCTGTGAACGAGCTGCCCCACGGCTGGAAGGCTTGGGGGAAATCGTACTTCGTCAGAGCAGCAAAGACAAATAACAGCTTTGTCGTGGATGGTGACACACTGAAGGAATGCCCACTCAAACATAGAGCATGGAACAGCTTTCTTGTGGAGGATCATGGGTTCGGGGTATTTCACACTAGTGTCTGGCTCAAGGTTAGAGAAGATTATTCATTAGAGTGTGATCCAGCCGTTATTGGAACAGCTGTTAAGGGAAAGGAGGCTGTACACAGTGATCTAGGCTACTGGATTGAGAGTGAGAAGAATGACACATGGAGGCTGAAGAGGGCCCATCTGATCGAGATGAAAACATGTGAATGGCCAAAGTCCCACACATTGTGGACAGATGGAATAGAAGAGAGTGATCTGATCATACCCAAGTCTTTAGCTGGGCCACTCAGCCATCACAATACCAGAGAGGGCTACAGGACCCAAATGAAAGGGCCATGGCACAGTGAAGAGCTTGAAATTCGGTTTGAGGAATGCCCAGGCACTAAGGTCCACGTGGAGGAAACATGTGGAACAAGAGGACCATCTCTGAGATCAACCACTGCAAGCGGAAGGGTGATCGAGGAATGGTGCTGCAGGGAGTGCACAATGCCCCCACTGTCGTTCCGGGCTAAAGATGGCTGTTGGTATGGAATGGAGATAAGGCCCAGGAAAGAACCAGAAAGCAACTTAGTAAGGTCAATGGTGACTGCAGGATCAACTGATCACATGGACCACTTCTCCCTTGGAGTGCTTGTGATCCTGCTCATGGTGCAGGAAGGGCTGAAGAAGAGAATGACCACAAAGATCATCATAAGCACATCAATGGCAGTGCTGGTAGCTATGATCCTGGGAGGATTTTCAATGAGTGACCTGGCTAAGCTTGCAATTTTGATGGGTGCCACCTTCGCGGAAATGAACACTGGAGGAGATGTAGCTCATCTGGCGCTGATAGCGGCATTCAAAGTCAGACCAGCGTTGCTGGTATCTTTCATCTTCAGAGCTAATTGGACACCCCGTGAAAGCATGCTGCTGGCCTTGGCCTCGTGTCTTTTGCAAACTGCGATCTCCGCCTTGGAAGGCGACCTGATGGTTCTCATCAATGGTTTTGCTTTGGCCTGGTTGGCAATACGAGCGATGGTTGTTCCACGCACTGATAACATCACCTTGGCAATCCTGGCTGCTCTGACACCACTGGCCCGGGGCACACTGCTTGTGGCGTGGAGAGCAGGCCTTGCTACTTGCGGGGGGTTTATGCTCCTCTCTCTGAAGGGAAAAGGCAGTGTGAAGAAGAACTTACCATTTGTCATGGCCCTGGGACTAACCGCTGTGAGGCTGGTCGACCCCATCAACGTGGTGGGACTGCTGTTACTCACAAGGAGTGGGAAGCGGAGCTGGCCCCCTAGCGAAGTACTCACAGCTGTTGGCCTGATATGCGCATTGGCTGGAGGGTTCGCCAAGGCAGATATAGAGATGGCTGGGCCCATGGCCGCGGTCGGTCTGCTAATTGTCAGTTACGTGGTCTCAGGAAAGAGTGTGGACATGTACATTGAAAGAGCAGGTGACATCACATGGGAAAAAGATGCGGAAGTCACTGGAAACAGTCCCCGGCTCGATGTGGCGCTAGATGAGAGTGGTGATTTCTCCCTGGTGGAGGATGACGGTCCCCCCATGAGAGAGATCATACTCAAGGTGGTCCTGATGACCATCTGTGGCATGAATCCAATAGCCATACCCTTTGCAGCTGGAGCGTGGTACGTATACGTGAAGACTGGAAAAAGGAGTGGTGCTCTATGGGATGTGCCTGCTCCCAAGGAAGTAAAAAAGGGGGAGACCACAGATGGAGTGTACAGAGTAATGACTCGTAGACTGCTAGGTTCAACACAAGTTGGAGTGGGAGTTATGCAAGAGGGGGTCTTTCACACTATGTGGCACGTCACAAAAGGATCCGCGCTGAGAAGCGGTGAAGGGAGACTTGATCCATACTGGGGAGATGTCAAGCAGGATCTGGTGTCATACTGTGGTCCATGGAAGCTAGATGCCGCCTGGGACGGGCACAGCGAGGTGCAGCTCTTGGCCGTGCCCCCCGGAGAGAGAGCGAGGAACATCCAGACTCTGCCCGGAATATTTAAGACAAAGGATGGGGACATTGGAGCGGTTGCGCTGGATTACCCAGCAGGAACTTCAGGATCTCCAATCCTAGACAAGTGTGGGAGAGTGATAGGACTTTATGGCAATGGGGTCGTGATCAAAAATGGGAGTTATGTTAGTGCCATCACCCAAGGGAGGAGGGAAGAAGAGACTCCTGTTGAGTGCTTCGAGCCCTCGATGCTGAAGAAGAAGCAGCTAACTGTCTTAGACTTGCATCCTGGAGCTGGGAAAACCAGGAGAGTTCTTCCTGAAATAGTCCGTGAAGCCATAAAAACAAGACTCCGTACTGTGATCTTAGCTCCAACCAGGGTTGTCGCTGCTGAAATGGAGGAGGCCCTTAGAGGGCTTCCAGTGCGTTATATGACAACAGCAGTCAATGTCACCCACTCTGGAACAGAAATCGTCGACTTAATGTGCCATGCCACCTTCACTTCACGTCTACTACAGCCAATCAGAGTCCCCAACTATAATCTGTATATTATGGATGAGGCCCACTTCACAGATCCCTCAAGTATAGCAGCAAGAGGATACATTTCAACAAGGGTTGAGATGGGCGAGGCGGCTGCCATCTTCATGACCGCCACGCCACCAGGAACCCGTGACGCATTTCCGGACTCCAACTCACCAATTATGGACACCGAAGTGGAAGTCCCAGAGAGAGCCTGGAGCTCAGGCTTTGATTGGGTGACGGATCATTCTGGAAAAACAGTTTGGTTTGTTCCAAGCGTGAGGAACGGCAATGAGATCGCAGCTTGTCTGACAAAGGCTGGAAAACGGGTCATACAGCTCAGCAGAAAGACTTTTGAGACAGAGTTCCAGAAAACAAAACATCAAGAGTGGGACTTTGTCGTGACAACTGACATTTCAGAGATGGGCGCCAACTTTAAAGCTGACCGTGTCATAGATTCCAGGAGATGCCTAAAGCCGGTCATACTTGATGGCGAGAGAGTCATTCTGGCTGGACCCATGCCTGTCACACATGCCAGCGCTGCCCAGAGGAGGGGGCGCATAGGCAGGAATCCCAACAAACCTGGAGATGAGTATCTGTATGGAGGTGGGTGCGCAGAGACTGACGAAGACCATGCACACTGGCTTGAAGCAAGAATGCTCCTTGACAATATTTACCTCCAAGATGGCCTCATAGCCTCGCTCTATCGACCTGAGGCCGACAAAGTAGCAGCCATTGAGGGAGAGTTCAAGCTTAGGACGGAGCAAAGGAAGACCTTTGTGGAACTCATGAAAAGAGGAGATCTTCCTGTTTGGCTGGCCTATCAGGTTGCATCTGCCGGAATAACCTACACAGATAGAAGATGGTGCTTTGATGGCACGACCAACAACACCATAATGGAAGATAGTGTGCCGGCAGAGGTGTGGACCAGACACGGAGAGAAAAGAGTGCTCAAACCGAGGTGGATGGACGCCAGAGTTTGTTCAGATCATGCGGCCCTGAAGTCATTCAAGGAGTTTGCCGCTGGGAAAAGAGGAGCGGCTTTTGGAGTGATGGAAGCCCTGGGAACACTGCCAGGACACATGACAGAGAGATTCCAGGAAGCCATTGACAACCTCGCTGTGCTCATGCGGGCAGAGACTGGAAGCAGGCCTTACAAAGCCGCGGCGGCCCAATTGCCGGAGACCCTAGAGACCATAATGCTTTTGGGGTTGCTGGGAACAGTCTCGCTGGGAATCTTCTTCGTCTTGATGAGGAACAAGGGCATAGGGAAGATGGGCTTTGGAATGGTGACTCTTGGGGCCAGCGCATGGCTCATGTGGCTCTCGGAAATTGAGCCAGCCAGAATTGCATGTGTCCTCATTGTTGTGTTCCTATTGCTGGTGGTGCTCATACCTGAGCCAGAAAAGCAAAGATCTCCCCAGGACAACCAAATGGCAATCATCATCATGGTAGCAGTAGGTCTTCTGGGCTTGATTACCGCCAATGAACTCGGATGGTTGGAGAGAACAAAGAGTGACCTAAGCCATCTAATGGGAAGGAGAGAGGAGGGGGCAACCATAGGATTCTCAATGGACATTGACCTGCGGCCAGCCTCAGCTTGGGCCATCTATGCTGCCTTGACAACTTTCATTACCCCAGCCGTCCAACATGCAGTGACCACCTCATACAACAACTACTCCTTAATGGCGATGGCCACGCAAGCTGGAGTGTTGTTTGGTATGGGCAAAGGGATGCCATTCTACGCATGGG-ACTTTGGAGTCCCGCTGCTAATGATAGGTTGCTACTCACAATTAACACCCCTGACCCTAATAGTGGCCATCATTTTGCTCGTGGCGCACTACATGTACTTGATCCCAGGGCTGCAGGCAGCAGCTGCGCGTGCTGCCCAGAAGAGAACGGCAGCTGGCATCATGAAGAACCCTGTTGTGGATGGAATAGTGGTGACTGACATTGACACAATGACAATTGACCCCCAAGTGGAGAAAAAGATGGGACAGGTGCTACTCATAGCAGTAGCCGTCTCCAGCGCCATACTGTCGCGGACCGCCTGGGGGTGGGGGGAGGCTGGGGCCCTGATCACAGCCGCAACTTCCACTTTGTGGGAAGGCTCTCCGAACAAGTACTGGAACTCCTCTACAGCCACTTCACTGTGTAACATTTTTAGGGGAAGTTACTTGGCTGGAGCTTCTCTAATCTACACAGTAACAAGAAACGCTGGCTTGGTCAAGAGACGTGGGGGTGGAACAGGAGAGACCCTGGGAGAGAAATGGAAGGCCCGCTTGAACCAGATGTCGGCCCTGGAGTTCTACTCCTACAAAAAGTCAGGCATCACCGAGGTGTGCAGAGAAGAGGCCCGCCGCGCCCTCAAGGACGGTGTGGCAACGGGAGGCCATGCTGTGTCCCGAGGAAGTGCAAAGCTGAGATGGTTGGTGGAGCGGGGATACCTGCAGCCCTATGGAAAGGTCATTGATCTTGGATGTGGCAGAGGGGGCTGGAGTTACTACGCCGCCACCATCCGCAAAGTTCAAGAAGTGAAAGGATACACAAAAGGAGGCCCTGGTCATGAAGAACCCGTGTTGGTGCAAAGCTATGGGTGGAACATAGTCCGTCTTAAGAGTGGGGTGGACGTCTTTCATATGGCGGCTGAGCCGTGTGACACGCTGCTGTGTGACATAGGTGAGTCATCATCTAGTCCTGAAGTGGAAGAAGCACGGACGCTCAGAGTCCTCTCCATGGTGGGGGATTGGCTTGAAAAAAGACCAGGAGCCTTTTGTATAAAAGTGTTGTGCCCATACACCAGCACTATGATGGAAACCCTGGAGCGACTGCAGCGTAGGTATGGGGGAGGACTGGTCAGAGTGCCACTCTCCCGCAACTCTACACATGAGATGTACTGGGTCTCTGGAGCGAAAAGCAACACCATAAAAAGTGTGTCCACCACGAGCCAGCTCCTCTTGGGGCGCATGGACGGGCCTAGGAGGCCAGTGAAATATGAGGAGGATGTGAATCTCGGCTCTGGCACGCGGGCTGTGGTAAGCTGCGCTGAAGCTCCCAACATGAAGATCATTGGTAACCGCATTGAAAGGATCCGCAGTGAGCACGCGGAAACGTGGTTCTTTGACGAGAACCACCCATATAGGACATGGGCTTACCATGGAAGCTATGAGGCCCCCACACAAGGGTCAGCGTCCTCTCTAATAAACGGGGTTGTCAGGCTCCTGTCAAAACCCTGGGATGTGGTGACTGGAGTCACAGGAATAGCCATGACCGACACCACACCGTATGGTCAGCAAAGAGTTTTCAAGGAAAAAGTGGACACTAGGGTGCCAGACCCCCAAGAAGGCACTCGTCAGGTTATGAGCATGGTCTCTTCCTGGTTGTGGAAAGAGCTAGGCAAACACAAACGGCCACGAGTCTGTACCAAAGAAGAGTTCATCAACAAGGTTCGTAGCAATGCAGCATTAGGGGCAATATTTGAAGAGGAAAAAGAGTGGAAGACTGCAGTGGAAGCTGTGAACGATCCAAGGTTCTGGGCTCTAGTGGACAAGGAAAGAGAGCACCACCTGAGAGGAGAGTGCCAGAGTTGTGTGTATAACATGATGGGAAAAAGAGAAAAGAAACAAGGGGAATTTGGAAAGGCCAAGGGCAGCCGCGCCATCTGGTATATGTGGCTAGGGGCTAGATTTCTAGAGTTCGAAGCCCTTGGATTCTTGAACGAGGATCACTGGATGGGGAGAGAGAACTCAGGAGGTGGTGTTGAAGGGCTGGGATTACAAAGACTCGGATATGTCCTAGAAGAGATGAGTCGTATACCAGGAGGAAGGATGTATGCAGATGACACTGCTGGCTGGGACACCCGCATTAGCAGGTTTGATCTGGAGAATGAAGCTCTAATCACCAACCAAATGGAGAAAGGGCACAGGGCCTTGGCATTGGCCATAATCAAGTACACATACCAAAACAAAGTGGTAAAGGTCCTTAGACCAGCTGAAAAAGGGAAAACAGTTATGGACATTATTTCGAGACAAGACCAAAGGGGGAGCGGACAAGTTGTCACTTACGCTCTTAACACATTTACCAACCTAGTGGTGCAACTCATTCGGAATATGGAGGCTGAGGAAGTTCTAGAGATGCAAGACTTGTGGCTGCTGCGGAGGTCAGAGAAAGTGACCAACTGGTTGCAGAGCAACGGATGGGATAGGCTCAAACGAATGGCAGTCAGTGGAGATGATTGCGTTGTGAAGCCAATTGATGATAGGTTTGCACATGCCCTCAGGTTCTTGAATGATATGGGAAAAGTTAGAAAGGACACACAAGAGTGGAAACCCTCAACTGGATGGGACAACTGGGAAGAAGTTCCGTTTTGCTCCCACCACTTCAACAAGCTCCATCTCAAGGACGGGAGGTCCATTGTGGTTCCCTGCCGCCACCAAGATGAACTGATTGGCCGGGCCCGCGTCTCTCCAGGGGCGGGATGGAGCATCCGGGAGACTGCTTGCCTAGCAAAATCATATGCGCAGATGTGGCAGCTCCTTTATTTCCACAGAAGGGACCTCCGACTGATGGCCAATGCCATTTGTTCATCTGTGCCAGTTGACTGGGTTCCAACTGGGAGAACTACCTGGTCAATCCATGGAAAGGGAGAATGGATGACCACTGAAGACATGCTTGTGGTGTGGAACAGAGTGTGGATTGAGGAGAACGACCACATGGAAGACAAGACCCCAGTTACGAAATGGACAGACATTCCCTATTTGGGAAAAAGGGAAGACTTGTGGTGTGGATCTCTCATAGGGCACAGACCGCGCACCACCTGGGCTGAGAACATTAAAAACACAGTCAACATGGTGCGCAGGATCATAGGTGATGAAGAAAAGTACATGGACTACCTATCCACCCAAGTTCGCTACTTGGGTGAAGAAGGGTCTACACCTGGAGTGCTGTAA

>ENA|ACD75819|ACD75819.1 Zika virus polyprotein

ATGAAAAACCCCAAAGAAGAAATCCGGAGGAT-CCGGATTGTCAATATGCTAAAACGCGGAGTAGCCCGTGTGAGCCCCTTTGGGGGCTTGAAGAGGCTGCCAGCCGGACTTCTGCTGGGTCATGGGCCCATCAGGATGGTCTTGGCGATACTAGCCTTTTTGAGATTCACGGCAATCAAGCCATCACTGGGTCTCATCAATAGATGGGGTTCAGTGGGGAAAAAAGAGGCTATGGAAATAATAAAGAAGTTCAAGAAAGATCTGGCTGCCATGCTGAGAATAATCAATGCTAGGAAGGAGAAGAAGAGACGAGGCACAGATACTAGTGTCGGAATTGTTGGCCTCCTGCTGACCACAGCCATGGCAGTGGAGGTCACTAGACGTGGGAGTGCATACTATATGTACTTGGACAGAAGCGATGCTGGGGAGGCCATATCTTTTCCAACCACACTGGGGATGAACAAGTGTTACATACAGATCATGGATCTTGGACACATGTGTGATGCCACCATGAGCTATGAATGCCCTATGTTGGATGAGGGGGTAGAACCAGATGACGTCGATTGTTGGTGCAACACGACATCAACTTGGGTTGTGTACGGAACCTGCCACCACAAAAAAGGTGAAGCACGGAGATCTAGAAGAGCTGTGACGCTCCCCTCCCATTCCACTAGGAAGCTGCAAACGCGGTCGCAGACCTGGTTGGAATCAAGAGAATATACAAAGCACCTGATTAGAGTCGAAAATTGGATATTCAGGAACCCTGGCTTCGCGTTAGCAGCAGCTGCCATCGCCTGGCTTTTGGGAAGTTCAACGAGCCAAAAAGTCATATACTTGGTCATGATACTGCTGATTGCCCCGGCATACAGCATCAGGTGCATAGGAGTCAGCAATAGGGACTTTGTGGAAGGTATGTCAGGTGGGACTTGGGTTGATGTTGTCTTGGAACATGGAGGTTGTGTTACCGTAATGGCACAGGACAAACCGGCTGTCGACATAGAGCTGGTTACAACAACAGTCAGCAACATGGCGGAGGTAAGATCCTATTGCTATGAGGCATCAATATCGGACATGGCTTCGGACAGCCGCTGCCCAACACAAGGTGAAGCCTACCTTGACAAGCAGTCAGACACTCAATATGTCTGCAAAAGAACGTTAGTGGACAGAGGCTGGGGAAATGGATGTGGACTTTTTGGCAAAGGGAGCCTGGTGACATGCGCTAAGTTTGCATGCTCCAAGAAAATGACCGGGAAGAGCATCCAGCCAGAGAATCTGGAGTACCGGATAATGCTGTCAGTTCATGGCTCCCAGCACAGTGGGATGATCGTTAATGACACAGGACATGAAACTGATGAGAATAGAGCGAAGGTTGAGATAACGCCCAATTCACCAAGAGCTGAAGCCACCCTGGGGGGTTTTGGAAGCCTAGGACTTGATTGTGAACCGAGGACAGGCCTTGACTTTTCAGATTTGTATTACTTGACTATGAATAACAAGCACTGGTTGGTTCACAAGGAGTGGTTCCACGACATTCCATTACCTTGGCATGCTGGGGCAGACACCGGAACTCCACATTGGAACAACAAAGAAGCATTGGTAGAGTTCAAGGACGCACATGCCAAAAGGCAAACTGTCGTGGTTCTAGGGAGTCAAGAAGGAGCAGTTCACACGGCCCTTGCTGGAGCTCTGGAGGCTGAGATGGATGGTGCAAAGGGAAGGCTGTCCTCTGGCCACTTGAAATGTCGCCTGAAAATGGATAAACTTAGATTGAAGGGCGTGTCATACTCCTTGTGTACCGCAGCGTTCACATTCACCAAGATCCCGGCTGAAACACTGCACGGGACAGTCACAGTGGAGGTACAGTACGCAGGGACAGATGGACCCTGCAAGGTTCCAGCTCAGATGGCGGTGGACATGCAAACTCTGACCCCAGTTGGGAGGCTGATAACCGCTAACCCTGTAATCACTGAAAGCACTGAGAACTCTAAGATGATGCTGGAACTTGATCCACCATTTGGGGACTCTTACATTGTCATAGGAGTCGGGGAGAAGAAGATCACCCATCACTGGCACAGGAGTGGCAGCACCATTGGAAAAGCATTTGAAGCCACTGTGAGAGGTGCCAAGAGAATGGCAGTCTTGGGAGACACAGCCTGGGATTTTGGATCAGTTGGAGGTGCTCTCAACTCATTGGGCAAGGGCATCCATCAAATTTTTGGAGCAGCTTTCAAATCATTGTTTGGAGGAATGTCCTGGTTCTCACAAATTCTCATTGGAACGTTGCTGGTGTGGTTGGGTCTGAATACAAAGAATGGATCTATTTCCCTTACGTGCTTGGCCTTAGGGGGAGTGTTGATCTTTTTATCCACAGCCGTCTCTGCTGATGTGGGGTGCTCGGTGGACTTCTCAAAGAAGGAAACGAGATGCGGTACGGGGGTGTTCGTCTATAACGACGTTGATGCCTGGAGGGACAGGTACAAGTACCATCCTGACTCCCCTCGTAGATTAGCAGCAGCAGTCAAGCAAGCCTGGGRAGATGGGATCTGTGGGATCTCCTCTGTTTCAAGAATGGAAAACATCATGTGGAGATCAGTAGAAGGGGAGCTCAACGCAATCCTGGAAGAGAATGGAGTTCAACTGACGGTCGTTGTGGGATCTGTAAAAAACCCCATGTGGAGAGGTCCACAGAGATTGCCCGTGCCTGTGAACGAGCTGCCCCACGGCTGGAAGGCTTGGGGGAAATCGTACTTCGTCAGAGCAGCAAAGACAAATAACAGCTTTGTCGTGGATGGTGACACACTGAAGGAATGCCCACTCAAACATAGAGCATGGAACAGCTTTCTTGTGGAGGATCATGGGTTCGGGGTATTTCACACTAGTGTCTGGCTCAAGGTTAGAGAAGATTATTCATTAGAGTGTGATCCAGCCGTTATTGGAACAGCTGCTAAGGGAAAGGAGGCTGTGCACAGTGATCTAGGCTACTGGATTGAGAGTGAGAAGAATGACACATGGAGGCTGAAGAGGGCCCACCTGATCGAGATGAAAACATGTGAATGGCCAAAGTCCCACACATTGTGGACAGATGGAATAGAAGAAAGTGATCTGATCATACCCAAGTCTTTAGCTGGGCCACTCAGCCATCACAACACCAGAGAGGGCTACAGGACTCAAATGAAAGGGCCATGGCACAGTGAAGAGCTTGAAATTCGGTTTGAGGAATGCCCAGGCACTAAGGTCCACGTGGAGGAAACATGTGGAACAAGAGGACCATCTCTGAGATCAACCACTGCAAGCGGAAGGGTGATCGAGGAATGGTGCTGCAGGGAATGCACAATGCCCCCACTGTCGTTCCGGGCTAAAGATGGCTGTTGGTATGGAATGGAGATAAGGCCCAGGAAAGAACCAGAAAGTAACTTAGTAAGGTCAATGGTGACTGCAGGATCAACTGATCACATGGATCACTTCTCCCTTGGAGTGCTTGTGATTCTGCTCATGGTGCAGGAAGGGCTGAAGAAGAGAATGACCACAAAGATCATCATAAGCACATCAATGGCAGTGCTGGTAGCTATGATCCTGGGAGGATTTTCAATGAGTGACCTGGCCAAGCTTGCAATTTTGATGGGTGCCACCTTTGCGGAAATGAACACTGGAGGAGATGTAGCTCATCTGGCGCTGATAGCGGCATTCAAAGTCAGACCTGCGTTGCTGGTATCTTTCATCTTCAGAGCTAATTGGACACCCCGTGAGAGCATGCTGCTGGCCCTGGCTTCGTGTCTTCTGCAAACTGCGATCTCCGCCTTGGAAGGCGACCTGATGGTTCTCATCAATGGTTTTGCTTTGGCCTGGTTGGCAATACGAGCGATGGTTGTTCCACGCACTGACAACATCACCTTGGCAATCCTGACTGCGCTGACACCACTGGCCCGGGGCACGCTGCTTGTGGCGTGGAGAGCAGGCCTTGCTACTTGCGGGGGGTTCATGCTTCTCTCTCTGAAGGGGAAGGGCAGTGTGAAGAAGAACCTACCATTTGTCATGGCCTTGGGACTCACCGCTGTGAGGCTGGTCGACCCCATCAACGTGGTGGGACTGCTGTTGCTCACAAGGAGTGGGAAGCGGAGCTGGCCCCCTAGTGAAGTACTCACAGCTGTTGGTCTGATATGCGCGTTGGCCGGAGGGTTCGCCAAGGCGGATATAGAGATGGCTGGGCCCATGGCCGCGGTCGGTCTGCTAATTGTCAGTTACGTGGTCTCAGGAAAGAGTGTGGACATGTACATTGAAAGAGCAGGTGACATCACATGGGAAAAAGATGCGGAAGTCACTGGAAACAGTCCCCGGCTCGATGTGGCACTGGATGAGAGTGGTGATTTCTCCCTAGTGGAGGATGATGGTCCCCCCATGAGAGAGATCATACTCAAAGTGGTCCTGATGACCATCTGTGGCATGAACCCAATAGCCATACCCTTTGCAGCTGGAGCGTGGTACGTGTATGTGAAGACTGGAAAAAGGAGTGGTGCTCTATGGGATGTGCCTGCTCCCAAGGAAGTAAAAAAGGGGGAGACCACAGATGGAGTGTACAGAGTAATGACTCGTAGACTGCTTGGTTCAACACAAGTTGGAGTGGGAGTCATGCAAGAGGGGGTCTTCCACACTATGTGGCACGTCACAAAAGGATCCGCGCTGAGAAGCGGTGAAGGGAGACTTGATCCATACTGGGGAGATGTCAAGCAGGATCTGGTGTCATATTGTGGTCCGTGGAAGCTAGACGCCGCCTGGGACGGGCACAGCGAGGTGCAGCTCTTGGCCGTGCCCCCCGGAGAGAGAGCGAGGAACATCCAGACTCCGCCCGGAATATTTAAGACAAAGGATGGGGACATTGGAGCAGTTGCGTTGGACTACCCAGCAGGAACTTCAGGATCTCCAATCCTAGACAAGTGTGGGAGAGTGATAGGACTCTATGGTAATGGGGTCGTGATAAAAAATGGGAGTTATGTTAGTGCCATCACCCAAGGGAGGAGGGAGGAAGAGACTCCTGTTGAGTGCTTCGAGCCTTCGATGTTGAAGAAGAAGCAGCTAACTGTCTTAGACCTGCATCCTGGAGCTGGGAAAACCAGGAGAGTTCTTCCTGAAATAGTCCGTGAAGCCATAAAAACAAGACTCCGTACTGTGATCTTAGCTCCAACCAGGGTTGTCGCTGCTGAAATGGAGGAAGCCCTTAGAGGGCTTCCAGTGCGTTATATGACAACAGCAGTCAATGTCACCCATTCTGGGACAGAAATTGTTGACTTAATGTGCCATGCCACCTTCACTTCACGTCTACTACAACCAATCAGAGTCCCCAACTATAATCTGTATATTATGGACGAGGCCCACTTCACAGATCCCTCAAGTATAGCAGCAAGAGGATACATTTCAACAAGGGTTGAGATGGGCGAGGCGGCCGCCATCTTCATGACCGCCACCCCACCAGGAACCCGTGACGCATTCCCGGACTCCAACTCACCAATTATGGACACCGAAGTGGAAGTCCCAGAGAGAGCCTGGAGCTCAGGCTTTGATTGGGTGACGGATCATTCTGGAAAAACAGTTTGGTTTGTTCCAAGCGTGAGGAACGGCAATGAGATCGCAGCTTGTCTGACAAAGGCTGGAAAACGGGTCATACAGCTCAGCAGAAAGACTTTTGAGACAGAGTTCCTGAAAACAAAAAATCAAGAGTGGGACTTCGTCGTGACAACTGACATTTCAGAGATGGGCGCCAACTTTAAAGCTGACCGTGTCATAGATTCCAGGAGATGCCTAAAGCCGGTCATACTTGATGGCGAGAGAGTCATTCTGGCTGGACCCATGCCTGTCACACATGCCAGCGCTGCCCAGAGGAGGGGGCGCATAGGCAGGAATCCCAACAAACCTGGAGATGAGTATCTGTATGGAGGTGGGTGCGCAGAGACTGATGAAGACCATGCACACTGGCTTGAAGCAAGAATGCTTCTTGACAACATTTACCTCCAAGATGGCCTCATAGCCTCGCTCTATCGACCTGAGGCCGACAAAGTAGCAGCTATTGAGGGAGAGTTCAAGCTTAGGACGGAGCAAAGGAAGACCTTTGTGGAACTCATGAAAAGAGGAGATCTTCCTGTTTGGCTGGCCTATCAGGTTGCATCTGCCGGAATAACCTACACAGATAGAAAATGGTGCTTTGATGGCACGACCAACAACACCATAATGGAAGACAGTGTGCCGGCAGAGGTGTGGACCAGATACGGAGAGAAAAGAGTGCTCAAACCAAGGTGGATGGACGCCAGAGTTTGTTCAGATCATGCGGCCCTGAAGTCATTCAAAGAGTTTGCCGCTGGGAAAAGAGGAGYGGCCTTTGGAGTGATGGAAGCCCTGGGAACATTGCCGGGACACATGACAGAGAGATTCCAGGAAGCCATTGACAACCTCGCTGTGCTCATGCGGGCAGAGACTGGAAGCAGGCCTTACGAAGCCGCGGCGGCCCAATTGCCGGAGACCTTAGAGACCATCATGCTTTTGGGGTTGCTGGGAACAGTCTCGCTGGGAATCTTTTTCGTCTTGATGCGGAATAAGGGCATCGGGAAGATGGGCTTTGGAATGGTGACTCTTGGGGCCAGCGCATGGCTTATGTGGCTCTCGGAAATTGAGCCAGCCAGAATTGCATGTGTCCTCATTGTTGTGTTCCTATTGCTGGTGGTGCTCATACCTGAGCCAGAAAAGCAAAGATCTCCCCAGGACAACCAAATGGCAATCATCATCATGATAGCAGTGGGTCTTCTGGGTTTGATTACCGCCAATGAACTTGGATGGTTGGAAAGAACAAAGAGTGACCTAAGCCATCTAATGGGAAGGAGAGAGGAGGGGGCAACCATAGGATTCTCAATGGACATTGACCTGCGGCCAGCCTCAGCTTGGGCTATCTATGCTGCTCTGACAACTTTCATCACCCCAGCCGTCCAACATGCGGTGACCACTTCATACAACAACTACTCCTTAATGGCGATGGCCACGCAAGCTGGAGTGTTGTTTGGTATGGGTAAAGGGATGCCATTCTACGCATGGG-ACTTTGGAGTCCCGCTGCTAATGATGGGTTGCTACTCACAATTAACACCCCTGACCCTAATAGTGGCCATCATTTTGCTCGTGGCGCACTACATGTACTTGATCCCAGGGCTGCAGGCAGCAGCTGCGCGTGCTGCCCAGAAGAGAACGGCAGCTGGCATCATGAAGAACCCTGTTGTGGATGGAATAGTGGTGACTGACATTGACACAATGACAATTGACCACCGAGTGGAGAAAAAGATGGGACAGGTGCTACTCATAGCAGTAGCCGTCTCCAGCGCCATACTGTCGCGGACCGCCTGGGGGTGGGGGGAGGCTGGGGCCCTGATCACAGCTGCAACTTCCACTTTGTGGGAAGGCTCTCCGAACAAGTACTGGAACTCCTCCACAGCCACTTCACTGTGTAACATTTTTAGGGGAAGTTACTTGGCTGGAGCTTCTCTAATCTACACAGTAACAAGAAACGCTGGCTTGGTCAAGAGACGTGGGGGTGGAACGGGAGAGACCCTGGGAGAGAAATGGAAGGCCCGCCTGAACCAGATGTCGGCCCTAGAGTTCTACTCCTACAAAAAGTCAGGCATCACCGAGGTGTGCAGAGAAGAGGCCCGCCGCGCCCTCAAGGACGGTGTGGCAACAGGAGGCCATGCTGTGTCCCGAGGAAGTGCAAAGCTTAGATGGTTGGTGGAGAGAGGATACCTGCAGCCCTATGGAAAGGTCATTGATCTTGGATGTGGCAGAGGGGGCTGGAGTTACTACGCCGCCACCATCCGCAAAGTTCAAGAAGTGAAAGGATACACAAAAGGAGGCCCTGGTCATGAAGAACCCATGTTGGTGCAAAGCTATGGGTGGAACATAGTCCGTCTTAAGAGTGGGGTGGACGTCTTTCATATGGCGGCTGAGCCGTGTGACACTTTGCTGTGTGATATAGGTGAGTCATCATCTAGTCCTGAAGTGGAAGAAGCACGGACGCTCAGAGTCCTTTCCATGGTGGGGGATTGGCTTGAAAAAAGACCAGGAGCCTTTTGTATAAAAGTGTTGTGCCCATACACCAGCACTATGATGGAAACCCTGGAGCGACTGCAGCGTAGGTATGGGGGAGGACTGGTCAGAGTGCCACTCTCCCGCAACTCTACACATGAGATGTACTGGGTCTCTGGAGCGAAAAGCAACACCATAAAAAGTGTGTCCACCACGAGCCAGCTCCTCTTGGGGCGCATGGACGGGCCCAGGAGGCCAGTGAAATATGAGGAGGATGTGAATCTCGGCTCCGGCACGCGGGCTGTGGTAAGCTGCGCTGAAGCTCCCAACATGAAGATCATTGGTAACCGCATTGAGAGGATCCGCAGTGAGCACGCGGAAACGTGGTTCTTTGACGAGAACCACCCATATAGGACATGGGCTTACCATGGAAGCTATGAGGCCCCTACACAAGGGTCAGCGTCCTCTCTAATAAACGGGGTTGTCAGGCTCCTGTCAAAACCCTGGGATGTGGTGACTGGAGTCACAGGAATAGCCATGACCGACACCACACCGTATGGTCAGCAAAGAGTTTTCAAGGAAAAAGTGGACACTAGGGTGCCAGACCCCCAAGAAGGCACTCGTCAGGTTATGAGCATGGTCTCTTCCTGGTTATGGAAAGAGCTAGGCAAACACAAACGGCCACGAGTCTGTACCAAAGAAGAGTTCATCAACAAGGTTCGTAGCAATGCAGCATTAGGGGCAATATTTGAAGAGGAAAAAGAGTGGAAGACTGCAGTGGAAGCTGTGAACGATCCAAGGTTCTGGGCTCTAGTGGACAAGGAAAGAGAGCACCACCTGAGAGGAGAGTGTCAGAGCTGTGTGTACAACATGATGGGAAAAAGAGAAAAGAAACAAGGGGAATTTGGAAAGGCCAAGGGCAGCCGCGCCATCTGGTATATGTGGCTAGGGGCTAGATTCCTGGAGTTCGAAGCCCTTGGATTCTTGAACGAGGATCACTGGATGGGGAGAGAGAATTCAGGAGGTGGTGTTGAAGGGCTGGGATTACAAAGACTCGGATATGTCCTAGAAGAGATGAGTCGCATACCAGGAGGAAGGATGTATGCTGATGACACAGCTGGCTGGGACACCCGCATCAGCAGGTTTGATCTGGAGAATGAAGCTCTAATCACCAACCAAATGGAGAAAGGGCACAGGGCCTTGGCATTGGCCATAATCAAGTACACATACCAAAACAAAGTGGTAAAGGTCCTCAGACCAGCTGAAAAAGGGAAGACAGTTATGGACATTATTTCAAGACAAGACCAAAGGGGGAGCGGACAAGTTGTCACTTACGCTCTTAATACATTCACCAACCTGGTGGTGCAGCTCATTCGGAATATGGAGGCTGAGGAAGTTCTAGAGATGCAAGACTTGTGGCTGCTGCGGAGGTCAGAGAAAGTGACCAACTGGTTGCAGAGCAACGGATGGGATAGGCTCAAACGAATGGCAGTCAGTGGAGATGATTGCGTTGTGAAACCAATTGATGATAGGTTTGCACATGCCCTCAGGTTCTTGAATGATATGGGAAAAGTCAGGAAGGACACACAAGAGTGGAAACCCTCAACTGGATGGGACAATTGGGAAGAAGTTCCGTTTTGCTCCCACCACTTCAACAAGCTCCATCTCAAGGACGGGAGGTCCATTGTGGTTCCCTGCCGCCACCAAGATGAACTGATTGGCCGAGCCCGCGTCTCACCAGGGGCGGGATGGAGCATCCGGGAGACTGCTTGCCTAGCAAAATCATATGCGCAAATGTGGCAGCTCCTTTATTTCCACAGAAGGGACCTCCGACTGATGGCCAATGCCATTTGTTCATCTGTGCCAGTTGACTGGGTTCCAACTGGGAGAACTACCTGGTCAATCCATGGAAAGGGAGAATGGATGACCACTGAAGACATGCTTGTGGTGTGGAACAGAGTGTGGATTGAGGAGAACGACCACATGGAAGACAAGACCCCAGTTACGAAATGGACAGACATTCCCTATTTGGGAAAAAGGGAAGACTTGTGGTGTGGATCTCTCATAGGGCACAGACCGCGCACTACCTGGGCTGAGAACATCAAAAACACAGTCAACATGATGCGCAGGATCATAGGTGATGAAGAAAAGTACATGGACTACCTATCCACCCAAGTTCGCTACTTGGGTGAAGAAGGGTCCACACCCGGAGTGTTGTAA

>ENA|AMA12084|AMA12084.1 Zika virus polyprotein

ATGAAAAACCC-AAAAAAGAAATCCGGAGGATTCCGGATTGTCAATATGCTAAAACGCGGAGTAGCCCGTGTGAGCCCCTTTGGGGGCTTGAAGAGGCTGCCAGCCGGACTTCTGCTGGGTCATGGGCCCATCAGGATGGTCTTGGCGATTCTAGCCTTTTTGAGATTCACGGCAATCAAGCCATCACTGGGTCTCATCAATAGATGGGGTTCAGTGGGGAAAAAAGAGGCTATGGAAATAATAAAGAAGTTCAAGAAAGATCTGGCTGCCATGCTGAGAATAATCAATGCTAGGAAGGAGAAGAAGAGACGAGGCGCAGATACTAGTGTCGGAATTGTTGGCCTCCTGCTGACCACAGCTATGGCAGCGGAGGTCACTAGACGTGGGAGTGCATACTATATGTACTTGGACAGAAACGATGCTGGGGAGGCCATATCTTTTCCAACCACATTGGGGATGAATAAGTGTTATATACAGATCATGGATCTTGGACACATGTGTGATGCCACCATGAGCTATGAATGCCCTATGCTGGATGAGGGGGTGGAACCAGATGACGTCGATTGTTGGTGCAACACGACGTCAACTTGGGTTGTGTACGGAACCTGCCATCACAAAAAAGGTGAAGCACGGAGATCTAGAAGAGCTGTGACGCTCCCCTCCCATTCCACTAGGAAGCTGCAAACGCGGTCGCAAACCTGGTTGGAATCAAGAGAATACACAAAGCACTTGATTAGAGTCGAAAATTGGATATTTAGGAACCCTGGCTTCGCGTTAGCAGCAGCTGCCATCGCTTGGCTTTTGGGAAGCTCAACGAGCCAAAAAGTCATATACTTGGTCATGATACTGCTGATTGCCCCGGCATACAGCATCAGGTGCATAGGAGTCAGCAATAGGGACTTTGTGGAAGGTATGTCAGGTGGGACCTGGGTTGATGTTGTCTTGGAACATGGAGGTTGTGTCACCGTAATGGCACAGGACAAACCGACTGTCGACATAGAGCTGGTTACAACAACAGTCAGCAACATGGCGGAGGTAAGATCCTACTGCTATGAGGCATCAATATCAGACATGGCTTCGGACAGCCGCTGCCCAACACAAGGTGAAGCCTACCTTGACAAGCAATCAGACACTCAATATGTCTGCAAAAGAACGTTAGTGGACAGAGGCTGGGGAAATGGATGTGGACTTTTTGGCAAAGGGAGCCTGGTGACATGCGCTAAGTTTGCATGCTCCAAGAAAATGACCGGGAAGAGCATCCAGCCAGAGAATCTGGAGTACCGGATAATGCTGTCAGTTCATGGCTCCCAGCACAGTGGGATGATTGTTAATGACACAGGACATGAAACTGATGAGAATAGAGCGAAAGTTGAGATAACGCCCAATTCACCAAGAGCCGAAGCCACCCTGGGGGGTTTTGGAAGCCTAGGACTTGATTGTGAACCGAGGACAGGCCTTGACTTTTCAGATTTGTATTACTTGACTATGAATAACAAGCACTGGTTGGTTCACAAGGAGTGGTTCCACGACATTCCATTACCTTGGCACGCTGGGGCAGACACCGGAACTCCACACTGGAACAACAAAGAAGCACTGGTAGAGTTCAAGGACGCACATGCCAAAAGGCAAACTGTCGTGGTTCTAGGGAGTCAAGAAGGAGCAGTTCACACGGCCCTTGCTGGAGCTCTGGAGGCTGAGATGGATGGTGCAAAGGGAAGGCTGTCCTCTGGCCACTTGAAATGTCGCCTGAAAATGGATAAACTTAGATTGAAGGGCGTGTCATACTCCTTGTGTACTGCAGCGTTCACATTCACCAAGATCCCGGCTGAAACACTGCACGGGACAGTCACAGTGGAGGTACAGTACGCAGGGACAGATGGACCTTGCAAGGTTCCAGCTCAGATGGCGGTGGACATGCAAACTCTGACCCCAGTTGGGAGGTTGATAACCGCTAACCCCGTAATCACTGAAAGCACTGAGAACTCTAAGATGATGCTGGAACTTGATCCACCATTTGGGGACTCTTACATTGTCATAGGAGTCGGGGAGAAGAAGATCACCCACCACTGGCACAGGAGTGGCAGCACCATTGGAAAAGCATTTGAAGCCACTGTGAGAGGTGCCAAGAGAATGGCAGTCTTGGGAGACACAGCCTGGGACTTTGGATCAGTTGGAGGCGCTCTCAACTCATTGGGCAAGGGCATCCATCAAATTTTTGGAGCAGCTTTCAAATCATTGTTTGGAGGAATGTCCTGGTTCTCACAAATTCTCATTGGAACGTTGCTGATGTGGTTGGGTCTGAACACAAAGAATGGATCTATTTCCCTTATGTGCTTGGCCTTAGGGGGAGTGTTGATCTTCTTATCCACAGCCGTCTCTGCTGATGTGGGGTGCTCGGTGGACTTCTCAAAGAAGGAGACGAGATGCGGTACAGGGGTGTTCGTCTATAACGACGTTGAAGCCTGGAGGGACAGGTACAAGTACCATCCTGACTCCCCCCGTAGATTGGCAGCAGCAGTCAAGCAAGCCTGGGAAGATGGTATCTGCGGGATCTCCTCTGTTTCAAGAATGGAAAACATCATGTGGAGATCAGTAGAAGGGGAGCTCAACGCAATCCTGGAAGAGAATGGAGTTCAACTGACGGTCGTTGTGGGATCTGTAAAAAACCCCATGTGGAGAGGTCCACAGAGATTGCCCGTGCCTGTGAACGAGCTGCCCCACGGCTGGAAGGCTTGGGGGAAATCGTACTTCGTCAGAGCAGCAAAGACAAATAACAGCTTTGTCGTGGATGGTGACACACTGAAGGAATGCCCACTCAAACATAGAGCATGGAACAGCTTTCTTGTGGAGGATCATGGGTTCGGGGTATTTCACACTAGTGTCTGGCTCAAGGTTAGAGAAGATTATTCATTAGAGTGTGATCCAGCCGTTATTGGAACAGCTGTTAAGGGAAAGGAGGCTGTACACAGTGATCTAGGCTACTGGATTGAGAGTGAGAAGAATGACACATGGAGGCTGAAGAGGGCCCATCTGATCGAGATGAAAACATGTGAATGGCCAAAGTCCCACACATTGTGGACAGATGGAATAGAAGAGAGTGATCTGATCATACCCAAGTCTTTAGCTGGGCCACTCAGCCATCACAATACCAGAGAGGGCTACAGGACCCAAATGAAAGGGCCATGGCACAGTGAAGAGCTTGAAATTCGGTTTGAGGAATGCCCAGGCACTAAGGTCCACGTGGAGGAAACATGTGGAACAAGAGGACCATCTCTGAGATCAACCACTGCAAGCGGAAGGGTGATCGAGGAATGGTGCTGCAGGGAGTGCACAATGCCCCCACTGTCGTTCCGGGCTAAAGATGGCTGTTGGTATGGAATGGAGATAAGGCCCAGGAAAGAACCAGAAAGCAACTTAGTAAGGTCAATGGTGACTGCAGGATCAACTGATCACATGGACCACTTCTCCCTTGGAGTGCTTGTGATTCTGCTCATGGTGCAGGAAGGGCTGAAGAAGAGAATGACCACAAAGATCATCATAAGCACATCAATGGCAGTGCTGGTAGCTATGATCCTGGGAGGATTTTCAATGAGTGACCTGGCTAAGCTTGCAATTTTGATGGGTGCCACCTTCGCGGAAATGAACACTGGAGGAGATGTAGCTCATCTGGCGCTGATAGCGGCATTCAAAGTCAGACCAGCGTTGCTGGTATCTTTCATCTTCAGAGCTAATTGGACACCCCGTGAAAGCATGCTGCTGGCCTTGGCCTCGTGTCTTTTGCAAACTGCGATCTCCGCCTTGGAAGGCGACCTGATGGTTCTCATCAATGGTTTTGCTTTGGCCTGGTTGGCAATACGAGCGATGGTTGTTCCACGCACTGATAACATCACCTTGGCAATCCTGGCTGCTCTGACACCACTGGCCCGGGGCACACTGCTTGTGGCGTGGAGAGCAGGCCTTGCTACTTGCGGGGGGTTTATGCTCCTCTCTCTGAAGGGAAAAGGCAGTGTGAAGAAGAACTTACCATTTGTCATGGCCCTGGGACTAACCGCTGTGAGGCTGGTCGACCCCATCAACGTGGTGGGACTGCTGTTGCTCACAAGGAGTGGGAAGCGGAGCTGGCCCCCTAGCGAAGTACTCACAGCTGTTGGCCTGATATGCGCATTGGCTGGAGGGTTCGCCAAGGCAGATATAGAGATGGCTGGGCCCATGGCCGCGGTCGGTCTGCTAATTGTCAGTTACGTGGTCTCAGGAAAGAGTGTGGACATGTACATTGAAAGAGCAGGTGACATCACATGGGAAAAAGATGCGGAAGTCACTGGAAACAGTCCCCGGCTCGATGTGGCGCTAGATGAGAGTGGTGATTTCTCCCTGGTGGAGGATGACGGTCCCCCCATGAGAGAGATCATACTCAAGGTGGTCCTGATGACCATCTGTGGCATGAACCCAATAGCCATACCCTTTGCAGCTGGAGCGTGGTACGTATACGTGAAGACTGGAAAAAGGAGTGGTGCTCTATGGGATGTGCCTGCTCCCAAGGAAGTAAAAAAGGGGGAGACCACAGATGGAGTGTACAGAGTAATGACTCGTAGACTGCTAGGTTCAACACAAGTTGGAGTGGGAGTTATGCAAGAGGGGGTCTTTCACACTATGTGGCACGTCACAAAAGGATCCGCGCTGAGAAGCGGTGAAGGGAGACTTGATCCATACTGGGGAGATGTCAAGCAGGATCTGGTGTCATACTGTGGTCCATGGAAGCTAGATGCCGCCTGGGACGGGCACAGCGAGGTGCAGCTCTTGGCCGTGCCCCCCGGAGAGAGAGCGAGGAACATCCAGACTCTGCCCGGAATATTTAAGACAAAGGATGGGGACATTGGAGCGGTTGCGCTGGATTACCCAGCAGGAACTTCAGGATCTCCAATCCTAGACAAGTGTGGGAGAGTGATAGGACTTTATGGCAATGGGGTCGTGATCAAAAATGGGAGTTATGTTAGTGCCATCACCCAAGGGAGGAGGGAGGAAGAGACTCCTGTTGAGTGCTTCGAGCCTTCGATGCTGAAGAAGAAGCAGCTAACTGTCTTAGACTTGCATCCTGGAGCTGGGAAAACCAGGAGAGTTCTTCCTGAAATAGTCCGTGAAGCCATAAAAACAAGACTCCGCACCGTGATCTTAGCTCCAACCAGGGTTGTCGCTGCTGAAATGGAGGAGGCCCTTAGAGGGCTTCCAGTGCGTTATATGACAACAGCAGTCAATGTCACCCACTCTGGAACAGAAATCGTCGACTTAATGTGCCATGCCACCTTCACTTCACGTCTACTACAGCCAATCAGAGTCCCCAACTATAATCTGTATATTATGGATGAGGCCCACTTCACAGATCCCTCAAGTATAGCAGCAAGAGGATACATTTCAACAAGGGTTGAGATGGGCGAGGCGGCTGCCATCTTCATGACCGCCACGCCACCAGGAACCCGTGACGCATTTCCGGACTCCAACTCACCAATTATGGACACCGAAGTGGAAGTCCCAGAGAGAGCCTGGAGCTCAGGCTTTGATTGGGTGACGGATCATTCTGGAAAAACAGTTTGGTTTGTTCCAAGCGTGAGGAACGGCAATGAGATCGCAGCTTGTCTGACAAAGGCTGGAAAACGGGTCATACAGCTCAGCAGAAAGACTTTTGAGACAGAGTTCCAGAAAACAAAACATCAAGAGTGGGACTTTGTCGTGACAACTGACATTTCAGAGATGGGCGCCAACTTTAAAGCTGACCGTGTCATAGATTCCAGGAGATGCCTAAAGCCGGTCATACTTGATGGCGAGAGAGTCATTCTGGCTGGACCCATGCCTGTCACACATGCCAGCGCTGCCCAGAGGAGGGGGCGCATAGGCAGGAATCCCAACAAACCTGGAGATGAGTATCTGTATGGAGGTGGGTGCGCAGAGACTGACGAAGACCATGCACACTGGCTTGAAGCAAGAATGCTCCTTGACAATATTTACCTCCAAGATGGCCTCATAGCCTCGCTCTATCGACCTGAGGCCGACAAAGTAGCAGCCATTGAGGGAGAGTTCAAGCTTAGGACGGAGCAAAGGAAGACCTTTGTGGAACTCATGAAAAGAGGAGATCTTCCTGTTTGGCTGGCCTATCAGGTTGCATCTGCCGGAATAACCTACACAGATAGAAGATGGTGCTTTGATGGCACGACCAACAACACCATAATGGAAGACAGTGTGCCGGCAGAGGTGTGGACCAGACACGGAGAGAAAAGAGTGCTCAAACCGAGGTGGATGGACGCCAGAGTTTGTTCAGATCATGCGGCCCTGAAGTCATTCAAGGAGTTTGCCGCTGGGAAAAGAGGAGCGGCTTTTGGAGTGATGGAAGCCCTGGGAACACTGCCAGGACACATGACAGAGAGATTCCAGGAAGCCATTGACAACCTCGCTGTGCTCATGCGGGCAGAGACTGGAAGCAGGCCTTACAAAGCCGCGGCGGCCCAATTGCCGGAGACCCTAGAGACCATTATGCTTTTGGGGTTGCTGGGAACAGTCTCGCTGGGAATCTTCTTCGTCTTGATGAGGAACAAGGGCATAGGGAAGATGGGCTTTGGAATGGTGACTCTTGGGGCCAGCGCATGGCTCATGTGGCTCTCGGAAATTGAGCCAGCCAGAATTGCATGTGTCCTCATTGTTGTGTTTCTATTGCTGGTGGTGCTCATACCTGAGCCAGAAAAGCAAAGATCTCCCCAGGACAACCAAATGGCAATCATCATCATGGTAGCAGTAGGTCTTCTGGGCTTGATTACCGCCAATGAACTCGGATGGTTGGAGAGAACAAAGAGTGACCTAAGCCATCTAATGGGAAGGAGAGAGGAGGGGGCAACCATAGGATTCTCAATGGACATTGACCTGCGGCCAGCCTCAGCTTGGGCCATCTATGCTGCCTTGACAACTTTCATTACCCCAGCCGTCCAACATGCAGTGACCACTTCATACAACAACTACTCCTTAATGGCGATGGCCACGCAAGCTGGAGTGTTGTTTGGTATGGGCAAAGGGATGCCATTCTACGCATGGG-ACTTTGGAGTCCCGCTGCTAATGATAGGTTGCTACTCACAATTAACACCCCTGACCCTAATAGTGGCCATCATTTTGCTCGTGGCGCACTACATGTACTTGATCCCAGGGCTGCAGGCAGCAGCTGCGCGTGCTGCCCAGAAGAGAACGGCAGCTGGCATCATGAAGAACCCTGTTGTGGATGGAATAGTGGTGACTGACATTGACACAATGACAATTGACCCCCAAGTGGAGAAAAAGATGGGACAGGTGCTACTCATAGCAGTAGCCGTCTCCAGCGCCATACTGTCGCGGACCGCCTGGGGGTGGGGGGAGGCTGGGGCCCTGATCACAGCCGCAACTTCCACTTTGTGGGAAGGCTCTCCGAACAAGTACTGGAACTCCTCTACAGCCACTTCACTGTGTAACATTTTTAGGGGAAGTTACTTGGCTGGAGCTTCTCTAATCTACACAGTAACAAGAAACGCTGGCTTGGTCAAGAGACGTGGGGGTGGAACAGGAGAGACCCTGGGAGAGAAATGGAAGGCCCGCTTGAACCAGATGTCGGCCCTGGAGTTCTACTCCTACAAAAAGTCAGGCATCACCGAGGTGTGCAGAGAAGAGGCCCGCCGCGCCCTCAAGGACGGTGTGGCAACGGGAGGCCATGCTGTGTCCCGAGGAAGTGCAAAGCTGAGATGGTTGGTGGAGCGGGGATACCTGCAGCCCTATGGAAAGGTCATTGATCTTGGATGTGGCAGAGGGGGCTGGAGTTACTACGCCGCCACCATCCGCAAAGTTCAAGAAGTGAAAGGATACACAAAAGGAGGCCCTGGTCATGAAGAACCCGTGTTGGTGCAAAGCTATGGGTGGAACATAGTCCGTCTTAAGAGTGGGGTGGACGTCTTTCATATGGCGGCTGAGCCGTGTGACACGTTGCTGTGTGACATAGGTGAGTCATCATCTAGTCCTGAAGTGGAAGAAGCACGGACGCTCAGAGTCCTCTCCATGGTGGGGGATTGGCTTGAAAAAAGACCAGGAGCCTTTTGTATAAAGGTGTTGTGCCCATACACCAGCACTATGATGGAAACCCTGGAGCGACTGCAGCGTAGGTATGGGGGAGGACTGGTCAGAGTGCCACTCTCCCGCAACTCTACACATGAGATGTATTGGGTCTCTGGAGCGAAAAGCAACACCATAAAAAGTGTGTCCACCACGAGCCAGCTCCTCTTGGGGCGCATGGACGGGCCTAGGAGGCCAGTGAAATATGAGGAGGATGTGGATCTCGGCTCTGGCACGCGGGCTGTGGTAAGCTGCGCTGAAGCTCCCAACATGAAGATCATTGGTAACCGCATTGAAAGGATCCGCAGTGAGCACGCGGAAACGTGGTTCTTTGACGAGAACCACCCATATAGGACATGGGCTTACCATGGAAGCTATGAGGCCCCCACACAAGGGTCAGCGTCCTCTCTAATAAACGGGGTTGTCAGGCTCCTGTCAAAACCCTGGGATGTGGTGACTGGAGTCACAGGAATAGCCATGACCGACACCACACCGTATGGTCAGCAAAGAGTTTTCAAGGAAAAAGTGGACACTAGGGTGCCAGACCCCCAAGAAGGCACTCGTCAGGTTATGAGCATGGTCTCTTCCTGGTTGTGGAAAGAGCTAGGCAAACACAAACGGCCACGAGTCTGTACCAAAGAAGAGTTCATCAACAAGGTTCGTAGCAATGCAGCATTAGGGGCAATATTTGAAGAGGAAAAAGAGTGGAAGACTGCAGTGGAAGCTGTGAACGATCCAAGGTTCTGGGCTCTAGTGGATAAGGAAAGAGAGCACCACCTGAGAGGAGAGTGCCAGAGTTGTGTGTACAACATGATGGGAAAAAGAGAAAAGAAACAAGGGGAATTTGGAAAGGCCAAGGGCAGCCGCGCCATCTGGTATATGTGGCTAGGGGCTAGATTTCTAGAGTTCGAAGCCCTTGGATTCTTGAACGAGGATCACTGGATGGGGAGAGAGAACTCAGGAGGTGGTGTTGAAGGGCTGGGATTACAAAGACTCGGATATGTCCTAGAAGAGATGAGTCGTATACCAGGAGGAAGGATGTATGCAGATGACACTGCTGGCTGGGACACCCGCATCAGCAGGTTTGATCTGGAGAATGAAGCTCTAATCACCAACCAAATGGAAAAAGGGCACAGGGCCTTGGCATTGGCCATAATCAAGTACACATACCAAAACAAAGTGGTAAAGGTCCTTAGACCAGCTGAAAAAGGGAAAACAGTTATGGACATTATTTCGAGACAAGACCAAAGGGGGAGCGGACAAGTTGTCACTTACGCTCTTAACACATTTACCAACCTAGTGGTGCAACTCATTCGGAATATGGAGGCTGAGGAAGTTCTAGAGATGCAAGACTTGTGGCTGCTGCGGAGGTCAGAGAAAGTGACCAACTGGTTGCAGAGCAACGGATGGGATAGGCTCAAACGAATGGCAGTCAGTGGAGATGATTGCGTTGTGAAGCCAATTGATGATAGGTTTGCACATGCCCTCAGGTTCTTGAATGATATGGGAAAAGTTAGGAAGGACACACAAGAGTGGAAACCCTCAACTGGATGGGACAACTGGGAAGAAGTTCCGTTTTGCTCCCACCACTTCAACAAGCTCCATCTCAAGGACGGGAGGTCCATTGTGGTTCCCTGCCGCCACCAAGATGAACTGATTGGCCGGGCCCGCGTCTCTCCAGGGGCGGGATGGAGCATCCGGGAGACTGCTTGCCTAGCAAAATCATATGCGCAAATGTGGCAGCTCCTTTATTTCCACAGAAGGGACCTCCGACTGATGGCCAATGCCATTTGTTCATCTGTGCCAGTTGACTGGGTTCCAACTGGGAGAACTACCTGGTCAATCCATGGAAAGGGAGAATGGATGACCACTGAAGACATGCTTGTGGTGTGGAACAGAGTGTGGATTGAGGAGAACGACCACATGGAAGACAAGACCCCAGTTACGAAATGGACAGACATCCCCTATTTGGGAAAAAGGGAAGACTTGTGGTGTGGATCTCTCATAGGGCACAGACCGCGCACCACCTGGGCTGAGAACATTAAAAACACAGTCAACATGGTGCGCAGGATCATAGGTGATGAAGAAAAGTACATGGACTACCTATCCACCCAAGTTCGCTACTTGGGTGAAGAAGGGTCTACACCTGGAGTGCTGTAA

>ENA|AMC13912|AMC13912.1 Zika virus polyprotein

ATGAAAAACCC-AAAAAAGAAATCCGGAGGATTCCGGATTGTCAATATGCTAAAACGCGGAGTAGCCCGTGTGAGCCCCTTTGGGGGCTTGAAGAGGCTGCCAGCCGGACTTCTGCTGGGTCATGGGCCCATCAGGATGGTCTTGGCGATTCTAGCCTTTTTGAGATTCACGGCAATCAAGCCATCACTGGGTCTCATCAATAGATGGGGTTCAGTGGGGAAAAAAGAGGCTATGGAAATAATAAAGAAGTTCAAGAAAGATCTGGCTGCCATGCTGAGAATAATCAATGCTAGGAAGGAGAAGAAGAGACGAGGCGCAGATACTAGTGTCGGAATTGTTGGCCTCCTGCTGACCACAGCTATGGCAGCGGAGGTCACTAGACGTGGGAGTGCATACTATATGTACTTGGACAGAAACGATGCTGGGGAGGCCATATCTTTTCCAACCACATTGGGGATGAATAAGTGTTATATACAGATCATGGATCTTGGACACATGTGTGATGCCACCATGAGCTATGAATGCCCTATGCTGGATGAGGGGGTGGAACCAGATGACGTCGATTGTTGGTGCAACACGACGTCAACTTGGGTTGTGTACGGAACCTGCCATCACAAAAAAGGTGAAGCACGGAGATCTAGAAGAGCTGTGACGCTCCCCTCCCATTCCACTAGGAAGCTGCAAACGCGGTCGCAAACCTGGTTGGAATCAAGAGAATACACAAAGCACTTGATTAGAGTCGAAAATTGGATATTCAGGAACCCTGGCTTCGCGTTAGCAGCAGCTGCCATCGCTTGGCTTTTGGGAAGCTCAACGAGCCAAAAAGTCATATACTTGGTCATGATACTGCTGATTGCCCCGGCATACAGCATCAGGTGCATAGGAGTCAGCAATAGGGACTTTGTGGAAGGTATGTCAGGTGGGACTTGGGTTGATGTTGTCTTGGAACATGGAGGTTGTGTCACCGTAATGGCACAGGACAAACCGACTGTCGACATAGAGCTGGTTACAACAACAGTCAGCAACATGGCGGAGATAAGATCCTACTGCTATGAGGCATCAATATCAGACATGGCTTCGGACAGCCGCTGCCCAACACAAGGTGAAGCCTACCTTGACAAGCAATCAGACACTCAATATGTCTGCAAAAGAACGTTAGTGGACAGAGGCTGGGGAAATGGATGTGGACTTTTTGGCAAAGGGAGCCTGGTGACATGCGCTAAGTTTGCATGCTCCAAGAAAATGACCGGGAAGAGCATCCAGCCAGAGAATCTGGAGTACCGGATAATGCTGTCAGTTCATGGCTCCCAGCACAGTGGGATGATCGTTAATGACACAGGACATGAAACTGATGAGAATAGAGCGAAGGTTGAGATAACGCCCAATTCACCAAGAGCCGAAGCCACCCTGGGGGGTTTTGGAAGCCTAGGACTTGATTGTGAACCGAGGACAGGCCTTGACTTTTCAGATTTGTATTACTTGACTATGAATAACAAGCACTGGTTGGTTCACAAGGAGTGGTTCCACGACATTCCATTACCTTGGCACGCTGGGGCAGACACCGGAACTCCACACTGGAACAACAAAGAAGCACTGGTGGAGTTCAAGGACGCACATGCCAAAAGGCAAACTGTCGTGGTTCTAGGGAGTCAAGAAGGAGCAGTTCACACGGCCCTTGCTGGAGCTCTGGAGGCTGAGATGGATGGTGCAAAGGGAAGGCTGTCCTCTGGCCACTTGAAATGTCGCCTGAAAATGGATAAACTTAGATTGAAGGGCGTGTCATACTCCTTGTGTACCGCAGCGTTCACATTCACCAAGATCCCGGCTGAAACACTGCACGGGACAGTCACAGTGGAGGTACAGTACGCAGGGACAGATGGACCTTGCAAGGTTCCAGCTCAGATGGCGGTGGACATGCAAACTCTGACCCCAGTTGGGAGGTTGATAACCGCTAACCCCGTAATCACTGAAAGCACTGAGAACTCTAAGATGATGCTGGAACTTGATCCACCATTTGGGGACTCTTACATTGTCATAGGAGTCGGGGAGAAGAAGATCACCCACCACTGGCACAGGAGTGGCAGCACCATTGGAAAAGCATTTGAAGCCACTGTGAGAGGTGCCAAGAGAATGGCAGTCTTGGGAGACACAGCCTGGGACTTTGGATCAGTTGGAGGCGCTCTCAACTCATTGGGCAAGGGCATCCATCAAATTTTTGGAGCTGCTTTCAAATCATTGTTTGGAGGAATGTCCTGGTTCTCACAAATTCTCATTGGAACGTTGCTGATGTGGTTGGGTCTGAACACAAAGAATGGATCTATTTCCCTTATGTGCTTGGCCTTAGGGGGAGTGTTGATCTTCTTATCCACAGCCGTCTCTGCTGATGTGGGGTGCTCGGTGGACTTCTCAAAGAAGGAGACGAGATGCGGTACAGGGGTGTTCGTCTATAACGACGTTGAAGCCTGGAGGGACAGGTACAAGTACCATCCTGACTCCCCCCGTAGATTGGCAGCAGCAGTCAAGCAAGCCTGGGAAGATGGTATCTGCGGGATCTCCTCTGTTTCAAGAATGGAAAACATCATGTGGAGATCAGTAGAAGGGGAGCTCAACGCAATCCTGGAAGAGAATGGAGTTCAACTGACGGTCGTTGTGGGATCTGTAAAAAACCCCATGTGGAGAGCTCCACAGAGATTGCCCGTGCCTGTGAACGAGCTGCCCCACGGCTGGAAGGCTTGGGGGAAATCGTACTTCGTCAGAGCAGCAAAGACAAATAACAGCTTTGTCGTGGATGGTGACACACTGAAGGAATGCCCACTCAAACATAGAGCATGGAACAGCTTTCTTGTGGAGGATCATGGGTTCGGGGTATTTCACACTAGTGTCTGGCTCAAGGTTAGAGAAGATTATTCATTAGAGTGTGATCCAGCCGTTATTGGAACAGCTGTTAAGGGAAAGGAGGCTGTACACAGTGATCTAGGCTACTGGATTGAGAGTGAGAAGAATGACACATGGAGGCTGAAGAGGGCCCATCTGATCGAGATGAAAACATGTGAATGGCCAAAGTCCCACACATTGTGGACAGATGGAATAGAAGAGAGTGATCTGATCATACCCAAGTCTTTAGCTGGGCCACTCAGCCATCACAATACCAGAGAGGGCTACAGGACCCAAATGAAAGGGCCATGGCACAGTGAAGAGCTTGAAATTCGGTTTGAGGAATGCCCAGGCACTAAGGTCCACGTGGAGGAAACATGTGGAACAAGAGGACCATCTCTGAGATCAACCACTGCAAGCGGAAGGGTGATCGAGGAATGGTGCTGCAGGGAGTGCACAATGCCCCCACTGTCGTTCCGGGCTAAAGATGGCTGTTGGTATGGAATGGAGATAAGGCCCAGGAAAGAACCAGAAAGCAACTTAGTAAGGTCAATGGTGACTGCAGGATCAACTGATCACATGGATCACTTCTCCCTTGGAGTGCTTGTGATTCTGCTCATGGTGCAGGAAGGGCTAAAGAAGAGAATGACCACAAAGATCATCATAAGCACATCAATGGCAGTGCTGGTAGCTATGATCCTGGGAGGATTTTCAATGAGTGACCTGGCTAAGCTTGCAATTTTGATGGGTGCCACCTTCGCGGAAATGAACACTGGAGGAGATGTAGCTCATCTGGCGCTGATAGCGGCATTCAAAGTCAGACCAGCGTTGCTGGTATCTTTCATCTTCAGAGCTAATTGGACACCCCGTGAAAGCATGCTACTGGCCTTGGCCTCGTGTCTTTTGCAAACTGCGATCTCCGCCTTGGAAGGCGACCTGATGGTTCTCATCAATGGTTTTGCTTTGGCCTGGTTGGCAATACGAGCGATGGTTGTTCCACGCACTGATAACATCACCTTGGCAATCCTGGCTGCTCTGACACCACTGGCCCGGGGCACACTGCTTGTGGCGTGGAGAGCAGGCCTTGCTACTTGCGGGGGGTTTATGCTCCTCTCTCTGAAGGGAAAAGGCAGTGTGAAGAAGAACTTACCATTTGTCATGGCCCTGGGACTAACCGCTGTGAGGCTGGTCGACCCCATCAACGTGGTGGGACTGCTGTTGCTCACAAGGAGTGGGAAGCGGAGCTGGCCCCCTAGCGAAGTACTCACAGCTGTTGGCCTGATATGCGCATTGGCTGGAGGGTTCGCCAAGGCAGATATAGAGATGGCTGGGCCCATGGCCGCGGTCGGTCTGCTAATTGTCAGTTACGTGGTCTCAGGAAAGAGTGTGGACATGTACATTGAAAGAGCAGGTGACATCACATGGGAAAAAGATGCGGAAGTCACTGGAAACAGTCCCCGGCTCGATGTGGCGCTAGATGAGAGTGGTGATTTCTCCCTGGTGGAGGATGACGGTCCCCCCATGAGAGAGATCATACTCAAGGTGGTCCTGATGACCATCTGTGGCATGAACCCAATAGCCATACCCTTTGCAGCTGGAGCGTGGTACGTATACGTGAAGACTGGAAAAAGGAGTGGTGCTCTATGGGATGTGCCTGCTCCCAAGGAAGTAAAAAAGGGGGAGACCACAGATGGAGTGTACAGAGTAATGACTCGTAGACTGCTAGGTTCAACACAAGTTGGAGTGGGAGTTATGCAAGAGGGGGTCTTTCACACTATGTGGCACGTCACAAAAGGATCCGCACTGAGAAGCGGTGAAGGGAGACTTGATCCATACTGGGGAGATGTCAAGCAGGATCTGGTGTCATACTGTGGTCCATGGAAGCTAGATGCCGCCTGGGACGGGCACAGCGAGGTGCAGCTCTTGGCCGTGCCCCCCGGAGAGAGAGCGAGGAACATCCAGACTCTGCCCGGAATATTTAAGACAAAGGATGGGGACATTGGAGCGGTTGCGCTGGATTACCCAGCAGGAACTTCAGGATCTCCAATCCTAGACAAGTGTGGGAGAGTGATAGGACTTTATGGCAATGGGGTCGTGATCAAAAATGGGAGTTATGTTAGTGCCATCACCCAAGGGAGGAGGGAGGAAGAGACTCCTGTTGAGTGCTTCGAGCCTTCGATGCTGAAGAAGAAGCAGCTAACTGTCTTAGACTTACATCCTGGAGCTGGGAAAACCAGGAGAGTTCTTCCTGAAATAGTCCGTGAAGCCATAAAAACAAGACTCCGTACTGTGATCTTAGCTCCAACCAGGGTTGTCGCTGCTGAAATGGAGGAGGCCCTTAGAGGGCTTCCAGTGCGTTATATGACAACAGCAGTCAATGTCACCCACTCTGGAACAGAAATCGTCGACTTAATGTGCCATGCCACCTTCACTTCACGTCTACTACAGCCAATCAGAGTCCCCAACTATAATCTGTATATTATGGATGAGGCCCACTTCACAGATCCCTCAAGTATAGCAGCAAGAGGATACATTTCAACAAGGGTTGAGATGGGCGAGGCGGCTGCCATCTTCATGACCGCCACGCCACCAGGAACCCGTGACGCATTTCCGGACTCCAACTCACCAATTATGGACACCGAAGTGGAAGTCCCAGAGAGAGCCTGGAGCTCAGGCTTTGATTGGGTGACGGATCATTCTGGAAAAACAGTTTGGTTTGTTCCAAGCGTGAGGAACGGCAATGAGATCGCAGCTTGTCTGACAAAGGCTGGAAAACGGGTCATACAGCTCAGCAGAAAGACTTTTGAGACAGAGTTCCAGAAAACAAAACATCAAGAGTGGGACTTTGTCGTGACAACTGACATTTCAGAGATGGGCGCCAACTTTAAAGCTGACCGTGTCATAGATTCCAGGAGATGCCTAAAGCCGGTCATACTTGATGGCGAGAGAGTCATTCTGGCTGGACCCATGCCTGTCACACATGCCAGCGCTGCCCAGAGGAGGGGGCGCATAGGCAGGAATCCCAACAAACCTGGAGATGAGTATCTGTATGGAGGTGGGTGCGCAGAGACTGACGAAGACCATGCACACTGGCTTGAAGCAAGAATGCTCCTTGACAATATTTACCTCCAAGATGGCCTCATAGCCTCGCTCTATCGACCTGAGGCCGACAAAGTAGCAGCCATTGAGGGAGAGTTCAAGCTTAGGACGGAGCAAAGGAAGACCTTTGTGGAACTCATGAAAAGAGGAGATCTTCCTGTTTGGCTGGCCTATCAGGTTGCATCTGCCGGAATAACCTACACAGATAGAAGATGGTGCTTTGATGGCACGACCAACAACACCATACTGGAAGACAGTGTGCCGGCAGAGGTGTGGACCAGACACGGAGAGAAAAGAGTGCTCAAACCGAGGTGGATGGACGCCAGAGTTTGTTCAGATCATGCGGCCCTGAAGTCATTCAAGGAGTTTGCCGCTGGGAAAAGAGGAGCGGCTTTTGGAGTGATGGAAGCCTTGGGAACACTGCCAGGACACATGACAGAGAGATTCCAGGAAGCCATTGACAACCTCGCTGTGCTCATGCGGGCAGAGACTGGAAGCAGGCCTTACAAAGCCGCGGCGGCCCAATTGCCGGAGACCCTAGAGACCATTATGCTTTTGGGGTTGCTGGGAACAGTCTCGCTGGGAATCTTTTTCGTCTTGATGAGGAACAAGGGCATAGGGAAGATGGGCTTTGGAATGGTGACCCTTGGGGCCAGTGCATGGCTCATGTGGCTCTCGGAAATTGAGCCAGCCAGAATTGCATGTGTCCTCATTGTTGTGTTCCTATTGCTGGTGGTGCTCATACCTGAGCCAGAAAAGCAAAGATCTCCCCAGGACAACCAAATGGCAATCATCATCATGGTAGCAGTAGGTCTTCTGGGCTTGATTACCGCCAATGAACTCGGATGGTTGGAGAGAACAAAGAGTGACCTAAGCCATCTAATGGGAAGGAGAGAGGAGGGAGCAACCATAGGATTCTCAATGGACATTGACCTGCGGCCAGCCTCAGCTTGGGCCATCTATGCTGCCTTGACAACTTTCATTACCCCAGCCGTCCAACATGCAGTGACCACTTCATACAACAACTACTCCTTAATGGCGATGGCCACGCAAGCTGGAGTGTTGTTTGGTATGGGCAAAGGGATGCCATTCTACGCATGGG-ACTTTGGAGTCCCGCTGCTAATGATAGGTTGCTACTCACAATTAACACCCCTGACCCTAATAGTGGCCATCATTTTGCTCGTGGCGCACTACATGTACTTGATCCCAGGGCTGCAGGCAGCAGCTGCGCGTGCTGCCCAGAAGAGAACGGCAGCTGGCATCATGAAGAACCCTGTTGTGGATGGAATAGTGGTGACTGACATTGACACAATGACAATTGACCCCCAAGTGGAGAAAAAGATGGGACAGGTGCTACTCATAGCAGTAGCCGTCTCCAGCGCCATACTGTCGCGGACCGCCTGGGGGTGGGGGGAGGCTGGGGCCCTGATCACAGCCGCAACTTCCACTTTGTGGGAAGGCTCTCCGAACAAGTACTGGAACTCCTCTACAGCCACTTCACTGTGTAACATTTTTAGGGGAAGTTATTTGGCTGGAGCTTCTCTAATCTACACAGTAACAAGAAACGCTGGCTTGGTCAAGAGACGTGGGGGTGGAACAGGAGAGACCCTGGGAGAGAAATGGAAGGCCCGCTTGAACCAGATGTCGGCCCTGGAGTTCTACTCCTACAAAAAGTCAGGCATCACTGAGGTGTGCAGAGAAGAGGCCCGCCGCGCCCTCAAGGACGGTGTGGCAACGGGAGGCCATGCTGTGTCCCGAGGAAGTGCAAAGCTGAGATGGTTGGTGGAGCGGGGATACCTGCAGCCCTATGGAAAGGTCATTGATCTTGGATGTGGCAGAGGGGGCTGGAGTTACTACGCCGCCACCATCCGCAAAGTTCAAGAAGTGAAAGGATACACAAAAGGAGGCCCTGGTCATGAAGAACCCGTGTTGGTGCAAAGCTATGGGTGGAACATAGTCCGTCTTAAGAGTGGGGTGGACGTCTTTCATATGGCGGCTGAGCCGTGTGACACGTTGCTGTGTGACATAGGCGAGTCATCATCTAGTCCTGAAGTGGAAGAAGCACGGACGCTCAGAGTCCTCTCCATGGTGGGGGATTGGCTTGAAAGAAGACCAGGAGCCTTTTGTATAAAAGTGTTGTGCCCATACACCAGCACTATGATGGAAACCCTGGAGCGACTGCAGCGTAGGTATGGGGGAGGACTGGTCAGAGTGCCACTCTCCCGCAACTCTACACATGAGATGTACTGGGTCTCTGGAGCGAAAAGCAACACCATAAAAAGTGTGTCCACCACGAGCCAGCTCCTCTTGGGGCGCATGGACGGGCCTAGGAGGCCAGTGAAATATGAGGAGGATGTGAATCTCGGCTCTGGCACGCGGGCTGTGGTAAGCTGCGCTGAAGCTCCCAACATGAAGATCATTGGTAACCGCATTGAAAGGATCCGCAGTGAGCACGCGGAAACGTGGTTCTTTGACGAGAACCACCCATATAGGACATGGGCTTACCATGGAAGCTATGAGGCCCCCACACAAGGGTCAGCGTCCTCTCTAATAAACGGGGTTGTCAGGCTCCTGTCAAAACCCTGGGATGTGGTGACTGGAGTCACAGGAATAGCCATGACCGACACCACACCGTATGGTCAGCAAAGAGTTTTCAAGGAAAAAGTGGACACTAGGGTGCCAGACCCCCAAGAAGGCACTCGTCAGGTTATGAGCATGGTCTCTTCCTGGTTGTGGAAAGAGCTAGGCAAACACAAACGGCCACGAGTCTGTACCAAAGAAGAGTTCATCAACAAGGTTCGTAGCAATGCAGCATTAGGGGCAATATTTGAAGAGGAAAAAGAGTGGAAGACTGCAGTGGAAGCTGTGAACGATCCAAGGTTCTGGGCTCTAGTGGACAAGGAAAGAGAGCACCACCTGAGAGGAGAGTGCCAGAGTTGTGTGTACAACATGATGGGAAAAAGAGAAAAGAAACAAGGGGAATTTGGAAAGGCCAAGGGCAGCCGCGCCATCTGGTATATGTGGCTAGGGGCTAGATTTCTAGAGTTCGAAGCCCTTGGATTCTTGAACGAGGATCACTGGATGGGGAGAGAGAACTCAGGAGGTGGTGTTGAAGGGCTGGGATTACAAAGACTCGGATATGTCCTAGAAGAGATGAGTTGCATACCAGGAGGAAGGATGTATGCAGATGACACTGCTGGCTGGGACACCCGCATCAGCAGGTTTGATCTGGAGAATGAAGCTCTAATCACCAACCAAATGGAGAAAGGGCACAGGGCCTTGGCATTGGCCATAATCAAGTACACATACCAAAACAAAGTGGTAAAGGTCCTTAGACCAGCTGAAAAAGGGAAAACAGTTATGGACATTATTTCAAGACAAGACCAAAGGGGGAGCGGACAAGTTGTCACTTACGCTCTTAACACATTTACCAACCTAGTGGTGCAACTCATTCGGAATATGGAGGCTGAGGAAGTTCTAGAGATGCAAGACTTGTGGCTGCTGCGGAGGTCAGAGAAAGTGACCAACTGGTTGCAGAGCAACGGATGGGATAGGCTCAAACGAATGGCAGTCAGTGGAGATGATTGCGTTGTGAAGCCAATTGATGATAGGTTTGCACATGCCCTCAGGTTCTTGAATGATATGGGAAAAGTTAGGAAGGACACACAAGAGTGGAAACCCTCAACTGGATGGGACAACTGGGAAGAAGTTCCGTTTTGCTCCCACCACTTCAACAAGCTCCATCTCAAGGACGGGAGGTCCATTGTGGTTCCCTGCCGCCACCAAGATGAACTGATTGGCCGGGCCCGCGTCTCTCCAGGGGCGGGATGGAGCATCCGGGAGACTGCTTGCCTAGCAAAATCATATGCGCAAATGTGGCAGCTCCTTTATTTCCACAGAAGGGACCTCCGACTGATGGCCAATGCCATTTGTTCATCTGTGCCAGTTGACTGGGTTCCAACTGGGAGAACTACCTGGTCAATCCATGGAAAGGGAGAATGGATGACCACTGAAGACATGCTTGTGGTGTGGAACAGAGTGTGGATTGAGGAGAACGACCACATGGAAGACAAGACCCCAGTTACGAAATGGACAGACATTCCCTATTTGGGAAAAAGGGAAGACTTGTGGTGTGGATCTCTCATAGGGCACAGACCGCGCACCACCTGGGCTGAGAACATTAAAAACACAGTCAACATGGTGCGCAGGATCATAGGTGATGAAGAAAAGTACATGGACTACCTATCCACCCAAGTTCGCTACTTGGGTGAAGAAGGGTCTACACCTGGAGTGCTGTAA

>ENA|AMB37295|AMB37295.3 Zika virus polyprotein

ATGAAAAACCC-AAAAAAGAAATCCGGAGGATTCCGGATTGTCAATATGCTAAAACGCGGAGTAGCCCGTGTGAGCCCCTTTGGGGGCTTGAAGAGGCTGCCAGCCGGACTTCTGCTGGGTCATGGGCCCATCAGGATGGTCTTGGCAATTCTAGCCTTTTTGAGATTCACGGCAATCAAGCCATCACTGGGTCTCATCAATAGATGGGGTTCAGTGGGGAAAAAAGAGGCTATGGAAATAATAAAGAAGTTCAAGAAAGATCTGGCTGCCATGCTGAGAATAATCAATGCTAGGAAGGAGAAGAAGAGACGAGGCGCAGATACTAGTGTCGGAATTGTTGGCCTCCTGCTGACCACAGCTATGGCAGCGGAGGTCACTAGACGTGGGAGTGCATACTATATGTACTTGGACAGAAACGATGCTGGGGAGGCCATATCTTTTCCAACCACATTGGGGATGAATAAGTGTTATATACAGATCATGGATCTTGGACACATGTGTGATGCCACCATGAGCTATGAATGCCCTATGCTGGATGAGGGGGTGGAACCAGATGACGTCGATTGTTGGTGCAACACGACGTCAACTTGGGTTGTGTACGGAACCTGCCATCACAAAAAAGGTGAAGCACGGAGATCTAGAAGAGCTGTGACGCTCCCCTCCCATTCCACTAGGAAGCTGCAAACGCGGTCGCAAACCTGGTTGGAATCAAGAGAATACACAAAGCACTTGATTAGAGTCGAAAATTGGATATTCAGGAACCCTGGCTTCGCGTTAGCAGCAGCTGCCATCGCTTGGCTTTTGGGAAGCTCAACGAGCCAAAAAGTCATATACTTGGTCATGATACTGCTGATTGCCCCGGCATACAGCATCAGGTGCATAGGAGTCAGCAATAGGGACTTTGTGGAAGGTATGTCAGGTGGGACTTGGGTTGATGTTGTCTTGGAACATGGAGGTTGTGTCACCGTAATGGCACAGGACAAACCGACTGTCGACATAGAGCTGGTTACAACAACAGTCAGCAACATGGCGGAGGTAAGATCCTACTGCTATGAGGCATCAATATCAGACATGGCTTCGGACAGCCGCTGCCCAACACAAGGTGAAGCCTACCTTGACAAGCAATCAGACACTCAATATGTCTGCAAAAGAACGTTAGTGGACAGAGGCTGGGGAAATGGATGTGGACTTTTTGGCAAAGGGAGTCTGGTGACATGCGCTAAGTTTGCATGCTCCAAGAAAATGACCGGGAAGAGCATCCAGCCAGAGAATCTGGAGTACCGGATAATGCTGTCAGTTCATGGCTCCCAGCACAGTGGGATGATCGTTAATGACACAGGACATGAAACTGATGAGAATAGAGCGAAGGTTGAGATAACGCCCAATTCACCAAGAGCCGAAGCCACCCTGGGGGGTTTTGGAAGCCTAGGACTTGATTGTGAACCGAGGACAGGCCTTGACTTTTCAGATTTGTATTACTTGACTATGAATAACAAGCACTGGTTGGTTCACAAGGAGTGGTTCCACGACATTCCATTACCTTGGCACGCTGGGGCAGACACCGGAACTCCACACTGGAACAACAAAGAAGCACTGGTAGAGTTCAAGGACGCACATGCCAAAAGGCAAACTGTCGTGGTTCTAGGGAGTCAAGAAGGAGCAGTTCACACGGCCCTTGCTGGAGCTCTGGAGGCTGAGATGGATGGTGCAAAGGGAAGGCTGTCCTCTGGCCACTTGAAATGTCGCCTGAAAATGGATAAACTTAGATTGAAGGGCGTGTCATACTCCTTGTGTACCGCAGCGTTCACATTCACCAAGATCCCGGCTGAAACACTGCACGGGACAGTCACAGTGGAGGTACAGTACGCAGGGACAGATGGACCTTGCAAGGTTCCAGCTCAGATGGCGGTGGACATGCAAACTCTGACCCCAGTTGGGAGGTTGATAACCGCTAACCCCGTAATCACTGAAAGCACTGAGAACTCTAAGATGATGCTGGAACTTGATCCACCATTTGGGGACTCTTACATTGTCATAGGAGTCGGGGAGAAGAAGATCACCCACCACTGGCACAGGAGTGGCAGCACCATTGGAAAAGCATTTGAAGCCACTGTGAGAGGTGCCAAGAGAATGGCAGTCTTGGGAGACACAGCCTGGGACTTTGGATCAGTTGGAGGCGCTCTCAACTCATTGGGCAAGGGCATCCATCAAATTTTTGGAGCAGCTTTCAAATCATTGTTTGGAGGAATGTCCTGGTTCTCACAAATTCTCATTGGAACGTTGCTGATGTGGTTGGGTCTGAACACAAAGAATGGATCTATTTCCCTTATGTGCTTGGCCTTAGGGGGAGTGTTGATCTTCTTATCCACAGCCGTCTCTGCTGATGTGGGGTGCTCGGTGGACTTCTCAAAGAAGGAGACGAGATGCGGTACAGGGGTGTTCGTCTATAACGACGTTGAAGCCTGGAGGGACAGGTACAAGTACCATCCTGACTCCCCCCGTAGATTGGCAGCAGCAGTCAAGCAAGCCTGGGAAGATGGTATCTGCGGGATCTCCTCTGTTTCAAGAATGGAAAACATCATGTGGAGATCAGTAGAAGGGGAGCTCAACGCAATCCTGGAAGAGAATGGAGTTCAACTGACGGTCGTTGTGGGATCTGTAAAAAACCCCATGTGGAGAGGTCCACAGAGATTGCCCGTGCCTGTGAACGAGCTGCCCCACGGCTGGAAGGCTTGGGGGAAATCGCACTTCGTCAGAGCAGCAAAGACAAATAACAGCTTTGTCGTGGATGGTGACACACTGAAGGAATGCCCACTCAAACATAGAGCATGGAACAGCTTTCTTGTGGAGGATCATGGGTTCGGGGTATTTCACACTAGTGTCTGGCTCAAGGTTAGAGAAGATTATTCATTAGAGTGTGATCCAGCCGTTATTGGAACAGCTGTTAAGGGAAAGGAGGCTGTACACAGTGATCTAGGCTACTGGATTGAGAGTGAGAAGAATGACACATGGAGGCTGAAGAGGGCCCATCTGATCGAGATGAAAACATGTGAATGGCCAAAGTCCCACACATTGTGGACAGATGGAATAGAAGAGAGTGATCTGATCATACCCAAGTCTTTAGCTGGGCCACTCAGCCATCACAATACCAGAGAGGGCTACAGGACCCAAATGAAAGGGCCATGGCACAGTGAAGAGCTTGAAATTCGGTTTGAGGAATGCCCAGGCACTAAGGTCCACGTGGAGGAAACATGTGGAACAAGAGGACCATCTCTGAGATCAACCACTGCAAGCGGAAGGGTGATCGAGGAATGGTGCTGCAGGGAGTGCACAATGCCCCCACTGTCGTTCCGGGCTAAAGATGGCTGTTGGTATGGAATGGAGATAAGGCCCAGGAAAGAACCAGAAAGCAACTTAGTAAGGTCAATGGTGACTGCAGGATCAACTGATCACATGGATCACTTCTCCCTTGGAGTGCTTGTGATTCTGCTCATGGTGCAGGAAGGGCTGAAGAAGAGAATGACCACAAAGATCATCATAAGCACATCAATGGCAGTGCTGGTAGCTATGATCCTGGGAGGATTTTCAATGAGTGACCTGGCTAAGCTTGCAATTTTGATGGGTGCCACCTTCGCGGAAATGAACACTGGAGGAGATGTAGCTCATCTGGCGCTGATAGCGGCATTCAAAGTCAGACCAGCGTTGCTGGTATCTTTCATCTTCAGAGCTAATTGGACACCCCGTGAAAGCATGCTGCTGGCCTTGGCCTCGTGTCTTTTGCAAACTGCGATCTCCGCCTTGGAAGGCGACCTGATGGTTCTCATCAATGGTTTTGCTTTGGCCTGGTTGGCAATACGAGCGATGGTTGTTCCACGCACTGATAACATCACCTTGGCAATCCTGGCTGCTCTGACACCACTGGCCCGGGGCACACTGCTTGTGGCGTGGAGAGCAGGCCTTGCTACTTGCGGGGGGTTTATGCTCCTCTCTCTGAAGGGAAAAGGCAGTGTGAAGAAGAACTTACCATTTGTCATGGCCCTGGGACTAACCGCTGTGAGGCTGGTCGACCCCATCAACGTGGTGGGGCTGCTGTTGCTCACAAGGAGTGGGAAGCGGAGCTGGCCCCCTAGCGAAGTACTCACAGCTGTTGGCCTGATATGCGCATTGGCTGGAGGGTTCGCCAAGGCAGATATAGAGATGGCTGGGCCCATGGCCGCGGTCGGTCTGCTAATTGTCAGTTACGTGGTCTCAGGAAAGAGTGTGGACATGTACATTGAAAGAGCAGGTGACATCACATGGGAAAAAGATGCGGAAGTCACTGGAAACAGTCCCCGGCTCGATGTGGCGCTAGATGAGAGTGGTGATTTCTCCCTGGTGGAGGATGACGGTCCCCCCATGAGAGAGATCATACTCAAGGTGGTCCTGATGACCATCTGTGGCATGAACCCAATAGCCATACCCTTTGCAGCTGGAGCGTGGTACGTATACGTGAAGACTGGAAAAAGGAGTGGTGCTCTATGGGATGTGCCTGCTCCCAAGGAAGTAAAAAAGGGGGAGACCACAGATGGAGTGTACAGAGTAATGACTCGTAGACTGCTAGGTTCAACACAAGTTGGAGTGGGAGTTATGCAAGAGGGGGTCTTTCACACTATGTGGCACGTCACAAAAGGATCCGCGCTGAGAAGCGGTGAAGGGAGACTTGATCCATACTGGGGAGATGTCAAGCAGGATCTGGTGTCATACTGTGGTCCATGGAAGCTAGATGCCGCCTGGGACGGGCACAGCGAGGTGCAGCTCTTGGCCGTGCCCCCCGGAGAGAGAGCGAGGAACATCCAGACTCTGCCCGGAATATTTAAGACAAAGGATGGGGACATTGGAGCGGTTGCGCTGGATTACCCAGCAGGAACTTCAGGATCTCCAATCCTAGACAAGTGTGGGAGAGTGATAGGACTTTATGGCAATGGGGTCGTGATCAAAAATGGGAGTTATGTTAGTGCCATCACCCAAGGGAGGAGGGAGGAAGAGACTCCTGTTGAGTGCTTCGAGCCTTCGATGCTGAAGAAGAAGCAGCTAACTGTCTTAGACTTGCATCCTGGAGCTGGGAAAACCAGGAGAGTTCTTCCTGAAATAGTCCGTGAAGCCATAAAAACAAGACTCCGTACTGTGATCTTAGCTCCAACCAGGGTTGTCGCTGCTGAAATGGAGGAAGCCCTTAGAGGGCTTCCAGTGCGTTATATGACAACAGCAGTCAATGTCACCCACTCTGGAACAGAAATCGTCGACTTAATGTGCCATGCCACCTTCACTTCACGTCTACTACAGCCAATCAGAGTCCCCAACTATAATCTGTATATTATGGATGAGGCCCACTTCACAGATCCCTCAAGTATAGCAGCAAGAGGATACATTTCAACAAGGGTTGAGATGGGCGAGGCGGCTGCCATCTTCATGACCGCCACGCCACCAGGAACCCGTGACGCATTTCCGGACTCCAACTCACCAATTATGGACACCGAAGTGGAAGTCCCAGAGAGAGCCTGGAGCTCAGGCTTTGATTGGGTGACGGATTATTCTGGAAAAACAGTTTGGTTTGTTCCAAGCGTGAGGAACGGCAATGAGATCGCAGCTTGTCTGACAAAGGCTGGAAAACGGGTCATACAGCTCAGCAGAAAGACTTTTGAGACAGAGTTCCAGAAAACAAAACATCAAGAGTGGGACTTTGTCGTGACAACTGACATTTCAGAGATGGGCGCCAACTTTAAAGCTGACCGTGTCATAGATTCCAGGAGATGCCTAAAGCCGGTCATACTTGATGGCGAGAGAGTCATTCTGGCTGGACCCATGCCTGTCACACATGCCAGCGCTGCCCAGAGGAGGGGGCGCATAGGCAGGAATCCCAACAAACCTGGAGATGAGTATCTGTATGGAGGTGGGTGCGCAGAGACTGACGAAGACCATGCACACTGGCTTGAAGCAAGAATGCTCCTTGACAATATTTACCTCCAAGATGGCCTCATAGCCTCGCTCTATCGACCTGAGGCCGACAAAGTAGCAGCCATTGAGGGAGAGTTCAAGCTTAGGACGGAGCAAAGGAAGACCTTTGTGGAACTCATGAAAAGAGGAGATCTTCCTGTTTGGCTGGCCTATCAGGTTGCATCTGCCGGAATAACCTACACAGATAGAAGATGGTGCTTTGATGGCACGACCAACAACACCATAATGGAAGACAGTGTGCCGGCAGAGGTGTGGACCAGACACGGAGAGAAAAGAGTGCTCAAACCGAGGTGGATGGACGCCAGAGTTTGTTCAGATCATGCGGCCCTGAAGTCATTCAAGGAGTTTGCCGCTGGGAAAAGAGGAGCGGCTTTTGGAGTGATGGAAGCCCTGGGAACACTGCCAGGACACATGACAGAGAGATTCCAGGAAGCCATTGACAACCTCGCTGTGCTCATGCGGGCAGAGACTGGAAGCAGGCCTTACAAAGCCGCGGCGGCCCAATTGCCGGAGACCCTAGAGACCATTATGCTTTTGGGGTTGCTGGGAACAGTCTCGCTGGGAATCTTTTTCGTCTTGATGAGGAACAAGGGCATAGGGAAGATGGGCTTTGGAATGGTGACTCTTGGGGCCAGCGCATGGCTCATGTGGCTCTCGGAAATTGAGCCAGCCAGAATTGCATGTGTCCTCATTGTTGTGTTCCTATTGCTGGTGGTGCTCATACCTGAGCCAGAAAAGCAAAGATCTCCCCAGGACAACCAAATGGCAATCATCATCATGGTAGCAGTAGGTCTTCTGGGCTTGATTACCGCCAATGAACTCGGATGGTTGGAGAGAACAAAGAGTGACCTAAGCCATCTAATGGGAAGGAGAGAGGAGGGGGCAACCATGGGATTCTCAATGGACATTGACCTGCGGCCAGCCTCAGCTTGGGCCATCTATGCTGCCTTGACAACTTTCATTACCCCAGCCGTCCAACATGCAGTGACCACTTCATACAACAACTACTCCTTAATGGCGATGGCCACGCAAGCTGGAGTGTTGTTTGGTATGGGCAAAGGGATGCCATTCTACGCATGGG-ACTTTGGAGTCCCGCTGCTAATGATAGGTTGCTACTCACAATTAACGCCCCTGACCCTAATAGTGGCCATCATTTTGCTCGTGGCGCACTACATGTACTTGATCCCAGGGCTGCAGGCAGCAGCTGCGCGTGCTGCCCAGAAGAGAACGGCAGCTGGCATCATGAAGAACCCTGTTGTGGATGGAATAGTGGTGACTGACATTGACACAATGACAATTGACCCCCAAGTGGAGAAAAAGATGGGACAGGTGCTACTCATGGCAGTAGCCGTCTCCAGCGCCATACTGTCGCGGACCGCCTGGGGGTGGGGGGAGGCTGGGGCCCTGATCACAGCCGCAACTTCCACTTTGTGGGAAGGCTCTCCGAACAAGTACTGGAACTCCTCTACAGCCACTTCACTGTGTAACATTTTTAGGGGAAGTTACTTGGCTGGAGCTTCTCTAATCTACACAGTAACAAGAAACGCTGGCTTGGTCAAGAGACGTGGGGGTGGAACAGGAGAGACCCTGGGAGAGAAATGGAAGGCCCGCTTGAACCAGATGTCGGCCCTGGAGTTCTACTCCTACAAAAAGTCAGGCATCACCGAGGTGTGCAGAGAAGAGGCCCGCCGCGCCCTCAAGGACGGTGTGGCAACGGGAGGCCATGCTGTGTCCCGAGGAAGTGCAAAGCTGAGATGGTTGGTGGAGCGGGGATACCTGCAGCCCTATGGAAAGGTCATTGATCTTGGATGTGGCAGAGGGGGCTGGAGTTACTACGCCGCCACCATCCGCAAAGTTCAAGAAGTGAAAGGATACACAAAAGGAGGCCCTGGTCATGAAGAACCCGTGTTGGTGCAAAGCTATGGGTGGAACATAGTCCGTCTTAAGAGTGGGGTGGACGTCTTTCATATGGCGGCTGAGCCGTGTGACACGTTGCTGTGTGACATAGGTGAGTCATCATCTAGTCCTGAAGTGGAAGAAGCACGGACGCTCAGAGTCCTCTCCATGGTGGGGGATTGGCTTGAAAAAAGACCAGGAGCCTTTTGTATAAAAGTGTTGTGCCCATACACCAGCACTATGATGGAAACCCTGGAGCGACTGCAGCGTAGGTATGGGGGAGGACTGGTCAGAGTGCCACTCTCCCGCAACTCTACACATGAGATGTACTGGGTCTCTGGAGCGAAAAGCAACACCATAAAAAGTGTGTCCACCACGAGCCAGCTCCTCTTGGGGCGCATGGACGGGCCTAGGAGGCCAGTGAAATATGAGGAGGATGTGAATCTCGGCTCTGGCACGCGGGCTGTGGTAAGCTGCGCTGAAGCTCCCAACATGAAGATCATTGGTAACCGCATTGAAAGGATCCGCAGTGAGCACGCGGAAACGTGGTTCTTTGACGAGAACCACCCATATAGGACATGGGCTTACCATGGAAGCTATGAGGCCCCCACACAAGGGTCAGCGTCCTCTCTAATAAACGGGGTTGTCAGGCTCCTGTCAAAACCCTGGGATGTGGTGACTGGAGTCACAGGAATAGCCATGACCGACACCACACCGTATGGTCAGCAAAGAGTTTTCAAGGAAAAAGTGGACACTAGGGTGCCAGACCCCCAAGAAGGCACTCGTCAGGTTATGAGCATGGTCTCTTCCTGGTTGTGGAAAGAGCTAGGCAAACACAAACGGCCACGAGTCTGTACCAAAGAAGAGTTCATCAACAAGGTTCGTAGCAATGCAGCATTAGGGGCAATATTTGAAGAGGAAAAAGAGTGGAAGACTGCAGTGGAAGCTGTGAACGATCCAAGGTTCTGGGCTCTAGTGGACAAGGAAAGAGAGCACCACCTGAGAGGAGAGTGCCAGAGTTGTGTGTACAACATGATGGGAAAAAGAGAAAAGAAACAAGGGGAATTTGGAAAGGCCAAGGGCAGCCGCGCCATCTGGTATATGTGGCTAGGGGCTAGATTTCTAGAGTTCGAAGCCCTTGGATTCTTGAACGAGGATCACTGGATGGGGAGAGAGAACTCAGGAGGTGGTGTTGAAGGGCTGGGATTACAAAGACTCGGATATGTCCTAGAAGAGATGAGTCGCATACCAGGAGGAAGGATGTATGCAGATGACACTGCTGGCTGGGACACCCGCATCAGCAGGTTTGATCTGGAGAATGAAGCTCTAATCACCAACCAAATGGAGAAAGGGCACAGGGCCTTGGCATTGGCCATAATCAAGTACACATACCAAAACAAAGTGGTAAAGGTCCTTAGACCAGCTGAAAAAGGGAAGACAGTTATGGACATTATTTCGAGACAAGACCAAAGGGGGAGCGGACAAGTTGTCACTTACGCTCTTAACACATTTACCAACCTAGTGGTGCAACTCATTCGGAATATGGAGGCTGAGGAAGTTCTAGAGATGCAAGACTTGTGGCTGCTGCGGAGGTCAGAGAAAGTGACCAACTGGTTGCAGAGCAACGGATGGGATAGGCTCAAACGAATGGCAGTCAGTGGAGATGATTGCGTTGTGAAGCCAATTGATGATAGGTTTGCACATGCCCTCAGGTTCTTGAATGATATGGGAAAAGTTAGGAAGGACACACAAGAGTGGAAACCCTCAACTGGATGGGACAACTGGGAAGAAGTTCCGTTTTGCTCCCACCACTTCAACAAGCTCCATCTCAAGGACGGGAGGTCCATTGTGGTTCCCTGCCGCCACCAAGATGAACTGATTGGCCGGGCCCGCGTCTCTCCAGGGGCGGGATGGAGCATCCGGGAGACTGCTTGCCTAGCAAAATCATATGCGCAAATGTGGCAGCTCCTTTATTTCCACAGAAGGGACCTCCGACTGATGGCCAATGCCATTTGTTCATCTGTGCCAGTTGACTGGGTTCCAACTGGGAGAACTACCTGGTCAATCCATGGAAAGGGAGAATGGATGACCACTGAAGACATGCTTGTGGTGTGGAACAGAGTGTGGATTGAGGAGAACGACCACATGGAAGACAAGACCCCAGTTACGAAATGGACAGACATTCCCTATTTGGGAAAAAGGGAAGACTTGTGGTGTGGATCTCTCATAGGGCACAGACCGCGCACCACCTGGGCTGAGAACATTAAAAACACAGTCAACATGGTGCGCAGGATCATAGGTGATGAAGAAAAGTACATGGACTACCTATCCACCCAAGTTCGCTACTTGGGTGAAGAAGGGTCTACACCTGGAGTGCTGTAA

>ENA|AMC33116|AMC33116.1 Zika virus polyprotein

ATGAAAAACCC-AAAAAAGAAATCCGGAGGATTCCGGATTGTCAATATGCTAAAACGCGGAGTAGCCCGTGTGAGCCCCTTTGGGGGCTTGAAGAGGCTGCCAGCCGGACTTCTGCTGGGTCATGGGCCCATCAGGATGGTCTTGGCGATTCTAGCCTTTTTGAGATTCACGGCAATCAAGCCATCACTGGGTCTCATCAATAGATGGGGTTCAGTGGGGAAAAAAGAGGCTATGGAAATAATAAAGAAGTTCAAGAAAGATCTGGCTGCCATGCTGAGAATAATCAATGCTAGGAAGGAGAAGAAGAGACGAGGCGCAGAAACTAGTGTCGGAATTGTTGGCCTCCTGCTGACCACAGCTATGGCAGCGGAGGTCACTAGACGTGGGAGTGCATACTATATGTACTTGGACAGAAACGATGCTGGGGAGGCCATATCTTTTCCAACCACATTGGGGATGAATAAGTGTTATATACAGATCATGGATCTTGGACACATGTGTGATGCCACCATGAGCTATGAATGCCCTATGCTGGATGAGGGGGTGGAACCAGATGACGTCGATTGTTGGTGCAACACGACGTCAACTTGGGTTGTGTACGGAACCTGCCATCACAAAAAAGGTGAAGCACGGAGATCTAGAAGAGCCGTGACGCTCCCCTCCCATTCCACTAGGAAGCTGCAAACGCGGTCGCAAACCTGGTTGGAATCAAGAGAATACACAAAGCACTTGATTAGAGTCGAAAATTGGATATTCAGGAACCCTGGTTTCGCTTTAGCAGCAGCTGCCATCGCGTGGCTTTTGGGAAGCTCAACGAGCCAAAAAGTCATATACTTGGTCATGATACTGCTGATTGCCCCGGCATACAGCATCAGGTGCATAGGAGTCAGCAATAGGGACTTTGTGGAAGGTATGTCAGGTGGGACTTGGGTTGATGTTGTCTTGGAACATGGAGGTTGTGTCACCGTAATGGCACAGGACAAACCGACTGTCGACATAGAGCTGGTTACAACAACAGTCAGCAACATGGCGGAGGTAAGATCCTACTGCTATGAGGCATCAATATCAGACATGGCTTCGGACAGCCGCTGCCCAACACAAGGTGAAGCCTACCTTGACAAGCAATCAGACACTCAATATGTCTGCAAAAGAACGTTAGTGGACAGAGGCTGGGGAAATGGATGTGGACTTTTTGGCAAAGGGAGCCTGGTGACATGCGCTAAGTTTGCATGCTCCAAGAAAATGACCGGGAAGAGCATCCAGCCAGAGAATCTGGAGTACCGGATAATGTTGTCAGTTCATGGCTCCCAGCACAGTGGGATGATCGTTAATGACACAGGACATGAAACTGATGAGAATAGAGCGAAGGTTGAGATAACGCCCAATTCACCAAGAGCCGAAGCCACCCTGGGGGGTTTTGGAAGCCTAGGACTTGATTGTGAACCGAGGACAGGCCTTGACTTTTCAGATTTGTATTACTTGACTATGAATAACAAGCACTGGTTGGTTCACAAGGAGTGGTTCCACGACATTCCATTACCTTGGCACGCTGGGGCAGACACCGGAACTCCACACTGGAACAACAAAGAAGCACTGGTAGAGTTCAAGGACGCACATGCCAAAAGGCAAACTGTCGTGGTTCTAGGGAGTCAAGAAGGAGCAGTTCACACGGCCCTTGCTGGAGCTCTGGAGGCTGAGATGGATGGTGCAAAGGGAAGGCTGTCCTCTGGCCACTTGAAATGTCGCCTGAAAATGGATAAACTTAGATTGAAGGGCGTGTCATACTCCTTGTGTACCGCAGCGTTCACATTCACCAAGATCCCGGCTGAAACACTGCACGGGACAGTCACAGTGGAGGTACAGTACGCAGGGACAGATGGACCTTGCAAGGTTCCAGCCCAGATGGCGGTGGACATGCAAACTCTGACCCCAGTTGGGAGGTTGATAACCGCTAACCCCGTAATCACTGAAAGCACTGAGAACTCTAAGATGATGCTGGAACTTGATCCACCATTTGGGGACTCTTACATTGTCATAGGAGTCGGGGAGAAGAAGATCACCCACCACTGGCACAGGAGTGGCAGCACCATTGGAAAAGCATTTGAAGCCACTGTGAGAGGTGCCAAGAGAATGGCAGTCTTGGGAGACACAGCCTGGGACTTTGGATCAGTTGGAGGCGCTCTCAACTCATTGGGCAAGGGCATCCATCAAATTTTTGGAGCAGCTTTCAAATCATTGTTTGGAGGAATGTCCTGGTTCTCACAAATTCTCATTGGAACGTTGCTGATGTGGTTGGGTCTGAACACAAAGAATGGATCTATTTCCCTTATGTGCTTGGCCTTAGGGGGAGTGTTGATCTTCTTATCCACAGCCGTCTCTGCTGATGTGGGGTGCTCGGTGGACTTCTCAAAGAAGGAGACGAGATGTGGTACAGGGGTGTTCGTCTATAACGACGTTGAAGCCTGGAGGGACAGGTACAAGTACCATCCTGACTCCCCCCGTAGATTGGCAGCAGCAGTCAAGCAAGCCTGGGAAGATGGTATCTGCGGGATCTCCTCTGTTTCAAGAATGGAAAACATCATGTGGAGATCAGTAGAAGGGGAGCTCAACGCAATCCTGGAAGAGAATGGAGTTCAACTGACGGTCGTTGTGGGATCTGTAAAAAACCCCATGTGGAGAGGTCCACAGAGATTGCCCGTGCCTGTGAACGAGCTGCCCCACGGCTGGAAGGCTTGGGGGAAATCGTACTTCGTCAGAGCAGCAAAGACAAATAACAGCTTTGTCGTGGATGGTGACACACTGAAGGAATGCCCACTCAAACATAGAGCATGGAACAGCTTTCTTGTGGAGGATCATGGGTTCGGGGTATTTCACACTAGTGTCTGGCTCAAGGTTAGAGAAGATTATTCATTAGAGTGTGATCCAGCCGTTATTGGAACAGCTGTTAAGGGAAAGGAGGCTGTACACAGTGATCTAGGCTACTGGATTGAGAGTGAGAAGAATGACACATGGAGGCTGAAGAGGGCCCATCTGATCGAGATGAAAACATGTGAATGGCCAAAATCCCACACATTGTGGACAGATGGAATAGAAGAGAGTGATCTGATCATACCCAAGTCTTTAGCTGGGCCACTCAGCCATCACAATACCAGAGAGGGCTACAGGACCCAAATGAAAGGGCCATGGCACAGTGAAGAGCTTGAAATTCGGTTTGAGGAATGCCCAGGCACTAAGGTCCACGTGGAGGAAACATGTGGAACAAGAGGACCATCTCTGAGATCAACCACTGCAAGCGGAAGGGTGATCGAGGAATGGTGCTGCAGGGAGTGCACAATGCCCCCACTGTCGTTCTGGGCTAAAGATGGCTGTTGGTATGGAATGGAGATAAGGCCCAGGAAAGAACCAGAAAGCAACTTAGTAAGGTCAATGGTGACTGCAGGATCAACTGATCACATGGATCACTTCTCCCTTGGAGTGCTTGTGATTCTGCTCATGGTGCAGGAAGGGCTGAAGAAGAGAATGACCACAAAGATCATCATAAGCACATCAATGGCAGTGCTGGTAGCTATGATCCTGGGAGGATTTTCAATGAGTGACCTGGCTAAGCTTGCAATTTTGATGGGTGCCACCTTCGCGGAAATGAACACTGGAGGAGATGTAGCTCATCTGGCGCTGACAGCGGCATTCAAAGTCAGACCAGCGTTGCTGGTATCTTTCATCTTCAGAGCTAATTGGACACCCCGTGAAAGCATGCTGCTGGCCTTGGCCTCGTGTCTTTTGCAAACTGCGATCTCTGCCTTGGAAGGCGACCTGATGGTTCTCATCAATGGTTTTGCTTTGGCCTGGTTGGCAATACGAGCGATGGTTGTTCCACGCACTGACAACATCACCTTGGCAATCCTGGCTGCTCTGACACCACTGGCCCGGGGCACACTGCTTGTGGCGTGGAGAGCAGGCCTTGCTACTTGCGGGGGGTTTATGCTCCTCTCTCTGAAGGGAAAAGGCAGTGTGAAGAAGAACTTACCATTTGTCATGGCCCTGGGACTAACCGCTGTGAGGCTGGTCGACCCCATCAACGTGGTGGGACTGCTGTTGCTCACAAGGAGTGGGAAGCGGAGCTGGCCCCCTAGCGAAGTACTCACAGCTGTTGGCCTGATATGTGCATTGGCTGGAGGGTTCGCCAAGGCAGATATAGAGATGGCTGGGCCCATGGCCGCGGTCGGTCTGCTAATTGTCAGTTACGTGGTCTCAGGAAAGAGTGTGGACATGTACATTGAAAGAGCAGGTGACATCACATGGGAAAAAGATGCGGAAGTCACTGGAAACAGTCCCCGGCTCGACGTGGCGCTAGATGAGAGTGGTGATTTCTCCCTGGTGGAGGATGACGGTCCCCCCATGAGAGAGATCATACTCAAGGTGGTCCTGATGACCATCTGTGGCATGAACCCAATAGCCATACCCTTTGCAGCTGGAGCGTGGTACGTATACGTGAAGACTGGAAAAAGGAGTGGTGCTCTATGGGATGTGCCTGCTCCCAAGGAAGTAAAAAAGGGGGAGACCACAGATGGAGTGTACAGAGTAATGACTCGTAGACTGCTAGGTTCAACACAAGTTGGAGTGGGAGTTATGCAAGAGGGGGTCTTTCACACTATGTGGCACGTCACAAAAGGATCCGCGCTGAGAAGCGGTGAAGGGAGACTTGATCCATACTGGGGAGATGTCAAGCAGGATCTGGTGTCATACTGTGGTCCATGGAAGCTAGATGCCGCCTGGGACGGGCACAGCGAGGTGCAGCTCTTGGCCGTGCCCCCCGGAGAGAGAGCGAGGAACATCCAGACTCTGCCCGGAATATTTAAGACAAAGGATGGGGACATTGGAGCGGTTGCGCTGGATTACCCAGCAGGAACTTCAGGATCTCCAATCCTAGACAAGTGTGGGAGAGTGATAGGACTTTATGGCAATGGGGTCGTGATCAAAAATGGGAGTTATGTTAGTGCCATCACCCAAGGGAGGAGGGAGGAAGAGACTCCTGTTGAGTGCTTCGAGCCTTCGATGCTGAAGAAGAAGCAGCTAACTGTCTTAGACTTGCATCCTGGAGCTGGGAAAACCAGGAGAGTTCTTCCTGAAATAGTCCGTGAAGCCATAAAAACAAGACTCCGTACTGTGATCTTAGCTCCAACCAGGGTTGTCGCTGCTGAAATGGAGGAAGCCCTTAGAGGGCTTCCAGTGCGTTATATGACAACAGCAGTCAATGTCACCCACTCTGGAACAGAAATCGTCGACTTAATGTGCCATGCCACCTTCACTTCACGTCTACTACAGCCAATCAGAGTCCCCAACTATAATCTGTATATTATGGATGAGGCCCACTTCACAGATCCCTCAAGTATAGCAGCAAGAGGATACATTTCAACAAGGGTTGAGATGGGCGAGGCGGCTGCCATCTTCATGACCGCCACGCCACCAGGAACCCGTGACGCATTTCCGGACTCCAACTCACCAATTATGGACACCGAAGTGGAAGTCCCAGAGAGAGCCTGGAGCTCAGGCTTTGATTGGGTGACGGATCATTCTGGAAAAACAGTTTGGTTTGTTCCAAGCGTGAGGAACGGCAATGAGATCGCAGCTTGTCTGACAAAGGCTGGAAAACGGGTCATACAGCTCAGCAGAAAGACTTTTGAGACAGAGTTCCAGAAAACAAAACATCAAGAGTGGGACTTTGTCGTGACAACTGACATTTCAGAGATGGGCGCCAACTTTAAAGCTGACCGTGTCATAGATTCCAGGAGATGCCTAAAGCCGGTCATACTTGATGGCGAGAGAGTCATTCTGGCTGGACCCATGCCTGTCACACATGCCAGCGCTGCCCAGAGGAGGGGGCGCATAGGCAGGAATCCCAATAAACCTGGAGATGAGTATCTGTATGGAGGTGGGTGCGCAGAGACTGACGAAGACCATGCACACTGGCTTGAAGCAAGAATGCTCCTTGACAATATTTACCTCCAAGATGGCCTCATAGCCTCGCTCTATCGACCTGAGGCCGACAAAGTAGCAGCCATTGAGGGAGAGTTCAAGCTTAGGACGGAGCAAAGGAAGACCTTTGTGGAACTCATGAAAAGAGGAGATCTTCCTGTTTGGCTGGCCTATCAGGTTGCATCTGCCGGAATAACCTACACAGATAGAAGATGGTGCTTTGATGGCACGACCAACAACACCATAATGGAAGACAGTGTGCCGGCAGAGGTGTGGACCAGACACGGAGAGAAAAGAGTGCTCAAACCGAGGTGGATGGACGCCAGAGTTTGTTCAGATCATGCGGCCCTGAAGTCATTCAAGGAGTTTGCCGCTGGGAAAAGAGGAGCGGCTTTTGGAGTGATGGAAGCCCTGGGAACACTGCCAGGACACATGACAGAGAGATTCCAGGAAGCCATTGACAACCTCGCTGTGCTCATGCGGGCAGAGACTGGAAGCAGGCCTTACAAAGCCGCGGCGGCCCAATTGCCGGAGACCCTAGAGACCATTATGCTTTTGGGGTTGCTGGGAACAGTCTCGCTGGGAATCTTTTTCGTCTTGATGAGGAACAAGGGCATAGGGAAGATGGGCTTTGGAATGGTGACTCTTGGGGCCAGCGCATGGCTCATGTGGCTCTCGGAAATTGAGCCAGCCAGAATTGCATGTGTCCTCATTGTTGTGTTCCTATTGCTGGTGGTGCTCATACCTGAGCCAGAAAAGCAAAGATCTCCCCAGGACAACCAAATGGCAATCATCATCATGGTAGCAGTAGGTCTTCTGGGCTTGATTACCGCCAATGAACTCGGATGGTTGGAGAGAACAAAGAGTGACCTAAGCCATCTAATGGGAAGGAGAGAGGAGGGGGCAACCATAGGATTCTCAATGGACATTGACCTGCGGCCAGCCTCAGCTTGGGCCATCTATGCTGCCTTGACAACTTTCATTACCCCAGCCGTCCAACATGCAGTGACCACTTCATACAACAACTACTCCTTAATGGCGATGGCCACGCAAGCTGGAGTGTTGTTTGGTATGGGCAAAGGGATGCCATTCTACGCATGGG-ACTTTGGAGTCCCACTGCTAATGATAGGTTGCTACTCACAATTAACACCCCTGACCCTAATAGTGGCCATCATTTTGCTCGTGGCGCACTACATGTACTTGATCCCAGGGCTGCAGGCAGCAGCTGCGCGTGCTGCCCAGAAGAGAACGGCAGCTGGCATCATGAAGAACCCTGTTGTGGATGGAATAGTGGTGACTGACATTGACACAATGACAATTGACCCCCAAGTGGAGAAAAAGATGGGACAGGTGCTACTCATAGCAGTAGCCGTCTCCAGCGCCATACTGTCGCGGACCGCCTGGGGGTGGGGGGAGGCTGGGGCCCTGATCACAGCCGCAACTTCCACTTTGTGGGAAGGCTCTCCGAACAAGTACTGGAACTCCTCTACAGCCACTTCACTGTGTAACATTTTTAGGGGAAGTTACTTGGCTGGAGCTTCTCTAATCTACACAGTAACAAGAAACGCTGGCTTGGTCAAGAGACGTGGGGGTGGAACAGGAGAGACCCTGGGAGAGAAATGGAAGGCCCGCTTGAACCAGATGTCGGCCCTGGAGTTCTACTCCTACAAAAAGTCAGGCATCACCGAGGTGTGCAGAGAAGAGGCCCGCCGCGCCCTCAAGGACGGTGTGGCAACGGGAGGCCATGCTGTGTCCCGAGGAAGTGCAAAGCTGAGATGGTTGGTGGAGCGGGGATACCTGCAGCCCTATGGAAAGGTCATTGATCTTGGATGTGGCAGAGGGGGCTGGAGTTACTACGCCGCCACCATCCGCAAAGTTCAAGAAGTGAAAGGATACACAAAAGGAGGCCCTGGTCATGAAGAACCCGTGTTGGTGCAAAGCTATGGGTGGAACATAGTCCGTCTTAAGAGTGGGGTGGACGTCTTTCATATGGCGGCTGAGCCGTGTGACACGTTGCTGTGTGACATAGGTGAGTCATCATCTAGTCCTGAAGTGGAAGAAGCACGGACGCTCAGAGTCCTCTCCATGGTGGGGGATTGGCTTGAAAAAAGACCAGGAGCCTTTTGTATAAAAGTGTTGTGCCCATACACCAGCACTATGATGGAAACCCTGGAGCGACTGCAGCGTAGGTATGGGGGAGGACTGGTCAGAGTGCCACTCTCCCGCAACTCTACACATGAGATGTACTGGGTCTCTGGAGCGAAAAGCAACACCATAAAAAGTGTGTCCACCACGAGCCAGCTCCTCTTGGGGCGCATGGACGGGCCTAGGAGGCCAGTGAAATATGAGGAGGATGTGAATCTCGGCTCTGGCACGCGGGCTGTGGTAAGCTGCGCTGAAGCTCCCAACATGAAGATCATTGGTAACCGCATTGAAAGGATCCGCAGTGAGCACGCGGAAACGTGGTTCTTTGACGAGAACCACCCATATAGGACATGGGCTTACCATGGAAGCTATGAGGCCCCCACACAAGGGTCAGCGTCCTCTCTAATAAACGGGGTTGTCAGGCTCCTGTCAAAACCCTGGGATGTGGTGACTGGAGTCACAGGAATAGCCATGACCGACACCACACCGTATGGTCAGCAAAGAGTTTTCAAGGAAAAAGTGGACACTAGGGTGCCAGACCCCCAAGAAGGCACTCGTCAGGTTATGAGCATGGTCTCTTCCTGGTTGTGGAAAGAGCTAGGCAAACACAAACGGCCACGAGTCTGTACCAAAGAAGAGTTCATCAACAAGGTTCGTAGCAATGCAGCATTAGGGGCAATATTTGAAGAGGAAAAAGAGTGGAAGACTGCAGTGGAAGCTGTGAACGATCCAAGGTTCTGGGCTCTAGTGGACAAGGAAAGAGAGCACCACCTGAGAGGAGAGTGCCAGAGTTGTGTGTACAACATGATGGGAAAAAGAGAAAAGAAACAAGGGGAATTTGGAAAGGCCAAGGGCAGCCGCGCCATCTGGTATATGTGGCTAGGGGCTAGATTTCTAGAGTTCGAAGCCCTTGGATTCTTGAACGAGGATCACTGGATGGGGAGAGAGAACTCAGGAGGTGGTGTTGAAGGGCTGGGATTACAAAGACTCGGATATGTCCTAGAAGAGATGAGTCGCATACCAGGAGGAAGGATGTATGCAGATGACACTGCTGGCTGGGACACCCGCATCAGCAGGTTTGATCTGGAGAATGAAGCTCTAATCACCAACCAAATGGAGAAAGGGCACAGGGCCTTGGCATTGGCCATAATCAAGTACACATACCAAAACAAAGTGGTAAAGGTCCTTAGACCAGCTGAAAAAGGGAAAACAGTTATGGACATTATTTCGAGACAAGACCAAAGGGGGAGCGGACAAGTTGTCACTTACGCTCTTAACACATTTACCAACCTAGTGGTGCAACTCATTCGGAATATGGAGGCTGAGGAAGTTCTAGAGATGCAAGACTTGTGGCTGCTGCGGAGGTCAGAGAAAGTGACCAACTGGTTGCAGAGCAACGGATGGGATAGGCTCAAACGAATGGCAGTCAGTGGAGATGATTGCGTTGTGAAGCCAATTGATGATAGGTTTGCACATGCCCTCAGGTTCTTGAATGATATGGGAAAAGTTAGGAAGGACACACAAGAGTGGAAACCCTCAACTGGATGGGACAACTGGGAAGAAGTTCCGTTTTGCTCCCACCACTTCAACAAGCTCCATCTCAAGGACGGGAGGTCCATTGTGGTTCCCTGCCGCCACCAAGATGAACTGATTGGCCGGGCCCGCGTCTCTCCAGGGGCGGGATGGAGCATCCGGGAGACTGCTTGCCTAGCAAAATCATATGCGCAAATGTGGCAGCTCCTTTATTTCCACAGAAGGGACCTCCGACTGATGGCCAATGCCATTTGTTCATCTGTGCCAGTTGACTGGGTTCCAACTGGGAGAACTACCTGGTCAATCCATGGAAAGGGAGAATGGATGACCACTGAAGACATGCTTGTGGTGTGGAACAGAGTGTGGATTGAGGAGAACGACCACATGGAAGACAAGACCCCAGTTGCGAAATGGACAGACATTCCCTATTTGGGAAAAAGGGAAGACTTGTGGTGTGGATCTCTCATAGGGCACAGACCGCGCACCACCTGGGCTGAGAACATTAAAAACACAGTCAACATGGTGCGCAGGATCATAGGTGATGAAGAAAAGTACATGGACTACCTATCCACCCAAGTTCGCTACTTGGGTGAAGAAGGGTCTACACCTGGAGTGCTGTAA

>ENA|AMC13911|AMC13911.1 Zika virus polyprotein

ATGAAAAACCC-AAAAAAGAAATCCGGAGGATTCCGGATTGTCAATATGCTAAAACGCGGAGTAGCCCGTGTGAGCCCCTTTGGGGGCTTGAAGAGGCTGCCAGCCGGACTTCTGCTGGGTCATGGGCCCATCAGGATGGTCTTGGCGATTCTAGCCTTTTTGAGATTCACGGCAATCAAGCCATCACTGGGTCTCATCAATAGATGGGGTTCAGTGGGGAAAAAAGAGGCTATGGAAACAATAAAGAAGTTCAAGAAAGATCTGGCTGCCATGCTGAGAATAATCAATGCTAGGAAGGAGAAGAAGAGACGAGGCGCAGATACTAGTGTCGGAATTGTTGGCCTCCTGCTGACCACAGCTATGGCAGCGGAGGTCACTAGACGTGGGAGTGCATACTATATGTACTTGGACAGAAACGATGCTGGGGAGGCCATATCTTTTCCAACCACATTGGGGATGAATAAGTGTTATATACAGATCATGGATCTTGGACACATGTGTGATGCCACCATGAGCTATGAATGCCCTATGCTGGATGAGGGGGTGGAACCAGATGACGTCGATTGTTGGTGCAACACGACGTCAACTTGGGTTGTGTACGGAACCTGCCATCACAAAAAAGGTGAAGCACGGAGATCTAGAAGAGCTGTGACGCTCCCCTCCCATTCCACCAGGAAGCTGCAAACGCGGTCGCAAACCTGGTTGGAATCAAGAGAATACACAAAGCACTTGATTAGAGTCGAAAATTGGATATTCAGGAACCCTGGCTTCGCGTTAGCAGCAGCTGCCATCGCTTGGCTTTTGGGAAGCTCAACGAGCCAAAAAGTCATATACTTGGTCATGATACTGCTGATTGCCCCGGCATACAGCATCAGGTGCATAGGAGTCAGCAATAGGGACTTTGTGGAAGGTATGTCAGGTGGGACTTGGGTTGATGTTGTCTTGGAACATGGAGGTTGTGTCACCGTAATGGCACAGGACAAACCGACTGTCGACATAGAGCTGGTTACAACAACAGTCAGCAACATGGCGGAGGTAAGATCCTACTGCTATGAGGCATCAATATCAGACATGGCTTCTGACAGCCGCTGCCCAACACAAGGTGAAGCCTACCTTGACAAGCAATCAGACACTCAATATGTCTGCAAAAGAACGTTAGTGGACAGAGGCTGGGGAAATGGATGTGGACTTTTTGGCAAAGGGAGCCTGGTGACATGCGCTAAGTTTGCATGCTCCAAGAAAATGACCGGGAAGAGCATCCAGCCAGAGAATCTGGAGTACCGGATAATGCTGTCAGTTCATGGCTCCCAGCACAGTGGGATGATCGTTAATGACACAGGACATGAAACTGATGAGAATAGAGCGAAAGTTGAGATAACGCCCAATTCACCGAGAGCCGAAGCCACCCTGGGGGGTTTTGGAAGCCTAGGACTTGATTGTGAACCGAGGACAGGCCTTGACTTTTCAGATTTGTATTACTTGACTATGAATAACAAGCACTGGTTGGTTCACAAGGAGTGGTTCCACGACATTCCATTACCTTGGCACGCTGGGGCAGACACCGGAACTCCACACTGGAACAACAAAGAAGCACTGGTAGAGTTCAAGGACGCACATGCCAAAAGGCAAACTGTCGTGGTTCTAGGGAGTCAAGAAGGAGCAGTTCACACGGCCCTTGCTGGAGCTCTGGAGGCTGAGATGGATGGTGCAAAGGGAAGGCTGTCCTCTGGCCACTTGAAATGTCGCCTGAAAATGGATAAACTTAGATTGAAGGGCGTGTCATACTCCTTGTGTACTGCAGCGTTCACATTCACCAAGATCCCGGCTGAAACACTGCACGGGACAGTCACAGTGGAGGTACAGTACGCAGGGACAGATGGACCTTGCAAGGTTCCAGCTCAGATGGCGGTGGACATGCAAACTCTGACCCCAGTTGGGAGGTTGATAACCGCTAACCCCGTAATCACTGAAAGCACTGAGAACTCTAAGATGATGCTGGAACTTGATCCACCATTTGGGGACTCTTACATTGTCATAGGAGTCGGGGAGAAGAAGATCACCCACCACTGGCACAGGAGTGGCAGCACCATTGGAAAAGCATTTGAAGCCACTGTGAGAGGTGCCAAGAGAATGGCAGTCTTGGGAGACACAGCCTGGGACTTTGGATCAGTTGGAGGCGCTCTCAACTCATTGGGCAAGGGCATCCATCAAATTTTTGGAGCAGCTTTCAAATCATTGTTTGGAGGAATGTCCTGGTTCTCACAAATTCTCATTGGAACGTTGCTGATGTGGTTGGGTCTGAACACAAAGAATGGATCTATTTCCCTTATGTGCTTGGCCTTAGGGGGAGTGTTGATCTTCTTATCCACAGCCGTCTCTGCTGATGTGGGGTGCTCGGTGGACTTCTCAAAGAAGGAGACGAGATGCGGTACAGGGGTGTTCGTCTATAACGACGTTGAAGCCTGGAGGGACAGGTACAAGTACCATCCTGACTCCCCCCGTAGATTGGCAGCAGCAGTCAAGCAAGCCTGGGAAGATGGTATCTGCGGGATCTCCTCTGTTTCAAGAATGGAAAACATCATGTGGAGATCAGTAGAAGGGGAGCTCAACGCAATCCTGGAAGAGAATGGAGTTCAACTGACGGTCGTTGTGGGATCTGTAAAAAACCCCATGTGGAGAGGTCCACAGAGATTGCCCGTGCCTGTGAACGAGCTGCCCCACGGCTGGAAGGCTTGGGGGAAATCGTATTTCGTCAGAGCAGCAAAGACAAATAACAGCTTTGTCGTGGATGGTGACACACTGAAGGAATGCCCACTCAAACATAGAGCATGGAACAGCTTTCTTGTGGAGGATCATGGGTTCGGGGTATTTCACACTAGTGTCTGGCTCAAGGTTAGAGAAGATTATTCATTAGAGTGTGATCCAGCCGTTATTGGAACAGCTGTTAAGGGAAAGGAGGCTGTACACAGTGATCTAGGCTACTGGATTGAGAGTGAGAAGAATGACACATGGAGGCTGAAGAGGGCCCATCTGATCGAGATGAAAACATGTGAATGGCCAAAGTCCCACACATTGTGGACAGATGGAATAGAAGAGAGTGATCTGATCATACCCAAGTCTTTAGCTGGGCCACTCAGCCATCACAATACCAGAGAGGGCTACAGGACCCAAATGAAAGGGCCATGGCACAGTGAAGAGCTTGAAATTCGGTTTGAGGAATGCCCAGGCACTAAGGTCCACGTGGAGGAAACATGTGGAACAAGAGGACCATCTCTGAGATCAACCACTGCAAGCGGAAGGGTGATCGAGGAATGGTGCTGCAGGGAGTGCACAATGCCCCCACTGTCGTTCCGGGCTAAAGATGGCTGTTGGTATGGAATGGAGATAAGGCCCAGGAAAGAACCAGAAAGCAACTTAGTAAGGTCAATGGTGACTGCAGGATCAACTGATCACATGGACCACTTCTCCCTTGGAGTGCTTGTGATCCTGCTCATGGTGCAGGAAGGGCTGAAGAAGAGAATGACCACAAAGATCATCATAAGCACATCAATGGCAGTGCTGGTAGCTATGATCCTGGGAGGATTTTCAATGAGTGACCTGGCTAAGCTTGCAATTTTGATGGGTGCCACCTTCGCGGAAATGAACACTGGAGGAGATGTAGCTCATCTGGCGCTGATAGCGGCATTCAAAGTCAGACCAGCGTTGCTGGTATCTTTCATCTTCAGAGCTAATTGGACACCCCGTGAAAGCATGCTGCTGGCCTTGGCCTCGTGTCTTTTGCAAACTGCGATCTCCGCCTTGGAAGGCGACCTGATGGTTCTCATCAATGGTTTTGCTTTGGCCTGGTTGGCAATACGAGCGATGGTTGTTCCACGCACTGATAACATCACCTTGGCAATCCTGGCTGCTCTGACACCACTGGCCCGGGGCACACTGCTTGTGGCGTGGAGAGCAGGCCTTGCTACTTGCGGGGGGTTTATGCTCCTCTCTCTGAAGGGAAAAGGCAGTGTGAAGAAGAACTTACCATTTGTCATGGCCCTGGGACTAACCGCTGTGAGGCTGGTCGACCCCATCAACGTGGTGGGACTGCTGTTGCTCACAAGGAGTGGGAAGCGGAGCTGGCCCCCTAGCGAAGTACTCACAGCTGTTGGCCTGATATGCGCATTGGCTGGAGGGTTCGCCAAGGCAGATATAGAGATGGCTGGGCCCATGGCCGCGGTCGGTCTGCTAATTGTCAGTTACGTGGTCTCAGGAAAGAGTGTGGACATGTACATTGAAAGAGCAGGTGACATCACATGGGAAAAAGATGCGGAAGTCACTGGAAACAGTCCCCGGCTCGATGTGGCGCTAGATGAGAGTGGTGATTTCTCCCTGGTGGAGGATGACGGTCCCCCCATGAGAGAGATCATACTCAAGGTGGTCCTGATGACCATCTGTGGCATGAACCCAATAGCCATACCCTTTGCAGCTGGAGCGTGGTACGTATACGTGAAGACTGGAAAAAGGAGTGGTGCTCTATGGGATGTGCCTGCTCCCAAGGAAGTAAAAAAGGGGGAGACCACAGATGGAGTGTACAGAGTAATGACTCGTAGACTGCTAGGTTCAACACAAGTTGGAGTGGGAGTTATGCAAGAGGGGGTCTTTCACACTATGTGGCACGTCACAAAAGGATCCGCGCTGAGAAGCGGTGAAGGGAGACTTGATCCATACTGGGGAGATGTCAAGCAGGATCTGGTGTCATACTGTGGTCCATGGAAGCTAGATGCCGCCTGGGATGGGCACAGCGAGGTGCAGCTCTTGGCCGTGCCCCCCGGAGAGAGAGCGAGGAACATCCAGACTCTGCCCGGAATATTTAAGACAAAGGATGGGGACATTGGAGCGGTTGCGCTGGATTACCCAGCAGGAACTTCAGGATCTCCAATCCTAGACAAGTGTGGGAGAGTGATAGGACTTTATGGCAATGGGGTCGTGATCAAAAACGGGAGTTATGTTAGTGCCATCACCCAAGGGAGGAGGGAGGAAGAGACTCCTGTTGAGTGCTTCGAGCCCTCGATGCTGAAGAAGAAGCAGCTAACTGTCTTAGACTTGCATCCTGGAGCTGGGAAAACCAGGAGAGTTCTTCCTGAAATAGTCCGTGAAGCCATAAAAACAAGACTCCGTACTGTGATCTTAGCTCCAACCAGGGTTGTCGCTGCTGAAATGGAGGAGGCCCTTAGAGGGCTTCCAGTGCGTTATATGACAACAGCAGTCAATGTCACCCACTCTGGAACAGAAATCGTCGACTTAATGTGCCATGCCACCTTCACTTCACGTCTACTACAGCCAATCAGAGTCCCCAACTATAATCTGTATATTATGGATGAGGCCCACTTCACAGATCCCTCAAGTATAGCAGCAAGAGGATACATTTCAACAAGGGTTGAGATGGGCGAGGCGGCTGCCATCTTCATGACCGCCACGCCACCAGGAACCCGTGACGCATTTCCGGACTCCAACTCACCAATTATGGACACCGAAGTGGAAGTCCCAGAGAGAGCCTGGAGCTCAGGCTTTGATTGGGTGACGGATCATTCTGGAAAAACAGTTTGGTTTGTTCCAAGCGTGAGGAACGGCAATGAGATCGCAGCTTGTCTGACAAAGGCTGGAAAACGGGTCATACAGCTCAGCAGAAAGACTTTTGAGACAGAGTTCCAGAAAACAAAACATCAAGAGTGGGACTTTGTCGTGACAACTGACATTTCAGAGATGGGCGCCAACTTTAAAGCTGACCGTGTCATAGATTCCAGGAGATGCCTAAAGCCGGTCATACTTGATGGCGAGAGAGTCATTCTGGCTGGACCCATGCCTGTCACACATGCCAGCGCTGCCCAGAGGAGGGGGCGCATAGGCAGGAATCCCAACAAACCTGGAGATGAGTATCTGTATGGAGGTGGGTGCGCAGAGACTGACGAAGACCATGCACACTGGCTTGAAGCAAGAATGCTCCTTGACAATATTTACCTCCAAGATGGCCTCATAGCCTCGCTCTATCGACCTGAGGCCGACAAAGTAGCAGCCATTGAGGGAGAGTTCAAGCTTAGGACGGAGCAAAGGAAGACCTTTGTGGAACTCATGAAAAGAGGAGATCTTCCTGTTTGGCTGGCCTATCAGGTTGCATCTGCCGGAATAACCTACACAGATAGAAGATGGTGCTTTGATGGCACGACCAACAACACCATAATGGAAGACAGTGTGCCGGCAGAGGTGTGGACCAGACACGGAGAGAAAAGAGTGCTCAAACCGAGGTGGATGGACGCCAGAGTTTGTTCAGATCATGCGGCCCTGAAGTCATTCAAGGAGTTTGCCGCTGGGAAAAGAGGAGCGGCTTTTGGAGTGATGGAAGCCCTGGGAACACTGCCAGGACACATGACAGAGAGATTCCAGGAAGCCATTGACAACCTCGCTGTGCTCATGCGGGCAGAGACTGGAAGCAGGCCTTACAAAGCCGCGGCGGCCCAATTGCCGGAGACCCTAGAGACCATAATGCTTTTGGGGTTGCTGGGAACAGTCTCGCTGGGAATCTTCTTCGTCTTGATGAGGAACAAGGGCATAGGGAAGATGGGCTTTGGAATGGTGACTCTTGGGGCCAGCGCATGGCTCATGTGGCTCTCGGAAATTGAGCCAGCCAGAATTGCATGTGTCCTCATTGTTGTGTTCCTATTGCTGGTGGTGCTCATACCTGAGCCAGAAAAGCAAAGATCTCCCCAGGACAACCAAATGGCAATCATCATCATGGTAGCAGTAGGTCTTCTGGGCTTGATTACCGCCAATGAACTCGGATGGTTGGAGAGAACAAAGAGTGACCTAAGCCATCTAATGGGAAGGAGAGAGGAGGGGGCAACCATAGGATTCTCAATGGACATTGACCTGCGGCCAGCCTCAGCTTGGGCCATCTATGCTGCCTTGACAACTTTCATTACCCCAGCCGTCCAACATGCAGTGACCACCTCATACAACAACTACTCCTTAATGGCGATGGCCACGCAAGCTGGAGTGTTGTTTGGCATGGGCAAAGGGATGCCATTCTACGCATGGG-ACTTTGGAGTCCCGCTGCTAATGATAGGTTGCTACTCACAATTAACACCCCTGACCCTAATAGTGGCCATCATTTTGCTCGTGGCGCACTACATGTACTTGATCCCAGGGCTGCAGGCAGCAGCTGCGCGTGCTGCCCAGAAGAGAACGGCAGCTGGCATCATGAAGAACCCTGTTGTGGATGGAATAGTGGTGACTGACATTGACACAATGACAATTGACCCCCAAGTGGAGAAAAAGATGGGACAGGTGCTACTCATAGCAGTAGCCGTCTCCAGCGCCATACTGTCGCGGACCGCCTGGGGGTGGGGGGAGGCTGGGGCTCTGATCACAGCCGCAACTTCCACTTTGTGGGAAGGCTCTCCGAACAAGTACTGGAACTCCTCTACAGCCACTTCACTGTGTAACATTTTTAGGGGAAGTTACTTGGCTGGAGCTTCTCTAATCTACACAGTAACAAGAAACGCTGGCTTGGTCAAGAGACGTGGGGGTGGAACAGGAGAGACCCTGGGAGAGAAATGGAAGGCCCGCTTGAACCAGATGTCGGCCCTGGAGTTCTACTCCTACAAAAAGTCAGGCATCACCGAGGTGTGCAGAGAAGAGGCCCGCCGCGCCCTCAAGGACGGTGTGGCAACGGGAGGCCATGCTGTGTCCCGAGGAAGTGCAAAGCTGAGATGGTTGGTGGAGCGGGGATACCTGCAGCCCTATGGAAAGGTCATTGATCTTGGATGTGGCAGAGGGGGCTGGAGTTACTACGTCGCCACCATCCGCAAAGTTCAAGAAGTGAAAGGATACACAAAAGGAGGCCCTGGTCATGAAGAACCCGTGTTGGTGCAAAGCTATGGGTGGAACATAGTCCGTCTTAAGAGTGGGGTGGACGTCTTTCATATGGCGGCTGAGCCGTGTGACACGTTGCTGTGTGACATAGGTGAGTCATCATCTAGTCCTGAAGTGGAAGAAGCACGGACGCTCAGAGTCCTCTCCATGGTGGGGGATTGGCTTGAAAAAAGACCAGGAGCCTTTTGTATAAAAGTGTTGTGCCCATACACCAGCACTATGATGGAAACCCTGGAGCGACTGCAGCGTAGGTATGGGGGAGGACTGGTCAGAGTGCCACTCTCCCGCAACTCTACACATGAGATGTACTGGGTCTCTGGAGCGAAAAGCAACACCATAAAAAGTGTGTCCACCACGAGCCAGCTCCTCTTGGGGCGCATGGACGGGCCTAGGAGGCCAGTGAAATATGAGGAGGATGTGAATCTCGGCTCTGGCACGCGGGCTGTGGTAAGCTGCGCTGAAGCTCCCAACATGAAGATCATTGGTAACCGCATTGAAAGGATCCGCAGTGAGCACGCGGAAACGTGGTTCTTTGACGAGAACCACCCATATAGGACATGGGCTTACCATGGAAGCTATGAGGCCCCCACACAAGGGTCAGCGTCCTCTCTAATAAACGGGGTTGTCAGGCTCCTGTCAAAACCCTGGGATGTGGTGACTGGAGTCACAGGAATAGCCATGACCGACACCACACCGTATGGTCAGCAAAGAGTTTTCAAGGAAAAAGTGGACACTAGGGTGCCAGACCCCCAAGAAGGCACTCGTCAGGTTATGAGCATGGTCTCTTCCTGGTTGTGGAAAGAGCTAGGCAAACACAAACGGCCACGAGTCTGCACCAAAGAAGAGTTCATCAACAAGGTTCGTAGCAATGCAGCATTAGGGGCAATATTTGAAGAGGAAAAAGAGTGGAAGACTGCAGTGGAAGCTGTGAACGATCCAAGGTTCTGGGCTCTAGTGGACAAGGAAAGAGAGCACCACCTGAGAGGAGAGTGCCAGAGCTGTGTGTACAACATGATGGGAAAAAGAGAAAAGAAACAAGGGGAATTTGGAAAGGCCAAGGGCAGCCGCGCCATCTGGTATATGTGGCTAGGGGCTAGATTTCTAGAGTTCGAAGCCCTTGGATTCTTGAACGAGGATCACTGGATGGGGAGAGAGAACTCAGGAGGTGGTGTTGAAGGGCTGGGATTACAAAGACTCGGATATGTCCTAGAAGAGATGAGTCGTATACCAGGAGGAAGGATGTATGCAGATGACACTGCTGGCTGGGACACCCGCATTAGCAGGTTTGATCTGGAGAATGAAGCTCTAATCACCAACCAAATGGAGAAAGGGCACAGGGCCTTGGCATTGGCCATAATCAAGTACACATACCAAAACAAAGTGGTAAAGGTCCTTAGACCAGCTGAAAAAGGGAAAACAGTTATGGACATTATTTCGAGACAAGACCAAAGGGGGAGCGGACAAGTTGTCACTTACGCTCTTAACACATTTACCAACCTAGTGGTGCAACTCATTCGGAATATGGAGGCTGAGGAAGTTCTAGAGATGCAAGACTTGTGGCTGCTGCGGAGGTCAGAGAAAGTGACCAACTGGTTGCAGAGCAACGGATGGGATAGGCTCAAACGAATGGCAGTCAGTGGAGATGATTGCGTTGTGAAGCCAATTGATGATAGGTTTGCACATGCCCTCAGGTTCTTGAATGATATGGGAAAAGTTAGGAAGGACACACAAGAGTGGAAACCCTCAACTGGATGGGACAACTGGGAAGAAGTTCCGTTTTGCTCCCACCACTTCAACAAGCTCCATCTCAAGGACGGGAGGTCCATTGTGGTTCCCTGCCGCCACCAAGATGAACTGATTGGCCGGGCCCGCGTCTCTCCAGGGGCGGGATGGAGCATCCGGGAGACTGCTTGCCTAGCAAAATCATATGCGCAAATGTGGCAGCTCCTTTATTTCCACAGAAGGGACCTCCGACTGATGGCCAATGCCATTTGTTCATCTGTGCCAGTTGACTGGGTTCCAACTGGGAGAACTACCTGGTCAATCCATGGAAAGGGAGAATGGATGACCACTGAAGACATGCTTGTGGTGTGGAACAGAGTGTGGATTGAGGAGAACGACCACATGGAAGACAAGACCCCAGTTACGAAATGGACAGACATTCCCTATTTGGGAAAAAGGGAAGACTTGTGGTGTGGATCTCTCATAGGGCACAGACCGCGCACCACCTGGGCTGAGAACATTAAAAACACAGTCAACATGGTGCGCAGGATCATAGGTGATGAAGAAAAGTACATGGACTACCTATCCACCCAAGTTCGCTACTTGGGTGAAGAAGGGTCTACACCTGGAGTGCTGTAA

>ENA|AHL43500|AHL43500.1 Zika virus polyprotein

ATGAAAAACCC-AAAGAAGAAATCCGGAGGATTCCGGATTGTCAATATGCTAAAACGCGGAGTAGCCCGTGTAAACCCCTTGGGAGGTTTGAAGAGGCTGCCAGCCGGACTTCTGCTGGGTCATGGACCCATTAGAATGGTTTTGGCGATACTAGCCTTTTTGAGATTTACAGCAATCAAGCCATCACTGGGCCTCATCAATAGATGGGGTTCAGTGGGGAAAAAAGAGGCTATGGAGATAATAAAGAAGTTTAAGAAAGATCTTGCTGCCATGTTGAGAATAATCAATGCTAGGAAGGAGAGGAAGAGACGTGGTGCAGACACCAGTATCGGAATCATTGGCCTCCTGCTGACTACAGCCATGGCAGCAGAGATTACTAGACGAGGAAGTGCATACTACATGTACTTGGACAGGAGCGATGCTGGGAAGGCCATTTCCTTTGCTACCACATTGGGAGTGAACAAATGCCATGTACAGATCATGGATCTCGGGCACATGTGTGACGCCACCATGAGTTATGAGTGCCCTATGCTAGACGAAGGAGTGGAACCAGATGATGTCGATTGCTGGTGCAACACGACATCAACTTGGGTTGTGTACGGAACCTGTCATCACAAAAAAGGTGAAGCACGGCGATCTAGAAGAGCTGTGACGCTCCCTTCTCACTCCACAAGGAAGTTGCAAACGCGGTCGCAGACCTGGTTAGAATCAAGAGAATACACGAAGCACTTGATCAAGGTTGAAAACTGGATATTCAGGAACCCCGGGTTTGCGCTAGCGGCTGTTGCCATTGCTTGGCTTCTGGGAAGTTCGACGAGCCAAAAAGTCATATACTTGATCATGATACTGCTGATTGCCCCGGCATACAGCATCAGGTGCATTGGAGTCAGCAACAGAGACTTCGTGGAAGGCATGTCAGGTGGGACCTGGGTTGATGTCGTCTTGGAACATGGAGGTTGCGTCACCGTAATGGCACAGGACAAGCCAACAGTTGACATAGAGTTGGTCACGACAACGGTTAGCAACATGGCTGAGGTAAGATCCTACTGCTACGAGGCATCAATATCGGACATGGCTTCGGATAGTCGCTGCCCAACACAAGGTGAAGCCTACCTTGACAAGCAATCAGACACTCAATATGTTTGCAAAAGAACATTGGTGGACAGAGGTTGGGGAAATGGGTGTGGACTTTTTGGCAAAGGGAGCCTGGTGACATGTGCCAAGTTCACGTGTTCCAAGAAGATGACCGGGAAGAGCATCCAGCCGGAGAATCTGGAGTATCGAATAATGCTATCGGTGCATGGCTCCCAGCACAGCGGGATGATTGTTAATGACNNNNNNNNNNNNNNNNNNNNNAACAGAGCGGAAGTCGAGGTCACGCCTAATTCACCAAGAGCAGAAGCAACCTTGGGAGGCTTTGGAAGTTTAGGACTTGATTGTGAACCAAGGACAGGCCTTGACTTTTCAGATTTGTATTACCTGACCATGAACAATAAGCACTGGTTAGTCCACAAAGAGTGGTTTCATGACATCCCATTGCCTTGGCATGCTGGGGCAGACACCGGAACTCCACACTGGAACAACAAAGAGGCATTGGTAGAGTTCAAGGATGCCCACGCCAAGAGGCAAACCGTCGTCGTTCTGGGGAGCCAGGAAGGAGCTGTTCACACGGCTCTTGCTGGAGCTTTGGAGGCTGAGATGGATGGTGCAAAAGGAAGGCTGTTCTCTGGCCATTTGAAATGCCGCCTAAAAATGGACAAGCTTAGATTGAAGGGCGTGTCATATTCCTTGTGCACCGCAGCGTTCACATTCACCAAGGTCCCAGCTGAAACACTGCATGGAACAGTCACAGTGGAGGTGCAGTATGCAGGGACAGATGGACCCTGCAAGGTCCCAGCCCAGATGGCGGTGGACATGCAGACTTTGACCCCAGTCGGAAGGCTGATAACTGCCAACCCCGTGATTACTGAAAGCACTGAAAATTCAAAGATGATGTTGGAGCTCGACCCACCATTTGGGGATTCTTACATTGTCATAGGAGTTGGAGATAAGAAAATCACCCACCACTGGCATAGGAGTGGTAGCACCATCGGAAAAGCATTTGAGGCCACTGTGAGAGGTGCCAAGAGAATGGCAGTTTTGGGGGACACAGCTTGGGACTTTGGATCCGTCGGGGGTGTGTTCAACTCATTGGGCAAGGGCATCCACCAGATCTTTGGAGCAGCCTTCAAATCACTGTTTGGAGGAATGTCCTGGTTCTCACAGATCCTCATAGGCACGCTGCTAGTGTGGTTGGGCTTGAACACAAAGAATGGATCCATCTCCCTCACATGCTTGGCCCTGGGGGGAGTGATGATCTTCCTCTCCACGGCTGTTTCTGCTGATGTGGGGTGCTCGGTGGACTTCTCAAAAAGAGAAACGAGATGTGGCACGGGGGTGTTCGTCTACAATGACGTTGAAGCCTGGAGGGATAGGTACAAGTACCATCCTGACTCCCCCCGCAGATTGGCAGCAGCAGTCAAGCAGGCTTGGGAAGAGGGGATTTGTGGGATCTCCTCCGTTTCAAGAATGGAAAACATCATGTGGAAATCAGTGGAAGGGGAGCTCAATGCTATCTTAGAGGAGAATGGAGTTCAACTGACAGTTGTTGTGGGATCTGTAAAAAACCCCATGTGGAGAGGTCCACAAAGATTGCCAGTGCCTGTAAATGAGCTGCCCCATGGCTGGAAAGCCTGGGGGAAATCATATTTTGTCAGAGCGGCAAAGACCAACAACAGTTTTGTTGTCGACGGTGACACACTGAAGGAATGTCCGCTCAAACACAGAGCATGGAACAGTTTCCTTGTGGAGGATCACGGGTTTGGGGTCTTCCACACTAGTGTTTGGCTTAAGGTTAGAGAAGATTACTCACTAGAGTGTGACCCAGCCGTCATAGGAACAGCTGTTAAGGGAAAGGAGGCCGCGCACAGTGATCTAGGCTACTGGATTGAAAGTGAAAAGAATGACACATGGAGGTTGAAGAGGGCCCACCTGATTGAGATGAAAACATGTGAGTGGCCAAAGTCTCACACACTGTGGACAGATGGAGTAGAAGAAAGTGATCTGATCATACCCAAGTCTTTAGCTGGTCCACTCAGCCACCACAACACCAGAGAGGGTTACAGAACTCAGGCAAAAGGGCCATGGCATAGTGAAGAGCTTGAAATCCGGTTTGAGGAATGTCCAGGCACCAAGGTTCACGTGGAGGAGACATGCGGAACTAGAGGACCATCTCTGAGATCAACCACTGCAAGTGGAAGGGTCATTGAGGAATGGTGCTGTAGAGAATGCACAATGCCCCCACTATCATTTCGAGCGAAAGACGGCTGCTGGTATGGAATGGAGATAAGGCCCAGGAAAGAACCAGAGAGCAATTTAGTGAGGTCAATGGTGACAGCGGGATCAACCGATCACATGGACCACTTCTCTCTTGGAGTGCTTGTGATTCTACTCATGGTGCAGGAAGGGCTGAAGAAGAGAATGACCACGAAGATCATTGTGAGCACATCAATGGCAGTGCTGGTAGCCATGATCTTGGGAGGATTCTCAATGAGTGACCTGGCTAAGCTTGTGATCCTGATGGGTGCTACTTTCGCAGAAATGAATACTGGAGGAGATGTGGCTCACTTGGCATTGGTCGCGGCATTCAAAGTCAGACCAGCCTTGCTGGTCTCCTTCATTTTCAGAGCCAATTGGACACCCCGTGAGAGCATGCTGCTAGCCCTGGCTTCGTGTCTTTTGCAAACTGCAATTTCAGCTCTTGAAGGTGACTTGATGGTCCTCGTTAATGGATTTGCTTTGGCCTGGTTGGCAATACGAGCAATGGCCGTGCCACGCACTGACAACATTGCTCTAGCAATCCTGGCTGCTCTGACACCACTAGCCCGAGGCACACTGCTCGTGGCATGGAGAGCGGGCCTCGCCACTTGTGGAGGGTTCATGCTCCTCTCCTTGAAAGGAAAAGGTAGTGTGAAGAAGAACCTGCCATTTGTCATGGCCTTGGGAATGACCGCTGTGAGGATAGTGGACCCCATTAACGTGGTGGGACTACTACTACTCACAAGGAGTGGGAAGCGGAGCTGGCCCCCCAGTGAAGTGCTTACAGCTGTTGGCCTGATATGTGCACTGGCCGGAGGGTTTGCCAAGGCAGACATTGAGATGGCTGGGCCCATGGCTGCAGTAGGCTTGCTAATTGTCAGTTATGTGGTCTCAGGAAAGAGTGTGGATATGTACATTGAAAGAGCAGGTGACATCACATGGGAAAAAGACGCGGAAGTTACTGGAAATAGTCCTCGGCTTGACGTGGCACTAGATGAGAGTGGTGATTTCTCTCTGGTAGAGGAAGATGGTCCACCCATGAGAGAGATCATACTCAAGGTGGTCCTGATGGCCATCTGTGGCATGAACCCAATAGCCATACCTTTTGCTGCAGGAGCGTGGTATGTGTATGTGAAGACTGGGAAAAGGAGTGGTGCCCTCTGGGATGTGCCAGCTCCTAAAGAAGTGAAAAAAGGAGAGACCACAGATGGAGTATACAGAGTGATGACCCGCAGATTGCTGGGTTCAACACAGGTTGGAGTGGGAGTCATGCAGGAGGGAGTCTTCCACACCATGTGGCACGTCACAAAAGGAGCCGCACTGAGGAGCGGTGAAGGGAGACTTGATCCATACTGGGGTGATGTCAAGCAGGACTTGGTGTCATACTGTGGGCCTTGGAAGCTGGATGCAGCTTGGGATGGACTTAGTGAGGTGCAGCTCTTGGCAGTACCTCCCGGAGAGAGGGCCAGAAACATCCAGACTCTGCCTGGAATATTCAAGACAAAGGATGGGGACATCGGAGCAGTTGCTCTGGACTACCCCGCAGGAACCTCAGGATCTCCGATCCTAGACAAATGCGGAAGAGTGATAGGACTTTATGGCAATGGGGTTGTGATCAAGAATGGAAGCTATGTTAGTGCCATAACCCAGGGAAAGAGGGAGGAGGAGACTCCGGTTGAATGTTTCGAACCCTCGATGCTGAAGAAGAAGCAGCTAACTGTCCTGGACCTGCATCCAGGAGCCGGAAAGACTAGGAGAGTTCTTCCTGAAATAGTCCGTGAAGCCATAAAAAAGAGACTCCGCACAGTGATCTTGGCTCCAACCAGGGTTGTCGCTGCTGAGATGGAGGAAGCCTTGAGAGGACTTCNNGTNCGTTACATGACAACAGCAGTCAACGTCACCCATTCTGGGACAGAAATCGTTGATTTGATGTGCCATGCCACTTTCACTTCACGCTTACTACAACCCATCAGAGTCCCTAATTACAATCTCAACATCATGGATGAAGCCCACTTCACAGACCCCTCAAGTATAGCTGCAAGAGGATACATATCAACAAGGGTTGAAATGGGCGAGGCGGCTGCCATTTTTATGACTGCCACACCACCAGGAACCCGTGATGCGTTTCCTGACTCTAACTCACCAATCATGGACACAGAAGTGGAAGTCCCAGAGAGAGCCTGGAGCTCAGGCTTTGATTGGGTGACAGACCATTCTGGGAAAACAGTTTGGTTCGTTCCAAGTGTGAGAAATGGAAATGAAATCACAGCCTGCCTGACAAAGGCTGGAAAGCGGGTCATACAGCTCAGCAGGAAGACTTTTGAGACAGAGTTCCAGAAAACAAAAAACCAAGAGTGGGACTTTGTCATAACAACTGACATCTCAGAGATGGGCGCCAATTTCAAGGCTGACCGGGTCATAGACTCTAGGAGATGCCTAAAGCCAGTCATACTTGATGGTGAGAGAGTCATCTTGGCTGGGCCCATGCCTGTCACGCATGCTAGTGCTGCTCAGAGGAGAGGACGTATAGGCAGGAACCCTAACAAACCTGGAGATGAGTACATGTATGGAGGTGGGTGTGCAGAGACTGATGAAGACCACGCACACTGGCTTGAAGCAAGAATGCTTCTTGACAACATCTACCTCCAGGATGGCCTCATAGCCTCGCTCTATCGGCCTGAGGCCGATAAGGTGGCCGCCATTGAGGGAGAGTTTAAGCTGAGGACAGAGCAAAGGAAGACCTTTGTGGAACTTATGAAGAGAGGAGACCTTCCCGTTTGGCTAGCCTATCAGGTTGCATCTGCCGGAATAACTTACACAGACAGAAGATGGTGCTTTGATGGCACAACTAACAACACCATAATGGAAGACAGTGTACCAGCAGAGGTGTGGACAAAATATGGAGAGAAGAGAGTGCTCAAACCAAGATGGATGGATGCTAGAGTCTGTTCAGATCATGCCGCCCTGAAGTCGTTCAAAGAGTTTGCTGCCGGGAAAAGAGGAGCGGCTTTGGGAGTAATGGAGGCCCTGGGAACATTGCCAGGACACATGACAGAGAGGTTTCAGGAAGCCATTGACAATCTCGCTGTGCTTATGCGAGCAGAAACTGGAAGCAGGCCTTACAAGGCAGCGGCAGCCCCACTGCCGGAGACCCTAGAGACCATCATGCTCTTAGGTTTGCTGGGAACAGTTTCGTTGGGGATATTCTTTGTCTTGATGCGGAACAAGGGCATCGGGAAGATGGGGTTTGGAATGGTAACTCTTGGGGCCAGTGCATGGCTCATGTGGCTTTCGGAAATTGAACCAGCCAGAATTGCATGTGTCCTCATTGTTGTGTTTTTGTTACTGGTGGTACTCATACCTGAGCCAGAGAAGCAAAGATCTCCCCAGGATAACCAGATGGCAATCATCATCATGGTGGCAGTGGGCCTTCTGGGGTTGATAACTGCAAATGAACTTGGATGGCTGGAAAGAACAAAAAATGACATAGCTCATCTAATGGGAAGGAGAGAAGAAGGAGTAACTATGGGATTCTCAATGGACATCGATTTGAGGCCGGCCTCCGCCTGGGCTATCTATGCTGCATTGACAACTCTCATCACCCCAGCCGTCCAACATGCGGTAACCACTTCATATAACAACTACTCCTTAATGGCGATGGCCACACAAGCTGGAGTGCTGTTTGGCATGGGCAAAGGGATGCCATTTTATGCATGGG-ACTTTGGAGTCCCGCTGCTAATGATGGGTTGCTACTCACAATTAACACCCCTGACCCTGATAGTGGCTATCATTCTGCTTGTGGCACACTACATGTACTTGATCCCAGGCCTACAGGCAGCAGCAGCGCGTGCTGCCCAGAAGAGAACAGCAGCTGGCATCATGAAGAATCCCGTTGTGGATGGAATAGTGGTAACTGACATTGACACAATGACAATTGACCCCCAAGTGGAGAAGAAGATGGGACAAGTGTTACTCATAGCAGTAGCCATCTCCAGTGCTGTGCTGCTGCGGACTGCTTGGGGATGGGGGGAAGCTGGAGCTCTGATCACAGCAGCAACCTCCACTTTGTGGGAAGGTTCTCCAAACAAATACTGGAACTCCTCTACAGCCACCTCACTGTGCAACATCTTTAGAGGAAGTTACCTGGCAGGCGCTTCTCTTATTTACACAGTGACAAGAAATGCTGGCCTGGTCAAGAGACGTGGAGGTGGAACGGGAGAGACTCTAGGAGAGAAGTGGAAAGCTCGCCTGAATCAGATGTCGGCCCTGGAGTTCTACTCTTACAAAAAGTCGGGCATCACTGAAGTGTGTAGAGAGGAGGCTCGCCGCGCCCTCAAGGATGGAGTGGCCACAGGAGGACATGCTGTATCCCGGGGAAGCGCAAAGCTCAGATGGCTGGTGGAGAGAGGATACCTGCAGCCCCATGGAAAGGTTGTTGACCTCGGATGTGGCAGAGGGGGTTGGAGCTATTATGCCGCTACCATCCGGAAAGTGCAGGAGGTGAAAGGATACACAAAGGGAGGTCCCGGTCATGAAGAACCCATGCTGGTGCAAAGCTATGGATGGAACATAGTTCGTCTCAAGAGTGGAGTGGACGTCTTCCACATGGCGGCTGAGCCGTGTGATACTCTGCTGTGTGACATAGGTGAGTCATCATCCAGTCCTGAAGTAGAAGAGACGCGAACACTCAGAGTGCTCTCTATGGTGGGGGACTGGCTTGAAAAAAGACCAGGGGCCTTCTGCATAAAGGTGCTGTGCCCATACACCAGCACTATGATGGAGACCATGGAGCGACTGCAACGTAGGTATGGGGGAGGATTAGTCAGAGTGCCATTGTCTCGCAACTCCACACATGAGATGTATTGGGTCTCTGGAGCAAAAAGCAACATCATAAAAAGTGTGTCCACCACAAGCCAGCTCCTCCTGGGACGCATGGATGGGCCCAGGAGGCCAGTGAAATATGAGGAGGATGTGAACCTCGGCTCGGGCACACGAGCTGTGGCAAGCTGTGCTGAGGCTCCCAACATGAAGATCATTGGTAAGCGCATTGAGAGAATCCGCAGTGAACATGCAGAAACATGGTTCTTTGATGAAAACCACCCATACAGGACATGGGCTTACCATGGGAGCTACGAAGCCCCCACGCAAGGGTCAGCGTCTTCCCTCGTGAATGGGGTTGTTAGGCTCCTGTCAAAGCCCTGGGATGTAGTGACTGGAGTTACAGGAATAGCTATGACTGACACCACACCATATGGCCAACAAAGAGTCTTCAAAGAAAAAGTGGACACCAGGGTGCCAGATCCCCAAGAAGGCACTCGCCAGACAATGAACATGGTCTCGTCCTGGCTGTGGAAGGAGCTGGGAAAACGCAAGCGGCCACGCGTCTGCACCAAAGAAGAGTTCATCAACAAGGTGCGCAGCAATGCAGCGCTGGGAGCAATATTTGAAGAGGAAAAAGAATGGAAGACGGCCGTGGAAGCTGTGAATGATCCAAGGTTTTGGGCCCTAGTGGATAAGGAGAGAGAGCATCACCTAAGAGGAGAGTGTCATAGCTGTGTGTACAACATGATGGGAAAAAGAGAGAAGAAGNAAGGAGAATTCGGGAAAGCAAAAGGCAGCCGCGCCATCTGGTACATGTGGTTGGGAGCCAGGTTTCTGGAGTTTGAATCACTGGGGTTTCTGAATGAAGATCACTGGATGGGAAGAGAGAACTCAGGAGGTGGTGTCGAAGGGCTAGGACTGCAAAGACTTGGATATATTCTAGAAGAAATGAATCGGGCACCAGGAGGAAAGATGTATGCAGATGACACCGCTGGTTGGGATACCCGCATTAGTAAGTTTGATCTGGAGAATGAAGCCTTGATTACCAACCAAATGGAGGAAGGGCACAGAGCTCTGGCGTTGGCTGTGATTAAATACACATACCAAAACAAAGTGGTGAAGGTCCTTAGACCAGCTGAAGGAGGAAAAACAGTTATGGACATCATTTCAAGACAAGACCAGAGAGGGAGTGGGCAAGTTGTCACTTATGCTCTCAACACGTTTACCAACTTAGTGGTGCAGCTTATTCGGAACATGGAGGCTGAGGAAGTGTTAGAGATGCAAGACTTATGGCTGTTGAGGAAGCCAGAGAAAGTGACTAGATGGTTGCAGAGTAATGGATGGGACAGACTCAAACGAATGGCAGTCAGTGGAGATGACTGTGTTGTGAAGCCAATTGATGATAGGTTTGCATGCTGCCTCAGGTTCCTAAACGACATGGGGAAAGTTAGGAAAGACACACAGGAATGGAAACCCTCGACTGGATGGAGCAATTGGGAAGAAGTCCCTTTCTGCTCCCACCACTTCAACAAGCTGCACCTCAAGGATGGTAGATCCATTGTGGTCCCTTGCCGCCACCAAGATGAACTGATTGGCCGAGCCCGTGTATCACCAGGGGCAGGATGGAGCATCCGGGAGACTGCCTGTCTTGCAAAATCATATGCACAGATGTGGCAGCTTCTTTACTTCCACAGAAGAGACCTTCGACTGATGGCCAACGCCATTTGTTCGGCTGTGCCAGTCGACTGGGTACCAACCGGGAGAACCACCTGGTCAATCCACGGGAAGGGAGAATGGATGACCACTGAGGACATGCTCATGGTGTGGAATAGAGTGTGGATTGAGGAGAACGACCATATGGAGGACAAGACTCCTGTAACAAAGTGGACTGACATTCCCTACCTGGGAAAAAGGGAGGACTTATGGTGTGGATCCCTTATAGGGCACAGACCCCGCACCACTTGGGCTGAGAACATCAAAGACACAGTCAACATGGTGCGCAGGATCATAGGTGATGAAGAAAAATACATGGACTACCTATCCACCCAAGTCCGCTACTTGGGTGAGGAAGGGTCCACACCTGGAGTGTTGTAA

>ENA|ALX35659|ALX35659.1 Zika virus polyprotein

ATGAAAAACCC-AAAAAAGAAATCCGGAGGATTCCGGATTGTCAATATGCTAAAACGCGGAGTAGCCCGTGTGAGCCCCTTTGGGGGCTTGAAGAGGCTGCCAGCCGGACTTCTGCTGGGTCATGGGCCCATCAGGATGGTCTTGGCGATTCTAGCCTTTTTGAGATTCACGGCAATCAAGCCATCACTGGGTCTCATCAATAGATGGGGTTCAGTGGGGAAAAAAGAGGCTATGGAAATAATAAAGAAGTTCAAGAAAGATCTGGCTGCCATGCTGAGAATAATCAATGCTAGGAAGGAGAAGAAGAGACGAGGCGCAGATACTAGTGTCGGAATTGTTGGCCTCCTGCTGACCACAGCTATGGCAGCGGAGGTCACTAGACGTGGGAGTGCATACTATATGTACTTGGACAGAAACGATGCTGGGGAGGCCATATCTTTTCCAACCACATTGGGGATGAATAAGTGTTATATACAGATCATGGATCTTGGACACACGTGTGATGCCACCATGAGCTATGAATGCCCTATGCTGGATGAGGGGGTGGAACCAGATGACGTCGATTGTTGGTGCAACACGACGTCAACTTGGGTTGTGTACGGAACCTGCCATCACAAAAAAGGTGAAGCACGGAGATCTAGAAGAGCTGTGACGCTCCCCTCCCATTCCACTAGGAAGCTGCAAACGCGGTCGCAAACCTGGTTGGAATCAAGAGAATACACAAAGCACTTGATTAGAGTCGAAAATTGGATATTCAGGAACCCTGGCTTCGCGTTAGCAGCAGCTGCCATCGCTTGGCTTTTGGGAAGCTCAACGAGCCAAAAAGTCATATACTTGGTCATGATACTGCTGATTGCCCCGGCATACAGCATCAGGTGCATAGGAGTCAGCAATAGGGACTTTGTGGAAGGTATGTCAGGTGGGACTTGGGTTGATGTTGTCTTGGAACATGGAGGTTGTGTCACTGTAATGGCACAGGACAAACCGACTGTCGACATAGAGCTGGTTACAACAACAGTCAGCAACATGGCGGAGGTAAGATCCTACTGCTATGAGGCATCAATATCAGACATGGCTTCGGACAGCCGCTGCCCAACACAAGGTGAAGCCTACCTTGACAAGCAATCAGACACTCAATATGTCTGCAAAAGAACGTTAGTGGACAGAGGCTGGGGAAATGGATGTGGACTTTTTGGCAAAGGGAGCCTGGTGACATGCGCTAAGTTTGCATGCTCCAAGAAAATGACCGGGAAGAGCATCCAGCCAGAGAATCTGGAGTACCGGATAATGCTGTCAGTTCATGGCTCCCAGCACAGTGGGATGATCGTTAATGACACAGGACATGAAACTGATGAGAATAGAGCGAAAGTTGAGATAACGCCCAATTCACCAAGAGCCGAAGCCACCCTGGGGGGGTTTGGAAGCCTAGGACTTGATTGTGAACCGAGGACAGGCCTTGACTTTTCAGATTTGTATTACTTGACTATGAATAACAAGCACTGGCTGGTTCACAAGGAGTGGTTCCACGACATTCCATTACCTTGGCACGCTGGGGCAGACACCGGAACTCCACACTGGAACAACAAAGAAGCACTGGTAGAGTTCAAGGACGCACATGCCAAAAGGCAAACTGTCGTGGTTCTAGGGAGTCAAGAAGGAGCAGTTCACACGGCCCTTGCTGGAGCTCTGGAGGCTGAGATGGATGGTGCAAAGGGAAGGCTGTCCTCTGGCCACTTGAAATGTCGCCTGAAAATGGATAAACTTAGATTGAAGGGCGTGTCATACTCCTTGTGTACTGCAGCGTTCACATTCACCAAGATCCCGGCTGAAACACTGCACGGGACAGTCACAGTGGAGGTACAGTACGCAGGGACAGATGGACCTTGCAAGGTTCCAGCTCAGATGGCGGTGGACATGCAAACTCTGACCCCAGTTGGGAGGTTGATAACCGCTAACCCCGTAATCACTGAAAGCACTGAGAACTCTAAGATGATGCTGGAACTTGATCCACCATTTGGGGACTCTTACATTGTCATAGGAGTCGGGGAGAAGAAGATCACCCACCACTGGCACAGGAGTGGCAGCACCATTGGAAAAGCATTTGAAGCCACTGTGAGAGGTGCCAAGAGAATGGCAGTCTTGGGAGACACAGCCTGGGACTTTGGATCAGTTGGAGGCGCTCTCAACTCATTGGGCAAGGGCATCCATCAAATCTTTGGAGCAGCTTTCAAATCATTGTTTGGAGGAATGTCCTGGTTCTCACAAATTCTCATTGGAACGTTGCTGATGTGGTTGGGTCTGAACGCAAAGAATGGATCTATTTCCCTTATGTGCTTGGCCTTAGGGGGAGTGTTGATCTTCTTATCCACAGCCGTCTCTGCTGATGTGGGGTGCTCGGTGGACTTCTCAAAGAAGGAGACGAGATGCGGTACAGGGGTGTTCGTCTATAACGACGTTGAAGCCTGGAGGGACAGGTACAAGTACCATCCTGACTCCCCCCGTAGATTGGCAGCAGCAGTCAAGCAAGCCTGGGAAGATGGTATCTGCGGGATCTCCTCTGTTTCAAGAATGGAAAACATCATGTGGAGATCAGTAGAAGGGGAGCTCAACGCAATCCTGGAAGAGAATGGAGTTCAACTGACGGTCGTTGTGGGATCTGTAAAAAACCCCATGTGGAGAGGTCCACAGAGATTGCCCGTGCCTGTGAACGAGCTGCCCCACGGCTGGAAGGCTTGGGGGAAATCGTACTTCGTCAGAGCAGCAAAGACAAATAACAGCTTTGTCGTGGATGGTGACACACTGAAGGAATGCCCACTCAAACATAGAGCATGGAACAGCTTTCTTGTGGAGGATCATGGGTTCGGGGTATTTCACACTAGTGTCTGGCTCAAGGTTAGAGAAGATTATTCATTAGAGTGTGATCCAGCCGTTATTGGAACAGCTGTTAAGGGAAAGGAGGCTGTACACAGTGATCTAGGCTACTGGATTGAGAGTGAGAAGAATGACACATGGAGGCTGAAGAGGGCCCATCTGATCGAGATGAAAACATGTGAATGGCCAAAGTCCCACACATTGTGGACAGATGGAATAGAAGAGAGTGATCTGATCATACCCAAGTCTTTAGCTGGGCCACTCAGCCATCACAATACCAGAGAGGGCTACAGGACCCAAATGAAAGGGCCATGGCACAGTGAAGAGCTTGAAATTCGGTTTGAGGAATGCCCAGGCACTAAGGTCCACGTGGAGGAAACATGTGGAACGAGAGGACCATCTCTGAGATCAACCACTGCAAGCGGAAGGGTGATCGAGGAATGGTGCTGCAGGGAGTGCACAATGCCCCCACTGTCGTTCCGGGCTAAAGATGGCTGTTGGTATGGAATGGAGATAAGGCCCAGGAAAGAACCAGAAAGCAACTTAGTAAGGTCAATGGTGACTGCAGGATCAACTGATCACATGGACCACTTCTCCCTTGGAGTGCTTGTGATTCTGCTCATGGTGCAGGAAGGGTTGAAGAAGAGAATGACCACAAAGATCATCATAAGCACATCAATGGCAGTGCTGGTAGCTATGATCCTGGGAGGATTTTCAATGAGTGACCTGGCTAAGCTTGCAATTTTGATGGGTGCCACCTTCGCGGAAATGAACACTGGAGGAGATGTAGCTCATCTGGCGCTGATAGCGGCATTCAAAGTCAGACCAGCGTTGCTGGTATCTTTCATCTTCAGAGCTAATTGGACACCCCGTGAAAGCATGCTGCTGGCCTTGGCCTCGTGTCTTTTGCAAACTGCGATCTCCGCCTTGGAAGGCGACCTGATGGTTCTCATCAATGGTTTTGCTTTGGCCTGGTTGGCAATACGAGCGATGGTTGTTCCACGCACTGATAACATCACCTTGGCAATCCTGGCTGCTCTGACACCACTGGCCCGGGGCACACTGCTTGTGGCGTGGAGAGCAGGCCTTGCTACTTGCGGGGGGTTTATGCTCCTCTCTCTGAAGGGAAAAGGCAGTGTGAAGAAGAACTTACCATTTGTCATGGCCCTGGGACTAACCGCTGTGAGGCTGGTCGACCCCATCAACGTGGTGGGACTGCTGTTGCTCACAAGGAGTGGGAAGCGGAGCTGGCCCCCTAGCGAAGTACTCACAGCTGTTGGCCTGATATGCGCATTGGCTGGAGGGTTCGCCAAGGCAGATATAGAGATGGCTGGGCCCATGGCCGCGGTCGGTCTGCTAATTGTCAGTTACGTGGTCTCAGGAAAGAGTGTGGACATGTACATTGAAAGAGCAGGTGACATCACATGGGAAAAAGATGCGGAAGTCACTGGAAACAGTCCCCGGCTCGATGTGGCGCTAGATGAGAGTGGTGATTTCTCCCTGGTGGAGGATGACGGTCCCCCCATGAGAGAGATCATACTCAAGGTGGTCCTGATGACCATCTGTGGCATGAACCCAATAGCCATACCCTTTGCAGCTGGAGCGTGGTACGTATACGTGAAGACTGGAAAAAGGAGTGGTGCTCTATGGGATGTGCCTGCTCCCAAGGAAGTAAAAAAGGGGGAGACCACAGATGGAGTGTACAGAGTAATGACTCGTAGACTGCTAGGTTCAACACAAGTTGGAGTGGGAGTTATGCAAGAGGGGGTCTTTCACACTATGTGGCACGTCACAAAAGGATCCGCGCTGAGAAGCGGTGAAGGGAGACTTGATCCATACTGGGGAGATGTCAAGCAGGATCTGGTGTCATACTGTGGTCCATGGAAGCTAGATGCCGCCTGGGACGGGCACAGCGAGGTGCAGCTCTTGGCCGTGCCCCCCGGAGAGAGAGCGAGGAACATCCAGACTCTGCCCGGAATATTTAAGACAAAGGATGGGGACATTGGAGCGGTTGCGCTGGATTACCCAGCAGGAACTTCAGGATCTCCTATCCTAGACAAGTGTGGGAGAGTGATAGGACTTTATGGCAATGGGGTCGTGATCAAAAATGGGAGTTATGTTAGTGCCATCACCCAAGGGAGGAGGGAGGAAGAGACTCCTGTTGAGTGCTTCGAGCCTTCGATGCTGAAGAAGAAGCAGCTAACTGTCTTAGACTTGCATCCTGGAGCTGGGAAAACCAGGAGAGTTCTTCCTGAAATAGTCCGTGAAGCCATAAAAACAAGACTCCGTACTGTGATCTTAGCTCCAACCAGGGTTGTCGCTGCTGAAATGGAGGAGGCCCTTAGAGGGCTTCCAGTGCGTTATATGACAACAGCAGTCAATGTCACCCACTCTGGAACAGAAATCGTCGACTTAATGTGCCATGCCACCTTCACTTCGCGTCTACTACAGCCAATCAGAGTCCCCAACTATAATCTGTATATTATGGATGAGGCCCACTTCACAGATCCCTCAAGTATAGCAGCAAGAGGATACATTTCAACAAGGGTTGAGATGGGCGAGGCGGCCGCCATCTTCATGACCGCCACGCCACCAGGAACCCGTGACGCATTTCCGGACTCCAACTCACCAATTATGGACACCGAAGTGGAAGTCCCAGAGAGAGCCTGGAGCTCAGGCTTTGATTGGGTGACGGATCATTCTGGAAAAACAGTTTGGTTTGTTCCAAGCGTGAGGAACGGCAATGAGATCGCAGCTTGTCTGACAAAGGCTGGAAAACGGGTCATACAGCTCAGCAGAAAGACTTTTGAGACAGAGTTCCAGAAAACAAAACATCAAGAGTGGGACTTTGTCGTGACAACTGACATTTCAGAGATGGGCGCCAACTTTAAAGCTGACCGTGTCATAGATTCCAGGAGATGCCTAAAGCCGGTCATACTTGATGGCGAGAGAGTCATTCTGGCTGGACCCATGCCTGTCACACATGCCAGCGCTGCCCAGAGGAGGGGGCGCATAGGCAGGAATCCCAACAAACCTGGAGATGAGTATCTGTATGGAGGTGGGTGCGCAGAGACTGACGAAGACCATGCACACTGGCTTGAAGCAAGAATGCTCCTTGACAATATTTACCTCCAAGATGGCCTCATAGCCTCGCTCTATCGACCTGAGGCCGACAAAGTAGCAGCCATTGAGGGAGAGTTCAAGCTTAGGACGGAGCAAAGGAAGACCTTTGTGGAACTCATGAAAAGAGGAGATCTTCCTGTTTGGCTGGCCTATCAGGTTGCATCTGCCGGAATAACCTACACAGATAGAAGATGGTGCTTTGATGGCACGACCAACAACACCATAATGGAAGACAGTGTGCCGGCAGAAGTGTGGACCAGACACGGAGAGAAAAGAGTGCTCAAACCGAGGTGGATGGACGCCAGAGTTTGTTCAGATCATGCGGCCCTGAAGTCATTCAAGGAGTTTGCCGCTGGGAAAAGAGGAGCGGCTTTTGGAGTGATGGAAGCCCTGGGAACACTGCCAGGACACATGACAGAGAGATTCCAGGAAGCCATTGACAACCTCGCTGTGCTCATGCGGGCAGAGACTGGAAGCAGGCCTTACAAAGCCGCGGCGGCCCAATTGCCGGAGACCCTAGAGACCATTATGCTTTTGGGGTTGCTGGGAACAGTCTCGCTGGGAATCTTCTTCGTCTTGATGAGGAACAAGGGCATAGGGAAGATGGGCTTTGGAATGGTGACTCTTGGGGCCAGCGCATGGCTCATGTGGCTCTCGGAAATTGAGCCAGCCAGAATTGCATGTGTCCTCATTGTTGTGTTCCTATTGCTGGTGGTGCTCATACCTGAGCCAGAAAAGCAAAGATCTCCCCAGGACAACCAAATGGCAATCATCATCATGGTAGCAGTAGGTCTTCTGGGCTTGATTACCGCCAATGAACTCGGATGGTTGGAGAGAACAAAGAGTGACCTAAGCCATCTAATGGGAAGGAGAGAGGAGGGGGCAACCATAGGATTCTCAATGGACATTGACCTGCGGCCAGCCTCAGCTTGGGCCATCTATGCTGCCTTGACAACTTTCATTACCCCAGCCGTCCAACATGCAGTGACCACCTCATACAACAACTACTCCTTAATGGCGATGGCCACGCAAGCTGGAGTGTTGTTTGGTATGGGCAAAGGGATGCCATTCTACGCATGGG-ACTTTGGAGTCCCGCTGCTAATGATAGGTTGCTACTCACAATTAACACCCCTGACCCTAATAGTGGCCATCATTTTGCTCGTGGCGCACTACATGTACTTGATCCCAGGGCTGCAGGCAGCAGCTGCGCGTGCTGCCCAGAAGAGAACGGCAGCTGGCATCATGAAGAACCCTGTTGTGGATGGAATAGTGGTGACTGACATTGACACAATGACAATTGACCCCCAAGTGGAGAAAAAGATGGGACAGGTGCTACTCATAGCAGTAGCCGTCTCCAGCGCCATACTGTCGCGGACCGCCTGGGGGTGGGGGGAGGCTGGGGCCCTGATCACAGCCGCAACTTCCACTTTGTGGGAAGGCTCTCCGAACAAGTACTGGAACTCCTCTACAGCCACTTCACTGTGTAACATTTTTAGGGGAAGTTACTTGGCTGGAGCTTCTCTAATCTACACAGTAACAAGAAACGCTGGCTTGGTCAAGAGACGTGGGGGTGGAACAGGAGAGACCCTGGGAGAGAAATGGAAGGCCCGCTTGAACCAGATGTCGGCCCTGGAGTTCTACTCCTACAAAAAGTCAGGCATCACCGAGGTGTGCAGAGAAGAGGCCCGCCGCGCCCTCAAGGACGGTGTGGCAACGGGAGGCCATGCTGTGTCCCGAGGAAGTGCAAAGCTGAGATGGTTGGTGGAGCGGGGATACCTGCAGCCCTATGGAAAGGTCATTGATCTTGGATGTGGCAGAGGGGGCTGGAGTTACTACGCCGCCACCATCCGCAAAGTTCAAGAAGTGAAAGGATACACAAAAGGAGGCCCTGGTCATGAAGAACCCGTGTTGGTGCAAAGCTATGGGTGGAACATAGTCCGTCTTAAGAGTGGGGTGGACGTCTTTCATATGGCGGCTGAGCCGTGTGACACGTTGCTGTGTGACATAGGTGAGTCATCATCTAGTCCTGAAGTGGAAGAAGCACGGACGCTCAGAGTCCTCTCCATGGTGGGGGATTGGCTTGAAAAAAGACCAGGAGCCTTTTGTATAAAAGTGTTGTGCCCATACACCAGCACTATGATGGAAACCCTGGAGCGACTGCAGCGTAGGTATGGGGGAGGACTGGTCAGAGTGCCACTCTCCCGCAACTCTACACATGAGATGTACTGGGTCTCTGGAGCGAAAAGCAACACCATAAAAAGTGTGTCCACCACGAGCCAGCTCCTCTTGGGGCGCATGGACGGGCCTAGGAGGCCAGTGAAATATGAGGAGGATGTGAATCTCGGCTCTGGCACGCGGGCTGTGGTAAGCTGCGCTGAAGCTCCCAACATGAAGATCATTGGTAACCGCATTGAAAGGATCCGCAGTGAGCACGCGGAAACGTGGTTCTTTGACGAGAACCACCCATATAGGACATGGGCTTACCATGGAAGCTATGAGGCCCCCACACAAGGGTCAGCGTCCTCTCTAATAAACGGGGTTGTCAGGCTCCTGTCAAAACCCTGGGATGTGGTGACTGGAGTCACAGGAATAGCCATGACCGACACCACACCGTATGGTCAGCAAAGAGTTTTCAAGGAAAAAGTGGACACTAGGGTGCCAGACCCCCAAGAAGGCACTCGTCAGGTTATGAGCATGGTCTCTTCCTGGTTGTGGAAAGAGCTAGGCAAACACAAACGGCCACGAGTCTGTACCAAAGAAGAGTTCATCAACAAGGTTCGTAGCAATGCAGCATTAGGGGCAATATTTGAAGAGGAAAAAGAGTGGAAGACTGCAGTGGAAGCTGTGAACGATCCAAGGTTCTGGGCTCTAGTGGACAAGGAAAGAGAGCACCACCTGAGAGGAGAGTGCCAGAGTTGTGTGTACAACATGATGGGAAAAAGAGAAAAGAAACAAGGGGAATTTGGAAAGGCCAAGGGCAGCCGCGCCATCTGGTATATGTGGCTAGGGGCTAGATTTCTAGAGTTCGAAGCCCTTGGATTCTTGAACGAGGATCACTGGATGGGGAGAGAGAACTCAGGAGGTGGTGTTGAAGGGCTGGGATTACAAAGACTCGGATATGTCCTAGAAGAGATGAGTCGTATACCAGGAGGAAGGATGTATGCAGATGACACTGCTGGCTGGGACACCCGCATTAGCAGGTTTGATCTGGAGAATGAAGCTCTAATCACCAACCAAATGGAGAAAGGGCACAGGGCCTTGGCATTGGCCATAATCAAGTACACATACCAAAACAAAGTGGTAAAGGTCCTTAGACCAGCTGAAAAAGGGAAAACAGTTATGGACATTATTTCGAGACAAGACCAAAGGGGGAGCGGACAAGTTGTCACTTACGCTCTTAACACATTTACCAACCTAGTGGTGCAACTCATTCGGAATATGGAGGCTGAGGAAGTTCTAGAGATGCAAGACTTGTGGCTGCTGCGGAGGTCAGAGAAAGTGACTAACTGGTTGCAGAGCAACGGATGGGATAGGCTCAAACGAATGGCAGTCAGTGGAGATGATTGCGTTGTGAAGCCAATTGATGATAGGTTTGCACATGCCCTCAGGTTCTTGAATGATATGGGAAAAGTTAGGAAGGACACACAAGAGTGGAAACCCTCAACTGGATGGGACAACTGGGAAGAAGTTCCGTTTTGCTCCCACCACTTCAACAAGCTCCATCTCAAGGACGGGAGGTCCATTGTGGTTCCCTGCCGCCACCAAGATGAACTGATTGGCCGGGCCCGCGTCTCTCCAGGGGCGGGATGGAGCATCCGGGAGACTGCTTGCCTAGCAAAATCATATGCGCAAATGTGGCAGCTCCTTTATTTCCACAGAAGGGACCTCCGACTGATGGCCAATGCCATTTGTTCATCTGTGCCAGTTGACTGGGTTCCAACTGGGAGAACTACCTGGTCAATCCATGGAAAGGGAGAATGGATGACCACTGAAGACATGCTTGTGGTGTGGAACAGAGTGTGGATTGAGGAGAACGACCACATGGAAGACAAGACCCCAGTTACGAAATGGACAGACATTCCCTATTTGGGAAAAAGGGAAGACTTGTGGTGTGGATCTCTCATAGGGCACAGACCGCGCACCACCTGGGCTGAGAACATTAAAAACACAGTCAACATGGTGCGCAGGATCATAGGTGATGAAGAAAAGTACATGGACTACCTATCCACCCAAGTTCGCTACTTGGGTGAAGAAGGGTCTACACCTGGAGTGCTGTAA

>ENA|AHL43504|AHL43504.1 Zika virus polyprotein

ATGAAAAACCC-AAAGAAGAAATCCGGAGGATTCCGGATTGTCAATATGCTAAAACGCGGAGTAGCCCGTGTAAACCCCTTGGGAGGTTTGAAGAGGTTGCCAGCCGGACTTCTGCTGGGTCATGGACCCATCAGAATGGTTTTGGCGATACTAGCCTTTTTGAGATTTACAGCAATCAAGCCATCACTGGGCCTTATCAACAGATGGGGTTCCGTGGGGAAAAAAGAGGCTATGGAAATAATAAAGAAGTTCAAGAAAGATCTTGCTGCCATGTTGAGAATAATCAATGCTAGGAAAGAGAGGAAGAGACGTGGCGCAGACACCAGCATCGGAATCATTGGCCTCCTGCTGACTACAGCCATGGCAGCAGAGATCACTAGACGCGGGAGTGCATACTACATGTACTTGGATAGGAGCGATGCCGGGAAGGCCATTTCGTTTGCTACCACATTGGGAGTGAACAAGTGCCACGTACAGATCATGGACCTCGGGCACATGTGTGACGCCACCATGAGTTATGAGTGCCCTATGCTGGATGAGGGAGTGGAACCAGATGATGTCGATTGCTGGTGCAACACGACATCAACTTGGGTTGTGTACGGAACCTGTCATCACAAAAAAGGTGAGGCACGGCGATCTAGAAGAGCCGTGACGCTCCCTTCTCACTCTACAAGGAAGTTGCAAACGCGGTCGCAGACCTGGTTAGAATCAAGAGAATACACGAAGCACTTGATCAAGGTTGAAAACTGGATATTCAGGAACCCCGGGTTTGCGCTAGTGGCCGTTGCCATTGCCTGGCTTTTGGGAAGCTCGACGAGCCAAAAAGTCATATACTTGGTCATGATACTGCTGATTGCCCCGGCATACAGTATCAGGTGCATTGGAGTCAGCAATAGAGACTTCGTGGAGGGCATGTCAGGTGGGACCTGGGTTGATGTTGTCTTGGAACATGGAGGCTGCGTTACCGTGATGGCACAGGACAAGCCAACAGTTGACATAGAGTTGGTCACGACGACGGTTAGTAACATGGCCGAGGTAAGATCCTATTGCTACGAGGCATCGATATCGGACATGGCTTCGGACAGTCGTTGCCCAACACAAGGTGAAGCCTACCTTGACAAGCAATCAGACACTCAATATGTCTGCAAAAGAACATTAGTGGACAGAGGTTGGGGAAATGGTTGTGGACTTTTTGGCAAAGGGAGCTTGGTGACATGTGCCAAGTTCACGTGTTCTAAGAAGATGACCGGGAAGAGCATTCAACCGGAAAATCTGGAGTATCGGATAATGCTATCAGTGCATGGCTCCCAGCATAGCGGGATGATTGTCAATGATATAGGACATGAAACTGACGAAAACAGAGCGAAAGTCGAGGTTACGCCTAATTCACCAAGAGCGGAAGCAACCTTGGGAGGCTTTGGAAGCTTAGGACTTGACTGTGAACCAAGGACAGGCCTTGACTTTTCAGATCTGTATTACCTGACCATGAACAATAAGCATTGGTTGGTGCACAAAGAGTGGTTTCATGACATCCCATTGCCTTGGCATGCTGGGGCAGACACCGGAACTCCACACTGGAACAACAAAGAGGCATTGGTAGAATTCAAGGATGCCCACGCCAAGAGGCAAACCGTCGTCGTTCTGGGGAGCCAGGAAGGAGCCGTTCACACGGCTCTCGCTGGAGCTCTAGAGGCTGAGATGGATGGTGCAAAGGGAAGGCTGTTCTCTGGCCATTTGAAATGCCGCCTAAAAATGGACAAGCTTAGATTGAAGGGCGTGTCATATTCCTTGTGCACTGCGGCATTCACATTCACCAAGGTCCCAGCTGAAACACTGCATGGAACAGTCACAGTGGAGGTGCAGTATGCAGGGACAGATGGACCCTGCAAGGTCCCAGCCCAGATGGCGGTGGACATGCAGACCCTGACCCCAGTTGGAAGGCTGATAACCGCCAACCCCGTGATTACTGAAAGCACTGAGAACTCAAAGATGATGTTGGAGCTTGACCCACCATTTGGGGATTCTTACATTGTCATAGGAGTTGGGGACAAGAAAATCACCCACCACTGGCATAGGAGTGGTAGCACCATCGGAAAGGCATTTGAAGCCACTGTGAGAGGCGCCAAGAGAATGGCAGTCCTGGGGGATACAGCCTGGGACTTCGGATCAGTCGGGGGTGTGTTCAACTCACTGGGTAAGGGCATTCACCAGATTTTTGGAGCAGCCTTCAAATCACTGTTTGGAGGAATGTCCTGGTTCTCACAGATCCTCATAGGCACGCTGCTAGTGTGGTTAGGTTTGAACACAAAGAATGGATCTATCTCCCTCACATGCTTGGCCCTGGGGGGAGTGATGATCTTCCTCTCCACGGCTGTTTCTGCTGACGTGGGGTGCTCAGTGGACTTCTCAAAAAGAGAAACGAGATGTGGCACGGGGGTATTCATCTATAATGATGTTGAAGCCTGGAGGGACCGGTACAAGTACCATCCTGACTCCCCCCGCAGATTGGCAGCAGCAGTCAAGCAGGCCTGGGAAGAGGGGATCTGTGGGATCTCATCCGTTTCAAGAATGGAAAACATCATGTGGAAATCAGTAGAAGGGGAGCTCAATGCTATCCTAGAGGAGAATGGAGTTCAACTGACAGTTGTTGTGGGATCTGTAAAAAACCCCATGTGGAGAGGTCCACAAAGATTGCCAGTGCCTGTGAATGGGCTGCCCCATGGCTGGAAAGCCTGGGGGAAATCGTATTTTGTTAGGGCGGCAAAGACCAACAACAGTTTTGTTGTCGACGGTGACACACTGAAGGAATGTCCGCTTAAGCACAGAGCATGGAATAGTTTTCTTGTGGAGGATCACGGGTTTGGAGTCTTCCACACCAGTGTTTGGCTTAAGGTCAGAGAAGATTACTCATTAGAATGTGACCCAGCCGTCATAGGAACAGCTGTTAAGGGAAGGGAGGCCGCGCACAGTGATCTGGGCTATTGGATTGAAAGTGAAAAGAATGACACATGGAGGCTGAAGAGGGCCCACCTGATTGAGATGAAAACATGTGAATGGCCAAAGTCTCACACATTGTGGACAGATGGAGTAGAAGAAAGTGATCTTATCATACCCAAGTCTTTAGCTGGTCCACTCAGCCACCACAACACCAGAGAGGGTTACAGAACCCAAGTGAAAGGGCCATGGCACAGTGAAGAGCTTGAAATCCGGTTTGAGGAATGTCCAGGCACCAAGGTTCACGTGGAGGAGACATGCGGAACTAGAGGACCATCTCTGAGATCAACTACTGCAAGTGGAAGGGTCATTGAGGAATGGTGCTGTAGGGAATGCACAATGCCCCCACTATCGTTTCGAGCAAAAGACGGCTGCTGGTATGGAATGGAGATAAGGCCCAGGAAAGAACCAGAGAGCAACTTAGTGAGGTCAATGGTGACAGCGGGGTCAACCGATCATATGGACCACTTCTCTCTTGGAGTGCTTGTGATTCTACTCATGGTGCAGGAGGGGTTGAAGAAGAGAATGACCACAAAGATCATCATGAGCACATCAATGGCAGTGCTGGTAGTCATGATCTTGGGAGGATTTTCAATGAGTGACCTGGCCAAGCTTGTGATCCTGATGGGTGCTACTTTCGCAGAAATGAACACTGGAGGAGATGTAGCTCACTTGGCATTGGTAGCGGCATTTAAAGTCAGACCAGCCTTGCTGGTCTCCTTCATTCTCAGAGCCAATTGGACACCCCGTGAGAGCATGCTGCTAGCCCTGGCTTCGTGTCTTCTGCAAACTGCGATCTCTGCTCTTGAAGGTGACTTGATGGTCCTCGTTAATGGATTTGCTTTGGCCTGGTTGGCAATTCGAGCAATGGCCGTGCCACGCACTGACAACATCGCTCTAGCAATCTTGGCTGCTCTAACACCACTAGCTCGAGGCACACTGCTCGTGGCATGGAGAGCGGGCCTGGCTACTTGTGGAGGGTTCATGCTCCTCTCCCTGAAAGGGAAAGGTAGTGTGAAGAAGAACCTGCCATTTGTCATGGCCCTGGGATTGACAGCTGTGAGGGTAGTAGACCCTATTAATGTGGTAGGACTACTGTTACTCACAAGGAGTGGGAAGCGGAGCTGGCCCCCTAGTGAAGTTCTCACAGCCGTTGGCCTGATATGTGCACTGGCCGGAGGGTTTGCCAAGGCAGACATTGAGATGGCTGGACCCATGGCTGCAGTAGGCTTGCTAATTGTCAGCTATGTGGTCTCGGGAAAGAGTGTGGACATGTACATTGAAAGAGCAGGTGACATCACATGGGAAAAGGACGCGGAAGTCACTGGAAACAGTCCTCGGCTTGACGTGGCACTGGATGAGAGTGGTGATTTCTCCTTGGTAGAGGAAGATGGTCCACCCATGAGAGAGATCATACTCAAGGTGGTCCTGATGGCCATCTGTGGCATGAACCCAATAGCTATACCTTTTGCTGCAGGAGCGTGGTATGTGTATGTGAAGACTGGGAAAAGGAGTGGCGCCCTCTGGGACGTGCCTGCTCCCAAAGAAGTGAAGAAAGGAGAGACCACAGATGGAGTGTACAGAGTGATGACTCGCAGACTGCTAGGTTCAACACAGGTTGGAGTGGGAGTCATGCAAGAGGGAGTCTTCCACACCATGTGGCACGTTACAAAAGGAGCCGCACTGAGGAGCGGTGAGGGAAGACTTGATCCATACTGGGGGGATGTCAAGCAGGACTTGGTGTCATACTGTGGGCCTTGGAAGTTGGATGCAGCTTGGGATGGACTTAGCGAGGTACAGCTTTTGGCCGTACCTCCCGGAGAGAGGGCCAGAAACATTCAGACCCTGCCTGGAATATTCAAGACAAAGGACGGGGACATCGGAGCAGTTGCTCTGGACTACCCTGCAGGGACCTCAGGATCTCCGATCCTAGACAAATGTGGAAGAGTGATAGGACTCTATGGCAATGGGGTTGTGATCAAGAATGGAAGCTATGTTAGTGCTATAACCCAGGGAAAGAGGGAGGAGGAGACTCCGGTTGAATGTTTCGAACCCTCGATGCTGAAGAAGAAGCAGCTAACTGTCCTGGATCTGCATCCAGGAGCCGGAAAAACCAGGAGAGTTCTTCCTGAAATAGTCCGTGAAGCCATAAAAAAGAGACTCCGGACAGTGATCTTGGCACCAACTAGGGTTGTCGCTGCTGAGATGGAGGAAGCCTTGAGAGGACTTCCGGTGCGTTACATGACAACAGCAGTCAACGTCACCCATTCTGGGACAGAAATCGTTGATTTGATGTGCCATGCCACTTTCACTTCACGCTTACTACAACCCATCAGAGTCCCTAATTACAATCTCAACATCATGGATGAAGCCCACTTCACAGACCCCTCAAGTATAGCTGCAAGAGGATACATATCAACAAGGGTTGAAATGGGCGAGGCGGCTGCCATTTTTATGACTGCCACACCACCAGGAACCCGTGATGCGTTTCCTGACTCTAACTCACCAATCATGGACACAGAAGTGGAAGTCCCAGAGAGAGCCTGGAGCTCAGGCTTTGATTGGGTGACAGACCATTCTGGGAAAACAGTTTGGTTCGTTCCAAGCGTGAGAAACGGAAATGAAATCGCAGCCTGTCTGACAAAGGCTGGAAAGCGGGTCATACAGCTCAGCAGGAAGACTTTTGAGACAGAATTTCAGAAAACAAAAAATCAAGAGTGGGACTTTGTCATAACAACTGACATCTCAGAGATGGGCGCCAACTTCAAGGCTGACCGGGTCATAGACTCTAGGAGATGCCTAAAACCAGTCATACTTGATGGTGAGAGAGTCATCTTGGCTGGGCCCATGCCTGTCACGCATGCTAGTGCTGCTCAGAGGAGAGGACGTATAGGCAGGAACCCTAACAAACCTGGAGATGAGTACATGTATGGAGGTGGGTGTGCAGAGACTGATGAAGGCCATGCACACTGGCTTGAAGCAAGAATGCTTCTTGACAACATCTACCTCCAGGATGGCCTCATAGCCTCGCTCTATCGGCCTGAGGCCGATAAGGTAGCCGCCATTGAGGGAGAGTTTAAGCTGAGGACAGAGCAAAGGAAGACCTTCGTGGAACTCATGAAGAGAGGAGACCTTCCCGTCTGGCTAGCCTATCAGGTTGCATCTGCCGGAATAACTTACACAGACAGAAGATGGTGCTTTGATGGCACAACCAACAACACCATAATGGAAGACAGTGTACCAGCAGAGGTTTGGACAAAGTATGGAGAGAAGAGAGTGCTCAAACCGAGATGGATGGATGCTAGGGTCTGTTCAGACCATGCGGCCCTGAAGTCGTTCAAAGAATTCGCCGCTGGAAAAAGAGGAGCGGCTTTGGGAGTAATGGAGGCCCTGGGAACACTGCCAGGACACATGACAGAGAGGTTTCAGGAAGCCATTGACAACCTCGCCGTGCTCATGCGAGCAGAGACTGGAAGCAGGCCTTATAAGGCAGCGGCAGCCCAACTGCCGGAGACCCTAGAGACCATTATGCTCTTAGGTTTGCTGGGAACAGTTTCACTGGGGATCTTCTTTGTCTTGATGCGGAATAAGGGCATCGGGAAGATGGGCTTTGGAATGGTAACCCTTGGGGCCAGTGCATGGCTCATGTGGCTTTCGGAAATTGAACCAGCCAGAATTGCATGTGTCCTCATTGTTGTGTTTTTATTACTGGTGGTGCTCATACCCGAGCCAGAGAAGCAAAGATCTCCCCAAGATAACCAGATGGCAATTATCATCATGGTGGCAGTGGGCCTTCTAGGTTTGATAACTGCAAACGAACTTGGATGGCTGGAAAGAACAAAAAATGACATAGCTCATCTAATGGGAAGGAGAGAAGAAGGAGCAACCATGGGATTCTCAATGGACATTGATCTGCGGCCAGCCTCCGCCTGGGCTATCTATGCCGCATTGACAACTCTCATCACCCCAGCTGTCCAACATGCGGTAACCACTTCATACAACAACTACTCCTTAATGGCGATGGCCACACAAGCTGGAGTGCTGTTTGGCATGGGCAAAGGGATGCCATTTTATGCATGGG-ACTTTGGAGTCCCGCTGCTAATGATGGGTTGCTATTCACAATTAACACCCCTGACTCTGATAGTAGCTATCATTCTGCTTGTGGCGCACTACATGTACTTGATCCCAGGCCTACAAGCGGCAGCAGCGCGTGCTGCCCAGAAAAGGACAGCAGCTGGCATCATGAAGAATCCCGTTGTGGATGGAATAGTGGTAACTGACATTGACACAATGACAATAGACCCCCAGGTGGAGAAGAAGATGGGACAAGTGTTACTCATAGCAGTAGCCATCTCCAGTGCTGTGCTGCTGCGGACCGCCTGGGGATGGGGGGAGGCTGGAGCTCTGATCACAGCAGCGACCTCCACCTTGTGGGAAGGCTCTCCAAACAAATACTGGAACTCCTCTACAGCCACCTCACTGTGCAACATCTTCAGAGGAAGCTATCTGGCAGGAGCTTCCCTTATCTATACAGTGACGAGAAACGCTGGCCTGGTTAAGAGACGTGGAGGTGGGACGGGAGAGACTCTGGGAGAGAAGTGGAAAGCTCGTCTGAATCAGATGTCGGCCCTGGAGTTCTACTCTTATAAAAAGTCAGGTATCACTGAAGTGTGTAGAGAGGAGGCTCGCCGTGCCCTCAAGGATGGAGTGGCCACAGGAGGACATGCCGTATCCCGGGGAAGTGCAAAGCTCAGATGGTTGGTGGAGAGAGGATATCTGCAGCCCCATGGGAAGGTTGTTGACCTCGGATGTGGCAGAGGGGGCTGGAGCTATTATGCCGCCACCATCCGCAAAGTGCAGGAGGTGAGAGGATACACAAAGGGAGGTCCCGGTCATGAAGAACCCATGCTGGTGCAAAGCTATGGGTGGAACATAGTTCGTCTCAAGAGTGGAGTGGACGTCTTCCACATGGCGGCTGAGCCGTGTGACACTCTGCTGTGTGACATAGGTGAGTCATCATCTAGTCCTGAAGTGGAAGAGACACGAACACTCAGAGTGCTCTCTATGGTGGGGGACTGGCTTGAAAAAAGACCAGGGGCCTTCTGTATAAAGGTGCTGTGCCCATACACCAGCACTATGATGGAAACCATGGAGCGACTGCAACGTAGGCATGGGGGAGGATTAGTCAGAGTGCCATTGTCTCGCAACTCCACACATGAGATGTACTGGGTCTCTGGGGCAAAAAGCAACATCATAAAAAGTGTGTCCACCACAAGTCAGCTCCTCCTGGGACGCATGGATGGCCCCAGGAGGCCAGTGAAATATGAGGAGGATGTGAACCTCGGCTCGGGTACACGAGCTGTGGCAAGCTGTGCTGAGGCTCCTAACATGAAAATCATCGGCAGGCGCATTGAGAGAATCCGCAATGAACATGCAGAAACATGGTTTCTTGATGAAAACCACCCATACAGGACATGGGCCTACCATGGGAGCTACGAAGCCCCCACGCAAGGATCAGCGTCTTCCCTCGTGAACGGGGTTGTTAGACTCCTGTCAAAGCCTTGGGACGTGGTGACTGGAGTTACAGGAATAGCCATGACTGACACCACACCATACGGCCAACAAAGAGTCTTCAAAGAAAAAGTGGACACCAGGGTGCCAGATCCCCAAGAAGGCACTCGCCAGGTAATGAACATGGTCTCTTCCTGGCTGTGGAAGGAGCTGGGGAAACGCAAGCGGCCACGCGTCTGCACCAAAGAAGAGTTTATCAACAAGGTGCGCAGCAATGCAGCACTGGGAGCAATATTTGAAGAGGAAAAAGAATGGAAGACGGCTGTGGAAGCTGTGAATGATCCAAGGTTTTGGGCCCTAGTGGATAGGGAGAGAGAACACCACCTGAGAGGAGAGTGTCACAGCTGTGTGTACAACATGATGGGAAAAAGAGAAAAGAAGCAAGGAGAGTTCGGGAAAGCAAAAGGTAGCCGCGCCATCTGGTACATGTGGTTGGGAGCCAGATTCTTGGAGTTTGAAGCCCTTGGATTCTTGAACGAGGACCATTGGATGGGAAGAGAGAACTCTGGAGGCGGAGTTGAAGGACTGGGACTGCAGAGACTGGGCTATGTCCTTGAGGAGATGAGCCAGGCACCAGGAGGGAAGATGTACGCAGATGACACTGCTGGCTGGGACACCCGCATTAGTAAGTTTGATCTGGAGAATGAAGCTTTGATTACCAACCAAATGGAGGAAGGGCACAGAACTCTGGCGTTGGCCGTGATTAAATACACATACCAAAACAAAGTGGTGAAGGTCCTCAGACCAGCTGAAGGAGGAAAAACAGTTATGGACATCATTTCAAGACAAGACCAGAGAGGGAGTGGACAAGTTGTCACTTATGCTCTCAACACATTCACCAACTTGGTGGTGCAGCTTATCCGGAACATGGAAGCTGAGGAAGTGTTAGAGATGCAAGACTTATGGCTGTTGAGGAAGCCAGAGAAAGTGACCAGATGGTTGCAGAGCAATGGATGGGATGGACTCAAACGAATGGCGGTCAGTGGAGATGACTGCGTTGTGAAGCCAATCGATGATAGGTTTGCACATGCCCTCAGGTTCTTGAATGACATGGGAAAAGTTAGGAAAGACACACAGGAGTGGAAACCCTCGACTGGATGGAGCAATTGGGAAGAAGTCCCGTTCTGCTCCCACCACTTCAACAAGCTGTACCTCAAGGATGGGAGATCCATTGTGGTCCCTTGCCGCCACCAAGATGAACTGATTGGCCGAGCTCGCGTTTCACCAGGGGCAGGATGGAGCATCCGGGAGACTGCCTGTCTTGCAAAATCATATGCGCAGATGTGGCAGCTCCTTTATTTCCACAGAAGAGACCTTCGACTGATGGCTAATGCCATTTGCTCGGCTGTACCAGTTGACTGGGTACCAACTGGGAGAACCACCTGGTCAATCCATGGAAAGGGAGAATGGATGACCACTGAGGACATGCTCATGGTGTGGAATAGAGTGTGGATTGAGGAGAACGACCATATGGAGGACAAGACTCCTGTAACAAAATGGACAGACATTCCCTATCTAGGAAAAAGGGAGGACTTATGGTGTGGATCCCTTATAGGGCACAGACCCCGCACCACTTGGGCTGAAAACATCAAAGACACAGTCAACATGGTGCGCAGGATCATAGGTGATGAAGAAAAGTACATGGACTATCTATCCACCCAAGTCCGCTACTTGGGTGAGGAAGGGTCCACACCCGGAGTGTTGTAA

>ENA|AMB18850|AMB18850.1 Zika virus polyprotein

ATGAAAAACCC-AAAAAAGAAATCCGGAGGATTCCGGATTGTCAATATGCTAAAACGCGGAGTAGCCCGTGTGAGCCCCTTTGGGGGCTTGAAGAGGCTGCCAGCCGGACTTCTGCTGGGTCATGGGCCCATCAGGATGGTCTTGGCAATTCTAGCCTTTTTGAGATTCACGGCAATCAAGCCATCACTGGGTCTCATCAATAGATGGGGTTCAGTGGGGAAAAAAGAGGCTATGGAAATAATAAAGAAGTTCAAGAAAGATCTGGCTGCCATGCTGAGAATAATCAATGCTAGGAAGGAGAAGAAGAGACGAGGCGCAGATACTAGTGTCGGAATTGTTGGCCTCCTGCTGACCACAGCTATGGCAGCGGAGGTCACTAGACGTGGGAGTGCATACTATATGTACTTGGACAGAAACGATGCTGGGGAGGCCATATCTTTTCCAACCACATTGGGGATGAATAAGTGTTATATACAGATCATGGATCTTGGACACATGTGTGATGCCACCATGAGCTATGAATGCCCTATGCTGGATGAGGGGGTGGAACCAGATGACGTCGATTGTTGGTGCAACACGACGTCAACTTGGGTTGTGTACGGAACCTGCCATCACAAAAAAGGTGAAGCACGGAGATCTAGAAGAGCTGTGACGCTCCCCTCCCATTCCACTAGGAAGCTGCAAACGCGGTCGCAAACCTGGTTGGAATCAAGAGAATACACAAAGCACTTGATTAGAGTCGAAAATTGGATATTCAGGAACCCTGGCTTCGCGTTAGCAGCAGCTGCCATCGCTTGGCTTTTGGGAAGCTCAACGAGCCAAAAAGTCATATACTTGGTCATGATACTGCTGATTGCCCCGGCATACAGCATCAGGTGCATAGGAGTCAGCAATAGGGACTTTGTGGAAGGTATGTCAGGTGGGACTTGGGTTGATGTTGTCTTGGAACATGGAGGTTGTGTCACCGTAATGGCACAGGACAAACCGACTGTCGACATAGAGCTGGTTACAACAACAGTCAGCAACATGGCGGAGGTAAGATCCTACTGCTATGAGGCATCAATATCAGACATGGCTTCGGACAGCCGCTGCCCAACACAAGGTGAAGCCTACCTTGACAAGCAATCAGACACTCAATATGTCTGCAAAAGAACGTTAGTGGACAGAGGCTGGGGAAATGGATGTGGACTTTTTGGCAAAGGGAGCCTGGTGACATGCGCTAAGTTTGCATGCTCCAAGAAAATGACCGGGAAGAGCATCCAGCCAGAGAATCTGGAGTACCGGATAATGCTGTCAGTTCATGGCTCCCAGCACAGTGGGATGATCGTTAATGACACAGGACATGAAACTGATGAGAATAGAGCGAAGGTTGAGATAACGCCCAATTCACCAAGAGCCGAAGCCACCCTGGGGGGTTTTGGAAGCCTAGGACTTGATTGTGAACCGAGGACAGGCCTTGACTTTTCAGATTTGTATTACTTGACTATGAATAACAAGCACTGGTTGGTCCACAAGGAGTGGTTCCACGACATTCCATTACCTTGGCACGCTGGGGCAGACACCGGAACTCCACACTGGAACAACAAAGAAGCACTGGTAGAGTTCAAGGACGCACATGCCAAAAGGCAAACTGTCGTGGTTCTAGGGAGTCAAGAAGGAGCAGTTCACACGGCCCTTGCTGGAGCTCTGGAGGCTGAGATGGATGGTGCAAAGGGAAGGCTGTCCTCTGGCCACTTGAAATGTCGCCTGAAAATGGATAAACTTAGATTGAAGGGCGTGTCATACTCCTTGTGTACCGCAGCGTTCACATTCACCAAGATCCCGGCTGAAACACTGCACGGGACAGTCACAGTGGAGGTACAGTACGCAGGGACAGATGGACCTTGCAAGGTTCCAGCTCAGATGGCGGTGGACATGCAAACTCTGACCCCAGTTGGGAGGTTGATAACCGCTAACCCCGTAATCACTGAAAGCACTGAGAACTCTAAGATGATGCTGGAACTTGATCCACCATTTGGGGACTCTTACATTGTCATAGGAGTCGGGGAGAAGAAGATCACCCACCACTGGCACAGGAGTGGCAGCACCATTGGAAAAGCATTTGAAGCCACTGTGAGAGGTGCCAAGAGAATGGCAGTCTTGGGAGACACAGCCTGGGACTTTGGATCAGTTGGAGGCGCTCTCAACTCATTGGGCAAGGGCATCCATCAAATTTTTGGAGCAGCTTTCAAATCATTGTTTGGAGGAATGTCCTGGTTCTCACAAATCCTCATTGGAACGTTGCTGATGTGGTTGGGTCTGAACACAAAGAATGGATCTATTTCCCTTATGTGCTTGGCCTTAGGGGGAGTGTTGATCTTCTTATCCACAGCCGTCTCTGCTGATGTGGGGTGCTCGGTGGACTTCTCAAAGAAGGAGACGAGATGCGGTACAGGGGTGTTCGTCTATAACGACGTTGAAGCCTGGAGGGACAGGTACAAGTACCATCCTGACTCCCCCCGTAGATTGGCAGCAGCAGTCAAGCAAGCCTGGGAAGATGGTATCTGCGGGATCTCCTCTGTTTCAAGAATGGAGAACATCATGTGGAGATCAGTAGAAGGGGAGCTCAACGCAATCTTGGAAGAGAATGGAGTTCAACTGACGGTCGTTGTGGGATCTGTAAAAAACCCCATGTGGAGAGGTCCACAGAGATTGCCCGTGCCTGTGAACGAGCTGCCCCACGGCTGGAAGGCTTGGGGGAAATCGTACTTCGTCAGAGCAGCAAAGACAAATAACAGCTTTGTCGTGGATGGTGACACACTGAAGGAATGCCCACTCGAACATAGAGCATGGAACAGCTTTCTTGTGGAGGATCATGGGTTCGGGGTATTTCACACTAGTGTCTGGCTCAAGGTTAGAGAAGATTATTCATTAGAGTGTGATCCAGCCGTTATTGGAACAGCTGTTAAGGGGAAGGAGGCTGTACACAGTGATCTAGGCTACTGGATTGAGAGTGAGAAGAATGACACATGGAGGCTGAAGAGGGCCCATCTAATCGAGATGAAAACATGTGAATGGCCAAAGTCCCACACATTGTGGGCAGATGGAATAGAAGAGAGTGATCTGATCATTCCCAAGTCTTTAGCTGGGCCACTCAGCCATCACAATACCAGAGAGGGCTACAGGACCCAAATGAAAGGGCCATGGCACAGTGAAGAGCTTGAAATTCGGTTTGAGGAATGCCCGGGCACTAAGGTCCACGTGGAGGAAACATGTGGAACAAGAGGACCATCTCTGAGATCAACCACTGCAAGCGGAAGGGTGATCGAGGAATGGTGCTGCAGGGAGTGCACAATGCCCCCACTGTCGTTCCGGGCTAAAGATGGCTGTTGGTATGGAATGGAGATAAGGCCCAGGAAAGAACCAGAAAGCAACTTAGTAAGGTCAGTGGTGACTGCAGGATCAACTGATCACATGGATCACTTCTCCCTTGGAGTGCTTGTGATTCTGCTCATGGTGCAGGAAGGGCTGAAGAAGAGAATGACCACAAAGATCATCATAAGCACATCAATGGCAGTGCTGGTAGCTATGATCCTGGGAGGATTTTCAATGAGTGACCTGGCTAAGCTTGCAATTTTGATGGGCGCCACCTTCGCGGAAATGAACACTGGAGGAGATGTAGCTCATCTGGCGCTGATAGCGGCATTCAAAGTCAGACCAGCGTTGCTGGTATCTTTCATCTTCAGAGCTAATTGGACACCCCGTGAAAGCATGCTGCTGGCCTTGGCCTCGTGTCTTTTGCAAACTGCGATCTCCGCCTTGGAAGGCGACCTGATGGTTCTCATCAATGGTTTTGCTTTGGCCTGGTTGGCAATACGAGCGATGGTTGTTCCACGCACTGATAACATCACCTTGGCAATCCTGGCTGCTCTGACACCACTGGCCCGGGGCACACTGCTTGTGGCGTGGAGAGCAGGCCTTGCTACTTGCGGGGGGTTTATGCTCCTCTCTCTGAAGGGAAAAGGCAGTGTGAAGAAGAACTTACCATTTGTCATGGCCCTGGGACTAACCGCTGTGAGGCTGGTCGACCCCATCAACGTGGTGGGACTGCTGTTGCTCACAAGGAGTGGGAAGCGGAGCTGGCCCCCTAGCGAAGTACTCACAGCTGTTGGCCTGATATGCGCATTGGCTGGAGGGTTCGCCAAGGCAGATATAGAGATGGCTGGGCCCATGGCCGCGGTCGGTCTGCTAATTGTCAGTTACGTGGTCTCAGGAAAGAGTGTGGACATGTACATTGAAAGAGCAGGTGACATCACATGGGAAAAAGATGCGGAAGTCACTGGAAACAGTCCCCGGCTCGATGTGGCGCTAGATGAGAGTGGTGATTTCTCCCTGGTGGAGGATGACGGTCCCCCCATGAGAGAGATCATACTCAAGGTGGTCCTGATGACCATCTGTGGCATGAACCCAATAGCCATACCCTTTGCAGCTGGAGCGTGGTACGTATACGTGAAGACTGGAAAAAGGAGTGGTGCTCTATGGGATGTGCCTGCTCCCAAGGAAGTAAAAAAGGGGGAGACCACAGATGGAGTGTACAGAGTAATGACTCGTAGACTGCTAGGTTCAACACAAGTTGGAGTGGGAGTTATGCAAGAGGGGGTCTTTCACACTATGTGGCACGTCACAAAAGGATCCGCGCTGAGAAGCGGTGAAGGGAGACTTGATCCATACTGGGGAGATGTCAAGCAGGATCTGGTGTCATACTGTGGTCCATGGAAGCTAGATGCCGCCTGGGACGGGCACAGCGAGGTGCAGCTCTTGGCCGTGCCCCCCGGAGAGAGAGCGAGGAACATCCAGACTCTGCCCGGAATATTTAAGACAAAGGATGGGGACATTGGAGCGGTTGCGCTGGATTACCCAGCAGGAACTTCAGGATCTCCAATCCTAGACAAGTGTGGGAGAGTGATAGGACTTTATGGCAATGGGGTCGTGATCAAAAATGGGAGTTATGTTAGTGCCATCACCCAAGGGAGGAGGGAGGAAGAGACTCCTGTTGAGTGCTTCGAGCCTTCGATGCTGAAGAAGAAGCAGCTAACTGTCTTAGACTTGCATCCTGGAGCTGGGAAAACCAGGAGAGTTCTTCCTGAAATAGTCCGTGAAGCCATAAAAACAAGACTCCGTACTGTGATCTTAGCTCCAACCAGGGTTGTCGCTGCTGAAATGGAGGAAGCCCTTAGAGGGCTTCCAGTGCGTTATATGACAACAGCAGTCAATGTCACCCACTCTGGAACAGAAATCGTCGACTTAATGTGCCATGCCACCTTCACTTCACGTCTACTACAGCCAATCAGAGTCCCCAACTATAATCTGTATATTATGGATGAGGCCCACTTCACAGATCCCTCAAGTATAGCAGCAAGAGGATACATTTCAACAAGGGTTGAGATGGGCGAGGCGGCTGCCATCTTCATGACCGCCACGCCACCAGGAACCCGTGACGCATTTCCGGACTCCAACTCACCAATTATGGACACCGAAGTGGAAGTCCCAGAGAGAGCCTGGAGCTCAGGCTTTGATTGGGTGACGGATCATTCTGGAAAAACAGTTTGGTTTGTTCCAAGCGTGAGGAACGGCAATGAGATCGCAGCTTGTCTGACAAAGGCTGGAAAACGGGTCATACAGCTCAGCAGAAAGACTTTTGAGACAGAGTTCCAGAAAACAAAACATCAAGAGTGGGACTTTGTCGTGACAACTGACATTTCAGAGATGGGCGCCAACTTTAAAGCTGACCGTGTCATAGATTCCAGGAGATGCCTAAAGCCGGTCATACTTGATGGCGAGAGAGTCATTTTGGCTGGACCCATGCCTGTCACACATGCCAGCGCTGCCCAGAGGAGGGGGCGCATAGGCAGGAATCCCAACAAACCTGGAGATGAGTATCTGTATGGAGGTGGGTGCGCAGAGACTGACGAAGACCATGCACACTGGCTTGAAGCAAGAATGCTCCTTGACAATATTTACCTCCAAGATGGCCTCATAGCCTCGCTCTATCGACCTGAGGCCGACAAAGTAGCAGCCATTGAGGGAGAGTTCAAGCTTAGGACGGAGCAAAGGAAGACCTTTGTGGAACTCATGAAAAGAGGAGATCTTCCTGTTTGGCTGGCCTATCAGGTTGCATCTGCCGGAATAACCTACACAGATAGAAGATGGTGCTTTGATGGCACGACCAACAACACCATAATGGAAGACAGTGTGCCGGCAGAGGTGTGGACCAGACACGGAGAGAAAAGAGTGCTCAAACCGAGGTGGATGGACGCCAGAGTTTGTTCAGATCATGCGGCCCTGAAGTCATTCAAGGAGTTTGCCGCTGGGAAAAGAGGAGCGGCTTTTGGAGTGATGGAAGCCCTGGGAACACTGCCAGGACACATGACAGAGAGATTCCAGGAAGCCATTGACAACCTCGCTGTGCTCATGCGGGCAGAGACTGGAAGCAGGCCTTACAAAGCCGCGGCGGCCCAATTGCCGGAGACCCTAGAGACCATTATGCTTTTGGGGTTGCTGGGAACAGTCTCGCTGGGAATCTTTTTCGTCTTGATGAGGAACAAGGGCATAGGGAAGATGGGCTTTGGAATGGTGACTCTTGGGGCCAGCGCATGGCTCATGTGGCTCTCGGAAATTGAGCCAGCCAGAATTGCATGTGTCCTCATTGTTGTGTTCCTATTGCTGGTGGTGCTCATACCTGAGCCAGAAAAGCAAAGATCTCCCCAGGACAACCAAATGGCAATCATCATCATGGTAGCAGTAGGTCTTCTGGGCTTGATTACCGCCAATGAACTCGGATGGTTGGAGAGAACAAAGAGTGACCTAAGCCATCTAATGGGAAGGAGAGAGGAGGGAGCAACCATAGGATTCTCAATGGACATTGACCTGCGGCCAGCCTCAGCTTGGGCCATCTATGCTGCCTTGACAACTTTCATTACCCCAGCCGTCCAACATGCAGTGACCACTTCATACAACAACTACTCCTTAATGGCGATGGCCACGCAAGCTGGAGTGTTGTTTGGTATGGGCAAAGGGATGCCATTCTACGCATGGG-ACTTTGGAGTCCCGCTGCTAATGATAGGTTGCTACTCACAATTAACACCCCTGACCCTAATAGTGGCCATCATTTTGCTCGTGGCGCACTACATGTACTTGATCCCAGGGCTGCAGGCAGCAGCTGCGCGTGCTGCCCAGAAGAGAACGGCAGCTGGCATCATGAAGAACCCTGTTGTGGATGGAATAGTGGTGACTGACATTGACACAATGACAATTGACCCCCAAGTGGAGAAAAAGATGGGACAGGTGCTACTCATAGCAGTAGCAGTCTCCAGCGCCATACTGTCGCGGACCGCCTGGGGGTGGGGGGAGGCTGGGGCCCTGATCACAGCCGCAACTTCCACTTTGTGGGAAGGCTCTCCGAACAAGTACTGGAACTCCTCTACAGCCACTTCACTGTGTAACATTTTTAGGGGAAGTTACTTGGCTGGAGCTTCTCTAATCTACATAGTAACAAGAAACGCTGGCTTGGTCAAGAGACGTGGGGGTGGAACAGGAGAGACCCTGGGAGAGAAATGGAAGGCCCGCTTGAACCAGATGTCGGCCCTGGAGTTCTACTCCTACAAAAAGTCAGGCATCACCGAGGTGTGCAGAGAAGAGGCCCGCCGCGCCCTCAAGGATGGTGTGGCAACGGGAGGCCATGCTGTGTCCCGAGGAAGTGCAAAGCTGAGATGGTTGGTGGAGCGGGGATACCTGCAGCCCTATGGAAAGGTCATTGATCTTGGATGTGGCAGAGGGGGCTGGAGTTACTACGCCGCCACCATCCGCAAAGTTCAAGAAGTGAAAGGATACACAAAAGGAGGCCCTGGTCATGAAGAACCCGTGTTGGTGCAAAGCTATGGGTGGAACATAGTCCGTCTTAAGAGTGGGGTGGACGTCTTTCATATGGCGGCTGAGCCGTGTGACACGTTGCTGTGTGACATAGGTGAGTCATCATCTAGTCCTGAAGTGGAAGAAGCACGGACGCTCAGAGTCCTCTCCATGGTGGGGGATTGGCTTGAAAAAAGACCAGGAGCCTTTTGTATAAAAGTGTTGTGCCCATACACCAGCACTATGATGGAAACCCTGGAGCGACTGCAGCGTAGGTATGGGGGAGGACTGGTCAGAGTGCCACTCTCCCGCAACTCTACACATGAGATGTACTGGGTCTCTGGAGCGAAAAGCAACACCATAAAAAGTGTGTCCACCACGAGCCAGCTCCTCTTGGGGCGCATGGACGGGCCTAGGAGGCCAGTGAAATATGAGGAGGATGTGAATCTCGGCTCTGGCACGCGGGCTGTGGTAAGCTGCGCTGAAGCTCCCAACATGAAGATCATTGGTAACCGCATTGAAAGGATCCGCAGTGAGCACGCGGAAACGTGGTTCTTTGACGAGAACCACCCATATAGGACATGGGCTTACCATGGAAGCTATGAGGCCCCCACACAAGGGTCAGCGTCCTCTCTAATAAACGGGGTTGTCAGGCTCCTGTCAAAACCCTGGGATGTGGTGACTGGAGTCACAGGAATAGCCATGACCGACACCACACCGTATGGTCAGCAAAGAGTTTTCAAGGAAAAAGTGGACACTAGGGTGCCAGACCCCCAAGAAGGCACTCGTCAGGTTATGAGCATGGTCTCTTCCTGGTTGTGGAAAGAGCTAGGCAAACACAAACGGCCACGAGTCTGTACCAAAGAAGAGTTCATCAACAAGGTTCGTAGCAATGCAGCATTAGGGGCAATATTTGAAGAGGAAAAAGAGTGGAAGACTGCAGTGGAAGCTGTGAACGATCCAAGGTTCTGGGCTCTAGTGGACAAGGAAAGAGAGCACCACCTGAGAGGAGAGTGCCAGAGTTGTGTGTACAACATGATGGGAAAAAGAGAAAAGAAACAAGGGGAATTTGGAAAGGCCAAGGGCAGCCGCGCCATCTGGTATATGTGGCTAGGGGCTAGATTTCTAGAGTTCGAAGCCCTTGGATTCTTGAACGAGGATCACTGGATGGGGAGAGAGAACTCAGGAGGTGGTGTTGAAGGGCTGGGATTACAAAGACTCGGATATGTCCTAGAAGAGATGAGTCGCATACCAGGAGGAAGGATGTATGCAGATGACACTGCTGGCTGGGACACCCGCATCAGCAGGTTCGATCTGGAGAATGAAGCTCTAATCACCAACCAAATGGAGAAAGGGCATAGGGCCTTGGCATTGGCCATAATCAAGTACACATACCAAAACAAAGTGGTAAAGGTCCTTAGACCAGCTGAAAAAGGGAAAACAGTTATGGACATTATTTCGAGACAAGACCAAAGGGGGAGCGGACAAGTTGTCACTTACGCTCTTAACACATTTACCAACCTAGTGGTGCAACTCATTCGGAATATGGAGGCTGAGGAAGTTCTAGAGATGCAAGACTTGTGGCTGCTGCGGAGGTCAGAGAAAGTGACCAACTGGTTGCAGAGCAACGGATGGGATAGGCTCAAACGAATGGCAGTCAGTGGAGATGATTGCGTTGTGAAGCCAATTGATGATAGGTTTGCACATGCCCTCAGGTTCTTGAATGATATGGGAAAAGTTAGGAAGGACACACAAGAGTGGAAACCCTCAACTGGATGGGACAACTGGGAAGAAGTTCCGTTTTGCTCCCACCACTTCAACAAGCTCCATCTCAAGGACGGGAGGTCCATTGTGGTTCCCTGCCGCCACCAAGATGAACTGATTGGCCGGGCCCGCGTCTCTCCAGGGGCGGGATGGAGCATCCGGGAGACTGCTTGCCTAGCAAAATCATATGCGCAAATGTGGCAGCTCCTTTATTTCCACAGAAGGGACCTCCGACTGATGGCCAATGCCATTTGTTCATCTGTGCCAGTTGACTGGGTTCCAACTGGGAGAACTACCTGGTCAATCCATGGAAAGGGAGAATGGATGACCACTGAAGACATGCTTGTGGTGTGGAACAGAGTGTGGATTGAGGAGAACGACCACATGGAAGACAAGACCCCAGTTACGAAATGGACAGACATTCCCTATTTGGGAAAAAGGGAAGACTTGTGGTGTGGATCTCTCATAGGGCACAGACCGCGCACCACCTGGGCTGAGAACATTAAAAACACAGTCAACATGGTGCGCAGGATCATAGGTGATGAAGAAAAGTACATGGACTACCTATCCACCCAAGTTCGCTACTTGGGTGAAGAAGGGTCTACACCTGGAGTGCTGTAA

>ENA|AHL43502|AHL43502.1 Zika virus polyprotein

ATGAAAAACCC-AAAGAGGAAATCCGGAGGATTCCGGATTGTCAATATGCTAAAACGCGGAGTAGCCCGTGTAAACCCCTTGGGGGGTTTGAAGAGGCTGCCAGCCGGACTTCTGCTGGGTCATGGACCCATTAGAATGGTTTTGGCGATACTAGCCTTTTTGAGATTTACAGCAATCAAGCCATCACTGGGCCTCATCAATAGATGGGGTTCAGTGGGGAAAAAAGAGGCTATGGAGATAATAAAGAAGTTTAAGAAAGATCTTGCTGCCATGTTGAGAATAATCAATGCTAGGAAGGAGAGGAAGAGACGTGGTGCAGACACCAGTATCGGAATCATTGGCCTCCTGCTGACTACAGCCATGGCAGCAGAGATTACTAGACGAGGAAGTGCATACTACATGTACTTGGACAGGAGCGATGCTGGGAAGGCCATTTCCTTTGCTACCACATTGGGAGTGAACAAATGCCATGTACAGATCATGGATCTCGGGCACATGTGTGACGCCACCATGAGTTATGAGTGCCCTATGCTAGACGAAGGAGTGGAACCAGATGATGTCGATTGCTGGTGCAACACGACATCAACTTGGGTTGTGTACGGAACCTGTCATCACAAAAAAGGTGAAGCACGGCGATCTAGAAGAGCTGTGACGCTCCCTTCTCACTCCACAAGGAAGTTGCAAACGCGGTCGCAGACCTGGTTAGAATCAAGAGAATACACGAAGCACTTGATCAAGGTTGAAAACTGGATATTCAGGAACCCCGGGTTTGCGCTAGCGGCTGTTGCCATTGCTTGGCTTCTGGGAAGCTCGACGAGCCAAAAAGTCATATACTTGGTCATGATATTGTTGATTGCCCCGGCATACAGCATCAGGTGCATAGGAGTTAGCAATAGAGATTTCGTAGAGGGCATGTCAGGTGGGACCTGGGTTGATGTCGTCTTGGAACATGGGGGTTGCGTCACCGTGATGGCACAGGACAAGCCAACAGTTGACATCGAGTTGGTCACGACAACGGTTAGCAACATGGCCGAGGTAAGATCCTACTGCTATGAGGCATCAATATCGGACATGGCCTCGGACAGCCGCTGTCCAACACAAGGTGAAGCCTACCTTGACAAGCAATCAGACACTCAATATGTCTGCAAAAGAACATTAGTGGACAGAGGTTGGGGAAACGGTTGTGGACTTTTTGGCAAAGGGAGCTTGGTGACATGTGCCAAGTTTACGTGTTCTAAGAAGATGACCGGGAAGAGCATTCAGCCGGAGAACTTGGAGTACCGGATAATGCTATCAGTGCATGGATCCCAGCACAGTGGGATGNNNNNNNNNNNNNNNGGACATGAAACTGACGAAAACAGAGCAAAGGTCGAGGTCACACCCAATTCACCAAGAGCAGAAGCAACCTTGGGAGGTTTTGGAAGCTTGGGACTTGACTGTGAACCAAGGACAGGCCTTGACTTCTCAGATCTGTATTATCTGACCATGAACAACAAGCATCGGTTGGTGCGCAAGGAGTGGTTTCATGACATCCCATTACCTTGGCATGCTGGTGCAGACACTGGAACTCCACACTGGAACAACAAAGAGGCATTGGTGGAGTTCAAGGACGCTCACGCCAAGAGGCAAACTGTTGTGGTTCTGGGGAGCCAAGAGGGAGCTGTTCACACGGCCCTCGCTGGAGCTCTGGAGGCTGAGATGGATGGTGCAAAGGGAAGGCTATTCTCTGGCCATTTGAAATGCCGCCTAAAAATGGACAAGCTTAGGTTGAAGGGTGTGTCATATTCCCTGTGTACTGCAGCGTTCACATTTACCAAGGTCCCAGCTGAAACATTGCATGGAACAGTTACAGTGGAGGTGCAGTATGCAGGGACAGATGGACCTTGCAAGGTCCCAGCCCAGATGGCGGTGGACATGCAGACTCTGACCCCAGTTGGAAGGCTGATAACCGCCAACCCTGTGATCACTGAAAGTACTGAGAATTCAAAGATGATGTTGGAACTTGACCCACCATTTGGGGATTCTTACATTGTCATAGGAGTCGGGGACAAGAAAATCACCCATCACTGGCTCAAGAAAGGGAGCTCCATCGGAAAGGCATTTGAAGCCACTGTGAGAGGTGCCAAGAGAATGGCAGTCTTGGGGGACACAGCCTGGGACTTTGGATCAGTTGGAGGTGTGTTTAATTCATTGGGTAAGGGTGTTCATCAGATCTTTGGAGCAGCTTTCAAATCACTGTTTGGAGGAATGTCCTGGTTCTCACAGATCCTCATAGGCACACTGTTGGTGTGGTTGGGTCTGAACACAAAGAATGGATCTATCTCCCTCACATGCTTAGCCCTGGGGGGAGTGATGATTTTCCTTTCCACGGCTGTTTCTGCTGATGTTGGGTGCTCGGTGGACTTCTCAAAAAAGGAAACGAGATGTGGCACGGGGGTGTTCGTCTACAATGACGTTGAAGCCTGGAGGGACCGTTACAAGTATCATCCTGACTCCCCCCGCAGATTGGCAGCAGCTGTTAAGCAGGCTTGGGAAGAGGGGATTTGTGGGATCTCCTCCGTTTCGAGAATGGAAAACATCATGTGGAAATCAGTGGAAGGGGAGCTTAATGCAATCCTAGAGGAGAATGGAGTCCAACTGACAGTTGTAGTGGGGTCTGTAAAAAACCCCATGTGGAGAGGTCCACAAAGATTGCCAGTGCCCGTGAATGAGCTGCCCCATGGCTGGAAAGCCTGGGGGAAATCGTACTTTGTCAGAGCGGCAAAGACCAACAACAGTTTTGTTGTCGACGGAGACACACTGAAGGAGTGTCCGCTCAAACATAGAGCATGGAATAGCTTCCTTGTGGAGGATCACGGGTTTGGGATTTTCCACACCAGTGTTTGGCTGAAGGTCAGAGAGGACTACTCATTAGAGTGTGACCCAGCCGTCATAGGAACAGCTGTCAAGGGAAAGGAAGCTGCACACAGTGATCTATGCTATTGGATTGAGAGTGAAAAGAATGACACATGGAGGCTGAGGAGGGCCCACCTGATTGAGATGAAAACATGTGAATGGCCAAAGTCTCACACATTGTGGACAGATGGAGTAGAAGAAAGTGATCTTATCATACCCAAGTCTTTAGCTGGTCCACTCAGCCACCACAACACCAGAGAGGGTTACAGAACCCAAGTGAAAGGGCCATGGCACAGTGAAGAGCTTGAAATCCGGTTTGAGGAATGTCCAGGCACCAAGGTTTACGTGGAGGAGACATGCGGAACTAGAGGACCATCTCTGAGATCAACTACTGCAAGTGGAAGGGTCATTGAGGAATGGTGCTGTAGGGAATGCACAATGCCCCCACTATCGTTTCGAGCAAAAGACGGCTGCTGGTATGGAATGGAGATAAGGCCCAGGAAAGAACCAGAGAGCAACTTAGTGAGGTCAATGGTGACAGCGGGGTCAACCGATCATATGGACCACTTCTCTCTTGGAGTGCTTGTGATTCTACTCATGGTGCAGGAGGGGTTGAAGAAGAGAATGACCACAAAGATCATCATGAGCACATCAATGGCAGTGCTGGTAGCCATGGTCTTGGGAGGATTCTCAATGAGTGATCTGGCTAAGCTTGTGATCCTGATGGGTGCCACTTTCGCAGAAATGAACACTGGAGGAGATGTGGCTCACTTGGCATTGGTTGCGGCATTTAAAGTCAGACCAGCCTTGTTGGTTTCCTTTATCTTCAGAGCCAACTGGACACCCCGTGAGAGCATGCTGCTAGCCCTGGCTTCGTGTCTCCTGCAGACTGCGATCTCCGCTCTTGAAGGCGAGCTGATGGTCCTCGTTAATGGATTTGCTTTGGCCTGGTTGGCAATACGAGCAATGGCCGTGCCACGCACTGACAATATCGCTCTAGCAACTCTGGCCGCTTTAACACCATTGGCCAGAGGCACACTGCTCGTGGCATGGAGAGCGGGCCTCGCCACTTGTGGAGGGTTCATGCTTCTTTCCCTGAAAGGGAAAGGTAGTGTGAAGAAGAACCTGCCTTTTGTCATGGCCTTGGGGTTGGCCGCTGTGAGGATAGTGGACCCCATCAATGTGGTAGGACTACTGTTACTCACAAGGAGTGGAAAACGGAGCTGGCCCCCCAGTGAAGTGCTTACAGCTGTTGGCCTGATATGTGCACTGGCCGGAGGGTTTGCCAAGGCAGACATAGAGATGGCTGGGCCCATGGCTGCAGTAGGCCTGCTAATTGTCAGTTATGTGGTCTCGGGAAAGAGTGTGGACATGTACATCGAAAGAGCAGGTGATATCACATGGGAAAAAGACGCGGAAGTCACTGGAAACAGTCCCCGGCTTGACGTGGCACTGGATGAGAGTGGTGATTTCTCCTTGGTAGAGGAGGATGGCCCACCCATGAGAGAGATCATACTCAAGGTGGTCCTGATGGCCATCTGTGGCATGAACCCAATAGCCATACCCTTCGCTGCAGGAGCGTGGTATGTGTATGTAAAGACTGGGAAAAGGAGTGGTGCCCTCTGGGACGTGCCTGCTCCCAAAGAAGTGAAAAAAGGAGAGACTACAGATGGAGTGTACAGAGTGATGACTCGCAGACTGCTGGGTTCAACACAGGTTGGAGTGGGGGTCATGCAAGAGGGAGTCTTCCACACCATGTGGCACGTCACAAAAGGAGCTGCATTGAGGAGCGGTGAAGGAAGACTTGATCCATATTGGGGGGACGTCAAGCAGGACTTGGTGTCATATTGTGGGCCGTGGAAGTTGGACGCAACCTGGGATGGACTAAGTGAAGTGCAGCTTTTGGCCGTACCCCCCGGAGAGAGGGCTAGAAACATTCAGACTCTGCCTGGAATATTCAAGACAAAGGATGGGGACATCGGAGCGGTTGCTCTAGACTACCCCGCAGGAACCTCAGGATCTCCAATCCTAGACAAATGCGGAAGAGTGATAGGACTTTATGGCAATGGGGTTGTGATCAAGAATGGAAGCTATGTCAGTGCAATAACCCAGGGAAAAAGGGAGGAGGAGGCCCCAGTTGAGTGCTTTGAACCCTCGATGCTGAGGAAGAAGCAGCTAACAGTCTTGGATCTGCATCCAGGAGCCGGAAAAACCAGGAGGGTTCTTCCTGAAATAGTCCGTGAAGCCATAAAAAAGAGACTCCGGACAGTGATCTTGGCACCAACTAGGGTTGTCGCTGCTGAGATGGAGGAGGCCTTGAGAGGACTTCCGGTGCGTTACATGACAACAGCAGTCAACGTCACCCATTCTGGGACAGAAATCGTTGATTTGATGTGCCATGCCACTTTCACTTCACGCTTACTACAACCCATCAGAGTCCCTAATTACAATCTCAACATCATGGATGAAGCCCACTTCACAGACCCCTCAAGTATAGCTGCAAGAGGATACATATCAACAAGGGTTGAAATGGGCGAGGCGGCTGCCATTTTTATGACTGCCACACCACCAGGAACCCGTGATGCGTTTCCTGACTCTAACTCACCAATCATGGACACAGAAGTGGAAGTCCCAGAGAGAGCCTGGAGCTCAGGCTTTGATTGGGTGACAGACCATTCTGGGAAAACAATTTGGTTTGTTCCAAGTGTGAGAAACGGAAATGAAATCGCAGCCTGTCTGACAAAGGCTGGAAAGCGGGTTATACAGCTCAGCAGGAAGACTTTTGAGACAGAGTTTCAGAAGACAAAAAATCAAGAGTGGGACTTTGTCATAACAACTGACATTTCAGAGATGGGTGCCAATTTCAAGGCTGACCGGGTCATCGATTCCAGGAGATGCCTAAAGCCAGTCATACTTGATGGTGAGAGAGTCATCCTGGCTGGGCCCATGCCTGTCACGCATGCCAGTGCTGCTCAGAGGAGAGGACGTATAGGCAGGAACCCCAACAAACCCGGAGATGAGTACATGTATGGAGGTGGGTGTGCAGAGACTGATGAAGACCATGCACACTGGCTTGAAGCAAGAATGCTTCTTGACAACATTTACCTCCAAGATGGCCTCATAGCCTCGCTCTATCGGCCTGAGGCCGACAAGGTAGCCGCCATTGAGGGAGAGTTCAAGCTGAGGACAGAGCAAAGGAAGACCTTTGTGGAACTCATGAAGAGAGGAGACCTTCCCGTTTGGCTGGCCTATCAAGTAGCATCTGCCGGAATAACTTACACAGACAGAAGATGGTGCTTTGATGGCACTACCAACAACACCATAATGGAAGACAGTGTACCAGCAGAGGTGTGGACCAAGTATGGAGAGAAGAGAGTGCTTAAACCGAGGTGGATGGATGCTAGGGTCTGTTCAGATCATGCGGCTCTGAAGTCGTTCAAAGAATTTGCCGCTGGGAAGAGAGGAGCGGCTTTGGGAGTAATGGATGCCCTGGGAACATTGCCAGGACACATGACAGAGAGGTTTCAGGAAGCCATTGACAATCTTGCTGTGCTCATGCGAGCAGAGACTGGAAGTAGGCCCTACAAAGCAGCGGCAGCTCAACTGCCGGAGACCCTAGAGACCATCATGCTTTTGGGCTTATTGGGAACAGTTTCGCTAGGAATCTTCTTTGTCTTGATGCGGAACAAGGGCATCGGGAAGATGGGCTTCGGAATGGTAACCCTTGGGGCCAGCGCATGGCTCATGTGGCTCTCGGAAATTGAACCAGCCAGAATCGCATGTGTCCTCATTGTCGTGTTTTTGTTACTGGTGGTGCTCATACCCGAGCCAGAGAAGCAAAGATCTCCCCAGGATAATCAAATGGCAATCATCATCATGGTGGCAGTTGGCCTTTTGGGTTTGATAACTGCAAATGAACTCGGATGGCTGGAAAGGACAAAAAGTGATATAGCTTATCTGATGGGAAGGAAAGAAGAGGGGACAACCATAGGATTCTCAATGGATATTGATCTGCGGCCAGCCTCTGCCTGGGCTATTTATGCCGCATTGACAACTCTCATCACCCCAGCCGTCCAACATGCGGTAACCACCTCATACAACAACTACTCCCTGATGGCGATGGCCACACAAGCTGGAGTGCTGTTTGGCATGGGCAAAGGGATGCCATTTTATGCATGGG-ACTTTGGAGTCCCGCTGCTAATGATAGGTTGCTACTCACAGTTAACACCCCTGACCCTGATAGTAGCCATCATTCTGCTTGTAGCACACTACATGTACTTGATCCCAGGTTTGCAGGCAGCAGCAGCACGTGCTGCCCAGAAGAGGACAGCAGCTGGCATCATGAAAAATCCCGTTGTGGATGGAATAGTGGTGACTGACATTGACACAATGACAATTGACCCCCAAGTGGAGAAGAAGATGGGACAAGTGTTACTCATAGCAGTAGCTGTCTCCAGTGCTGTGCTGCTGCGGACCGCTGGGGGATGGGGGGAGGCTGGGGCTCTGATCACAGCAGCTACCTCCACCTTATGGGAAGGCTCTCCAAACAAATACTGGAACTCCTCTACAGCCACCTCACTATGCAACATCTTTAGAGGAAGTTATTTGGCAGGGGCTTCCCTTATTTACACAGTGACAAGAAATGCCGGTCTGGTTAAGAGACGTGGAGGTGGAACGGGAGAGACCCTGGGAGAGAAGTGGAAAGCCCGCCTGAACCAGATGTCGGCCCTGGAGTTTTACTCTTACAAAAAGTCAGGCATCACCGAAGTGTGTAGGGAGGAGGCGCGCCGCGCTCTCAAGGATGGAGTGGCCACAGGAGGGCATGCTGTATCCCGGGGAAGCGCAAAGCTTAGATGGTTGGTAGAGAGAGGATACCTGCAGCCCCATGGAAAGGTTGTTGACCTTGGATGTGGCAGAGGAGGCTGGAGTTATTACGCTGCCACTATCCGTAAAGTGCAGGAGGTCAGAGGATACACAAAGGGAGGTCCTGGTCATGAAGAACCCATGCTGGTGCAAAGCTATGGGTGGAACATAGTTCGCCTCAAGAGTGGAGTGGACGTCTTCCACATGGCGGCTGAGCCGTGTGACACTCTGCTGTGTGACATAGGCGAGTCATCATCTAGTCCTGAAGTGGAAGAGACGCGAACACTCAGAGTGCTCTCCATGGTAGAAGACTGGCTTGAGAAAAGACCAGGGGCCTTCTGCATAAAGGTGCTGTGCCCATACACCAGCACAATGATGGAGACCATGGAGCGACTGCAACGTAGGCATGGGGGAGGATTGGTCAGAGTGCCATTGTCCCGCAACTCCACACATGAGATGTATTGGGTCTCTGGAGCCAAAAGTAACATCATAAAGAGTGTGTCCACCACAAGTCAGCTCCTCTTGGGACGCATGGATGGGCCTAGGAGGCCAGTGAAATATGAGGAGGATGTGAACCTCGGCTCAGGCACACGAGCTGTGGCAAGCTGCGCTGAGGCTCCCAACATGAAGATCATTGGTAGGCGCATTGAGAGAATCCGCAATGAACATGCAGAGACATGGTTCTTTGATGAAAACCACCCATACAGGACATGGGCCTACCATGGGAGCTACGAAGCCCCCACGCAGGGGTCAGCGTCATCCCTCGTGAATGGGGTTGTTAGACTCTTGTCAAAGCCCTGGGATGTGGTGACTGGAGTTACAGGAATAGCTATGACTGACACCACGCCATACGGCCAACAAAGAGTCTTCAAGGAAAAAGTGGACACAAGGGTGCCAGATCCCCAAGAAGGCACTCGCCAAGTAATGAACATAGTTTCGTCTTGGCTATGGAAGGAGCTAGGAAAACGCAAGCGACCACGTGTCTGCACCAAAGAAGAGTTCATCAATAAGGTGCGCAGCAATGCAGCACTGGGAGCAATATTTGAAGAGGAAAAAGAATGGAAGACAGCTGTAGAAGCTGTGAATGATCCAAGATTTTGGGCTCTAGTGGACAAGGAAAGAGAACACCACCTGAGAGGAGAGTGTCACAGCTGTGTGTACAACATGATGGGTAAAAGAGAAAAGAAGCAAGGAGAATTCGGGAAAGCAAAAGGCAGCCGCGCAATCTGGTACATGTGGTTGGGAGCCAGATTTTTGGAGTTTGAAGCTCTTGGATTCTTGAACGAGGATCACTGGATGGGAAGAGAAAACTCAGGAGGTGGTGTTGAAGGGCTAGGACTGCAAAGGCTTGGATATATCCTAGAAGAAATGAACCGGGCACCAGGAGGAAAGATGTATGCAGATGACACCGCTGGCTGGGACACCCGTATTAGTAAGTTTGATCTAGAGAATGAAGCCCTGATCACTAACCAGATGGAAGAAGGGCACAGAGCTCTGGCGTTGGCCGTGATTAAATACACATACCAAAACAAAGTGGTGAAGGTTCTCAGACCAGCTGAAGGAGGGAAAACAGTCATGGACATCATCTCAAGACAAGACCAGAGAGGGAGTGGACAAGTTGTTACTTATGCCCTCAACACATTCACCAACCTGGTGGTGCAGCTTATCCGGAACATGGAGGCTGAGGAAGTGCTAGAGATGCATGATCTGTGGTTGTTGAGGAAACCAGAGAAAGTGACCAGATGGTTGCAGAGTAATGGATGGGACAGACTCAAACGGATGGCAGTTAGTGGAGATGACTGCGTTGTAAAGCCAATTGATGATAGGTTTGCACATGCCCTCAGGTTCTTGAATGACATGGGAAAGGTTAGGAAAGACACACAGGAATGGAAGCCCTCGACTGGATGGAGCAATTGGGAGGAGGTCCCGTTCTGTTCTCACCACTTCAACAAGCTGCACCTCAAGGACGGGAGATCCATTGTGGTCCCCTGCCGCCACCAAGATGAACTGATTGGCCGAGCCCGTGTCTCACCAGGGGCAGGATGGAGCATCCGTGAGACTGCTTGCCTTGCAAAATCATATGCACAGATGTGGCAGCTTCTTTATTTCCACAGAAGAGATCTCCGACTGATGGCCAATGCCATTTGTTCGGCTGTGCCAGTTGATTGGGTACCAACTGGGAGAACCACCTGGTCAATCCATGGAAAGGGAGAATGGATGACTACTGAGGACATGCTCATGGTGTGGAATAGAGTGTGGATTGAGGAGAACGACCACATGGAGGACAAGACCCCTGTTCCAAAATGGACAGAAATTCCCTACTTGGGGAAAAGGGAGTCCCTATGGTGTGGATACATTATAGGGCACAGACCTCGCACCACTTGGGCTGAGAACATCAAAGACACAGTCAACATGGTGCTTAGGCTCATAGGTGATGAAGAAAAGTACATGGACTACCTATCCACCCAAGTACGCTACTTGGGTGAGGAAGGGTCCACACCTGGAGTGCTGTAA

>ENA|BAP47441|BAP47441.1 Zika virus polyprotein

ATGAAAAACCC-AAAGAAGAAATCCGGAGGATTCCGGATTGTCAATATGCTAAAACGCGGAGTAGCCCGTGTAAACCCCTTGGGAGGTTTGAAGAGGTTGCCAGCCGGACTTCTGCTGGGTCATGGACCCATCAGAATGGTTTTGGCGATACTAGCCTTTTTGAGATTTACAGCAATCAAGCCATCACTGGGCCTTATCAACAGATGGGGTTCCGTGGGGAAAAAAGAGGCTATGGAAATAATAAAGAAGTTCAAGAAAGATCTTGCTGCCATGTTGAGAATAATCAATGCTAGGAAAGAGAGGAAGAGACGTGGCGCAGACACCAGCATCGGAATCATTGGCCTCCTGCTGACTACAGCCATGGCAGCAGAGATCACTAGACGCGGGAGTGCATACTACATGTACTTGGATAGGAGCGATGCCGGGAAGGCCATTTCGTTTGCTACCACATTGGGAGTGAACAAGTGCCACGTACAGATCATGGACCTCGGGCACATGTGTGACGCCACCATGAGTTATGAGTGCCCTATGCTGGATGAGGGAGTGGAACCAGATGATGTCGATTGCTGGTGCAACACGACATCAACTTGGGTTGTGTACGGAACCTGTCATCACAAAAAAGGTGAGGCACGGCGATCTAGAAGAGCCGTGACGCTCCCTTCTCACTCTACAAGGAAGTTGCAAACGCGGTCGCAGACCTGGTTAGAATCAAGAGAATACACGAAGCACTTGATCAAGGTTGAAAACTGGATATTCAGGAACCCCGGGTTTGCGCTAGTGGCCGTTGCCATTGCCTGGCTTTTGGGAAGCTCGACGAGCCAAAAAGTCATATACTTGGTCATGATACTGCTGATTGCCCCGGCATACAGTATCAGGTGCATTGGAGTCAGCAATAGAGACTTCGTGGAGGGCATGTCAGGTGGGACCTGGGTTGATGTTGTCTTGGAACATGGAGGCTGCGTTACCGTGATGGCACAGGACAAGCCAACAGTTGACATAGAGTTGGTCACGACGACGGTTAGTAACATGGCCGAGGTAAGATCCTATTGCTACGAGGCATCGATATCGGACATGGCTTCGGACAGTCGTTGCCCAACACAAGGTGAAGCCTACCTTGACAAGCAATCAGACACTCAATATGTCTGCAAAAGAACATTAGTGGACAGAGGTTGGGGAAACGGTTGTGGACTTTTTGGCAAAGGGAGCTTGGTGACATGTGCCAAGTTTACGTGTTCTAAGAAGATGACCGGGAAGAGCATTCAACCGGAAAATCTGGAGTATCGGATAATGCTATCAGTGCATGGCTCCCAGCATAGCGGGATGACTGTCAATGATATAGGATATGAAACTGACGAAAATAGAGCGAAAGTCGAGGTTACGCCTAATTCACCAAGAGCGGAAGCAACCTTGGGAGGCTTTGGAAGCTTAGGACTTGACTGTGAACCAAGGACAGGCCTTGACTTTTCAGATCTGTATTACCTGACCATGAACAATAAGCATTGGTTGGTGCACAAAGAGTGGTTTCATGACATCCCATTGCCTTGGCATGCTGGGGCAGACACTGGAACTCCACACTGGAACAACAAAGAGGCATTGGTAGAATTCAAGGATGCCCACGCCAAGAGGCAAACCGTCGTCGTTCTGGGGAGCCAGGAAGGAGCCGTTCACACGGCTCTCGCTGGAGCTCTAGAGGCTGAGATGGATGGTGCAAAGGGAAAGCTGTTCTCTGGCCATTTGAAATGCCGCCTAAAAATGGACAAGCTTAGATTGAAGGGCGTGTCATATTCCTTGTGCACTGCGGCATTCACATTCACCAAGGTCCCAGCTGAAACACTGCATGGAACAGTCACAGTGGAGGTGCAGTATGCAGGGACAGATGGACCCTGCAAGATCCCAGTCCAGATGGCGGTGGACATGCAGACCCTGACCCCAGTTGGAAGGCTGATAACCGCCAACCCCGTGATTACTGAAAGCACTGAGAACTCAAAGATGATGTTGGAGCTTGACCCACCATTTGGGGATTCTTACATTGTCATAGGAGTTGGGGACAAGAAAATCACCCACCACTGGCATAGGAGTGGTAGCACCATCGGAAAGGCATTTGAGGCCACTGTGAGAGGCGCCAAGAGAATGGCAGTCCTGGGGGATACAGCCTGGGACTTCGGATCAGTCGGGGGTGTGTTCAACTCACTGGGTAAGGGCATTCACCAGATTTTTGGAGCAGCCTTCAAATCACTGTTTGGAGGAATGTCCTGGTTCTCACAGATCCTCATAGGCACGCTGCTAGTGTGGTTAGGTTTGAACACAAAGAATGGATCTATCTCCCTCACATGCTTGGCCCTGGGGGGAGTGATGATCTTCCTCTCCACGGCTGTTTCTGCTGACGTGGGGTGCTCAGTGGACTTCTCAAAAAAGGAAACGAGATGTGGCACGGGGGTATTCATCTATAATGATGTTGAAGCCTGGAGGGACCGGTACAAGTACCATCCTGACTCCCCCCGCAGATTGGCAGCAGCAGTCAAGCAGGCCTGGGAAGAGGGGATCTGTGGGATCTCATCCGTTTCAAGAATGGAAAACATCATGTGGAAATCAGTAGAAGGGGAGCTCAATGCTATCCTAGAGGAGAATGGAGTTCAACTGACAGTTGTTGTGGGATCTGTAAAAAACCCCATGTGGAGAGGTCCACAAAGATTGCCAGTGCCTGTGAATGAGCTGCCCCATGGCTGGAAAGCCTGGGGGAAATCGTATTTTGTTAGGGCGGCAAAGACCAACAACAGTTTTGTTGTCGACGGTGACACACTGAAGGAATGTCCGCTTGAGCACAGAGCATGGAATAGTTTTCTTGTGGAGGATCACGGGTTTGGAGTCTTCCACACCAGTGTCTGGCTTAAGGTCAGAGAAGATTACTCATTAGAATGTGACCCAGCCGTCATAGGAACAGCTGTTAAGGGAAGGGAGGCCGCGCACAGTGATCTGGGCTATTGGATTGAAAGTGAAAAGAATGACACATGGAGGCTGAAGAGGGCCCACCTGATTGAGATGAAAACATGTGAATGGCCAAAGTCTCACACATTGTGGACAGATGGAGTAGAAGAAAGTGATCTTATCATACCCAAGTCTTTAGCTGGTCCACTCAGCCACCACAACACCAGAGAGGGTTACAGAACCCAAGTGAAAGGGCCATGGCACAGTGAAGAGCTTGAAATCCGGTTTGAGGAATGTCCAGGCACCAAGGTTTACGTGGAGGAGACATGCGGAACTAGAGGACCATCTCTGAGATCAACTACTGCAAGTGGAAGGGTCATTGAGGAATGGTGCTGTAGGGAATGCACAATGCCCCCACTATCGTTTCGAGCAAAAGACGGCTGCTGGTATGGAATGGAGATAAGGCCCAGGAAAGAACCAGAGAGCAACTTAGTGAGGTCAATGGTGACAGCGGGGTCAACCGATCATATGGACCACTTCTCTCTTGGAGTGCTTGTGATTCTACTCATGGTGCAGGAGGGGTTGAAGAAGAGAATGACCACAAAGATCATCATGAGCACATCAATGGCAGTGCTGGTAGTCATGATCTTGGGAGGATTTTCAATGAGTGACCTGGCCAAGCTTGTGATCCTGATGGGTGCTACTTTCGCAGAAATGAACACTGGAGGAGATGTAGCTCACTTGGCATTGGTAGCGGCATTTAAAGTCAGACCAGCCTTGCTGGTCTCCTTCATTTTCAGAGCCAATTGGACACCCCGTGAGAGCATGCTGCTAGCCCTGGCTTCGTGTCTTCTGCAAACTGCGATCTCTGCTCTTGAAGGTGACTTGATGGTCCTCATTAATGGATTTGCTTTGGCCTGGTTGGCAATTCGAGCAATGGCCGTGCCACGCACTGACAACATCGCTCTACCAATCTTGGCTGCTCTAACACCACTAGCTCGAGGCACACTGCTCGTGGCATGGAGAGCGGGCCTGGCTACTTGTGGAGGGATCATGCTCCTCTCCCTGAAAGGGAAAGGTAGTGTGAAGAAGAACCTGCCATTTGTCATGGCCCTGGGATTGACAGCTGTGAGGGTAGTAGACCCTATTAATGTGGTAGGACTACTGTTACTCACAAGGAGTGGGAAGCGGAGCTGGCCCCCTAGTGAAGTTCTCACAGCCGTTGGCCTGATATGTGCACTGGCCGGAGGGTTTGCCAAGGCAGACATTGAGATGGCTGGACCCATGGCTGCAGTAGGCTTGCTAATTGTCAGCTATGTGGTCTCGGGAAAGAGTGTGGACATGTACATTGAAAGAGCAGGTGACATCACATGGGAAAAGGACGCGGAAGTCACTGGAAACAGTCCTCGGCTTGACGTGGCACTGGATGAGAGTGGTGATTTCTCCTTGGTAGAGGAAGATGGTCCACCCATGAGAGAGATCATACTTAAGGTGGTCCTGATGGCCATCTGTGGCATGAACCCAATAGCTATACCTTTTGCTGCAGGAGCGTGGTATGTGTATGTGAAGACTGGGAAAAGGAGTGGCGCCCTCTGGGACGTGCCTGCTCCCAAAGAAGTGAAGAAAGGAGAGACCACAGATGGAGTGTACAGAGTGATGACTCGCAGACTGCTAGGTTCAACACAGGTTGGAGTGGGAGTCATGCAAGAGGGAGTCTTCCACACCATGTGGCACGTTACAAAAGGAGCCGCACTGAGGAGCGGTGAGGGAAGACTTGATCCATACTGGGGGGATGTCAAGCAGGACTTGGTGTCATACTGTGGGCCTTGGAAGTTGGATGCAGCTTGGGATGGACTCAGCGAGGTACAGCTTTTGGCCGTACCTCCCGGAGAGAGGGCCAGAAACATTCAGACCCTGCCTGGAATATTCAAGACAAAGGACGGGGACATCGGAGCAGTTGCTCTGGACTACCCTGCAGGGACCTCAGGATCTCCGATCCTAGACAAATGTGGAAGAGTGATAGGACTCTATGGCAATGGGGTTGTGATCAAGAATGGAAGCTATGTTAGTGCTATAACCCAGGGAAAGAGGGAGGAGGAGACTCCGGTTGAATGTTTCGAACCCTCGATGCTGAAGAAGAAGCAGCTAACTGTCTTGGATCTGCATCCAGGAGCCGGAAAAACCAGGAGAGTTCTTCCTGAAATAGTCCGTGAAGCCATAAAAAAGAGACTCCGGACAGTGATCTTGGCACCAACTAGGGTTGTCGCTGCTGAGATGGAGGAGGCCTTGAGAGGACTTCCGGTGCGTTACATGACAACAGCAGTCAACGTCACCCATTCTGGGACAGAAATCGTTGATTTGATGTGCCATGCCACTTTCACTTCACGCTTACTACAACCCATCAGAGTCCCTAATTACAATCTCTACATCATGGATGAAGCCCACTTCACAGACCCCTCAAGTATAGCTGCAAGAGGATATATATCAACAAGGGTTGAAATGGGCGAGGCGGCTGCCATTTTTATGACTGCCACACCACCAGGAACCCGTGATGCGTTTCCTGACTCTAACTCACCAATCATGGACACAGAAGTGGAAGTCCCAGAGAGAGCCTGGAGCTCAGGCTTTGATTGGGTGACAGACCATTCTGGGAAAACAGTTTGGTTCGTTCCAAGCGTGAGAAACGGAAATGAAATCGCAGCCTGTCTGACAAAGGCTGGAAAGCGGGTCATACAGCTCAGCAGGAAGACTTTTGAGACAGAATTTCAGAAAACAAAAAATCAAGAGTGGGACTTTGTCATAACAACTGACATCTCAGAGATGGGCGCCAACTTCAAGGCTGACCGGGTCATAGACTCTAGGAGATGCCTAAAACCAGTCATACTTGATGGTGAGAGAGTCATCTTGGCTGGGCCCATGCCTGTCACGCATGCTAGTGCTGCTCAGAGGAGAGGACGTATAGGCAGGAACCCTAACAAACCTGGAGATGAGTACATGTATGGAGGTGGGTGTGCAGAGACTGATGAAGGCCATGCACACTGGCTTGAAGCAAGAATGCTTCTTGACAACATCTACCTCCAGGATGGCCTCATAGCCTCGCTCTATCGGCCTGAGGCCGATAAGGTAGCCGCCATTGAGGGAGAGTTTAAGCTGAGGACAGAGCAAAGGAAGACCTTCGTGGAACTCATGAAGAGAGGAGACCTTCCCGTCTGGCTAGCCTATCAGGTTGCATCTGCCGGAATAACTTACACAGACAGAAGATGGTGCTTTGATGGCACAACCAACAACACCATAATGGAAGACAGCGTACCAGCAGAGGTGTGGACAAAGTATGGAGAGAAGAGAGTGCTCAAACCGAGATGGATGGATGCTAGGGTCTGTTCAGACCATGCGGCCCTGAAGTCGTTCAAAGAATTCGCCGCTGGAAAAAGAGGAGCGGCTTTGGGAGTAATGGAGGCCCTGGGAACACTGCCAGGACACATGACAGAGAGGTTTCAGGAAGCCATTGACAACCTCGCCGTGCTCATGCGAGCAGAGACTGGAAGCAGGCCTTATAAGGCAGCGGCAGCCCAACTGCCGGAGACCCTAGAGACCATTATGCTCTTAGGTTTGCTGGGAACAGTTTCACTGGGGATCTTCTTCGTCTTGATGCGGAATAAGGGCATCGGGAAGATGGGCTTTGGAATGGTAACCCTTGGGGCCAGTGCATGGCTCATGTGGCTTTCGGAAATTGAACCAGCCAGAATTGCATGTGTCCTCATTGTTGTGTTTTTATTACTGGTGGTGCTCATACCCGAGCCAGAGAAGCAAAGATCTCCCCAAGATAACCAGATGGCAATTATCATCATGGTGGCAGTGGGCCTTCTAGGTTTGATAACTGCAAACGAACTTGGATGGCTGGAAAGAACAAAAAATGACATAGCTCATCTAATGGGAAGGAGAGAAGAAGGAGCAACCATGGGATTCTCAATGGACATTGATCTGCGGCCAGCCTCCGCCTGGGCTATCTATGCCGCATTGACAACTCTCATCACCCCAGCTGTCCAACATGCGGTAACCACTTCATACAACAACTACTCCTTAATGGCGATGGCCACACAAGCTGGAGTGCTGTTTGGCATGGGCAAAGGGATGCCATTTTATGCATGGG-ACCTTGGAGTCCCGCTGCTAATGATGGGTTGCTATTCACAATTAACACCCCTGACTCTGATAGTAGCTATCATTCTGCTTGTGGCGCACTACATGTACTTGATCCCAGGCCTACAAGCGGCAGCAGCGCGTGCTGCCCAGAAAAGGACAGCAGCTGGCATCATGAAGAATCCCGTTGTGGATGGAATAGTGGTAACTGACATTGACACAATGACAATAGACCCCCAGGTGGAGAAGAAGATGGGACAAGTGTTACTCATAGCAGTAGCCATCTCCAGTGCTGTGCTGCTGCGGACCGCCTGGGGATGGGGGGAGGCTGGAGCTCTGATCACAGCAGCGACCTCCACCTTGTGGGAAGGCTCTCCAAACAAATACTGGAACTCCTCTACAGCCACCTCACTGTGCAACATCTTCAGAGGAAGCTATCTGGCAGGAGCTTCCCTTATCTATACAGTGACGAGAAACGCTGGCCTGGTTAAGAGACGTGGAGGTGGGACGGGAGAGACTCTGGGAGAGAAGTGGAAAGCTCGTCTGAATCAGATGTCGGCCCTGGAGTTCTACTCTTATAAAAAGTCAGGTATCACTGAAGTGTGTAGAGAGGAGGCTCGCCGTGCCCTCAAGGATGGAGTGGCCACAGGAGGACATGCCGTATCCCGGGGAAGTGCAAAGCTCAGATGGTTGGTGGAGAGAGGATATCTGCAGCCCTATGGGAAGGTTGTTGACCTCGGATGTGGCAGAGGGGGCTGGAGCTATTATGCCGCCACCATCCGCAAAGTGCAGGAGGTGAGAGGATACACAAAGGGAGGTCCCGGTCATGAAGAACCCATGCTGGTGCAAAGCTATGGGTGGAACATAGTTCGTCTCAAGAGTGGAGTGGACGTCTTCCACATGGCGGCTGAGCCGTGTGACACTCTGCTGTGTGACATAGGTGAGTCATCATCTAGTCCTGAAGTGGAAGAGACACGAACACTCAGAGTGCTCTCTATGGTGGGGGACTGGCTTGAAAAAAGACCAGGGGCCTTCTGTATAAAGGTGCTGTGCCCATACACCAGCACTATGATGGAAACCATGGAGCGACTGCAACGTAGGCATGGGGGAGGATTAGTCAGAGTGCCATTGTCTCGCAACTCCACACATGAGATGTACTGGGTCTCTGGGGCAAAGAGCAACATCATAAAAAGTGTGTCCACCACAAGTCAGCTCCTCCTGGGACGCATGGATGGCCCCAGGAGGCCAGTGAAATATGAGGAGGATGTGAACCTCGGCTCGGGTACACGAGCTGTGGCAAGCTGTGCTGAGGCTCCTAACATGAAAATCATCGGCAGGCGCATTGAGAGAATCCGCAATGAACATGCAGAAACATGGTTTCTTGATGAAAACCACCCATACAGGACATGGGCCTACCATGGGAGCTACGAAGCCCCCACGCAAGGATCAGCGTCTTCCCTCGTGAACGGGGTTGTTAGACTCCTGTCAAAGCCTTGGGACGTGGTGACTGGAGTTACAGGAATAGCCATGACTGACACCACACCATACGGCCAACAAAGAGTCTTCAAAGAAAAAGTGGACACCAGGGTGCCAGATCCCCAAGAAGGCACTCGCCAGGTAATGAACATAGTCTCTTCCTGGCTGTGGAAGGAGCTGGGGAAACGCAAGCGGCCACGCGTCTGCACCAAAGAAGAGTTTATCAACAAGGTGCGCAGCAATGCAGCACTGGGAGCAATATTTGAAGAGGAAAAAGAATGGAAGACGGCTGTGGAAGCTGTGAATGATCCAAGGTTTTGGGCCCTAGTGGATAGGGAGAGAGAACACCACCTGAGAGGAGAGTGTCACAGCTGTGTGTACAACATGATGGGAAAAAGAGAAAAGAAGCAAGGAGAGTTCGGGAAAGCAAAAGGTAGCCGCGCCATCTGGTACATGTGGTTGGGAGCCAGATTCTTGGAGTTTGAAGCCCTTGGATTCTTGAACGAGGACCATTGGATGGGAAGAGAAAACTCAGGAGGTGGAGTCGAAGGGTTAGGATTGCAAAGACTTGGATACATTCTAGAAGAAATGAATCGGGCACCAGGAGGAAAGATGTACGCAGATGACACTGCTGGCTGGGACACCCGCATTAGTAAGTTTGATCTGGAGAATGAAGCTCTGATTACCAACCAAATGGAGGAAGGGCACAGAACTCTGGCGTTGGCCGTGATTAAATACACATACCAAAACAAAGTGGTGAAGGTTCTCAGACCAGCTGAAGGAGGAAAAACAGTTATGGACATCATTTCAAGACAAGACCAGAGAGGGAGTGGACAAGTTGTCACTTATGCTCTCAACACATTCACCAACTTGGTGGTGCAGCTTATCCGGAACATGGAAGCTGAGGAAGTGTTAGAGATGCAAGACTTATGGTTGTTGAGGAAGCCAGAGAAAGTGACCAGATGGTTGCAGAGCAATGGATGGGATAGACTCAAACGAATGGCGGTCAGTGGAGATGACTGCGTTGTGAAGCCAATCGATGATAGGTTTGCACATGCCCTCAGGTTCTTGAATGACATGGGAAAAGTTAGGAAAGACACACAGGAGTGGAAACCCTCGACTGGATGGAGCAATTGGGAAGAAGTCCCGTTCTGCTCCCACCACTTCAACAAGCTGTACCTCAAGGATGGGAGATCCATTGTGGTCCCTTGCCGCCACCAAGATGAACTGATTGGCCGAGCTCGCGTCTCACCAGGGGCAGGATGGAGCATCCGGGAGACTGCCTGTCTTGCAAAATCATATGCGCAGATGTGGCAGCTCCTTTATTTCCACAGAAGAGACCTTCGACTGATGGCTAATGCCATTTGCTCGGCTGTGCCAGTTGACTGGGTACCAACTGGGAGAACCACCTGGTCAATCCATGGAAAGGGAGAATGGATGACCACTGAGGACATGCTCATGGTGTGGAATAGAGTGTGGATTGAGGAGAACGACCATATGGAGGACAAGACTCCTGTAACAAAATGGACAGACATTCCCTATCTAGGAAAAAGGGAGGACTTATGGTGTGGATCCCTTATAGGGCACAGACCCCGCACCACTTGGGCTGAAAACATCAAAGACACAGTCAACATGGTGCGCAGGATCATAGGTGATGAAGAAAAGTACATGGACTATCTATCCACCCAAGTCCGCTACTTGGGTGAGGAAGGGTCCACACCCGGAGTGTTGTAA

>ENA|AHL43501|AHL43501.1 Zika virus polyprotein

ATGAAAAACCC-AAAGAAGAGATCCGGAGGATTCCGGATTGTCAATATGCTAAAACGCGGAGTAGCCCGTGTAAACCCCTTGGGGGGTTTGAAGAGGCTGCCGGCCGGACTTCTGCTGGGCCATGGACCCATCAGAATGGTTTTGGCGATACTAGCCTTTTTGAGATTCACAGCAATCAAGCCATCACTGGGCCTCATCAATAGATGGGGTTCCGTGGGGAAGAAGGAGGCTATGGAAATAATAAAAAAGTTCAAGAAAGACCTTGCTGCCATGTTGAGAATTATCAATGCTAGGAAGGAGAGGAAGAGACGTGGCGCTGACACCAGCATCGGAATCGTCGGCCTCCTGCTGACCACAGCCATGGCAGCCGAGATCACCAGACGTGGGAGTGCATACTACATGTACTTGGACAGGAGCGATGCTGGTAAGGCCATTTCTTTTGCTACCACATTGGGGGTGAACAAATGCCATGTACAGATCATGGACCTCGGGCACATGTGTGACGCCACCATGAGTTATGAGTGCCCCATGCTGGACGAGGGAGTGGAGCCAGATGACGTCGATTGCTGGTGCAACACGACATCAACTTGGGTTGTGTACGGAACCTGTCAACATAAAAAAGGTGAAGCACGACGATCCAGAAGAGCCGTGACGCTTCCTTCTCACTCCACAAGGAAGCTGCAAACGCGATCACAGACTTGGCTAGAATCAAGAGAATACACAAAGCACCTGATCAAGGTTGAGAATTGGATATTCAGGAACCCCGGGTTTGCGCTAGTGGCTGTAGCTATTGCCTGGCTCCTGGGAAGCTCGACGAGCCAAAAAGTCATATACTTGGTAATGATATTGTTGATTGCCCCGGCATACAGCATCAGGTGCATAGGAGTTAGCAATAGAGACTTCGTGGAGGGCATGTCAGGTGGGACCTGGGTTGATGTTGTCTTGGAACATGGGGGTTGCGTCACCGTGATGGCACAGGACAAGCCAACAGTTGACATCGAGTTGGTCACGACAACGGTTAGCAACATGGCCGAGGTAAGATCCTACTGCTATGAGGCATCAATATCGGACATGGCTTCGGACAGCCGCTGTCCAACACAAGGTGAAGCCTACCTTGACAAGCAGTCAGACACTCAATATGTCTGCAAGAGAACATTGGTGGATAGAGGTTGGGGAAATGGGTGTGGACTTTTTGGCAAAGGGAGCTTGGTGACATGTGCCAAGTTTACGTGCTCCAAGAAAATGACAGGCAAGAGCATCCAGCCGGAGAACTTGGAGTACCGGATAATGCTATCAGTGCATGGATCCCAGCACAGTGGGATGATTGTGAATGACATAGGACATGAAACTGACGAAAACAGAGCAAAAGTCGAGGTCACACCCAATTCACCAAGAGCAGAAGCAACCTTGGGAGGTTTTGGAAGCTTGGGACTTGACTGTGAACCAAGGACAGGCCTTGACTTCTCAGATCTGTATTACCTGACCATGAACAATAAGCATTGGTTGGTGCACAAGGAGTGGTTTCATGACATCCCATTACCTTGGCATGCTGGTGCAGACACTGGAACTCCACACTGGAACAACAAAGAGGCATTGGTGGAGTTTAAGGACGCCCACGCCAAGAGGCAAACTGTTGTGGTTCTGGGGAGCCAAGAGGGAGCTGTTCACACGGCCCTCGCTGGAGCTTTGGAGGCTGAGATGGATGGTGCAAAGGGAAGGCTATTCTCTGGCCATTTGAAATGCCGCCTAAAAATGGACAAGCTTAGGTTGAAGGGTGTGTCATATTCCCTGTGTACCGCAGTGTGTACCGCAGCGAAGGTCCCAGCTGAAACATTGCATGGAACAGTCACAGTGGAGGTGCAGTATGCAGGGACAGATGGACCCTGCAAGGTCCCAGCCCAGATGGCGGTAGACATGCAGACCCTGACCCCAGTTGGAAGGCTGATAACCGCCAACCCTGTGATCACTGAAAGTACTGAGAATTCAAAGATGATGTTGGAGCTCGACCCACCATTTGGGGATTCTTACATTGTCATAGGAGTCGGGGACAAGAAAATCACCCATCACTGGCATCGGAGTGGTAGCACCATCGGAAAGGCATTTGAAGCCACTGTGAGAGGTGCCAAGAGAATGGCAGTCTTGGGGGACACAGCCTGGGACTTTGGATCAGTTGGGGGTGTGTTTAATTCATTGGGTAAGGGTATTCACCAGATCTTTGGAGCAGCTTTTAAATCACTGTTTGGAGGAATGTCCTGGTTCTCACAGATCCTCATAGGCACACTGTTGGTGTGGTTAGGTCTGAACACAAAGAATGGATCTATCTCCCTCACATGCTTAGCCCTGGGGGGAGTGATGATCTTCCTTTCCACGGCTGTTTCTGCTGATGTTGGGTGCTCGGTGGACTTCTCAAAAAAGGAAACGAGATGTGGCACGGGGGTGTTCGTCTACAATGACGTTGAAGCCTGGAGGGACCGGTACAAGTACCATCCTGACTCCCCCCGCAGATTGGCAGCAGCTGTCAAGCAGGCTTGGGAAGAGGGGATTTGTGGGATCTCCTCCGTTTCGAGAATGGAAAACATCATGTGGAAATCAGTGGAAGGGGAGCTTAATGCAATCCTAGAGGAGAATGGAGTCCAACTGACAGTTGTAGTGGGGTCTGTAAAAAACCCTATGTGGAGAGGTCCACAAAGATTGCCAGTGCCCGTGAATGAGCTGCCCCATGGCTGGAAAGCCTGGGGGAAATCGTACTTTGTCAGAGCGGCAAAGACCAACAACAGTTTTGTTGTCGACGGTGACACACTGAAGGAGTGTCCGCTCAAACATAGAGCATGGAATAGCTTCCTTGTGGAGGATCACGGGTTTGGGATCTTCCACACCAGTGTTTGGCTGAAGGTCAGAGAGGACTACTCATTAGAGTGTGACCCAGCCGTCATAGGAACAGCTGTCAAGGGAAAGGAAGCTGCACACAGTGATCTAGGCTATTGGATTGAGAGTGAAAAGAATGACACATGGAGGCTGAGGAGGGCTCATCTGATTGAGATGAAAACATGTGAGTGGCCAAAGTCTCACACACTGTGGACAGATGGAGTGGAAGAAAGTGATCTGATCATACCCAAGTCCTTAGCTGGTCCACTCAGCCACCACAACACCAGAGAGGGTTACAGAACTCAAGTGAAAGGGCCATGGCATAGTGAAGAGCTCGAAATCCGGTTTGAGGAGTGCCCAGGAACCAAGGTTCACGTGGAGGAGACATGCGGAACTAGAGGACCATCTCTGAGATCAACTACTGCAAGTGGAAGGGTCATAGAGGAATGGTGCTGTAGGGAATGCACAATGCCTCCACTATCGTTCCGGGCGAAAGACGGCTGCTGGTATGGAATGGAGATAAGGCCCAGAAAGGAACCAGAGAGCAACTTAGTAAGGTCTATGGTGACAGCAGGATCAACCGATCATATGGATCACTTCTCTCTTGGAGTGCTTGTGATTCTACTCATGGTGCAGGAAGGTTTGAAGAAGAGAATGACCACAAAGATCATAATGAGCACATCAATGGCAGTGCTGGTAGCCATGGTCTTGGGAGGATTCTCAATGAGTGACCTGGCTAAGCTTGTGATCCTGATGGGTGCCACTTTCGCAGAAATGAACACTGGAGGAGATGTGGCTCACTTGGCATTGGTTGCGGCATTTAAAGTCAGACCAGCCTTGTTGGTTTCCTTCATCTTCAGAGCCAACTGGACACCCCGTGAGAGCATGCTGCTAGCCCTGGCTTCGTGTCTCCTGCAGACTGCGATCTCCGCTCTTGAAGGCGAGCTGATGGTCCTCGTTAATGGATTTGCTTTGGCCTGGTTGGCAATACGAGCAATGGCCGTGCCACGCACTGACAATATCGCTCTAGCAATTCTGGCTGCTCTAACACCATTAGCCAGAGGCACACTGCTCGTGGCATGGAGAGCGGGCCTCGCCACTTGTGGAGGGTTCATGCTCCTTTCCCTGAAAGGGAAAGGTAGTGTGAAGAAGAACCTGCCTTTTGTCATGGCCTTGGGGTTGACCGCTGTGAGGATAGTGGACCCCATCAATGTGGTAGGACTACTGTTACTCACACGGAGTGGAAAACGGAGCTGGCCCCCTAGTGAAGTGCTTACAGCTGTTGGCCTGATATGTGCACTGGCCGGAGGGTTTGCCAAGGCAGATATAGAGATGGCTGGGCCCATGGCTGCAGTAGGCCTGCTAATTGTCAGTTATGTGGTCTCGGGAAAGAGTGTGGACATGTACATCGAAAGAGCAGGTGATATCACATGGGAAAAAGACNCGGAAGTCACTGGAAACAGTCCTCGGCTTGACGTGGCACTAGATGAGAGTGGTGATTTCTCCTTGGTAGAGGAGGATGGCCCACCCATGAGAGAGATCATACTCAAGGTGGTCCTGATGGCCATCTGTGGCATGAACCCAATAGCCATACCCTTCGCTGCAGGAGCGTGGTATGTGTATGTAAAGACTGGAAAAAGGAGTGGTGCCCTCTGGGACGTGCCTGCTCCCAAAGAAGTGAAAAAAGGAGAGACTACAGATGGAGTGTACAGAGTGATGACTCGCAGACTGCTGGGTTCAACACAGGTTGGAGTGGGAGTCATGCAAGAGGGAGTCTTCCACACCATGTGGCACGTCACAAAAGGAGCCGCATTGAGGAGCGGTGAAGGAAGACTTGATCCATATTGGGGGGACGTCAAGCAGGACTTGGTGTCATATTGTGGGCCGTGGAAGTTGGATGCAGCCTGGGATGGACTAAGTGAAGTGCAGCTTTTGGCCGTACCCCCCGGAGAGAGGGCTAGAAACATTCAGACTCTGCCTGGAATATTCAAGACAAAGGATGGGGACATCGGAGCAGTCGCTCTAGACTACCCCGCAGGAACCTCAGGATCTCCAATCCTAGACAAATGCGGAAGAGTGATAGGACTTTATGGCAATGGGGTTGTGATCAAGAATGGAAGCTATGTTAGTGCCATAACCCAGGGAAAAAGGGAGGAGGAGGCTCCAGTTGAGTGCTTTGAACCCTCGATGCTGAGGAAGAAGCAGCTAACAGTCTTGGATCTGCATCCAGGAGCCGGGAAAACCAGGAGGGTTCTTCCTGAAATAGTCCGTGAAGCCATAAAGAAGAGACTTCGCACATTGATCCTAGCACCAACCAGGGTCGTTGCTGCTGAGATGGAGGAAGCCCTAAGAGGACTTCNNGTGCGTTACATGACAACAGCAGTCAACGTCACCCATTCTGGGACAGAAATCGTTGATTTGATGTGCCATGCCACTTTCACTTCACGCTTACTACAACCCATCAGAGTCCCTAATTACAATCTCAACATCATGGATGAAGCCCACTTCACAGACCCCTCAAGTATAGCTGCAAGAGGATACATATCAACAAGGGTTGAAATGGGCGAGGCGGCTGCCATTTTTATGACTGCCACACCACCAGGAACCCGTGATGCGTTTCCTGACTCTAACTCACCAATCATGGACACAGAAGTGGAAGTCCCAGAGAGAGCCTGGAGCTCAGGCTTTGACTGGGTGACGGACCATTCTGGGAAAACAATTTGGTTTGTTCCAAGTGTGAGATCTGGAGAAGAAAGCGCAGCCTGTCTGACAAAGGCTGGAAAGCGGGTTATACAGCTCAGCAGGAAGACTTTTGAGACAGAGTTTCAGAAGACAAAAAATCAAGAGTGGGACTTTGTCATAACAACTGACATTTCAGAGATGGGTGCCAATTTCAAGGCTGACCGGGTCATAGATTCCAGGAGATGCCTAAAGCCAGTCATACTCGATGGTGAGAGAGTCATCCTGGCTGGGCCCATGCCTGTCACGCATGCCAGTGCTGCTCAGAGGAGAGGACGTATAGGCAGGAACCCCAACAAACCTGGAGATGAGTACATGTATGGAGGTGGGTGTGCAGAGACTGATGAAGACCATGCACACTGGCTTGAAGCAAGAATGCTTCTTGACAACATTTACCTCCAGGATGGCCTCATAGCCTCGCTCTATCGGCCTGAGGCCGACAAGGTAGCCGCCATTGAGGGAGAGTTCAAGCTGAGGACAGAGCAAAGGAAGACCTTTGTGGAACTCATGAAGAGAGGAGACCTTCCCGTTTGGCTGGCCTATCAAGTAGCATCTGCCGGAATAACTTACACAGACAGAAGATGGTGCTTTGATGGCACTACCAACAACACCATAATGGAAGACAGTGTACCAGCAGAGGTGTGGACCAAGTATGGAGAGAAGAGAGTGCTTAAACCGAGGTGGATGGATGCTAGGGTCTGTTCAGATCATGCGGCTTTGAAGTCGTTCAAAGAATTTGCCGCTGGGAAGAGAGGAGCGGCTTTGGGAGTGATGGATGCCCTGGGAACATTGCCAGGACACATGACAGAGAGGTTTCAGGAAGCCATTGACAATCTCGCTGTGCTCATGCGAGCAGAGACTGGAAGTAGGCCCTACAAAGCAGCGGCAGCTCAACTGCCGGAGACCCTAGAGACCATCATGCTTTTGGGCTTATTGGGAACAGTTTCGCTAGGAATCTTCTTTGTCTTGATGCGGAACAAGGGCATCGGGAAGATGGGCTTCGGAATGGTAACCCTTGGGGCCAGCGCATGGCTCATGTGGCTTTCGGAAATTGAACCAGCCAGAATCGCATGTGTCCTCATTGTCGTGTTTTTGTTACTGGTGGTGCTCATACCCGAGCCAGAGAAGCAAAGATCTCCCCAGGATAATCAAATGGCAATCATCATCATGGTGGCAGTTGGCCTTCTGGGTTTGATAACAGCAAATGAACTCGGATGGCTGGAAAGGACAAAAAGTGATATAGCTCATCTAATGGGAAGGAAAGAAGAGGGGACAACCATAGGATTCTCAATGGATATTGATCTGCGGCCAGCCTCCGCCTGGGCTATTTATGCCGCATTGACAACTCTCATCACCCCAGCCGTCCAACATGCGGTAACCACCTCATACAACAACTACTCCCTGATGGCGATGGCCACACAAGCTGGAGTGCTGTTTGGCATGGGCAAAGGGATGCCATTTTATGCATGGG-ACTTTGGAGTCCCGCTGCTAATGATGGGTTGCTACTCACAATTAACACCCCTGACCCTGATAGTGGCCATCATTCTGCTTGTGGCACACTACATGTACTTGATCCCAGGTTTGCAGGCAGCAGCAGCACGTGCTGCCCAGAAGAGGACAGCAGCTGGCATCATGAAGAATCCCGTTGTGGATGGAATAGTGGTGACTGACATTGACACAATGACAATTGACCCCCAAGTGGAGAAGAAGATGGGACAAGTGTTACTCATAGCAGTAGCTGTCTCCAGTGCTGTGCTGCTGCGGACCGCTTGGGGATGGGGGGAGGCTGGGGCTCTGATCACAGCAGCAACCTCCACCTTATGGGAAGGCTCTCCAAACAAATACTGGAACTCCTCCACAGCCACCTCACTGTGCAACATCTTTAGAGGAAGTTATTTGGCAGGGGCTTCCCTTATTTACACAGTGACAAGAAATGCCGGTCTGGTTAAGAGACGTGGAGGTGGAACGGGAGAGACCCTGGGAGAGAAGTGGAAAGCCCGCCTGAACCAGATGTCGGCCTTGGAGTTTTACTCTTACAAAAAGTCAGGCATCACCGAAGTGTGTAGGGAGGAGGCGCGCCGCGCCCTCAAGGATGGAGTGGCCACAGGAGGACATGCTGTATCCCGGGGAAGCGCAAAGCTTAGATGGTTGGTAGAGAGAGGATACCTGCAGCCCCATGGAAAGGTTGTTGACCTTGGATGTGGCAGAGGAGGCTGGAGTTATTACGCTGCCACCATCCGTAAAGTGCAGGAGGTCAGAGGATACACAAAGGGAGGTCCTGGTCATGAAGAACCCATGCTGGTGCAAAGCTATGGGTGGAACATAGTTCGCCTCAAGAGTGGAGTGGACGTCTTCCACATGGCGGCTGAGCCGTGTGACACTCTGCTGTGTGACATCGGCGAGTCATCATCTAGTCCTGAAGTGGAAGAGACGCGAACACTCAGAGTGCTCTCCATGGTGGGAGACTGGCTTGAGAAAAGACCAGGGGCCTTCTGCATAAAGGTGCTGTGCCCATACACCAGCACAATGATGGAGACCATGGAGCGACTGCAACGTAGGCATGGGGGAGGATTGGTCAGAGTGCCATTGTCCCGCAACTCCACACATGAGATGTATTGGGTCTCTGGAGCCAAAAGTAACATCATAAAGAGTGTGTCCACCACAAGTCAGCTCCTCTTGGGACGCATGGATGGGCCTAGGAGGCCAGTGAAATATGAGGAGGATGTGAACCTCGGCTCAGGCACACGAGCTGTGGCAAGCTGCGCTGAGGCTCCCAACATGAAGATCATTGGTAGGCGCATTGAGAGAATCCGCAATGAACATGCAGAGACATGGTTCTTTGATGAAAACCACCCATACAGGACATGGGCCTACCATGGGAGCTACGAAGCCCCCACGCAGGGGTCAGCGTCATCCCTCGTGAACGGGGTTGTTAGACTCTTGTCAAAACCCTGGGATGTGGTGACTGGAGTTACAGGAATAGCTATGACTGACACCACGCCATACGGCCAACAAAGAGTCTTCAAGGAAAAAGTGGACACTAGGGTGCCAGATCCCCAAGAAGGCACTCGCCAAGTAATGAACATGGTCTCGTCTTGGCTATGGAAGGAGCTGGGAAAACGCAAGCGGCCACGTGTCTGCACCAAAGAAGAGTTCATTAATAAGGTGCGCAGCAATGCAGCACTGGGAGCAATATTTGAAGAGGAAAAAGAATGGAAGACAGCTGTAGAAGCTGTGAATGATCCAAGATTTTGGGCTCTAGTGGACAAGGAAAGAGAACACCACCTGAGAGGAGAGTGTCACAGCTGTGTGTACAACATGATGGGTAAAAGAGAAAAGAAGCAAGGAGAATTCGGGAAAGCAAAAGGCAGCCGCGCAATCTGGTACATGTGGTTGGGAGCCAGATTTTTGGAGTTTGAAGCTCTTGGATTCTTGAACGAGGACCACTGGATGGGAAGAGAAAACTCAGGAGGTGGCGTTGAAGGGCTAGGACTGCAAAGGCTTGGATATATCCTAGAAGAAATGAACCGGGCACCAGGAGGAAAGATGTATGCAGATGACACCGCTGGCTGGGACACCCGCATTAGCAAGTTTGATCTAGAGAATGAAGCCCTGATCACTAACCAGATGGAAGAAGGGCACAGAGCTCTGGCGTTGGCCGTGATTAAATACACATACCAAAACAAAGTGGTGAAGGTTCTCAGACCAGCTGAAGGAGGGAAAACAGTCATGGACATCATCTCAAGACAAGACCAGAGAGGGAGCGGACAAGTTGTTACTTATGCTCTCAACACATTCACCAACCTGGTGGTGCAGCTTATCCGGAACATGGAGGCTGAGGAAGTGCTAGAGATGCATGATCTATGGTTGTTGAGGAAGCCAGAGAAAGTGACCAGATGGTTGCAGAGCAATGGATGGGACAGACTCAAACGGATGGCAGTTAGTGGAGATGACTGCGTTGTAAAGCCAATTGATGATAGGTTTGCACATGCCCTCAGGTTCTTGAATGACATGGGAAAGGTTAGGAAAGACACACAGGAATGGAAACCCTCGACTGGATGGAGCAATTGGGAAGAGGTCCCGTTCTGTTCTCACCACTTCAACAAGCTGCACCTCAAGGACGGGAGATCCATTGTGGTCCCCTGCCGCCACCAAGATGAACTGATTGGCCGAGCCCGTGTCTCACCAGGGGCAGGATGGAGCATCCGGGAGACTGCTTGTCTTGCAAAATCATATGCACAGATGTGGCAGCTTCTTTATTTCCACAAAAGAGACCTCCGACTGATGGCCAATGCCATTTGTTCGGCTGTGCCAGTTGACTGGGTACCAACTGGGAGAACCACCTGGTCAATCCATGGAAAGGGAGAATGGATGACTACTGAGGACATGCTCATGGTGTGGAATAGAGTGTGGATTGAGGAGAACGACCACATGGAGGACAAGACCCCTGTAACAAAATGGACAGACATTCCCTATTTGGGGAAAAGGGAGGACTTATGGTGTGGATCCCTTATAGGGCACAGACCTCGCACCACTTGGGCTGAGAACATCAAAGACACAGTCAACATGGTGCGTAGGATCATAGGTGATGAAGAAAAGTACATGGACTACCTATCCACCCAAGTACGCTACTTGGGTGAGGAAGGGTCCACACCTGGAGTGCTGTAA

>ENA|ALU33341|ALU33341.1 Zika virus polyprotein

ATGAAAAACCC-AAAAAAGAAATCCGGAGGATTCCGGATTGTCAATATGCTAAAACGCGGAGTAGCCCGTGTGAGCCCCTTTGGGGGCTTGAAGAGGCTGCCAGCCGGACTTCTGCTGGGTCATGGGCCCATCAGGATGGTCTTGGCAATTCTAGCCTTTTTGAGATTCACGGCAATCAAGCCATCACTGGGTCTCATCAATAGATGGGGTTCAGTGGGGAAAAAAGAGGCTATGGAAATAATAAAGAAGTTCAAGAAAGATCTGGCTGCCATGCTGAGAATAATCAATGCTAGGAAGGAGAAGAAGAGACGGGGCGCAGATACTAGTGTCGGAATTGTTGGCCTCCTGCTGACCACAGCTATGGCAGCGGAGGTCACTAGACGTGGGAGTGCATACTATATGTACTTGGACAGAAACGATGCTGGGGAGGCCATATCTTTTCCAACCACATTGGGGATGAATAAGTGTTATATACAGATCATGGATCTTGGACACATGTGTGATGCCACCATGAGCTATGAATGCCCTATGCTGGATGAGGGGGTGGAACCAGATGACGTCGATTGTTGGTGCAACACGACGTCAACTTGGGTTGTGTACGGAACCTGCCATCACAAAAAAGGTGAAGCACGGAGATCTAGAAGAGCTGTGACGCTCCCCTCCCATTCCACTAGGAAGCTGCAAACGCGGTCGCAAACCTGGTTGGAATCAAGAGAATACACAAAGCACTTGATTAGAGTCGAAAATTGGATATTCAGGAACCCTGGCTTCGCGTTAGCAGCAGCTGCCATCGCTTGGCTTTTGGGAAGCTCAACGAGCCAAAAAGTCATATACTTGGTCATGATACTGCTGATTGCCCCGGCATACAGCATCAGGTGCATAGGAGTCAGCAATAGGGACTTTGTGGAAGGTATGTCAGGTGGGACTTGGGTTGATATTGTCTTGGAACATGGAGGTTGTGTCACCGTAATGGCACAGGACAAACCGACTGTCGACATAGAGCTGGTTACAACAACAGTCAGCAACATGGCGGAGGTAAGATCCTACTGCTATGAGGCATCAATATCAGACATGGCTTCGGACAGCCGCTGCCCAACACAAGGTGAAGCCTACCTTGACAAGCAATCAGACACTCAATATGTCTGCAAAAGAACGTTAGTGGACAGAGGCTGGGGAAATGGATGTGGACTTTTTGGCAAAGGGAGTCTGGTGACATGCGCTAAGTTTGCATGCTCCAAGAAAATGACCGGGAAGAGCATCCAGCCAGAGAATCTGGAGTACCGGATAATGCTGTCAGTTCATGGCTCCCAGCACAGTGGGATGATCGTTAATGACACAGGACATGAAACTGATGAGAATAGAGCGAAGGTTGAGATAACGCCCAATTCACCAAGAGCCGAAGCCACCCTGGGGGGTTTTGGAAGCCTAGGACTTGATTGTGAACCGAGGACAGGCCTTGACTTTTCAGATTTGTATTACTTGACTATGAATAACAAGCACTGGTTGGTTCACAAGGAGTGGTTCCACGACATTCCATTACCTTGGCACGCTGGGGCAGACACCGGAACTCCACACTGGAACAACAAAGAAGCACTGGTAGAGTTCAAGGACGCACATGCCAAAAGGCAAACTGTCGTGGTTCTAGGGAGTCAAGAAGGAGCAGTTCACACGGCCCTTGCTGGAGCTCTGGAGGCTGAGATGGATGGTGCAAAGGGAAGGCTGTCCTCTGGCCACTTGAAATGTCGCCTGAAAATGGATAAACTTAGATTGAAGGGCGTGTCATACTCCTTGTGTACCGCAGCGTTCACATTCACCAAGATCCCGGCTGAAACACTGCACGGGACAGTCACAGTGGAGGTACAGTACGCAGGGACAGATGGACCTTGCAAGGTTCCAGCTCAGATGGCGGTGGACATGCAAACTCTGACCCCAGTTGGGAGGTTGATAACCGCTAACCCCGTAATCACTGAAAGCACTGAGAACTCTAAGATGATGCTGGAACTTGATCCACCATTTGGGGACTCTTACATTGTCATAGGAGTCGGGGAGAAGAAGATCACCCACCACTGGCACAGGAGTGGCAGCACCATTGGAAAAGCATTTGAAGCCACTGTGAGAGGTGCCAAGAGAATGGCAGTCTTGGGAGACACAGCCTGGGACTTTGGATCAGTTGGAGGCGCTCTCAACTCATTGGGCAAGGGCATCCATCAAATTTTTGGAGCAGCTTTCAAATCATTGTTTGGAGGAATGTCCTGGTTCTCACAAATTCTCATTGGAACGTTGCTGATGTGGTTGGGTCTGAACACAAAGAATGGATCTATTTCCCTTATGTGCTTGGCCTTAGGGGGAGTGTTGATCTTCTTATCCACAGCCGTCTCTGCTGATGTGGGGTGCTCGGTGGACTTCTCAAAGAAGGAGACGAGATGCGGTACAGGGGTGTTCGTCTATAACGACGTTGAAGCCTGGAGGGACAGGTACAAGTACCATCCTGACTCCCCCCGTAGATTGGCAGCAGCAGTCAAGCAAGCCTGGGAAGATGGTATCTGCGGGATCTCCTCTGTTTCAAGAATGGAAAACATCATGTGGAGATCAGTAGAAGGGGAGCTCAACGCAATCCTGGAAGAGAATGGAGTTCAACTGACGGTCGTTGTGGGATCTGTAAAAAACCCCATGTGGAGAGGTCCACAGAGATTGCCCGTGCCTGTGAACGAGCTGCCCCACGGCTGGAAGGCTTGGGGGAAATCGCACTTCGTCAGAGCAGCAAAGACAAATAACAGCTTTGTCGTGGATGGTGACACACTGAAGGAATGCCCACTCAAACATAGAGCATGGAACAGCTTTCTTGTGGAGGATCATGGGTTCGGGGTATTTCACACTAGTGTCTGGCTCAAGGTTAGAGAAGATTATTCATTAGAGTGTGATCCAGCCGTTATTGGAACAGCTGTTAAGGGAAAGGAGGCTGTACACAGTGATCTAGGCTACTGGATTGAGAGTGAGAAGAATGACACATGGAGGCTGAAGAGGGCCCATCTGATCGAGATGAAAACATGTGAATGGCCAAAGTCCCACACATTGTGGACAGATGGAATAGAAGAGAGTGATCTGATCATACCCAAGTCTTTAGCTGGGCCACTCAGCCATCACAATACCAGAGAGGGCTACAGGACCCAAATGAAAGGGCCATGGCACAGTGAAGAGCTTGAAATTCGGTTTGAGGAATGCCCAGGCACTAAGGTCCACGTGGAGGAAACATGTGGAACAAGAGGACCATCTCTGAGATCAACCACTGCAAGCGGAAGGGTGATCGAGGAATGGTGCTGCAGGGAGTGCACAATGCCCCCACTGTCGTTCCGGGCTAAAGATGGCTGTTGGTATGGAATGGAGATAAGGCCCAGGAAAGAACCAGAAAGCAACTTAGTAAGGTCAATGGTGACTGCAGGATCAACTGATCACATGGATCACTTCTCCCTTGGAGTGCTTGTGATTCTGCTCATGGTGCAGGAAGGGCTGAAGAAGAGAATGACCACAAAGATCATCATAAGCACATCAATGGCAGTGCTGGTAGCTATGATCCTGGGAGGATTTTCAATGAGTGACCTGGCTAAGCTTGCAATTTTGATGGGTGCCACCTTCGCGGAAATGAACACTGGAGGAGATGTAGCTCATCTGGCGCTGATAGCGGCATTCAAAGTCAGACCAGCGTTGCTGGTATCTTTCATCTTCAGAGCTAATTGGACACCCCGTGAAAGCATGCTGCTGGCCTTGGCCTCGTGTCTTTTGCAAACTGCGATCTCCGCCTTGGAAGGCGACCTGATGGTTCTCATCAATGGTTTTGCTTTGGCCTGGTTGGCAATACGAGCGATGGTTGTTCCACGCACTGATAACATCACCTTGGCAATCCTGGCTGCTCTGACACCACTGGCCCGGGGCACACTGCTTGTGGCGTGGAGAGCAGGCCTTGCTACTTGCGGGGGGTTTATGCTCCTCTCTCTGAAGGGAAAAGGCAGTGTGAAGAAGAACTTACCATTTGTCATGGCCCTGGGACTAACCGCTGTGAGGCTGGTCGACCCCATCAACGTGGTGGGGCTGCTGTTGCTCACAAGGAGTGGGAAGCGGAGCTGGCCCCCTAGCGAAGTACTCACAGCTGTTGGCCTGATATGCGCATTGGCTGGAGGGTTCGCCAAGGCAGATATAGAGATGGCTGGGCCCATGGCCGCGGTCGGTCTGCTAATTGTCAGTTACGTGGTCTCAGGAAAGAGTGTGGACATGTACATTGAAAGAGCAGGTGACATCACATGGGAAAAAGATGCGGAAGTCACTGGAAACAGTCCCCGGCTCGATGTGGCGCTAGATGAGAGTGGTGATTTCTCCCTGGTGGAGGATGACGGTCCCCCCATGAGAGAGATCATACTCAAGGTGGTCCTGATGACCATCTGTGGCATGAACCCAATAGCCATACCCTTTGCAGCTGGAGCGTGGTACGTATACGTGAAGACTGGAAAAAGGAGTGGTGCTCTATGGGATGTGCCTGCTCCCAAGGAAGTAAAAAAGGGGGAGACCACAGATGGAGTGTACAGAGTAATGACTCGTAGACTGCTAGGTTCAACACAAGTTGGAGTGGGAGTTATGCAAGAGGGGGTCTTTCACACTATGTGGCACGTCACAAAAGGATCCGCGCTGAGAAGCGGTGAAGGGAGACTTGATCCATACTGGGGAGATGTCAAGCAGGATCTGGTGTCATACTGTGGTCCATGGAAGCTAGATGCCGCCTGGGACGGGCACAGCGAGGTGCAGCTCTTGGCCGTGCCCCCCGGAGAGAGAGCGAGGAACATCCAGACTCTGCCCGGAATATTTAAGACAAAGGATGGGGACATTGGAGCGGTTGCGCTGGATTACCCAGCAGGAACTTCAGGATCTCCAATCCTAGACAAGTGTGGGAGAGTGATAGGACTTTATGGCAATGGGGTCGTGATCAAAAATGGGAGTTATGTTAGTGCCATCACCCAAGGGAGGAGGGAGGAAGAGACTCCTGTTGAGTGCTTCGAGCCTTCGATGCTGAAGAAGAAGCAGCTAACTGTCTTAGACTTGCATCCTGGAGCTGGGAAAACCAGGAGAGTTCTTCCTGAAATAGTCCGTGAAGCCATAAAAACAAGACTCCGTACTGTGATCTTAGCTCCAACCAGGGTTGTCGCTGCTGAAATGGAGGAAGCCCTTAGAGGGCTTCCAGTGCGTTATATGACAACAGCAGTCAATGTCACCCACTCTGGAACAGAAATCGTCGACTTAATGTGCCATGCCACCTTCACTTCACGTCTACTACAGCCAATCAGAGTCCCCAACTATAATCTGTATATTATGGATGAGGCCCACTTCACAGATCCCTCAAGTATAGCAGCAAGAGGATACATTTCAACAAGGGTTGAGATGGGCGAGGCGGCTGCCATCTTCATGACCGCCACGCCACCAGGAACCCGTGACGCATTTCCGGACTCCAACTCACCAATTATGGACACCGAAGTGGAAGTCCCAGAGAGAGCCTGGAGCTCAGGCTTTGATTGGGTGACGGATTATTCTGGAAAAACAGTTTGGTTTGTTCCAAGCGTGAGGAACGGCAATGAGATCGCAGCTTGTCTGACAAAGGCTGGAAAACGGGTCATACAGCTCAGCAGAAAGACTTTTGAGACAGAGTTCCAGAAAACAAAACATCAAGAGTGGGACTTTGTCGTGACAACTGACATTTCAGAGATGGGCGCCAACTTTAAAGCTGACCGTGTCATAGATTCCAGGAGATGCCTAAAGCCGGTCATACTTGATGGCGAGAGAGTCATTCTGGCTGGACCCATGCCTGTCACACATGCCAGCGCTGCCCAGAGGAGGGGGCGCATAGGCAGGAATCCCAACAAACCTGGAGATGAGTATCTGTATGGAGGTGGGTGCGCAGAGACTGACGAAGACCATGCACACTGGCTTGAAGCAAGAATGCTCCTTGACAATATTTACCTCCAAGATGGCCTCATAGCCTCGCTCTATCGACCTGAGGCCGACAAAGTAGCAGCCATTGAGGGAGAGTTCAAGCTTAGGACGGAGCAAAGGAAGACCTTTGTGGAACTCATGAAAAGAGGAGATCTTCCTGTTTGGCTGGCCTATCAGGTTGCATCTGCCGGAATAACCTACACAGATAGAAGATGGTGCTTTGATGGCACGACCAACAACACCATAATGGAAGACAGTGTGCCGGCAGAGGTGTGGACCAGACACGGAGAGAAAAGAGTGCTCAAACCGAGGTGGATGGACGCCAGAGTTTGTTCAGATCATGCGGCCCTGAAGTCATTCAAGGAGTTTGCCGCTGGGAAAAGAGGAGCGGCTTTTGGAGTGATGGAAGCCCTGGGAACACTGCCAGGACACATGACAGAGAGATTCCAGGAAGCCATTGACAACCTCGCTGTGCTCATGCGGGCAGAGACTGGAAGCAGGCCTTACAAAGCCGCGGCGGCCCAATTGCCGGAGACCCTAGAGACCATTATGCTTTTGGGGTTGCTGGGAACAGTCTCGCTGGGAATCTTTTTCGTCTTGATGAGGAACAAGGGCATAGGGAAGATGGGCTTTGGAATGGTGACTCTTGGGGCCAGCGCATGGCTCATGTGGCTCTCGGAAATTGAGCCAGCCAGAATTGCATGTGTCCTCATTGTTGTGTTCCTATTGCTGGTGGTGCTCATACCTGAGCCAGAAAAGCAAAGATCTCCCCAGGACAACCAAATGGCAATCATCATCATGGTAGCAGTAGGTCTTCTGGGCTTGATTACCGCCAATGAACTCGGATGGTTGGAGAGAACAAAGAGTGACCTAAGCCATCTAATGGGAAGGAGAGAGGAGGGGGCAACCATGGGATTCTCAATGGACATTGACCTGCGGCCAGCCTCAGCTTGGGCCATCTATGCTGCCTTGACAACTTTCATTACCCCAGCCGTCCAACATGCAGTGACCACTTCATACAACAACTACTCCTTAATGGCGATGGCCACGCAAGCTGGAGTGTTGTTTGGTATGGGCAAAGGGATGCCATTCTACGCATGGG-ACTTTGGAGTCCCGCTGCTAATGATAGGTTGCTACTCACAATTAACGCCCCTGACCCTAATAGTGGCCATCATTTTGCTCGTGGCGCACTACATGTACTTGATCCCAGGGCTGCAGGCAGCAGCTGCGCGTGCTGCCCAGAAGAGAACGGCAGCTGGCATCATGAAGAACCCTGTTGTGGATGGAATAGTGGTGACTGACATTGACACAATGACAATTGACCCCCAAGTGGAGAAAAAGATGGGACAGGTGCTACTCATGGCAGTAGCCGTCTCCAGCGCCATACTGTCGCGGACCGCCTGGGGGTGGGGGGAGGCTGGGGCCCTGATCACAGCCGCAACTTCCACTTTGTGGGAAGGCTCTCCGAACAAGTACTGGAACTCCTCTACAGCCACTTCACTGTGTAACATTTTTAGGGGAAGTTACTTGGCTGGAGCTTCTCTAATCTACACAGTAACAAGAAACGCTGGCTTGGTCAAGAGACGTGGGGGTGGAACAGGAGAGACCCTGGGAGAGAAATGGAAGGCCCGCTTGAACCAGATGTCGGCCCTGGAGTTCTACTCCTACAAAAAGTCAGGCATCACCGAGGTGTGCAGAGAAGAGGCCCGCCGCGCCCTCAAGGACGGTGTGGCAACGGGAGGCCATGCTGTGTCCCGAGGAAGTGCAAAGCTGAGATGGTTGGTGGAGCGGGGATACCTGCAGCCCTATGGAAAGGTCATTGATCTTGGATGTGGCAGAGGGGGCTGGAGTTACTACGCCGCCACCATCCGCAAAGTTCAAGAAGTGAAAGGATACACAAAAGGAGGCCCTGGTCATGAAGAACCCGTGTTGGTGCAAAGCTATGGGTGGAACATAGTCCGTCTTAAGAGTGGGGTGGACGTCTTTCATATGGCGGCTGAGCCGTGTGACACGTTGCTGTGTGACATAGGTGAGTCATCATCTAGTCCTGAAGTGGAAGAAGCACGGACGCTCAGAGTCCTCTCCATGGTGGGGGATTGGCTTGAAAAAAGACCAGGAGCCTTTTGTATAAAAGTGTTGTGCCCATACACCAGCACTATGATGGAAACCCTGGAGCGACTGCAGCGTAGGTATGGGGGAGGACTGGTCAGAGTGCCACTCTCCCGCAACTCTACACATGAGATGTACTGGGTCTCTGGAGCGAAAAGCAACACCATAAAAAGTGTGTCCACCACGAGCCAGCTCCTCTTGGGGCGCATGGACGGGCCTAGGAGGCCAGTGAAATATGAGGAGGATGTGAATCTCGGCTCTGGCACGCGGGCTGTGGTAAGCTGCGCTGAAGCTCCCAACATGAAGATCATTGGTAACCGCATTGAAAGGATCCGCAGTGAGCACGCGGAAACGTGGTTCTTTGACGAGAACCACCCATATAGGACATGGGCTTACCATGGAAGCTATGAGGCCCCCACACAAGGGTCAGCGTCCTCTCTAATAAACGGGGTTGTCAGGCTCCTGTCAAAACCCTGGGATGTGGTGACTGGAGTCACAGGAATAGCCATGACCGACACCACACCGTATGGTCAGCAAAGAGTTTTCAAGGAAAAAGTGGACACTAGGGTGCCAGACCCCCAAGAAGGTACTCGTCAGGTTATGAGCATGGTCTCTTCCTGGTTGTGGAAAGAGCTAGGCAAACACAAACGGCCACGAGTCTGTACCAAAGAAGAGTTCATCAACAAGGTTCGTAGCAATGCAGCATTAGGGGCAATATTTGAAGAGGAAAAAGAGTGGAAGACTGCAGTGGAAGCTGTGAACGATCCAAGGTTCTGGGCTCTAGTGGACAAGGAAAGAGAGCACCACCTGAGAGGAGAGTGCCAGAGTTGTGTGTACAACATGATGGGAAAAAGAGAAAAGAAACAAGGGGAATTTGGAAAGGCCAAGGGCAGCCGCGCCATCTGGTATATGTGGCTAGGGGCTAGATTTCTAGAGTTCGAAGCCCTTGGATTCTTGAACGAGGATCACTGGATGGGGAGAGAGAACTCAGGAGGTGGTGTTGAAGGGCTGGGATTACAAAGACTCGGATATGTCCTAGAAGAGATGAGTCGCATACCAGGAGGAAGGATGTATGCAGATGACACTGCTGGCTGGGACACCCGCATCAGCAGGTTTGATCTGGAGAATGAAGCTCTAATCACCAACCAAATGGAGAAAGGGCACAGGGCCTTGGCATTGGCCATAATCAAGTACACATACCAAAACAAAGTGGTAAAGGTCCTTAGACCAGCTGAAAAAGGGAAAACAGTTATGGACATTATTTCGAGACAAGACCAAAGGGGGAGCGGACAAGTTGTCACTTACGCTCTTAACACATTTACCAACCTAGTGGTGCAACTCATTCGGAATATGGAGGCTGAGGAAGTCCTAGAGATGCAAGACTTGTGGCTGCTGCGGAGGTCAGAGAAAGTGACCAACTGGTTGCAGAGCAACGGATGGGATAGGCTCAAACGAATGGCAGTCAGTGGAGATGATTGCGTTGTGAAGCCAATTGATGATAGGTTTGCACATGCCCTCAGGTTCTTGAATGATATGGGAAAAGTTAGGAAGGACACACAAGAGTGGAAACCCTCAACTGGATGGGACAACTGGGAAGAAGTTCCGTTTTGCTCCCACCACTTCAACAAGCTCCATCTCAAGGACGGGAGGTCCATTGTGGTTCCCTGCCGCCACCAAGATGAACTGATTGGCCGGGCCCGCGTCTCTCCAGGGGCGGGATGGAGCATCCGGGAGACTGCTTGCCTAGCAAAATCATATGCGCAAATGTGGCAGCTCCTTTATTTCCACAGAAGGGACCTCCGACTGATGGCCAATGCCATTTGTTCATCTGTGCCAGTTGACTGGGTTCCAACTGGGAGAACTACCTGGTCAATCCATGGAAAGGGAGAATGGATGACCACTGAAGACATGCTTGTGGTGTGGAACAGAGTGTGGATTGAGGAGAACGACCACATGGAAGACAAGACCCCAGTTACGAAATGGACAGACATTCCCTATTTGGGAAAAAGGGAAGACTTGTGGTGTGGATCTCTCATAGGGCACAGACCGCGCACCACCTGGGCTGAGAACATTAAAAACACAGTCAACATGGTGCGCAGGATCATAGGTGATGAAGAAAAGTACATGGACTACCTATCCACCCAAGTTCGCTACTTGGGTGAAGAAGGGTCTACACCTGGAGTGCTGTAA

>ENA|AFD30972|AFD30972.1 Zika virus partial polyprotein

ATGAAAAACCC-AAAGAAGAAATCCGGAGGATTCCGGATTGTCAATATGCTAAAACGCGGAGTAGCCCGTGTGAGCCCCTTTGGGGGCTTGAAGAGGCTGCCAGCCGGACTTCTGCTGGGTCATGGGCCCATCAGGATGGTCTTGGCGATTCTAGCCTTTTTGAGATTCACGGCAATCAAGCCATCACTGGGTCTCATCAATAGATGGGGTTCAGTGGGGAAAAAAGAGGCTATGGAAATAATAAAGAAGTTTAAGAAAGATCTGGCTGCCATGCTGAGAATAATCAATGCTAGGAAGGAGAAGAAGAGACGAGGCACAGATACTAGTGTCGGAATTGTTGGCCTCCTGCTGACCACAGCCATGGCAGTGGAGGTCACTAGACGTGGGAATGCATACTATATGTACTTGGACAGAAGCGATGCTGGGGAGGCCATATCTTTTCCAACCACAATGGGGATGAATAAGTGTTATATACAGATCATGGATCTTGGACACATGTGTGATGCCACCATGAGCTATGAATGCCCTATGCTGGATGAGGGGGTAGAACCAGATGACGTCGATTGTTGGTGCAACACGACGTCAACTTGGGTTGTGTACGGAACCTGCCACCACAAAAAAGGTGAAGCACGGAGATCTAGAAGAGCTGTGACGCTCCCCTCCCATTCCACTAGGAAGCTGCAAACGCGGTCGCAGACCTGGTTGGAATCAAGAGAATACACAAAGCACCTGATTAGAGTCGAAAATTGGATATTCAGGAACCCTGGCTTCGCGTTAGCAGCAGCTGCCATCGCTTGGCTTTTGGGAAGCTCAACGAGCCAAAAAGTCATATACTTGGTCATGATACTGCTGATTGCCCCGGCATACAGCATCAGGTGCATAGGAGTCAGCAATAGGGACTTTGTGGAAGGTATGTCAGGTGGGACTTGGGTTGATGTTGTCTTGGAACATGGAGGTTGTGTTACCGTAATGGCACAGGACAAACCGACTGTCGACATAGAGCTGGTTACAACAACAGTCAGCAACATGGCGGAGGTAAGATCCTACTGCTATGAGGCATCAATATCGGACATGGCTTCGGACAGCCGCTGCCCAACACAAGGTGAAGCCTACCTTGACAAGCAATCAGACACTCAATATGTCTGCAAAAGAACGTTAGTGGACAGAGGCTGGGGAAATGGATGTGGACTTTTTGGCAAAGGGAGCCTGGTGACATGCGCTAAGTTTGCTTGCTCTAAGAAAATGACCGGGAAGAGCATCCAGCCAGAGAATCTGGAGTACCGGATAATGCTGTCAGTTCATGGCTCCCAGCACAGTGGGATGATCGTTAATGATACAGGACATGAAACTGATGAGAATAGAGCGAAGGTTGAGATAACGCCCAATTCACCAAGAGCCGAAGCCACCCTGGGGGGTTTTGGAAGCCTAGGACTTGATTGTGAACCGAGGACAGGCCTTGACTTTTCAGATTTGTATTACTTGACTATGAATAACAAGCACTGGTTGGTTCACAAGGAGTGGTTCCACGACATTCCATTACCTTGGCATGCTGGGGCAGACACCGGAACTCCACACTGGAACAACAAAGAAGCACTGGTAGAGTTCAAGGACGCACATGCCAAAAGGCAGACTGTCGTGGTTCTAGGGAGTCAAGAAGGAGCAGTTCACACGGCCCTTGCTGGAGCTCTGGAGGCTGAGATGGATGGTGCAAAGGGAAGGCTGTCCTCTGGCCACTTGAAATGTCGCCTGAAAATGGATAAACTTAGATTGAAGGGCGTGTCATACTCCTTGTGTACCGCAGCGTTCACATTCACTAAGATCCCGGCTGAAACACTGCACGGGACAGTCACAGTGGAGGTACAGTACGCAGGGACAGATGGACCTTGCAAGGTTCCAGCTCAGATGGCGGTGGACATGCAAACTCTGACCCCAGTTGGGAGGTTGATAACCGCTAACCCTGTAATCACTGAAAGCACTGAGAACTCCAAGATGATGCTGGAACTGGATCCACCATTTGGGGACTCTTACATTGTCATAGGAGTCGGGGAAAAGAAGATCACCCACCACTGGCACAGGAGTGGCAGCACCATTGGAAAAGCATTTGAAGCCACTGTGAGAGGTGCCAAGAGAATGGCAGTCTTGGGAGACACAGCCTGGGACTTTGGATCAGTTGGGGGTGCTCTCAACTCACTGGGCAAGGGCATCCATCAAATTTTTGGAGCAGCTTTCAAATCATTGTTTGGAGGAATGTCCTGGTTCTCACAAATTCTCATTGGAACGTTGCTGGTGTGGTTGGGTCTGAATACAAAGAATGGATCTATTTCCCTTATGTGCTTGGCCTTAGGGGGAGTGTTGATCTTCTTATCCACAGCCGTCTCTGCTGATGTGGGGTGCTCGGTGGACTTCTCAAAGAAGGAAACGAGATGCGGTACAGGGGTGTTCGTCTATAACGACGTTGAAGCTTGGAGGGACAGGTACAAGTACCATCCTGACTCCCCTCGTAGATTGGCAGCAGCAGTCAAGCAAGCCTGGGAAGATGGGATCTGTGGGATCTCCTCTGTTTCAAGAATGGAAAACATCATGTGGAGATCAGTAGAAGGGGAGCTCAACGCAATCCTGGAAGAGAATGGAGTTCAACTGACGGTCGTTGTGGGATCTGTAAAAAACCCCATGTGGAGAGGTCCACAGAGATTGCCCGTGCCTGTGAACGAGCTGCCCCATGGCTGGAAGGCTTGGGGGAAATCGTACTTCGTCAGGGCAGCAAAGACAAATAACAGCTTTGTCGTGGATGGTGACACACTGAAGGAATGCCCACTCAAACATAGAGCATGGAACAGCTTTCTTGTGGAGGATCATGGGTTCGGGGTATTTCACACTAGTGTCTGGCTCAAGGTTAGAGAAGATTATTCATTAGAGTGTGATCCAGCCGTCATTGGAACAGCCGCTAAGGGAAAGGAGGCTGTGCACAGTGATCTAGGCTACTGGATTGAGAGTGAGAAGAACGACACATGGAGGCTGAAGAGGGCCCACCTGATCGAGATGAAAACATGTGAATGGCCAAAGTCCCACACATTGTGGACAGATGGAATAGAAGAAAGTGATCTGATCATACCCAAGTCTTTAGCTGGGCCACTCAGCCATCACAACACCAGAGAGGGCTACAGGACCCAAATGAAAGGGCCATGGCATAGTGAAGAGCTTGAAATTCGGTTTGAGGAATGCCCAGGCACTAAGGTCCACGTGGAGGAAACATGTGGAACAAGAGGACCATCTCTGAGATCAACCACTGCAAGCGGAAGGGTGATCGAGGAATGGTGCTGCAGGGAGTGCACAATGCCCCCACTGTCGTTCCGGGCTAAAGATGGTTGTTGGTATGGAATGGAGATAAGGCCCAGGAAAGAACCAGAAAGTAACTTAGTAAGGTCAATGGTGACTGCAGGATCAACTGATCACATGGATCACTTCTCCCTTGGAGTGCTTGTGATTCTGCTCATGGTACAGGAAGGGCTAAAGAAGAGAATGACCACAAAGATCATCATAAGCACATCAATGGCAGTGCTGGTAGCTATGATCCTGGGAGGATTTTCAATGAGTGACCTGGCTAAGCTTGCAATTTTGATGGGTGCCACCTTCGCGGAAATGAACACTGGAGGAGATGTTGCTCATCTGGCGCTGATAGCGGCATTCAAAGTCAGACCTGCGTTGCTGGTATCTTTCATTTTCAGAGCTAATTGGACACCCCGTGAGAGCATGCTGCTGGCCTTGGCCTCGTGTCTTCTGCAAACTGCGATCTCCGCCTTGGAAGGCGACCTGATGGTTCCCATCAATGGTTTTGCTTTGGCCTGGTTGGCAATACGAGCGATGGTTGTTCCACGCACTGACAACATCACCTTGGCAATCCTGGCTGCTCTGACACCACTGGCCCGGGGCACACTGCTTGTGGCGTGGAGAGCAGGCCTTGCTACTTGCGGGGGGTTCATGCTCCTTTCTCTGAAGGGGAAAGGCAGTGTGAAGAAGAACTTACCATTTGTCATGGCCCTGGGACTAACCGCTGTGAGGCTGGTCGACCCCATCAACGTGGTGGGACTGCTGTTGCTCACAAGGAGTGGGAAGCGGAGCTGGCCCCCTAGTGAAGTACTCACAGCTGTTGGCCTGATATGCGCATTGGCTGGAGGGTTCGCCAAGGCGGATATAGAGATGGCTGGGCCCATGGCCGCGGTCGGTCTGCTAATTGTCAGTTACGTGGTCTCAGGAAAGAGTGTGGACATGTACATTGAAAGAGCAGGTGACATCACATGGGAAAAAGATGCGGAAGTCACTGGAAACAGTCCCCGGCTCGATGTGGCACTAGATGAGAGTGGTGATTTCTCCCTAGTGGAGGATGATGGTCCCCCCATGAGAGAGATCATACTCAAAGTGGTCCTGATGNCCATCTGTGGCATGAACCCAATAGCCATACCCTTTGCAGCTGGAGCGTGGTACGTGTATGTGAAGACTGGAAAAAGGAGTGGTGCTCTATGGGATGTGCCTGCTCCCAAGGAAGTAAAAAAGGGGGAGACCACAGATGGAGTGTACAGAGTAATGACTCGTAGACTGCTAGGTTCAACACAAGTTGGAGTGGGAGTCATGCAAGAGGGGGTCTTCCACACTATGTGGCACGTCACAAAAGGATCCGCGCTGAGAAGCGGTGAAGGGAGACTTGATCCATACTGGGGAGATGTCAAGCAGGATCTGGTGTCATACTGTGGTCCATGGAAGCTAGATGCCGCCTGGGACGGGCACAGCGAGGTGCAGCTCTTGGCCGTGCCCCCCGGAGAGAGAGCGAGGAACATCCAGACTCTGCCCGGAATATTTAAGACAAAGGATGGGGACATTGGAGCAGTTGCGCTGGACTACCCAGCAGGAACTTCAGGATCTCCAATCCTAGATAAGTGTGGGAGAGTGATAGGACTCTATGGTAATGGGGTCGTGATCAAAAATGGGAGTTACGTTAGTGCCATCACCCAAGGGAGGAGGGAGGAAGAGACTCCTGTTGAGTGCTTCGAGCCTTCGATGCTGAAGAAGAAGCAGCTAACTGTCTTAGACTTGCATCCTGGAGCTGGGAAAACCAGGAGAGTTCTTCCTGAAATAGTCCGTGAAGCCATAAAAACAAGACTCCGCACTGTGATCTTAGCTCCAACCAGGGTTGTCGCTGCTGAAATGGAGGAAGCCCTTAGAGGGCTTCCAGTGCGTTATATGACAACAGCAGTCAATGTCACCCATTCTGGGACAGAAATCGTTGACTTAATGTGCCATGCCACCTTCACTTCACGTCTACTACAGCCAATCAGAGTCCCCAACTATAATCTGTATATTATGGATGAGGCCCACTTCACAGATCCCTCAAGTATAGCAGCAAGAGGATACATTTCAACAAGGGTTGAGATGGGCGAGGCGGCTGCCATCTTCATGACTGCCACGCCACCAGGAACCCGTGACGCATTCCCGGACTCCAACTCACCAATTATGGACACCGAAGTGGAAGTCCCAGAGAGAGCCTGGAGCTCAGGCTTTGATTGGGTGACGGATCATTCTGGAAAAACAGTTTGGTTTGTTCCAAGCGTGAGGAATGGCAATGAGATCGCAGCTTGTCTGACAAAGGCTGGAAAACGGGTCATACAGCTCAGCAGAAAGACTTTTGAGACAGAGTTCCAGAAAACAAAACATCAAGAGTGGGACTTCGTCGTGACAACTGACATTTCAGAGATGGGCGCCAACTTTAAAGCTGACCGTGTCATAGATTCCAGGAGATGCCTAAAGCCGGTCATACTTGATGGCGAGAGAGTCATTCTGGCTGGACCCATGCCTGTCACACATGCCAGCGCTGCCCAGAGGAGGGGGCGCATAGGCAGGAACCCCAACAAACCTGGAGATGAGTATCTGTATGGAGGTGGGTGCGCAGAGACTGATGAAGACCATGCACACTGGCTTGAAGCAAGAATGCTTCTTGACAACATTTACCTCCAAGATGGCCTCATAGCCTCGCTCTATCGACCTGAGGCCGACAAAGTAGCAGCTATTGAGGGAGAGTTCAAGCTTAGGACGGAGCAAAGGAAGACCTTTGTGGAACTCATGAAAAGAGGAGATCTTCCTGTTTGGCTGGCCTATCAGGTTGCATCTGCCGGAATAACCTACACAGATAGAAGATGGTGCTTTGATGGCACGACCAACAACACCATAATGGAAGACAGTGTGCCGGCAGAGGTGTGGACCAGATACGGAGAGAAAAGAGTGCTCAAACCGAGGTGGATGGACGCCAGAGTTTGTTCAGATCATGCGGCCCTGAAGTCATTCAAAGAGTTTGCCGCTGGGAAAAGAGGAGCGGCCTTTGGAGTGATGGAAGCCCTGGGAACACTGCCAGGACATATGACAGAGAGATTCCAGGAGGCCATTGACAACCTCGCTGTGCTCATGCGGGCAGAGACTGGAAGCAGGCCCTACAAAGCCGCGGCGGCCCAATTACCGGAGACCCTAGAGACTATCATGCTTTTGGGGTTGCTGGGAACAGTCTCGCTGGGAATCTTTTTCGTCTTGATGCGGAACAAGGGCATAGGGAAGATGGGCTTTGGAATGGTGACTCTTGGGGCCAGCGCATGGCTTATGTGGCTCTCGGAAATTGAGCCAGCCAGAATTGCATGTGTCCTCATTGTTGTGTTCCTATTGCTGGTGGTGCTCATACCTGAGCCAGAAAAGCAAAGATCTCCCCAGGACAACCAAATGGCAATCATCATCATGGTAGCAGTGGGTCTTCTGGGCTTGATTACCGCCAATGAACTCGGATGGTTGGAGAGAACAAAGAGTGACCTAAGCCATCTAATGGGAAGGAGAGAGGAGGGGGCAACNATAGGATTCTCAATGGACATTGACCTGCGGCCAGCCTCAGCTTGGGCTATCTATGCTGCTCTGACAACTTTCATTACCCCAGCCGTCCAACATGCAGTGACCACTTCATACAACAACTACTCCTTAATGGCGATGGCCACGCAAGCTGGAGTGTTGTTCGGTATGGGTAAAGGGATGCCATTCTATGCATGGG-ACTTTGGAGTCCCGCTGCTAATGATAGGTTGCTACTCACAATTAACACCCCTGACCCTAATAGTGGCCATCATTTTGCTCGTGGCGCACTACATGTACTTGATCCCAGGGCTGCAGGCAGCAGCTGCGCGTGCTGCCCAGAAGAGAACGGCAGCTGGCATCATGAAGAACCCTGTTGTGGATGGAATAGTGGTGACTGACATTGACACAATGACAATTGACCCCCAAGTGGAGAAAAAGATGGGACAGGTGCTACTCATAGCAGTAGCTGTCTCCAGCGCCATACTGTCGCGGACCGCCTGGGGGTGGGGTGAGGCTGGGGCCCTGATCACAGCTGCAACTTCCACTTTGTGGGAGGGCTCTCCGAACAAGTACTGGAACTCCTCCACAGCCACCTCACTGTGTAACATTTTTAGGGGAAGCTACTTGGCTGGAGCTTCTCTAATCTACACAGTAACAAGAAACGCTGGCTTGGTCAAGAGACGTGGGGGTGGAACGGGAGAGACCCTGGGAGAGAAATGGAAGGCCCGCCTGAACCAGATGTCGGCCCTGGAGTTCTACTCCTACAAAAAGTCAGGCATCACCGAGGTGTGCAGAGAAGAGGCCCGCCGCGCCCTCAAGGACGGTGTGGCAACGGGAGGCCACGCTGTGTCCCGAGGAAGTGCAAAGCTGAGATGGTTGGTGGAGAGGGGATACCTGCAGCCCTATGGAAAGGTCATTGATCTTGGATGTGGCAGAGGGGGCTGGAGTTACTATGCCGCCACCATCCGCAAAGTTCAAGAAGTGAAAGGATACACAAAAGGAGGCCCTGGTCATGAAGAACCCATGTTGGTGCAAAGCTATGGGTGGAACATAGTCCGTCTTAAGAGTGGGGTGGACGTCTTTCATATGGCGGCTGAGCCGTGTGACACGTTGCTGTGTGATATAGGTGAGTCATCATCTAGTCCTGAAGTGGAAGAAGCACGGACGCTCAGAGTCCTCTCCATGGTGGGGGATTGGCTTGAAAAAAGACCAGGAGCCTTTTGTATAAAAGTGTTGTGCCCATACACCAGCACTATGATGGAAACCCTGGAGCGACTGCAGCGTAGGTATGGGGGAGGACTGGTCAGAGTGCCACTCTCCCGCAACTCTACACATGAGATGTACTGGGTCTCTGGAGCGAAAAGCAACACCATAAAAAGTGTGTCCACCACGAGCCAGCTCCTTTTGGGGCGCATGGACGGGCCCAGGAGGCCAGTGAAATATGAAGAGGATGTGAATCTCGGCTCTGGCACGCGGGCTGTGGTAAGCTGCGCTGAAGCTCCCAACATGAAGATCATTGGTAACCGCATTGAGAGGATCCGCAGTGAGCACGCGGAAACGTGGTTCTTTGACGAGAACCACCCATATAGGACATGGGCTTACCATGGAAGCTACGAGGCCCCCACACAAGGGTCAGCGTCCTCTCTAATAAACGGGGTTGTCAGGCTCCTGTCAAAACCCTGGGATGTGGTGACTGGAGTCACAGGAATAGCCATGACCGACACCACACCGTATGGTCAGCAAAGAGTTTTCAAGGAAAAAGTGGACACTAGGGTGCCAGACCCCCAAGAAGGCACTCGTCAGGTTATGAGCATGGTCTCTTCCTGGTTGTGGAAAGAGTTAGGCAAACACAAACGGCCACGAGTCTGTACCAAAGAAGAGTTCATCAACAAGGTTCGTAGCAACGCAGCATTAGGGGCAATATTTGAAGAGGAAAAAGAGTGGAAGACTGCAGTGGAAGCTGTGAACGATCCAAGGTTCTGGGCTCTAGTGGACAAGGAAAGAGAGCACCACCTGAGAGGAGAGTGCCAGAGCTGTGTGTACAACATGATGGGAAAAAGAGAAAAGAAACAAGGGGAATTTGGAAAGGCCAAGGGCAGCCGCGCCATCTGGTACATGTGGCTAGGGGCTAGATTTCTAGAGTTCGAAGCCCTTGGATTCTTGAACGAGGATCACTGGATGGGGAGAGAGAATTCAGGAGGTGGTGTTGAAGGGCTAGGATTACAAAGACTCGGATATGTCTTAGAAGAGATGAGTCGCATACCAGGAGGAAGGATGTATGCAGATGATACTGCTGGCTGGGACACCCGCATCAGCAGGTTTGATCTGGAGAATGAAGCTCTAATCACCAACCAAATGGAGAAAGGGCACAGGGCCTTGGCATTGGCCATAATCAAGTACACATACCAAAACAAAGTGGTAAAGGTCCTTAGACCAGCTGAAAAAGGGAAGACAGTTATGGACATTATTTCAAGACAAGACCAAAGGGGGAGCGGACAAGTTGTCACTTACGCTCTTAATACATTTACCAACCTAGTGGTGCAGCTCATTCGGAATATGGAGGCTGAGGAAGTTCTAGAGATGCAAGACTTGTGGCTGCTGCGGAGGTCAGAGAAAGTGACCAACTGGTTGCAGAGCAATGGATGGGATAGGCTCAAACGAATGGCAGTCAGTGGAGATGATTGCGTTGTGAAACCAATTGATGATAGGTTTGCACATGCTCTCAGGTTCTTGAATGATATGGGAAAAGTTAGGAAGGACACACAAGAGTGGAAGCCCTCAACTGGATGGGACAACTGGGAAGAAGTTCCGTTTTGCTCCCACCACTTCAACAAGCTCCATCTCAAGGACGGGAGGTCCATTGTGGTTCCCTGCCGCCACCAAGATGAACTGATTGGCCGAGCTCGCGTCTCACCGGGGGCGGGATGGAGCATCCGGGAGACTGCTTGCCTAGCAAAATCATATGCGCAAATGTGGCAGCTCCTTTATTTCCACAGAAGGGACCTCCGACTGATGGCCAATGCCATTTGTTCATCTGTGCCAGTTGACTGGGTTCCAACTGGGAGAACTACCTGGTCAATCCATGGAAAGGGAGAATGGATGACCACTGAAGACATGCTTGTGGTGTGGAACAGAGTGTGGATTGAGGAGAACGACCACATGGAAGACAAGACCCCAGTTACGAAATGGACAGACATTCCCTATTTGGGAAAAAGGGAAGACTTGTGGTGTGGGTCTCTCATAGGGCACAGACCGCGCACCACCTGGGCTGAGAACATTAAAAACACAGTCAACATGATGCGTAGGATCATAGGTGATGAAGAAAAGTACGTGGACTACCTATCCACCCAAGTTCGCTACTTGGGCGAAGAAGGGTCCACACCTGGAGTGCTA---

>ENA|AEN75263|AEN75263.1 Zika virus partial polyprotein

ATGAAAAACCC-AAAGAAGAAATCCGGAGGATTCCGGATTGTCAATATGCTAAAACGCGGAGTAGCCCGTGTAAACCCCTTGGGAGGTTTGAAGAGGTTGCCAGCCGGACTTCTGCTGGGTCATGGACCCATCAGAATGGTTTTGGCGATACTAGCCTTTTTGAGATTTACAGCAATCAAGCCATCACTGGGCCTTATCAACAGATGGGGTTCCGTGGGGAAAAAAGAGGCTATGGAAATAATAAAGAAGTTCAAGAAAGATCTTGCTGCCATGTTGAGAATAATCAATGCTAGGAAAGAGAGGAAGAGACGTGGCGCAGACACCAGCATCGGAATCATTGGCCTCCTGCTGACTACAGCCATGGCAGCAGAGATCACTAGACGCGGGAGTGCATACTACATGTACTTGGATAGGAGCGATGCCGGGAAGGCCATTTCGTTTGCTACCACATTGGGAGTGAACAAGTGCCACGTACAGATCATGGACCTCGGGCACATGTGTGACGCCACCATGAGTTATGAGTGCCCTATGCTGGATGAGGGAGTGGAACCAGATGATGTCGATTGCTGGTGCAACACGACATCAACTTGGGTTGTGTACGGAACCTGTCATCACAAAAAAGGTGAGGCACGGCGATCTAGAAGAGCCGTGACGCTCCCTTCTCACTCTACAAGGAAGTTGCAAACGCGGTCGCAGACCTGGTTAGAATCAAGAGAATACACGAAGCACTTGATCAAGGTTGAAAACTGGATATTCAGGAACCCCGGGTTTGCGCTAGTGGCCGTTGCCATTGCCTGGCTTTTGGGAAGCTCGACGAGCCAAAAAGTCATATACTTGGTCATGATACTGCTGATTGCCCCGGCATACAGTATCAGGTGCATTGGAGTCAGCAATAGAGACTTCGTGGAGGGCATGTCAGGTGGGACCTGGGTTGATGTTGTCTTGGAACATGGAGGCTGCGTTACCGTGATGGCACAGGACAAGCCAACAGTTGACATAGAGTTGGTCACGACGACGGTTAGTAACATGGCCGAGGTAAGATCCTATTGCTACGAGGCATCGATATCGGACATGGCTTCGGACAGTCGTTGCCCAACACAAGGTGAAGCCTACCTTGACAAGCAATCAGACACTCAATATGTCTGCAAAAGAACATTAGTGGACAGAGGTTGGGGAAACGGTTGTGGACTTTTTGGCAAAGGGAGCTTGGTGACATGTGCCAAGTTTACGTGTTCTAAGAAGATGACCGGGAAGAGCATTCAACCGGAAAATCTGGAGTATCGGATAATGCTATCAGTGCATGGCTCCCAGCATAGCGGGATGATTGTCAATGATACAGGATATGAAACTGACGAAAATAGAGCGAAAGTCGAGGTTACGCCTAATTCACCAAGAGCGGAAGCAACCTTGGGAGGCTTTGGAAGCTTAGGACTTGACTGTGAACCAAGGACAGGCCTTGACTTTTCAGATCTGTATTACCTGACCATGAACAATAAGCATTGGTTGGTGCACAAAGAGTGGTTTCATGACATCCCATTGCCTTGGCATGCTGGGGCAGACACCGGAACTCCACACTGGAACAACAAAGAGGCATTGGTAGAATTCAAGGATGCCCACGCCAAGAGGCAAACCGTCGTCGTTCTGGGGAGCCAGGAAGGAGCCGTTCACACGGCTCTCGCTGGAGCTCTAGAGGCTGAGATGGATGGTGCAAAGGGAAAGCTGTTCTCTGGCCATTTGAAATGCCGCCTAAAAATGGACAAGCTTAGATTGAAGGGCGTGTCATATTCCTTGTGCACTGCGGCATTCACATTCACCAAGGTCCCAGCTGAAACACTGCATGGAACAGTCACAGTGGAGGTGCAGTATGCAGGGACAGATGGACCCTGCAAGATCCCAGTCCAGATGGCGGTGGACATGCAGACCCTGACCCCAGTTGGAAGGCTGATAACCGCCAACCCCGTGATTACTGAAAGCACTGAGAACTCAAAGATGATGTTGGAGCTTGACCCACCATTTGGGGATTCTTACATTGTCATAGGAGTTGGGGACAAGAAAATCACCCACCACTGGCATAGGAGTGGTAGCACCATCGGAAAGGCATTTGAGGCCACTGTGAGAGGCGCCAAGAGAATGGCAGTCCTGGGGGATACAGCCTGGGACTTCGGATCAGTCGGGGGTGTGTTCAACTCACTGGGTAAGGGCATTCACCAGATTTTTGGAGCAGCCTTCAAATCACTGTTTGGAGGAATGTCCTGGTTCTCACAGATCCTCATAGGCACGCTGCTAGTGTGGTTAGGTTTGAACACAAAGAATGGATCTATCTCCCTCACATGCTTGGCCCTGGGGGGAGTGATGATCTTCCTCTCCACGGCTGTTTCTGCTGACGTGGGGTGCTCAGTGGACTTCTCAAAAAAGGAAACGAGATGTGGCACGGGGGTATTCATCTATAATGATGTTGAAGCCTGGAGGGACCGGTACAAGTACCATCCTGACTCCCCCCGCAGATTGGCAGCAGCAGTCAAGCAGGCCTGGGAAGAGGGGATCTGTGGGATCTCATCCGTTTCAAGAATGGAAAACATCATGTGGAAATCAGTAGAAGGGGAGCTCAATGCTATCCTAGAGGAGAATGGAGTTCAACTGACAGTTGTTGTGGGATCTGTAAAAAACCCCATGTGGAGAGGTCCACAAAGATTGCCAGTGCCTGTGAATGAGCTGCCCCATGGCTGGAAAGCCTGGGGGAAATCGTATTTTGTTAGGGCGGCAAAGACCAACAACAGTTTTGTTGTCGACGGTGACACACTGAAGGAATGTCCGCTTGAGCACAGAGCATGGAATAGTTTTCTTGTGGAGGATCACGGGTTTGGAGTCTTCCACACCAGTGTCTGGCTTAAGGTCAGAGAAGATTACTCATTAGAATGTGACCCAGCCGTCATAGGAACAGCTGTTAAGGGAAGGGAGGCCGCGCACAGTGATCTGGGCTATTGGATTGAAAGTGAAAAGAATGACACATGGAGGCTGAAGAGGGCCCACCTGATTGAGATGAAAACATGTGAATGGCCAAAGTCTCACACATTGTGGACAGATGGAGTAGAAGAAAGTGATCTTATCATACCCAAGTCTTTAGCTGGTCCACTCAGCCACCACAACACCAGAGAGGGTTACAGAACCCAAGTGAAAGGGCCATGGCACAGTGAAGAGCTTGAAATCCGGTTTGAGGAATGTCCAGGCACCAAGGTTTACGTGGAGGAGACATGCGGAACTAGAGGACCATCTCTGAGATCAACTACTGCAAGTGGAAGGGTCATTGAGGAATGGTGCTGTAGGGAATGCACAATGCCCCCACTATCGTTTCGAGCAAAAGACGGCTGCTGGTATGGAATGGAGATAAGGCCCAGGAAAGAACCAGAGAGCAACTTAGTGAGGTCAATGGTGACAGCGGGGTCAACCGATCATATGGACCACTTCTCTCTTGGAGTGCTTGTGATTCTACTCATGGTGCAGGAGGGGTTGAAGAAGAGAATGACCACAAAGATCATCATGAGCACATCAATGGCAGTGCTGGTAGTCATGATCTTGGGAGGATTTTCAATGAGTGACCTGGCCAAGCTTGTGATCCTGATGGGTGCTACTTTCGCAGAAATGAACACTGGAGGAGATGTAGCTCACTTGGCATTGGTAGCGGCATTTAAAGTCAGACCAGCCTTGCTGGTCTCCTTCATTTTCAGAGCCAATTGGACACCCCGTGAGAGCATGCTGCTAGCCCTGGCTTCGTGTCTTCTGCAAACTGCGATCTCTGCTCTTGAAGGTGACTTGATGGTCCTCATTAATGGATTTGCTTTGGCCTGGTTGGCAATTCGAGCAATGGCCGTGCCACGCACTGACAACATCGCTCTACCAATCTTGGCTGCTCTAACACCACTAGCTCGAGGCACACTGCTCGTGGCATGGAGAGCGGGCCTGGCTACTTGTGGAGGGATCATGCTCCTCTCCCTGAAAGGGAAAGGTAGTGTGAAGAAGAACCTGCCATTTGTCATGGCCCTGGGATTGACAGCTGTGAGGGTAGTAGACCCTATTAATGTGGTAGGACTACTGTTACTCACAAGGAGTGGGAAGCGGAGCTGGCCCCCTAGTGAAGTTCTCACAGCCGTTGGCCTGATATGTGCACTGGCCGGAGGGTTTGCCAAGGCAGACATTGAGATGGCTGGACCCATGGCTGCAGTAGGCTTGCTAATTGTCAGCTATGTGGTCTCGGGAAAGAGTGTGGACATGTACATTGAAAGAGCAGGTGACATCACATGGGAAAAGGACGCGGAAGTCACTGGAAACAGTCCTCGGCTTGACGTGGCACTGGATGAGAGTGGTGATTTCTCCTTGGTAGAGGAAGATGGTCCACCCATGAGAGAGATCATACTCAAGGTGGTCCTGATGGCCATCTGTGGCATGAACCCAATAGCTATACCTTTTGCTGCAGGAGCGTGGTATGTGTATGTGAAGACTGGGAAAAGGAGTGGCGCCCTCTGGGACGTGCCTGCTCCCAAAGAAGTGAAGAAAGGAGAGACCACAGATGGAGTGTACAGAGTGATGACTCGCAGACTGCTAGGTTCAACACAGGTTGGAGTGGGAGTCATGCAAGAGGGAGTCTTCCACACCATGTGGCACGTTACAAAAGGAGCCGCACTGAGGAGCGGTGAGGGAAGACTTGATCCATACTGGGGGGATGTCAAGCAGGACTTGGTGTCATACTGTGGGCCTTGGAAGTTGGATGCAGCTTGGGATGGACTCAGCGAGGTACAGCTTTTGGCCGTACCTCCCGGAGAGAGGGCCAGAAACATTCAGACCCTGCCTGGAATATTCAAGACAAAGGACGGGGACATCGGAGCAGTTGCTCTGGACTACCCTGCAGGGACCTCAGGATCTCCGATCCTAGACAAATGTGGAAGAGTGATAGGACTCTATGGCAATGGGGTTGTGATCAAGAATGGAAGCTATGTTAGTGCTATAACCCAGGGAAAGAGGGAGGAGGAGACTCCGGTTGAATGTTTCGAACCCTCGATGCTGAAGAAGAAGCAGCTAACTGTCTTGGATCTGCATCCAGGAGCCGGAAAAACCAGGAGAGTTCTTCCTGAAATAGTCCGTGAAGCCATAAAAAAGAGACTCCGGACAGTGATCTTGGCACCAACTAGGGTTGTCGCTGCTGAGATGGAGGAGGCCTTGAGAGGACTTCCGGTGCGTTACATGACAACAGCAGTCAACGTCACCCATTCTGGGACAGAAATCGTTGATTTGATGTGCCATGCCACTTTCACTTCACGCTTACTACAACCCATCAGAGTCCCTAATTACAATCTCTACATCATGGATGAAGCCCACTTCACAGACCCCTCAAGTATAGCTGCAAGAGGATACATATCAACAAGGGTTGAAATGGGCGAGGCGGCTGCCATTTTTATGACTGCCACACCACCAGGAACCCGTGATGCGTTTCCTGACTCTAACTCACCAATCATGGACACAGAAGTGGAAGTCCCAGAGAGAGCCTGGAGCTCAGGCTTTGATTGGGTGACAGACCATTCTGGGAAAACAGTTTGGTTCGTTCCAAGCGTGAGAAACGGAAATGAAATCGCAGCCTGTCTGACAAAGGCTGGAAAGCGGGTCATACAGCTCAGCAGGAAGACTTTTGAGACAGAATTTCAGAAAACAAAAAATCAAGAGTGGGACTTTGTCATAACAACTGACATCTCAGAGATGGGCGCCAACTTCAAGGCTGACCGGGTCATAGACTCTAGGAGATGCCTAAAACCAGTCATACTTGATGGTGAGAGAGTCATCTTGGCTGGGCCCATGCCTGTCACGCATGCTAGTGCTGCTCAGAGGAGAGGACGTATAGGCAGGAACCCTAACAAACCTGGAGATGAGTACATGTATGGAGGTGGGTGTGCAGAGACTGATGAAGGCCATGCACACTGGCTTGAAGCAAGAATGCTTCTTGACAACATCTACCTCCAGGATGGCCTCATAGCCTCGCTCTATCGGCCTGAGGCCGATAAGGTAGCCGCCATTGAGGGAGAGTTTAAGCTGAGGACAGAGCAAAGGAAGACCTTCGTGGAACTCATGAAGAGAGGAGACCTTCCCGTCTGGCTAGCCTATCAGGTTGCATCTGCCGGAATAACTTACACAGACAGAAGATGGTGCTTTGATGGCACAACCAACAACACCATAATGGAAGACAGCGTACCAGCAGAGGTGTGGACAAAGTATGGAGAGAAGAGAGTGCTCAAACCGAGATGGATGGATGCTAGGGTCTGTTCAGACCATGCGGCCCTGAAGTCGTTCAAAGAATTCGCCGCTGGAAAAAGAGGAGCGGCTTTGGGAGTAATGGAGGCCCTGGGAACACTGCCAGGACACATGACAGAGAGGTTTCAGGAAGCCATTGACAACCTCGCCGTGCTCATGCGAGCAGAGACTGGAAGCAGGCCTTATAAGGCAGCGGCAGCCCAACTGCCGGAGACCCTAGAGACCATTATGCTCTTAGGTTTGCTGGGAACAGTTTCACTGGGGATCTTCTTCGTCTTGATGCGGAATAAGGGCATCGGGAAGATGGGCTTTGGAATGGTAACCCTTGGGGCCAGTGCATGGCTCATGTGGCTTTCGGAAATTGAACCAGCCAGAATTGCATGTGTCCTCATTGTTGTGTTTTTATTACTGGTGGTGCTCATACCCGAGCCAGAGAAGCAAAGATCTCCCCAAGATAACCAGATGGCAATTATCATCATGGTGGCAGTGGGCCTTCTAGGTTTGATAACTGCAAACGAACTTGGATGGCTGGAAAGAACAAAAAATGACATAGCTCATCTAATGGGAAGGAGAGAAGAAGGAGCAACCATGGGATTCTCAATGGACATTGATCTGCGGCCAGCCTCCGCCTGGGCTATCTATGCCGCATTGACAACTCTCATCACCCCAGCTGTCCAACATGCGGTAACCACTTCATACAACAACTACTCCTTAATGGCGATGGCCACACAAGCTGGAGTGCTGTTTGGCATGGGCAAAGGGATGCCATTTTATGCATGGG-ACCTTGGAGTCCCGCTGCTAATGATGGGTTGCTATTCACAATTAACACCCCTGACTCTGATAGTAGCTATCATTCTGCTTGTGGCGCACTACATGTACTTGATCCCAGGCCTACAAGCGGCAGCAGCGCGTGCTGCCCAGAAAAGGACAGCAGCTGGCATCATGAAGAATCCCGTTGTGGATGGAATAGTGGTAACTGACATTGACACAATGACAATAGACCCCCAGGTGGAGAAGAAGATGGGACAAGTGTTACTCATAGCAGTAGCCATCTCCAGTGCTGTGCTGCTGCGGACCGCCTGGGGATGGGGGGAGGCTGGAGCTCTGATCACAGCAGCGACCTCCACCTTGTGGGAAGGCTCTCCAAACAAATACTGGAACTCCTCTACAGCCACCTCACTGTGCAACATCTTCAGAGGAAGCTATCTGGCAGGAGCTTCCCTTATCTATACAGTGACGAGAAACGCTGGCCTGGTTAAGAGACGTGGAGGTGGGACGGGAGAGACTCTGGGAGAGAAGTGGAAAGCTCGTCTGAATCAGATGTCGGCCCTGGAGTTCTACTCTTATAAAAAGTCAGGTATCACTGAAGTGTGTAGAGAGGAGGCTCGCCGTGCCCTCAAGGATGGAGTGGCCACAGGAGGACATGCCGTATCCCGGGGAAGTGCAAAGCTCAGATGGTTGGTGGAGAGAGGATATCTGCAGCCCTATGGGAAGGTTGTTGACCTCGGATGTGGCAGAGGGGGCTGGAGCTATTATGCCGCCACCATCCGCAAAGTGCAGGAGGTGAGAGGATACACAAAGGGAGGTCCCGGTCATGAAGAACCCATGCTGGTGCAAAGCTATGGGTGGAACATAGTTCGTCTCAAGAGTGGAGTGGACGTCTTCCACATGGCGGCTGAGCCGTGTGACACTCTGCTGTGTGACATAGGTGAGTCATCATCTAGTCCTGAAGTGGAAGAGACACGAACACTCAGAGTGCTCTCTATGGTGGGGGACTGGCTTGAAAAAAGACCAGGGGCCTTCTGTATAAAGGTGCTGTGCCCATACACCAGCACTATGATGGAAACCATGGAGCGACTGCAACGTAGGCATGGGGGAGGATTAGTCAGAGTGCCATTGTCTCGCAACTCCACACATGAGATGTACTGGGTCTCTGGGGCAAAGAGCAACATCATAAAAAGTGTGTCCACCACAAGTCAGCTCCTCCTGGGACGCATGGATGGCCCCAGGAGGCCAGTGAAATATGAGGAGGATGTGAACCTCGGCTCGGGTACACGAGCTGTGGCAAGCTGTGCTGAGGCTCCTAACATGAAAATCATCGGCAGGCGCATTGAGAGAATCCGCAATGAACATGCAGAAACATGGTTTCTTGATGAAAACCACCCATACAGGACATGGGCCTACCATGGGAGCTACGAAGCCCCCACGCAAGGATCAGCGTCTTCCCTCGTGAACGGGGTTGTTAGACTCCTGTCAAAGCCTTGGGACGTGGTGACTGGAGTTACAGGAATAGCCATGACTGACACCACACCATACGGCCAACAAAGAGTCTTCAAAGAAAAAGTGGACACCAGGGTGCCAGATCCCCAAGAAGGCACTCGCCAGGTAATGAACATAGTCTCTTCCTGGCTGTGGAAGGAGCTGGGGAAACGCAAGCGGCCACGCGTCTGCACCAAAGAAGAGTTTATCAACAAGGTGCGCAGCAATGCAGCACTGGGAGCAATATTTGAAGAGGAAAAAGAATGGAAGACGGCTGTGGAAGCTGTGAATGATCCAAGGTTTTGGGCCCTAGTGGATAGGGAGAGAGAACACCACCTGAGAGGAGAGTGTCACAGCTGTGTGTACAACATGATGGGAAAAAGAGAAAAGAAGCAAGGAGAGTTCGGGAAAGCAAAAGGTAGCCGCGCCATCTGGTACATGTGGTTGGGAGCCAGATTCTTGGAGTTTGAAGCCCTTGGATTCTTGAACGAGGACCATTGGATGGGAAGAGAAAACTCAGGAGGTGGAGTCGAAGGGTTAGGATTGCAAAGACTTGGATACATTCTAGAAGAAATGAATCGGGCACCAGGAGGAAAGATGTACGCAGATGACACTGCTGGCTGGGACACCCGCATTAGTAAGTTTGATCTGGAGAATGAAGCTCTGATTACCAACCAAATGGAGGAAGGGCACAGAACTCTGGCGTTGGCCGTGATTAAATACACATACCAAAACAAAGTGGTGAAGGTTCTCAGACCAGCTGAAGGAGGAAAAACAGTTATGGACATCATTTCAAGACAAGACCAGAGAGGGAGTGGACAAGTTGTCACTTATGCTCTCAACACATTCACCAACTTGGTGGTGCAGCTTATCCGGAACATGGAAGCTGAGGAAGTGTTAGAGATGCAAGACTTATGGTTGTTGAGGAAGCCAGAGAAAGTGACCAGATGGTTGCAGAGCAATGGATGGGATAGACTCAAACGAATGGCGGTCAGTGGAGATGACTGCGTTGTGAAGCCAATCGATGATAGGTTTGCACATGCCCTCAGGTTCTTGAATGACATGGGAAAAGTTAGGAAAGACACACAGGAGTGGAAACCCTCGACTGGATGGAGCAATTGGGAAGAAGTCCCGTTCTGCTCCCACCACTTCAACAAGCTGTACCTCAAGGATGGGAGATCCATTGTGGTCCCTTGCCGCCACCAAGATGAACTGATTGGCCGAGCTCGCGTCTCACCAGGGGCAGGATGGAGCATCCGGGAGACTGCCTGTCTTGCAAAATCATATGCGCAGATGTGGCAGCTCCTTTATTTCCACAGAAGAGACCTTCGACTGATGGCTAATGCCATTTGCTCGGCTGTGCCAGTTGACTGGGTACCAACTGGGAGAACCACCTGGTCAATCCATGGAAAGGGAGAATGGATGACCACTGAGGACATGCTCATGGTGTGGAATAGAGTGTGGATTGAGGAGAACGACCATATGGAGGACAAGACTCCTGTAACAAAATGGACAGACATTCCCTATCTAGGAAAAAGGGAGGACTTATGGTGTGGATCCCTTATAGGGCACAGACCCCGCACCACTTGGGCTGAAAACATCAAAGACACAGTCAACATGGTGCGCAGGATCATAGGTGATGAAGAAAAGTACATGGACTATCTATCCACCCAAGTCCGCTACTTGGGTGAGGAAGGGTCCACACCCGGAGTGTTG---

>ENA|AEN75266|AEN75266.1 Zika virus partial polyprotein

ATGAAAAACCC-AAAGAGGAAATCCGGAGGATTCCGGATTGTCAATATGCTAAAACGCGGAGTAGCCCGTGTAAACCCCTTGGGGGGTTTGAAGAGGCTGCCGGCCGGACTTCTGCTGGGCCATGGACCCATCAGAATGGTTTTGGCGATACTAGCCTTCTTGAGATTCACAGCAATCAAGCCATCACTGGGCCTCATCAATAGATGGGGTTCCGTGGGGAAGAAGGAGGCTATGGAAATAATAAAAAAGTTCAAGAAAGACCTTGCTGCCATGTTGAGAATCATCAATGCTAGGAAGGAGAGGAAGAGACGTGGCGCTGACACCAGCATCGGAATCGTCGGCCTCCTGCTGACCACAGCCATGGCAGCCGAGATCACTAGACGTGGGAGTGCATACTACATGTACTTGGACAGGAGCGATGCTGGTAAGGCCATTTCTTTTGCCACCACATTGGGGGTGAACAAATGCCATGTACAGATCATGGACCTCGGGCACATGTGTGACGCCACCATGAGTTATGAGTGCCCCATGCTAGACGAGGGAGTGGAGCCAGATGACGTCGATTGCTGGTGCAACACGACATCGACTTGGGTTGTGTACGGAACCTGTCATCATAAAAAAGGTGAAGCACGACGATCCAGAAGAGCCGTGACGCTTCCTTCTCACTCCACGAGGAAGCTGCAAACGCGATCGCAGACTTGGCTAGAGTCAAGAGAATACACAAAGCACCTGATCAAGGTTGAAAATTGGATATTTAGGAACCCCGGGTTTGCGCTAGTGGCTGTAGCTATAGCCTGGCTCCTGGGAAGCTCGACGAGCCAAAAAGTTATATACTTGGTCATGATATTGTTGATTGCCCCGGCATACAGCATTAGGTGCATAGGAGTTAGCAATAGAGACTTCGTGGAGGGCATGTCAGGTGGGACCTGGGTTGATGTTGTCTTGGAACATGGGGGTTGCGTCACCGTGATGGCACAGGACAAGCCAACAGTTGATATCGAGTTGGTCACGACAACGGTTAGCAACATGGCCGAGGTAAGATCCTACTGCTATGAGGCATCAATATCGGACATGGCTTCGGACAGCCGTTGTCCAACACAAGGTGAAGCCTACCTTGACAAGCAGTCAGACACTCAATATGTCTGCAAGAGAACATTGGTGGATAGAGGTTGGGGAAATGGGTGTGGACTTTTTGGCAAAGGGAGCTTGGTGACATGTGCCAAGTTTACGTGCTCCAAGAAAATGACAGGCAAGAGCATCCAGCCGGAGAACTTGGAGTACCGGATAATGCTATCAGTGCATGGATCCCAGCACAGTGGGATGATTGTGAATGACACAGGACATGAAACTGACGAAAACAGAGCAAAAGTCGAGGTCACACCCAATTCACCAAGAGCAGAAGCAACCTTGGGAGGTTTTGGAAGCTTGGGACTTGACTGTGAACCAAGGACAGGCCTTGACTTCTCAGATCTGTATTACCTGACCATGAACAATAAGCATTGGTTGGTGCACAAGGAGTGGTTTCATGACATCCCATTACCTTGGCATGCTGGTGCAGACACTGGAACTCCACACTGGAACAACAAAGAGGCATTGGTGGAGTTCAAGGACGCCCACGCCAAGAGGCAAACTGTTGTGGTTCTGGGGAGCCAAGAGGGAGCTGTTCATACGGCCCTCGCTGGAGCTTTGGAGGCTGAGATGGATGGTGCAAAGGGAAGGCTATTCTCTGGCCATTTGAAATGCCGCCTAAAAATGGACAAGCTTAGGTTGAAGGGTGTGTCATATTCCCTGTGTACCGCAGCGTTCACATTTACCAAGGTCCCAGCTGAAACATTGCATGGAACAGTCACAGTGGAGGTGCAGTATGCAGGGACAGACGGACCCTGCAAAGTCCCAGCCCAGATGGCGGTGGACATGCAGACCCTGACCCCAGTTGGAAGGCTGATAACCGCCAATCCTGTGATCACTGAAAGTACTGAGAATTCAAAGATGATGTTGGAGCTCGACCCACCATTTGGGGATTCTTACATTGTCATAGGAGTCGGGGACAAGAAAATCACCCATCACTGGCATCGGAGTGGTAGCACCATCGGAAAGGCATTTGAAGCCACTGTGAGAGGTGCCAAGAGAATGGCAGTCTTGGGGGACACAGCCTGGGACTTTGGATCAGTTGGGGGTGTGTTTAATTCATTGGGTAAGGGTATTCACCAGATCTTTGGAGCAGCTTTCAAATCACTGTTTGGAGGAATGTCCTGGTTCTCACAGATCCTCATAGGCACACTGTTGGTGTGGTTAGGTTTGAACACAAAGAATGGATCTATCTCCCTCACATGCTTAGCCCTGGGGGGAGTGATGATCTTCCTTTCCACGGCTGTTTCTGCTGATGTTGGGTGCTCGGTGGACTTCTCAAAAAAGGAAACGAGATGTGGCACGGGGGTGTTCGTCTACAATGACGTTGAAGCCTGGAGGGACCGGTACAAGTACCATCCTGACTCCCCCCGCAGATTGGCAGCAGCTGTTAAGCAGGCTTGGGAAGAGGGGATTTGTGGGATCTCCTCCGTTTCGAGAATGGAAAACATCATGTGGAAATCAGTGGAAGGGGAGCTTAATGCAATCCTAGAGGAGAATGGAGTTCAACTGACAGTTGTAGTGGGGTCTGTAAAAAACCCTATGTGGAGAGGTCCACAGAGATTGCCAGTGCCTGTGAATGAGCTGCCCCATGGCTGGAAAGCCTGGGGGAAATCGTACTTTGTCAGAGCAGCAAAGACCAACAACAGTTTTGTTGTCGACGGTGACACACTGAAGGAGTGTCCGCTCAAACATAGAGCATGGAATAGCTTCCTTGTGGAGGATCACGGGTTTGGGATCTTCCACACCAGTGTTTGGCTGAAGGTCAGAGAGGACTACTCACTAGAGTGTGACCCAGCCGTCATAGGAACAGCTGTCAAGGGAAAGGAAGCTGCACACAGTGATCTAGGCTATTGGATTGAGAGTGAAAAGAATGACACATGGAGGCTGAGGAGGGCTCATCTGATTGAGATGAAAACATGTGAGTGGCCAAAGTCTCACACACTGTGGACAGATGGAGTGGAAGAAAGTGATCTGATCATACCCAAGTCCTTAGCTGGTCCACTCAGCCACCACAACACCAGAGAGGGTTACAGAACTCAATTGAAAGGGCCATGGCACAGTGAAGAGCTCGAAATCCGGTTTGAGGAGTGCCCAGGAACCAAGGTTCACGTGGAGGAGACATGCGGAACTAGAGGACCATCTCTGAGATCGACTACTGCAAGTGGAAGGGTCATAGAGGAATGGTGCTGTAGGGAATGCACAATGCCTCCACTATCGTTCCGGGCAAAAGACGGCTGCTGGTATGGAATGGAGATAAGGCCCAGAAAGGAACCAGAGAGCAACTTAGTAAGGTCTATGGTGACAGCAGGATCAACCGATCATATGGATCACTTCTCTCTTGGAGTGCTTGTGATTCTACTCATGGTGCAGGAAGGTTTGAAGAAGAGAATGACCACAAAGATCATAATGAGCACATCAATGGCAGTGCTGGTAGCCATGGTCTTGGGAGGATTCTCAATGAGTGACCTGGCTAAGCTCGTGATCCTGATGGGTGCCACTTTCGCAGAAATGAACACTGGAGGAGATGTGGCTCACTTGGCATTGGTTGCGGCATTTAAAGTCAGACCAGCCTTGTTGGTTTCCTTCATCTTCAGAGCCAACTGGACACCCCGTGAGAGCATGCTGCTAGCCCTGGCTTCGTGCCTCCTGCAGACTGCGATCTCTGCTCTTGAAGGCGAGCTGATGGTCCTCGTTAATGGATTTGCTTTGGCCTGGTTGGCAATACGAGCAATGGCCGTGCCACGCACTGACAATATCGCTCTGGCAATTCTGGCTGCTCTAACACCATTAGCCAGAGGCACACTGCTCGTGGCATGGAGAGCGGGCCTCGCCACTTGTGGAGGGTTCATGCTCCTTTCCCTGAAAGGGAAAGGTAGTGTGAAGAAGAACCTGCCTTTTGTCATGGCCTTGGGGTTGACCGCTGTGAGGATAGTGGACCCCATCAATGTGGTAGGACTACTGTTACTCACAAGGAGTGGAAAACGGAGCTGGCCTCCTAGTGAAGTGCTTACAGCTGTTGGCCTGATATGTGCACTGGCCGGAGGGTTTGCCAAGGCAGACATAGAGATGGCTGGGCCCATGGCTGCAGTAGGCCTGCTAATTGTCAGTTATGTGGTCTCGGGAAAGAGTGTGGACATGTACATCGAAAGAGCAGGTGATATCACATGGGAAAAAGACGCGGAAGTCACTGGAAACAGTCCCCGGCTTGACGTGGCACTGGATGAGAGTGGTGATTTCTCCTTGGTAGAGGAGGATGGCCCACCCATGAGAGAGATCATACTCAAGGTGGTCCTGATGGCCATCTGTGGCATGAACCCAATAGCCATACCCTTCGCTGCAGGAGCGTGGTATGTGTATGTAAAGACTGGAAAAAGGAGTGGTGCCCTCTGGGACGTGCCTGCTCCCAAAGAAGTGAAAAAAGGAGAGACTACAGATGGAGTGTACAGAGTGATGACTCGCAGACTGCTGGGTTCAACACAGGTTGGAGTGGGGGTCATGCAAGAGGGAGTCTTCCACACCATGTGGCACGTCACAAAAGGAGCCGCATTGAGGAGCGGTGAAGGAAGACTTGATCCATATTGGGGGGACGTCAAGCAGGACTTGGTGTCATATTGTGGGCCGTGGAAGTTGGATGCAGCCTGGGATGGACTAAGTGAAGTGCAGCTTTTGGCCGTACCCCCCGGAGAGAGGGCTAGAAACATTCAGACTCTGCCTGGAATATTCAAGACAAAGGATGGGGACATCGGAGCAGTCGCTCTAGACTACCCCGCAGGAACTTCAGGATCTCCAATCCTAGACAAATGCGGAAGAGTGATAGGACTTTATGGCAATGGGGTTGTGATCAAGAATGGAAGCTATGTTAGTGCAATAACCCAGGGAAAAAGGGAGGAGGAGGCTCCAGTTGAGTGCTTTGAACCTTCGATGCTGAGGAAGAAGCAGCTAACAGTCTTGGATCTGCATCCAGGAGCCGGAAAAACCAGGAGGGTCCTTCCTGAAATAGTCCGTGAAGCCATAAAGAAGAGACTTCGCACAGTGATCCTAGCACCAACCAGGGTCGTTGCTGCCGAGATGGAGGAAGCCCTAAGAGGACTTCCGGTGCGTTACATGACAACAGCAGTCAACGTCACCCACTCTGGGACAGAAATCGTCGATTTGATGTGCCATGCCACCTTCACTTCACGCCTACTACAACCCATCAGAGTCCCCAACTACAACCTTTACATCATGGATGAAGCTCATTTCACAGATCCCTCAAGCATAGCTGCACGAGGATATATATCAACAAGGGTTGAAATGGGCGAGGCGGCTGCTATCTTCATGACTGCTACACCACCAGGAACCCGCGATGCGTTTCCGGATTCCAACTCACCAATCATGGACACAGAAGTGGAAGTTCCAGAGAGAGCCTGGAGCTCAGGCTTTGACTGGGTGACGGATCATTCTGGGAAAACAATTTGGTTTGTTCCAAGTGTGAGAAACGGAAATGAAATCGCAGCCTGTCTGACAAAGGCTGGAAAGCGGGTCATACAGCTCAGCAGGAAGACTTTTGAGACAGAGTTTCAGAAGACAAAAAATCAAGAGTGGGACTTTGTCATAACAACTGACATTTCAGAGATGGGTGCCAATTTCAAGGCTGACCGGGTCATAGATTCCAGGAGATGCCTAAAGCCAGTCATACTTGATGGGGAGAGAGTCATCCTGGCTGGGCCCATGCCTGTCACGCATGCCAGTGCTGCTCAGAGGAGAGGACGTGTAGGCAGGAACCCCAACAAACCTGGAGATGAGTACATGTATGGAGGTGGGTGTGCAGAGACTGATGAAGACCATGCACACTGGCTTGAAGCAAGAATGCTTCTTGACAACATTTACCTCCAGGATGGCCTCATAGCCTCGCTCTATCGGCCTGAGGCCGACAAGGTAGCCGCCATTGAGGGAGAGTTCAAGCTGAGGACAGAGCAAAGGAAGACCTTTGTGGAACTCATGAAGAGAGGAGACCTTCCCGTTTGGCTGGCCTATCAAGTAGCATCTGCCGGAATAACTTACACAGATAGAAGATGGTGCTTTGATGGCACTACCAACAACACCATAATGGAAGACAGTGTACCAGCAGAGGTGTGGACCAAGTATGGAGAGAAGAGAGTGCTTAAACCGAGGTGGATGGATGCTAGGGTCTGTTCAGATCATGCGGCTCTGAAGTCGTTCAAAGAATTTGCCGCTGGGAAGAGAGGAGCGGCTTTGGGAGTAATGGATGCCCTGGGAACATTGCCAGGACACATGACAGAGAGGTTTCAGGAAGCCATTGACAATCTCGCTGTGCTCATGCGAGCAGAGACTGGAAGTAGGCCCTACAAAGCAGCGGCAGCTCAACTGCCGGAGACCTTAGAGACTATCATGCTTCTGGGCTTATTGGGAACAGTTTCGCTAGGAATCTTCTTTGTCTTGATGCGGAACAAGGGCATCGGGAAGATGGGCTTCGGAATGGTAACCCTTGGGGCCAGCGCATGGCTCATGTGGCTTTCGGAAATTGAACCAGCCAGAATCGCATGTGTCCTCATTGTCGTGTTTTTGTTGCTGGTGGTGCTCATACCCGAGCCAGAGAAGCAAAGATCTCCCCAGGATAATCAAATGGCAATCATCATCATGGTGGCAGTTGGCCTTCTGGGTTTGATAACTGCAAATGAACTCGGATGGCTGGAAAGGACAAAAAGTGATATAGCTCATCTAATGGGAAGGAAAGAAGAGGGGACAACCATGGGATTCTCAATGGATATTGATCTGCGGCCAGCCTCCGCCTGGGCTATTTATGCCGCATTGACAACTCTCATCACCCCAGCCGTCCAACATGCGGTAACCACCTCATACAACAACTACTCCCTGATGGCGATGGCCACACAAGCTGGAGTGCTGTTTGGCATGGGCAAAGGGATGCCATTTTATGCATGGG-ACTTTGGAGTCCCGCTGCTAATGATGGGTTGCTACTCACAATTAACACCCCTGACCCTGATAGTGGCCATCATTCTGCTTGTGGCACACTACATGTACTTGATCCCAGGTTTGCAGGCAGCAGCAGCACGTGCTGCCCAGAAGAGGACAGCAGCTGGCATCATGAAGAATCCCGTTGTGGATGGAATAGTGGTGACTGACATTGACACAATGACAATTGACCCCCAAGTGGAGAAGAAGATGGGACAAGTGTTACTCATAGCAGTAGCTGTCTCCAGTGCTGTGCTACTGCGGACCGCTTGGGGATGGGGGGAGGCTGGGGCTCTGATCACAGCAGCAACCTCCACCTTATGGGAAGGCTCTCCAAACAAATACTGGAACTCCTCCACAGCCACCTCACTGTGCAACATCTTTAGAGGAAGTTATTTGGCAGGGGCTTCCCTTATTTACACAGTGACAAGAAATGCCGGTCTGGTTAAGAGACGTGGAGGTGGAACGGGAGAGACCCTGGGAGAGAAGTGGAAAGCCCGCCTGAACCAGATGTCGGCCTTGGAGTTCTACTCTTACAAAAAGTCAGGCATCACCGAAGTGTGTAGGGAGGAGGCGCGCCGCGCCCTCAAGGATGGAGTGGCCACAGGAGGACATGCTGTATCCCGGGGAAGCGCAAAGCTTAGATGGTTGGTAGAGAGAGGATACCTGCAGCCCCATGGAAAGGTTGTTGACCTTGGATGTGGCAGAGGAGGCTGGAGTTATTACGCTGCCACCATCCGTAAAGTGCAGGAGGTCAGAGGATACACAAAGGGAGGTCCTGGCCATGAAGAGCCCATGCTGGTGCAAAGCTATGGGTGGAACATAGTTCGCCTCAAGAGTGGAGTGGACGTCTTCCACATGGCGGCTGAACCGTGTGACACTCTGCTGTGTGACATAGGCGAGTCATCATCCAGTCCTGAAGTGGAAGAGACGCGAACACTCAGAGTGCTCTCCATGGTGGGAGACTGGCTTGAAAAAAGACCAGGGGCCTTCTGCATAAAGGTGCTGTGCCCATACACCAGCACAATGATGGAGACCATGGAGCGACTGCAACGTAGGCATGGGGGAGGATTAGTCAGAGTGCCATTGTCCCGCAACTCTACACATGAGATGTATTGGGTCTCTGGAGCCAAAAGTAACATCATAAAGAGTGTGTCCACCACAAGTCAGCTCCTCTTGGGACGCATGGAAGGGCCTAGGAGGCCAGTGAAATATGAGGAGGATGTGAACCTCGGCTCAGGCACACGAGCTGTGGCAAGCTGCGCTGAGGCTCCCAACATGAAGATCATTGGTAGGCGCATTGAGAGAATCCGCAATGAACATGCAGAGACATGGTTCTTTGATGAAAACCACCCATACAGGACATGGGCCTACCATGGGAGCTACGAAGCCCCCACGCAGGGGTCAGCGTCATCCCTCGTGAACGGGGTTGTTAGACTCTTGTCAAAGCCCTGGGATGTGGTGACTGGAGTTACAGGAATAGCTATGACTGACACCACGCCATACGGCCAACAAAGAGTCTTCAAGGAAAAAGTGGACACCAGGGTGCCAGATCCCCAAGAAGGCACTCGCCAAGTGATGAACATGGTATCGTCTTGGCTATGGAAGGAGCTGGGAAAACGCAAGCGACCACGTGTCTGCACCAAAGAAGAGTTCATTAATAAGGTGCGCAGCAATGCAGCACTGGGAGCAATATTTGAAGAGGAAAAAGAATGGAAGACAGCTGTAGAAGCTGTGAATGATCCAAGATTTTGGGCTCTAGTGGACAAGGAAAGAGAACACCACCTGAGAGGAGAGTGTCATAGCTGTGTGTACAACATGATGGGCAAAAGAGAAAAGAAGCAAGGAGAATTCGGGAAAGCAAAAGGCAGCCGCGCAATCTGGTACATGTGGTTGGGAGCCAGATTTTTGGAGTTTGAAGCTCTTGGGTTCTTGAACGAGGACCACTGGATGGGGAGAGAAAACTCAGGAGGTGGCGTTGAAGGGCTAGGACTGCAAAGGCTTGGATATATCCTAGAAGAAATGAACCGGGCACCAGGAGGAAAGATGTATGCAGATGACACTGCTGGCTGGGACACCCGCATTAGCAAGTTTGATCTAGAGAATGAAGCCCTGATCACTAACCAGATGGAAGAAGGGCACAGAGCTCTGGCGTTGGCCGTGATTAAATACACATACCAAAACAAAGTGGTGAAGGTTCTCAGACCAGCTGAAGGAGGGAAAACAGTCATGGACATCATCTCAAGACAAGACCAGAGAGGGAGTGGACAAGTTGTTACTTATGCCCTCAACACATTCACCAACCTGGTGGTGCAGCTTATCCGGAACATGGAGGCTGAGGAAGTGCTAGAGATGCATGATCTGTGGTTGTTGAGGAAACCAGAGAAAGTGACCAGATGGTTGCAGAGTAATGGATGGGACAGACTCAAACGGATGGCAGTTAGTGGAGATGACTGCGTTGTGAAGCCAATTGATGATAGGTTTGCACATGCCCTCAGGTTCTTGAATGACATGGGAAAGGTTAGGAAAGACACACAGGAATGGAAACCCTCGACTGGATGGAGCAATTGGGAAGAGGTCCCGTTCTGTTCTCACCACTTTAACAAGCTGCACCTCAAGGACGGGAGATCCATTGTGGTCCCCTGCCGCCACCAAGATGAACTGATTGGCCGAGCCCGTGTCTCACCAGGGGCAGGATGGAGCATCCGTGAGACTGCTTGCCTTGCAAAATCATATGCACAGATGTGGCAGCTTCTTTATTTCCACAGAAGAGATCTCCGACTGATGGCCAATGCCATTTGTTCGGCTGTGCCAGTTGATTGGGTACCAACTGGGAGAACCACCTGGTCAATCCATGGAAAGGGAGAATGGATGACTACTGAGGACATGCTCATGGTGTGGAATAGAGTGTGGATTGAGGAGAACGACCACATGGAGGACAAGACCCCTGTAACAAAATGGACAGACATTCCCTATTTGGGGAAAAGGGAGGACTTATGGTGTGGATCCCTTATAGGGCACAGACCTCGCACCACTTGGGCTGAGAACATCAAAGACACAGTCAACATGGTGCGTAGGATCATAGGTGATGAAGAAAAGTNCATGGACTACCTATCCACCCAAGTACGTTACTTGGGTGAGGAAGGGTCCACACCTGGAGTGCTG---

>ENA|AEN75264|AEN75264.1 Zika virus partial polyprotein

ATGAAAAACCC-AAAAAAGAAATCCGGAGGATTCCGGATTGTCAATATGCTAAAACGCGGAGTAGCCCGTGTGAGCCCCTTTGGGGGCTTGAAGAGGCTACCAGCTGGACTTCTGCTGGGTCATGGACCCATCAGGATGGTCTTGGCGATACTAGCCTTCTTGAGATTCACGGCAATCAAGCCATCACTGGGTCTCATCAATAGATGGGGTTCCGTGGGGAAAAAAGAGGCTATGGAAATAATAAAGAAGTTCAAGAAAGATCTGGCTGCCATGCTGAGAATAATCAATGCTAGGAAGGAGAAGAAGAGACGTGGCGCAGACACCAGTGTCGGAATTGTTGGCCTCCTGCTGACCACAGCCATGGCAGTGGAGGTCACCAGACGTGGGAGTGCATACTATATGTACTTAGACAGAAGCGATGCTGGGGAGGCCATATCTTTTCCAACCACACTGGGGGTGAATAAGTGTTACATACAGATCATGGATCTTGGACACATGTGTGATGCCACAATGAGCTATGAATGCCCTATGTTGGATGAGGGGGTAGAACCAGATGACGTCGATTGCTGGTGCAACACGACATCGACTTGGGTTGTGTACGGAACCTGCCATCACAAAAAAGGTGAGGCACGGAGATCTAGAAGAGCTGTGACGCTCCCCTCTCATTCCACTAGGAAGCTGCAAACGCGGTCGCAGACCTGGTTGGAATCAAGAGAATACACAAAGCACTTGATCAGAGTCGAAAATTGGATATTCAGGAACCCTGGCTTTGCGTTGGCAGCAGCTGCCATTGCTTGGCTTTTGGGAAGCTCAACGAGCCAAAAAGTCATATACTTGGTCATGATACTGTTGATTGCCCCGGCATACAGTATCAGGTGCATAGGAGTCAGCAATAGGGATTTTGTGGAAGGTATGTCAGGTGGGACCTGGGTTGATGTTGTCTTGGAACATGGAGGTTGTGTTACCGTAATGGCACAGGACAAGCCAACTGTTGATATAGAGTTGGTCACAACAACGGTTAGCAACATGGCGGAGGTAAGATCCTACTGCTACGAGGCATCAATATCGGACATGGCTTCGGACAGCCGCTGCCCAACACAAGGTGAAGCCTACCTTGACAAGCAGTCAGACACTCAATATGTTTGCAAAAGAACGTTAGTGGACAGAGGTTGGGGAAATGGATGTGGACTCTTTGGCAAAGGGAGCCTGGTGACATGCGCCAAGTTTGCATGCTCCAAGAAAATGACTGGGAAGAGCATCCAGCCAGAGAACCTGGAGTACCGGATAATGCTGTCAGTTCATGGCTCCCAGCACAGTGGGATGATTGTTAATGACANAGGACATGAAACTGATGAGAATAGAGCGAAGGTTGAGATAACGCCCAATTCACCAAGAGCCGAAGCCACCCTGGGAGGTTTTGGAAGCCTAGGACTTGATTGTGAACCGAGGACAGGCCTTGACTTTTCAGATTTGTATTACTTGACTATGAATAACAAGCATTGGTTGGTGCACAAGGAGTGGTTCCATGACATTCCACTACCTTGGCATGCTGGGGCAGACACCGGAACTCCACATTGGAACAACAAAGAAGCATTGGTAGAGTTCAAGGACGCACATGCCAAAAGGCAAACTGTCGTGGTTCTAGGGAGTCAAGAAGGAGCCGTTCACACGGCTCTTGCTGGAGCCCTGGAGGCTGAGATGGATGGTGCAAAGGGAAGGCTGTCCTCTGGCCACTTGAAATGTCGCTTGAAAATGGACAAACTTAGATTGAAGGGCGTGTCATACTCCTTATGTACCGCGGCGTTCACATTCACCAAGATCCCGGCTGAAACGCTGCATGGGACAGTCACAGTGGAGGTACAGTATGCAGGGACAGATGGACCCTGCAAGGTTCCAGCTCAGATGGCGGTGGATATGCAAACTCTGACCCCAGTTGGGAGGTTGATAACCGCTAACCCTGTGATCACTGAAAGCACTGAGAATTCAAAGATGATGTTGGAACTTGACCCACCATTTGGGGATTCTTACATTGTCATAGGAGTTGGGGATAAGAAGATCACCCACCACTGGNACAGGAGTGGCAGCACCATCGGAAAAGCATTTGAAGCCACTGTGAGAGGCGCCAAGAGAATGGCAGTCTTGGGAGACACAGCCTGGGACTTTGGATCAGTCGGAGGTGCTCTCAACTCATTGGGCAAGGGCATCCATCAAATTTTTGGAGCAGCTTTCAAATCATTGTTTGGAGGAATGTCCTGGTTCTCACAAATCCTCATAGGAACGTTGCTGGTGTGGTTGGGTCTGAACACAAAGAATGGATCTATTTCCCTTACGTGCTTGGCCTTAGGGGGAGTGTTGATCTTCCTATCTACAGCCGTCTCTGCTGATGTGGGGTGTTCGGTGGACTTCTCAAAGAAGGAAACGAGATGCGGTACGGGGGTGTTCGTCTATAACGACGTTGAAGCCTGGAGGGACAGGTACAAGTACCATCCTGACTCCCCTCGTAGATTGGCAGCAGCAGTCAAGCAGGCCTGGGAAGATGGGATCTGTGGGATCTCCTCTGTTTCAAGAATGGAAAACATTATGTGGAGATCAGTAGAAGGGGAGCTCAACGCAATTCTGGAAGAGAATGGAGTTCAACTGACGGTCGTTGTGGGATCTGTAAAAAACCCCATGTGGAGAGGTCCGCAGAGGTTGCCTGTGCCTGTGAATGAGCTGCCCCACGGTTGGAAGGCCTGGGGGAAATCGTACTTTGTCAGGGCAGCAAAGACCAACAACAGCTTTGTTGTGGATGGTGACACACTGAAGGAATGCCCGCTCAAACACAGAGCATGGAACAGCTTTCTTGTGGAGGATCACGGGTTCGGGGTATTTCACACTAGTGTCTGGCTTAAAGTCAGAGAGGATTACTCATTAGAGTGTGATCCAGCCGTCATAGGAACAGCTGCTAAGGGAAAGGAGGCCGTGCACAGTGATCTAGGCTACTGGATTGAGAGTGAAAAGAACGACACATGGAGGCTGAAGAGGGCTCACCTGATCGAGATGAAAACATGTGAATGGCCAAAGTCCCACACACTGTGGACAGATGGAATAGAAGAAAGTGATCTGATCATACCTAAGTCTTTAGCTGGGCCACTCAGCCACCACAACACCAGAGAGGGCTACAGGACTCAAGTGAAAGGGCCGTGGCATAGTGAAGAGCTTGAAATCCGGTTTGAGGAATGTCCAGGCACCAAGGTCCACGTGGAGGAAACATGTGGAACGAGAGGACCGTCCCTGAGATCAACCACTGCAAGCGGAAGGGTGATCGAGGAATGGTGCTGCAGGGAATGCACAATGCCCCCATTGTCGTTCCGGGCAAAAGATGGCTGTTGGTATGGAATGGAGATAAGGCCCAGGAAGGAACCAGAGAGTAACCTAGTAAGGTCAATGGTGACTGCAGGATCAACTGATCACATGGATCACTTCTCCCTTGGAGTGCTTGTGATTCTGCTCATGGTGCAGGAAGGGCTGAAGAAGAGAATGACCACAAAGATCATCATAAGCACATCAATGGCAGTGTTGGTAGCTATGATCCTGGGAGGATTTTCAATGAGTGACTTGGCTAAGCTTGCAATTCTGATGGGTGCCACCTTCGCGGAAATGAACACTGGAGGAGATGTAGCTCATCTGGCGCTGATAGCGGCATTCAAAGTCAGACCCGCGTTGCTGGTCTCTTTCATCTTCAGAGCCAATTGGACACCCCGTGAGAGCATGCTGCTGGCCTTGGCCTCGTGCCTTCTGCAAACTGNGATCTCCGCCCTGGAAGGCGACCTGATGGTTCTCATCAATGGTTTTGCTTTGGCCTGGTTGGCAATACGAGCGATGGCTGTTCCACGCACTGACAACATCACCTTGGCAATCCTGGCTGCTCTGACACCACTGGCCCGAGGCACACTGCTTGTAGCGTGGAGAGCAGGCCTTGCTACTTGTGGGGGGTTCATGCTCCTCTCTCTGAAGGGGAAAGGTAGTGTGAAGAAGAACCTACCATTTGTCATGGCCTTGGGACTAACCGCTGTGAGGCTGGTTGACCCCATCAACGTGGTGGGACTGCTGTTGCTCACAAGGAGTGGGAAGCGGAGCTGGCCCCCTAGTGAAGTACTCACAGCTGTTGGCCTGATATGTGCACTGGCCGGAGGGTTCGCCAAAGCAGATATAGAGATGGCTGGGCCCATGGCTGCAGTTGGCCTGCTAATTGTTAGTTACGTGGTCTCAGGAAAGAGTGTGGACATGTACATTGAAAGAGCAGGTGACATCACATGGGAAAAAGATGCGGAAGTTACTGGAAACAGCCCCCGGCTCGATGTGGCACTAGATGAGAGTGGTGATTTCTCCCTGGTGGAGGATGATGGTCCCCCCATGAGAGAGATCATACTCAAGGTGGTCCTGATGACCATCTGTGGCATGAACCCAATAGCCATACCCTTTGCAGCTGGAGCGTGGTATGTGTATGTGAAGACTGGAAAGAGGAGTGGTGCTCTATGGGATGTGCCTGCTCCCAAGGAAGTAAAAAAGGGGGAGACCACAGATGGAGTGTATAGAGTGATGACTCGCAGACTGCTAGGTTCAACACAAGTTGGAGTGGGAGTCATGCAAGAGGGGGTCTTCCACACTATGTGGCACGTCACAAAAGGATCCGCGCTGAGGAGCGGTGAAGGGAGACTTGATCCATACTGGGGAGATGTTAAGCAGGATCTGGTGTCATACTGTGGCCCGTGGAAGCTAGATGCCGCTTGGGACGGACACAGCGAGGTGCAGCTTTTGGCCGTGCCCCCCGGAGAGAGAGCGAGGAACATCCAGACTCTGCCCGGAATATTCAAGACAAAGGATGGGGACATCGGAGCAGTTGCTCTGGACTACCCAGCAGGAACTTCAGGATCTCCGATCCTAGACAAGTGTGGGAGAGTGATAGGACTCTATGGCAATGGGGTCGTGATCAAAAATGGAAGTTATGTTAGTGCCATCACCCAAGGGAGGAGGGAGGAAGAGACTCCTGTTGAATGCTTCGAACCTTCGATGCTGAAGAAGAAGCAGCTAACTGTCTTGGATCTGCATCCTGGAGCTGGGAAAACCAGGAGAGTTCTTCCTGAAATAGTCCGTGAAGCCATAAAAACAAGACTCCGCACGGTGATCCTGGCTCCAACCAGGGTTGTCGCTGCTGAAATGGAGGAAGCCCTTAGAGGGCTTCCAGTGCGTTACATGACAACAGCAGTTAATGTCACCCACTCTGGGACAGAAATCGTTGATTTAATGTGCCATGCCACCTTCACTTCACGCCTACTACAACCCATTAGAGTCCCCAACTACAATCTTTACATTATGGATGAGGCCCACTTCACAGATCCCTCAAGTATAGCAGCAAGAGGATACATATCAACAAGGGTTGAGATGGGCGAGGCGGCTGCCATCTTCATGACCGCCACACCACCAGGAACCCGCGACGCATTTCCGGACTCTAACTCACCAATCATGGACACAGAAGTGGAAGTCCCAGAGAGAGCCTGGAGCTCAGGCTTTGATTGGGTGACGGATCATTCTGGAAAAACAGTTTGGTTTGTTCCAAGCGTGAGGAACGGCAACGAGATCGCGGCTTGTCTGACAAAAGCTGGAAAACGGGTCATACAGCTCAGCAGAAAGACTTTTGAGACAGAGTTCCAGAAAACAAAAAATCAAGAGTGGGACTTCGTCGTAACAACTGACATCTCAGAGATGGGCGCCAACTTCAAAGCTGACCGGGTCATAGATTCCAGGAGATGCCTGAAGCCGGTCATACTTGATGGCGAGAGAGTCATTCTGGCTGGACCCATGCCTGTCACACATGCCAGCGCTGCCCAGAGGAGGGGGCGCATAGGCAGGAATCCCAACAAACCTGGAGATGAGTATATGTATGGAGGTGGGTGCGCAGAGACTGATGAAGACCATGCACACTGGCTTGAAGCAAGAATGCTTCTTGATAACATTTACCTCCAAGATGGCCTCATAGCCTCGCTCTATCGACCTGAGGCCGATAAGGTAGCAGCCATTGAGGGAGAGTTCAAGCTTAGGACGGAGCAAAGGAAGACCTTTGTGGAACTCATGAAAAGAGGAGATCTTCCTGTTTGGCTGGCCTATCAGGTTGCATCTGCCGGAATAACCTACACAGATAGAAGATGGTGTTTTGATGGCACGACCAACAACACCATAATGGAAGACAGTGTGCCGGCAGAGGTGTGGACCAGATACGGAGAGAAAAGAGTGCTCAAACCGAGGTGGATGGACGCCAGAGTTTGTTCAGATCATGCGGCCCTGAAGTCATTCAAAGAATTTGCCGCTGGGAAAAGAGGAGCGGCCTTTGGAGTGATGGAAGCCCTGGGAACACTGCCAGGACACATGACAGAGAGGTTTCAGGAAGCCATTGACAACCTCGCTGTGCTCATGCGGGCAGAGACTGGAAGCAGGCCCTACAAAGCCGCGGCGGCCCAATTACCGGAGACCTTAGAGACCATCATGCTTTTGGGTTTGCTGGGAACAGTCTCGCTGGGAATCTTCTTTGTCTTGATGCGGAACAAGGGCATAGGGAAGATGGGCTTTGGAATGGTGACCCTTGGGGCCAGTGCATGGCTTATGTGGCTCTCGGAAATTGAGCCAGCCAGAATTGCATGTGTCCTCATTGTCGTGTTTCTATTGCTGGTGGTGCTCATACCTGAGCCAGAAAAGCAGAGATCTCCCCAGGACAACCAAATGGCAATTATCATCATGGTAGCAGTGGGTCTTCTGGGCTTGATAACCGCCAATGAACTCGGATGGTTGGAGAGAACAAAAAGTGACCTAGGCCATCTAATGGGAAGGAGAGAGGAGGGGGCAACCATGGGATTCTCAATGGACATTGACTTGCGGCCAGCCTCAGCTTGGGCTATCTATGCCGCTCTGACAACTCTCATCACCCCAGCCGTCCAACATGCGGTAACCACTTCATACAACAACTACTCCTTAATGGCGATGGCCACGCAAGCCGGAGTGTTGTTTGGCATGGGCAAAGGGATGCCATTCTATGCGTGGG-ACTTCGGAGTCCCGCTGCTAATGATGGGTTGCTACTCACAATTAACACCCTTGACCTTAATAGTGGCCATCATTCTGCTCGTGGCGCACTACATGTACTTGATCCCAGGTCTACAGGCAGCAGCGGCGCGCGCTGCCCAGAAGAGAACGGCAGCTGGCATCATGAAGAACCCTGTTGTGGATGGAATAGTGGTGACTGACATTGACACAATGACAATTGACCCCCAAGTGGAGAAAAAGATGGGACAAGTGCTACTCATAGCAGTAGCCATCTCCAGTGCCGTTCTGCTGCGCACCGCCTGGGGGTGGGGGGAGGCTGGGGCCCTGATCACAGCCGCAACTTCCACTTTGTGGGAAGGCTCTCCGAATAAATACTGGAACTCCTCCACAGCCACTTCACTGTGTAACATTTTTAGGGGAAGTTACTTGGCTGGAGCTTCTCTTATTTACACAGTAACAAGAAACGCTGGCCTGGTCAAGAGACGTGGAGGTGGAACGGGAGAGACCCTGGGGGAGAAATGGAAGGCCCGCCTGAACCAGATGTCGGCCCTGGAGTTTTACTCCTACAAAAAGTCAGGCATCACCGAAGTGTGCAGAGAAGAAGCCCGCCGCGCCCTCAAGGACGGAGTGGCAACAGGAGGCCATGCTGTGTCCCGAGGAAGCGCAAAGCTTAGATGGTTGGTGGAGAGAGGATACCTGCAGCCCTATGGAAAGGTCATTGATCTTGGATGTGGCAGAGGGGGCTGGAGTTACTACGCCGCCACCATCCGCAAAGTTCAAGAGGTGAAAGGATACACAAAGGGAGGCCCTGGTCATGAAGAACCCACGTTGGTGCAAAGCTATGGATGGAACATAGTCCGTCTTAAGAGTGGGGTGGACGTCTTTCACATGGCGGCGGAGTCGTGTGACACTTTGCTGTGTGACATAGGTGAGTCATCATCTAGTCCTGAAGTGGAAGAAGCACGGACGCTCAGAGTACTCTCCATGGTGGGGGATTGGCTTGAAAAAAGACCAGGGGCCTTTTGTATAAAGGTGTTGTGCCCATACACCAGCACCATGATGGAAACCCTAGAGCGACTGCAGCGTAGGTATGGGGGAGGACTGGTCAGAGTGCCACTCTCCCGCAACTCTACACATGAGATGTACTGGGTCTCTGGAGCGAAAAGCAACATCATAAAAAGTGTGTCCACCACGAGCCAGCTCCTCTTGGGACGCATGGACGGGCCCAGGAGGCCAGTGAAATATGAGGAGGATGTGAATCTCGGCTCCGGCACGCGAGCTGTGGCAAGCTGCGCCGAAGCTCCCAACCTGAAGATCATTGGTAACCGCGTTGAGAGGATCCGCAGTGAGCATGCGGAAACGTGGTTCTTTGATGAGAACCACCCATACAGGACATGGGCTTACCATGGGAGCTACGAGGCCCCTACACAAGGGTCAGCGTCTTCTCTCATAAACGGGGTTGTCAGGCTCCTGTCAAAGCCCTGGGATGTGGTGACTGGAGTCACAGGAATAGCCATGACCGACACCACACCGTATGGCCAGCAAAGAGTTTTCAAGGAAAAAGTGGACACTAGGGTGCCAGACCCCCAGGAAGGCACTCGTCAGGTGATGAACATGGTCTCTTCCTGGCTATGGAAGGAGCTAGGTAAACACAAACGGCCACGAGTTTGCACCAAAGAAGAGTTCATCAATAAGGTTCGCAGCAATGCAGCACTGGGGGCAATATTTGAAGAGGAGAAAGAATGGAAGACTGCAGTGGAAGCTGTGAACGATCCAAGGTTCTGGGCCCTAGTGGACAAGGAAAGAGAGCACCACTTGAGAGGAGAGTGTCAGAGCTGTGTGTACAACATGATGGGAAAAAGAGAAAAGAAGCAAGGGGAATTTGGAAAGGCCAAGGGCAGCCGCGCCATTTGGTACATGTGGCTAGGGGCTAGATTTCTAGAGTTTGAAGCCCTTGGATTCTTGAACGAGGATCACTGGATGGGGAGAGAGAATTCAGGAGGTGGTGTTGAAGGGCTGGGATTACAAAGACTTGGATATGTTCTAGAAGAAATGAGCCGCACACCAGGAGGAAAGATGTATGCAGATGATACCGCTGGCTGGGACACCCGCATCAGTAGGTTTGATCTGGAGAATGAAGCTCTGATCACCAACCAAATGGAGAAAGGGCACAGGGCCTTGGCGTTGGCCATAATCAAGTACACATACCAAAACAAAGTGGTAAAGGTCCTTAGACCAGCTGAAAGAGGGAAGACAGTTATGGACATCATCTCAAGACAAGACCAAAGAGGGAGCGGACAAGTTGTTACTTACGCTCTTAATACATTCACCAACCTGGTGGTGCAGCTCATTCGGAACATGGAGGCTGAGGAAGTTCTAGAGATGCAAGACTTGTGGCTGTTGAGGAGGCCAGAGAAGGTGACCAGCTGGTTGCAGAGCAACGGATGGGATAGGCTCAAACGAATGGCAGTCAGTGGAGATGATTGTGTTGTGAAACCAATTGATGATAGGTTTGCACATGCCCTCAGGTTTTTGAATGACATGGGGAAAGTTAGGAAGGACACACAGGAGTGGAAACCCTCAACTGGATGGAGCAACTGGGAAGAAGTTCCGTTTTGCTCCCATCACTTCAACAAGCTTTACCTCAAGGACGGGAGGTCCATTGTGGTCCCCTGTCGCCACCAAGATGAACTGATTGGCCGAGCCCGCGTCTCACCAGGGGCGGGATGGAGCATCCGGGAGACTGCTTGCCTAGCAAAATCATATGCACAAATGTGGCAGCTTCTTTATTTCCACAGAAGGGACCTCCGACTGATGGCCAACGCCATTTGTTCATCTGTGCCAGTTGACTGGGTTCCAACTGGGAGAACCACCTGGTCAATCCATGGAAAGGGAGAATGGATGACCACTGAGGACATGCTTGTGGTGTGGAACAGAGTGTGGATTGAGGAGAACGACCACATGGAGGACAAGACCCCAGTCACGAAATGGACAGACATTCCCTATTTGGGAAAAAGGGAAGACTTATGGTGTGGATCTCTTATAGGGCACAGACCACGCACTACTTGGGCTGAGAACATTAAAGACACAGTCAACATGGTGCGCAGGATCATAGGTGATGAAGAAAAGTACATGGACTACCTATCCACTCAAGTTCGCTACTTGGGTGAAGAAGGGTCCACACCTGGAGTGTTA---

>ENA|AHL43503|AHL43503.1 Zika virus polyprotein

ATGAAAAACCC-AAAGAAGAAATCCGGAAGATTCCGGATTGTCAATATGCTAAAACGCGGAGTAGCCCGTGTAAACCCCTTGGGAGGTTTGAAGAGGTTGCCAGCCGGACTTCTGCTGGGTCATGGACCCATCAGAATGGTTTTGGCGATACTAGCCTTTTTGAGATTTACAGCAATCAAGCCATCACTGGGCCTTATCAACAGATGGGGTTCCGTGGGGAAAAAAGAGGCTATGGAAATAATAAAGAAGTTCAAGAAAGATCTTGCTGCCATGTTGAGAATAATCAATGCTAGGAAAGAGAGGAAGAGACGTGGCGCAGACACCAGCATCGGAATCATTGGCCTCCTGCTGACTACAGCCATGGCAGCAGAGATCACTAGACGCGGGAGTGCATACTACATGTACTTGGATAGGAGCGATGCCGGGAAGGCCATTTCGTTTGCTACCACATTGGGAGTGAACAAGTGCCACGTACAGATCATGGACCTCGGGCACATGTGTGACGCCACCATGAGTTATGAGTGCCCTATGCTGGATGAGGGAGTGGAACCAGATGATGTCGATTGCTGGTGCAACACGACATCAACTTGGGTTGTGTACGGAACCTGTCATCACAAAAAAGGTGAGACACGGCGATCTAGAAGATCTGTGTCGCTCCGTTATCACTATACAAGGAAGTTGCAAACGCGGTCGCAGACATGGTTAGAATCAAGAGAATACAAGAAGCACTTGATCATGGTCGAAAACTGGATATTCAGGAACCCCGGGTTTGCCATAGTGTCCGTTGCCATTACCTGGCTGATGGGAAGCTTGACGAGCCAAAAAGTCATATACTTGGTCATGATAGTGTTGATTGTCCCGGCATACAGTATCAGCTGCATTGGAGTCAGCAATAGAGACTTAGTGGAGGGCATGTCAGGTGGGACCTGGGTTGATGTTGTCTTGGAACATGGAGGGTGCGTTACCGAGATGGCACAGGACAAGCCAACAGTTGACATAGAGTTGGTCACGATGACGGTTAGTAACATGGCCGAGGTAAGATCCTATTGCTACGAGGCATCGTTATCCGACATGGCTTCGGCCAGTCGTTGCCCAACACAAGGCGAACCCTCCCTCGACAAGCAATCAGACACTCAATCTGTATGCAAAAGAACATTAGGAGACAGAGGTTGGGGAAATGGTTGTGGGATTTTTGGCAAAGGGAGCTTGGTGACATGTTCCAAGTTCACGTGTTGTAAGAAGATGCCCGGGAAGAGCATTCAACCGGAAAATCTGGAGTATCGGATAATGCTCCCAGTGCATGGCTCCCAGCATAGCGGGATGATTGTGAATGACATAGGACATGAAACTGACGAAAACAGAGCGAAAGTCGAGGTCACACCCAATTCACCAAGAGCAGAAGCAACCTTGGGAGGTTTTGGAAGCTTGGGACTTGACTGTGAACCAAGGACAGGCCTTGACTTCTCAGATCTGTATTATCTGACCATGAACAACAAGCATTGGTTGGTGCACAAGGAGTGGTTTCATGACATCCCATTACCTTGGCATGCTGGTGCAGACACTGGAACTCCACACTGGAACAACAAAGAGGCATTGGTGGAGTTCAAGGACGCCCACGCCAAGAGGCAAACTGTTGTGGTTCTGGGGAGCCAAGAGGGAGCTGTTCACACGGCCCTCGCTGGAGCTCTGGAGGCTGAGATGGATGGTGCAAAGGGAAGGCTATTCTCTGGCCATTTGAAATGCCGCCTAAAAATGGACAAGCTTAGGTTGAAGGGTGTGTCATATTCCCTGTGTACTGCAGCGTTCACATTTACCAAGGTCCCAGCTGAAACATTGCATGGAACAGTTACAGTGGAGGTGCAGTCTGCAGGGACAGATGGACCCTGCAAGGTCCCAGCCCAGATGGCGGTGGACATGCAGACCCTGACCCCAGTTGGAAGGCTGATAACCGCCAACCCCGTGATTACTGAAAGCACTGAGAACTCAAAGATGATGTTGGAGCTTGACCCACCATTTGGGGATTCTTACATTGTCATAGGAGTTGGGGACAAGAAAATCACCCACCACTGGCATAGGAGTGGTAGCACCATCGGAAAGGCATTTGAAGCCACTGTGAGAGGCGCCAAGAGAATGGCAGTCCTGGGGGATACAGCCTGGGACTTCGGATCAGTCGGGGGTGTGTTCAACTCACTGGGTAAGGGCATTCACCAGATTTTTGGAGCAGCCTTCAAATCACTGTTTGGAGGAATGTCCTGGTTCTCACAGATCCTCATAGGCACGCTGCTAGTGTGGTTAGGTTTGAACACAAAGAATGGATCTATCTCCCTCACATGCTTGGCCCTGGGGGGAGTGATGATCTTCCTCTCCACGGCTGTTTCTGCTGACGTGGGGTGCTCAGTGGACTTCTCAAAAAAGGAAACGAGATGTGGCACGGGGGTATTCATCTATAATGATGTTGAAGCCTGGAGGGACCGGTACAAGTACCATCCTGACTCCCCCCGCAGATTGGCAGCAGCAGTCAAGCAGGCCTGGGAAGAGGGGATCTGTGGGATCTCATCCGTTTCAAGAATGGAAAACATCATGTGGAAATCAGTAGAAGGGGAGCTCAATGCTATCCTAGAGGAGAATGGAGTTCAACTGACAGTTGTTGTGGGATCTGTAAAAAACCCCATGTGGAGAGGTCCACAAAGATTGCCAGTGCCTGTGAATGNGCTGCCCCATGGCTGGAAAGCCTGGGGGAAATCGTATTTTGTTAGGGCGGCAAAGACCAACAACAGTTTTGTTGTCGACGGTGACACACTGAAGGAATGTCCGCTTAAGCACAGAGCATGGAATAGTTTTCTTGTGGAGGATCACGGGTTTGGAGTCTTCCACACCAGTGTTTGGCTTAAGGTCAGAGAAGATTACTCATTAGAATGTGACCCAGCCGTCATAGGAACAGCTGTTAAGGGAAGGGAGGCCGCGCACAGTGATCTGGGCTATTGGATTGAAAGTGAAAAGAATGACACATGGAGGCTGAAGAGGGCCCACCTGATTGAGATGAAAACATGTGAATGGCCAAAGTCTCACACATTGTGGACAGATGGAGTAGAAGAAAGTGATCTTATCATACCCAAGTCTTTAGCTGGTCCACTCAGCCACCACAACACCAGAGAGGGTTACAGAACCCAAGTGAAAGGGCCATGGCACAGTGAAGAGCTTGAAATCCGGTTTGAGGAATGTCCAGGCACCAAGGTTCACGTGGAGGAGACATGCGGAACTAGAGGACCATCTCTGAGATCAACTACTGCAAGTGGAAGGGTCATTGAGGAATGGTGCTGTAGGGAATGCACAATGCCCCCACTATCGTTTCGAGCAAAAGACGGCTGCTGGTATGGAATGGAGATAAGGCCCAGGAAAGAACCAGAGAGCAACTTAGTGAGGTCAATGGTGACAGCGGGGTCAACCGATCATATGGACCACTTCTCTCTTGGAGTGCTTGTGATTCTACTCATGGTGCAGGAGGGGTTGAAGAAGAGAATGACCACAAAGATCATCATGAGCACATCAATGGCAGTGCTGGTAGTCATGATCTTGGGAGGATTTTCAATGAGTGACCTGGCCAAGCTTGTGATCCTGATGGGTGCTACTTTCGCAGAAATGAACACTGGAGGAGATGTAGCTCACTTGGCATTGGTAGCGGCATTTAAAGTCAGACCAGCCTTGCTGGTCTCCTTCATTCTCAGAGCCAATTGGACACCCCGTGAGAGCATGCTGCTAGCCCTGGCTTCGTGTCTTCTGCAAACTGCGATCTCTGCTCTTGAAGGTGACTTGATGGTCCTCGTTAATGGATTTGCTTTGGCCTGGTTGGCAATTCGAGCAATGGCCGTGCCACGCACTGACAACATCGCTCTAGCAATCTTGGCTGCTCTAACACCACTAGCTCGAGGCACACTGCTCGTGGCATGGAGAGCGGGCCTGGCTACTTGTGGAGGGTTCATGCTCCTCTCCCTGAAAGGGAAAGGTAGTGTGAAGAAGAACCTGCCATTTGTCATGGCCCTGGGATTGACAGCTGTGAGGGTAGTAGACCCTATTAATGTGGTAGGACTACTGTTACTCACAAGGAGTGGGAAGCGGAGCTGGCCCCCTAGTGAAGTTCTCACAGCCGTTGGCCTGATATGTGCACTGGCCGGAGGGTTTGCCAAGGCAGACATTGAGATGGCTGGACCCATGGCTGCAGTAGGCTTGCTAATTGTCAGCTATGTGGTCTCGGGAAAGAGTGTGGACATGTACATTGAAAGAGCAGGTGACATCACATGGGAAAAGGACGCGGAAGTCACTGGAAACAGTCCTCGGCTTGACGTGGCACTGGATGAGAGTGGTGATTTCTCCTTGGTAGAGGAAGATGGTCCACCCATGAGAGAGATCATACTCAAGGTGGTCCTGATGGCCATCTGTGGCATGAACCCAATAGCTATACCTTTTGCTGCAGGAGCGTGGTATGTGTATGTGAAGACTGGGAAAAGGAGTGGCGCCCTCTGGGACGTGCCTGCTCCCAAAGAAGTGAAGAAAGGAGAGACCACAGATGGAGTGTACAGAGTGATGACTCGCAGACTGCTAGGTTCAACACAGGTTGGAGTGGGAGTCATGCAAGAGGGAGTCTTCCACACCATGTGGCACGTTACAAAAGGAGCCGCACTGAGGAGCGGTGAGGGAAGACTTGATCCATACTGGGGGGATGTCAAGCAGGACTTGGTGTCATACTGTGGGCCTTGGAAGTTGGATGCAGCTTGGGATGGACTTAGCGAGGTACAGCTTTTGGCCGTACCTCCCGGAGAGAGGGCCAGAAACATTCAGACCCTGCCTGGAATATTCAAGACAAAGGACGGGGACATCGGAGCAGTTGCTCTGGACTACCCTGCAGGGACCTCAGGATCTCCGATCCTAGACAAATGTGGAAGAGTGATAGGACTCTATGGCAATGGGGTTGTGATCAAGAATGGAAGCTATGTTAGTGCTATAACCCAGGGAAAGAGGGAGGAGGAGACTCCGGTTGAATGTTTCGAACCCTCGATGCTGAAGAAGAAGCAGCTAACTGTCCTGGATCTGCATCCAGGAGCCGGAAAAACCAGGAGAGTTCTTCCTGAAATAGTCCGTGAAGCCATAAAAAAGAGACTCCGGACAGTGATCTTGGCACCAACTCCAGTTGAGAGATCATACTCAAGGTGGTCCTTGTGGCCATCTGTGGCACGTACCCAGACAACAGCAGTCAACGTCACCCATTCTGGGACAGAAATCGTTGATTTGATGTGCCATGCCACTTTCACTTCACGCTTACTACAACCCATCAGAGTCCCTAATTACAATCTCAACATCATGGATGAAGCCCACTTCACAGACCCCTCAAGTATAGCTGCAAGAGGATACATATCAACAAGGGTTGAAATGGGCGAGGCGGCTGCCATTTTTATGACTGCCACACCACCAGGAACCCGTGATGCGTTTCCTGACTCTAACTCACCAATCATGGACACAGAAGTGGAAGTCCCAGAGAGAGCCTGGAGCTCAGGCTTTGATTGGGTGACAGACCATTCTGGGAAAACAGTTTGGTTCGTTCCAAGCGTGAGATCTGGAGAAGAAAGCGCAGCCTGTCTGACAAAGGCTGGAAAGCGGGTCATACAGCTCAGCAGGAAGACTTTTGAGACAGAATTTCAGAAAACAAAAAATCAAGAGTGGGACTTTGTCATAACAACTGACATCTCAGAGATGGGCGCCAACTTCAAGGCTGACCGGGTCATAGACTCTAGGAGATGCCTAAAGCCAGTCATACTTGATGGTGAGAGAGTCATCTTGGCTGGGCCCATGCCTGTCACGCATGCTAGTGCTGCTCAGAGGAGAGGACGTATAGGCAGGAACCCTAACAAACCTGGAGATGAGTACATGTATGGAGGTGGGTGTGCAGAGACTGATGAAGACCATGCACACTGGCTTGAAGCAAGAATGCTTCTTGACAACATCTACCTCCAGGATGGCCTCATAGCCTCGCTCTATCGGCCTGAGGCCGATAAGGTAGCCGCCATTGAGGGAGAGTTTAAGCTGAGGACAGAGCAAAGGAAGACCTTCGTGGAACTCATGAAGAGAGGAGACCTTCCCGTCTGGCTAGCCTATCAGGTTGCATCTGCCGGAATAACTTACACAGACAGAAGATGGTGCTTTGATGGCACAACCAACAACACCATAATGGAAGACAGCGTACCAGCAGAGGTGTGGACAAAGTATGGAGAGAAGAGAGTGCTCAAACCGAGATGGATGGATGCTAGGGTCTGTTCAGACCATGCGGCCCTGAAGTCGTTCAAAGAATTCGCCGCTGGAAAAAGAGGAGCGGCTTTGGGAGTAATGGAGGCCCTGGGAACACTGCCAGGACACATGACAGAGAGGTTTCAGGAAGCCATTGACAACCTCGCCGTGCTCATGCGAGCAGAGACTGGAAGCAGGCCTTATAAGGCAGCGGCAGCCCAACTGCCGGAGACCCTAGAGACCATTATGCTCTTAGGTTTGCTGGGAACAGTTTCACTGGGGATCTTCTTCGTCTTGATGCGGAATAAGGGCATCGGGAAGATGGGCTTTGGAATGGTAACCCTTGGGGCCAGTGCATGGCTCATGTGGCTTTCGGAAATTGAACCAGCCAGAATTGCATGTGTCCTCATTGTTGTGTTTTTATTACTGGTGGTGCTCATACCCGAGCCAGAGAAGCAAAGATCTCCCCAAGATAACCAGATGGCAATTATCATCATGGTGGCAGTGGGCCTTCTAGGTTTGATAACTGCAAACGAACTTGGATGGCTGGAAAGAACAAAAAATGACATAGCTCATCTAATGGGAAGGAGAGAAGAAGGAGCAACCATGGGATTCTCAATGGACATTGATCTGCGGCCAGCCTCCGCCTGGGCTATCTATGCCGCATTGACAACTCTCATCACCCCAGCTGTCCAACATGCGGTAACCACTTCATACAACAACTACTCCTTAATGGCGATGGCCACACAAGCTGGAGTGCTGTTTGGCATGGGCAAAGGGATGCCATTT-ATGCATGGGGACCTTGGAGTCCCGCTGCTAATGATGGGTTGCTATTCACAATTAACACCCCTGACTCTGATAGTAGCTATCATTCTGCTTGTGGCGCACTACATGTACTTGATCCCAGGCCTACAAGCGGCAGCAGCGCGTGCTGCCCAGAAAAGGACAGCAGCTGGCATCATGAAGAATCCCGTTGTGGATGGAATAGTGGTAACTGACATTGACACAATGACAATAGACCCCCAGGTGGAGAAGAAGATGGGACAAGTGTTACTCATAGCAGTAGCCATCTCCAGTGCTGTGCTGCTGCGGACCGCCTGGGGATGGGGGGAGGCTGGAGCTCTGATCACAGCAGCGACCTCCACCTTGTGGGAAGGCTCTCCAAACAAATACTGGAACTCCTCTACAGCCACCTCACTGTGCAACATCTTCAGAGGAAGCTATCTGGCAGGAGCTTCCCTTATCTATACAGTGACGAGAAACGCTGGCCTGGTTAAGAGACGTGGAGGTGGGACGGGAGAGACTCTGGGAGAGAAGTGGAAAGCTCGTCTGAATCAGATGTCGGCCCTGGAGTTCTACTCTTATAAAAAGTCAGGTATCACTGAAGTGTGTAGAGAGGAGGCTCGCCGTGCCCTCAAGGATGGAGTGGCCACAGGAGGACATGCCGTATCCCGGGGAAGTGCAAAGCTCAGATGGTTGGTGGAGAGAGGATATCTGCAGCCCCATGGGAAGGTTGTTGACCTCGGATGTGGCAGAGGGGGCTGGAGCTATTATGCCGCCACCATCCGCAAAGTGCAGGAGGTGAGAGGATACACAAAGGGAGGTCCCGGTCATGAAGAACCCATGCTGGTGCAAAGCTATGGGTGGAACATAGTTCGTCTCAAGAGTGGAGTGGACGTCTTCCACATGGCGGCTGAGCCGTGTGACACTCTGCTGTGTGACATAGGTGAGTCATCATCTAGTCCTGAAGTGGAAGAGACACGAACACTCAGAGTGCTCTCTATGGTGGGGGACTGGCTTGAAAAAAGACCAGGGGCCTTCTGTATAAAGGTGCTGTGCCCATACACCAGCACTATGATGGAAACCATGGAGCGACTGCAACGTAGGCATGGGGGAGGATTAGTCAGAGTGCCATTGTCTCGCAACTCCACACATGAGATGTACTGGGTCTCTGGGGCAAAAAGCAACATCATAAAAAGTGTGTCCACCACAAGTCAGCTCCTCCTGGGACGCATGGATGGCCCCAGGAGGCCAGTGAAATATGAGGAGGATGTGAACCTCGGCTCGGGTACACGAGCTGTGGCAAGCTGTGCTGAGGCTCCTAACATGAAAATCATCGGCAGGCGCATTGAGAGAATCCGCAATGAACATGCAGAAACATGGTTTCTTGATGAAAACCACCCATACAGGACATGGGCCTACCATGGGAGNTACGAAGCCCCCACGCAAGGATCAGCGTCTTCCCTCGTGAACGGGGTTGTTAGACTCCTGTCAAAGCCTTGGGACGTGGTGACTGGAGTTACAGGAATAGCCATGACTGACACCACACCATACGGCCAACAAAGAGTCTTCAAAGAAAAAGTGGACACCAGGGTGCCAGATCCCCAAGAAGGCACTCGCCAGGTAATGAACATGGTCTCTTCCTGGCTGTGGAAGGAGCTGGGGAAACGCAAGCGGCCACGCGTCTGCACCAAAGAAGAGTTTATCAACAAGGTGCGCAGCAATGCAGCACTGGGAGCAATATTTGAAGAGGAAAAAGAATGGAAGACGGCTGTGGAAGCTGTGAATGATCCAAGGTTTTGGGCCCTAGTGGATAGGGAGAGAGAACACCACCTGAGAGGAGAGTGTCACAGCTGTGTGTACAACATGATGGGAAAAAGAGAAAAGAAGCAAGGAGAGTTCGGGAAAGCAAAAGGTAGCCGCGCCATCTGGTACATGTGGTTGGGAGCCAGGTTTCTGGAGTTTGAATCACTGGGGTTTCTGAATGAAGATCACTGGATGGGAAGAGAGAACTCTGGAGGCGGAGTTGAAGGACTGGGACTGCAGAGACTGGGCTATGTCCTTGAGGAGATGAGCCAGGCACCAGGAGGGAAGATGTACGCAGATGACACTGCTGGCTGGGACACCCGCATTAGTAAGTTTGATCTGGAGAATGAAGCTTTGATTACCAACCAAATGGAGGAAGGGCACAGAACTCTGGCGTTGGCCGTGATTAAATACACATACCAAAACAAAGTGGTGAAGGTCCTCAGACCAGCTGAAGGAGGAAAAACAGTTATGGACATCATTTCAAGACAAGACCAGAGAGGGAGTGGACAAGTTGTCACTTATGCTCTCAACACATTCACCAACTTGGTGGTGCAGCTTATCCGGAACATGGAAGCTGAGGAAGTGTTAGAGATGCAAGACTTATGGCTGTTGAGGAAGCCAGAGAAAGTGACCAGATGGTTGCAGAGCAATGGATGGGATAGACTCAAACGAATGGCGGTCAGTGGAGATGACTGCGTTGTGAAGCCAATCGATGATAGGTTTGCACATGCCCTCAGGTTCTTGAATGACATGGGAAAAGTTAGGAAAGACACACAGGAGTGGAAACCCTCGACTGGATGGAGCAATTGGGAAGAAGTCCCGTTCTGCTCCCACCACTTCAACAAGCTGTACCTCAAGGATGGGAGATCCATTGTGGTCCCTTGCCGCCACCAAGATGAACTGATTGGCCGAGCTCGCGTTTCACCAGGGGCAGGATGGAGCATCCGGGAGACTGCCTGTCTTGCAAAATCATATGCGCAGATGTGGCAGCTCCTTTATTTCCACAGAAGAGACCTTCGACTGATGGCTAATGCCATTTGCTCGGCTGTACCAGTTGACTGGGTACCAACTGGGAGAACCACCTGGTCAATCCATGGAAAGGGAGAATGGATGACCACTGAGGACATGCTCATGGTGTGGAATAGAGTGTGGATTGAGGAGAACGACCATATGGAGGACAAGACTCCTGTAACAAAATGGACAGACATTCCCTATCTAGGAAAAAGGGAGGACTTATGGTGTGGATCCCTTATAGGGCACAGACCCCGCACCACTTGGGCTGAAAACATCAAAGACACAGTCAACATGGTGCGCAGGATCATAGGTGATGAAGAAAAGTACATGGACTATCTATCCACCCAAGTCCGCTACTTGGGTGAGGAAGGGTCCACACCCGGAGTGTTGTAA

>ENA|AML82110|AML82110.1 Zika virus polyprotein

ATGAAAAACCC-AAAAAAGAAATCCGGAGGATTCCGGATTGTCAATATGCTAAAACGCGGAGTAGCCCGTGTGAGCCCCTTTGGGGGCTTGAAGAGGCTGCCAGCCGGACTTCTGCTGGGTCATGGGCCCATCAGGATGGTCTTGGCGATTCTAGCCTTCTTGAGATTCACGGCAATCAAGCCATCACTGGGTCTCATCAATAGATGGGGTTCAGTGGGGAAAAAAGAGGCTATGGAAATAATAAAGAAGTTCAAGAAAGATCTGGCTGCCATGCTGAGAATAATCAATGCTAGGAAGGAGAAGAAGAGACGAGGCGCAGATACTAGTGTCGGAATTGTTGGCCTCCTGCTGACCACAGCTATGGCAGCGGAGGTCACTAGACGTGGGAGTGCATACTATATGTACTTGGACAGAAACGATGCTGGGGAGGCCATATCTTTTCCAACCACATTGGGGATGAATAAGTGTTATATACAGATCATGGATCTTGGACACATGTGTGATGCCACCATGAGCTATGAATGCCCTATGCTGGATGAGGGGGTGGAACCAGATGACGTCGATTGTTGGTGCAACACGACGTCAACTTGGGTTGTGTACGGAACCTGCCATCACAAAAAAGGTGAAGCACGGAGATCTAGAAGAGCTGTGACGCTCCCTTCCCATTCCACTAGGAAGCTGCAAACGCGGTCGCAAACCTGGTTGGAATCAAGAGAATACACAAAGCACTTGATTAGAGTCGAAAATTGGATATTCAGGAACCCTGGCTTCGCGTTAGCAGCAGCTGCTATCGCTTGGCTTTTGGGAAGCTCAACGAGCCAAAAAGTCATATACTTGGTCATGATACTGCTGATTGCCCCGGCATACAGCATCAGGTGCATAGGAGTCAGCAATAGGGACTTTGTGGAAGGTATGTCAGGTGGGACTTGGGTTGATGTTGTCTTGGAACATGGAGGTTGTGTCACCGTAATGGCACAGGACAAACCGACTGTCGACATAGAGCTGGTTACAACAACAGTCAGCAACATGGCGGAGGTAAGATCCTACTGCTATGAGGCATCAATATCAGACATGGCTTCGGACAGCCGCTGCCCAACACAAGGTGAAGCCTACCTTGACAAGCAATCAGACACTCAATATGTCTGCAAAAGAACGTTAGTGGACAGAGGCTGGGGAAATGGATGTGGACTTTTTGGCAAAGGGAGCCTGGTGACATGCGCTAAGTTTGCATGCTCCAAGAAAATGACCGGGAAAAGCATCCAGCCAGAGAATCTGGAGTACCGGATAATGCTGTCAGTTCATGGCTCCCAGCACAGTGGGATGATCGTTAATGACACAGGACATGAAACTGATGAGAATAGAGCGAAAGTTGAGATAACGCCCAATTCACCAAGAGCCGAAGCCACCCTGGGGGGTTTTGGAAGCCTAGGACTTGATTGTGAACCGAGGACAGGCCTTGACTTTTCAGATTTGTATTACTTGACTATGAATAACAAGCACTGGTTGGTTCACAAGGAGTGGTTCCACGACATTCCATTACCTTGGCACGCTGGGGCAGACACCGGAACTCCACACTGGAACAACAAAGAAGCACTGGTAGAGTTCAAGGACGCACATGCCAAAAGGCAAACTGTCGTGGTTCTAGGGAGTCAAGAAGGAGCAGTTCACACGGCCCTTGCTGGAGCTCTGGAGGCTGAGATGGATGGAGCAAAGGGAAGGCTGTCCTCTGGCCACTTGAAATGTCGCCTGAAAATGGATAAACTTAGATTGAAGGGCGTGTCATACTCCTTGTGTACTGCAGCGTTCACATTCACCAAGATCCCGGCTGAAACACTGCACGGGACAGTCACAGTGGAGGTACAGTACGCAGGGACAGATGGACCTTGCAAGGTTCCAGCTCAGATGGCGGTGGACATGCAAACTCTGACCCCAGTTGGGAGGTTGATAACCGCTAACCCCGTAATCACTGAAAGCACTGAGAACTCTAAGATGATGCTGGAACTTGATCCACCATTTGGGGACTCTTACATTGTCATAGGAGTCGGGGAGAAGAAGATCACCCACCACTGGCACAGGAGTGGCAGCACCATTGGAAAAGCATTTGAAGCCACTGTGAGAGGTGCCAAGAGAATGGCAGTCTTGGGAGACACAGCCTGGGACTTTGGATCAGTTGGAGGCGCTCTCAACTCATTGGGCAAGGGCATCCATCAAATTTTTGGAGCAGCTTTCAAATCATTGTTTGGAGGAATGTCCTGGTTCTCACAAATTCTCATTGGAACGTTGCTGATGTGGTTGGGTCTGAACACAAAGAATGGATCTATTTCCCTTATGTGCTTGGCCTTAGGGGGAGTGTTGATCTTCTTATCCACAGCCGTCTCTGCTGATGTGGGGTGCTCGGTGGACTTCTCAAAGAAGGAGACGAGATGCGGTACAGGGGTGTTTGTCTATAACGACGTTGAAGCCTGGAGGGACAGGTACAAGTACCATCCTGACTCCCCCCGTAGATTGGCAGCAGCAGTCAAGCAAGCCTGGGAAGATGGTATCTGCGGGATCTCCTCTGTTTCAAGAATGGAAAACATCATGTGGAGATCAGTAGAAGGGGAGCTCAACGCAATCCTGGAAGAGAATGGAGTTCAACTGACGGTCGTTGTGGGATCTGTAAAAAACCCCATGTGGAGAGGTCCACAGAGATTGCCCGTGCCTGTGAACGAGCTGCCCCACGGCTGGAAGGCTTGGGGGAAATCGTACTTCGTCAGAGCAGCAAAGACAAATAACAGCTTTGTCGTGGATGGTGACACACTGAAGGAATGCCCACTCAAACATAGAGCATGGAACAGCTTTCTTGTGGAGGATCATGGGTTCGGGGTATTTCACACTAGTGTCTGGCTCAAGGTTAGAGAAGACTATTCATTAGAGTGTGATCCAGCCGTTATTGGAACAGCTGTTAAGGGAAAGGAGGCTGTACACAGTGATCTAGGCTACTGGATTGAGAGTGAGAAGAATGACACATGGAGGCTGAAGAGGGCCCATCTGATCGAGATGAAAACATGTGAATGGCCAAAGTCCCACACATTGTGGACAGATGGAATAGAAGAGAGTGATCTGATCATACCCAAGTCTTTAGCTGGGCCACTCAGCCATCACAATACCAGAGAGGGCTACAGGACCCAAATGAAAGGGCCATGGCACAGTGAGGAGCTTGAAATTCGGTTTGAGGAATGCCCAGGCACTAAGGTCCACGTGGAGGAAACATGTGGAACAAGAGGACCATCTCTGAGATCAACCACTGCAAGCGGAAGGGTGATCGAGGAATGGTGCTGCAGGGAGTGCACAATGCCCCCACTGTCGTTCCGGGCTAAAGATGGCTGTTGGTATGGAATGGAGATAAGGCCCAGGAAAGAACCAGAAAGCAACTTAGTAAGGTCAATGGTGACTGCAGGATCAACTGATCACATGGACCACTTCTCCCTTGGAGTGCTTGTGATCCTGCTCATGGTGCAGGAAGGGCTGAAGAAGAGAATGACCACAAAGATCATCATAAGCACATCAATGGCAGTGCTGGTAGCTATGATCCTGGGAGGATTTTCAATGAGTGACCTGGCTAAGCTTGCAATTTTGATGGGTGCCACCTTCGCGGAAATGAACACTGGAGGAGATGTAGCTCATCTGGCGCTGATAGCGGCATTCAAAGTCAGACCAGCGTTGCTGGTATCTTTCATCTTCAGAGCTAATTGGACACCCCGTGAAAGCATGCTGCTGGCCTTGGCCTCGTGTCTTTTGCAAACTGCGATCTCCGCCTTGGAAGGCGACCTGATGGTTCTCATCAATGGTTTTGCTTTGGCCTGGTTGGCAGTACGAGCGATGGTTGTTCCACGCACTGATAACATCACCTTAGCAATCCTGGCTGCTCTGACACCACTGGCCCGGGGCACACTGCTTGTGGCGTGGAGAGCAGGCCTTGCTACTTGCGGGGGGTTTATGCTCCTCTCTCTGAAGGGAAAAGGCAGTGTGAAGAAGAACTTACCATTTGTCATGGCCCTGGGACTAACCGCTGTGAGGCTGGTCGACCCCATCAACGTGGTGGGACTGCTGTTGCTCACAAGGAGTGGGAAGCGGAGCTGGCCCCCTAGCGAAGTACTCACAGCTGTTGGCCTGATATGCGCATTGGCTGGAGGGTTCGCCAAGGCAGATATAGAGATGGCTGGGCCCATGGCCGCGGTCGGTCTGCTAATTGTCAGTTACGTGGTCTCAGGAAAGAGTGTGGACATGTACATTGAAAGAGCAGGTGACATCACATGGGAAAAAGATGCGGAAGTCACTGGAAACAGTCCCCGGCTCGATGTGGCGCTAGATGAGAGTGGTGATTTCTCCCTGGTGGAGGATGACGGTCCCCCCATGAGAGAGATCATACTCAAGGTGGTCCTGATGACCATCTGTGGCATGAACCCAATAGCCATACCCTTTGCAGCTGGAGCGTGGTACGTATACGTGAAGACTGGAAAAAGGAGTGGTGCTCTATGGGATGTGCCTGCTCCCAAGGAAGTAAAAAAGGGGGAGACCACAGATGGAGTGTACAGAGTAATGACTCGCAGACTGCTAGGTTCAACACAAGTTGGAGTGGGAGTTATGCAAGAGGGGGTCTTTCACACTATGTGGCACGTCACAAAAGGATCCGCGCTGAGAAGCGGTGAAGGGAGACTTGATCCATACTGGGGAGATGTCAAGCAGGATCTGGTGTCATACTGTGGTCCATGGAAGCTAGATGCCGCCTGGGACGGGCACAGCGAGGTGCAGCTCTTGGCCGTGCCCCCCGGAGAGAGAGCGAGGAACATCCAGACTCTGCCCGGAATATTTAAGACAAAGGATGGGGACATTGGAGCGGTTGCGCTGGATTACCCAGCAGGAACTTCAGGATCTCCAATCCTAGACAAGTGTGGGAGAGTGATAGGACTTTATGGCAATGGGGTCGTGATCAAAAATGGGAGTTATGTTAGTGCCATCACCCAAGGGAGGAGGGAGGAAGAGACTCCTGTTGAGTGCTTCGAGCCTTCGATGCTGAAGAAGAAGCAGCTAACTGTCTTAGACTTGCATCCTGGAGCTGGGAAAACCAGGAGAGTTCTTCCTGAAATAGTCCGTGAAGCCATAAAAACAAGACTCCGTACTGTGATCTTAGCTCCAACCAGGGTTGTCGCTGCTGAAATGGAGGAGGCCCTTAGAGGGCTTCCAGTGCGTTATATGACAACAGCAGTCAATGTCACCCACTCTGGAACAGAAATCGTCGACTTAATGTGCCATGCCACCTTCACTTCACGTCTACTACAGCCAATTAGAGTCCCCAACTATAATCTGTATATTATGGATGAGGCCCACTTCACAGATCCCTCAAGTATAGCAGCAAGAGGATACATTTCAACAAGGGTTGAGATGGGCGAGGCGGCTGCCATCTTCATGACCGCCACGCCACCAGGAACCCGTGACGCATTTCCGGACTCCAACTCACCAATTATGGACACCGAAGTGGAAGTCCCAGAGAGAGCCTGGAGCTCAGGCTTTGATTGGGTGACGGATCATTCTGGAAAAACAGTTTGGTTTGTTCCAAGCGTGAGGAACGGCAATGAGATCGCAGCTTGTCTGACAAAGGCTGGAAAACGGGTCATACAGCTCAGCAGAAAGACTTTTGAGACAGAGTTCCAGAAAACAAAACATCAAGAGTGGGACTTTGTCGTGACAACTGACATTTCAGAGATGGGCGCCAACTTTAAAGCTGACCGTGTCATAGATTCCAGGAGATGCCTAAAGCCGGTCATACTTGATGGCGAGAGAGTCATTCTGGCTGGACCCATGCCTGTCACACATGCCAGCGCTGCCCAGAGGAGGGGGCGCATAGGCAGGAATCCCAACAAACCTGGAGATGAGTATCTGTATGGAGGTGGGTGCGCAGAGACTGACGAAGACCATGCACACTGGCTTGAAGCAAGAATGCTCCTTGACAATATTTACCTCCAAGATGGCCTCATAGCCTCGCTCTATCGACCTGAGGCCGACAAAGTAGCAGCCATTGAGGGAGAGTTCAAGCTTAGGACGGAGCAAAGGAAGACCTTTGTGGAACTCATGAAAAGAGGAGATCTTCCTGTTTGGCTGGCCTATCAGGTTGCATCTGCCGGAATAACCTACACAGATAGAAGATGGTGCTTTGATGGCACGACCAACAACACCATAATGGAAGACAGTGTGCCGGCAGAGGTGTGGACCAGACACGGAGAGAAAAGAGTGCTCAAACCGAGGTGGATGGACGCCAGAGTTTGTTCAGATCATGCGGCCCTGAAGTCATTCAAGGAGTTTGCCGCTGGGAAAAGAGGAGCGGCTTTTGGAGTGATGGAAGCCCTGGGAACACTGCCAGGACACATGACAGAGAGATTCCAGGAAGCCATTGACAACCTCGCTGTGCTCATGCGGGCAGAGACTGGAAGCAGGCCTTACAAAGCCGCGGCAGCCCAATTGCCGGAGACCCTAGAGACCATTATGCTTTTGGGGTTGCTGGGAACAGTCTCGCTGGGAATCTTCTTCGTCTTGATGAGGAACAAGGGCATAGGGAAGATGGGCTTTGGAATGGTGACTCTTGGGGCCAGCGCATGGCTCATGTGGCTCTCGGAAATTGAGCCAGCCAGAATTGCATGTGTCCTCATTGTTGTGTTCCTATTGCTGGTGGTGCTCATACCTGAGCCAGAAAAGCAAAGATCTCCCCAGGACAACCAAATGGCAATCATCATCATGGTAGCAGTAGGTCTTCTGGGCTTGATTACCGCCAATGAACTCGGATGGTTGGAGAGAACAAAGAGTGACCTAAGCCATCTAATGGGAAGGAGAGAGGAGGGGGCAACCATAGGATTCTCAATGGACATTGACCTGCGGCCAGCCTCAGCTTGGGCCATCTATGCTGCCTTGACAACTTTCATTACCCCAGCCGTCCAACATGCAGTGACCACCTCATACAACAACTACTCCTTAATGGCGATGGCCACGCAAGCTGGAGTGTTGTTTGGTATGGGCAAAGGGATGCCATTCTACGCATGGG-ACTTTGGAGTCCCGCTGCTAATGATAGGCTGCTACTCACAATTAACACCCCTGACCCTAATAGTGGCCATCATTTTGCTCGTGGCGCACTACATGTACTTGATCCCAGGGCTGCAGGCAGCAGCTGCGCGTGCTGCCCAGAAGAGAACGGCAGCTGGCATCATGAAGAACCCTGTTGTGGATGGAATAGTGGTGACTGACATTGACACAATGACAATTGACCCCCAAGTGGAGAAAAAGATGGGACAGGTGCTACTCATAGCAGTAGCCGTCTCCAGCGCCATACTGTCGCGGACCGCCTGGGGGTGGGGGGAGGCTGGGGCCCTGATCACAGCCGCAACTTCCACTTTGTGGGAAGGCTCTCCGAACAAGTACTGGAACTCCTCTACAGCCACTTCACTGTGTAACATTTTTAGGGGAAGTTACTTGGCTGGAGCTTCTCTAATCTACACAGTAACAAGAAACGCTGGCTTGGTCAAGAGACGTGGGGGTGGAACAGGAGAGACCCTGGGAGAGAAATGGAAGGCCCGCTTGAACCAGATGTCGGCCCTGGAGTTCTACTCCTACAAAAAGTCAGGCATCACCGAGGTGTGCAGAGAAGAGGCCCGCCGCGCCCTCAAGGACGGTGTGGCAACGGGAGGCCATGCTGTGTCCCGAGGAAGTGCAAAGCTGAGATGGTTGGTGGAGCGGGGATACCTGCAGCCCTATGGAAAGGTCATTGATCTTGGATGTGGCAGAGGGGGCTGGAGTTACTACGCCGCCACCATCCGCAAAGTTCAAGAAGTGAAAGGATACACAAAAGGAGGCCCTGGTCATGAAGAACCCGTGTTGGTGCAAAGCTATGGGTGGAACATAGTCCGTCTTAAGAGTGGGGTGGACGTCTTTCATATGGCGGCTGAGCCGTGTGACACGTTGCTGTGTGACATAGGTGAGTCATCATCTAGTCCTGAAGTGGAAGAAGCACGGACGCTCAGAGTCCTCTCCATGGTGGGGGATTGGCTTGAAAAAAGACCAGGAGCCTTTTGCATAAAAGTGTTGTGCCCATACACCAGCACTATGATGGAAACCCTGGAGCGACTGCAGCGTAGGTATGGGGGAGGACTGGTCAGAGTGCCACTCTCCCGCAACTCTACACATGAGATGTACTGGGTCTCTGGAGCGAAAAGCAACATCATAAAAAGTGTGTCCACCACGAGCCAGCTCCTCTTGGGGCGCATGGACGGGCCTAGGAGGCCAGTGAAATATGAGGAGGATGTGAATCTCGGCTCTGGCACGCGGGCTGTGGCAAGCTGCGCTGAAGCTCCCAACATGAAGATCATTGGTAACCGCATTGAAAGGATCCGCAGTGAGCACGCGGAAACGTGGTTCTTTGACGAGAACCACCCATATAGGACATGGGCTTACCATGGAAGCTATGAGGCCCCCACACAAGGGTCAGCGTCCTCTCTAATAAACGGGGTTGTCAGGCTCCTGTCAAAACCCTGGGATGTGGTGACTGGAGTCACAGGAATAGCCATGACCGACACCACACCGTATGGTCAGCAAAGAGTTTTCAAGGAAAAAGTGGACACTAGGGTGCCAGACCCCCAAGAAGGCACTCGTCAGGTTATGAGCATGGTCTCTTCCTGGTTGTGGAAAGAGCTAGGCAAACACAAACGGCCACGAGTCTGTACCAAAGAAGAGTTCATCAACAAGGTTCGTAGCAATGCAGCATTAGGGGCAATATTTGAAGAGGAAAAAGAGTGGAAGACTGCAGTGGAAGCTGTGAACGATCCAAGGTTCTGGGCTCTAGTGGACAAGGAAAGAGAGCACCACCTGAGAGGAGAGTGCCAGAGTTGTGTGTACAACATGATGGGAAAAAGAGAAAAGAAACAAGGGGAATTTGGAAAGGCCAAGGGCAGCCGCGCCATCTGGTATATGTGGCTGGGGGCTAGATTTCTAGAGTTCGAAGCCCTTGGATTCTTGAACGAGGATCACTGGATGGGGAGAGAGAACTCAGGAGGTGGTGTTGAAGGGCTGGGATTACAAAGACTCGGATATGTCCTAGAAGAGATGAGTCGTATACCAGGAGGAAGGATGTATGCAGATGACACTGCTGGCTGGGACACCCGCATTAGCAGGTTTGATCTGGAGAATGAAGCTCTAATCACCAACCAAATGGAGAAAGGGCACAGGGCCTTGGCATTGGCCATAATCAAGTACACATACCAAAACAAAGTGGTAAAGGTCCTTAGACCAGCTGAAAAAGGGAAAACAGTTATGGACATTATTTCGAGACAAGACCAAAGGGGGAGCGGACAAGTTGTCACTTACGCTCTTAACACATTTACCAACCTAGTGGTGCAACTCATTCGGAATATGGAGGCTGAGGAAGTTCTAGAGATGCAAGACTTGTGGCTGCTGCGGAGGTCAGAGAAAGTGACCAACTGGTTGCAGAGCAACGGATGGGATAGGCTCAAACGAATGGCAGTCAGTGGAGATGATTGCGTTGTGAAGCCAATTGATGATAGGTTTGCACATGCCCTCAGGTTCTTGAATGATATGGGAAAAGTTAGGAAGGACACACAAGAGTGGAAACCCTCAACTGGATGGGACAACTGGGAAGAAGTTCCGTTTTGCTCCCACCATTTCAACAAGCTCCATCTCAAGGACGGGAGGTCCATTGTGGTTCCCTGCCGCCACCAAGATGAACTGATTGGCCGGGCCCGCGTCTCTCCAGGGGCGGGATGGAGCATCCGGGAGACTGCTTGCCTAGCAAAATCATATGCGCAAATGTGGCAGCTCCTTTATTTCCACAGAAGGGACCTCCGACTGATGGCCAATGCCATTTGTTCATCTGTGCCAGTTGACTGGGTTCCAACTGGGAGAACTACCTGGTCAATCCATGGAAAGGGAGAATGGATGACCACTGAAGACATGCTTGTGGTGTGGAACAGAGTGTGGATTGAGGAGAACGACCACATGGAAGACAAGACCCCAGTTACGAAATGGACAGACATTCCCTATTTGGGAAAAAGGGAAGACTTGTGGTGTGGATCTCTCATAGGGCACAGACCGCGCACCACCTGGGCTGAGAACATTAAAAACACAGTCAACATGGTGCGCAGGATCATAGGTGATGAAGAAAAGTACATGGATTACCTATCCACCCAAGTTCGCTACTTGGGTGAAGAAGGGTCTACACCTGGAGTGCTGTAA

>ENA|AMM39806|AMM39806.2 Zika virus polyprotein

ATGAAAAACCC-AAAAAAGAAATCCGGAGGATTCCGGATTGTCAATATGCTAAAACGCGGAGTAGCCCGTGTGAGCCCCTTTGGGGGCTTGAAGAGGCTGCCAGCCGGACTTCTGCTGGGTCATGGGCCCATCAGGATGGTCTTGGCGATTCTAGCCTTCTTGAGATTCACGGCAATCAAGCCATCACTGGGTCTCATCAATAGATGGGGTTCAGTGGGGAAAAAAGAGGCTATGGAAATAATAAAGAAGTTCAAGAAAGATCTGGCTGCCATGCTGAGAATAATCAATGCTAGGAAGGAGAAGAAGAGACGAGGCGCAGATACTAATGTCGGAATTGTTGGCCTCCTGCTGACCACAGCTATGGCAGCGGAGGTCACTAGACGTGGGAGTGCATACTATATGTACTTGGACAGAAACGATGCTGGGGAGGCCATATCTTTTCCAACCACATTGGGGATGAATAAGTGTTATATACAGATCATGGATCTTGGACACATGTGTGATGCCACCATGAGCTATGAATGCCCTATGCTGGATGAGGGGGTGGAACCAGATGACGTCGATTGTTGGTGCAACACGACGTCAACTTGGGTTGTGTACGGAACCTGCCATCACAAAAAAGGTGAAGCACGGAGATCTAGAAGAGCTGTGACGCTCCCCTCCCATTCCACTAGGAAGCTGCAAACGCGGTCGCAAACTTGGTTGGAATCAAGAGAATACACAAAGCACTTGATTAGAGTCGAAAATTGGATATTCAGGAACCCTGGCTTCGCGTTAGCAGCAGCTGCCATCGCTTGGCTTTTGGGAAGCTCAACGAGCCAAAAAGTCATATACTTGGTCATGATACTGCTGATTGCCCCGGCATACAGCATCAGGTGCATAGGAGTCAGCAATAGGGACTTTGTGGAAGGTATGTCAGGTGGGACTTGGGTTGATGTTGTCTTGGAACATGGAGGTTGTGTCACCGTAATGGCACAGGACAAACCGACTGTCGACATAGAGCTGGTTACAACAACAGTCAGCAACATGGCGGAGGTAAGATCCTACTGCTATGAGGCATCAATATCGGACATGGCTTCGGACAGCCGCTGCCCAACACAAGGTGAAGCCTACCTTGACAAGCAATCAGACACTCAATATGTCTGCAAAAGAACGTTAGTGGACAGAGGCTGGGGAAATGGATGTGGACTTTTTGGCAAAGGGAGCCTGGTGACATGCGCTAAGTTTGCATGCTCCAAGAAAATGACCGGGAAGAGCATCCAGCCAGAGAATCTGGAGTACCGGATAATGCTGTCAGTTCATGGCTCCCAGCACAGTGGGATGATCGTTAATGACACAGGACATGAAACTGATGAGAATAGAGCGAAGGTTGAGATAACGCCCAATTCACCAAGAGCCGAAGCCACCCTGGGGGGTTTTGGAAGCCTAGGACTTGATTGTGAACCGAGGACAGGCCTTGACTTTTCAGATTTGTATTACTTGACTATGAATAACAAGCACTGGTTGGTTCACAAGGAGTGGTTCCACGACATTCCATTACCTTGGCACGCTGGGGCAGACACCGGAACTCCACACTGGAACAACAAAGAAGCACTGGTAGAGTTCAAGGACGCACATGCCAAAAGGCAAACTGTCGTGGTTCTAGGGAGTCAAGAAGGAGCAGTTCACACGGCCCTTGCTGGAGCTCTGGAGGCTGAGATGGATGGTGCAAAGGGAAGGCTGTCCTCTGGCCACTTGAAATGTCGCCTGAAAATGGATAAACTTAGATTGAAGGGCGTGTCATACTCCTTGTGTACCGCAGCGTTCACATTCACCAAGATCCCGGCTGAAACACTGCACGGGACAGTCACAGTGGAGGTACAGTACGCAGGGACAGATGGACCTTGCAAGGTTCCAGCTCAGATGGCGGTGGACATGCAAACTCTGACCCCAGTTGGGAGGCTGATAACCGCTAACCCCGTAATCACTGAAAGCACTGAGAACTCCAAGATGATGCTGGAACTTGATCCACCATTTGGGGACTCTTACATTGTCATAGGAGTCGGGGAGAAGAAGATCACCCACCACTGGCACAGGAGTGGCAGCACCATTGGAAAAGCATTTGAAGCCACTGTGAGAGGTGCCAGGAGAATGGCAGTCTTGGGAGACACAGCCTGGGACTTTGGATCAGTTGGAGGCGCTCTCAACTCATTGGGCAAGGGCATCCATCAAATTTTTGGAGCAGCTTTCAAATCATTGTTTGGAGGAATGTCCTGGTTCTCACAAATTCTCATTGGAACGTTGCTGATGTGGTTGGGTCTGAACACAAAGAATGGATCTATTTCCCTTATGTGCTTGGCCTTAGGGGGAGTGTTGATCTTCTTATCCACAGCCGTCTCTGCTGATGTGGGGTGCTCGGTGGACTTCTCAAAGAAGGAGACGAGATGCGGTACAGGGGTGTTCGTCTATAACGACGTTGAAGCCTGGAGGGACAGGTACAAGTACCATCCTGACTCCCCCCGTAGATTGGCAGCAGCAGTCAAGCAAGCCTGGGAAGATGGTATCTGTGGGATCTCCTCTGTTTCAAGAATGGAAAACATCATGTGGAGATCAGTAGAAGGGGAGCTCAACGCAATCCTGGAAGAGAATGGAGTTCAACTGACGGTCGTTGTGGGATCTGTAAAAAACCCCATGTGGAGAGGTCCACAGAGATTGCCCGTGCCTGTGAACGAGCTGCCCCACGGCTGGAAGGCTTGGGGGAAATCGTACTTCGTCAGAGCAGCAAAGACAAATAACAGCTTTGTCGTGGATGGTGACACACTGAAGGAATGCCCACTCAAACATAGAGCATGGAACAGCTTTCTTGTGGAGGATCATGGGTTCGGGGTATTTCACACTAGTGTCTGGCTCAAGGTTAGAGAAGATTATTCATTAGAGTGTGATCCAGCCGTTATTGGAACAGCTGTTAAGGGAAAGGAGGCTGTACACAGTGATCTAGGCTACTGGATTGAGAGTGAGAAGAATGACACATGGAGGCTGAAGAGGGCCCATCTGATCGAGATGAAAACATGTGAATGGCCAAAGTCCCACACATTGTGGACAGATGGAATAGAAGAGAGTGATCTGATCATACCCAAGTCTTTAGCTGGGCCACTCAGCCATCACAATACCAGAGAGGGCTACAGGACCCAAATGAAAGGGCCATGGCACAGTGAAGAGCTTGAAATTCGGTTTGAGGAATGCCCAGGCACCAAGGTCCACGTGGAGGAAACATGTGGAACAAGAGGACCATCTCTGAGATCAACCACTGCAAGCGGAAGGGTGATCGAGGAATGGTGCTGCAGGGAGTGCACAATGCCCCCACTGTCGTTCCAGGCTAAAGATGGCTGTTGGTATGGAATGGAGATAAGGCCCAGGAAAGAACCAGAAAGTAACTTAGTAAGGTCAATGGTGACTGCAGGATCAACTGATCACATGGATCACTTCTCCCTTGGAGTGCTTGTGATTCTGCTCATGGTGCAGGAAGGGCTGAAGAAGAGAATGACCACAAAGATCATCATAAGCACATCAATGGCAGTGCTGGTAGCTATGATCCTGGGAGGATTTTCAATGAGTGACCTGGCTAAGCTTGCAATTTTGATGGGTGCCACCTTCGCGGAAATGAACACTGGAGGAGATGTAGCTCATCTGGCGCTGATAGCGGCATTCAAAGTCAGACCAGCGTTGCTGGTATCTTTCATCTTCAGAGCTAATTGGACACCCCGTGAAAGCATGCTGCTGGCCTTGGCCTCGTGTCTTTTGCAAACTGCGATCTCCGCCTTGGAAGGCGACCTGATGGTTCTCATCAATGGTTTTGCTTTGGCCTGGTTGGCAATACGAGCGATGGTTGTTCCACGCACTGATAACATCACCTTGGCAATCCTGGCTGCTCTGACACCACTGGCCCGGGGCACACTGCTTGTGGCGTGGAGAGCAGGCCTTGCTACTTGCGGGGGGTTTATGCTCCTCTCTCTGAAGGGAAAAGGCAGTGTGAAGAAGAACTTACCATTTGTCATGGCCCTGGGACTAACCGCTGTGAGGCTGGTCGACCCCATCAACGTGGTGGGACTGCTGTTGCTCACAAGGAGTGGGAAGCGGAGCTGGCCCCCTAGCGAAGTACTCACAGCTGTTGGCCTGATATGCGCATTGGCTGGAGGGTTCGCCAAGGCAGATATAGAGATGGCTGGGCCCATGGCCGCGGTCGGTCTGCTAATTGTCAGTTACGTGGTCTCAGGAAAGAGTGTGGACATGTACATTGAAAGAGCAGGTGACATCACATGGGAAAAAGATGCGGAAGTCACTGGAAACAGTCCCCGGCTTGATGTGGCGCTAGATGAGAGTGGTGATTTCTCCCTGGTGGAGGATGACGGTCCCCCCATGAGAGAGATCATACTCAAGGTGGTCCTGATGACCATCTGTGGCATGAACCCAATAGCCATACCCTTTGCAGCTGGAGCGTGGTACGTATACGTGAAGACTGGAAAAAGGAGTGGAGCTCTATGGGATGTGCCTGCTCCCAAGGAAGTAAAAAAGGGGGAGACCACAGATGGAGTGTACAGAGTGATGACTCGTAGACTGCTAGGTTCAACACAAGTTGGAGTGGGAGTTATGCAAGAGGGGGTCTTTCACACCATGTGGCACGTCACAAAAGGATCCGCGCTGAGAAGCGGTGAAGGGAGACTTGATCCATACTGGGGAGATGTCAAGCAGGATCTGGTGTCATACTGTGGTCCATGGAAGCTAGATGCCGCCTGGGACGGGCACAGCGAGGTGCAGCTCTTGGCCGTGCCCCCCGGAGAGAGAGCGAGGAACATCCAGACTCTGCCCGGAATATTTAAGACAAAGGATGGGGACATTGGAGCGGTTGCGCTGGATTACCCAGCAGGAACTTCAGGATCTCCAATCCTAGACAAGTGTGGGAGAGTGATAGGACTTTATGGCAATGGGGTCGTGATCAAAAATGGGAGTTATGTTAGTGCCATCACCCAAGGGAGGAGGGAGGAAGAGACTCCTGTTGAGTGCTTCGAGCCTTCGATGCTGAAGAAGAAGCAGCTAACTGTCTTAGACTTGCATCCTGGAGCTGGGAAAACCAGGAGAGTTCTTCCTGAAATAGTCCGTGAAGCCATAAAAACAAGACTCCGTACTGTGATCTTAGCTCCAACCAGGGTTGTCGCTGCCGAAATGGAGGAAGCCCTTAGAGGGCTTCCAGTGCGTTATATGACAACAGCAGTCAATGTCACCCACTCTGGAACAGAAATCGTCGACTTAATGTGCCATGCCACCTTCACTTCACGTCTACTACAGCCAATCAGAGTCCCCAACTATAATCTGTATATTATGGATGAGGCCCACTTCACAGATCCCTCAAGTATAGCAGCAAGAGGATACATTTCAACAAGGGTTGAGATGGGCGAGGCGGCTGCCATCTTCATGACCGCCACGCCACCAGGAACCCGTGACGCATTTCCGGACTCCAACTCACCAATTATGGACACCGAAGTGGAAGTCCCAGAGAGAGCCTGGAGCTCAGGCTTTGATTGGGTGACGGATCATTCTGGAAAAACAGTCTGGTTTGTTCCAAGCGTGAGGAACGGCAATGAGATCGCAGCTTGTCTGACAAAGGCTGGAAAACGGGTCATACAGCTCAGCAGAAAGACTTTTGAGACAGAGTTCCAGAAAACAAAACATCAAGAGTGGGACTTTGTCGTGACAACTGACATTTCAGAGATGGGCGCCAACTTTAAAGCTGACCGTGTCATAGATTCCAGGAGATGCCTAAAGCCGGTCATACTTGATGGCGAGAGAGTCATTCTGGCTGGACCCATGCCTGTCACACATGCCAGCGCTGCCCAGAGGAGGGGGCGCATAGGCAGGAATCCCAACAAACCTGGAGATGAGTATCTGTATGGAGGTGGGTGCGCAGAGACTGACGAAGACCATGCACACTGGCTTGAAGCAAGAATGCTCCTTGACAATATTTACCTCCAAGATGGCCTCATAGCCTCGCTCTATCGACCTGAGGCCGACAAAGTAGCAGCCATTGAGGGAGAGTTCAAGCTTAGGACGGAGCAAAGGAAGACCTTTGTGGAACTCATGAAAAGAGGAGATCTTCCTGTTTGGCTGGCCTATCAGGTTGCATCTGCCGGAATAACCTACACAGATAGAAGATGGTGCTTTGATGGCACGACCAACAACACCATAATGGAAGACAGTGTGCCGGCAGAGGTGTGGACCAGACACGGAGAGAAAAGAGTGCTCAAACCGAGGTGGATGGACGCCAGAGTTTGTTCAGATCACGCGGCCCTGAAGTCATTCAAGGAGTTTGCCGCTGGGAAAAGAGGAGCGGCTTTTGGAGTGATGGAAGCCTTGGGAACACTGCCAGGACACATGACAGAGAGATTCCAGGAAGCCATTGACAACCTCGCTGTGCTCATGCGGGCAGAGACTGGAAGCAGGCCTTACAAAGCCGCGGCGGCCCAATTGCCGGAGACCCTAGAGACCATTATGCTTTTGGGGTTGCTGGGAACAGTCTCGCTGGGAATCTTTTTCGTCTTGATGAGGAACAAGGGCATAGGGAAGATGGGCTTTGGAATGGTGACTCTTGGGGCCAGCGCATGGCTCATGTGGCTCTCGGAAATTGAGCCAGCCAGAATTGCATGTGTCCTCATTGTTGTGTTCCTATTGCTGGTGGTGCTCATACCTGAGCCAGAAAAGCAAAGATCTCCCCAGGACAACCAAATGGCAATCATCATCATGGTAGCAGTAGGTCTTCTGGGCTTGATTACCGCCAATGAACTCGGATGGTTGGAGAGAACAAAGAGTGACCTAAGCCATCTAATGGGAAGGAGAGAGGAGGGGGCAACCATAGGATTCTCAATGGACATTGACCTGCGGCCAGCCTCAGCTTGGGCCATCTACGCTGCCTTGACAACTTTCATTACCCCAGCCGTCCAACATGCAGTGACCACTTCATACAACAACTACTCCTTAATGGCGATGGCCACGCAAGCTGGAGTGTTGTTTGGTATGGGCAAAGGGATGCCATTCTACGCATGGG-ACTTTGGAGTCCCGCTGCTAATGATAGGTTGCTACTCACAATTAACACCCCTGACCCTAATAGTAGCCATCATTTTGCTCGTGGCGCACTACATGTACTTGATCCCAGGGCTGCAGGCAGCAGCTGCGCGTGCTGCCCAGAAGAGAACGGCAGCTGGCATCATGAAGAACCCTGTTGTGGATGGAATAGTGGTGACTGACATTGACACAATGACAATTGACCCCCAAGTGGAGAAAAAGATGGGACAGGTGCTACTCATAGCAGTAGCCGTCTCCAGCGCCATACTGTCGCGGACCGCCTGGGGGTGGGGGGAGGCTGGGGCCCTGATCACAGCTGCAACTTCCACTTTGTGGGAAGGCTCTCCGAACAAGTACTGGAACTCCTCTACAGCCACTTCACTGTGTAACATTTTTAGGGGAAGTTACTTGGCTGGAGCTTCTCTAATCTACACAGTAACAAGAAACGCTGGCTTGGTCAAGAGACGTGGGGGTGGAACAGGAGAGACCCTGGGAGAGAAATGGAAGGCCCGCTTGAACCAGATGTCGGCCCTGGAGTTCTACTCCTACAAAAAGTCAGGCATCACCGAGGTGTGCAGAGAAGAGGCCCGCCGCGCCCTCAAGGACGGTGTGGCAACGGGAGGCCATGCTGTGTCCCGAGGAAGTGCAAAGCTGAGATGGTTGGTGGAGCGGGGATACCTGCAGCCCTATGGAAAGGTCATTGATCTTGGATGTGGCAGAGGGGGCTGGAGTTACTACGCCGCCACCATCCGCAAAGTTCAAGAAGTGAAAGGATACACAAAAGGAGGCCCTGGTCATGAAGAACCCATGTTGGTGCAAAGCTATGGGTGGAACATAGTCCGTCTTAAGAGTGGGGTGGACGTCTTTCATATGGCGGCTGAGCCGTGTGACACGTTGCTGTGTGACATAGGTGAGTCATCATCTAGTCCTGAAGTGGAAGAAGCACGGACGCTCAGAGTCCTTTCCATGGTGGGGGATTGGCTTGAAAAAAGACCAGGAGCCTTTTGTATAAAAGTGTTGTGCCCATACACCAGCACTATGATGGAAACCCTGGAGCGACTGCAGCGTAGGTATGGAGGAGGACTGGTCAGAGTGCCACTCTCCCGCAACTCTACACATGAGATGTACTGGGTCTCTGGAGCGAAAAGCAACACCATAAAAAGTGTGTCCACCACGAGCCAGCTCCTCTTGGGGCGCATGGACGGGCCCAGGAGGCCAGTGAAATATGAGGAGGATGTGAATCTCGGCTCTGGCACGCGGGCTGTGGTAAGCTGCGCTGAAGCTCCCAACATGAAGATCATTGGTAACCGCATTGAAAGGATCCGCAGTGAGCACGCGGAAACGTGGTTCTTTGACGAGAACCACCCATATAGGACATGGGCTTACCATGGAAGCTATGAGGCCCCCACACAAGGGTCAGCGTCCTCTCTAATAAACGGGGTTGTCAGGCTCCTGTCAAAACCCTGGGATGTGGTGACTGGAGTCACAGGAATAGCCATGACCGACACCACACCGTATGGTCAGCAAAGAGTTTTCAAGGAAAAAGTGGACACTAGGGTGCCAGATCCCCAAGAAGGCACTCGTCAGGTTATGAGCATGGTCTCTTCCTGGTTGTGGAAAGAGCTAGGCAAACACAAACGGCCACGAGTCTGTACCAAAGAAGAGTTCATCAACAAGGTTCGTAGCAATGCAGCATTAGGGGCAATATTTGAAGAGGAAAAAGAGTGGAAGACTGCAGTGGAAGCTGTGAACGATCCAAGGTTCTGGGCTCTAGTGGACAAGGAAAGAGAGCACCACCTGAGAGGAGAGTGCCAGAGTTGTGTGTACAACATGATGGGAAAAAGAGAAAAGAAACAAGGGGAATTTGGAAAGGCCAAGGGCAGCCGCGCCATCTGGTATATGTGGCTAGGGGCTAGATTTCTAGAGTTCGAAGCCCTTGGATTCTTGAACGAGGATCACTGGATGGGGAGAGAGAACTCAGGAGGTGGTGTTGAAGGGCTGGGATTACAAAGACTCGGATATGTCCTAGAAGAGATGAGTCGCATACCAGGAGGAAGGATGTATGCAGATGACACTGCTGGCTGGGACACCCGCATCAGCAGGTTTGATCTGGAGAATGAAGCTCTAATCACCAACCAAATGGAGAAAGGGCACAGGGCCTTGGCATTGGCCATAATCAAGTACACATACCAAAACAAAGTGGTAAAGGTCCTTAGACCAGCTGAAAAAGGGAAGACAGTTATGGACATTATTTCGAGACAAGACCAAAGGGGGAGCGGACAAGTTGTCACTTACGCTCTTAACACATTTACCAACCTAGTGGTGCAACTCATTCGGAGTATGGAGGCTGAGGAAGTTCTAGAGATGCAAGACTTGTGGCTGCTGCGGAGGTCAGAGAAAGTGACCAACTGGCTGCAGAGCAACGGATGGGATAGGCTCAAACGAATGGCAGTCAGTGGAGATGATTGCGTTGTGAGGCCAATTGATGATAGGTTTGCACATGCCCTCAGGTTCTTGAATGATATGGGGAAAGTTAGGAAGGACACACAAGAGTGGAAACCCTCAACTGGATGGGACAACTGGGAGGAAGTTCCGTTTTGCTCCCACCACTTCAACAAGCTCCATCTCAAGGACGGGAGGTCCATTGTGGTTCCCTGCCGCCACCAAGATGAACTGATTGGCCGGGCCCGCGTCTCTCCAGGGGCGGGATGGAGCATCCGGGAGACTGCTTGCCTAGCAAAATCATATGCGCAAATGTGGCAGCTCCTTTATTTCCACAGAAGGGACCTCCGACTGATGGCCAATGCCATTTGTTCATCTGTGCCAGTTGACTGGGTTCCAACTGGGAGAACTACCTGGTCAATCCATGGAAAGGGAGAATGGATGACCACTGAAGACATGCTTGTGGTGTGGAACAGAGTGTGGATTGAGGAGAACGACCACATGGAAGACAAGACCCCAGTTACGAAATGGACAGACATTCCCTATTTGGGAAAAAGGGAAGACTTGTGGTGTGGATCTCTCATAGGGCACAGACCGCGCACCACCTGGGCTGAGAACATTAAAAACACAGTCAACATGGTGCGCAGGATCATAGGTGATGAAGAAAAGTACATGGACTACCTATCCACCCAAGTTCGCTACTTGGGTGAAGAAGGGTCTACACCTGGAGTGCTGTAA

>ENA|AMN14620|AMN14620.1 Zika virus polyprotein

ATGAAAAACCC-AAAAAAGAAATCCGGAGGATTCCGGATTGTCAATATGCTAAAACGCGGAGTAGCCCGTGTGAGCCCCTTTGGGGGCTTGAAGAGGCTGCCAGCCGGACTTCTGCTGGGTCATGGGCCCATCAGGATGGTCTTGGCGATTCTAGCCTTTTTGAGATTCACGGCAATCAAGCCATCACTGGGTCTCATCAATAGATGGGGTTCAGTGGGGAAAAAAGAGGCTATGGAAATAATAAAGAAGTTCAAGAAAGATCTGGCTGCCATGCTGAGAATAATCAATGCTAGGAAGGAGAAGAAGAGACGAGGCGCAGATACTAGTGTCGGAATTGTTGGCCTCCTGCTGACCACAGCTATGGCAGCGGAGGTCACTAGACGTGGGAGTGCATACTACATGTACTTGGACAGAAACGATGCTGGGGAGGCCATATCTTTTCCAACCACATTGGGGATGAATAAGTGTTATATACAGATCATGGATCTTGGACACATGTGTGATGCCACCATGAGCTATGAATGCCCTATGCTGGATGAGGGGGTGGAACCAGATGACGTCGATTGTTGGTGCAACACGACGTCAACTTGGGTTGTGTACGGAACCTGCCATCACAAAAAAGGTGAAGCACGGAGATCTAGAAGAGCTGTGACGCTCCCCTCCCATTCCACTAGGAAGCTGCAAACGCGGTCGCAAACCTGGTTGGAATCAAGAGAATACACAAAGCACTTGATTAGAGTCGAAAATTGGATATTCAGGAACCCTGGCTTCGCGTTAGCAGCAGCTGCCATCGCTTGGCTTTTGGGAAGCTCAACGAGCCAAAAAGTCATATACTTGGTCATGATACTGCTGATTGCCCCGGCATACAGCATCAGGTGCATAGGAGTCAGCAATAGGGACTTTGTGGAAGGTATGTCAGGTGGGACTTGGGTTGATGTTGTCTTGGAACATGGAGGTTGTGTCACCGTAATGGCACAGGACAAACCGACTGTCGACATAGAGCTGGTTACAACAACAGTCAGCAACATGGCGGAGGTAAGATCCTACTGCTATGAGGCATCAATATCAGACATGGCTTCGGACAGCCGCTGCCCAACACAAGGTGAAGCCTACCTTGACAAGCAATCAGACACTCAATATGTCTGCAAAAGAACGTTAGTGGACAGAGGCTGGGGAAATGGATGTGGACTTTTTGGCAAAGGGAGCCTGGTGACATGCGCTAAGTTTGCATGCTCCAAGAAAATGACCGGGAAGAGCATCCAGCCAGAGAATCTGGAGTACCGGATAATGCTGTCAGTTCATGGCTCCCAGCACAGTGGGATGATCGTTAATGACACTGGACATGAAACTGATGAGAATAGAGCGAAGGTTGAGATAACGCCCAATTCACCAAGAGCCGAAGCCACCCTGGGGGGTTTTGGAAGCCTAGGACTTGATTGTGAACCGAGGACAGGCCTTGACTTTTCAGATTTGTATTACTTGACTATGAATAACAAGCACTGGTTGGTTCACAAGGAGTGGTTCCACGACATTCCATTACCTTGGCACGCTGGGGCAGACACCGGAACTCCACACTGGAACAACAAAGAAGCACTGGTAGAGTTCAAGGACGCACATGCCAAAAGGCAAACTGTCGTGGTTCTAGGGAGTCAAGAAGGAGCAGTTCACACGGCCCTTGCTGGAGCTCTGGAGGCTGAGATGGATGGTGCAAAGGGAAGGCTGTCCTCTGGCCACTTGAAATGTCGCCTGAAAATGGATAAACTTAGATTGAAGGGCGTGTCATACTCCTTGTGTACCGCAGCGTTCACATTCACCAAGATCCCGGCTGAAACACTGCACGGGACAGTCACAGTGGAGGTACAGTACGCAGGGACAGATGGACCTTGCAAGGTTCCAGCTCAGATGGCGGTGGACATGCAAACTCTGACCCCAGTTGGGAGGTTGATAACCGCCAACCCCGTAATCACTGAAAGCACTGAGAACTCTAAGATGATGCTGGAACTTGATCCACCATTTGGGGACTCTTACATTGTCATAGGAGTCGGGGAGAAGAAGATCACCCACCACTGGCACAGGAGTGGCAGCACCATTGGAAAAGCATTTGAAGCCACTGTGAGAGGTGCCAAGAGAATGGCAGTCTTGGGAGACACAGCCTGGGACTTTGGATCAGTTGGAGGCGCTCTCAACTCATTGGGCAAGGGCATCCATCAAATTTTTGGAGCAGCTTTCAAATCATTGTTTGGAGGAATGTCCTGGTTCTCACAAATCCTCATTGGAACGTTGCTGATGTGGTTGGGTCTGAACACAAAGAATGGATCTATTTCCCTCATGTGCTTGGCCTTAGGGGGAGTGTTGATCTTCTTATCCACAGCCGTCTCTGCTGATGTGGGGTGCTCGGTGGACTTCTCAAAGAAGGAGACGAGATGCGGTACAGGGGTGTTCGTCTATAACGACGTTGAAGCCTGGAGGGACAGGTACAAGTACCATCCTGACTCCCCCCGTAGATTGGCAGCAGCAGTCAAGCAAGCCTGGGAAGATGGTATCTGCGGGATCTCCTCTGTTTCAAGAATGGAAAACATCATGTGGAGATCAGTAGAAGGGGAGCTCAATGCAATCCTGGAAGAGAATGGAGTTCAACTGACGGTCGTTGTGGGATCTGTAAAAAACCCCATGTGGAGAGGTCCACAGAGATTGCCCGTGCCTGTGAACGAGCTGCCCCACGGCTGGAAGGCTTGGGGGAAATCGTACTTCGTTAGAGCAGCAAAGACAAATAACAGCTTTGTCGTGGATGGTGACACACTGAAGGAATGCCCACTCAAACATAGAGCATGGAACAGCTTTCTTGTGGAGGATCATGGGTTCGGGGTATTTCACACTAGTGTCTGGCTCAAGGTTAGAGAAGATTATTCATTAGAGTGTGATCCAGCCGTTATTGGAACAGCTGTTAAGGGAAAGGAGGCTGTACACAGTGATCTAGGCTACTGGATTGAGAGTGAGAAGAATGACACATGGAGGCTGAAGAGGGCCCATCTGATCGAGATGAAAACATGTGAATGGCCAAAGTCCCACACATTGTGGACAGATGGAATAGAAGAGAGTGATCTGATCATACCCAAGTCTTTAGCTGGGCCACTCAGCCATCACAATACCAGAGAGGGCTACAGGACCCAAATGAAAGGGCCATGGCACAGTGAAGAGCTTGAAATTCGGTTTGAGGAATGCCCAGGCACTAAGGTCCACGTGGAGGAAACATGTGGAACAAGAGGACCATCTCTGAGATCAACCACTGCAAGCGGAAGGGTGATCGAGGAATGGTGCTGCAGGGAGTGCACAATGCCCCCACTGTCGTTCCGGGCTAAAGATGGCTGTTGGTATGGAATGGAGATAAGGCCCAGGAAAGAACCAGAAAGCAACTTAGTAAGGTCAGTGGTGACTGCAGGATCAACTGATCACATGGATCACTTCTCCCTTGGAGTGCTTGTGATTCTGCTCATGGTGCAGGAAGGGCTGAAGAAGAGAATGACCACAAAGATCATCATAAGCACATCAATGGCAGTGCTGGTAGCTATGATCCTGGGAGGATTTTCAATGAGTGACCTGGCTAAGCTTGCAATTTTGATGGGTGCCACCTTCGCGGAAATGAACACTGGAGGAGATGTAGCTCATCTGGCGCTGATAGCGGCATTCAAAGTCAGACCAGCGTTGCTGGTATCTTTCATCTTCAGAGCTAATTGGACACCCCGTGAAAGCATGCTGCTGGCCTTGGCCTCGTGTCTTTTGCAAACTGCGATCTCCGCCTTGGAAGGCGACCTGATGGTTCTCATCAATGGTTTTGCTTTGGCCTGGTTGGCAATACGAGCGATGGTTGTTCCACGCACTGATAACATCACCTTGGCAATCCTGGCTGCTCTGACACCACTGGCCCGGGGCACACTGCTTGTGGCGTGGAGAGCAGGCCTTGCTACTTGCGGGGGGTTTATGCTCCTCTCTCTGAAGGGAAAAGGCAGTGTGAAGAAGAACTTACCATTTGTCATGGCCCTGGGACTAACCGCTGTGAGGCTGGTCGACCCCATCAACGTGGTGGGACTGCTGTTGCTCACAAGGAGTGGGAAGCGGAGCTGGCCCCCTAGCGAAGTACTCACAGCTGTTGGCCTGATATGCGCATTGGCTGGAGGGTTCGCCAAGGCAGATATAGAGATGGCTGGGCCCATGGCCGCAGTCGGTCTGCTAATTGTCAGTTACGTGGTCTCAGGAAAGAGTGTGGACATGTACATTGAAAGAGCAGGTGACATCACATGGGAAAAAGATGCGGAAGTCACTGGAAACAGTCCCCGGCTCGATGTGGCGCTAGATGAGAGTGGTGATTTCTCCCTGGTGGAGGATGACGGTCCCCCCATGAGAGAGATCATACTCAAGGTGGTCCTGATGACCATCTGTGGCATGAACCCAATAGCCATACCCTTTGCAGCTGGAGCGTGGTACGTATACGTGAAGACTGGAAAAAGGAGTGGTGCTCTATGGGATGTGCCTGCTCCCAAGGAAGTAAAAAAGGGGGAGACCACAGATGGAGTGTACAGAGTAATGACTCGTAGACTGCTAGGTTCAACACAAGTTGGAGTGGGAGTTATGCAAGAGGGGGTCTTTCACACTATGTGGCACGTCACAAAAGGATCCGCGCTGAGAAGCGGTGAAGGGAGACTTGATCCATACTGGGGAGATGTCAAGCAGGATCTGGTGTCATACTGTGGTCCATGGAAGCTAGATGCCGCCTGGGACGGGCACAGCGAGGTGCAGCTCTTGGCCGTGCCCCCCGGAGAGAGAGCGAGGAACATCCAGACTCTGCCCGGAATATTTAAGACAAAGGATGGGGACATTGGAGCGGTTGCGCTGGATTACCCAGCAGGAACTTCAGGATCTCCAATCCTAGACAAGTGTGGGAGAGTGATAGGACTTTATGGCAATGGGGTCGTGATCAAAAATGGGAGTTATGTTAGTGCCATCACCCAAGGGAGGAGGGAGGAAGAGACTCCTGTTGAGTGCTTCGAGCCTTCGATGCTGAAGAAGAAGCAGCTAACTGTCTTAGACTTGCATCCTGGAGCTGGGAAAACCAGGAGAGTTCTTCCTGAAATAGTCCGTGAAGCTATAAAAACAAGACTCCGTACTGTGATCTTAGCTCCAACCAGGGTTGTCGCTGCTGAAATGGAGGAAGCCCTTAGAGGGCTTCCAGTGCGTTATATGACAACAGCAGTCAATGTCACCCATTCTGGAACAGAAATCGTCGACTTAATGTGCCATGCCACCTTCACTTCACGTCTACTACAGCCAATCAGAGTCCCCAACTATAATCTGTACATTATGGATGAGGCCCACTTCACAGATCCCTCAAGTATAGCAGCAAGAGGATACATTTCAACAAGGGTTGAGATGGGCGAGGCGGCTGCCATCTTCATGACCGCCACGCCACCAGGAACCCGTGACGCATTTCCGGACTCCAACTCACCAATTATGGACACCGAAGTGGAAGTCCCAGAGAGAGCCTGGAGCTCAGGCTTTGATTGGGTGACGGATCATTCTGGAAAAACAGTTTGGTTTGTTCCAAGCGTGAGGAACGGCAATGAGATCGCAGCTTGTCTGACAAAGGCTGGAAAACGGGTCATACAGCTCAGCAGAAAGACTTTTGAGACAGAGTTCCAGAAAACAAAACATCAAGAGTGGGACTTTGTCGTGACAACCGACATTTCAGAGATGGGCGCCAACTTTAAAGCTGACCGTGTCATAGATTCCAGGAGATGCCTAAAGCCGGTCATACTTGATGGCGAGAGAGTCATTCTGGCTGGACCCATGCCTGTCACACATGCCAGCGCTGCCCAGAGGAGGGGGCGCATAGGCAGGAATCCCAACAAACCTGGAGATGAGTATCTGTATGGAGGTGGGTGCGCAGAGACTGACGAAGACCATGCACACTGGCTTGAAGCAAGAATGCTCCTTGACAATATTTACCTCCAAGATGGCCTCATAGCCTCGCTCTATCGACCTGAGGCCGACAAAGTAGCAGCCATTGAGGGAGAGTTCAAGCTTAGGACGGAGCAAAGGAAGACCTTTGTGGAACTCATGAAAAGAGGAGATCTTCCTGTTTGGCTGGCCTATCAGGTTGCATCTGCCGGAATAACTTACACAGATAGAAGATGGTGCTTTGATGGCACGACCAACAACACCATAATGGAAGACAGTGTGCCGGCAGAGGTGTGGACCAGACACGGAGAGAAAAGAGTGCTCAAACCGAGGTGGATGGACGCCAGAGTTTGTTCAGATCATGCGGCCCTGAAGTCATTCAAGGAGTTTGCCGCTGGGAAAAGAGGAGCGGCTTTTGGAGTGATGGAAGCCCTGGGAACACTGCCAGGACACATGACAGAGAGATTCCAGGAAGCCATTGACAACCTCGCTGTGCTCATGCGGGCAGAGACTGGAAGCAGGCCTTACAAAGCCGCGGCGGCCCAATTGCCGGAGACCCTAGAGACCATTATGCTTTTGGGGTTGCTGGGAACAGTCTCGCTGGGAATCTTTTTCGTCTTGATGAGGAACAAGGGCATAGGGAAGATGGGCTTTGGAATGGTGACTCTTGGGGCCAGCGCATGGCTCATGTGGCTCTCGGAAATTGAGCCAGCCAGAATTGCATGTGTCCTCATTGTTGTGTTCCTACTGCTGGTGGTGCTCATACCTGAGCCAGAAAAGCAAAGATCTCCCCAGGACAACCAAATGGCAATCATCATCATGGTAGCAGTAGGTCTTCTGGGCTTGATCACCGCCAATGAACTCGGATGGTTGGAGAGAACAAAGAGTGACCTAAGCCATCTAATGGGAAGAAGAGAGGAGGGAGCAACCATAGGATTCTCAATGGACATTGACCTGCGGCCAGCCTCAGCTTGGGCCATCTATGCTGCCTTGACAACTTTCATTACCCCAGCCGTCCAACATGCAGTGACCACTTCATACAACAACTACTCCTTAATGGCGATGGCCACGCAAGCTGGAGTGTTGTTTGGTATGGGCAAAGGGATGCCATTCTACGCATGGG-ACTTTGGAGTCCCGCTGCTAATGATAGGTTGCTACTCACAATTAACACCCCTGACCCTAATAGTGGCCATCATTTTGCTCGTGGCGCACTACATGTACTTGATCCCAGGGCTGCAGGCAGCAGCTGCGCGTGCTGCCCAGAAGAGAACGGCAGCTGGCATCATGAAGAACCCTGTTGTGGATGGAATAGTGGTGACTGACATTGACACAATGACAATTGACCCCCAAGTGGAGAAAAAGATGGGACAGGTGCTACTCATAGCAGTAGCCGTCTCCAGCGCCATACTGTCGCGGACCGCCTGGGGGTGGGGGGAGGCTGGGGCCCTGATCACAGCCGCAACTTCCACTTTGTGGGAAGGCTCTCCGAACAAGTACTGGAACTCCTCTACAGCCACTTCACTGTGTAACATTTTTAGGGGAAGTTACTTGGCTGGAGCTTCTCTAATCTACACAGTAACAAGAAACGCTGGCTTGGTCAAGAGACGTGGGGGTGGAACAGGAGAGACCCTGGGAGAGAAATGGAAGGCCCGCTTGAACCAGATGTCGGCCCTGGAGTTCTACTCCTACAAAAAGTCAGGCATCACCGAGGTGTGCAGAGAAGAGGCCCGCCGCGCCCTCAAGGACGGTGTGGCAACGGGAGGCCATGCTGTGTCCCGAGGAAGTGCAAAGCTGAGATGGTTGGTGGAGCGGGGATACCTGCAGCCCTATGGAAAGGTCATTGATCTTGGATGTGGCAGAGGGGGCTGGAGTTACTACGCCGCCACCATCCGCAAAGTTCAAGAAGTGAAAGGATACACAAAAGGAGGCCCTGGTCATGAAGAACCCGTGTTGGTGCAAAGCTATGGGTGGAACATAGTCCGTCTCAAGAGTGGGGTGGACGTCTTTCATATGGCGGCTGAGCCGTGTGACACGTTGCTGTGTGACATAGGTGAGTCATCATCTAGTCCTGAAGTGGAAGAAGCACGGACGCTCAGAGTCCTCTCCATGGTGGGGGATTGGCTTGAAAAAAGACCAGGAGCCTTTTGTATAAAAGTGTTGTGCCCATACACCAGCACTATGATGGAAACCCTGGAGCGACTGCAGCGTAGGTATGGGGGAGGACTGGTCAGAGTGCCACTCTCCCGCAACTCTACACATGAGATGTACTGGGTCTCTGGAGCGAAAAGCAACACCATAAAAAGTGTGTCCACCACGAGCCAGCTCCTCTTGGGGCGCATGGACGGGCCTAGGAGGCCAGTGAAATATGAGGAGGATGTGAATCTCGGCTCTGGCACGCGGGCTGTGGTAAGCTGCGCTGAAGCTCCCAACATGAAGATCATTGGTAACCGCATTGAAAGGATCCGCAGTGAGCACGCGGAAACGTGGTTCTTTGACGAGAACCACCCATATAGGACATGGGCTTACCATGGAAGCTATGAGGCCCCCACACAAGGGTCAGCATCCTCTCTAGTAAACGGGGTTGTCAGGCTCCTGTCAAAACCCTGGGATGTGGTGACTGGAGTCACAGGAATAGCCATGACCGACACCACACCGTATGGTCAGCAAAGAGTTTTCAAGGAAAAAGTGGACACTAGGGTGCCAGACCCCCAAGAAGGCACTCGTCAGGTTATGAGCATGGTCTCTTCCTGGTTGTGGAAAGAGCTAGGCAAACACAAACGGCCACGAGTCTGTACCAAAGAAGAGTTCATCAACAAGGTTCGTAGCAATGCAGCATTAGGGGCAATATTTGAAGAGGAAAAAGAGTGGAAGACTGCAGTGGAAGCTGTGAACGATCCAAGGTTCTGGGCTCTAGTGGACAAGGAAAGAGAGCACCACCTGAGAGGAGAGTGCCAGAGTTGTGTGTACAACATGATGGGAAAAAGAGAAAAGAAACAAGGGGAATTTGGAAAGGCCAAGGGCAGCCGCGCCATCTGGTATATGTGGCTAGGGGCTAGATTTCTAGAGTTCGAAGCCCTTGGATTCTTGAACGAGGATCACTGGATGGGGAGAGAGAACTCAGGAGGTGGTGTTGAAGGGCTGGGATTACAAAGACTCGGATATGTCCTAGAAGAGATGAGTCGCATACCAGGAGGAAGGATGTATGCAGATGACACTGCTGGCTGGGATACCCGCATCAGCAGGTTTGATCTAGAGAATGAAGCTCTAATCACCAACCAAATGGAGAAAGGGCACAGGGCCTTGGCATTGGCCATAATCAAGTACACATACCAAAACAAAGTGGTAAAGGTCCTTAGACCAGCTGAAAAAGGGAAAACAGTTATGGACATTATTTCGAGACAAGACCAAAGGGGGAGCGGACAAGTTGTCACTTACGCTCTTAACACATTTACCAACCTAGTGGTGCAACTCATTCGGAATATGGAGGCTGAGGAAGTTCTAGAGATGCAAGACTTGTGGCTGCTGCGGAGGTCAGAGAAAGTGACCAACTGGTTGCAGAGCAACGGATGGGATAGGCTCAAACGAATGGCAGTCAGTGGAGATGATTGCGTTGTGAAGCCAATTGATGATAGGTTTGCACATGCCCTCAGGTTCTTGAATGATATGGGAAAAGTTAGGAAGGACACACAAGAGTGGAAACCCTCAACTGGATGGGACAACTGGGAAGAAGTTCCGTTTTGCTCCCACCACTTCAACAAGCTCCATCTCAAGGACGGGAGGTCCATTGTGGTTCCCTGCCGCCACCAAGATGAACTGATTGGCCGGGCCCGCGTCTCTCCAGGGGCGGGATGGAGCATCCGGGAGACTGCTTGCCTAGCAAAATCATATGCGCAAATGTGGCAGCTCCTTTATTTCCACAGAAGGGACCTCCGACTGATGGCCAATGCCATTTGTTCATCTGTGCCAGTTGACTGGGTTCCAACTGGGAGAACTACCTGGTCAATCCATGGAAAGGGAGAATGGATGACCACTGAAGACATGCTTGTGGTGTGGAACAGAGTGTGGATTGAGGAGAACGACCACATGGAAGACAAGACCCCAGTTACGAAATGGACAGACATTCCCTATTTGGGGAAAAGGGAAGACTTGTGGTGTGGATCTCTCATAGGGCACAGACCGCGCACCACCTGGGCTGAGAACATTAAAAACACAGTCAACATGGTGCGCAGGATCATAGGTGAGGAAGAAAAGTACATGGACTACCTATCCACCCAAGTTCGCTACTTGGGTGAAGAAGGGTCTACACCTGGAGTGCTGTAA

>ENA|AMD16557|AMD16557.1 Zika virus polyprotein

ATGAAAAACCC-AAAAAAGAAATCCGGAGGATTCCGGATTGTCAATATGCTAAAACGCGGAGTAGCCCGTGTGAGCCCCTTTGGGGGCTTGAAGAGGCTGCCAGCCGGACTTCTGCTGGGTCATGGGCCCATCAGGATGGTCTTGGCGATTCTAGCCTTTTTGAGATTCACGGCAATCAAGCCATCACTGGGTCTCATCAATAGATGGGGTTCAGTGGGGAAAAAAGAGGCTATGGAAATAATAAAGAAGTTCAAGAAAGATCTGGCTGCCATGCTGAGAATAATCAATGCCAGGAAGGAGAAGAAGAGACGAGGCGCAGATACTAGTGTCGGAATCGTTGGCCTCCTGCTGACCACAGCTATGGCAGCGGAGGTCACTAGACGTGGGAGTGCATACTATATGTACTTGGACAGAAACGATGCTGGGGAGGCCATATCTTTTCCAACCACATTGGGGATGAATAAGTGTTATATACAGATCATGGATCTTGGACACATGTGTGATGCCACCATGAGCTATGAATGCCCTATGCTGGATGAGGGGGTGGAACCAGATGACGTCGATTGTTGGTGCAACACGACGTCAACTTGGGTTGTGTACGGAACCTGCCATCACAAAAAAGGTGAAGCACGGAGATCTAGAAGAGCTGTGACGCTCCCCTCCCATTCCACTAGGAAGCTGCAAACGCGGTCGCAAACCTGGTTGGAATCAAGAGAATACACAAAGCACTTGATTAGAGTCGAAAATTGGATATTCAGGAACCCTGGCTTCGCGTTAGCAGCAGCTGCCATCGCTTGGCTTTTGGGAAGCTCAACGAGCCAAAAAGTCATATACTTGGTCATGATACTGCTGATTGCCCCGGCATACAGCATCAGGTGCATAGGAGTCAGCAATAGGGACTTTGTGGAAGGTATGTCAGGTGGGACTTGGGTTGATGTTGTCTTGGAACATGGGGGTTGTGTCACCGTAATGGCACAGGACAAACCGACTGTCGACATAGAGCTGGTTACAACAACAGTCAGCAACATGGCGGAGGTAAGATCCTACTGCTATGAGGCATCAATATCAGACATGGCTTCGGACAGCCGCTGCCCAACACAAGGTGAAGCCTACCTTGACAAGCAATCAGACACTCAATATGTCTGCAAAAGAACGTTAGTGGACAGAGGCTGGGGAAATGGATGTGGACTTTTTGGCAAAGGGAGCCTGGTGACATGCGCTAAGTTTGCATGCTCCAAGAAAATGACCGGGAAGAGCATCCAGCCAGAGAATCTGGAGTACCGGATAATGCTGTCAGTTCATGGCTCCCAGCACAGTGGGATGATCGTTAATGACACAGGACATGAAACTGATGAGAATAGAGCGAAGGTTGAGATAACGCCCAATTCACCAAGAGCCGAAGCCACCCTGGGGGGTTTTGGAAGCTTAGGACTTGATTGTGAACCGAGGACAGGCCTTGACTTTTCAGATTTGTATTACTTGACTATGAATAACAAGCACTGGTTGGTTCACAAGGAGTGGTTCCACGACATTCCATTACCTTGGCACGCTGGGGCAGACACCGGAACTCCACACTGGAACAACAAAGAAGCACTGGTAGAGTTCAAGGACGCACATGCCAAAAGGCAAACTGTCGTGGTTCTAGGGACTCAAGAAGGAGCAGTTCACACGGCCCTTGCTGGAGCTCTGGAGGCTGAGATGGATGGTGCAAAGGGAAGGCTGTCCTCTGGCCACTTGAAATGTCGCCTGAAAATGGATAAACTTAGATTGAAGGGCGTGTCATACTCCTTGTGTACCGCAGCGTTCACATTCACCAAGATCCCGGCTGAAACACTGCACGGGACAGTCACAGTGGAGGTACAGTACGCAGGGACAGATGGACCTTGCAAGGTTCCAGCTCAGATGGCGGTGGACATGCAAACTCTGACCCCAGTTGGGAGGTTGATAACCGCTAACCCCGTAATCACTGAAAGCACTGAGAACTCTAAGATGATGCTGGAACTTGATCCACCATTTGGGGACTCTTACATTGTCATAGGAGTCGGGGAGAAGAAGATCACCCACCACTGGCACAGGAGTGGCAGCACCATTGGAAAAGCATTTGAAGCCACTGTGAGAGGTGCCAAGAGAATGGCAGTCTTGGGAGACACAGCCTGGGACTTTGGATCAGTTGGAGGCGCTCTCAACTCATTGGGCAAGGGCATCCATCAAATTTTTGGAGCAGCTTTCAAATCATTGTTTGGAGGAATGTCCTGGTTCTCACAAATTCTCATTGGAACGTTGCTGATGTGGTTGGGTCTGAACACAAAGAATGGATCTATTTCCCTTATGTGCTTGGCCTTAGGGGGAGTGTTGATCTTCTTATCCACAGCCGTCTCTGCTGATGTGGGGTGCTCGGTGGACTTCTCAAAGAAGGAGACGAGATGTGGTACAGGGGTGTTCGTCTATAACGACGTTGAAGCCTGGAGGGACAGGTACAAGTACCATCCTGACTCTCCCCGTAGATTGGCAGCAGCAGTCAAGCAAGCCTGGGAAGATGGTATCTGCGGGATCTCCTCTGTTTCAAGAATGGAAAACATCATGTGGAGATCAGTAGAAGGGGAGCTTAACGCAATCCTGGAAGAGAATGGAGTTCAACTGACGGTCGTTGTGGGATCTGTAAAAAACCCCATGTGGAGAGGTCCACAGAGATTGCCCGTGCCTGTGAACGAGCTGCCCCACGGCTGGAAGGCTTGGGGGAAATCGTACTTCGTCAGAGCAGCAAAGACAAATAACAGCTTTGTCGTGGATGGTGACACACTGAAGGAATGCCCACTCAAACATAGAGCATGGAACAGCTTTCTTGTGGAGGATCATGGGTTCGGGGTATTTCACACTAGTGTCTGGCTCAAGGTTAGAGAAGATTATTCATTAGAGTGTGATCCAGCCGTTATTGGAACAGCTGTTAAGGGAAAGGAGGCTGTACACAGTGATCTAGGCTACTGGATTGAGAGTGAGAAGAATGACACATGGAGGCTGAAGAGGGCCCATCTGATCGAGATGAAAACATGTGAATGGCCAAAGTCCCACACATTGTGGACAGATGGAATAGAAGAGAGTGATCTGATCATACCCAAGTCTTTAGCTGGGCCACTCAGCCATCACAATACCAGAGAGGGCTACAGGACCCAAATGAAAGGGCCATGGCACAGTGAAGAGCTTGAAATTCGGTTTGAGGAATGCCCAGGCACTAAGGTCCACGTGGAGGAAACATGTGGAACAAGAGGACCATCTCTGAGATCAACCACTGCAAGCGGAAGGGTGATCGAGGAATGGTGCTGCAGGGAGTGCACAATGCCCCCACTGTCGTTCCGGGCTAAAGATGGCTGTTGGTATGGAATGGAGATAAGGCCCAGGAAAGAACCAGAAAGCAACTTAGTAAGGTCAATGGTGACTGCAGGATCAACTGATCACATGGATCACTTCTCCCTTGGAGTGCTTGTGATTCTGCTCATGGTGCAGGAAGGGCTGAAGAAGAGAATGACCACAAAGATCATCATAAGCACATCAATGGCAGTGCTGGTAGCTATGATCCTGGGAGGATTTTCAATGAGTGACCTGGCTAAGCTTGCAATTTTGATGGGTGCCACCTTCGCGGAAATGAACACTGGAGGAGATGTAGCTCATCTGGCGCTGATAGCGGCATTCAAAGTCAGACCAGCGTTGCTGGTATCTTTCATCTTCAGAGCTAATTGGACACCCCGTGAAAGCATGCTGCTGGCCTTGGCCTCGTGTTTTTTGCAAACTGCGATCTCCGCCTTGGAAGGCGACCTGATGGTTCTCATCAATGGTTTTGCTTTGGCCTGGTTGGCAATACGAGCGATGGTTGTTCCACGCACTGACAACATCACCTTGGCAATCCTGGCTGCTCTGACACCACTGGCCCGGGGCACACTGCTTGTGGCGTGGAGAGCAGGCCTTGCTACTTGCGGGGGGTTTATGCTCCTCTCTCTGAAGGGAAAAGGCAGTGTGAAGAAGAACTTACCATTTGTCATGGCCCTGGGACTAACCGCTGTGAGGCTGGTCGACCCCATCAACGTGGTGGGACTGCTGTTGCTCACAAGGAGTGGGAAGCGGAGCTGGCCCCCTAGCGAAGTACTCACAGCTGTTGGCCTGATATGCGCATTGGCTGGAGGGTTCGCCAAGGCAGATATAGAGATGGCTGGGCCCATGGCCGCGGTCGGTCTGCTAATTGTCAGTTACGTGGTCTCAGGAAAGAGTGTGGACATGTACATTGAAAGAGCAGGTGACATCACATGGGAAAAAGATGCGGAAGTCACTGGAAACAGTCCCCGGCTCGATGTGGCGCTAGATGAGAGTGGTGACTTCTCCCTGGTGGAGGATGACGGTCCCCCCATGAGAGAGATCATACTCAAGGTGGTCCTGATGACCATCTGTGGCATGAACCCAATAGCCATACCCTTTGCAGCTGGAGCGTGGTACGTATACGTGAAGACTGGAAAAAGGAGTGGTGCTCTATGGGATGTGCCTGCTCCCAAGGAAGTAAAAAAGGGGGAGACCACAGATGGAGTGTACAGAGTAATGACTCGTAGACTGCTAGGTTCAACACAAGTTGGAGTGGGAGTTATGCAAGAGGGGGTCTTTCACACTATGTGGCACGTCACAAAAGGATCCGCGCTGAGAAGCGGTGAAGGGAGACTTGATCCATACTGGGGAGATGTCAAGCAGGATCTGGTGTCATACTGTGGTCCATGGAAGCTAGATGCCGCCTGGGACGGGCACAGCGAGGTGCAGCTCTTGGCCGTGCCCCCCGGAGAGAGAGCGAGGAACATCCAGACTCTGCCCGGAATATTTAAGACAAAGGATGGGGACATTGGAGCGGTTGCGCTGGATTACCCAGCAGGAACTTCAGGATCTCCAATCCTAGACAAGTGTGGGAGAGTGATAGGACTTTATGGCAATGGGGTCGTGATAAAAAATGGGAGTTATGTTAGTGCCATCACCCAAGGGAGGAGGGAGGAAGAGACTCCTGTTGAGTGCTTCGAGCCTTCGATGCTGAAGAAGAAGCAGCTAACTGTCTTAGACTTGCATCCTGGAGCTGGGAAAACCAGGAGAGTTCTTCCTGAAATAGTCCGTGAAGCCATAAAAACAAGACTCCGTACTGTGATCTTAGCTCCAACCAGGGTTGTCGCTGCTGAAATGGAGGAAGCCCTTAGAGGGCTTCCAGTGCGTTATATGACAACAGCAGTCAATGTCACCCACTCTGGAACAGAAATCGTCGACTTAATGTGCCATGCCACCTTCACTTCACGTCTACTACAGCCAATCAGAGTCCCCAACTATAATCTGTATATTATGGATGAGGCCCACTTCACAGATCCCTCAAGCATAGCAGCAAGAGGATACATTTCAACAAGGGTTGAGATGGGCGAGGCGGCTGCCATCTTCATGACCGCCACGCCACCAGGAACCCGTGACGCATTTCCGGACTCCAACTCACCAATTATGGACACCGAAGTGGAAGTCCCAGAGAGAGCCTGGAGCTCAGGCTTTGATTGGGTGACGGATCATTCTGGAAAAACAGTTTGGTTTGTTCCAAGCGTGAGGAACGGCAATGAGATCGCAGCTTGTCTGACAAAGGCTGGAAAACGGGTCATACAGCTCAGCAGAAAGACTTTTGAGACAGAGTTCCAGAAAACAAAACATCAAGAGTGGGACTTTGTCGTGACAACTGACATTTCAGAGATGGGCGCCAACTTTAAAGCTGACCGTGTCATAGATTCCAGGAGATGCCTAAAGCCGGTCATACTTGATGGCGAGAGAGTCATTCTGGCTGGACCCATGCCTGTCACACATGCCAGCGCTGCCCAGAGGAGGGGGCGCATAGGCAGGAATCCCAACAAACCTGGAGATGAGTACCTGTATGGAGGTGGGTGCGCAGAGACTGACGAAGACCATGCACACTGGCTTGAAGCAAGAATGCTCCTTGACAATATTTACCTCCAAGATGGCCTCATAGCCTCGCTCTATCGACCTGAGGCCGACAAAGTAGCAGCCATTGAGGGAGAGTTCAAGCTTAGGACGGAGCAAAGGAAGACCTTTGTGGAACTCATGAAAAGAGGAGATCTTCCTGTTTGGCTGGCCTATCAGGTTGCATCTGCCGGAATAACCTACACAGATAGAAGATGGTGCTTTGATGGCACGACCAACAACACCATAATGGAAGACAGTGTGCCGGCAGAGGTGTGGACCAGACACGGAGAGAAAAGAGTGCTCAAACCGAGGTGGATGGACGCCAGAGTTTGTTCAGATCATGCGGCCCTGAAGTCATTCAAGGAGTTTGCCGCTGGGAAAAGAGGAGCGGCTTTTGGAGTGATGGAAGCCCTGGGAACACTGCCAGGACACATGACAGAGAGATTCCAGGAAGCCATTGACAACCTCGCTGTGCTCATGCGGGCAGAGACTGGAAGCAGGCCTTACAAAGCCGCGGCGGCCCAATTGCCGGAGACCCTAGAGACCATTATGCTTTTGGGGTTGCTGGGAACAGTCTCGCTGGGAATCTTTTTCGTCTTGATGAGGAACAAGGGCATAGGGAAGATGGGCTTTGGAATGGTGACTCTTGGGGCCAGCGCATGGCTCATGTGGCTCTCGGAAATTGAGCCAGCCAGAATTGCATGTGTCCTCATTGTTGTGTTCCTATTGCTGGTGGTGCTCATACCTGAGCCAGAAAAGCAAAGATCTCCCCAGGACAACCAAATGGCAATCATCATCATGGTAGCAGTAGGTCTTCTGGGCTTGATTACCGCCAATGAACTCGGATGGTTGGAGAGAACAAAGAGTGACCTAAGCCATCTAATGGGAAGGAGAGAGGAGGGGGCAACCATAGGATTCTCAATGGACATTGACCTGCGGCCAGCCTCAGCTTGGGCCATCTATGCTGCCTTGACAACTTTCATTACCCCAGCCGTCCAACATGCAGTGACCACTTCATACAACAACTACTCCTTAATGGCGATGGCCACGCAAGCTGGAGTGTTGTTTGGTATGGGCAAAGGGATGCCATTCTACGCATGGG-ACTTTGGAGTCCCGCTGCTAATGATAGGTTGCTACTCACAATTAACACCCCTGACCCTAATAGTGGCCATCATTTTGCTCGTGGCGCACTACATGTACTTGATCCCAGGGCTGCAGGCAGCAGCTGCGCGTGCTGCCCAGAAGAGAACGGCAGCTGGCATCATGAAGAACCCTGTTGTGGATGGAATAGTGGTGACTGACATTGACACAATGACAATTGACCCCCAAGTGGAGAAAAAGATGGGACAGGTGCTACTCATAGCAGTAGCCGTCTCCAGCGCCATACTGTCGCGGACCGCCTGGGGGTGGGGGGAGGCTGGGGCCCTGATCACAGCCGCAACTTCCACTTTGTGGGAAGGCTCTCCGAACAAGTACTGGAACTCCTCTACAGCCACTTCACTGTGTAACATTTTTAGGGGAAGTTACTTGGCTGGAGCTTCTCTAATCTACACAGTAACAAGAAACGCTGGCTTGGTCAAGAGACGTGGGGGTGGAACAGGAGAGACCCTGGGAGAGAAATGGAAGGCCCGCTTGAACCAGATGTCGGCCCTGGAGTTCTACTCCTACAAAAAGTCAGGCATCACCGAGGTGTGCAGAGAAGAGGCCCGCCGCGCCCTCAAGGACGGTGTGGCAACGGGAGGCCATGCTGTGTCCCGAGGAAGTGCAAAGCTGAGATGGTTGGTGGAGCGGGGATACCTGCAGCCCTATGGAAAGGTCATTGATCTTGGATGTGGCAGAGGGGGCTGGAGTTACTACGCCGCCACCATCCGCAAAGTTCAAGAAGTGAAAGGATACACAAAAGGAGGCCCTGGTCATGAAGAACCCGTGTTGGTGCAAAGCTATGGGTGGAACATAGTCCGTCTTAAGAGTGGGGTGGACGTCTTTCATATGGCGGCTGAGCCGTGTGACACGTTGCTGTGTGACATAGGTGAGTCATCATCTAGTCCTGAAGTGGAAGAAGCACGGACGCTCAGAGTCCTCTCCATGGTGGGGGATTGGCTTGAAAAAAGACCAGGAGCCTTTTGCATAAAAGTGTTGTGCCCATACACCAGCACTATGATGGAAACCCTGGAGCGACTGCAGCGTAGGTATGGGGGAGGACTGGTCAGAGTGCCACTCTCCCGCAACTCTACACATGAGATGTACTGGGTCTCTGGAGCGAAAAGCAACACCATAAAAAGTGTGTCCACCACGAGCCAGCTCCTCTTGGGGCGCATGGACGGGCCTAGGAGGCCAGTGAAATATGAGGAGGATGTGAATCTCGGCTCTGGCACGCGGGCTGTGGTAAGCTGCGCTGAAGCTCCCAACATGAAGATCATTGGTAACCGCATTGAAAGGATCCGCAGTGAGCACGCGGAAACGTGGTTCTTTGACGAAAACCACCCATATAGGACATGGGCTTACCATGGAAGCTATGTGGCCCCCACACAAGGGTCAGCGTCCTCTCTAATAAACGGGGTTGTCAGGCTCCTGTCAAAACCCTGGGATGTGGTGACTGGAGTCACAGGAATAGCCATGACCGACACCACACCGTATGGTCAGCAAAGAGTTTTCAAGGAAAAAGTGGACACTAGGGTGCCAGACCCCCAAGAAGGCACTCGTCAGGTTATGAGCATGGTCTCTTCCTGGTTGTGGAAAGAGCTAGGCAAACACAAACGACCACGAGTCTGTACCAAAGAAGAGTTCATCAACAAGGTTCGTAGCAATGCAGCATTAGGGGCAATATTTGAAGAGGAAAAAGAGTGGAAGACTGCAGTGGAAGCTGTGAACGATCCAAGGTTCTGGGCTCTAGTGGACAAGGAAAGAGAGCACCACCTGAGAGGAGAGTGCCAGAGTTGTGTGTACAACATGATGGGAAAAAGAGAAAAGAAACAAGGGGAATTTGGAAAGGCCAAGGGCAGCCGCGCCATCTGGTATATGTGGCTAGGGGCTAGATTTCTAGAGTTCGAAGCCCTTGGATTCTTGAACGAGGATCACTGGATGGGGAGAGAGAACTCAGGAGGTGGTGTTGAAGGGCTGGGATTACAAAGACTCGGATATGTCCTAGAAGAGATGAGTCGCATACCAGGAGGAAGGATGTATGCAGATGACACTGCTGGCTGGGACACCCGCATCAGCAGGTTTGATCTGGAGAATGAAGCTCTAATCACCAACCAAATGGAGAAAGGGCACAGGGCCTTGGCATTGGCCATAATCAAGTACACATACCAAAACAAAGTGGTAAAGGTCCTTAGACCAGCTGAAAAAGGGAAAACAGTTATGGACATTATTTCGAGACAAGACCAAAGGGGGAGCGGACAAGTTGTCACTTACGCTCTTAACACATTTACCAACCTAGTGGTGCAACTCATTCGGAATATGGAGGCTGAGGAAGTTCTAGAGATGCAAGACTTGTGGCTGCTGCGGAGGTCAGAGAAAGTGACCAACTGGTTGCAGAGCAACGGATGGGATAGGCTCAAACGAATGGCAGTCAGTGGAGATGATTGCGTTGTGAAGCCAATTGATGATAGGTTTGCACATGCCCTCAGGTTCTTGAATGATATGGGAAAAGTTAGGAAGGACACACAAGAGTGGAAACCCTCAACTGGATGGGACAACTGGGAAGAAGTTCCGTTTTGCTCCCACCACTTCAACAAGCTCCATCTCAAGGACGGGAGGTCCATTGTGGTTCCCTGCCGCCACCAAGATGAACTGATTGGCCGGGCCCGCGTCTCTCCAGGGGCGGGATGGAGCATCCGGGAGACTGCTTGCCTAGCAAAATCATATGCGCAAATGTGGCAGCTCCTTTATTTCCACAGAAGGGACCTCCGACTGATGGCCAATGCCATTTGTTCATCTGTGCCAGTTGACTGGGTTCCAACTGGGAGAACTACCTGGTCAATCCATGGAAAGGGAGAATGGATGACCACTGAAGACATGCTTGTGGTGTGGAACAGAGTGTGGATTGAGGAGAACGACCACATGGAAGACAAGACCCCAGTTACGAAATGGACAGACATTCCCTATTTGGGAAAAAGGGAAGACTTGTGGTGTGGATCTCTCATAGGGCACAGACCGCGCACCACCTGGGCTGAGAACATTAAAAATACAGTCAACATGGTGCGCAGGATCATAGGTGATGAAGAAAAGTACATGGACTACCTATCCACCCAAGTTCGCTACTTGGGTGAAGAAGGGTCTACACCTGGAGTGCTGTGA

>ENA|AMK49165|AMK49165.1 Zika virus polyprotein

ATGAAAAACCC-AAAAAAGAAATCCGGAGGATTCCGGATTGTCAATATGCTAAAACGCGGAGTAGCCCGTGTGAGCCCCTTTGGGGGCTTGAAGAGGCTGCCAGCCGGACTTCTGCTGGGTCATGGGCCCATCAGGATGGTCTTGGCGATTCTAGCCTTTTTGAGATTCACGGCAATCAAGCCATCACTGGGTCTCATCAATAGATGGGGTTCAGTGGGGAAAAAAGAGGCTATGGAAATAATAAAGAAGTTCAAGAAAGATCTGGCTGCCATGCTGAGAATAATCAATGCTAGGAAGGAGAAGAAGAGACGAAGCGCAGATACTAGTGTCGGAATTGTTGGCCTCCTGCTGACCACAGCTATGGCAGCGGAGGTCACTAGACGTGGGAGTGCATACTATATGTACTTGGACAGAAACGATGCTGGGGAGGCCATATCTTTTCCAACCACATTGGGGATGAATAAGTGTTATATACAGATCATGGATCTTGGACACATGTGTGATGCCACTATGAGCTATGAATGCCCTATGCTGGATGAGGGGGTGGAACCAGATGACGTCGATTGTTGGTGCAACACGACGTCAACTTGGGTTGTGTACGGAACCTGCCATCACAAAAAAGGTGAAGCACGGAGATCTAGAAGAGCTGTGACGCTCCCCTCCCATTCCACTAGGAAGCTGCAAACGCGGTCGCAAACCTGGTTGGAATCAAGAGAATACACAAAGCACTTGATTAGAGTCGAAAATTGGATATTCAGGAACCCTGGCTTCGCGTTAGCAGCAGCTGCCATCGCTTGGCTTTTGGGAAGCTCAACGAGCCAAAAAGTCATATACTTGGTCATGATACTGCTGATTGCCCCGGCATACAGCATCAGGTGCATAGGAGTCAGCAATAGGGACTTTGTGGAAGGTATGTCAGGTGGGACTTGGGTTGATGTTGTCTTGGAACATGGAGGTTGTGTCACCGTAATGGCACAGGACAAACCGACTGTCGACATAGAGCTGGTTACAACAACAGTCAGCAACATGGCGGAGGTAAGATCCTACTGCTATGAGGCATCAATATCAGACATGGCTTCGGACAGCCGCTGCCCAACACAAGGTGAAGCCTACCTTGACAAGCAATCAGACACTCAATATGTCTGCAAAAGAACGTTAGTGGACAGAGGCTGGGGAAATGGATGTGGACTTTTTGGCAAAGGGAGCCTGGTGACATGCGCTAAGTTTGCATGCTCCAAGAAAATGACCGGGAAGAGCATCCAGCCAGAGAATCTGGAGTACCGGATAATGCTGTCAGTTCATGGCTCCCAGCACAGTGGGATGATCGTTAATGACACAGGACATGAAACTGATGAGAATAGAGCGAAGGTTGAGATAACGCCCAATTCACCAAGAGCCGAAGCCACCCTGGGGGGTTTTGGAAGCCTAGGACTTGATTGTGAACCGAGGACAGGCCTTGACTTTTCAGATTTGTATTACTTGACTATGAATAACAAGCACTGGTTGGTTCACAAGGAGTGGTTCCACGACATTCCATTACCTTGGCACGCTGGGGCAGACACCGGAACTCCACACTGGAACAATAAAGAAGCACTGGTAGAGTTCAAGGACGCACATGCCAAAAGGCAAACTGTCGTGGTTCTAGGGAGTCAAGAAGGAGCAGTTCACACGGCCCTTGCTGGAGCTCTGGAGGCTGAGATGGATGGTGCAAAGGGAAGGCTGTCCTCTGGCCACTTGAAATGTCGCCTGAAAATGGATAAACTTAGATTGAAGGGCGTGTCATACTCCTTGTGTACCGCAGCGTTCACATTCACCAAGATCCCGGCTGAAACACTGCACGGGACAGTCACAGTGGAGGTACAGTACGCAGGGACAGATGGACCTTGCAAGGTTCCAGCTCAGATGGCGGTGGACACGCAAACTCTGACCCCAGTTGGGAGGTTGATAACCGCTAACCCCGTAATCACTGAAAGCACTGAGAACTCTAAGATGATGCTGGAACTTGATCCACCATTTGGGGACTCTTACATTGTCATAGGAGTCGGGGAGAAGAAGATCACCCACCACTGGCACAGGAGTGGCAGCACCATTGGAAAAGCATTTGAAGCCACTGTGAGAGGTGCCAAGAGAATGGCAGTCTTGGGAGACACAGCCTGGGACTTTGGATCAGTTGGAGGCGCTCTCAACTCATTGGGCAAGGGCATCCATCAAATTTTTGGAGCAGCTTTCAAATCATTGTTTGGAGGAATGTCCTGGTTCTCACAAATTCTCATTGGAACGTTGCTGATGTGGTTGGGTCTGAACACAAAGAATGGATCTATTTCCCTTATGTGCTTGGCCTTAGGGGGAGTGTTGATCTTCTTATCCACAGCCGTCTCTGCTGATGTGGGGTGCTCGGTGGACTTCTCAAAGAAGGAGACGAGATGCGGTACAGGGGTGTTCGTCTATAACGACGTTGAAGCCTGGAGGGACAGGTACAAGTACCATCCTGACTCCCCCCGTAGATTGGCAGCAGCAGTCAAGCAAGCCTGGGAAGATGGTATCTGCGGGATCTCCTCTGTTTCAAGAATGGAAAACATCATGTGGAGATCAGTAGAAGGGGAGCTCAACGCAATCCTGGAAGAGAATGGAGTTCAACTGACGGTCGTTGTGGGATCTGTAAAAAACCCCATGTGGAGAGGTCCACAGAGATTGCCCGTGCCTGTGAACGAGCTGCCCCACGGCTGGAAGGCTTGGGGGAAATCGTACTTCGTCAGAGCAGCAAAGACAAATAACAGCTTTGTCGTGGATGGTGACACACTGAAGGAATGCCCACTCAAACATAGAGCATGGAACAGCTTTCTTGTGGAGGATCATGGGTTCGGGGTATTTCACACTAGTGTCTGGCTCAAGGTTAGAGAAGATTATTCATTAGAGTGTGATCCAGCCGTTATTGGAACAGCTGTTAAGGGAAAGGAGGCTGTACACAGTGATCTAGGCTACTGGATTGAGAGTGAGAAGAATGACACATGGAGGCTGAAGAGGGCCCATCTGATCGAGATGAAAACATGTGAATGGCCAAAGTCCCACACATTGTGGACAGATGGAATAGAAGAGAGTGATCTGATCATACCCAAGTCTTTAGCTGGGCCACTCAGCCATCACAATACCAGAGAGGGCTACAGGACCCAAATGAAAGGGCCATGGCACAGTGAAGAGCTTGAAATTCGGTTTGAGGAATGCCCAGGCACTAAGGTCCACGTGGAGGAAACATGTGGAACAAGAGGACCATCTCTGAGATCAACCACTGCAAGCGGAAGGGTGATCGAGGAATGGTGCTGCAGGGAGTGCACAATGCCCCCACTGTCGTTCCGGGCTAAAGATGGCTGTTGGTATGGAATGGAGATAAGGCCCAGGAAAGAACCAGAAAGCAACTTAGTAAGGTCAATGGTGACTGCAGGATCAACTGATCACATGGATCACTTCTCCCTTGGAGTGCTTGTGATTCTGCTCATGGTGCAGGAAGGGCTGAAGAAGAGAATGACCACAAAGATCATCATAAGCACATCAATGGCAGTGCTGGTAGCTATGATCCTGGGAGGATTTTCAATGAGTGACCTGGCTAAGCTTGCAATTTTGATGGGTGCCACCTTCGCGGAAATGAACACTGGAGGAGATGTAGCTCATCTGGCGCTGATAGCGGCATTCAAAGTCAGACCAGCGTTGCTGGTATCTTTCATCTTCAGAGCTAATTGGACACCCCGTGAAAGCATGCTGCTGGCCTTGGCCTCGTGTCTTTTGCAAACTGCGATCTCCGCCTTGGAAGGCGACCTGATGGTTCTCATCAATGGTTTTGCTTTGGCCTGGTTGGCAATACGAGCGATGGTTGTTCCACGCACTGATAACATCACCTTGGCAATCCTGGCTGCTCTGACACCACTGGCCCGGGGCACACTGCTTGTGGCGTGGAGAGCAGGCCTTGCTACTTGCGGGGGGTTTATGCTCCTCTCTCTGAAGGGAAAAGGCAGTGTGAAGAAGAACTTACCATTTGTCATGGCCCTGGGACTAACCGCTGTGAGGCTGGTCGACCCCATCAACGTGGTGGGACTGCTGTTGCTCACAAGGAGTGGGAAGCGGAGCTGGCCCCCTAGCGAAGTACTCACAGCTGTTGGCCTGATATGCGCATTGGCTGGAGGGTTCGCCAAGGCAGATATAGAGATGGCTGGGCCCATGGCCGCGGTCGGTCTGCTAATTGTCAGTTACGTGGTCTCAGGAAAGAGTGTGGACATGTACATTGAAAGAGCAGGTGACATCACATGGGAAAAAGATGCGGAAGTCACTGGAAACAGTCCCCGGCTCGATGTGGCGCTAGATGAGAGTGGTGATTTCTCCCTGGTGGAGGATGACGGTCCCCCCATGAGAGAGATCATACTCAAGGTGGTCCTGATGACCATCTGTGGCATGAACCCAATAGCCATACCCTTTGCAGCTGGAGCGTGGTACGTATACGTGAAGACTGGAAAAAGGAGTGGTGCTCTATGGGATGTGCCTGCTCCCAAGGAAGTAAAAAAGGGGGAGACCACAGATGGAGTGTACAGAGTAATGACTCGTAGACTGCTAGGTTCAACACAAGTTGGAGTGGGAGTTATGCAAGAGGGGGTCTTTCACACTATGTGGCACGTCACAAAAGGATCCGCGCTGAGAAGCGGTGAAGGGAGACTTGATCCATACTGGGGAGATGTCAAGCAGGATCTGGTGTCATACTGTGGTCCATGGAAGCTAGATGCCGCCTGGGACGGGCACAGCGAGGTGCAGCTCTTGGCCGTGCCCCCCGGAGAGAGAGCGAGGAACATCCAGACTCTGCCCGGAATATTTAAGACAAAGGATGGGGACATTGGAGCGGTTGCGCTGGATTACCCAGCAGGAACTTCAGGATCTCCAATCCTAGACAAGTGCGGGAGAGTGATAGGACTTTATGGCAATGGGGTCGTGATCAAAAATGGGAGTTATGTTAGTGCCATCACCCAAGGGAGGAGGGAGGAAGAGACTCCTGTTGAGTGCTTCGAGCCTTCGATGCTGAAGAAGAAGCAGCTAACTGTCTTAGACTTGCATCCTGGAGCTGGGAAAACCAGGAGAGTTCTTCCTGAAATAGTCCGTGAAGCCATAAAAACAAGACTCCGTACTGTGATCTTAGCTCCAACCAGGGTTGTCGCTGCTGAAATGGAGGAAGCCCTTAGAGGGCTTCCAGTGCGTTATATGACAACAGCAGTCAATGTCACCCACTCTGGAACAGAAATCGTCGACTTAATGTGCCATGCCACCTTCACTTCACGTCTACTACAGCCAATCAGAGTCCCCAATTATAATCTGTATATTATGGATGAGGCCCACTTCACAGATCCCTCAAGTATAGCAGCAAGAGGATACATTTCAACAAGGGTTGAGATGGGCGAGGCGGCTGCCATCTTCATGACCGCCACGCCACCAGGAACCCGTGACGCATTTCCGGACTCCAACTCACCAATTACGGACACCGAAGTGGAAGTCCCAGAGAGAGCCTGGAGCTCAGGCTTTGATTGGGTGACGGATCATTCTGGAAAAACAGTTTGGTTTGTTCCAAGCGTGAGGAACGGCAATGAGATCGCAGCTTGTCTGACAAAGGCTGGAAAACGGGTCATACAGCTCAGCAGAAAGACTTTTGAGACAGAGTTCCAGAAAACAAAACATCAAGAGTGGGACTTTGTCGTGACAACTGACATTTCAGAGATGGGCGCCAACTTTAAAGCTGACCGTGTCATAGATTCCAGGAGATGCCTAAAGCCGGTCATACTTGATGGCGAGAGAGTCATTCTGGCTGGACCCATGCCTGTCACACATGCCAGCGCTGCCCAGAGGAGGGGGCGCATAGGCAGGAATCCCAACAAACCTGGAGATGAGTATCTGTATGGAGGTGGGTGCGCAGAGACTGACGAAGACCATGCACACTGGCTTGAAGCAAGAATGCTCCTTGACAATATTTACCTCCAAGATGGCCTCATAGCCTCGCTCTATCGACCTGAGGCCGACAAAGTAGCAGCCATTGAGGGAGAGTTCAAGCTTAGGACGGAGCAAAGGAAGACCTTTGTGGAACTCATGAAAAGAGGAGATCTTCCTGTTTGGCTGGCCTATCAGGTTGCATCTGCCGGAATAACCTACACAGATAGAAGATGGTGCTTTGATGGCACGACCAACAACACCATAATGGAAGACAGTGTGCCGGCAGAGGTGTGGACCAGACACGGAGAGAAAAGAGTGCTCAAACCGAGGTGGATGGACGCCAGAGTTTGTTCAGATCATGCGGCCCTGAAGTCATTCAAGGAGTTTGCCGCTGGGAAAAGAGGAGCGGCTTTTGGAGTGATGGAAGCCCTGGGAACACTGCCAGGACACATGACAGAGAGATTCCAGGAAGCCATTGACAACCTCGCTGTGCTCATGCGGGCAGAGACTGGAAGCAGGCCTTACAAAGCCGCGGCGGCCCAATTGCCGGAGACCCTAGAGACCATTATGCTTTTGGGGTTGCTGGGAACAGTCTCGCTGGGAATCTTTTTCGTCTTGATGAGGAACAAGGGCATAGGGAAGATGGGCTTTGGAATGGTGACTCTTGGGGCCAGCGCATGGCTCATGTGGCTCTCGGAAATTGAGCCAGCCAGAATTGCATGTGTCCTCATTGTTGTGTTCCTATTGCTGGTGGTGCTCATACCTGAGCCAGAAAAGCAAAGATCTCCCCAGGACAACCAAATGGCAATCATCATCATGGTAGCAGTAGGTCTTCTGGGCTTGATTACCGCCAATGAACTCGGATGGTTGGAGAGAACAAAGAGTGACCTAAGCCATCTAATGGGAAGGAGAGAGGAGGGGGCAACCATAGGATTCTCAATGGACATTGACCTGCGGCCAGCCTCAGCTTGGGCCATCTATGCTGCCTTGACAACTTTCATTACCCCAGCCGTCCAACATGCAGTGACCACTTCATACAACAACTACTCCTTAATGGCGATGGCCACGCAAGCTGGAGTGTTGTTTGGTATGGGCAAAGGGATGCCATTCTACGCATGGG-ACTTTGGAGTCCCGCTGCTAATGATAGGTTGCTACTCACAATTAACACCCCTGACCCTAATAGTGGCCATCATTTTGCTCGTGGCGCACTACATGTACTTGATCCCAGGGCTGCAGGCAGCAGCTGCGCGTGCTGCCCAGAAGAGAACGGCAGCTGGCATCATGAAGAACCCTGTTGTGGATGGAATAGTGGTGACTGACATTGACACAATGACAATTGACCCCCAAGTGGAGAAAAAGATGGGACAGGTGCTACTCATAGCAGTAGCCGTCTCCAGCGCCATACTGTCGCGGACCGCCTGGGGGTGGGGGGAGGCTGGGGCCCTGATCACAGCCGCAACTTCCACTTTGTGGGAAGGCTCTCCGAACAAGTACTGGAACTCCTCTACAGCCACTTCACTGTGTAACATTTTTAGGGGAAGTTACTTGGCTGGAGCTTCTCTGATCTACACAGTAACAAGAAACGCTGGCTTGGTCAAGAGACGTGGGGGTGGAACAGGAGAGACCCTGGGAGAGAAATGGAAGGCCCGCTTGAACCAGATGTCGGCTCTGGAGTTCTACTCCTACAAAAAGTCAGGCATCACCGAGGTGTGCAGAGAAGAGGCCCGCCGCGCCCTCAAGGACGGTGTGGCAACGGGAGGCCATGCTGTGTCCCGAGGAAGTGCAAAGCTGAGATGGTTGGTGGAGCGGGGATACCTGCAGCCCTATGGAAAGGTCATTGATCTTGGATGTGGCAGAGGGGGCTGGAGTTACTACGCCGCCACCATCCGCAAAGTTCAAGAAGTGAAAGGATACACAAAAGGAGGCCCTGGTCATGAAGAACCCGTGTTGGTGCAAAGCTATGGGTGGAACATAGTCCGTCTTAAGAGTGGGGTGGACGTCTTTCATATGGCGGCTGAGCCGTGTGACACGTTGCTGTGTGACATAGGTGAGTCATCATCTAGTCCTGAAGTGGAAGAAGCACGGACGCTCAGAGTCCTCTCCATGGTGGGGGATTGGCTTGAAAAAAGACCAGGAGCCTTTTGTATAAAAGTGTTGTGCCCATACACCAGCACTATGATGGAAACCCTGGAGCGACTGCAGCGTAGGTATGGGGGAGGACTGGTCAGAGTGCCACTCTCCCGCAACTCCACACATGAGATGTACTGGGTCTCTGGAGCGAAAAGCAACACCATAAAAAGTGTGTCCACCACGAGCCAGCTCCTCTTGGGGCGCATGGACGGGCCTAGGAGGCCAGTGAAATATGAGGAGGATGTGAATCTCGGCTCTGGCACGCGGGCTGTGGTAAGCTGCGCTGAAGCTCCCAACATGAAGATCATTGGTAACCGCATTGAAAGGATCCGCAGTGAGCACGCGGAAACGTGGTTCTTTGACGAGAACCACCCATATAGGACATGGGCTTACCATGGAAGCTATGAGGCCCCCACACAAGGGTCAGCGTCCTCTCTAATAAACGGGGTTGTCAGGCTCCTGTCAAAACCCTGGGATGTGGTGACTGGAGTCACAGGAATAGCCATGACCGACACCACACCGTATGGTCAGCAAAGAGTTTTCAAGGAAAAAGTGGACACTAGGGTGCCAGACCCCCAAGAAGGCACTCGTCAGGTTATGAGCATGGTCTCTTCCTGGTTGTGGAAAGAGCTAGGCAAACACAAACGGCCACGAGTCTGTACCAAAGAAGAGTTCATCAACAAGGTTCGTAGCAATGCAGCATTAGGGGCAATATTTGAAGAGGAAAAAGAGTGGAAGACTGCAGTGGAAGCTGTGAACGATCCAAGGTTCTGGGCTCTAGTGGACAAGGAAAGAGAGCACCACCTGAGAGGAGAGTGCCAGAGTTGTGTGTACAACATGATGGGGAAAAGAGAAAAGAAACAAGGGGAATTTGGAAAGGCCAAGGGCAGCCGCGCCATCTGGTATATGTGGCTAGGGGCTAGATTTCTAGAGTTCGAAGCCCTTGGATTCTTGAACGAGGATCACTGGATGGGGAGAGAGAACTCAGGAGGTGGTGTTGAAGGGCTGGGATTACAAAGACTCGGATATGTCCTAGAAGAGATGAGTCGCATACCAGGAGGAAGGATGTATGCAGATGACACTGCTGGCTGGGACACCCGCATCAGCAGGTTTGATCTGGAGAATGAAGCTCTAATCACCAACCAAATGGAGAAAGGGCACAGGGCCTTGGCATTGGCTATAATCAAGTACACATACCAAAACAAAGTGGTAAAGGTCCTTAGACCAGCTGAAAAAGGGAAAACAGTTATGGACATTATTTCGAGACAAGACCAAAGGGGGAGCGGACAAGTTGTCACTTACGCTCTTAACACATTTACCAACCTAGTGGTGCAACTCATTCGGAATATGGAGGCTGAGGAAGTTCTAGAGATGCAAGACTTGTGGCTGCTGCGGAGGTCAGAGAAAGTGACCAACTGGTTGCAGAGCAACGGATGGGATAGGCTCAAACGAATGGCAGTCAGTGGGGATGATTGCGTTGTGAAGCCAATTGATGATAGGTTTGCACATGCCCTTAGGTTCTTGAATGATATGGGAAAAGTTAGGAAGGACACACAAGAGTGGAAACCCTCAACTGGATGGGACAACTGGGAAGAAGTTCCGTTTTGCTCCCACCACTTCAACAAGCTCCATCTCAAGGACGGGAGGTCCATTGTGGTTCCCTGCCGCCACCAAGATGAACTGATTGGCCGGGCCCGCGTCTCTCCAGGGGCGGGATGGAGCATCCGGGAGACTGCTTGCCTAGCAAAATCATATGCGCAAATGTGGCAGCTCCTTTATTTCCACAGAAGGGACCTCCGACTGATGGCCAATGCCATTTGTTCATCTGTGCCAGTTGACTGGGTTCCAACTGGGAGAACTACCTGGTCAATCCATGGAAAGGGAGAATGGATGACCACTGAAGACATGCTTGTGGTGTGGAACAGAGTGTGGATTGAGGAGAACGACCACATGGAAGACAAGACCCCAGTTACGAAATGGACAGACATTCCCTATTTGGGAAAAAGGGAAGACTTGTGGTGTGGATCTCTCATAGGGCACAGACCGCGCACCACCTGGGCTGAGAACATTAAAAACACAGTCAACATGGTGCGCAGGATCATAGGTGATGAAGAAAAGTACATGGACTACCTATCCACCCAAGTTCGCTACTTGGGTGAAGAAGGGTCTACACCTGGAGTGCTGTAA

>ENA|AMN14619|AMN14619.1 Zika virus polyprotein

ATGAAAAACCC-AAAAAAGAAATCCGGAGGATTCCGGATTGTCAATATGCTAAAACGCGGAGTAGCCCGTGTGAGCCCCTTTGGGGGCTTGAAGAGGCTGCCAGCCGGACTTCTGCTGGGTCATGGGCCCATCAGGATGGTCTTGGCGATTCTAGCCTTTTTGAGATTCACGGCAATCAAGCCATCACTGGGTCTCATCAATAGATGGGGTTCAGTGGGGAAAAAAGAGGCTATGGAAATAATAAAGAAGTTCAAGAAAGATCTGGCTGCCATGCTGAGAATAATCAATGCTAGGAAGGAGAAGAAGAGACGAGGCGCAGATACTAGTGTCGGAATTGTTGGCCTCCTGCTGACCACAGCTATGGCAGCGGAGGTCACTAGACGTGGGAGTGCATACTACATGTACTTGGACAGAAACGATGCTGGGGAGGCCATATCTTTTCCAACCACATTGGGGATGAATAAGTGTTATATACAGATCATGGATCTTGGACACATGTGTGATGCCACCATGAGCTATGAATGCCCTATGCTGGATGAGGGGGTGGAACCAGATGACGTCGATTGTTGGTGCAACACGACGTCAACTTGGGTTGTGTACGGAACCTGCCATCACAAAAAAGGTGAAGCACGGAGATCTAGAAGAGCTGTGACGCTCCCCTCCCATTCCACTAGGAAGCTGCAAACGCGGTCGCAAACCTGGTTGGAATCAAGAGAATACACAAAGCACTTGATTAGAGTCGAAAATTGGATATTCAGGAACCCTGGCTTCGCGTTAGCAGCAGCTGCCATCGCTTGGCTTTTGGGAAGCTCAACGAGCCAAAAAGTCATATACTTGGTCATGATACTGCTGATTGCCCCGGCATACAGCATCAGGTGCATAGGAGTCAGCAATAGGGACTTTGTGGAAGGTATGTCAGGTGGGACTTGGGTTGATGTTGTCTTGGAACATGGAGGTTGTGTCACCGTAATGGCACAGGACAAACCGACTGTCGACATAGAGCTGGTTACAACAACAGTCAGCAACATGGCGGAGGTAAGATCCTACTGCTATGAGGCATCAATATCAGACATGGCTTCGGACAGCCGCTGCCCAACACAAGGTGAAGCCTACCTTGACAAGCAATCAGACACTCAATATGTCTGCAAAAGAACGTTAGTGGACAGAGGCTGGGGAAATGGATGTGGACTTTTTGGCAAAGGGAGCCTGGTGACATGCGCTAAGTTTGCATGCTCCAAGAAAATGACCGGGAAGAGCATCCAGCCAGAGAATCTGGAGTACCGGATAATGCTGTCAGTTCATGGCTCCCAGCACAGTGGGATGATCGTTAATGACACTGGACATGAAACTGATGAGAATAGAGCGAAGGTTGAGATAACGCCCAATTCACCAAGAGCCGAAGCCACCCTGGGGGGTTTTGGAAGCCTAGGACTTGATTGTGAACCGAGGACAGGCCTTGACTTTTCAGATTTGTATTACTTGACTATGAATAACAAGCACTGGTTGGTTCACAAGGAGTGGTTCCACGACATTCCATTACCTTGGCACGCTGGGGCAGACACCGGAACTCCACACTGGAACAACAAAGAAGCACTGGTAGAGTTCAAGGACGCACATGCCAAAAGGCAAACTGTCGTGGTTCTAGGGAGTCAAGAAGGAGCAGTTCACACGGCCCTTGCTGGAGCTCTGGAGGCTGAGATGGATGGTGCAAAGGGAAGGCTGTCCTCTGGCCACTTGAAATGTCGCCTGAAAATGGATAAACTTAGATTGAAGGGCGTGTCATACTCCTTGTGTACCGCAGCGTTCACATTCACCAAGATCCCGGCTGAAACACTGCACGGGACAGTCACAGTGGAGGTACAGTACGCAGGGACAGATGGACCTTGCAAGGTTCCAGCTCAGATGGCGGTGGACATGCAAACTCTGACCCCAGTTGGGAGGTTGATAACCGCCAACCCCGTAATCACTGAAAGCACTGAGAACTCTAAGATGATGCTGGAACTTGATCCACCATTTGGGGACTCTTACATTGTCATAGGAGTCGGGGAGAAGAAGATCACCCACCACTGGCACAGGAGTGGCAGCACCATTGGAAAAGCATTTGAAGCCACTGTGAGAGGTGCCAAGAGAATGGCAGTCTTGGGAGACACAGCCTGGGACTTTGGATCAGTTGGAGGCGCTCTCAACTCATTGGGCAAGGGCATCCATCAAATTTTTGGAGCAGCTTTCAAATCATTGTTTGGAGGAATGTCCTGGTTCTCACAAATCCTCATTGGAACGTTGCTGATGTGGTTGGGTCTGAACACAAAGAATGGATCTATTTCCCTCATGTGCTTGGCCTTAGGGGGAGTGTTGATCTTCTTATCCACAGCCGTCTCTGCTGATGTGGGGTGCTCGGTGGACTTCTCAAAGAAGGAGACGAGATGCGGTACAGGGGTGTTCGTCTATAACGACGTTGAAGCCTGGAGGGACAGGTACAAGTACCATCCTGACTCCCCCCGTAGATTGGCAGCAGCAGTCAAGCAAGCCTGGGAAGATGGTATCTGCGGGATCTCCTCTGTTTCAAGAATGGAAAACATCATGTGGAGATCAGTAGAAGGGGAGCTCAATGCAATCCTGGAAGAGAATGGAGTTCAACTGACGGTCGTTGTGGGATCTGTAAAAAACCCCATGTGGAGAGGTCCACAGAGATTGCCCGTGCCTGTGAACGAGCTGCCCCACGGCTGGAAGGCTTGGGGGAAATCGTACTTCGTTAGAGCAGCAAAGACAAATAACAGCTTTGTCGTGGATGGTGACACACTGAAGGAATGCCCACTCAAACATAGAGCATGGAACAGCTTTCTTGTGGAGGATCATGGGTTCGGGGTATTTCACACTAGTGTCTGGCTCAAGGTTAGAGAAGATTATTCATTAGAGTGTGATCCAGCCGTTATTGGAACAGCTGTTAAGGGAAAGGAGGCTGTACACAGTGATCTAGGCTACTGGATTGAGAGTGAGAAGAATGACACATGGAGGCTGAAGAGGGCCCATCTGATCGAGATGAAAACATGTGAATGGCCAAAGTCCCACACATTGTGGACAGATGGAATAGAAGAGAGTGATCTGATCATACCCAAGTCTTTAGCTGGGCCACTCAGCCATCACAATACCAGAGAGGGCTACAGGACCCAAATGAAAGGGCCATGGCACAGTGAAGAGCTTGAAATTCGGTTTGAGGAATGCCCAGGCACTAAGGTCCACGTGGAGGAAACATGTGGAACAAGAGGACCATCTCTGAGATCAACCACTGCAAGCGGAAGGGTGATCGAGGAATGGTGCTGCAGGGAGTGCACAATGCCCCCACTGTCGTTCCGGGCTAAAGATGGCTGTTGGTATGGAATGGAGATAAGGCCCAGGAAAGAACCAGAAAGCAACTTAGTAAGGTCAGTGGTGACTGCAGGATCAACTGATCACATGGATCACTTCTCCCTTGGAGTGCTTGTGATTCTGCTCATGGTGCAGGAAGGGCTGAAGAAGAGAATGACCACAAAGATCATCATAAGCACATCRATGGCAGTGCTGGTAGCTATGATCCTGGGAGGATTTTCAATGAGTGACCTGGCTAAGCTTGCAATTTTGATGGGTGCCACCTTCGCGGAAATGAACACTGGAGGAGATGTAGCTCATCTGGCGCTGATAGCGGCATTCAAAGTCAGACCAGCGTTGCTGGTATCTTTCATCTTCAGAGCTAATTGGACACCCCGTGAAAGCATGCTGCTGGCCTTGGCCTCGTGTCTTTTGCAAACTGCGATCTCCGCCTTGGAAGGCGACCTGATGGTTCTCATCAATGGTTTTGCTTTGGCCTGGTTGGCAATACGAGCGATGGTTGTTCCACGCACTGATAACATCACCTTGGCAATCCTGGCTGCTCTGACACCACTGGCCCGGGGCACACTGCTTGTGGCGTGGAGAGCAGGCCTTGCTACTTGCGGGGGGTTTATGCTCCTCTCTCTGAAGGGAAAAGGCAGTGTGAAGAAGAACTTACCATTTGTCATGGCCCTGGGACTAACCGCTGTGAGGCTGGTCGACCCCATCAACGTGGTGGGACTGCTGTTGCTCACAAGGAGTGGGAAGCGGAGCTGGCCCCCTAGCGAAGTACTCACAGCTGTTGGCCTGATATGCGCATTGGCTGGAGGGTTCGCCAAGGCAGATATAGAGATGGCTGGGCCCATGGCCGCAGTCGGTCTGCTAATTGTCAGTTACGTGGTCTCAGGAAAGAGTGTGGACATGTACATTGAAAGAGCAGGTGACATCACATGGGAAAAAGATGCGGAAGTCACTGGAAACAGTCCCCGGCTCGATGTGGCGCTAGATGAGAGTGGTGATTTCTCCCTGGTGGAGGATGACGGTCCCCCCATGAGAGAGATCATACTCAAGGTGGTCCTGATGACCATCTGTGGCATGAACCCAATAGCCATACCCTTTGCAGCTGGAGCGTGGTACGTATACGTGAAGACTGGAAAAAGGAGTGGTGCTCTATGGGATGTGCCTGCTCCCAAGGAAGTAAAAAAGGGGGAGACCACAGATGGAGTGTACAGAGTAATGACTCGTAGACTGCTAGGTTCAACACAAGTTGGAGTGGGAGTTATGCAAGAGGGGGTCTTTCACACTATGTGGCACGTCACAAAAGGATCCGCGCTGAGAAGCGGTGAAGGGAGACTTGATCCATACTGGGGAGATGTCAAGCAGGATCTGGTGTCATACTGTGGTCCATGGAAGCTAGATGCCGCCTGGGACGGGCACAGCGAGGTGCAGCTCTTGGCCGTGCCCCCCGGAGAGAGAGCGAGGAACATCCAGACTCTGCCCGGAATATTTAAGACAAAGGATGGGGACATTGGAGCGGTTGCGCTGGATTACCCAGCAGGAACTTCAGGATCTCCAATCCTAGACAAGTGTGGGAGAGTGATAGGACTTTATGGCAATGGGGTCGTGATCAAAAATGGGAGTTATGTTAGTGCCATCACCCAAGGGAGGAGGGAGGAAGAGACTCCTGTTGAGTGCTTCGAGCCTTCGATGCTGAAGAAGAAGCAGCTAACTGTCTTAGACTTGCATCCTGGAGCTGGGAAAACCAGGAGAGTTCTTCCTGAAATAGTCCGTGAAGCTATAAAAACAAGACTCCGTACTGTGATCTTAGCTCCAACCAGGGTTGTCGCTGCTGAAATGGAGGAAGCCCTTAGAGGGCTTCCAGTGCGTTATATGACAACAGCAGTCAATGTCACCCATTCTGGAACAGAAATCGTCGACTTAATGTGCCATGCCACCTTCACTTCACGTCTACTACAGCCAATCAGAGTCCCCAACTATAATCTGTACATTATGGATGAGGCCCACTTCACAGATCCCTCAAGTATAGCAGCAAGAGGATACATTTCAACAAGGGTTGAGATGGGCGAGGCGGCTGCCATCTTCATGACCGCCACGCCACCAGGAACCCGTGACGCATTTCCGGACTCCAACTCACCAATTATGGACACCGAAGTGGAAGTCCCAGAGAGAGCCTGGAGCTCAGGCTTTGATTGGGTGACGGATCATTCTGGAAAAACAGTTTGGTTTGTTCCAAGCGTGAGGAACGGCAATGAGATCGCAGCTTGTCTGACAAAGGCTGGAAAACGGGTCATACAGCTCAGCAGAAAGACTTTTGAGACAGAGTTCCAGAAAACAAAACATCAAGAGTGGGACTTTGTCGTGACAACCGACATTTCAGAGATGGGCGCCAACTTTAAAGCTGACCGTGTCATAGATTCCAGGAGATGCCTAAAGCCGGTCATACTTGATGGCGAGAGAGTCATTCTGGCTGGACCCATGCCTGTCACACATGCCAGCGCTGCCCAGAGGAGGGGGCGCATAGGCAGGAATCCCAACAAACCTGGAGATGAGTATCTGTATGGAGGTGGGTGCGCAGAGACTGACGAAGACCATGCACACTGGCTTGAAGCAAGAATGCTCCTTGACAATATTTACCTCCAAGATGGCCTCATAGCCTCGCTCTATCGACCTGAGGCCGACAAAGTAGCAGCCATTGAGGGAGAGTTCAAGCTTAGGACGGAGCAAAGGAAGACCTTTGTGGAACTCATGAAAAGAGGAGATCTTCCTGTTTGGCTGGCCTATCAGGTTGCATCTGCCGGAATAACTTACACAGATAGAAGATGGTGCTTTGATGGCACGACCAACAACACCATAATGGAAGACAGTGTGCCGGCAGAGGTGTGGACCAGACACGGAGAGAAAAGAGTGCTCAAACCGAGGTGGATGGACGCCAGAGTTTGTTCAGATCATGCGGCCCTGAAGTCATTCAAGGAGTTTGCCGCTGGGAAAAGAGGAGCGGCTTTTGGAGTGATGGAAGCCCTGGGAACACTGCCAGGACACATGACAGAGAGATTCCAGGAAGCCATTGACAACCTCGCTGTGCTCATGCGGGCAGAGACTGGAAGCAGGCCTTACAAAGCCGCGGCGGCCCAATTGCCGGAGACCCTAGAGACCATTATGCTTTTGGGGTTGCTGGGAACAGTCTCGCTGGGAATCTTTTTCGTCTTGATGAGGAACAAGGGCATAGGGAAGATGGGCTTTGGAATGGTGACTCTTGGGGCCAGCGCATGGCTCATGTGGCTCTCGGAAATTGAGCCAGCCAGAATTGCATGTGTCCTCATTGTTGTGTTCCTACTGCTGGTGGTGCTCATACCTGAGCCAGAAAAGCAAAGATCTCCCCAGGACAACCAAATGGCAATCATCATCATGGTAGCAGTAGGTCTTCTGGGCTTGATCACCGCCAATGAACTCGGATGGTTGGAGAGAACAAAGAGTGACCTAAGCCATCTAATGGGAAGGAGAGAGGAGGGAGCAACCATAGGATTCTCAATGGACATTGACCTGCGGCCAGCCTCAGCTTGGGCCATCTATGCTGCCTTGACAACTTTCATTACCCCAGCCGTCCAACATGCAGTGACCACTTCATACAACAACTACTCCTTAATGGCGATGGCCACGCAAGCTGGAGTGTTGTTTGGTATGGGCAAAGGGATGCCATTCTACGCATGGG-ACTTTGGAGTCCCGCTGCTAATGATAGGTTGCTACTCACAATTAACACCCCTGACCCTAATAGTGGCCATCATTTTGCTCGTGGCGCACTACATGTACTTGATCCCAGGGCTGCAGGCAGCAGCTGCGCGTGCTGCCCAGAAGAGAACGGCAGCTGGCATCATGAAGAACCCTGTTGTGGATGGAATAGTGGTGACTGACATTGACACAATGACAATTGACCCCCAAGTGGAGAAAAAGATGGGACAGGTGCTACTCATAGCAGTAGCCGTCTCCAGCGCCATACTGTCGCGGACCGCCTGGGGGTGGGGGGAGGCTGGGGCCCTGATCACAGCCGCAACTTCCACTTTGTGGGAAGGCTCTCCGAACAAGTACTGGAACTCCTCTACAGCCACTTCACTGTGTAACATTTTTAGGGGAAGTTACTTGGCTGGAGCTTCTCTAATCTACACAGTAACAAGAAACGCTGGCTTGGTCAAGAGACGTGGGGGTGGAACAGGAGAGACCCTGGGAGAGAAATGGAAGGCCCGCTTGAACCAGATGTCGGCCCTGGAGTTCTACTCCTACAAAAAGTCAGGCATCACCGAGGTGTGCAGAGAAGAGGCCCGCCGCGCCCTCAAGGACGGTGTGGCAACGGGAGGCCATGCTGTGTCCCGAGGAAGTGCAAAGCTGAGATGGTTGGTGGAGCGGGGATACCTGCAGCCCTATGGAAAGGTCATTGATCTTGGATGTGGCAGAGGGGGCTGGAGTTACTACGCCGCCACCATCCGCAAAGTTCAAGAAGTGAAAGGATACACAAAAGGAGGCCCTGGTCATGAAGAACCCGTGTTGGTGCAAAGCTATGGGTGGAACATAGTCCGTCTCAAGAGTGGGGTGGACGTCTTTCATATGGCGGCTGAGCCGTGTGACACGTTGCTGTGTGACATAGGTGAGTCATCATCTAGTCCTGAAGTGGAAGAAGCACGGACGCTCAGAGTCCTCTCCATGGTGGGGGATTGGCTTGAAAAAAGACCAGGAGCCTTTTGTATAAAAGTGTTGTGCCCATACACCAGCACTATGATGGAAACCCTGGAGCGACTGCAGCGTAGGTATGGGGGAGGACTGGTCAGAGTGCCACTCTCCCGCAACTCTACACATGAGATGTACTGGGTCTCTGGAGCGAAAAGCAACACCATAAAAAGTGTGTCCACCACGAGCCAGCTCCTCTTGGGGCGCATGGACGGGCCTAGGAGGCCAGTGAAATATGAGGAGGATGTGAATCTCGGCTCTGGCACGCGGGCTGTGGTAAGCTGCGCTGAAGCTCCCAACATGAAGATCATTGGTAACCGCATTGAAAGGATCCGCAGTGAGCACGCGGAAACGTGGTTCTTTGACGAGAACCACCCATATAGGACATGGGCTTACCATGGAAGCTATGAGGCCCCCACACAAGGGTCAGCATCCTCTCTAGTAAACGGGGTTGTCAGGCTCCTGTCAAAACCCTGGGATGTGGTGACTGGAGTCACAGGAATAGCCATGACCGACACCACACCGTATGGTCAGCAAAGAGTTTTCAAGGAAAAAGTGGACACTAGGGTGCCAGACCCCCAAGAAGGCACTCGTCAGGTTATGAGCATGGTCTCTTCCTGGTTGTGGAAAGAGCTAGGCAAACACAAACGGCCACGAGTCTGTACCAAAGAAGAGTTCATCAACAAGGTTCGTAGCAATGCAGCATTAGGGGCAATATTTGAAGAGGAAAAAGAGTGGAAGACTGCAGTGGAAGCTGTGAACGATCCAAGGTTCTGGGCTCTAGTGGACAAGGAAAGAGAGCACCACCTGAGAGGAGAGTGCCAGAGTTGTGTGTACAACATGATGGGAAAAAGAGAAAAGAAACAAGGGGAATTTGGAAAGGCCAAGGGCAGCCGCGCCATCTGGTATATGTGGCTAGGGGCTAGATTTCTAGAGTTCGAAGCCCTTGGATTCTTGAACGAGGATCACTGGATGGGGAGAGAGAACTCAGGAGGTGGTGTTGAAGGGCTGGGATTACAAAGACTCGGATATGTCCTAGAAGAGATGAGTCGCATACCAGGAGGAAGGATGTATGCAGATGACACTGCTGGCTGGGATACCCGCATCAGCAGGTTTGATCTAGAGAATGAAGCTCTAATCACCAACCAAATGGAGAAAGGGCACAGGGCCTTGGCATTGGCCATAATCAAGTACACATACCAAAACAAAGTGGTAAAGGTCCTTAGACCAGCTGAAAAAGGGAAAACAGTTATGGACATTATTTCGAGACAAGACCAAAGGGGGAGCGGACAAGTTGTCACTTACGCTCTTAACACATTTACCAACCTAGTGGTGCAACTCATTCGGAATATGGAGGCTGAGGAAGTTCTAGAGATGCAAGACTTGTGGCTGCTGCGGAGGTCAGAGAAAGTGACCAACTGGTTGCAGAGCAACGGATGGGATAGGCTCAAACGAATGGCAGTCAGTGGAGATGATTGCGTTGTGAAGCCAATTGATGATAGGTTTGCACATGCCCTCAGGTTCTTGAATGATATGGGAAAAGTTAGGAAGGACACACAAGAGTGGAAACCCTCAACTGGATGGGACAACTGGGAAGAAGTTCCGTTTTGCTCCCACCACTTCAACAAGCTCCATCTCAAGGACGGGAGGTCCATTGTGGTTCCCTGCCGCCACCAAGATGAACTGATTGGCCGGGCCCGCGTCTCTCCAGGGGCGGGATGGAGCATCCGGGAGACTGCTTGCCTAGCAAAATCATATGCGCAAATGTGGCAGCTCCTTTATTTCCACAGAAGGGACCTCCGACTGATGGCCAATGCCATTTGTTCATCTGTGCCAGTTGACTGGGTTCCAACTGGGAGAACTACCTGGTCAATCCATGGAAAGGGAGAATGGATGACCACTGAAGACATGCTTGTGGTGTGGAACAGAGTGTGGATTGAGGAGAACGACCACATGGAAGACAAGACCCCAGTTACGAAATGGACAGACATTCCCTATTTGGGGAAAAGGGAAGACTTGTGGTGTGGATCTCTCATAGGGCACAGACCGCGCACCACCTGGGCTGAGAACATTAAAAACACAGTCAACATGGTGCGCAGGATCATAGGTGAGGAAGAAAAGTACATGGACTACCTATCCACCCAAGTTCGCTACTTGGGTGAAGAAGGGTCTACACCTGGARTGCTGTAA

>ENA|AMK02027|AMK02027.1 Zika virus polyprotein

ATGAAAAACCC-AAAGAAGAAATCCGGAGGATTCCGGATTGTCAATATGCTAAAACGCGGAGTAGCCCGTGTAAACCCCTTGGGAGGTTTGAAGAGGTTGCCAGCCGGACTTCTGCTGGGTCATGGACCCATCAGAATGGTTTTGGCGATACTAGCCTTTTTGAGATTTACAGCAATCAAGCCATCACTGGGCCTTATCAACAGATGGGGTTCCGTGGGGAAAAAAGAGGCTATGGAAATAATAAAGAAGTTCAAGAAAGATCTTGCTGCCATGTTGAGAATAATCAATGCTAGGAAAGAGAGGAAGAGACGTGGCGCAGACACCAGCATCGGAATCATTGGCCTCCTGCTGACTACAGCCATGGCAGCAGAGATCACTAGACGCGGGAGTGCATACTACATGTACTTGGATAGGAGCGATGCCGGGAAGGCCATTTCGTTTGCTACCACATTGGGAGTGAACAAGTGCCACGTACAGATCATGGACCTCGGGCACATGTGTGACGCCACCATGAGTTATGAGTGCCCTATGCTGGATGAGGGAGTGGAACCAGATGATGTCGATTGCTGGTGCAACACGACATCAACTTGGGTTGTGTACGGAACCTGTCATCACAAAAAAGGTGAGGCACGGCGATCTAGAAGAGCCGTGACGCTCCCTTCTCACTCTACAAGGAAGTTGCAAACGCGGTCGCAGACCTGGTTAGAATCAAGAGAATACACGAAGCACTTGATCAAGGTTGAAAACTGGATATTCAGGAACCCCGGGTTTGCGCTAGTGGCCGTTGCCATTGCCTGGCTTTTGGGAAGCTCGACGAGCCAAAAAGTCATATACTTGGTCATGATACTGCTGATTGCCCCGGCATACAGTATCAGGTGCATTGGAGTCAGCAATAGAGACTTCGTGGAGGGCATGTCAGGTGGGACCTGGGTTGATGTTGTCTTGGAACATGGAGGCTGCGTTACCGTGATGGCACAGGACAAGCCAACAGTTGACATAGAGTTGGTCACGACGACGGTTAGTAACATGGCCGAGGTAAGATCCTATTGCTACGAGGCATCGATATCGGACATGGCTTCGGACAGTCGTTGCCCAACACAAGGTGAAGCCTACCTTGACAAGCAATCAGACACTCAATATGTCTGCAAAAGAACATTAGTGGACAGAGGTTGGGGAAACGGTTGTGGACTTTTTGGCAAAGGGAGCTTGGTGACATGTGCCAAGTTTACGTGTTCTAAGAAGATGACCGGGAAGAGCATTCAACCGGAAAATCTGGAGTATCGGATAATGCTATCAGTGCATGGCTCCCAGCATAGCGGGATGATTGTCAATGATACAGGATATGAAACTGACGAAAATAGAGCGAAAGTCGAGGTTACGCCTAATTCACCAAGAGCGGAAGCAACCTTGGGAGGCTTTGGAAGCTTAGGACTTGACTGTGAACCAAGGACAGGCCTTGACTTTTCAGATCTGTATTACCTGACCATGAACAATAAGCATTGGTTGGTGCACAAAGAGTGGTTTCATGACATCCCATTGCCTTGGCATGCTGGGGCAGACACCGGAACTCCACACTGGAACAACAAAGAGGCATTGGTAGAATTCAAGGATGCCCACGCCAAGAGGCAAACCGTCGTCGTTCTGGGGAGCCAGGAAGGAGCCGTTCACACGGCTCTCGCTGGAGCTCTAGAGGCTGAGATGGATGGTGCAAAGGGAAAGCTGTTCTCTGGCCATTTGAAATGCCGCCTAAAAATGGACAAGCTTAGATTGAAGGGCGTGTCATATTCCTTGTGCACTGCGGCATTCACATTCACCAAGGTCCCAGCTGAAACACTGCATGGAACAGTCACAGTGGAGGTGCAGTATGCAGGGACAGATGGACCCTGCAAGATCCCAGTCCAGATGGCGGTGGACATGCAGACCCTGACCCCAGTTGGAAGGCTGATAACCGCCAACCCCGTGATTACTGAAAGCACTGAGAACTCAAAGATGATGTTGGAGCTTGACCCACCATTTGGGGATTCTTACATTGTCATAGGAGTTGGGGACAAGAAAATCACCCACCACTGGCATAGGAGTGGTAGCACCATCGGAAAGGCATTTGAGGCCACTGTGAGAGGCGCCAAGAGAATGGCAGTCCTGGGGGATACAGCCTGGGACTTCGGATCAGTCGGGGGTGTGTTCAACTCACTGGGTAAGGGCATTCACCAGATTTTTGGAGCAGCCTTCAAATCACTGTTTGGAGGAATGTCCTGGTTCTCACAGATCCTCATAGGCACGCTGCTAGTGTGGTTAGGTTTGAACACAAAGAATGGATCTATCTCCCTCACATGCTTGGCCCTGGGGGGAGTGATGATCTTCCTCTCCACGGCTGTTTCTGCTGACGTGGGGTGCTCAGTGGACTTCTCAAAAAAGGAAACGAGATGTGGCACGGGGGTATTCATCTATAATGATGTTGAAGCCTGGAGGGACCGGTACAAGTACCATCCTGACTCCCCCCGCAGATTGGCAGCAGCAGTCAAGCAGGCCTGGGAAGAGGGGATCTGTGGGATCTCATCCGTTTCAAGAATGGAAAACATCATGTGGAAATCAGTAGAAGGGGAGCTCAATGCTATCCTAGAGGAGAATGGAGTTCAACTGACAGTTGTTGTGGGATCTGTAAAAAACCCCATGTGGAGAGGTCCACAAAGATTGCCAGTGCCTGTGAATGAGCTGCCCCATGGCTGGAAAGCCTGGGGGAAATCGTATTTTGTTAGGGCGGCAAAGACCAACAACAGTTTTGTTGTCGACGGTGACACACTGAAGGAATGTCCGCTTGAGCACAGAGCATGGAATAGTTTTCTTGTGGAGGATCACGGGTTTGGAGTCTTCCACACCAGTGTCTGGCTTAAGGTCAGAGAAGATTACTCATTAGAATGTGACCCAGCCGTCATAGGAACAGCTGTTAAGGGAAGGGAGGCCGCGCACAGTGATCTGGGCTATTGGATTGAAAGTGAAAAGAATGACACATGGAGGCTGAAGAGGGCCCACCTGATTGAGATGAAAACATGTGAATGGCCAAAGTCTCACACATTGTGGACAGATGGAGTAGAAGAAAGTGATCTTATCATACCCAAGTCTTTAGCTGGTCCACTCAGCCACCACAACACCAGAGAGGGTTACAGAACCCAAGTGAAAGGGCCATGGCACAGTGAAGAGCTTGAAATCCGGTTTGAGGAATGTCCAGGCACCAAGGTTTACGTGGAGGAGACATGCGGAACTAGAGGACCATCTCTGAGATCAACTACTGCAAGTGGAAGGGTCATTGAGGAATGGTGCTGTAGGGAATGCACAATGCCCCCACTATCGTTTCGAGCAAAAGACGGCTGCTGGTATGGAATGGAGATAAGGCCCAGGAAAGAACCAGAGAGCAACTTAGTGAGGTCAATGGTGACAGCGGGGTCAACCGATCATATGGACCACTTCTCTCTTGGAGTGCTTGTGATTCTACTCATGGTGCAGGAGGGGTTGAAGAAGAGAATGACCACAAAGATCATCATGAGCACATCAATGGCAGTGCTGGTAGTCATGATCTTGGGAGGATTTTCAATGAGTGACCTGGCCAAGCTTGTGATCCTGATGGGTGCTACTTTCGCAGAAATGAACACTGGAGGAGATGTAGCTCACTTGGCATTGGTAGCGGCATTTAAAGTCAGACCAGCCTTGCTGGTCTCCTTCATTTTCAGAGCCAATTGGACACCCCGTGAGAGCATGCTGCTAGCCCTGGCTTCGTGTCTTCTGCAAACTGCGATCTCTGCTCTTGAAGGTGACTTGATGGTCCTCATTAATGGATTTGCTTTGGCCTGGTTGGCAATTCGAGCAATGGCCGTGCCACGCACTGACAACATCGCTCTACCAATCTTGGCTGCTCTAACACCACTAGCTCGAGGCACACTGCTCGTGGCATGGAGAGCGGGCCTGGCTACTTGTGGAGGGATCATGCTCCTCTCCCTGAAAGGGAAAGGTAGTGTGAAGAAGAACCTGCCATTTGTCATGGCCCTGGGATTGACAGCTGTGAGGGTAGTAGACCCTATTAATGTGGTAGGACTACTGTTACTCACAAGGAGTGGGAAGCGGAGCTGGCCCCCTAGTGAAGTTCTCACAGCCGTTGGCCTGATATGTGCACTGGCCGGAGGGTTTGCCAAGGCAGACATTGAGATGGCTGGACCCATGGCTGCAGTAGGCTTGCTAATTGTCAGCTATGTGGTCTCGGGAAAGAGTGTGGACATGTACATTGAAAGAGCAGGTGACATCACATGGGAAAAGGACGCGGAAGTCACTGGAAACAGTCCTCGGCTTGACGTGGCACTGGATGAGAGTGGTGATTTCTCCTTGGTAGAGGAAGATGGTCCACCCATGAGAGAGATCATACTCAAGGTGGTCCTGATGGCCATCTGTGGCATGAACCCAATAGCTATACCTTTTGCTGCAGGAGCGTGGTATGTGTATGTGAAGACTGGGAAAAGGAGTGGCGCCCTCTGGGACGTGCCTGCTCCCAAAGAAGTGAAGAAAGGAGAGACCACAGATGGAGTGTACAGAGTGATGACTCGCAGACTGCTAGGTTCAACACAGGTTGGAGTGGGAGTCATGCAAGAGGGAGTCTTCCACACCATGTGGCACGTTACAAAAGGAGCCGCACTGAGGAGCGGTGAGGGAAGACTTGATCCATACTGGGGGGATGTCAAGCAGGACTTGGTGTCATACTGTGGGCCTTGGAAGTTGGATGCAGCTTGGGATGGACTCAGCGAGGTACAGCTTTTGGCCGTACCTCCCGGAGAGAGGGCCAGAAACATTCAGACCCTGCCTGGAATATTCAAGACAAAGGACGGGGACATCGGAGCAGTTGCTCTGGACTACCCTGCAGGGACCTCAGGATCTCCGATCCTAGACAAATGTGGAAGAGTGATAGGACTCTATGGCAATGGGGTTGTGATCAAGAATGGAAGCTATGTTAGTGCTATAACCCAGGGAAAGAGGGAGGAGGAGACTCCGGTTGAATGTTTCGAACCCTCGATGCTGAAGAAGAAGCAGCTAACTGTCTTGGATCTGCATCCAGGAGCCGGAAAAACCAGGAGAGTTCTTCCTGAAATAGTCCGTGAAGCCATAAAAAAGAGACTCCGGACAGTGATCTTGGCACCAACTAGGGTTGTCGCTGCTGAGATGGAGGAGGCCTTGAGAGGACTTCCGGTGCGTTACATGACAACAGCAGTCAACGTCACCCATTCTGGGACAGAAATCGTTGATTTGATGTGCCATGCCACTTTCACTTCACGCTTACTACAACCCATCAGAGTCCCTAATTACAATCTCTACATCATGGATGAAGCCCACTTCACAGACCCCTCAAGTATAGCTGCAAGAGGATACATATCAACAAGGGTTGAAATGGGCGAGGCGGCTGCCATTTTTATGACTGCCACACCACCAGGAACCCGTGATGCGTTTCCTGACTCTAACTCACCAATCATGGACACAGAAGTGGAAGTCCCAGAGAGAGCCTGGAGCTCAGGCTTTGATTGGGTGACAGACCATTCTGGGAAAACAGTTTGGTTCGTTCCAAGCGTGAGAAACGGAAATGAAATCGCAGCCTGTCTGACAAAGGCTGGAAAGCGGGTCATACAGCTCAGCAGGAAGACTTTTGAGACAGAATTTCAGAAAACAAAAAATCAAGAGTGGGACTTTGTCATAACAACTGACATCTCAGAGATGGGCGCCAACTTCAAGGCTGACCGGGTCATAGACTCTAGGAGATGCCTAAAACCAGTCATACTTGATGGTGAGAGAGTCATCTTGGCTGGGCCCATGCCTGTCACGCATGCTAGTGCTGCTCAGAGGAGAGGACGTATAGGCAGGAACCCTAACAAACCTGGAGATGAGTACATGTATGGAGGTGGGTGTGCAGAGACTGATGAAGGCCATGCACACTGGCTTGAAGCAAGAATGCTTCTTGACAACATCTACCTCCAGGATGGCCTCATAGCCTCGCTCTATCGGCCTGAGGCCGATAAGGTAGCCGCCATTGAGGGAGAGTTTAAGCTGAGGACAGAGCAAAGGAAGACCTTCGTGGAACTCATGAAGAGAGGAGACCTTCCCGTCTGGCTAGCCTATCAGGTTGCATCTGCCGGAATAACTTACACAGACAGAAGATGGTGCTTTGATGGCACAACCAACAACACCATAATGGAAGACAGCGTACCAGCAGAGGTGTGGACAAAGTATGGAGAGAAGAGAGTGCTCAAACCGAGATGGATGGATGCTAGGGTCTGTTCAGACCATGCGGCCCTGAAGTCGTTCAAAGAATTCGCCGCTGGAAAAAGAGGAGCGGCTTTGGGAGTAATGGAGGCCCTGGGAACACTGCCAGGACACATGACAGAGAGGTTTCAGGAAGCCATTGACAACCTCGCCGTGCTCATGCGAGCAGAGACTGGAAGCAGGCCTTATAAGGCAGCGGCAGCCCAACTGCCGGAGACCCTAGAGACCATTATGCTCTTAGGTTTGCTGGGAACAGTTTCACTGGGGATCTTCTTCGTCTTGATGCGGAATAAGGGCATCGGGAAGATGGGCTTTGGAATGGTAACCCTTGGGGCCAGTGCATGGCTCATGTGGCTTTCGGAAATTGAACCAGCCAGAATTGCATGTGTCCTCATTGTTGTGTTTTTATTACTGGTGGTGCTCATACCCGAGCCAGAGAAGCAAAGATCTCCCCAAGATAACCAGATGGCAATTATCATCATGGTGGCAGTGGGCCTTCTAGGTTTGATAACTGCAAACGAACTTGGATGGCTGGAAAGAACAAAAAATGACATAGCTCATCTAATGGGAAGGAGAGAAGAAGGAGCAACCATGGGATTCTCAATGGACATTGATCTGCGGCCAGCCTCCGCCTGGGCTATCTATGCCGCATTGACAACTCTCATCACCCCAGCTGTCCAACATGCGGTAACCACTTCATACAACAACTACTCCTTAATGGCGATGGCCACACAAGCTGGAGTGCTGTTTGGCATGGGCAAAGGGATGCCATTTTATGCATGGG-ACCTTGGAGTCCCGCTGCTAATGATGGGTTGCTATTCACAATTAACACCCCTGACTCTGATAGTAGCTATCATTCTGCTTGTGGCGCACTACATGTACTTGATCCCAGGCCTACAAGCGGCAGCAGCGCGTGCTGCCCAGAAAAGGACAGCAGCTGGCATCATGAAGAATCCCGTTGTGGATGGAATAGTGGTAACTGACATTGACACAATGACAATAGACCCCCAGGTGGAGAAGAAGATGGGACAAGTGTTACTCATAGCAGTAGCCATCTCCAGTGCTGTGCTGCTGCGGACCGCCTGGGGATGGGGGGAGGCTGGAGCTCTGATCACAGCAGCGACCTCCACCTTGTGGGAAGGCTCTCCAAACAAATACTGGAACTCCTCTACAGCCACCTCACTGTGCAACATCTTCAGAGGAAGCTATCTGGCAGGAGCTTCCCTTATCTATACAGTGACGAGAAACGCTGGCCTGGTTAAGAGACGTGGAGGTGGGACGGGAGAGACTCTGGGAGAGAAGTGGAAAGCTCGTCTGAATCAGATGTCGGCCCTGGAGTTCTACTCTTATAAAAAGTCAGGTATCACTGAAGTGTGTAGAGAGGAGGCTCGCCGTGCCCTCAAGGATGGAGTGGCCACAGGAGGACATGCCGTATCCCGGGGAAGTGCAAAGCTCAGATGGTTGGTGGAGAGAGGATATCTGCAGCCCTATGGGAAGGTTGTTGACCTCGGATGTGGCAGAGGGGGCTGGAGCTATTATGCCGCCACCATCCGCAAAGTGCAGGAGGTGAGAGGATACACAAAGGGAGGTCCCGGTCATGAAGAACCCATGCTGGTGCAAAGCTATGGGTGGAACATAGTTCGTCTCAAGAGTGGAGTGGACGTCTTCCACATGGCGGCTGAGCCGTGTGACACTCTGCTGTGTGACATAGGTGAGTCATCATCTAGTCCTGAAGTGGAAGAGACACGAACACTCAGAGTGCTCTCTATGGTGGGGGACTGGCTTGAAAAAAGACCAGGGGCCTTCTGTATAAAGGTGCTGTGCCCATACACCAGCACTATGATGGAAACCATGGAGCGACTGCAACGTAGGCATGGGGGAGGATTAGTCAGAGTGCCATTGTCTCGCAACTCCACACATGAGATGTACTGGGTCTCTGGGGCAAAGAGCAACATCATAAAAAGTGTGTCCACCACAAGTCAGCTCCTCCTGGGACGCATGGATGGCCCCAGGAGGCCAGTGAAATATGAGGAGGATGTGAACCTCGGCTCGGGTACACGAGCTGTGGCAAGCTGTGCTGAGGCTCCTAACATGAAAATCATCGGCAGGCGCATTGAGAGAATCCGCAATGAACATGCAGAAACATGGTTTCTTGATGAAAACCACCCATACAGGACATGGGCCTACCATGGGAGCTACGAAGCCCCCACGCAAGGATCAGCGTCTTCCCTCGTGAACGGGGTTGTTAGACTCCTGTCAAAGCCTTGGGACGTGGTGACTGGAGTTACAGGAATAGCCATGACTGACACCACACCATACGGCCAACAAAGAGTCTTCAAAGAAAAAGTGGACACCAGGGTGCCAGATCCCCAAGAAGGCACTCGCCAGGTAATGAACATAGTCTCTTCCTGGCTGTGGAAGGAGCTGGGGAAACGCAAGCGGCCACGCGTCTGCACCAAAGAAGAGTTTATCAACAAGGTGCGCAGCAATGCAGCACTGGGAGCAATATTTGAAGAGGAAAAAGAATGGAAGACGGCTGTGGAAGCTGTGAATGATCCAAGGTTTTGGGCCCTAGTGGATAGGGAGAGAGAACACCACCTGAGAGGAGAGTGTCACAGCTGTGTGTACAACATGATGGGAAAAAGAGAAAAGAAGCAAGGAGAGTTCGGGAAAGCAAAAGGTAGCCGCGCCATCTGGTACATGTGGTTGGGAGCCAGATTCTTGGAGTTTGAAGCCCTTGGATTCTTGAACGAGGACCATTGGATGGGAAGAGAAAACTCAGGAGGTGGAGTCGAAGGGTTAGGATTGCAAAGACTTGGATACATTCTAGAAGAAATGAATCGGGCACCAGGAGGAAAGATGTACGCAGATGACACTGCTGGCTGGGACACCCGCATTAGTAAGTTTGATCTGGAGAATGAAGCTCTGATTACCAACCAAATGGAGGAAGGGCACAGAACTCTGGCGTTGGCCGTGATTAAATACACATACCAAAACAAAGTGGTGAAGGTTCTCAGACCAGCTGAAGGAGGAAAAACAGTTATGGACATCATTTCAAGACAAGACCAGAGAGGGAGTGGACAAGTTGTCACTTATGCTCTCAACACATTCACCAACTTGGTGGTGCAGCTTATCCGGAACATGGAAGCTGAGGAAGTGTTAGAGATGCAAGACTTATGGTTGTTGAGGAAGCCAGAGAAAGTGACCAGATGGTTGCAGAGCAATGGATGGGATAGACTCAAACGAATGGCGGTCAGTGGAGATGACTGCGTTGTGAAGCCAATCGATGATAGGTTTGCACATGCCCTCAGGTTCTTGAATGACATGGGAAAAGTTAGGAAAGACACACAGGAGTGGAAACCCTCGACTGGATGGAGCAATTGGGAAGAAGTCCCGTTCTGCTCCCACCACTTCAACAAGCTGTACCTCAAGGATGGGAGATCCATTGTGGTCCCTTGCCGCCACCAAGATGAACTGATTGGCCGAGCTCGCGTCTCACCAGGGGCAGGATGGAGCATCCGGGAGACTGCCTGTCTTGCAAAATCATATGCGCAGATGTGGCAGCTCCTTTATTTCCACAGAAGAGACCTTCGACTGATGGCTAATGCCATTTGCTCGGCTGTGCCAGTTGACTGGGTACCAACTGGGAGAACCACCTGGTCAATCCATGGAAAGGGAGAATGGATGACCACTGAGGACATGCTCATGGTGTGGAATAGAGTGTGGATTGAGGAGAACGACCATATGGAGGACAAGACTCCTGTAACAAAATGGACAGACATTCCCTATCTAGGAAAAAGGGAGGACTTATGGTGTGGATCCCTTATAGGGCACAGACCCCGCACCACTTGGGCTGAAAACATCAAAGACACAGTCAACATGGTGCGCAGGATCATAGGTGATGAAGAAAAGTACATGGACTATCTATCCACCCAAGTCCGCTACTTGGGTGAGGAAGGGTCCACACCCGGAGTGTTGTAA

>ENA|AMK49164|AMK49164.2 Zika virus polyprotein

ATGAAAAACCC-AAAAAAGAAATCCGGAGGATTCCGGATTGTCAATATGCTAAAACGCGGAGTAGCCCGTGTGAGCCCCTTTGGGGGCTTGAAGAGGCTGCCAGCCGGACTTCTGCTGGGTCATGGGCCCATCAGGATGGTCTTGGCGATTCTAGCCTTTTTGAGATTCACGGCAATCAAGCCATCACTGGGTCTCATCAATAGATGGGGTTCAGTGGGGAAAAAAGAGGCTATGGAAATAATAAAGAAGTTCAAGAAAGATCTGGCTGCCATGCTGAGAATAATCAATGCTAGGAAGGAGAAGAAGAGACGAGGCGCAGATACTAGTGTCGGAATTGTTGGCCTCCTGCTGACCACAGCTATGGCAGCGGAGGTCACTAGACGTGGGAGTGCATACTATATGTACTTGGACAGAAACGATGCTGGGGAGGCCATATCTTTTCCAACCACATTGGGGATGAATAAGTGTTATATACAGATCATGGATCTTGGACACATGTGTGATGCCACCATGAGCTATGAATGCCCTATGCTGGATGAGGGGGTGGAACCAGATGACGTCGATTGTTGGTGCAACACGACGTCAACTTGGGTTGTGTACGGAACCTGCCATCACAAAAAAGGTGAAGCACGGAGATCTAGAAGAGCTGTGACGCTCCCCTCCCATTCCACTAGGAAGCTGCAAACGCGGTCGCAAACCTGGTTGGAATCAAGAGAATACACAAAGCACTTGATTAGAGTCGAAAATTGGATATTCAGGAACCCTGGCTTCGCGTTAGCAGCAGCTGCCATCGCTTGGCTTTTGGGAAGCTCAACGAGCCAAAAAGTCATATACTTGGTCATGATACTGCTGATTGCCCCGGCATACAGCATCAGGTGCATAGGAGTCAGCAATAGGGACTTTGTGGAAGGTATGTCAGGTGGGACTTGGGTTGATGTTGTCTTGGAACATGGAGGTTGTGTCACCGTAATGGCACAGGACAAACCGACTGTCGACATAGAGCTGGTTTCAACAACAGTCAGCAACATGGCGGAGGTAAGATCCTACTGCTATGAGGCAACAATATCAGACATAGCTTCGGACAGCCGCTGCCCAACACAAGGTGAAGCCTACCTTGACAAGCAATCAGACACTCAATATGTCTGCAAAAGAACGTTAGTGGACAGAGGCTGGGGAAATGGATGTGGACTTTTTGGCAAAGGGAGCCTGGTGACATGCGCTAAGTTTGCATGCTCCAAGAAAATGACCGGGAAGAGCATCCAGCCAGAGAATCTGGAGTACCGGATAATGCTGTCAGTTCATGGCTCCCAGCACAGTGGGATGATCGTTAATGACACAGGACATGAAACTGATGAGAATAGAGCGAAGGTTGAGATAACGCCCAATTCACCAAGAGCCGAAGCCACCCTGGGGGGTTTTGGAAGCCTAGGACTTGATTGTGAACCGAGGACAGGCCTTGACTTTTCAGATTTGTATTACTTGACTATGAATAACAAGCACTGGTTGGTTCACAAGGAGTGGTTCCACGACATTCCATTACCTTGGCACGCTGGGGCAGACACCGGAACTCCACACTGGAACAACAAAGAAGCACTGGTAGAGTTCAAGGACGCACATGCCAAAAGGCAAACTGCCGTGGTTCTAGGGAGTCAAGAAGGAGCAGTTCACACGGCCCTTGCTGGAGCTCTGGAGGCTGAGATGGATGGTGCAAAGGGAAGGCTGTCCTCTGGCCACTTGAAATGTCGCCTGAAAATGGATAAACTTAGATTGAAGGGCGTGTCATACTCCTTGTGTACCGCAGCGTTCACATTCACCAAGATCCCGGCTGAAACACTGCACGGGACAGTCACAGTGGAGGTACAGTACGCAGGGACAGATGGACCTTGCAAGGTTCCAGCTCAGATGGCGGTGGACATGCAAACTCTGACCCCAGTTGGGAGGTTGATAACCGCTAACCCCGTAATCACTGAAAGCACTGAGAACTCTAAGATGATGCTGGAACTTGATCCACCATTTGGGGACTCTTACATTGTCATAGGAGTCGGGGAGAAGAAGATCACCCACCACTGGCACAGGAGTGGCAGCACCATTGGAAAAGCATTTGAAGCCACTGTGAGAGGTGCCAAGAGAATGGCAGTCTTGGGAGACACAGCCTGGGACTTTGGATCAGTTGGAGGCGCTCTCAACTCATTGGGCAAGGGCATCCATCAAATTTTTGGAGCAGCTTTCAAATCATTGTTTGGAGGAATGTCCTGGTTCTCACAAATCCTCATTGGAACGTTGCTGATGTGGTTGGGTCTGAACACAAAGAATGGATCTATTTCCCTTATGTGCTTGGCCTTAGGGGGAGTGTTGATCTTCTTATCCACAGCCGTCTCTGCTGATGTGGGGTGCTCGGTGGACTTCTCAAAGAAGGAGACGAGATGCGGTACAGGGGTGTTCGTCTATAACGACGTTGAAGCCTGGAGGGACAGGTACAAGTACCATCCTGACTCCCCCCGTAGATTGGCAGCAGCAGTCAAGCAAGCCTGGGAAGATGGTATCTGCGGGATCTCCTCTGTTTCAAGAATGGAAAACATCATGTGGAGATCAGTAGAAGGGGAGCTCAATGCAATCCTGGAAGAGAATGGAGTTCAACTGACGGTCGTTGTGGGATCTGTGAAAAACCCCATGTGGAGAGGTCCACAGAGATTGCCCGTGCCTGTGAACGAGCTGCCCCACGGCTGGAAGGCTTGGGGGAAATCGTACTTCGTCAGAGCAGCAAAGACAAATAACAGCTTTGTCGTGGATGGTGACACACTGAAGGAATGCCCACTCAAACATAGAGCATGGAACAGCTTTCTTGTGGAGGATCATGGGTTCGGGGTATTTCACACTAGTGTCTGGCTCAAGGTTAGAGAAGATTATTCATTAGAGTGTGATCCAGCCGTTATTGGAACAGCTGTTAAGGAAAAGGAGGCTGTACACAGTGATCTAGGCTACTGGATTGAGAGTGAGAAGAATGACACATGGAGGCTGAAGAGGGCCCATCTGATCGAGATGAAAACATGTGAATGGCCAAAGTCCCACACATTGTGGACAGATGGAATAGAAGAGAGTGATCTGATCATACCCAAGTCTTTAGCTGGGCCACTCAGCCATCACAATACCAGAGAGGGCTACAGGACCCAAATGAAAGGGCCATGGCACAGTGAAGAGCTTGAAATTCGGTTTGAGGAATGCCCAGGCACTAAGGTCCACGTGGAGGAAACATGTGGAACAAGAGGACCATCTCTGAGATCAACCACTGCAAGCGGAAGGGTGATCGAGGAATGGTGCTGCAGGGAGTGCACAATGCCCCCACTGTCGTTCCGGGCTAAAGATGGCTGTTGGTATGGAATGGAGATAAGACCCAGGAAAGAACCAGAAAGCAACTTAGTAAGGTCAGTGGTGACTGCAGGATCAACTGATCACATGGATCACTTCTCCCTTGGAGTGCTTGTGATTCTGCTCATGGTGCAGGAAGGGCTGAAGAAGAGAATGACCACAAAGATCATCATAAGCACATCAATGGCAGTGCTGGTAGCTATGATCCTGGGAGGATTTTCAATGAGTGACCTGGCTAAGCTTGCAATTTTGATGGGTGCCACCTTCGCGGAAATGAACACTGGAGGAGATGTAGCTCATCTGGCGCTGATAGCGGCATTCAAAGTCAGACCAGCGTTGCTGGTATCTTTCATCTTCAGAGCTAATTGGACACCCCGTGAAAGCATGCTGCTGGCCTTGGCCTCGTGTCTTTTGCAAACTGCGATCTCCGCCTTGGAAGGCGACCTGATGGTTCTCATCAATGGTTTTGCTTTGGCCTGGTTGGCAATACGAGCGATGGTTGTTCCACGCACTGATAACATCACCTTGGCAATCCTGGCTGCTCTGACACCACTGGCCCGGGGCACACTGCTTGTGGCGTGGAGAGCAGGCCTTGCTACTTGCGGGGGGTTTATGCTCCTCTCTCTGAAGGGAAAAGGCAGTGTGAAGAAGAACTTACCATTTGTCATGGCCCTGGGACTAACCGCTGTGAGGCTGGTCGACCCCATCAACGTGGTGGGACTGCTGTTGCTCACAAGGAGTGGGAAGCGGAGCTGGCCCCCTAGCGAAGTACTCACAGCTGTTGGCCTGATATGCGCATTGGCTGGAGGGTTCGCCAAGGCAGATATAGAGATGGCTGGGCCCATAGCCGCGGTCGGTCTGCTAATTGTCAGTTACGTGGTCTCAGGAAAGAGTGTGGACATGTACATTGAAAGAGCAGGTGACATCACATGGGAAAAAGATGCGGAAGTCACTGGAAACAGTCCCCGGCTCGATGTGGCGCTAGATGAGAGTGGTGATTTCTCCCTGGTGGAGGATGACGGTCCCCCCATGAGAGAGATCATACTCAAGGTGGTCCTGATGACCATCTGTGGCATGAACCCAATAGCCATACCCTTTGCAGCTGGAGCGTGGTACGTATACGTGAAGACTGGAAAAAGGAGTGGTGCTCTATGGGATGTGCCTGCTCCCAAGGAAGTAAAAAAGGGGGAGACCACAGATGGAGTGTACAGAGTAATGACTCGTAGACTGCTAGGTTCAACACAAGTTGGAGTGGGAGTTATGCAAGAGGGGGTCTTTCACACTATGTGGCACGTCACAAAAGGATCCGCGCTGAGAAGCGGTGAAGGGAGACTTGATCCATACTGGGGAGATGTCAAGCAGGATCTGGTGTCATACTGTGGTCCATGGAAGCTAGATGCCGCCTGGGACGGGCACAGCGAGGTGCAGCTCTTGGCCGTGCCCCCCGGAGAGAGAGCGAGGAACATCCAGACTCTGCCCGGAATATTTAAGACAAAGGATGGGGACATTGGAGCGGTTGCGCTGGATTACCCAGCAGGAACTTCAGGATCTCCAATCCTAGACAAGTGTGGGAGAGTGATAGGACTTTATGGCAATGGGGTCGTGATCAAAAATGGGAGTTATGTTAGTGCCATCACCCAAGGGAGGAGGGAGGAAGAGACTCCTGTTGAGTGCTTCGAGCCTTCGATGCTGAAGAAGAAGCAGCTAACTGTCTTAGACTTGCATCCTGGAGCTGGGAAAACCAGGAGAGTTCTTCCTGAAATAGTCCGTGAAGCCATAAAAACAAGACTCCGTACTGTGATCTTAGCTCCAACCAGGGTTGTCGCTGCTGAAATGGAGGAAGCCCTTAGAGGGCTTCCAGTGCGTTATATGACAACAGCAGTCAATGTCACCCACTCTGGAACAGAAATCGTCGACTTAATGTGCCATGCCACCTTCACTTCACGCCTACTACAGCCAATCAGAGTCCCCAACTATAATCTGTATATTATGGATGAGGCCCACTTCACAGATCCCTCAAGTATAGCAGCAAGAGGATACATTTCAACAAGGGTTGAGATGGGCGAGGCGGCTGCCATCTTCATGACCGCCACGCCACCAGGAACCCGTGACGCATTTCCGGACTCCAACTCACCAATTATGGACACCGAAGTGGAAGTCCCAGAGAGAGCCTGGAGCTCAGGCTTTGATTGGGTGACGGATCATTCTGGAAAAACAGTTTGGTTTGTTCCAAGCGTGAGGAACGGCAATGAGATCGCAGCTTGTCTGACAAAGGCTGGAAAACGGGTCATACAGCTCAGCAGAAAGACTTTTGAGACAGAGTTCCAGAAAACAAAACATCAAGAGTGGGACTTTGTCGTGACAACTGACATTTCAGAGATGGGCGCCAACTTTAAAGCTGACCGTGTCATAGATTCCAGGAGATGCCTAAAGCCGGTCATACTTGATGGCGAGAGAGTCATTCTGGCTGGACCCATGCCTGTCACACATGCCAGCGCTGCCCAGAGGAGGGGGCGCATAGGCAGGAATCCCAACAAACCTGGAGATGAGTATCTGTATGGAGGTGGGTGCGCAGAGACTGACGAAGACCATGCACACTGGCTTGAAGCAAGAATGCTCCTTGACAATATTTACCTCCAAGATGGCCTCATAGCCTCGCTCTATCGACCTGAGGCCGACAAAGTAGCAGCCATTGAGGGAGAGTTCAAGCTTAGGACGGAGCAAAGGAAGACCTTTGTGGAACTCATGAAAAGAGGAGATCTTCCTGTTTGGCTGGCCTATCAGGTTGCATCTGCCGGAATAACCTACACAGATAGAAGATGGTGCTTTGATGGCACGACCAACAACACCATAATGGAAGACAGTGTGCCGGCAGAGGTGTGGACCAGACACGGAGAGAAAAGAGTGCTCAAACCGAGGTGGATGGACGCCAGAGTTTGTTCAGATCATGCGGCCCTGAAGTCATTCAAGGAGTTTGCCGCTGGGAAAAGAGGAGCGGCTTTTGGAGTGATGGAAGCCCTGGGAACACTGCCAGGACACATGACAGAGAGATTCCAGGAAGCCATTGACAACCTCGCTGTGCTCATGCGGGCAGAGACTGGAAGCAGGCCTTACAAAGCCGCGGCGGCCCAATTGCCGGAGACCCTAGAGACCATTATGCTTTTGGGGTTGCTGGGAACAGTCTCGCTGGGAATCTTTTTCGTCTTGATGAGGAACAAGGGCATAGGGAAGATGGGCTTTGGAATGGTGACTCTTGGGGCCAGCGCATGGCTCATGTGGCTCTCGGAAATTGAGCCAGCCAGAATTGCATGTGTCCTCATTGTTGTGTTCCTATTGCTGGTGGTGCTCATACCTGAGCCAGAAAAGCAAAGATCTCCCCAGGACAACCAAATGGCAATCATCATCATGGTAGCAGTAGGTCTTCTGGGCTTGATTACCGCCAATGAACTCGGATGGTTGGAGAGAACAAAGAGTGACCTAAGCCATCTAATGGGAAGGAGAGAGGAGGGAGCAACCATAGGATTCTCAATGGACATTGACCTGCGGCCAGCCTCAGCTTGGGCCATCTATGCTGCCTTGACAACTTTCATTACCCCAGCCGTCCAACATGCAGTGACCACTTCATACAACAACTACTCCTTAATGGCGATGGCCACGCAAGCTGGAGTGTTGTTTGGTATGGGCAAAGGGATGCCATTCTACGCATGGG-ACTTTGGAGTCCCGCTGCTAATGATAGGTTGCTACTCACAATTAACACCCCTGACCCTAATAGTGGCCATCATTTTGCTCGTGGCGCACTACATGTACTTGATCCCAGGGCTGCAGGCAGCAGCTGCGCGTGCTGCCCAGAAGAGAACGGCAGCTGGCATCATGAAGAACCCTGTTGTGGATGGAATAGTGGTGACTGACATTGACACAATGACAATTGACCCCCAAGTGGAGAAAAAGATGGGACAGGTGCTACTCATAGCAGTAGCCGTCTCCAGCGCCATACTGTCGCGGACCGCCTGGGGGTGGGGGGAGGCTGGGGCCCTGATTACAGCCGCAACTTCCACTTTGTGGGAAGGCTCTCCGAACAAGTACTGGAACTCCTCTACAGCCACTTCACTGTGTAACATTTTTAGGGGAAGTTACTTGGCTGGAGCTTCTCTAATCTACACAGTAACAAGAAACGCTGGCTTGGTCAAGAGACGTGGGGGTGGAACAGGAGAGACCCTGGGAGAGAAATGGAAGGCCCGCTTGAACCAGATGTCGGCCCTGGAGTTCTACTCCTACAAAAAGTCAGGCATCACCGAGGTGTGCAGAGAAGAGGCCCGCCGCGCCCTCAAGGACGGTGTGGCAACGGGAGGCCATGCTGTGTCCCGAGGAAGTGCAAAGCTGAGATGGTTGGTGGAGCGGGGATACCTGCAGCCCTATGGAAAGGTCATTGATCTTGGATGTGGCAGAGGGGGCTGGAGTTACTACGCCGCCACCATCCGCAAAGTTCAAGAAGTGAAAGGATACACAAAAGGAGGCCCTGGTCATGAAGAACCCGTGTTGGTGCAAAGCTATGGGTGGAACATAGTCCGTCTCAAGAGTGGGGTGGACGTCTTTCATATGGCGGCTGAGCCGTGTGACACGTTGCTGTGTGACATAGGTGAGTCATCATCTAGTCCTGAAGTGGAAGAAGCACGGACGCTCAGAGTCCTCTCCATGGTGGGGGATTGGCTTGAAAAAAGACCAGGAGCCTTTTGCATAAAAGTGTTGTGCCCATACACCAGCACTATGATGGAAACCTTGGAGCGACTGCAGCGTAGGTATGGGGGAGGACTGGTCAGAGTGCCACTCTCCCGCAACTCTACACATGAGATGTACTGGGTCTCTGGAGCGAAAAGCAACACCATAAAAAGTGTGTCCACCACGAGCCAGCTCCTCTTGGGGCGCATGGACGGGCCTAGGAGGCCAGTGAAATATGAGGAGGATGTGAATCTCGGCTCTGGCACGCGGGCTGTGGTAAGCTGCGCTGAAGCTCCCAACATGAAGATCATTGGTGACCGCATTGAAAGGATCCGCAGTGAGCACGCGGAAACGTGGTTCTTTGACGAGAACCACCCATATAGGACATGGGCTTACCATGGAAGCTATGAGGCCCCCACACAAGGGTCAGCGTCCTCTCTAATAAACGGGGTTGTCAGGCTCCTGTCAAAACCCTGGGATGTGGTGACTGGAGTCACAGGAATAGCCATGACCGACACCACACCGTATGGTCAGCAAAGAGTTTTCAAGGAAAAAGTGGACACTAGGGTGCCAGACCCCCAAGAAGGCACTCGTCAGGTTATGAGCATGGTCTCTTCCTGGTTGTGGAAAGAGCTAGGCAAACACAAACGGCCACGAGTCTGTACCAAAGAAGAGTTCATCAACAAGGTTCGTAGCAATGCAGCATTAGGGGCAATATTTGAAGAGGAAAAAGAGTGGAAGACTGCAGTGGAGGCTGTGAACGATCCAAGGTTCTGGGCTCTAGTGGACAAGGAAAGAGAGCACCACCTGAGAGGAGAGTGCCAGAGTTGTGTGTACAACATGATGGGAAAAAGAGAAAAGAAACAAGGGGAATTTGGAAAGGCCAAGGGCAGCCGCGCCATCTGGTATATGTGGCTAGGGGCTAGATTTCTAGAGTTCGAAGCCCTTGGATTCTTGAACGAGGATCACTGGATGGGGAGAGAGAACTCAGGAGGTGGTGTTGAAGGGCTGGGATTACAAAGACTCGGATATGTCCTAGAAGAGATGAGTCGCATACCAGGAGGAAGGATGTATGCAGATGACACTGCTGGCTGGGACACCCGCATCAGCAGGTTTGATCTGGAGAATGAAGCTCTAATCACCAACCAAATGGAGAAAGGGCACAGGGCCTTGGCACTGGCCATAATCAAGTACACATACCAAAACAAAGTGGTAAAGGTCCTTAGACCAGCTGAAAAAGGGAAAACAGTTATGGACATTATTTCGAGACAAGACCAAAGGGGGAGCGGACAAGTTGTCACTTACGCTCTTAACACATTTACCAACCTAGTGGTGCAACTCATTCGGAATATGGAGGCTGAGGAAGTTCTAGAGATGCAAGACTTGTGGCTGCTGCGGAGGTCAGAGAAAGTGACCAACTGGTTGCAGAGCAACGGATGGGATAGGCTCAAACGAATGGCAGTCAGTGGAGATGATTGCGTTGTGAAGCCAATTGATGATAGGTTTGCACATGCCCTCAGGTTCTTGAATGATATGGGAAAAGTTAGGAAGGACACACAAGAGTGGAAACCCTCAACTGGATGGGACAACTGGGAAGAAGTTCCGTTTTGCTCCCACCACTTCAACAAGCTCCATCTCAAGGACGGGAGGTCCATTGTGGTTCCCTGCCGCCACCAAGATGAACTGATTGGCCGGGCCCGCGTCTCTCCAGGGGCGGGATGGAGCATCCGGGAGACTGCTTGCCTAGCAAAATCATATGCGCAAATGTGGCAGCTCCTTTATTTCCACAGAAGGGACCTCCGACTGATGGCCAATGCCATTTGTTCATCTGTGCCAGTTGACTGGGTTCCAACTGGGAGAACTACCTGGTCAATCCATGGAAAGGGAGAATGGATGACCACTGAAGACATGCTTGTGGTGTGGAACAGAGTGTGGATTGAGGAGAACGACCACATGGAAGACAAGACCCCAGTTACGAAATGGACAGACATTCCCTATTTGGGAAAAAGGGAAGACTTGTGGTGTGGATCTCTCATAGGGCACAGACCGCGCACCACCTGGGCTGAGAACATTAAAAACACAGTCAACATGGTGCGCAGGATCATAGGTGATGAAGAAAAGTACATGGACTACCTATCCACCCAAGTTCGCTACTTGGGTGAAGAAGGGTCTACACCTGGAGTGCTGTAA

>ENA|AMK79469|AMK79469.1 Zika virus polyprotein

ATGAAAAACCC-AAAAAAGAAATCCGGAGGATTCCGGATTGTCAATATGCTAAAACGCGGAGTAGCCCGTGTGAGCCCCTTTGGGGGCTTGAAGAGGCTGCCAGCCGGACTTCTGCTGGGTCATGGGCCCATCAGGATGGTCTTGGCAATTCTAGCCTTTTTGAGATTCACGGCAATCAAGCCATCACTGGGTCTCATCAATAGATGGGGTTCAGTGGGGAAAAAAGATGCTATGGAAATAATAAAGAAGTTCAAGAAAGATCTGGCTGCCATGCTGAGAATAATCAATGCTAGGAAGGAGAAGAAGAGACGAGGCGCAGATACTAGTGTCGGAATTGTTGGCCTCCTGCTGACCACAGCTATGGCAGCGGAGGTCACTAGACGTGGGAGTGCATACTATATGTACTTGGACAGAAACGATGCTGGGGAGGCCATATCTTTTCCAACCACATTGGGGATGAATAAGTGTTATATACAGATCATGGATCTTGGACACATGTGTGATGCCACCATGAGCTATGAATGCCCTATGCTGGATGAGGGGGTGGAACCAGATGACGTCGATTGTTGGTGCAACACGACGTCAACTTGGGTTGTGTACGGAACCTGCCATCACAAAAAAGGTGAAGCACGGAGATCTAGAAGAGCTGTGACGCTCCCTTCCCATTCCACTAGGAAGCTGCAAACGCGGTCGCAAACCTGGTTGGAATCAAGAGAATACACAAAGCACTTGATTAGAGTCGAAAATTGGATATTCAGGAACCCTGGCTTCGCGTTAGCAGCAGCTGCCATCGCTTGGCTTTTGGGAAGCTCAACGAGCCAAAAAGTCATATACTTGGTCATGATACTGCTGATTGCCCCGGCATACAGCATCAGGTGCATAGGAGTCAGCAATAGGGACTTTGTGGAAGGTATGTCAGGTGGGACTTGGGTTGATGTTGTCTTGGAACATGGAGGTTGTGTCACCGCAATGGCACAGGACAAACCGACTGTCGACATAGAGCTGGTTACAACAACAGTCAGCAACATGGCGGAGGTAAGATCCTACTGCTATGAGGCATCAATATCAGACATGGCTTCGGACAGCCGCTGCCCAACACAAGGTGAAGCCTACCTTGACAAGCAATCAGACACTCAATATGTTTGCAAAAGAACGTTAGTGGACAGAGGCTGGGGAAATGGATGTGGACTTTTTGGCAAAGGGAGTCTGGTGACATGCGCTAAGTTTGCATGCTCCAAGAAAATGACCGGGAAGAGCATCCAGCCAGAGAATCTGGAGTACCGGATAATGCTGTCAGTTCATGGCTCCCAGCACAGTGGGATGCTCGTTAATGACACAGGACATGAAACTGATGAGAATAGAGCGAAGGTTGAGATAACGCCCAATTCACCAAGAGCCGAAGCCACCCTGGGGGGTTTTGGAAGCCTAGGACTTGATTGTGAACCGAGGACAGGCCTTGACTTTTCAGATTTGTATTACTTGACTATGAATAACAAGCACTGGTTGGCTCACAAGGAGTGGTTCCACGACATTCCATTACCTTGGCACGCTGGGGCAGCCACCGGAACTCCACACTGGAACAACAAAGAAGCACTGGTAGAGTTCAAGGACGCACATGCCAAAAGGCAAACTGTCGTGGTTCTAGGGAGTCAAGAAGGAGCAGTTCACACGGCCCTTGCTGGAGCTCTGGAGGCTGAGATGGATGGTGCAAAGGGAAGGCTGTCCTCTGGCCACTTGAAATGTCGCCTGAAAATGGATAAACTTAGATTGAAGGGCGTGTCATACTCCTTGTGTACCGCAGCGTTCACATTCACCAAGATCCCGGCTGAAACAGTGGACGGGACAGTCACAGTGGAGGGACAGTACGGAGGGACAGATGGACCTTGCAAGGTTCCAGCTCAGATGGCGGTGGACATGCAGACTCTGACCCCAGTTGGGAGGTTGATAACCGCTAACCCCGTAATCACTGAAAGCACTGAGAACTCTAAGATGATGCTGGAACTTGATCCACCATTTGGGGACTCTTACATTGTCATAGGAGTCGGGGAGAAGAAGATCACCCACCACTGGCACAGGAGTGGCAGCACCATTGGAAAAGCATTTGAAGCCACTGTGAGAGGTGCCAAGAGAATGGCAGTCTTGGGAGACACAGCCTGGGACTTTGGATCAGTTGGAGGCGCTCTCAACTCATTGGGCAAGGGCATCCATCAAATTATTGGAGCAGCTTTCAAATCATTGTTTGGAGGAATGTCCTGGTTCTCACAAATTCTCATTGGGACGTTGCTGATGTGGTTGGGTCTGAACACAAAGAATGGATCTATTTCCCTTATGTGCTTGGCCTTAGGGGGAGTGTTGATCTTCTTATCCACAGCCGTCTCAGGTGGTGTGGGGTGCTCGGTGGACTTCTCAAAGAAGGAGACGAGATGCGGTACAGGGGTGTTCGTCTATAACGATGTTGAAGCCTGGAGGGACAGGTACAAGTACCATCCTGACTCCCCCCGTAGATTGGCAGCAGCAGTCAAGCAAGCCTGGGAAGATGGTATCTGCGGGATCTCCTCTGTTTCAAGAATGGAAAACATCATGTGGAGATCAGTAGAAGGGGAGCTCAACGCAATCCTGGAAGAGAATGGAGTTCAACTGACGGTCGTTGTGGGATCTGTAAAAAACCCCATGTGGAGAGGTCCACAGAGATTGCCCGTGCCTGTGAACGAGCTGCCCCACGGCTGGAAGGCTTGGGGGAAATCGTACTTCGTCAGAGCAGCAAAGACAAATAACAGCTTTGTCGTGGATGGTGACACACTGAAGGAATGCCCACTCAAACATAGAGCATGGAACAGCTTTCTTGTGGAGGATCATGGGTTCGGGGTATTTCACACTAGTGTCTGGCTCAAGGTTAGAGAAGACTATTGGTTAGAGTGTGATCCAGCCGTTATTGGAACAGCTGTTAAGGGAAAGGAGGCTGTACACAGTGATCTAGGCTACTGGATTGAGAGTGAGAAGAATGACACATGGTGGCTGAAGAGGGCCCATCTGATCGAGATGAAAACATGTGAATGGCCAAAGTCCCACACATTGTGGACAGATGGAATAGAAGAGAGTGATCTGATCATACCCAAGTCTTTAGCTGGGCCACTCAGCCATCACAATGCCAGAGAGGGCTACAGGACCCAAATGAAAGGGCCATGGCACAGTGAAGAGCTTGAAATTCGGTTTGAGGAATGCCCAGGCACTAAGGTCCACGTGGAGGAAACATGTGGAACAAGAGGACCATCTCTGAGATCAACCACTGCAAGCGGAAGGGTGATCGAGGAATGGTGCTCCAGGGAGTGCACAATGCCCCCACTGTCCTTCCAGGCTAAAGATGGCTGTTGGTATGGAATGGAGATAAGGCCCAGGAAAGAACCAGAAAGCAACTTAGTAAGGTCAATGGTGACTGCAGGATCAACTGATCACATGGATCACTTCTCCCTTGGAGTGCTTGTGATTCTGCTCATGGTGCAGGAAGGGCTGAAGAAGAGAATGACCACAAAGATCATCATAAGCACATCAATGGCAGTGCTGGTAGCTATGATCCTGGGAGGATTTTCAATGAGTGACCTGGCTAAGCTTGCAATTTTGATGGGTGCCACCTTCGCGGAAATGAACACTGGAGGAGATGTAGCTCATCTGGCGCTGATAGCGGCATTCAAAGTCAGACCAGCGTTGCTGGTATCTTTCATCTTCAGAGCTAATTGGACACCCCGTGAAAGCATGCTGCTGGCCTTGGCCTCGTGTCTTTTGCAAACTGCGATCTCCGCCTTGGAAGGCGACCTGATGGTTCTCATCAATGGTTTTGCTTTGGCCTGGTTGGCAATACGAGCGATGGTTGTTCCACGCACTGATAACATCACCTTAGCAATCCTGGCTGCTCTGACACCACTGGCCCGGGGCACACTGCTTGTGGCGTGGAGAGCAGGCCTTGCTACTTGCGGGGGGTTTATGCTCCTCTCTCTGAAGGGAAAAGGCAGTGTGAAGAAGAACTTACCATTTGTCATGGCCCTGGGACTAACCGCTGTGAGGCTGGTCGACCCCATCAACGTGGTGGGACTGCTGTTGCTCACAAGGAGTGGGAAGCGGAGCTGGCCCCCTAGCGAAGTACTCACAGCTGTTGGCCTGATATGCGCATTGGCTGGAGGGTTCGCCAAGGCAGATATAGAGATGGCTGGGCCCATGGCCGCGGTCGGTCTGCTAATTGTCAGTTACGTGGTCTCAGGAAAGAGTGTGGACATGTACATTGAAAGAGCAGGTGACATCACATGGGAAAAAGATGCGGAAGTCACTGGAAACAGTCCCCGGCTCGATGTGGCGCTAGATGAGAGTGGTGATTTCTCCCTGGTGGAGGATGACGGTCCCCCCATGAGAGAGATCATACTCAAGGTGGTCCTGATGACCATCTGTGGCATGAACCCAATAGCCATACCCTTTGCAGCTGGAGCGTGGTACGTATACGTGAAGACTGGAAAAAGGAGTGGTGCTCTATGGGATGTGCCTGCTCCCAAGGAAGTAAAAAAGGGGGAGACCACAGATGGAGTGTACAGAGTAATGACTCGCAGACTGCTAGGTTCAACACAAGTTGGAGTGGGAGTTATGCAAGAGGGGGTCTTTCACACTATGTGGCACGTCACAAAAGGATCCGCGCTGAGAAGCGGTGAAGGGAGACTTGATCCATACTGGGGAGATGTCAAGCAGGATCTGGTGTCATACTGTGGTCCATGGAAGCTAGATGCCGCCTGGGACGGGCACAGCGAGGTGCAGCTCTTGGCCGTGCCCCCCGGAGAGAGAGCGAGGAACATCCAGACTCTGCCCGGAATATTTAAGACAAAGGATGGGGACATTGGAGCGGTTGCACTGGATTACCCAGCAGGAACTTCAGGATCTCCAATCCTAGACAAGTGTGGGAGAGTGATAGGACTTTATGGCAATGGGGTCGTGATCAAAAATGGGAGTTATGTTAGTGCCATCACCCAAGGGAGGAGGGAGGAAGAGACTCCTGTTGAGTGCTTCGAGCCTTCGATGCTGAAGAAGAAGCAGCTAACTGTCTTAGACTTGCATCCTGGAGCTGGGAAAACCAGGAGAGTTCTTCCTGAAATAGTCCGTGAAGCCATAAAAACAAGACTCCGTACTGTGATCTTGGCTCCAACCAGGGTTGTCGCTGCTGAAATGGAGGAGGCCCTTAGAGGGCTTCCAGTGCGTTATATGACAACAGCAGTCAATGTCACCCACTCTGGAACAGAAATCGTCGACTTAATGTGCCATGCCACCTTCACTTCACGTCTACTACAGCCAATTAGAGTCCCCAACTATAATCTGTATATTATGGATGAGGCCCACTTCACAGATCCCTCAAGTATAGCAGCAAGAGGATACATTTCAACAAGGGTTGAGATGGGCGAGGCGGCTGCCATCTTCATGACCGCCACGCCACCAGGAACCCGTGACGCATTTCCGGACTCCAACTCACCAATTATGGACACCGAAGTGGAAGTCCCAGAGAGAGCCTGGAGCTCAGGCTTTGATTGGGTGACGGAGTATTCTGGAAAAACAGTTTGGTTTGTTCCACGCGTGAGGAACGGCAATGAGATCGCAGCTTGTCTGACAAAGGCTGGAAAACGGGTCATACAGCTCAGCAGAAAGACTTTTGAGACAGAGTTCCAGAAAACAAAACATCAAGAGTGGGACTTTGTCGTGACAACTGACATTTCAGAGATGGGCGCCAACTTTAAAGCTGACCGTGTCATAGATTCCAGGAGATGCCTAAAGCCGGTCATACTTGGTGGCGAGAGAGTCATTCTGGCTGGACCCATGCCTGTCACACATGCCAGCGCTGCCCAGAGGAGGGGGCGCATAGGCAGGAATCCCAACAAACCTGGAGATGAGTATCTGTATGGAGGTGGGTGCGCAGAGACTGACGAAGACCATGCACACTGGCTTGAAGCAAGAATGCTCCTTGACAATATTTACCTCCAAGATGGCCTCATAGCCTCGCTCTATCGACCTGAGGCCGACAAAGTAGCAGCCATTGAGGGAGAGTTCAAGCTTAGGACGGAGCAAAGGAAGACCTTTGTGGAACTCATGAAAAGAGGAGATCTTCCTGTTTGGCTGGCCTATCAGGTTGCATCTGCCGGAATAACCTACACAGATAGAAGATGGTGCTTTGATGGCACGACCAACAACACCATAATGGAAGACAGTGTGCCGGCAGAGGTGTGGACCAGACACGGAGAGAAAAGAGTGCTCAAACCGAGGTGGATGGACGCCAGAGTTTGTTCAGATCATGCGGCCCTGAAGTCATTCAAGGAGTTTGCCGCTGGGAAAAGAGGAGCGGCTTTTGGAGTGATGGAAGCCCTGGGAACACTGCCAGGACACATGACAGAGAGATTCCAGGAAGCCATTGACAACCTCGCTGTGCTCATGCGGGCAGAGACTGGAAGCAGGCCTTACAAAGCCGCGGCGGCCCAATTGCCGGAGACCCTAGAGACCATTATGCTTTTGGGGTTGCTGGGAACAGTCTCGCTGGGAATCTTTTTCGTCTTGATGAGGAACAAGGGCATAGGGAAGATGGGCTTTGGAATGGTGACTCTTGGGGCCAGCGCATGGCTCATGTGGCTCTCGGAAATTGAGCCAGCCAGAATTGCATGTGTCCTCATTGTTGTGTTCCTATTGCTGGTGGTGCTCATACCTGAGCCAGAAAAGCAAAGATCTCCCCAGGACAACCAAATGGCCATCATCATCATGGTAGCAGTAGGTCTTCTGGGCTTGATTACCGCCAATGAACTCGGATGGTTGGAGAGAACAAAGAGTGACCTAAGCCATCTAATGGGAAGGAGAGAGGAGGGGGCAACCATGGGATTCTCAATGGACATTGACCTGCGGCCAGCCTCAGCTTGGGCCATCTATCCTGCCTTGACATCTTTCATTACCCCAGCCGTCCAACATGCAGTGACCACTTCATACAACAACTACTCCTTAATGGCGATGGCCACGCAAGCTGGAGTGTTGTTTGGTATGGGCAAAGGGATGCCATTCTACGCATGGG-ACTTTGGAGTCCCGCTGCTAATGATAGGTTGCTACTCACAATTAACGCCCCTGACCCTAATAGTGGCCATCATTTTGCTCGTGGCGCACTACATGTACTTGATCCCAGGGCTGCAGGCAGCAGCTGCGCGTGCTGCCCAGAAGAGAACGGCAGCTGGCATCATGAAGAACCCTGTTGTGGAGGGAATAGTGGTGACTGACATTGACACAATGACAATTGACCCCCAAGTGGAGAAAAAGATGGGACAGGTGCTACTCATGGCAGTAGCCGTCTCCAGCGCCATACTGTCGAGGACCGCCTGGGGGTGGGGGGAGGCTGGGGCCCTGATCACAGCCGCAACTTCCACTTTGTGGGAAGGCTCTCCGAACAAGTACTGGAACTCCTCTACAGCCACCTCACTGTGTAACATTTTTAGGGGAAGTTACTTGGCTGGAGCTTCTCTAATCTACACAGTAACAAGAAACGCTGGCTTGGTCAAGAGACGTGGGGGTGGAACAGGAGAGACCCTGGGAGAGAAATGGAAGGCCCGCTTGAACCAGATGTCGGCCCTGGAGTTCTACTCCTACAAAAAGTCAGGCATCACCGAGGTGTGCAGAGAAGAGGCCCGCCGCGCCCTCAAGGACGGTGTGGCAACGGGAGGCCATGCTGTGTCCCGAGGAAGTGCAAAGCTGAGATGGTTGGTGGAGCGGGGATACCTGCAGCCCTATGGAAAGGTCATTGATCTTGGATGTGGCAGAGGGGGCTGGAGTTACTACGCCGCCACCATCCGCAAAGTTCAAGAAGTGAAAGGATACACAAAAGGAGGCCCTGGTCATGAAGAACCCGTGTTGGTGCAAAGCTATGGGTGGAACATAGTCCGTCTTAAGAGTGGGGTGGACGTCTTTCATATGGCGGCTGAGCCGTGTGACACGTTGCTGTGTGACATAGGTGAGTCATCATCTAGTCCTGAAGTGGAAGAAGCACGGACGCTCAGAGTCCTCTCCATGGTGGGGGATTGGCTTGAAAAAAGACCAGGAGCCTTTTGTATAAAAGTGTTGTGCCCATACACCAGCACTATGATGGAAACCCTGGAGCGACTGCAGCGTAGGTATGGGGGAGGACTGGTCAGAGTGCCACTCTCCCGCAACTCTACACATGAGATGTACTGGGTCTCTGGAGCGAAAAGCAACACCATAAAAAGTGTGTCCACCACGAGCCAGCTCCTCTTGGGGCGCATGGACGGGCCTAGGAGGCCAGTGAAATATGAGGAGGATGTGAATCTCGGCTCTGGCACGCGGGCTGTGGTAAGCTGCGCTGAAGCTCCCAACATGAAGATCATTGGTAACCGCATTGAAAGGATCCGCGCTGAGAAAGCGGAAACGTGGTTCTTTGACGAGAACCACCCATATAGGACATGGGCTTACCATGGAAGCTATGATGCCGCCACACAAGGGTCAGCGTCCTCTCTAATAAACGGGGTTGTCAGGCTCCTGTCAAAACCCTGGGATGTGGTGACTGGAGTCACAGGAATAGCCATGACCGACACCACACCGTATGGTCAGCAAAGAGTTTTCAAGGAAAAAGTGGACACTAGGGTGCCAGACCCCCAAGAAGGCACTCGTCAGGTTATGAGCATGGTCTCTTCCTGGTTGTGGAAAGAGCTAGGCAAACACAAACGGCCACGAGTCTGTACCAAAGAAGAGTTCATCAACAAGGTTCGTAGCAATGCAGCATTAGGGGCAATATTTGAAGAGGAAAAAGAGTGGAAGACTGCAGTGGAAGCTGTGAACGATCCAAGGTTCTGGGCTCTAGTGGACAAGGAAAGAGAGCACCACCTGAGAGGAGAGTGCCAGAGTTGTGTGTACATCACAATGGGAAAAAGAGAAAAGAAACAAGGGGAATTTGGAAAGGCCAAGGGCAGCCGCGCCATCTGGTATATGTGGCTAGGGGCTAGATTTCTAGAGTTCGAAGCCCTTGGATTCTTGAACGAGGATCACTGGATGGGGAGAGAGAACTCAGGAGGTGGTGTTGAAGGGCTGGGATTACAAAGACTCGGATATGTCCTAGAAGAGATGAGTCGCATACCAGGAGGAAGGATGTATGCAGATGACACTGCTGGCTGGGACACCCGCATCAGCAGGTTTGATCTGGAGAATGAAGCTCTAATCACCAACCAAATGGAGAAAGGGCACAGGGCCTTGGCATTGGCCATAATCAAGTACACATACCAAAACAAAGTGGTAAAGGTCCTTAGACCAGCTGAAAAAGGGAAGACAGTTATGGACATTATTTCGAGACAAGACCAAAGGGGGAGCGGACAAGTTGTCACTTACGCTCTCAACACATTTACCAACCTAGTGGTGCAACTCATTCGGAATATGGAGGCTGAGGAAGTTCTAGAGATGCAAGACTTGTGGCTGCTGCGGAGGTCAGAGAAAGTGACCAACTGGTTGCAGAGCAACGGATGGGATAGGCTCAAACGAATGGCGGTCAGTGGAGATGATTGCGTTGTGAAACCAATTGATGATAGGTTTGCACATGCCCTCAGGTTCTTGAATGATATGGGAAAAGTTAGGAAGGACACACAAGAGTGGAAACCCTCAACTGGATGGGACAACTGGGAAGAAGTTCCCTTCTGCTCCCACCACTTCAACAAGCTCCATCTCAAGGACGGGAGGTCCATTGTGGTTCCCTGCCGCCACCAAGATGAACTGATTGGCCGGGCCCGCGTCTCTCCAGGGGCGGGATGGAGCATCCGGGAGACTGCTTGCCTAGCAAAATCATATGCGCAAATGTGGCAGCTCCTTTATTTCCACAGAAGGGACCTCCGACTGATGGCCAATGCCATTTGTTCATCTGTGCCAGTTGACTGGGTTCCAACTGGGAGAACTACCTGGTCAATCCATGGAAAGGGAGAATGGATGACCACTGAAGACATGCTTGTGGCGTGGAACAGAGTGTGGATTGAGGAGAACGACCACATGGAAGACAAGACCCCAGTCACGAAATGGACAGACATTCCCTATTTGGGAAAAAGGGAAGACTTGTGGTGTGGATCTCTCATAGGGCACAGACCGCGCACCACCTGGGCTGAGAACATTAAAAACACAGTCAACATGGTGCGCAGGATCATAGGTGATGAAGAAAAGTACATGGACTACCTATCCACCCAAGTTCGCTACTTGGGTGAAGAAGGGTCTACACCTGGAGTGCTGTAA

>ENA|AMM39804|AMM39804.2 Zika virus polyprotein

ATGAAAAACCC-AAAAAAGAAATCCGGAGGATTCCGGATTGTCAATATGCTAAAACGCGGAGTAGCCCGTGTGAGCCCCTTTGGGGGCTTGAAGAGGCTGCCAGCCGGACTTCTGCTGGGTCATGGGCCCATCAGGATGGTCTTGGCGATTCTAGCCTTTTTGAGATTCACGGCAATCAAGCCATCACTGGGTCTCATCAATAGATGGGGTTCAGTGGGGAAAAAAGAGGCTATGGAAATAATAAAGAAGTTCAAGAAAGATCTGGCTGCCATGCTGAGAATAATCAATGCTAGGAAGGAGAAGAAGAGACGAGGCGCAGAAACTAGTGTCGGAATTGTTGGCCTCCTGCTGACCACAGCTATGGCAGCGGAGGTCACTAGACGTGGGAGTGCATACTATATGTACTTGGACAGAAACGATGCTGGGGAGGCCATATCTTTTCCAACCACATTGGGGATGAATAAGTGTTATATACAGATCATGGATCTTGGACACATGTGTGATGCCACCATGAGCTATGAATGCCCTATGCTGGATGAGGGGGTGGAACCAGATGACGTCGATTGTTGGTGCAACACGACGTCAACTTGGGTTGTGTACGGAACCTGCCATCACAAAAAAGGTGAAGCACGGAGATCTAGAAGAGCCGTGACGCTCCCCTCCCATTCCACTAGGAAGCTGCAAACGCGGTCGCAAACCTGGTTGGAATCAAGAGAATACACAAAGCACTTGATTAGAGTCGAAAATTGGATATTCAGGAACCCTGGTTTCGCTTTAGCAGCAGCTGCCATCGCTTGGCTTTTGGGAAGCTCAACGAGCCAAAAAGTCATATACTTGGTCATGATACTGCTGATTGCCCCGGCATACAGCATCAGGTGCATAGGAGTCAGCAATAGGGACTTTGTGGAAGGTATGTCAGGTGGGACTTGGGTTGATGTCGTCTTGGAACATGGAGGTTGTGTCACCGTAATGGCACAGGACAAACCGACTGTCGACATAGAGCTGGTTACAACAACAGTCAGCAACATGGCGGAGGTAAGATCCTACTGCTATGAGGCATCAATATCAGACATGGCTTCGGACAGCCGCTGCCCAACACAAGGTGAAGCCTACCTTGACAAGCAATCAGACACTCAATATGTCTGCAAAAGAACGTTAGTGGACAGAGGCTGGGGAAATGGATGTGGACTTTTTGGCAAAGGGAGCCTGGTGACATGCGCTAAGTTTGCATGCTCCAAGAAAATGACCGGGAAGAGCATCCAGCCAGAGAATCTGGAGTACCGGATAATGTTGTCAGTTCATGGCTCCCAGCACAGTGGGATGATCGTTAATGACACAGGACATGAAACTGATGAGAATAGAGCGAAGGTTGAGATAACGCCCAATTCACCAAGAGCCGAAGCCACCCTGGGGGGCTTTGGAAGCCTAGGACTTGATTGTGAACCGAGGACAGGCCTTGACTTTTCAGATTTGTATTACTTGACTATGAATAACAAGCACTGGTTGGTTCACAAGGAGTGGTTCCACGACATTCCATTACCTTGGCACGCTGGGGCAGACACCGGAACTCCACACTGGAACAACAAAGAAGCACTGGTAGAGTTCAAGGACGCACATGCCAAAAGGCAAACTGTCGTGGTTCTAGGGAGTCAAGAAGGAGCAGTTCACACGGCCCTTGCTGGAGCTCTGGAGGCTGAGATGGATGGTGCAAAGGGAAGGCTGTCCTCTGGCCACTTGAAATGTCGCCTGAAAATGGATAAACTTAGATTGAAGGGCGTGTCATACTCCTTGTGTACCGCAGCGTTCACATTCACCAAGATCCCGGCTGAAACACTGCACGGGACAGTCACAGTGGAGGTACAGTACGCAGGGACAGATGGACCTTGCAAGGTTCCAGCTCAGATGGCGGTGGACATGCAAACTCTGACCCCAGTTGGGAGGTTGATAACCGCTAACCCCGTAATCACTGAAAGCACTGAGAACTCTAAGATGATGCTGGAACTTGATCCACCATTTGGGGACTCTTACATTGTCATAGGAGTCGGGGAGAAGAAGATCACCCACCACTGGCACAGGAGTGGCAGCACCATTGGAAAAGCATTTGAAGCCACTGTGAGAGGTGCCAAGAGAATGGCAGTCTTGGGAGACACAGCCTGGGACTTTGGATCAGTTGGAGGCGCTCTCAACTCATTGGGCAAGGGCATCCATCAAATTTTTGGAGCAGCTTTCAAATCATTGTTTGGAGGAATGTCCTGGTTCTCACAAATTCTCATTGGAACGTTGCTGATGTGGTTGGGTCTGAACACAAAGAATGGATCTATTTCCCTTATGTGCTTGGCCTTAGGGGGAGTGTTGATCTTCTTATCCACAGCCGTCTCTGCTGATGTGGGGTGCTCGGTGGACTTCTCAAAGAAGGAGACGAGATGTGGTACAGGGGTGTTCGTCTATAACGACGTTGAAGCCTGGAGGGACAGGTACAAGTACCATCCTGACTCCCCCCGTAGATTGGCAGCAGCAGTCAAGCAAGCCTGGGAAGATGGTATCTGCGGGATCTCCTCTGTTTCAAGAATGGAAAACATCATGTGGAGATCAGTAGAAGGGGAGCTCAACGCAATCCTGGAAGAGAATGGAGTTCAACTGACGGTCGTTGTGGGATCTGTAAAAAACCCCATGTGGAGAGGTCCACAGAGATTGCCCGTGCCTGTGAACGAGCTGCCCCACGGCTGGAAGGCTTGGGGGAAATCGTACTTCGTCAGAGCAGCAAAGACAAATAACAGCTTTGTCGTGGATGGTGACACACTGAAAGAATGCCCACTCAAACATAGAGCATGGAACAGCTTTCTTGTGGAGGATCATGGGTTCGGGGTATTTCACACTAGTGTCTGGCTCAAGGTTAGAGAAGATTATTCATTAGAGTGTGATCCAGCCGTTATTGGAACAGCTGTTAAGGGAAAGGAGGCTGTACACAGTGATCTAGGCTACTGGATTGAGAGTGAGAAGAATGACACATGGAGGCTGAAGAGGGCCCATCTGATCGAGATGAAAACATGTGAATGGCCAAAGTCCCACACATTGTGGACAGATGGAATAGAAGAGAGTGATCTGATCATACCCAAGTCTTTAGCTGGGCCACTCAGCCATCACAATACCAGAGAGGGCTACAGGACCCAAATGAAAGGGCCATGGCACAGTGAAGAGCTTGAAATTCGGTTTGAGGAATGCCCAGGCACTAAGGTCCACGTGGAGGAAACATGTGGAACAAGAGGACCATCTCTGAGATCAACCACTGCAAGCGGAAGGGTGATCGAGGAATGGTGCTGCAGGGAGTGCACAATGCCCCCACTGTCGTTCTGGGCTAAAGATGGCTGTTGGTATGGAATGGAGATAAGGCCCAGGAAAGAACCAGAAAGCAACTTAGTAAGGTCAATGGTGACTGCAGGATCAACTGATCACATGGATCACTTCTCCCTTGGAGTGCTTGTGATTCTGCTCATGGTGCAGGAAGGGCTGAAGAAGAGAATGACCACAAAGATCATCATAAGCACATCAATGGCAGTGCTGGTAGCTATGATCCTGGGAGGATTTTCAATGAGTGACCTGGCTAAGCTTGCAATCTTGATGGGTGCCACCTTCGCGGAAATGAACACTGGAGGAGATGTAGCTCATCTGGCGCTGATAGCGGCATTCAAAGTCAGACCAGCGTTGCTGGTATCCTTCATCTTCAGAGCTAATTGGACACCCCGTGAAAGCATGCTGCTGGCCTTGGCCTCGTGTCTTTTGCAAACTGCGATCTCCGCCTTGGAGGGCGACCTGATGGTTCTCATCAATGGTTTTGCTTTGGCCTGGTTGGCAATACGAGCGATGGTTGTTCCACGCACTGACAACATCACCTTGGCAATCCTGGCTGCTCTGACACCACTGGCCCGGGGCACACTGCTTGTGGCGTGGAGAGCAGGCCTTGCTACTTGCGGGGGGTTTATGCTCCTCTCTCTGAAGGGAAAAGGCAGTGTGAAGAAGAACTTACCATTTGTCATGGCCCTGGGACTAACCGCTGTGAGGCTGGTCGACCCCATCAACGTGGTGGGACTGCTGTTGCTCACAAGGAGTGGGAAGCGGAGCTGGCCCCCTAGCGAAGTACTCACAGCTGTTGGCCTGATATGCGCATTGGCTGGAGGGTTCGCCAAGGCAGATATAGAGATGGCTGGGCCCATGGCCGCGGTTGGTCTGCTAATTGTCAGTTACGTGGTCTCAGGAAAGAGTGTGGACATGTACATTGAAAGAGCAGGTGACATCACATGGGAAAAAGATGCGGAAGTCACTGGAAACAGTCCCCGGCTCGATGTGGCGCTAGATGAGAGTGGTGATTTCTCCCTGGTGGAGGATGACGGTCCCCCCATGAGAGAGATCATACTCAAGGTGGTCCTGATGACCATCTGTGGCATGAACCCAATAGCCATACCCTTTGCAGCTGGAGCGTGGTACGTATACGTGAAGACTGGAAAAAGGAGTGGTGCGCTATGGGATGTGCCTGCTCCCAAGGAAGTAAAAAAGGGGGAGACCACAGATGGAGTGTACAGAGTAATGACTCGTAGACTGCTAGGTTCAACACAAGTTGGAGTGGGAGTTATGCAAGAGGGGGTCTTTCACACTATGTGGCACGTCACAAAAGGATCCGCGCTGAGAAGCGGTGAAGGGAGACTTGATCCATACTGGGGAGATGTCAAGCAGGATCTGGTGTCATACTGTGGTCCATGGAAGCTAGATGCCGCCTGGGACGGGCACAGCGAGGTGCAGCTCTTGGCCGTGCCCCCCGGAGAGAGAGCGAGGAACATCCAGACTCTGCCCGGAATATTTAAGACAAAGGATGGGGACATTGGAGCGGTTGCGCTGGATTACCCAGCAGGAACTTCAGGATCTCCAATCCTAGACAAGTGTGGGAGAGTGATAGGACTTTATGGCAATGGGGTCGTGATCAAAAATGGGAGTTATGTTAGTGCCATCACCCAAGGGAGGAGGGAGGAAGAGACTCCTGTTGAGTGCTTCGAGCCTTCGATGCTGAAGAAGAAGCAGCTAACTGTCTTAGACTTGCATCCTGGAGCTGGGAAAACCAGGAGAGTTCTTCCTGAAATAGTCCGTGAAGCCATAAAAACAAGACTCCGTACTGTGATCTTAGCTCCAACCAGGGTTGTCGCTGCTGAAATGGAGGAAGCCCTTAGAGGGCTTCCAGTGCGTTATATGACAACAGCAGTCAATGTCACCCACTCTGGAACAGAAATCGTCGACTTAATGTGCCATGCCACCTTCACTTCACGTCTACTACAGCCAATCAGAGTCCCCAACTATAATCTGTATATTATGGATGAGGCCCACTTCACAGATCCCTCAAGTATAGCAGCAAGAGGATACATTTCAACAAGGGTTGAGATGGGCGAGGCGGCTGCCATCTTCATGACCGCCACGCCACCAGGAACCCGTGACGCATTTCCGGACTCCAACTCACCAATTATGGACACCGAAGTGGAAGTCCCAGAGAGAGCCTGGAGCTCAGGCTTTGATTGGGTGACGGATCATTCTGGAAAAACAGTTTGGTTTGTTCCAAGCGTGAGGAACGGCAATGAGATCGCAGCTTGTCTGACAAAGGCTGGAAAACGGGTCATACAGCTCAGCAGAAAGACTTTTGAGACAGAGTTCCAGAAAACAAAACATCAAGAGTGGGACTTTGTCGTGACAACTGACATTTCAGAGATGGGCGCCAACTTTAAAGCTGACCGTGTCATAGATTCCAGGAGATGCCTAAAGCCGGTCATACTTGATGGCGAGAGAGTCATTCTGGCTGGACCCATGCCTGTCACACATGCCAGCGCTGCCCAGAGGAGGGGGCGCATAGGCAGGAATCCCAATAAACCTGGAGATGAGTATCTGTATGGAGGTGGGTGCGCAGAGACTGACGAAGACCATGCACACTGGCTTGAAGCAAGAATGCTCCTTGACAATATTTACCTCCAAGATGGCCTCATAGCCTCGCTCTATCGACCTGAGGCCGACAAAGTAGCAGCCATTGAGGGAGAGTTCAAGCTTAGGACGGAGCAAAGGAAGACCTTTGTGGAACTCATGAAAAGAGGAGATCTTCCTGTTTGGCTGGCCTATCAGGTTGCATCTGCCGGAATAACCTACACAGATAGAAGATGGTGCTTTGATGGCACGACCAACAACACCATAATGGAAGACAGTGTGCCGGCAGAGGTGTGGACCAGACACGGAGAGAAAAGAGTGCTCAAACCGAGGTGGATGGACGCCAGAGTTTGTTCAGATCATGCGGCCCTGAAGTCATTCAAGGAGTTTGCCGCTGGGAAAAGAGGAGCGGCTTTTGGAGTGATGGAAGCCCTGGGAACACTGCCAGGACACATGACAGAGAGATTCCAGGAAGCCATTGACAACCTCGCTGTGCTCATGCGGGCAGAGACTGGAAGCAGGCCTTACAAAGCCGCGGCGGCCCAATTGCCGGAGACCCTAGAGACCATTATGCTTTTGGGGTTGCTGGGAACAGTCTCGTTGGGAATCTTTTTCGTCTTGATGAGGAACAAGGGCATAGGGAAGATGGGCTTTGGAATGGTGACTCTTGGGGCCAGCGCATGGCTCATGTGGCTCTCGGAAATTGAGCCAGCCAGAATTGCATGTGTCCTCATTGTTGTGTTCCTATTGCTGGTGGTGCTCATACCTGAGCCAGAAAAGCAAAGATCTCCCCAGGACAACCAAATGGCAATCATCATCATGGTAGCAGTAGGTCTTCTGGGCTTGATTACCGCCAATGAACTCGGATGGTTGGAGAGAACAAAGAGTGACCTAAGCCATCTAATGGGAAGGAGAGAGGAGGGGGCAACCATAGGATTCTCAATGGACATTGACCTGCGGCCAGCCTCAGCTTGGGCCATCTATGCTGCCTTGACAACTTTCATTACCCCAGCCGTCCAACATGCAGTGACCACTTCATACAACAACTACTCCTTAATGGCGATGGCCACGCAAGCTGGAGTGTTGTTTGGTATGGGCAAAGGGATGCCATTCTACGCATGGG-ACTTTGGAGTCCCGCTGCTAATGATAGGTTGCTACTCACAATTAACACCCCTGACCCTAATAGTGGCCATCATTTTGCTCGTGGCGCACTACATGTACTTGATCCCAGGGCTGCAGGCAGCAGCTGCGCGTGCTGCCCAGAAGAGAACGGCAGCTGGCATCATGAAGAACCCTGTTGTGGATGGAATAGTGGTGACTGACATTGACACAATGACAATTGACCCCCAAGTGGAGAAAAAGATGGGACAGGTGCTACTCATAGCAGTAGCCGTCTCCAGCGCCATACTGTCGCGGACCGCCTGGGGGTGGGGGGAGGCTGGGGCCCTGATCACAGCCGCAACTTCCACTTTGTGGGAAGGCTCTCCGAACAAGTACTGGAACTCCTCTACAGCCACTTCACTGTGTAACATTTTTAGGGGAAGTTACTTGGCTGGAGCTTCTCTAATCTACACAGTAACAAGAAACGCTGGCTTGGTCAAGAGACGTGGGGGTGGAACAGGAGAGACCCTGGGAGAGAAATGGAAGGCCCGCTTGAACCAGATGTCGGCCCTGGAGTTCTACTCCTACAAAAAGTCAGGCATCACCGAGGTGTGCAGAGAAGAGGCCCGCCGCGCCCTCAAGGACGGTGTGGCAACGGGAGGCCATGCTGTGTCCCGAGGAAGTGCAAAGCTGAGATGGTTGGTGGAGCGGGGATACCTGCAGCCCTATGGAAAGGTCATTGATCTTGGATGTGGCAGAGGGGGCTGGAGTTACTACGCCGCCACCATCCGCAAAGTTCAAGAAGTGAAAGGATACACAAAAGGAGGCCCTGGTCATGAAGAACCCGTGTTGGTGCAAAGCTATGGGTGGAACATAGTCCGTCTTAAGAGTGGGGTGGACGTCTTTCATATGGCGGCTGAGCCGTGTGACACGTTGCTGTGTGACATAGGTGAGTCATCATCTAGTCCTGAAGTGGAAGAAGCACGGACGCTCAGAGTCCTCTCCATGGTGGGGGATTGGCTTGAAAAAAGACCAGGAGCCTTTTGTATAAAAGTGTTGTGCCCATACACCAGCACTATGATGGAAACCCTGGAGCGACTGCAGCGTAGGTATGGGGGAGGACTGGTCAGAGTGCCACTCTCCCGCAACTCTACACATGAGATGTACTGGGTCTCTGGAGCGAAAAGCAACACCATAAAAAGTGTGTCCACCACGAGCCAGCTCCTCTTGGGGCGCATGGACGGGCCTAGGAGGCCAGTGAAATATGAGGAGGATGTGAATCTCGGCTCTGGCACGCGGGCTGTGGTAAGCTGCGCTGAAGCTCCCAACATGAAGATCATTGGTAACCGCATTGAAAGGATCCGCAGTGAGCACGCGGAAACGTGGTTCTTTGACGAGAACCACCCATATAGGACATGGGCTTACCATGGAAGCTATGAGGCCCCCACACAAGGGTCAGCGTCCTCTCTAATAAACGGGGTTGTCAGGCTCCTGTCAAAACCCTGGGATGTGGTGACTGGAGTCACAGGAATAGCCATGACCGACACCACACCGTATGGTCAGCAAAGAGTTTTCAAGGAAAAAGTGGACACTAGGGTGCCAGACCCCCAAGAAGGCACTCGTCAGGTTATGAGCATGGTCTCTTCCTGGTTGTGGAAAGAGCTAGGCAAACACAAACGGCCACGAGTCTGTACCAAAGAAGAGTTCATCAACAAGGTGCGTAGCAATGCAGCATTAGGGGCAATATTTGAAGAGGAAAAAGAGTGGAAGACTGCAGTGGAAGCTGTGAACGATCCAAGGTTCTGGGCTCTAGTGGACAAGGAAAGAGAGCACCACCTGAGAGGAGAGTGCCAGAGTTGTGTGTACAACATGATGGGAAAAAGAGAAAAGAAACAAGGGGAATTTGGAAAGGCCAAGGGCAGCCGCGCCATCTGGTATATGTGGCTAGGGGCTAGATTTCTAGAGTTCGAAGCCCTTGGATTCTTGAACGAGGATCACTGGATGGGGAGAGAGAACTCAGGAGGTGGTGTTGAAGGGCTGGGATTACAAAGACTCGGATATGTCCTAGAAGAGATGAGTCGCATACCAGGAGGAAGGATGTATGCAGATGACACTGCTGGCTGGGACACCCGCATTAGCAGGTTTGATCTGGAGAATGAAGCTCTAATCACCAACCAAATGGAGAAAGGGCACAGGGCCTTGGCATTGGCCATAATCAAGTACACATACCAAAACAAAGTGGTAAAGGTCCTTAGACCAGCTGAAAAAGGGAAAACAGTTATGGACATTATTTCGAGACAAGACCAAAGGGGGAGCGGACAAGTTGTCACTTACGCTCTTAACACATTTACCAACCTAGTGGTGCAACTCATTCGGAATATGGAGGCTGAGGAAGTTCTAGAGATGCAAGACTTGTGGCTGCTGCGGAGGTCAGAGAAAGTGACCAACTGGTTGCAGAGCAACGGATGGGATAGGCTCAAACGAATGGCAGTCAGTGGAGATGATTGCGTTGTGAAGCCAATTGATGATAGGTTTGCACATGCCCTCAGGTTCTTGAATGATATGGGAAAAGTTAGGAAGGACACACAAGAGTGGAAACCCTCAACTGGATGGGACAACTGGGAAGAAGTTCCGTTTTGCTCCCACCACTTCAACAAGCTCCATCTCAAGGACGGGAGGTCCATTGTGGTTCCCTGCCGCCACCAAGATGAACTGATTGGCCGGGCCCGCGTCTCTCCAGGGGCGGGATGGAGCATCCGGGAGACTGCTTGCCTAGCAAAATCATATGCGCAAATGTGGCAGCTCCTTTATTTCCACAGAAGGGACCTCCGACTGATGGCCAATGCCATTTGTTCATCTGTGCCAGTTGACTGGGTTCCAACTGGGAGAACTACCTGGTCAATCCATGGAAAGGGAGAATGGATGACCACTGAAGACATGCTTGTGGTGTGGAACAGAGTGTGGATTGAGGAGAACGACCACATGGAAGACAAGACCCCAGTTGCGAAATGGACAGACATTCCCTATTTGGGAAAAAGGGAAGACTTGTGGTGTGGATCTCTCATAGGGCACAGACCGCGCACCACCTGGGCTGAGAACATTAAAAACACAGTCAACATGGTGCGCAGGATCATAGGTGATGAAGAAAAGTACATGGACTACCTATCCACCCAAGTTCGCTACTTGGGTGAAGAAGGGTCTACACCTGGAGTGCTGTAA

>ENA|AMX81918|AMX81918.1 Zika virus polyprotein

ATGAAAAACCC-AAAAAAGAAATCCGGAGGATTCCGGATTGTCAATATGCTAAAACGCGGAGTAGCCCGTGTGAGCCCCTTTGGGGGCTTGAAGAGGCTGCCAGCCGGACTTCTGCTGGGTCATGGGCCCATCAGGATGGTCTTGGCGATTCTAGCCTTTTTGAGATTCACGGCAATCAAGCCATCACTGGGTCTCATCAATAGATGGGGTTCAGTGGGGAAAAAAGAGGCTATGGAAATAATAAAGAAGTTCAAGAAAGATCTGGCTGCCATGCTGAGAATAATCAATGCTAGGAAGGAGAAGAAGAGACGAGGCGCAGATACTAGTGTCGGAATTGTTGGCCTCCTGCTGACCACAGCTATGGCAGCGGAGGTCACTAGACGTGGGAGTGCATACTATATGTACTTGGACAGAAACGATGCTGGGGAGGCCATATCTTTTCCAACCACATTGGGGATGAATAAGTGTTATATACAGATCATGGATCTTGGACACATGTGTGATGCCACCATGAGCTATGAATGCCCTATGCTGGATGAGGGGGTGGAACCAGATGACGTCGATTGTTGGTGCAACACGACGTCAACTTGGGTTGTGTACGGAACCTGCCATCACAAAAAAGGTGAAGCACGGAGATCTAGAAGAGCTGTGACGCTCCCTTCCCATTCCACTAGGAAGCTGCAAACGCGGTCGCAAACCTGGTTGGAATCAAGAGAATACACAAAGCACTTGATTAGAGTCGAAAATTGGATATTCAGGAACCCTGGCTTCGCGTTAGCAGCAGCTGCCATCGCTTGGCTTTTGGGAAGCTCAACGAGCCAAAAAGTCATATACTTGGTCATGATACTGCTGATTGCCCCGGCATACAGCATCAGGTGCATAGGAGTCAGCAATAGGGACTTTGTGGAAGGTATGTCAGGTGGGACTTGGGTCGATGTTGTCTTGGAACATGGAGGTTGTGTCACCGTAATGGCACAGGACAAACCGACTGTCGACATAGAGCTGGTTACAACAACAGTCAGCAACATGGCGGAGGTAAGATCCTACTGCTATGAGGCATCAATATCAGACATGGCTTCGGACAGCCGCTGCCCAACACAAGGTGAAGCCTACCTTGACAAGCAATCAGACACTCAATATGTCTGCAAAAGAACGTTAGTGGACAGAGGCTGGGGAAATGGATGTGGACTTTTTGGCAAAGGGAGCCTGGTGACATGCGCTAAGTTTGCATGCTCCAAGAAAATGACCGGGAAGAGCATCCAGCCAGAGAATCTGGAGTACCGGATAATGCTGTCAGTTCATGGCTCCCAGCACAGTGGGATGATCGTTAATGACACAGGACATGAAACTGATGAGAATAGAGCGAAAGTTGAGATAACGCCCAATTCACCAAGAGCCGAAGCCACCCTGGGGGGTTTTGGAAGCCTAGGACTTGATTGTGAACCGAGGACAGGCCTTGACTTTTCAGATTTGTATTACTTGACTATGAATAACAAGCACTGGTTGGTTCACAAGGAGTGGTTCCACGACATTCCATTACCTTGGCACGCAGGGGCAGACACCGGAACTCCACACTGGAACAACAAAGAAGCACTGGTAGAGTTCAAGGACGCACATGCCAAAAGGCAAACTGTCGTGGTTCTAGGGAGTCAAGAAGGAGCAGTTCACACGGCCCTTGCTGGAGCTCTGGAGGCTGAGATGGATGGAGCAAAGGGAAGGCTGTCCTCTGGCCACTTGAAATGTCGCCTGAAAATGGATAAACTTAGATTGAAGGGCGTGTCATACTCCTTGTGTACTGCAGCGTTCACATTCACCAAGATCCCGGCTGAAACACTGCACGGGACAGTCACAGTGGAGGTACAGTACGCAGGGACAGATGGACCTTGCAAGGTTCCAGCTCAGATGGCGGTGGACATGCAAACTCTGACCCCAGTTGGGAGGTTGATAACCGCTAACCCCGTAATCACTGAAAGCACTGAGAACTCTAAGATGATGCTGGAACTTGATCCACCATTTGGGGACTCTTACATTGTCATAGGAGTCGGGGAGAAGAAGATCACCCACCACTGGCACAGGAGTGGCAGCACCATTGGAAAAGCATTTGAAGCCACTGTGAGAGGTGCCAAGAGAATGGCAGTCTTGGGAGACACAGCCTGGGACTTTGGATCAGTTGGAGGCGCTCTCAACTCATTGGGCAAGGGCATCCATCAAATTTTTGGAGCAGCTTTCAAATCATTGTTTGGAGGAATGTCCTGGTTCTCACAAATTCTCATTGGAACGTTGCTGATGTGGTTGGGTCTGAACACAAAGAATGGATCTATTTCCCTTATGTGCTTGGCCTTAGGGGGAGTGTTGATCTTCTTATCCACAGCCGTCTCTGCTGATGTGGGGTGCTCGGTGGACTTCTCAAAGAAGGAGACGAGATGCGGTACAGGGGTGTTTGTCTATAACGACGTTGAAGCCTGGAGGGACAGGTACAAGTACCATCCTGACTCCCCCCGTAGATTGGCAGCAGCAGTCAAGCAAGCCTGGGAAGATGGTATCTGCGGGATCTCCTCTGTTTCAAGAATGGAAAACATCATGTGGAGATCAGTAGAAGGGGAGCTCAACGCAATCCTGGAAGAGAATGGAGTTCAACTGACGGTCGTTGTGGGATCTGTAAAAAACCCCATGTGGAGAGGTCCACAGAGATTGCCCGTGCCTGTGAACGAGCTGCCCCACGGCTGGAAGGCTTGGGGGAAATCGTACTTCGTCAGAGCAGCAAAGACAAATAACAGCTTTGTCGTGGATGGTGACACACTGAAGGAATGCCCACTCAAACATAGAGCATGGAACAGCTTTCTTGTGGAGGATCATGGGTTCGGGGTATTTCACACTAGTGTCTGGCTCAAGGTTAGAGAAGATTATTCATTAGAGTGTGATCCAGCCGTTATTGGAACAGCTGTTAAGGGAAAGGAGGCTGTACACAGTGATCTAGGCTACTGGATTGAGAGTGAGAAGAATGACACATGGAGGCTGAAGAGGGCCCATCTGATCGAGATGAAAACATGTGAATGGCCAAAGTCCCACACATTGTGGACAGATGGAATAGAAGAGAGTGATCTGATCATACCCAAGTCTTTAGCTGGGCCACTCAGCCATCACAATACCAGAGAGGGCTACAGGACCCAAATGAAAGGGCCATGGCACAGTGAAGAGCTTGAAATTCGGTTTGAGGAATGCCCAGGCACTAAGGTCCACGTGGAGGAAACATGTGGAACAAGAGGACCATCTCTGAGATCAACCACTGCAAGCGGAAGGGTGATCGAGGAATGGTGCTGCAGGGAGTGCACAATGCCCCCACTGTCGTTCCGGGCTAAAGATGGCTGTTGGTATGGAATGGAGATAAGGCCCAGGAAAGAACCAGAAAGCAACTTAGTAAGGTCAATGGTGACTGCAGGATCAACTGATCACATGGACCACTTCTCCCTTGGAGTGCTTGTGATCCTGCTCATGGTGCAGGAAGGGCTGAAGAAGAGAATGACCACAAAGATCATCATAAGCACATCAATGGCAGTGCTGGTAGCTATGATCCTGGGAGGATTTTCAATGAGTGACCTGGCTAAGCTTGCAATTTTGATGGGTGCCACCTTCGCGGAAATGAACACTGGAGGAGATGTAGCTCATCTGGCGCTGATAGCGGCATTCAAAGTCAGACCAGCGTTGCTGGTATCTTTCATCTTCAGAGCTAATTGGACACCCCGTGAAAGCATGCTGCTGGCCTTGGCCTCGTGTCTTTTGCAAACTGCGATCTCCGCCTTGGAAGGCGACCTGATGGTTCTCATCAATGGTTTTGCTTTGGCCTGGTTGGCAGTACGAGCGATGGTTGTTCCACGCACTGATAACATCACCCTAGCAATCCTGGCTGCTCTGACACCACTGGCCCGGGGCACACTGCTTGTGGCGTGGAGAGCAGGCCTTGCTACTTGCGGGGGGTTTATGCTCCTCTCTCTGAAGGGAAAAGGCAGTGTGAAGAAGAACTTACCATTTGTCATGGCCCTGGGACTAACCGCTGTGAGGCTGGTCGACCCCATCAACGTGGTGGGACTGCTGTTGCTCACAAGGAGTGGGAAGCGGAGCTGGCCCCCTAGCGAAGTACTCACAGCTGTTGGCCTGATATGCGCATTGGCTGGAGGGTTCGCCAAGGCAGATATAGAGATGGCTGGGCCCATGGCCGCGGTCGGTCTGCTAATTGTCAGTTACGTGGTCTCAGGAAAGAGTGTGGACATGTACATTGAAAGAGCAGGTGACATCACATGGGAAAAAGATGCGGAAGTCACTGGAAACAGTCCCCGGCTCGATGTGGCGCTAGATGAGAGTGGTGATTTCTCCCTGGTGGAGGATGACGGTCCCCCCATGAGAGAGATCATACTCAAGGTGGTCCTGATGACCATCTGTGGCATGAACCCAATAGCCATACCCTTTGCAGCTGGAGCGTGGTACGTATACGTGAAGACTGGAAAAAGGAGTGGTGCTCTATGGGATGTGCCTGCTCCCAAGGAAGTAAAAAAGGGGGAGACCACAGATGGAGTGTACAGAGTAATGACTCGTAGACTGCTAGGTTCAACACAAGTTGGAGTGGGAGTTATGCAAGAGGGGGTCTTTCACACTATGTGGCACGTCACAAAAGGATCCGCGCTGAGAAGCGGTGAAGGGAGACTTGATCCATACTGGGGAGATGTCAAGCAGGATCTGGTGTCATACTGTGGTCCATGGAAGCTAGATGCCGCCTGGGACGGGCACAGCGAGGTGCAGCTCTTGGCCGTGCCCCCCGGAGAGAGAGCGAGGAACATCCAGACTCTGCCCGGAATATTTAAGACAAAGGATGGGGACATTGGAGCGGTTGCGCTGGATTACCCAGCAGGAACTTCAGGATCTCCAATCCTAGACAAGTGTGGGAGAGTGATAGGACTTTATGGCAATGGGGTCGTGATCAAAAATGGGAGTTATGTTAGTGCCATCACCCAAGGGAGGAGGGAGGAAGAGACTCCTGTTGAGTGCTTCGAGCCTTCGATGCTGAAGAAGAAGCAGCTAACTGTCTTAGACTTGCATCCTGGAGCTGGGAAAACCAGGAGAGTTCTTCCTGAAATAGTCCGTGAAGCCATAAAAACAAGACTCCGTACTGTGATCTTAGCTCCAACCAGGGTTGTCGCTGCTGAAATGGAGGAGGCCCTTAGAGGGCTTCCAGTGCGTTATATGACAACAGCAGTCAATGTCACCCACTCTGGAACAGAAATCGTCGACTTAATGTGCCATGCCACCTTCACTTCACGTCTACTACAGCCAATTAGAGTCCCCAACTATAATCTGTATATTATGGATGAGGCCCACTTTACAGATCCCTCAAGTATAGCAGCAAGAGGATACATTTCAACAAGGGTTGAGATGGGCGAGGCGGCTGCCATCTTCATGACCGCCACGCCACCAGGAACCCGTGACGCATTTCCGGACTCCAACTCACCAATTATGGACACCGAAGTGGAAGTCCCAGAGAGAGCCTGGAGCTCAGGCTTTGATTGGGTGACGGATCATTCTGGAAAAACAGTTTGGTTTGTTCCAAGCGTGAGGAACGGCAATGAGATCGCAGCTTGTCTGACAAAGGCTGGAAAACGGGTCATACAGCTCAGCAGAAAGACTTTTGAGACAGAGTTCCAGAAAACAAAACATCAAGAGTGGGACTTTGTCGTGACAACTGACATTTCAGAGATGGGCGCCAACTTTAAAGCTGACCGTGTCATAGATTCCAGGAGATGCCTAAAGCCGGTCATACTTGATGGCGAGAGAGTCATTCTGGCTGGACCCATGCCTGTCACACATGCCAGCGCTGCCCAGAGGAGGGGGCGCATAGGCAGGAATCCCAACAAACCTGGAGATGAGTATCTGTATGGAGGTGGGTGCGCAGAGACTGACGAAGACCATGCACATTGGCTTGAAGCAAGAATGCTCCTTGACAATATTTACCTCCAAGATGGCCTCATAGCCTCGCTCTATCGACCTGAGGCCGACAAAGTAGCAGCCATTGAGGGAGAGTTCAAGCTTAGGACGGAGCAAAGGAAGACCTTTGTGGAACTCATGAAAAGAGGAGATCTTCCTGTTTGGCTGGCCTATCAGGTTGCATCTGCCGGAATAACCTACACAGATAGAAGATGGTGCTTTGATGGCACGACCAACAACACCATAATGGAAGACAGTGTGCCGGCAGAGGTGTGGACCAGACACGGAGAGAAAAGAGTGCTCAAACCGAGGTGGATGGACGCCAGAGTTTGTTCAGATCATGCGGCCCTGAAGTCATTCAAGGAGTTTGCCGCTGGGAAAAGAGGAGCGGCTTTTGGAGTGATGGAAGCCCTGGGAACACTGCCAGGACACATGACAGAGAGATTCCAGGAAGCCATTGACAACCTCGCTGTGCTCATGCGGGCAGAGACTGGAAGCAGGCCTTACAAAGCCGCGGCAGCCCAATTGCCGGAGACCCTAGAGACCATTATGCTTTTGGGGTTGCTGGGAACAGTCTCGCTGGGAATCTTCTTCGTCTTGATGAGGAACAAGGGCATAGGGAAGATGGGCTTTGGAATGGTGACTCTTGGGGCCAGCGCATGGCTCATGTGGCTCTCGGAAATTGAGCCAGCCAGAATTGCATGTGTCCTCATTGTTGTGTTCCTATTGCTGGTGGTGCTCATACCTGAGCCAGAAAAGCAAAGATCTCCCCAGGACAACCAAATGGCAATCATCATTATGGTAGCAGTAGGTCTTCTGGGCTTGATTACCGCCAATGAACTCGGATGGTTGGAGAGAACAAAGAGTGACCTAAGCCATCTAATGGGAAGGAGAGAGGAGGGGGCAACCATAGGATTCTCAATGGACATTGACCTGCGGCCAGCCTCAGCTTGGGCCATCTATGCTGCCTTGACAACTTTCATTACCCCAGCCGTCCAACATGCAGTGACCACCTCATACAACAACTACTCCTTAATGGCGATGGCCACGCAAGCTGGAGTGTTGTTTGGTATGGGCAAAGGGATGCCATTCTACGCATGGG-ACTTTGGAGTCCCGCTGCTAATGATAGGTTGCTACTCACAATTAACACCCCTGACCCTAATAGTGGCCATCATTTTGCTCGTGGCGCACTACATGTACTTGATCCCAGGGCTGCAGGCAGCAGCTGCGCGTGCTGCCCAGAAGAGAACGGCAGCTGGCATCATGAAGAACCCTGTTGTGGATGGAATAGTGGTGACTGACATTGACACAATGACAATTGACCCCCAAGTGGAGAAAAAGATGGGACAGGTGCTACTCATAGCAGTAGCCGTCTCCAGCGCCATACTGTCGCGGACCGCCTGGGGGTGGGGGGAGGCTGGGGCCCTGATCACAGCCGCAACTTCCACTTTGTGGGAAGGCTCTCCGAACAAGTACTGGAACTCCTCTACAGCCACTTCACTGTGTAACATTTTTAGGGGAAGTTACTTGGCTGGAGCTTCTCTAATCTACACAGTAACAAGAAACGCTGGCTTGGTCAAGAGACGTGGGGGTGGAACAGGAGAGACCCTGGGAGAGAAATGGAAGGCCCGCTTGAACCAGATGTCGGCCCTGGAGTTCTACTCCTACAAAAAGTCAGGCATCACCGAGGTGTGCAGAGAAGAGGCCCGCCGCGCCCTCAAGGACGGTGTGGCAACGGGAGGCCATGCTGTGTCCCGAGGAAGTGCAAAGCTGAGATGGTTGGTGGATCGGGGATACCTGCAGCCCTATGGAAAGGTCATTGATCTTGGATGTGGCAGAGGGGGCTGGAGTTACTACGCCGCCACCATCCGCAAAGTTCAAGAAGTGAAAGGATACACAAAAGGAGGCCCTGGTCATGAAGAACCCGTGTTGGTGCAAAGCTATGGGTGGAACATAGTCCGTCTTAAGAGTGGGGTGGACGTCTTTCATATGGCGGCTGAGCCGTGTGACACGTTGCTGTGTGACATAGGTGAGTCATCATCTAGTCCTGAAGTGGAAGAAGCACGGACGCTCAGAGTCCTCTCCATGGTGGGGGATTGGCTTGAAAAAAGACCAGGAGCCTTTTGCATAAAAGTGTTGTGCCCATACACCAGCACTATGATGGAAACCCTGGAGCGACTGCAGCGTAGGTATGGGGGAGGACTGGTCAGAGTGCCACTCTCCCGCAACTCTACACATGAGATGTACTGGGTCTCTGGAGCGAAAAGCAACACCATAAAAAGTGTGTCCACCACGAGCCAGCTCCTCTTGGGGCGCATGGACGGGCCTAGGAGGCCAGTGAAATATGAGGAGGATGTGAATCTCGGCTCTGGCACGCGGGCTGTGGCAAGCTGCGCTGAAGCTCCCAACATGAAGATCATTGGTAACCGCATTGAAAGGATCCGCAGTGAGCACGCGGAAACGTGGTTCTTTGACGAGAACCACCCATATAGGACATGGGCTTACCATGGAAGCTATGAGGCCCCCACACAAGGGTCAGCGTCCTCTCTAATAAACGGGGTTGTCAGGCTCCTGTCAAAACCCTGGGATGTGGTGACTGGAGTCACAGGAATAGCCATGACCGACACCACACCGTATGGTCAGCAAAGAGTTTTCAAGGAAAAAGTGGACACTAGGGTGCCAGACCCCCAAGAAGGCACTCGTCAGGTTATGAGCATGGTCTCTTCCTGGTTGTGGAAAGAGCTAGGCAAACACAAACGGCCACGAGTCTGTACCAAAGAAGAGTTCATCAACAAGGTTCGTAGCAATGCAGCATTAGGGGCAATATTTGAAGAGGAAAAAGAGTGGAAGACTGCAGTGGAAGCTGTGAACGATCCAAGGTTCTGGGCTCTAGTGGACAAGGAAAGAGAGCACCACCTGAGAGGAGAGTGCCAGAGTTGTGTGTACAACATGATGGGAAAAAGAGAAAAGAAACAAGGGGAATTTGGAAAGGCCAAGGGCAGCCGCGCCATCTGGTATATGTGGCTGGGGGCTAGATTTCTAGAGTTCGAAGCCCTTGGATTCTTGAACGAGGATCACTGGATGGGGAGAGAGAACTCAGGAGGTGGTGTTGAAGGGCTGGGATTACAAAGACTCGGATATGTCCTAGAAGAGATGAGTCGTATACCAGGAGGAAGGATGTATGCAGATGACACTGCTGGCTGGGACACCCGCATTAGCAGGTTTGATCTGGAGAATGAAGCTCTAATCACCAACCAAATGGAGAAAGGGCACAGGGCCTTGGCATTGGCCATAATCAAGTACACATACCAAAACAAAGTGGTAAAGGTCCTTAGACCAGCTGAAAAAGGGAAAACAGTTATGGACATTATTTCGAGACAAGACCAAAGGGGGAGCGGACAAGTTGTCACTTACGCTCTTAACACATTTACCAACCTAGTGGTGCAACTCATTCGGAATATGGAGGCTGAGGAAGTTCTAGAGATGCAAGACTTGTGGCTGCTGCGGAGGTCAGAGAAAGTGACCAACTGGTTGCAGAGCAACGGATGGGATAGGCTCAAACGAATGGCAGTCAGTGGAGATGATTGCGTTGTGAAGCCAATTGATGATAGGTTTGCACATGCCCTCAGGTTCTTGAATGATATGGGAAAAGTTAGGAAGGACACACAAGAGTGGAAACCCTCAACTGGATGGGACAACTGGGAAGAAGTTCCGTTTTGCTCCCACCATTTCAACAAGCTCCATCTCAAGGACGGGAGGTCCATTGTGGTTCCCTGCCGCCACCAAGATGAACTGATTGGCCGGGCCCGCGTCTCTCCAGGGGCGGGATGGAGCATCCGGGAGACTGCTTGCCTAGCAAAATCATATGCGCAAATGTGGCAGCTCCTTTATTTCCACAGAAGGGACCTCCGACTGATGGCCAATGCCATTTGTTCATCTGTGCCAGTTGACTGGGTTCCAACTGGGAGAACTACCTGGTCAATCCATGGAAAGGGAGAATGGATGACCACTGAAGACATGCTTGTGGTGTGGAACAGAGTGTGGATTGAGGAGAACGACCACATGGAAGACAAGACCCCAGTTACGAAATGGACAGACATTCCCTATTTGGGAAAAAGGGAAGACTTGTGGTGTGGATCTCTCATAGGGCACAGACCGCGCACCACCTGGGCTGAGAACATTAAAAACACAGTCAACATGGTGCGCAGGATCATAGGTGATGAAGAAAAGTACATGGATTACCTATCCACCCAAGTTCGCTACTTGGGTGAAGAAGGGTCTACACCTGGAGTGCTGTAA

>ENA|AMR39834|AMR39834.1 Zika virus polyprotein

ATGAAAAACCC-AAAGAAGAAATCCGGAGGATTCCGGATTGTCAATATGCTAAAACGCGGAGTAGCCCGTGTGAGCCCCTTTGGGGGCTTGAAGAGGCTGCCAGCCGGACTTCTGCTGGGTCATGGGCCCATCAGGATGGTCTTGGCGATTCTAGCCTTTTTGAGATTCACGGCAATCAAGCCATCACTGGGTCTCATCAATAGATGGGGTTCAGTGGGGAAAAAAGAGGCTATGGAAATAATAAAGAAGTTTAAGAAAGATCTGGCTGCCATGCTGAGAATAATCAATGCTAGGAAGGAGAAGAAGAGACGAGGCACAGATACTAGTGTCGGAATTGTTGGCCTCCTGCTGACCACAGCCATGGCAGTGGAGGTCACTAGACGTGGGAATGCATACTATATGTACTTGGACAGAAGCGATGCTGGGGAGGCCATATCTTTTCCAACCACAATGGGGATGAATAAGTGTTATATACAGATCATGGATCTTGGACACATGTGTGATGCCACCATGAGCTATGAATGCCCTATGCTGGATGAGGGGGTAGAACCAGATGACGTCGATTGTTGGTGCAACACGACGTCAACTTGGGTTGTGTACGGAACCTGCCACCACAAAAAAGGTGAAGCACGGAGATCTAGAAGAGCTGTGACGCTCCCCTCCCATTCCACTAGGAAGCTGCAAACGCGGTCGCAGACCTGGTTGGAATCAAGAGAATACACAAAGCACCTGATTAGAGTCGAAAATTGGATATTCAGGAACCCTGGCTTCGCGTTAGCAGCAGCTGCCATCGCTTGGCTTTTGGGAAGCTCAACGAGCCAAAAAGTCATATACTTGGTCATGATACTGCTGATTGCCCCGGCATACAGCATCAGGTGCATAGGAGTCAGCAATAGGGACTTTGTGGAAGGTATGTCAGGTGGGACTTGGGTTGATGTTGTCTTGGAACATGGAGGTTGTGTTACCGTAATGGCACAGGACAAACCGACTGTCGACATAGAGCTGGTTACAACAACAGTCAGCAACATGGCGGAGGTAAGATCCTACTGCTATGAGGCATCAATATCGGACATGGCTTCGGACAGCCGCTGCCCAACACAAGGTGAAGCCTACCTTGACAAGCAATCAGACACTCAATATGTCTGCAAAAGAACGTTAGTGGACAGAGGCTGGGGAAATGGATGTGGACTTTTTGGCAAAGGGAGCCTGGTGACATGCGCTAAGTTTGCTTGCTCTAAGAAAATGACCGGGAAGAGCATCCAGCCAGAGAATCTGGAGTACCGGATAATGCTGTCAGTTCATGGCTCCCAGCACAGTGGGATGATCGTTAATGATACAGGACATGAAACTGATGAGAATAGAGCGAAGGTTGAGATAACGCCCAATTCACCAAGAGCCGAAGCCACCCTGGGGGGTTTTGGAAGCCTAGGACTTGATTGTGAACCGAGGACAGGCCTTGACTTTTCAGATTTGTATTACTTGACTATGAATAACAAGCACTGGTTGGTTCACAAGGAGTGGTTCCACGACATTCCATTACCTTGGCATGCTGGGGCAGACACCGGAACTCCACACTGGAACAACAAAGAAGCACTGGTAGAGTTCAAGGACGCACATGCCAAAAGGCAGACTGTCGTGGTTCTAGGGAGTCAAGAAGGAGCAGTTCACACGGCCCTTGCTGGAGCTCTGGAGGCTGAGATGGATGGTGCAAAGGGAAGGCTGTCCTCTGGCCACTTGAAATGTCGCCTGAAAATGGATAAACTTAGATTGAAGGGCGTGTCATACTCCTTGTGTACCGCAGCGTTCACATTCACTAAGATCCCGGCTGAAACACTGCACGGGACAGTCACAGTGGAGGTACAGTACGCAGGGACAGATGGACCTTGCAAGGTTCCAGCTCAGATGGCGGTGGACATGCAAACTCTGACCCCAGTTGGGAGGTTGATAACCGCTAACCCTGTAATCACTGAAAGCACTGAGAACTCCAAGATGATGCTGGAACTGGATCCACCATTTGGGGACTCTTACATTGTCATAGGAGTCGGGGAAAAGAAGATCACCCACCACTGGCACAGGAGTGGCAGCACCATTGGAAAAGCATTTGAAGCCACTGTGAGAGGTGCCAAGAGAATGGCAGTCTTGGGAGACACAGCCTGGGACTTTGGATCAGTTGGGGGTGCTCTCAACTCACTGGGCAAGGGCATCCATCAAATTTTTGGAGCAGCTTTCAAATCATTGTTTGGAGGAATGTCCTGGTTCTCACAAATTCTCATTGGAACGTTGCTGGTGTGGTTGGGTCTGAATACAAAGAATGGATCTATTTCCCTTATGTGCTTGGCCTTAGGGGGAGTGTTGATCTTCTTATCCACAGCCGTCTCTGCTGATGTGGGGTGCTCGGTGGACTTCTCAAAGAAGGAAACGAGATGCGGTACAGGGGTGTTCGTCTATAACGACGTTGAAGCTTGGAGGGACAGGTACAAGTACCATCCTGACTCCCCTCGTAGATTGGCAGCAGCAGTCAAGCAAGCCTGGGAAGATGGGATCTGTGGGATCTCCTCTGTTTCAAGAATGGAAAACATCATGTGGAGATCAGTAGAAGGGGAGCTCAACGCAATCCTGGAAGAGAATGGAGTTCAACTGACGGTCGTTGTGGGATCTGTAAAAAACCCCATGTGGAGAGGTCCACAGAGATTGCCCGTGCCTGTGAACGAGCTGCCCCATGGCTGGAAGGCTTGGGGGAAATCGTACTTCGTCAGGGCAGCAAAGACAAATAACAGCTTTGTCGTGGATGGTGACACACTGAAGGAATGCCCACTCAAACATAGAGCATGGAACAGCTTTCTTGTGGAGGATCATGGGTTCGGGGTATTTCACACTAGTGTCTGGCTCAAGGTTAGAGAAGATTATTCATTAGAGTGTGATCCAGCCGTCATTGGAACAGCCGCTAAGGGAAAGGAGGCTGTGCACAGTGATCTAGGCTACTGGATTGAGAGTGAGAAGAACGACACATGGAGGCTGAAGAGGGCCCACCTGATCGAGATGAAAACATGTGAATGGCCAAAGTCCCACACATTGTGGACAGATGGAATAGAAGAAAGTGATCTGATCATACCCAAGTCTTTAGCTGGGCCACTCAGCCATCACAACACCAGAGAGGGCTACAGGACCCAAATGAAAGGGCCATGGCATAGTGAAGAGCTTGAAATTCGGTTTGAGGAATGCCCAGGCACTAAGGTCCACGTGGAGGAAACATGTGGAACAAGAGGACCATCTCTGAGATCAACCACTGCAAGCGGAAGGGTGATCGAGGAATGGTGCTGCAGGGAGTGCACAATGCCCCCACTGTCGTTCCGGGCTAAAGATGGTTGTTGGTATGGAATGGAGATAAGGCCCAGGAAAGAACCAGAAAGTAACTTAGTAAGGTCAATGGTGACTGCAGGATCAACTGATCACATGGATCACTTCTCCCTTGGAGTGCTTGTGATTCTGCTCATGGTACAGGAAGGGCTAAAGAAGAGAATGACCACAAAGATCATCATAAGCACATCAATGGCAGTGCTGGTAGCTATGATCCTGGGAGGATTTTCAATGAGTGACCTGGCTAAGCTTGCAATTTTGATGGGTGCCACCTTCGCGGAAATGAACACTGGAGGAGATGTTGCTCATCTGGCGCTGATAGCGGCATTCAAAGTCAGACCTGCGTTGCTGGTATCTTTCATTTTCAGAGCTAATTGGACACCCCGTGAGAGCATGCTGCTGGCCTTGGCCTCGTGTCTTCTGCAAACTGCGATCTCCGCCTTGGAAGGCGACCTGATGGTTCCCATCAATGGTTTTGCTTTGGCCTGGTTGGCAATACGAGCGATGGTTGTTCCACGCACTGACAACATCACCTTGGCAATCCTGGCTGCTCTGACACCACTGGCCCGGGGCACACTGCTTGTGGCGTGGAGAGCAGGCCTTGCTACTTGCGGGGGGTTCATGCTCCTTTCTCTGAAGGGGAAAGGCAGTGTGAAGAAGAACTTACCATTTGTCATGGCCCTGGGACTAACCGCTGTGAGGCTGGTCGACCCCATCAACGTGGTGGGACTGCTGTTGCTCACAAGGAGTGGGAAGCGGAGCTGGCCCCCTAGTGAAGTACTCACAGCTGTTGGCCTGATATGCGCATTGGCTGGAGGGTTCGCCAAGGCGGATATAGAGATGGCTGGGCCCATGGCCGCGGTCGGTCTGCTAATTGTCAGTTACGTGGTCTCAGGAAAGAGTGTGGACATGTACATTGAAAGAGCAGGTGACATCACATGGGAAAAAGATGCGGAAGTCACTGGAAACAGTCCCCGGCTCGATGTGGCACTAGATGAGAGTGGTGATTTCTCCCTAGTGGAGGATGATGGTCCCCCCATGAGAGAGATCATACTCAAAGTGGTCCTGATGGCCATCTGTGGCATGAACCCAATAGCCATACCCTTTGCAGCTGGAGCGTGGTACGTGTATGTGAAGACTGGAAAAAGGAGTGGTGCTCTATGGGATGTGCCTGCTCCCAAGGAAGTAAAAAAGGGGGAGACCACAGATGGAGTGTACAGAGTAATGACTCGTAGACTGCTAGGTTCAACACAAGTTGGAGTGGGAGTCATGCAAGAGGGGGTCTTCCACACTATGTGGCACGTCACAAAAGGATCCGCGCTGAGAAGCGGTGAAGGGAGACTTGATCCATACTGGGGAGATGTCAAGCAGGATCTGGTGTCATACTGTGGTCCATGGAAGCTAGATGCCGCCTGGGACGGGCACAGCGAGGTGCAGCTCTTGGCCGTGCCCCCCGGAGAGAGAGCGAGGAACATCCAGACTCTGCCCGGAATATTTAAGACAAAGGATGGGGACATTGGAGCAGTTGCGCTGGACTACCCAGCAGGAACTTCAGGATCTCCAATCCTAGATAAGTGTGGGAGAGTGATAGGACTCTATGGTAATGGGGTCGTGATCAAAAATGGGAGTTACGTTAGTGCCATCACCCAAGGGAGGAGGGAGGAAGAGACTCCTGTTGAGTGCTTCGAGCCTTCGATGCTGAAGAAGAAGCAGCTAACTGTCTTAGACTTGCATCCTGGAGCTGGGAAAACCAGGAGAGTTCTTCCTGAAATAGTCCGTGAAGCCATAAAAACAAGACTCCGCACTGTGATCTTAGCTCCAACCAGGGTTGTCGCTGCTGAAATGGAGGAAGCCCTTAGAGGGCTTCCAGTGCGTTATATGACAACAGCAGTCAATGTCACCCATTCTGGGACAGAAATCGTTGACTTAATGTGCCATGCCACCTTCACTTCACGTCTACTACAGCCAATCAGAGTCCCCAACTATAATCTGTATATTATGGATGAGGCCCACTTCACAGATCCCTCAAGTATAGCAGCAAGAGGATACATTTCAACAAGGGTTGAGATGGGCGAGGCGGCTGCCATCTTCATGACTGCCACGCCACCAGGAACCCGTGACGCATTCCCGGACTCCAACTCACCAATTATGGACACCGAAGTGGAAGTCCCAGAGAGAGCCTGGAGCTCAGGCTTTGATTGGGTGACGGATCATTCTGGAAAAACAGTTTGGTTTGTTCCAAGCGTGAGGAATGGCAATGAGATCGCAGCTTGTCTGACAAAGGCTGGAAAACGGGTCATACAGCTCAGCAGAAAGACTTTTGAGACAGAGTTCCAGAAAACAAAACATCAAGAGTGGGACTTCGTCGTGACAACTGACATTTCAGAGATGGGCGCCAACTTTAAAGCTGACCGTGTCATAGATTCCAGGAGATGCCTAAAGCCGGTCATACTTGATGGCGAGAGAGTCATTCTGGCTGGACCCATGCCTGTCACACATGCCAGCGCTGCCCAGAGGAGGGGGCGCATAGGCAGGAACCCCAACAAACCTGGAGATGAGTATCTGTATGGAGGTGGGTGCGCAGAGACTGATGAAGACCATGCACACTGGCTTGAAGCAAGAATGCTTCTTGACAACATTTACCTCCAAGATGGCCTCATAGCCTCGCTCTATCGACCTGAGGCCGACAAAGTAGCAGCTATTGAGGGAGAGTTCAAGCTTAGGACGGAGCAAAGGAAGACCTTTGTGGAACTCATGAAAAGAGGAGATCTTCCTGTTTGGCTGGCCTATCAGGTTGCATCTGCCGGAATAACCTACACAGATAGAAGATGGTGCTTTGATGGCACGACCAACAACACCATAATGGAAGACAGTGTGCCGGCAGAGGTGTGGACCAGATACGGAGAGAAAAGAGTGCTCAAACCGAGGTGGATGGACGCCAGAGTTTGTTCAGATCATGCGGCCCTGAAGTCATTCAAAGAGTTTGCCGCTGGGAAAAGAGGAGCGGCCTTTGGAGTGATGGAAGCCCTGGGAACACTGCCAGGACATATGACAGAGAGATTCCAGGAGGCCATTGACAACCTCGCTGTGCTCATGCGGGCAGAGACTGGAAGCAGGCCCTACAAAGCCGCGGCGGCCCAATTACCGGAGACCCTAGAGACTATCATGCTTTTGGGGTTGCTGGGAACAGTCTCGCTGGGAATCTTTTTCGTCTTGATGCGGAACAAGGGCATAGGGAAGATGGGCTTTGGAATGGTGACTCTTGGGGCCAGCGCATGGCTTATGTGGCTCTCGGAAATTGAGCCAGCCAGAATTGCATGTGTCCTCATTGTTGTGTTCCTATTGCTGGTGGTGCTCATACCTGAGCCAGAAAAGCAAAGATCTCCCCAGGACAACCAAATGGCAATCATCATCATGGTAGCAGTGGGTCTTCTGGGCTTGATTACCGCCAATGAACTCGGATGGTTGGAGAGAACAAAGAGTGACCTAAGCCATCTAATGGGAAGGAGAGAGGAGGGGGCAACTATAGGATTCTCAATGGACATTGACCTGCGGCCAGCCTCAGCTTGGGCTATCTATGCTGCTCTGACAACTTTCATTACCCCAGCCGTCCAACATGCAGTGACCACTTCATACAACAACTACTCCTTAATGGCGATGGCCACGCAAGCTGGAGTGTTGTTCGGTATGGGTAAAGGGATGCCATTCTATGCATGGG-ACTTTGGAGTCCCGCTGCTAATGATAGGTTGCTACTCACAATTAACACCCCTGACCCTAATAGTGGCCATCATTTTGCTCGTGGCGCACTACATGTACTTGATCCCAGGGCTGCAGGCAGCAGCTGCGCGTGCTGCCCAGAAGAGAACGGCAGCTGGCATCATGAAGAACCCTGTTGTGGATGGAATAGTGGTGACTGACATTGACACAATGACAATTGACCCCCAAGTGGAGAAAAAGATGGGACAGGTGCTACTCATAGCAGTAGCTGTCTCCAGCGCCATACTGTCGCGGACCGCCTGGGGGTGGGGTGAGGCTGGGGCCCTGATCACAGCTGCAACTTCCACTTTGTGGGAGGGCTCTCCGAACAAGTACTGGAACTCCTCCACAGCCACCTCACTGTGTAACATTTTTAGGGGAAGCTACTTGGCTGGAGCTTCTCTAATCTACACAGTAACAAGAAACGCTGGCTTGGTCAAGAGACGTGGGGGTGGAACGGGAGAGACCCTGGGAGAGAAATGGAAGGCCCGCCTGAACCAGATGTCGGCCCTGGAGTTCTACTCCTACAAAAAGTCAGGCATCACCGAGGTGTGCAGAGAAGAGGCCCGCCGCGCCCTCAAGGACGGTGTGGCAACGGGAGGCCACGCTGTGTCCCGAGGAAGTGCAAAGCTGAGATGGTTGGTGGAGAGGGGATACCTGCAGCCCTATGGAAAGGTCATTGATCTTGGATGTGGCAGAGGGGGCTGGAGTTACTATGCCGCCACCATCCGCAAAGTTCAAGAAGTGAAAGGATACACAAAAGGAGGCCCTGGTCATGAAGAACCCATGTTGGTGCAAAGCTATGGGTGGAACATAGTCCGTCTTAAGAGTGGGGTGGACGTCTTTCATATGGCGGCTGAGCCGTGTGACACGTTGCTGTGTGATATAGGTGAGTCATCATCTAGTCCTGAAGTGGAAGAAGCACGGACGCTCAGAGTCCTCTCCATGGTGGGGGATTGGCTTGAAAAAAGACCAGGAGCCTTTTGTATAAAAGTGTTGTGCCCATACACCAGCACTATGATGGAAACCCTGGAGCGACTGCAGCGTAGGTATGGGGGAGGACTGGTCAGAGTGCCACTCTCCCGCAACTCTACACATGAGATGTACTGGGTCTCTGGAGCGAAAAGCAACACCATAAAAAGTGTGTCCACCACGAGCCAGCTCCTTTTGGGGCGCATGGACGGGCCCAGGAGGCCAGTGAAATATGAAGAGGATGTGAATCTCGGCTCTGGCACGCGGGCTGTGGTAAGCTGCGCTGAAGCTCCCAACATGAAGATCATTGGTAACCGCATTGAGAGGATCCGCAGTGAGCACGCGGAAACGTGGTTCTTTGACGAGAACCACCCATATAGGACATGGGCTTACCATGGAAGCTACGAGGCCCCCACACAAGGGTCAGCGTCCTCTCTAATAAACGGGGTTGTCAGGCTCCTGTCAAAACCCTGGGATGTGGTGACTGGAGTCACAGGAATAGCCATGACCGACACCACACCGTATGGTCAGCAAAGAGTTTTCAAGGAAAAAGTGGACACTAGGGTGCCAGACCCCCAAGAAGGCACTCGTCAGGTTATGAGCATGGTCTCTTCCTGGTTGTGGAAAGAGTTAGGCAAACACAAACGGCCACGAGTCTGTACCAAAGAAGAGTTCATCAACAAGGTTCGTAGCAACGCAGCATTAGGGGCAATATTTGAAGAGGAAAAAGAGTGGAAGACTGCAGTGGAAGCTGTGAACGATCCAAGGTTCTGGGCTCTAGTGGACAAGGAAAGAGAGCACCACCTGAGAGGAGAGTGCCAGAGCTGTGTGTACAACATGATGGGAAAAAGAGAAAAGAAACAAGGGGAATTTGGAAAGGCCAAGGGCAGCCGCGCCATCTGGTACATGTGGCTAGGGGCTAGATTTCTAGAGTTCGAAGCCCTTGGATTCTTGAACGAGGATCACTGGATGGGGAGAGAGAATTCAGGAGGTGGTGTTGAAGGGCTAGGATTACAAAGACTCGGATATGTCTTAGAAGAGATGAGTCGCATACCAGGAGGAAGGATGTATGCAGATGATACTGCTGGCTGGGACACCCGCATCAGCAGGTTTGATCTGGAGAATGAAGCTCTAATCACCAACCAAATGGAGAAAGGGCACAGGGCCTTGGCATTGGCCATAATCAAGTACACATACCAAAACAAAGTGGTAAAGGTCCTTAGACCAGCTGAAAAAGGGAAGACAGTTATGGACATTATTTCAAGACAAGACCAAAGGGGGAGCGGACAAGTTGTCACTTACGCTCTTAATACATTTACCAACCTAGTGGTGCAGCTCATTCGGAATATGGAGGCTGAGGAAGTTCTAGAGATGCAAGACTTGTGGCTGCTGCGGAGGTCAGAGAAAGTGACCAACTGGTTGCAGAGCAATGGATGGGATAGGCTCAAACGAATGGCAGTCAGTGGAGATGATTGCGTTGTGAAACCAATTGATGATAGGTTTGCACATGCTCTCAGGTTCTTGAATGATATGGGAAAAGTTAGGAAGGACACACAAGAGTGGAAGCCCTCAACTGGATGGGACAACTGGGAAGAAGTTCCGTTTTGCTCCCACCACTTCAACAAGCTCCATCTCAAGGACGGGAGGTCCATTGTGGTTCCCTGCCGCCACCAAGATGAACTGATTGGCCGAGCTCGCGTCTCACCGGGGGCGGGATGGAGCATCCGGGAGACTGCTTGCCTAGCAAAATCATATGCGCAAATGTGGCAGCTCCTTTATTTCCACAGAAGGGACCTCCGACTGATGGCCAATGCCATTTGTTCATCTGTGCCAGTTGACTGGGTTCCAACTGGGAGAACTACCTGGTCAATCCATGGAAAGGGAGAATGGATGACCACTGAAGACATGCTTGTGGTGTGGAACAGAGTGTGGATTGAGGAGAACGACCACATGGAAGACAAGACCCCAGTTACGAAATGGACAGACATTCCCTATTTGGGAAAAAGGGAAGACTTGTGGTGTGGGTCTCTCATAGGGCACAGACCGCGCACCACCTGGGCTGAGAACATTAAAAACACAGTCAACATGATGCGTAGGATCATAGGTGATGAAGAAAAGTACGTGGACTACCTATCCACCCAAGTTCGCTACTTGGGCGAAGAAGGGTCCACACCTGGAGTGCTATAA

>ENA|AMR39836|AMR39836.1 Zika virus polyprotein

ATGAAAAACCC-AAAGAAGAAATCCGGAGGATTCCGGATTGTCAATATGCTAAAACGCGGAGTAGCCCGTGTAAACCCCTTGGGGGGTTTGAAGAGGCTGCCAGCCGGACTTCTGCTGGGCCATGGACCCATCAGAATGGTTTTGGCGATACTAGCCTTCTTGAGATTCACAGCAATCAAACCATCACTGGGCCTCATCAATAGATGGGGTTCCGTGGGGAAGAAGGAGGCTATGGAAATAATAAAAAAGTTCAAGAAAGACCTTGCTGCCATGTTGAGAATTATCAATGCTAGGAAGGAGAGGAAGAGACGTGGCGCTGACACCAGCATCGGAATCGTCGGCCTCCTGCTGACCACAGCCATGGCAGCCGAGATCACTAGACGTGGGAGTGCATACTACATGTACTTGGACAGGAGCGATGCTGGTAAGGCCATTTCTTTTGCCACCACATTGGGGGTGAACAAATGCCATGTACAGATCATGGACCTCGGGCACATGTGTGACGCCACCATGAGTTATGAGTGCCCCATGCTAGACGAGGGAGTGGAGCCAGATGACGTCGATTGCTGGTGCAACACGACATCGACTTGGGTTGTGTACGGAACCTGTCATCATAAAAAAGGTGAAGCACGACGATCCAGAAGAGCCGTGACGCTTCCTTCTCACTCCACGAGGAAGCTGCAAACGCGATCGCAGACTTGGCTAGAGTCAAGAGAATACACAAAGCACCTGATCAAGGTTGAAAATTGGATATTCAGGAACCCCGGGTTTGCGCTAGTGGCTGTAGCTATAGCCTGGCTCCTGGGAAGCTCGACGAGCCAAAAAGTTATATACTTGGTCATGATATTGTTGATTGCCCCGGCATACAGCATTAGGTGCATAGGAGTTAGCAATAGAGACTTCGTGGAGGGCATGTCAGGTGGGACCTGGGTTGATGTTGTCTTGGAACATGGGGGTTGCGTCACCGTGATGGCACAGGACAAGCCAACAGTTGATATCGAGTTGGTCACGACAACGGTTAGCAACATGGCCGAGGTAAGATCCTACTGCTATGAGGCATCAATATCGGACATGGCTTCGGACAGCCGTTGTCCAACACAAGGTGAAGCCTACCTTGACAAGCAGTCAGACACTCAATATGTCTGCAAGAGAACATTGGTGGATAGAGGTTGGGGAAATGGGTGTGGACTTTTTGGCAAAGGGAGCTTGGTGACATGTGCCAAGTTTACGTGCTCCAAGAAAATGACAGGCAAGAGCATCCAGCCGGAGAACTTGGAGTACCGGATAATGCTATCAGTGCATGGATCCCAGCACAGTGGGATGATTGTGAATGACACAGGACATGAAACTGACGAAAACAGAGCAAAAGTCGAGGTCACACCCAATTCACCAAGAGCAGAAGCAACCTTGGGAGGTTTTGGAAGCTTGGGACTTGACTGTGAACCAAGGACAGGCCTTGACTTCTCAGATCTGTATTACTTGACCATGAACAATAAGCATTGGTTGGTGCACAAGGAGTGGTTTCATGACATCCCATTACCTTGGCATGCTGGTGCAGACACTGGAACTCCACACTGGAACAACAAAGAGGCATTGGTGGAGTTCAAGGACGCCCACGCCAAGAGGCAAACTGTTGTGGTTCTGGGGAGCCAAGAGGGAGCTGTTCATACGGCCCTCGCTGGAGCTTTGGAGGCTGAGATGGATGGTGCAAAGGGAAGGCTATTCTCTGGCCATTTGAAATGCCGCCTAAAAATGGACAAGCTTAGGTTGAAGGGTGTGTCATATTCCCTGTGTACCGCAGCGTTCACATTTACCAAGGTCCCAGCTGAAACATTGCATGGAACAGTCACAGTGGAGGTGCAGTATGCAGGGACAGACGGACCCTGCAAAGTCCCAGCCCAGATGGCGGTGGACATGCAGACCCTGACCCCAGTTGGAAGGCTGATAACCGCCAATCCTGTGATCACTGAAAGTACTGAGAATTCAAAGATGATGTTGGAGCTCGACCCACCATTTGGGGATTCTTACATTGTCATAGGAGTCGGGGACAAGAAAATCACCCATCACTGGCATCGGAGTGGTAGCACCATCGGAAAGGCATTTGAAGCCACTGTGAGAGGTGCCAAGAGAATGGCAGTCTTGGGGGACACAGCCTGGGACTTTGGATCAGTTGGGGGTGTGTTTAATTCATTGGGTAAGGGTATTCACCAGATCTTTGGAGCAGCTTTCAAATCACTGTTTGGAGGAATGTCCTGGTTCTCACAGATCCTCATAGGCACACTGTTGGTGTGGTTAGGTTTGAACACAAAGAATGGATCTATCTCCCTCACATGCTTAGCCCTGGGGGGAGTGATGATCTTCCTTTCCACGGCTGTTTCTGCTGATGTTGGGTGCTCGGTGGACTTCTCAAAAAAGGAAACGAGATGTGGCACGGGGGTGTTCGTCTACAATGACGTTGAAGCCTGGAGGGACCGGTACAAGTACCATCCTGACTCCCCCCGCAGATTGGCAGCAGCTGTTAAGCAGGCTTGGGAAGAGGGGATTTGTGGGATCTCCTCCGTTTCGAGAATGGAAAACATCATGTGGAAATCAGTGGAAGGGGAGCTTAATGCAATCCTAGAGGAGAATGGAGTTCAACTGACAGTTGTAGTGGGGTCTGTAAAAAACCCTATGTGGAGAGGTCCACAGAGATTGCCAGTGCCTGTGAATGAGCTGCCCCATGGCTGGAAAGCCTGGGGGAAATCGTACTTTGTCAGAGCAGCAAAGACCAACAACAGTTTTGTTGTCGACGGTGACACACTGAAGGAGTGTCCGCTCAAACATAGAGCATGGAATAGCTTCCTTGTGGAGGATCACGGGTTTGGGATCTTCCACACCAGTGTTTGGCTGAAGGTCAGAGAGGACTACTCACTAGAGTGTGACCCAGCCGTCATAGGAACAGCTGTCAAGGGAAAGGAAGCTGCACACAGTGATCTAGGCTATTGGATTGAGAGTGAAAAGAATGACACATGGAGGCTGAGGAGGGCTCATCTGATTGAGATGAAAACATGTGAGTGGCCAAAGTCTCACACACTGTGGACAGATGGAGTGGAAGAAAGTGATCTGATCATACCCAAGTCCTTAGCTGGTCCACTCAGCCACCACAACACCAGAGAGGGTTACAGAACTCAAGTGAAAGGGCCATGGCATAGTGAAGAGCTCGAAATCCGGTTTGAGGAGTGCCCAGGAACCAAGGTTCACGTGGAGGAGACATGCGGAACTAGAGGACCATCTCTGAGATCAACTACTGCTAGTGGAAGGGTCATAGAGGAATGGTGCTGTAGGGAATGCACAATGCCTCCACTATCGTTCCGGGCGAAAGACGGCTGCTGGTATGGAATGGAGATAAGGCCCAGAAAGGAACCAGAGAGCAACTTAGTGAGGTCCATGGTGACAGCAGGATCAACCGATCATATGGATCACTTCTCTCTTGGAGTGCTTGTGATTCTACTCATGGTGCAGGAAGGTTTGAAGAAGAGAATGACCACAAAGATCATAATGAGCACATCAATGGCAGTGCTGGTAGCCATGGTCTTGGGAGGATTCTCAATGAGTGACCTGGCTAAGCTTGTGATCCTGATGGGTGCTACTTTCGCAGAAATGAACACTGGAGGAGATGTAGCTCACTTGGCATTGGTTGCGGCATTTAAAGTCAGACCAGCCTTGTTGGTTTCCTTCATCTTCAGAGCCAACTGGACACCCCGTGAGAGCATGCTGCTAGCCCTGGCTTCGTGCCTCCTGCAGACTGCGATCTCTGCTCTTGAAGGCGAGCTGATGGTCCTCGTTAATGGATTTGCTTTGGCCTGGTTGGCAATACGAGCAATGGCCGTGCCACGCACTGACAATATCGCTCTGGCAATTCTGGCTGCTCTAACACCATTAGCCAGAGGCACACTGCTCGTGGCATGGAGAGCGGGCCTCGCCACTTGTGGAGGGTTCATGCTCCTTTCCCTGAAAGGGAAAGGTAGTGTGAAGAAGAACCTGCCTTTTGTCATGGCCTTGGGGTTGACCGCTGTGAGGATAGTGGACCCCATCAATGTGGTAGGACTACTGTTACTCACAAGGAGTGGAAAACGGAGCTGGCCTCCTAGTGAAGTGCTTACAGCTGTTGGCCTGATATGTGCACTGGCCGGAGGGTTTGCCAAGGCAGACATAGAGATGGCTGGGCCCATGGCTGCAGTGGGCCTGCTAATTGTCAGTTATGTGGTCTCGGGAAAGAGTGTGGACATGTACATCGAAAGAGCAGGTGATATCACATGGGAAAAAGACGCGGAAGTCACTGGAAACAGTCCTCGGCTTGACGTGGCACTAGATGAGAGTGGTGATTTCTCCTTGGTAGAGGAGGATGGCCCACCCATGAGAGAGATCATACTCAAGGTGGTTCTGATGGCCATCTGTGGCATGAACCCAATAGCCATACCCTTCGCTGCAGGAGCGTGGTATGTGTATGTAAAGACTGGAAAAAGGAGTGGTGCCCTCTGGGACGTGCCTGCTCCCAAAGAAGTGAAAAAAGGAGAGACTACAGATGGAGTGTACAGAGTGATGACTCGCAGACTGCTGGGTTCAACACAGGTTGGAGTGGGAGTCATGCAAGAGGGAGTCTTCCACACCATGTGGCACGTCACAAAAGGAGCCGCATTGAGGAGCGGTGAAGGAAGACTTGATCCATATTGGGGGGACGTCAAGCAGGACTTGGTGTCATATTGTGGGCCTTGGAAGTTGGATGCAGCCTGGGATGGACTAAGTGAAGTGCAGCTTTTGGCCGTACCCCCCGGAGAGAGGGCTAGAAACATTCAGACTCTGCCTGGAATATTCAAGACAAAGGATGGGGACATCGGAGCAGTCGCTCTAGACTACCCCGCAGGAACCTCAGGATCTCCAATCCTAGACAAATGCGGAAGAGTGATAGGACTTTATGGCAATGGGGTTGTGATCAAGAATGGAAGCTATGTTAGTGCCATAACCCAGGGAAAAAGGGAGGAGGAGGCTCCAGTTGAGTGCTTTGAACCCTCGATGCTGAGGAAGAAGCAGCTAACAGTCTTGGATCTGCATCCAGGAGCCGGGAAAACCAGGAGGGTTCTTCCTGAAATAGTCCGTGAAGCCATAAAGAAGAGACTTCGCACAGTGATCCTAGCACCAACCAGGGTCGTTGCTGCTGAGATGGAGGAAGCCCTAAGAGGACTGCCGGTGCGTTACATGACAACAGCAGTCAACGTCACCCACTCTGGGACAGAAATCGTCGATTTGATGTGCCATGCCACCTTCACTTCACGCCTACTACAACCCATCAGAGTCCCCAACTACAACCTTTACATCATGGATGAAGCTCATTTCACAGATCCCTCAAGCATAGCTGCACGAGGATATATATCAACAAGGGTTGAAATGGGCGAGGCGGCTGCTATCTTCATGACTGCTACACCACCAGGAACCCGCGATGCGTTTCCGGATTCCAACTCACCAATCATGGACACAGAAGTGGAAGTTCCAGAGAGAGCCTGGAGCTCAGGCTTTGACTGGGTGACGGATCATTCTGGGAAAACAATTTGGTTTGTTCCAAGTGTGAGAAACGGAAATGAAATCGCAGCCTGTCTGACAAAGGCTGGAAAGCGGGTCATACAGCTCAGCAGGAAGACTTTTGAGACAGAGTTTCAGAAGACAAAAAATCAAGAGTGGGACTTTGTCATAACAACTGACATTTCAGAGATGGGTGCCAATTTCAAGGCTGACCGGGTCATAGATTCCAGGAGATGCCTAAAGCCAGTCATACTCGATGGCGAGAGAGTCATCCTGGCTGGGCCCATGCCTGTCACGCATGCCAGTGCTGCTCAGAGGAGAGGACGTATAGGCAGGAACCCCAACAAACCTGGAGATGAGTACATGTATGGAGGTGGGTGTGCAGAGACTGATGAAGACCATGCACACTGGCTTGAAGCAAGAATGCTTCTTGACAACATTTACCTCCAGGATGGCCTCATAGCCTCGCTCTATCGGCCTGAGGCCGACAAGGTAGCCGCCATTGAGGGAGAGTTCAAGCTGAGGACAGAGCAAAGGAAGACCTTTGTGGAACTCATGAAGAGAGGAGACCTTCCCGTTTGGCTGGCCTATCAAGTAGCATCTGCCGGAATAACTTACACAGATAGAAGATGGTGCTTTGATGGCACTACCAACAACACCATAATGGAAGACAGTGTACCAGCAGAGGTGTGGACCAAGTATGGAGAGAAGAGAGTGCTTAAACCGAGGTGGATGGATGCTAGGGTCTGTTCAGATCATGCGGCTCTGAAGTCGTTCAAAGAATTTGCCGCTGGGAAGAGAGGAGCGGCTTTGGGAGTAATGGATGCCCTGGGAACATTGCCAGGACACATGACAGAGAGGTTTCAGGAAGCCATTGACAATCTCGCTGTGCTCATGCGAGCAGAGACTGGAAGTAGGCCCTACAAAGCAGCGGCAGCTCAACTGCCGGAGACCTTAGAGACTATCATGCTTCTGGGCTTATTGGGAACAGTTTCGCTAGGAATCTTCTTTGTCTTGATGCGGAACAAGGGCATCGGGAAGATGGGCTTCGGAATGGTAACCCTTGGGGCCAGCGCATGGCTCATGTGGCTTTCGGAAATTGAACCAGCCAGAATCGCATGTGTCCTCATTGTCGTGTTTTTGTTGCTGGTGGTGCTCATACCCGAGCCAGAGAAGCAAAGATCTCCCCAGGATAATCAAATGGCAATCATCATCATGGTGGCAGTTGGCCTTCTGGGTTTGATAACTGCAAATGAACTCGGATGGCTGGAAAGGACAAAAAGTGATATAGCTCATCTAATGGGAAGGAAAGAAGAGGGGACAACCATGGGATTCTCAATGGATATTGATCTGCGGCCAGCCTCCGCCTGGGCTATTTATGCCGCATTGACAACTCTCATCACCCCAGCCGTCCAACATGCGGTAACCACCTCATACAACAACTACTCCCTGATGGCGATGGCCACACAAGCTGGAGTGCTGTTTGGCATGGGCAAAGGGATGCCATTTTATGCATGGG-ACTTTGGAGTCCCGCTGCTAATGATGGGTTGCTACTCACAATTAACACCCCTGACCCTGATAGTGGCCATCATTCTGCTTGTGGCACACTACATGTACTTGATCCCAGGTTTGCAGGCAGCAGCAGCACGTGCTGCCCAGAAGAGGACAGCAGCTGGCATCATGAAGAATCCCGTTGTGGATGGAATAGTGGTGACTGACATTGACACAATGACAATTGACCCCCAAGTGGAGAAGAAGATGGGACAAGTGTTACTCATAGCAGTAGCTGTCTCCAGTGCTGTGCTACTGCGGACCGCTTGGGGATGGGGGGAGGCTGGGGCTCTGATCACAGCAGCAACCTCCACCTTATGGGAAGGCTCTCCAAACAAATACTGGAACTCCTCCACAGCCACCTCACTGTGCAACATCTTTAGAGGAAGTTATTTGGCAGGGGCTTCCCTTATTCACACAGTGACAAGAAATGCCGGTCTGGTTAAGAGACGTGGAGGTGGAACGGGAGAGACCCTGGGAGAGAAGTGGAAAGCCCGCCTGAACCAGATGTCGGCCTTGGAGTTCTACTCTTACAAAAAGTCAGGCATCACCGAAGTGTGTAGGGAGGAGGCGCGCCGCGCCCTCAAGGATGGAGTGGCCACAGGAGGACATGCTGTATCCCGGGGAAGCGCAAAGCTTAGATGGTTGGTAGAGAGAGGATACCTGCAGCCCCATGGAAAGGTTGTTGACCTTGGATGTGGCAGAGGAGGCTGGAGTTATTACGCTGCCACCATCCGTAAAGTGCAGGAGGTCAGAGGATACACAAAGGGAGGTCCTGGCCATGAAGAGCCCATGCTGGTGCAAAGCTATGGGTGGAACATAGTTCGCCTCAAGAGTGGAGTGGACGTCTTCCACATGGCGGCTGAACCGTGTGACACTCTGCTGTGTGACATAGGCGAGTCATCATCCAGTCCTGAAGTGGAAGAGACGCGAACACTCAGAGTGCTCTCCATGGTGGGAGACTGGCTTGAAAAAAGACCAGGGGCCTTCTGCATAAAGGTGCTGTGCCCATACACCAGCACAATGATGGAGACCATGGAGCGACTGCAACGTAGGCATGGGGGAGGATTAGTCAGAGTGCCATTGTCCCGCAACTCTACACATGAGATGTATTGGGTCTCTGGAGCCAAAAGCAACATCATAAAGAGTGTGTCCACCACAAGTCAGCTCCTCTTGGGACGCATGGAAGGGCCTAGGAGGCCAGTGAAATATGAGGAGGATGTGAACCTCGGCTCAGGCACACGAGCTGTGGCAAGCTGCGCTGAGGCTCCCAACATGAAGATCATTGGTAGGCGCATTGAGAGAATCCGCAATGAACATGCAGAGACATGGTTCTTTGATGAAAACCACCCATACAGGACATGGGCCTACCATGGGAGCTACGAAGCCCCCACGCAGGGGTCAGCGTCATCCCTCGTGAACGGGGTTGTTAGACTCTTGTCAAAGCCCTGGGATGTGGTGACTGGAGTTACAGGAATAGCTATGACTGACACCACGCCATACGGCCAACAAAGAGTCTTCAAGGAAAAAGTGGACACCAGGGTGCCAGATCCCCAAGAAGGCACTCGCCAAGTGATGAACATGGTATCGTCTTGGTTATGGAAGGAGCTGGGAAAACGCAAGCGGCCACGTGTCTGCACCAAAGAAGAGTTCATTAATAAGGTGCGCAGCAATGCAGCACTGGGAGCAATATTTGAAGAGGAAAAAGAATGGAAGACAGCTGTAGAAGCTGTGAATGATCCAAGATTTTGGGCTCTAGTGGACAAGGAAAGAGAACACCACCTGAGAGGAGAGTGTCATAGCTGTGTGTACAACATGATGGGCAAAAGAGAAAAGAAGCAAGGAGAATTCGGGAAAGCAAAAGGCAGCCGCGCAATCTGGTACATGTGGTTGGGAGCCAGATTTTTGGAGTTTGAAGCTCTTGGGTTCTTGAACGAGGACCACTGGATGGGGAGAGAAAACTCAGGAGGTGGCGTTGAAGGGCTAGGACTGCAAAGGCTTGGATATATCCTAGAAGAAATGAACCGGGCACCAGGAGGAAAGATGTATGCAGATGACACTGCTGGCTGGGACACCCGCATTAGCAAGTTTGATCTAGAGAATGAAGCCCTGATCACTAACCAGATGGAAGAAGGGCACAGAGCTCTGGCGTTGGCCGTGATTAAATACACATACCAAAACAAAGTGGTGAAGGTTCTCAGACCAGCTGAAGGAGGGAAAACAGTCATGGACATCATCTCAAGACAAGACCAGAGAGGGAGCGGACAAGTTGTTACTTATGCTCTCAACACATTCACCAACCTGGTGGTGCAGCTTATCCGGAACATGGAGGCTGAGGAAGTGCTAGAGATGCATGATCTGTGGTTGTTGAGGAAGCCAGAGAAAGTGACCAGATGGTTGCAGAGCAATGGATGGGACAGACTCAAACGGATGGCAGTTAGTGGAGATGACTGCGTTGTAAAGCCAATTGATGATAGGTTTGCACATGCCCTCAGGTTCTTGAATGACATGGGAAAGGTTAGGAAAGACACACAGGAATGGAAACCCTCGACTGGATGGAGCAATTGGGAAGAGGTCCCGTTCTGTTCTCACCACTTCAACAAGCTGCACCTCAAGGACGGGAGATCCATTGTGGTCCCCTGCCGCCACCAAGATGAACTGATTGGCCGAGCCCGTGTCTCACCAGGGGCAGGATGGAGCATCCGGGAGACTGCTTGTCTTGCAAAATCATATGCACAGATGTGGCAGCTTCTTTATTTCCACAGAAGAGACCTCCGACTGATGGCCAATGCCATTTGTTCGGCTGTGCCAATTGACTGGGTACCAACTGGGAGAACCACCTGGTCAATCCATGGAAAGGGAGAATGGATGACTACTGAGGACATGCTCATGGTGTGGAATAGAGTGTGGATTGAGGAGAACGACCACATGGAGGACAAGACCCCTGTAACAAAATGGACAGACATTCCCTATTTGGGAAAAAGGGAGGACTTATGGTGTGGATCCCTTATAGGGCATAGACCTCGCACCACTTGGGCTGAGAACATCAAAGACACAGTCAACATGGTGCGTAGGATCATAGGTGATGAAGAAAAGTTCATGGACTACCTATCCACCCAAGTACGCTACTTGGGTGAGGAAGGGTCCACACCTGGAGTGCTGTAA

>ENA|AMR39833|AMR39833.1 Zika virus polyprotein

ATGAAAAACCC-AAAGAAGAAATCCGGAGGATTCCGGATTGTCAATATGCTAAAACGCGGAGTAGCCCGTGTAAACCCCTTGGGGGGTTTGAAGAGGCTGCCAGCCGGACTTCTGCTGGGCCATGGACCCATCAGAATGGTTTTGGCGATACTAGCCTTCTTGAGATTCACAGCAATCAAACCATCACTGGGCCTCATCAATAGATGGGGTTCCGTGGGGAAGAAGGAGGCTATGGAAATAATAAAAAAGTTCAAGAAAGACCTTGCTGCCATGTTGAGAATTATCAATGCTAGGAAGGAGAGGAAGAGACGTGGCGCTGACACCAGCATCGGAATCGTCGGCCTCCTGCTGACCACAGCCATGGCAGCCGAGATCACTAGACGTGGGAGTGCATACTACATGTACTTGGACAGGAGCGATGCTGGTAAGGCCATTTCTTTTGCCACCACATTGGGGGTGAACAAATGCCATGTACAGATCATGGACCTCGGGCACATGTGTGACGCCACCATGAGTTATGAGTGCCCCATGCTAGACGAGGGAGTGGAGCCAGATGACGTCGATTGCTGGTGCAACACGACATCGACTTGGGTTGTGTACGGAACCTGTCATCATAAAAAAGGTGAAGCACGACGATCCAGAAGAGCCGTGACGCTTCCTTCTCACTCCACGAGGAAGCTGCAAACGCGATCGCAGACTTGGCTAGAGTCAAGAGAATACACAAAGCACCTGATCAAGGTTGAAAATTGGATATTCAGGAACCCCGGGTTTGCGCTAGTGGCTGTAGCTATAGCCTGGCTCCTGGGAAGCTCGACGAGCCAAAAAGTTATATACTTGGTCATGATATTGTTGATTGCCCCGGCATACAGCATTAGGTGCATAGGAGTTAGCAATAGAGACTTCGTGGAGGGCATGTCAGGTGGGACCTGGGTTGATGTTGTCTTGGAACATGGGGGTTGCGTCACCGTGATGGCACAGGACAAGCCAACAGTTGATATCGAGTTGGTCACGACAACGGTTAGCAACATGGCCGAGGTAAGATCCTACTGCTATGAGGCATCAATATCGGACATGGCTTCGGACAGCCGTTGTCCAACACAAGGTGAAGCCTACCTTGACAAGCAGTCAGACACTCAATATGTCTGCAAGAGAACATTGGTGGATAGAGGTTGGGGAAATGGGTGTGGACTTTTTGGCAAAGGGAGCTTGGTGACATGTGCCAAGTTTACGTGCTCCAAGAAAATGACAGGCAAGAGCATCCAGCCGGAGAACTTGGAGTACCGGATAATGCTATCAGTGCATGGATCCCAGCACAGTGGGATGATTGTGAATGACACAGGACATGAAACTGACGAAAACAGAGCAAAAGTCGAGGTCACACCCAATTCACCAAGAGCAGAAGCAACCTTGGGAGGTTTTGGAAGCTTGGGACTTGACTGTGAACCAAGGACAGGCCTTGACTTCTCAGATCTGTATTACCTGACCATGAACAATAAGCATTGGTTGGTGCACAAGGAGTGGTTTCATGACATCCCATTACCTTGGCATGCTGGTGCAGACACTGGAACTCCACACTGGAACAACAAAGAGGCATTGGTGGAGTTCAAGGACGCCCACGCCAAGAGGCAAACTGTTGTGGTTCTGGGGAGCCAAGAGGGAGCTGTTCATACGGCCCTCGCTGGAGCTTTGGAGGCTGAGATGGATGGTGCAAAGGGAAGGCTATTCTCTGGCCATTTGAAATGCCGCCTAAAAATGGACAAGCTTAGGTTGAAGGGTGTGTCATATTCCCTGTGTACCGCAGCGTTCACATTTACCAAGGTCCCAGCTGAAACATTGCATGGAACAGTCACAGTGGAGGTGCAGTATGCAGGGACAGACGGACCCTGCAAAGTCCCAGCCCAGATGGCGGTGGACATGCAGACCCTGACCCCAGTTGGAAGGCTGATAACCGCCAATCCTGTGATCACTGAAAGTACTGAGAATTCAAAGATGATGTTGGAGCTCGACCCACCATTTGGGGATTCTTACATTGTCATAGGAGTCGGGGACAAGAAAATCACCCATCACTGGCATCGGAGTGGTAGCACCATCGGAAAGGCATTTGAAGCCACTGTGAGAGGTGCCAAGAGAATGGCAGTCTTGGGGGACACAGCCTGGGACTTTGGATCAGTTGGGGGTGTGTTTAATTCATTGGGTAAGGGTATTCACCAGATCTTTGGAGCAGCTTTCAAATCACTGTTTGGAGGAATGTCCTGGTTCTCACAGATCCTCATAGGCACACTGTTGGTGTGGTTAGGTTTGAACACAAAGAATGGATCTATCTCCCTCACATGCTTAGCCCTGGGGGGAGTGATGATCTTCCTTTCCACGGCTGTTTCTGCTGATGTTGGGTGCTCGGTGGACTTCTCAAAAAAGGAAACGAGATGTGGCACGGGGGTGTTCGTCTACAATGACGTTGAAGCCTGGAGGGACCGGTACAAGTACCATCCTGACTCCCCCCGCAGATTGGCAGCAGCTGTTAAGCAGGCTTGGGAAGAGGGGATTTGTGGGATCTCCTCCGTTTCGAGAATGGAAAACATCATGTGGAAATCAGTGGAAGGGGAGCTTAATGCAATCCTAGAGGAGAATGGAGTTCAACTGACAGTTGTAGTGGGGTCTGTAAAAAACCCTATGTGGAGAGGTCCACAGAGATTGCCAGTGCCTGTGAATGAGCTGCCCCATGGCTGGAAAGCCTGGGGGAAATCGTACTTTGTCAGAGCAGCAAAGACCAACAACAGTTTTGTTGTCGACGGTGACACACTGAAGGAGTGTCCGCTCAAACATAGAGCATGGAATAGCTTCCTTGTGGAGGATCACGGGTTTGGGATCTTCCACACCAGTGTTTGGCTGAAGGTCAGAGAGGACTACTCACTAGAGTGTGACCCAGCCGTCATAGGAACAGCTGTCAAGGGAAAGGAAGCTGCACACAGTGATCTAGGCTATTGGATTGAGAGTGAAAAGAATGACACATGGAGGCTGAGGAGGGCTCATCTGATTGAGATGAAAACATGTGAGTGGCCAAAGTCTCACACACTGTGGACAGATGGAGTGGAAGAAAGTGATCTGATCATACCCAAGTCCTTAGCTGGTCCACTCAGCCACCACAACACCAGAGAGGGTTACAGAACTCAAGTGAAAGGGCCATGGCATAGTGAAGAGCTCGAAATCCGGTTTGAGGAGTGCCCAGGAACCAAGGTTCACGTGGAGGAGACATGCGGAACTAGAGGACCATCTCTGAGATCAACTACTGCTAGTGGAAGGGTCATAGAGGAATGGTGCTGTAGGGAATGCACAATGCCTCCACTATCGTTCCGGGCGAAAGACGGCTGCTGGTATGGAATGGAGATAAGGCCCAGAAAGGAACCAGAGAGCAACTTAGTGAGGTCCATGGTGACAGCAGGATCAACCGATCATATGGATCACTTCTCTCTTGGAGTGCTTGTGATTCTACTCATGGTGCAGGAAGGTTTGAAGAAGAGAATGACCACAAAGATCATAATGAGCACATCAATGGCAGTGCTGGTAGCCATGGTCTTGGGAGGATTCTCAATGAGTGACCTGGCTAAGCTTGTGATCCTGATGGGTGCTACTTTCGCAGAAATGAACACTGGAGGAGATGTAGCTCACTTGGCATTGGTTGCGGCATTTAAAGTCAGACCAGCCTTGTTGGTTTCCTTCATCTTCAGAGCCAACTGGACACCCCGTGAGAGCATGCTGCTAGCCCTGGCTTCGTGCCTCCTGCAGACTGCGATCTCTGCTCTTGAAGGCGAGCTGATGGTCCTCGTTAATGGATTTGCTTTGGCCTGGTTGGCAATACGAGCAATGGCCGTGCCACGCACTGACAATATCGCTCTGGCAATTCTGGCTGCTCTAACACCATTAGCCAGAGGCACACTGCTCGTGGCATGGAGAGCGGGCCTCGCCACTTGTGGAGGGTTCATGCTCCTTTCCCTGAAAGGGAAAGGTAGTGTGAAGAAGAACCTGCCTTTTGTCATGGCCTTGGGGTTGACCGCTGTGAGGATAGTGGACCCCATCAATGTGGTAGGACTACTGTTACTCACAAGGAGTGGAAAACGGAGCTGGCCTCCTAGTGAAGTGCTTACAGCTGTTGGCCTGATATGTGCACTGGCCGGAGGGTTTGCCAAGGCAGACATAGAGATGGCTGGGCCCATGGCTGCAGTGGGCCTGCTAATTGTCAGTTATGTGGTCTCGGGAAAGAGTGTGGACATGTACATCGAAAGAGCAGGTGATATCACATGGGAAAAAGACGCGGAAGTCACTGGAAACAGTCCTCGGCTTGACGTGGCACTAGATGAGAGTGGTGATTTCTCCTTGGTAGAGGAGGATGGCCCACCCATGAGAGAGATCATACTCAAGGTGGTTCTGATGGCCATCTGTGGCATGAACCCAATAGCCATACCCTTCGCTGCAGGAGCGTGGTATGTGTATGTAAAGACTGGAAAAAGGAGTGGTGCCCTCTGGGACGTGCCTGCTCCCAAAGAAGTGAAAAAAGGAGAGACTACAGATGGAGTGTACAGAGTGATGACTCGCAGACTGCTGGGTTCAACACAGGTTGGAGTGGGAGTCATGCAAGAGGGAGTCTTCCACACCATGTGGCACGTCACAAAAGGAGCCGCATTGAGGAGCGGTGAAGGAAGACTTGATCCATATTGGGGGGACGTCAAGCAGGACTTGGTGTCATATTGTGGGCCTTGGAAGTTGGATGCAGCCTGGGATGGACTAAGTGAAGTGCAGCTTTTGGCCGTACCCCCCGGAGAGAGGGCTAGAAACATTCAGACTCTGCCTGGAATATTCAAGACAAAGGATGGGGACATCGGAGCAGTCGCTCTAGACTACCCCGCAGGAACCTCAGGATCTCCAATCCTAGACAAATGCGGAAGAGTGATAGGACTTTATGGCAATGGGGTTGTGATCAAGAATGGAAGCTATGTTAGTGCCATAACCCAGGGAAAAAGGGAGGAGGAGGCTCCAGTTGAGTGCTTTGAACCCTCGATGCTGAGGAAGAAGCAGCTAACAGTCTTGGATCTGCATCCAGGAGCCGGGAAAACCAGGAGGGTTCTTCCTGAAATAGTCCGTGAAGCCATAAAGAAGAGACTTCGCACAGTGATCCTAGCACCAACCAGGGTCGTTGCTGCTGAGATGGAGGAAGCCCTAAGAGGACTGCCGGTGCGTTACATGACAACAGCAGTCAACGTCACCCACTCTGGGACAGAAATCGTCGATTTGATGTGCCATGCCACCTTCACTTCACGCCTACTACAACCCATCAGAGTCCCCAACTACAACCTTTACATCATGGATGAAGCTCATTTCACAGATCCCTCAAGCATAGCTGCACGAGGATATATATCAACAAGGGTTGAAATGGGCGAGGCGGCTGCTATCTTCATGACTGCTACACCACCAGGAACCCGCGATGCGTTTCCGGATTCCAACTCACCAATCATGGACACAGAAGTGGAAGTTCCAGAGAGAGCCTGGAGCTCAGGCTTTGACTGGGTGACGGATCATTCTGGGAAAACAATTTGGTTTGTTCCAAGTGTGAGAAACGGAAATGAAATCGCAGCCTGTCTGACAAAGGCTGGAAAGCGGGTCATACAGCTCAGCAGGAAGACTTTTGAGACAGAGTTTCAGAAGACAAAAAATCAAGAGTGGGACTTTGTCATAACAACTGACATTTCAGAGATGGGTGCCAATTTCAAGGCTGACCGGGTCATAGATTCCAGGAGATGCCTAAAGCCAGTCATACTCGATGGCGAGAGAGTCATCCTGGCTGGGCCCATGCCTGTCACGCATGCCAGTGCTGCTCAGAGGAGAGGACGTATAGGCAGGAACCCCAACAAACCTGGAGATGAGTACATGTATGGAGGTGGGTGTGCAGAGACTGATGAAGACCATGCACACTGGCTTGAAGCAAGAATGCTTCTTGACAACATTTACCTCCAGGATGGCCTCATAGCCTCGCTCTATCGGCCTGAGGCCGACAAGGTAGCCGCCATTGAGGGAGAGTTCAAGCTGAGGACAGAGCAAAGGAAGACCTTTGTGGAACTCATGAAGAGAGGAGACCTTCCCGTTTGGCTGGCCTATCAAGTAGCATCTGCCGGAATAACTTACACAGATAGAAGATGGTGCTTTGATGGCACTACCAACAACACCATAATGGAAGACAGTGTACCAGCAGAGGTGTGGACCAAGTATGGAGAGAAGAGAGTGCTTAAACCGAGGTGGATGGATGCTAGGGTCTGTTCAGATCATGCGGCTCTGAAGTCGTTCAAAGAATTTGCCGCTGGGAAGAGAGGAGCGGCTTTGGGAGTAATGGATGCCCTGGGAACATTGCCAGGACACATGACAGAGAGGTTTCAGGAAGCCATTGACAATCTCGCTGTGCTCATGCGAGCAGAGACTGGAAGTAGGCCCTACAAAGCAGCGGCAGCTCAACTGCCGGAGACCTTAGAGACTATCATGCTTCTGGGCTTATTGGGAACAGTTTCGCTAGGAATCTTCTTTGTCTTGATGCGGAACAAGGGCATCGGGAAGATGGGCTTCGGAATGGTAACCCTTGGGGCCAGCGCATGGCTCATGTGGCTTTCGGAAATTGAACCAGCCAGAATCGCATGTGTCCTCATTGTCGTGTTTTTGTTGCTGGTGGTGCTCATACCCGAGCCAGAGAAGCAAAGATCTCCCCAGGATAATCAAATGGCAATCATCATCATGGTGGCAGTTGGCCTTCTGGGTTTGATAACTGCAAATGAACTCGGATGGCTGGAAAGGACAAAAAGTGATATAGCTCATCTAATGGGAAGGAAAGAAGAGGGGACAACCATGGGATTCTCAATGGATATTGATCTGCGGCCAGCCTCCGCCTGGGCTATTTATGCCGCATTGACAACTCTCATCACCCCAGCCGTCCAACATGCGGTAACCACCTCATACAACAACTACTCCCTGATGGCGATGGCCACACAAGCTGGAGTGCTGTTTGGCATGGGCAAAGGGATGCCATTTTATGCATGGG-ACTTTGGAGTCCCGCTGCTAATGATGGGTTGCTACTCACAATTAACACCCCTGACCCTGATAGTGGCCATCATTCTGCTTGTGGCACACTACATGTACTTGATCCCAGGTTTGCAGGCAGCAGCAGCACGTGCTGCCCAGAAGAGGACAGCAGCTGGCATCATGAAGAATCCCGTTGTGGATGGAATAGTGGTGACTGACATTGACACAATGACAATTGACCCCCAAGTGGAGAAGAAGATGGGACAAGTGTTACTCATAGCAGTAGCTGTCTCCAGTGCTGTGCTACTGCGGACCGCTTGGGGATGGGGGGAGGCTGGGGCTCTGATCACAGCAGCAACCTCCACCTTATGGGAAGGCTCTCCAAACAAATACTGGAACTCCTCCACAGCCACCTCACTGTGCAACATCTTTAGAGGAAGTTATTTGGCAGGGGCTTCCCTTATTTACACAGTGACAAGAAATGCCGGTCTGGTTAAGAGACGTGGAGGTGGAACGGGAGAGACCCTGGGAGAGAAGTGGAAAGCCCGCCTGAACCAGATGTCGGCCTTGGAGTTCTACTCTTACAAAAAGTCAGGCATCACCGAAGTGTGTAGGGAGGAGGCGCGCCGCGCCCTCAAGGATGGAGTGGCCACAGGAGGACATGCTGTATCCCGGGGAAGCGCAAAGCTTAGATGGTTGGTAGAGAGAGGATACCTGCAGCCCCATGGAAAGGTTGTTGACCTTGGATGTGGCAGAGGAGGCTGGAGTTATTACGCTGCCACCATCCGTAAAGTGCAGGAGGTCAGAGGATACACAAAGGGAGGTCCTGGCCATGAAGAGCCCATGCTGGTGCAAAGCTATGGGTGGAACATAGTTCGCCTCAAGAGTGGAGTGGACGTCTTCCACATGGCGGCTGAACCGTGTGACACTCTGCTGTGTGACATAGGCGAGTCATCATCCAGTCCTGAAGTGGAAGAGACGCGAACACTCAGAGTGCTCTCCATGGTGGGAGACTGGCTTGAAAAAAGACCAGGGGCCTTCTGCATAAAGGTGCTGTGCCCATACACCAGCACAATGATGGAGACCATGGAGCGACTGCAACGTAGGCATGGGGGAGGATTAGTCAGAGTGCCATTGTCCCGCAACTCTACACATGAGATGTATTGGGTCTCTGGAGCCAAAAGTAACATCATAAAGAGTGTGTCCACCACAAGTCAGCTCCTCTTGGGACGCATGGAAGGGCCTAGGAGGCCAGTGAAATATGAGGAGGATGTGAACCTCGGCTCAGGCACACGAGCTGTGGCAAGCTGCGCTGAGGCTCCCAACATGAAGATCATTGGTAGGCGCATTGAGAGAATCCGCAATGAACATGCAGAGACATGGTTCTTTGATGAAAACCACCCATACAGGACATGGGCCTACCATGGGAGCTACGAAGCCCCCACGCAGGGGTCAGCGTCATCCCTCGTGAACGGGGTTGTTAGACTCTTGTCAAAGCCCTGGGATGTGGTGACTGGAGTTACAGGAATAGCTATGACTGACACCACGCCATACGGCCAACAAAGAGTCTTCAAGGAAAAAGTGGACACCAGGGTGCCAGATCCCCAAGAAGGCACTCGCCAAGTGATGAACATGGTATCGTCTTGGTTATGGAAGGAGCTGGGAAAACGCAAGCGGCCACGTGTCTGCACCAAAGAAGAGTTCATTAATAAGGTGCGCAGCAATGCAGCACTGGGAGCAATATTTGAAGAGGAAAAAGAATGGAAGACAGCTGTAGAAGCTGTGAATGATCCAAGATTTTGGGCTCTAGTGGACAAGGAAAGAGAACACCACCTGAGAGGAGAGTGTCATAGCTGTGTGTACAACATGATGGGCAAAAGAGAAAAGAAGCAAGGAGAATTCGGGAAAGCAAAAGGCAGCCGTGCAATCTGGTACATGTGGTTGGGAGCCAGATTTTTGGAGTTTGAAGCTCTTGGGTTCTTGAACGAGGACCACTGGATGGGGAGAGAAAACTCAGGAGGTGGCGTTGAAGGGCTAGGACTGCAAAGGCTTGGATATATCCTAGAAGAAATGAACCGGGCACCAGGAGGAAAGATGTATGCAGATGACACTGCTGGCTGGGACACCCGCATTAGCAAGTTTGATCTAGAGAATGAAGCCCTGATCACTAACCAGATGGAAGAAGGGCACAGAGCTCTGGCGTTGGCCGTGATTAAATACACATACCAAAACAAAGTGGTGAAGGTTCTCAGACCAGCTGAAGGAGGGAAAACAGTCATGGACATCATCTCAAGACAAGACCAGAGAGGGAGCGGACAAGTTGTTACTTATGCTCTCAACACATTCACCAACCTGGTGGTGCAGCTTATCCGGAACATGGAGGCTGAGGAAGTGCTAGAGATGCATGATCTGTGGTTGTTGAGGAAGCCAGAGAAAGTGACCAGATGGTTGCAGAGCAATGGATGGGACAGACTCAAACGGATGGCAGTTAGTGGAGATGACTGCGTTGTAAAGCCAATTGATGATAGGTTTGCACATGCCCTCAGGTTCTTGAATGACATGGGAAAGGTTAGGAAAGACACACAGGAATGGAAACCCTCGACTGGATGGAGCAATTGGGAAGAGGTCCCGTTCTGTTCTCACCACTTCAACAAGCTGCACCTCAAGGACGGGAGATCCATTGTGGTCCCCTGCCGCCACCAAGATGAACTGATTGGCCGAGCCCGTGTCTCACCAGGGGCAGGATGGAGCATCCGGGAGACTGCTTGTCTTGCAAAATCATATGCACAGATGTGGCAGCTTCTTTATTTCCACAGAAGAGACCTCCGACTGATGGCCAATGCCATTTGTTCGGCTGTGCCAATTGACTGGGTACCAACTGGGAGAACCACCTGGTCAATCCATGGAAAGGGAGAATGGATGACTACTGAGGACATGCTCATGGTGTGGAATAGAGTGTGGATTGAGGAGAACGACCACATGGAGGACAAGACCCCTGTAACAAAATGGACAGACATTCCCTATTTGGGAAAAAGGGAGGACTTATGGTGTGGATCCCTTATAGGGCATAGACCTCGCACCACTTGGGCTGAGAACATCAAAGACACAGTCAACATGGTGCGTAGGATCATAGGTGATGAAGAAAAGTTCATGGACTACCTATCCACCCAAGTACGCTACTTGGGTGAGGAAGGGTCCACACCTGGAGTGCTGTAA

>ENA|AMR39832|AMR39832.1 Zika virus polyprotein

ATGAAAAACCC-AAAGAAGAAATCCGGAGGATTCCGGATTGTCAATATGCTAAAACGCGGAGTAGCCCGTGTAAACCCCTTGGGGGGTTTGAAGAGGCTGCCAGCCGGACTTCTGCTGGGCCATGGACCCATCAGAATGGTTTTGGCGATACTAGCCTTCTTGAGATTCACAGCAATCAAACCATCACTGGGCCTCATCAATAGATGGGGTTCCGTGGGGAAGAAGGAGGCTATGGAAATAATAAAAAAGTTCAAGAAAGACCTTGCTGCCATGTTGAGAATTATCAATGCTAGGAAGGAGAGGAAGAGACGTGGCGCTGACACCAGCATCGGAATCGTCGGCCTCCTGCTGACCACAGCCATGGCAGCCGAGATCACTAGACGTGGGAGTGCATACTACATGTACTTGGACAGGAGCGATGCTGGTAAGGCCATTTCTTTTGCCACCACATTGGGGGTGAACAAATGCCATGTACAGATCATGGACCTCGGGCACATGTGTGACGCCACCATGAGTTATGAGTGCCCCATGCTAGACGAGGGAGTGGAGCCAGATGACGTCGATTGCTGGTGCAACACGACATCGACTTGGGTTGTGTACGGAACCTGTCATCATAAAAAAGGTGAAGCACGACGATCCAGAAGAGCCGTGACGCTTCCTTCTCACTCCACGAGGAAGCTGCAAACGCGATCGCAGACTTGGCTAGAGTCAAGAGAATACACAAAGCACCTGATCAAGGTTGAAAATTGGATATTTAGGAACCCCGGGTTTGCGCTAGTGGCTGTAGCTATAGCCTGGCTCCTGGGAAGCTCGACGAGCCAAAAAGTTATATACTTGGTCATGATATTGTTGATTGCCCCGGCATACAGCATTAGGTGCATAGGAGTTAGCAATAGAGACTTCGTGGAGGGCATGTCAGGTGGGACCTGGGTTGATGTTGTCTTGGAACATGGGGGTTGCGTCACCGTGATGGCACAGGACAAGCCAACAGTTGATATCGAGTTGGTCACGACAACGGTTAGCAACATGGCCGAGGTAAGATCCTACTGCTATGAGGCATCAATATCGGACATGGCTTCGGACAGCCGTTGTCCAACACAAGGTGAAGCCTACCTTGACAAGCAGTCAGACACTCAATATGTCTGCAAGAGAACATTGGTGGATAGAGGTTGGGGAAATGGGTGTGGACTTTTTGGCAAAGGGAGCTTGGTGACATGTGCCAAGTTTACGTGCTCCAAGAAAATGACAGGCAAGAGCATCCAGCCGGAGAACTTGGAGTACCGGATAATGCTATCAGTGCATGGATCCCAGCACAGTGGGATGATTGTGAATGACACAGGACATGAAACTGACGAAAACAGAGCAAAAGTCGAGGTCACACCCAATTCACCAAGAGCAGAAGCAACCTTGGGAGGTTTTGGAAGCTTGGGACTTGACTGTGAACCAAGGACAGGCCTTGACTTCTCAGATCTGTATTACCTGACCATGAACAATAAGCATTGGTTGGTGCACAAGGAGTGGTTTCATGACATCCCATTACCTTGGCATGCTGGTGCAGACACTGGAACTCCACACTGGAACAACAAAGAGGCATTGGTGGAGTTCAAGGACGCCCACGCCAAGAGGCAAACTGTTGTGGTTCTGGGGAGCCAAGAGGGAGCTGTTCATACGGCCCTCGCTGGAGCTTTGGAGGCTGAGATGGATGGTGCAAAGGGAAGGCTATTCTCTGGCCATTTGAAATGCCGCCTAAAAATGGACAAGCTTAGGTTGAAGGGTGTGTCATATTCCCTGTGTACCGCAGCGTTCACATTTACCAAGGTCCCAGCTGAAACATTGCATGGAACAGTCACAGTGGAGGTGCAGTATGCAGGGACAGACGGACCCTGCAAAGTCCCAGCCCAGATGGCGGTGGACATGCAGACCCTGACCCCAGTTGGAAGGCTGATAACCGCCAATCCTGTGATCACTGAAAGTACTGAGAATTCAAAGATGATGTTGGAGCTCGACCCACCATTTGGGGATTCTTACATTGTCATAGGAGTCGGGGACAAGAAAATCACCCATCACTGGCATCGGAGTGGTAGCACCATCGGAAAGGCATTTGAAGCCACTGTGAGAGGTGCCAAGAGAATGGCAGTCTTGGGGGACACAGCCTGGGACTTTGGATCAGTTGGGGGTGTGTTTAATTCATTGGGTAAGGGTATTCACCAGATCTTTGGAGCAGCTTTCAAATCACTGTTTGGAGGAATGTCCTGGTTCTCACAGATCCTCATAGGCACACTGTTGGTGTGGTTAGGTTTGAACACAAAGAATGGATCTATCTCCCTCACATGCTTAGCCCTGGGGGGAGTGATGATCTTCCTTTCCACGGCTGTTTCTGCTGATGTTGGGTGCTCGGTGGACTTCTCAAAAAAGGAAACGAGATGTGGCACGGGGGTGTTCGTCTACAATGACGTTGAAGCCTGGAGGGACCGGTACAAGTACCATCCTGACTCCCCCCGCAGATTGGCAGCAGCTGTTAAGCAGGCTTGGGAAGAGGGGATTTGTGGGATCTCCTCCGTTTCGAGAATGGAAAACATCATGTGGAAATCAGTGGAAGGGGAGCTTAATGCAATCCTAGAGGAGAATGGAGTTCAACTGACAGTTGTAGTGGGGTCTGTAAAAAACCCTATGTGGAGAGGTCCACAGAGATTGCCAGTGCCTGTGAATGAGCTGCCCCATGGCTGGAAAGCCTGGGGGAAATCGTACTTTGTCAGAGCAGCAAAGACCAACAACAGTTTTGTTGTCGACGGTGACACACTGAAGGAGTGTCCGCTCAAACATAGAGCATGGAATAGCTTCCTTGTGGAGGATCACGGGTTTGGGATCTTCCACACCAGTGTTTGGCTGAAGGTCAGAGAGGACTACTCACTAGAGTGTGACCCAGCCGTCATAGGAACAGCTGTCAAGGGAAAGGAAGCTGCACACAGTGATCTAGGCTATTGGATTGAGAGTGAAAAGAATGACACATGGAGGCTGAGGAGGGCTCATCTGATTGAGATGAAAACATGTGAGTGGCCAAAGTCTCACACACTGTGGACAGATGGAGTGGAAGAAAGTGATCTGATCATACCCAAGTCCTTAGCTGGTCCACTCAGCCACCACAACACCAGAGAGGGTTACAGAACTCAAGTGAAAGGGCCATGGCATAGTGAAGAGCTCGAAATCCGGTTTGAGGAGTGCCCAGGAACCAAGGTTCACGTGGAGGAGACATGCGGAACTAGAGGACCATCTCTGAGATCAACTACTGCTAGTGGAAGGGTCATAGAGGAATGGTGCTGTAGGGAATGCACAATGCCTCCACTATCGTTCCGGGCGAAAGACGGCTGCTGGTATGGAATGGAGATAAGGCCCAGAAAGGAACCAGAGAGCAACTTAGTGAGGTCCATGGTGACAGCAGGATCAACCGATCATATGGATCACTTCTCTCTTGGAGTGCTTGTGATTCTACTCATGGTGCAGGAAGGTTTGAAGAAGAGAATGACCACAAAGATCATAATGAGCACATCAATGGCAGTGCTGGTAGCCATGGTCTTGGGAGGATTCTCAATGAGTGACCTGGCTAAGCTTGTGATCCTGATGGGTGCTACTTTCGCAGAAATGAACACTGGAGGAGATGTAGCTCACTTGGCATTGGTTGCGGCATTTAAAGTCAGACCAGCCTTGTTGGTTTCCTTCATCTTCAGAGCCAACTGGACACCCCGTGAGAGCATGCTGCTAGCCCTGGCTTCGTGCCTCCTGCAGACTGCGATCTCTGCTCTTGAAGGCGAGCTGATGGTCCTCGTTAATGGATTTGCTTTGGCCTGGTTGGCAATACGAGCAATGGCCGTGCCACGCACTGACAATATCGCTCTGGCAATTCTGGCTGCTCTAACACCATTAGCCAGAGGCACACTGCTCGTGGCATGGAGAGCGGGCCTCGCCACTTGTGGAGGGTTCATGCTCCTTTCCCTGAAAGGGAAAGGTAGTGTGAAGAAGAACCTGCCTTTTGTCATGGCCTTGGGGTTGACCGCTGTGAGGATAGTGGACCCCATCAATGTGGTAGGACTACTGTTACTCACAAGGAGTGGAAAACGGAGCTGGCCTCCTAGTGAAGTGCTTACAGCTGTTGGCCTGATATGTGCACTGGCCGGAGGGTTTGCCAAGGCAGACATAGAGATGGCTGGGCCCATGGCTGCAGTGGGCCTGCTAATTGTCAGTTATGTGGTCTCGGGAAAGAGTGTGGACATGTACATCGAAAGAGCAGGTGATATCACATGGGAAAAAGACGCGGAAGTCACTGGAAACAGTCCTCGGCTTGACGTGGCACTAGATGAGAGTGGTGATTTCTCCTTGGTAGAGGAGGATGGCCCACCCATGAGAGAGATCATACTCAAGGTGGTTCTGATGGCCATCTGTGGCATGAACCCAATAGCCATACCCTTCGCTGCAGGAGCGTGGTATGTGTATGTAAAGACTGGAAAAAGGAGTGGTGCCCTCTGGGACGTGCCTGCTCCCAAAGAAGTGAAAAAAGGAGAGACTACAGATGGAGTGTACAGAGTGATGACTCGCAGACTGCTGGGTTCAACACAGGTTGGAGTGGGAGTCATGCAAGAGGGAGTCTTCCACACCATGTGGCACGTCACAAAAGGAGCCGCATTGAGGAGCGGTGAAGGAAGACTTGATCCATATTGGGGGGACGTCAAGCAGGACTTGGTGTCATATTGTGGGCCTTGGAAGTTGGATGCAGCCTGGGATGGACTAAGTGAAGTGCAGCTTTTGGCCGTACCCCCCGGAGAGAGGGCTAGAAACATTCAGACTCTGCCTGGAATATTCAAGACAAAGGATGGGGACATCGGAGCAGTCGCTCTAGACTACCCCGCAGGAACCTCAGGATCTCCAATCCTAGACAAATGCGGAAGAGTGATAGGACTTTATGGCAATGGGGTTGTGATCAAGAATGGAAGCTATGTTAGTGCCATAACCCAGGGAAAAAGGGAGGAGGAGGCTCCAGTTGAGTGCTTTGAACCCTCGATGCTGAGGAAGAAGCAGCTAACAGTCTTGGATCTGCATCCAGGAGCCGGGAAAACCAGGAGGGTTCTTCCTGAAATAGTCCGTGAAGCCATAAAGAAGAGACTTCGCACAGTGATCCTAGCACCAACCAGGGTCGTTGCCGCTGAGATGGAGGAAGCCCTAAGAGGACTGCCGGTGCGTTACATGACAACAGCAGTCAACGTCACCCACTCTGGGACAGAAATCGTCGATTTGATGTGCCATGCCACCTTCACTTCACGCCTACTACAACCCATCAGAGTCCCCAACTACAACCTTTACATCATGGATGAAGCTCATTTCACAGATCCCTCAAGCATAGCTGCACGAGGATATATATCAACAAGGGTTGAAATGGGCGAGGCGGCTGCTATCTTCATGACTGCTACACCACCAGGAACCCGCGATGCGTTTCCGGATTCCAACTCACCAATCATGGACACAGAAGTGGAAGTTCCAGAGAGAGCCTGGAGCTCAGGCTTTGACTGGGTGACGGATCATTCTGGGAAAACAATTTGGTTTGTTCCAAGTGTGAGAAACGGAAATGAAATCGCAGCCTGTCTGACAAAGGCTGGAAAGCGGGTCATACAGCTCAGCAGGAAGACTTTTGAGACAGAGTTTCAGAAGACAAAAAATCAAGAGTGGGACTTTGTCATAACAACTGACATTTCAGAGATGGGTGCCAATTTCAAGGCTGACCGGGTCATAGATTCCAGGAGATGCCTAAAGCCAGTCATACTCGATGGCGAGAGAGTCATCCTGGCTGGGCCCATGCCTGTCACGCATGCCAGTGCTGCTCAGAGGAGAGGACGTGTAGGCAGGAACCCCAACAAACCTGGAGATGAGTACATGTATGGAGGTGGGTGTGCAGAGACTGATGAAGACCATGCACACTGGCTTGAAGCAAGAATGCTTCTTGACAACATTTACCTCCAGGATGGCCTCATAGCCTCGCTCTATCGGCCTGAGGCCGACAAGGTAGCCGCCATTGAGGGAGAGTTCAAGCTGAGGACAGAGCAAAGGAAGACCTTTGTGGAACTCATGAAGAGAGGAGACCTTCCCGTTTGGCTGGCCTATCAAGTAGCATCTGCCGGAATAACTTACACAGATAGAAGATGGTGCTTTGATGGCACTACCAACAACACCATAATGGAAGACAGTGTACCAGCAGAGGTGTGGACCAAGTATGGAGAGAAGAGAGTGCTTAAACCGAGGTGGATGGATGCTAGGGTCTGTTCAGATCATGCGGCTCTGAAGTCGTTCAAAGAATTTGCCGCTGGGAAGAGAGGAGCGGCTTTGGGAGTAATGGATGCCCTGGGAACATTGCCAGGACACATGACAGAGAGGTTTCAGGAAGCCATTGACAATCTCGCTGTGCTCATGCGAGCAGAGACTGGAAGTAGGCCCTACAAAGCAGCGGCAGCTCAACTGCCGGAGACCTTAGAGACTATCATGCTTCTGGGCTTATTGGGAACAGTTTCGCTAGGAATCTTCTTTGTCTTGATGCGGAACAAGGGCATCGGGAAGATGGGCTTCGGAATGGTAACCCTTGGGGCCAGCGCATGGCTCATGTGGCTTTCGGAAATTGAACCAGCCAGAATCGCATGTGTCCTCATTGTCGTGTTTTTGTTGCTGGTGGTGCTCATACCCGAGCCAGAGAAGCAAAGATCTCCCCAGGATAATCAAATGGCAATCATCATCATGGTGGCAGTTGGCCTTCTGGGTTTGATAACTGCAAATGAACTCGGATGGCTGGAAAGGACAAAAAGTGATATAGCTCATCTAATGGGAAGGAAAGAAGAGGGGACAACCATGGGATTCTCAATGGATATTGATCTGCGGCCAGCCTCCGCCTGGGCTATTTATGCCGCATTGACAACTCTCATCACCCCAGCCGTCCAACATGCGGTAACCACCTCATACAACAACTACTCCCTGATGGCGATGGCCACACAAGCTGGAGTGCTGTTTGGCATGGGCAAAGGGATGCCATTTTATGCATGGG-ACTTTGGAGTCCCGCTGCTAATGATGGGTTGCTACTCACAATTAACACCCCTGACCCTGATAGTGGCCATCATTCTGCTTGTGGCACACTACATGTACTTGATCCCAGGTTTGCAGGCAGCAGCAGCACGTGCTGCCCAGAAGAGGACAGCAGCTGGCATCATGAAGAATCCCGTTGTGGATGGAATAGTGGTGACTGACATTGACACAATGACAATTGACCCCCAAGTGGAGAAGAAGATGGGACAAGTGTTACTCATAGCAGTAGCTGTCTCCAGTGCTGTGCTACTGCGGACCGCTTGGGGATGGGGGGAGGCTGGGGCTCTGATCACAGCAGCAACCTCCACCTTATGGGAAGGCTCTCCAAACAAATACTGGAACTCCTCCACAGCCACCTCACTGTGCAACATCTTTAGAGGAAGTTATTTGGCAGGGGCTTCCCTTATTTACACAGTGACAAGAAATGCCGGTCTGGTTAAGAGACGTGGAGGTGGAACGGGAGAGACCCTGGGAGAGAAGTGGAAAGCCCGCCTGAACCAGATGTCGGCCTTGGAGTTCTACTCTTACAAAAAGTCAGGCATCACCGAAGTGTGTAGGGAGGAGGCGCGCCGCGCCCTCAAGGATGGAGTGGCCACAGGAGGACATGCTGTATCCCGGGGAAGCGCAAAGCTTAGATGGTTGGTAGAGAGAGGATACCTGCAGCCCCATGGAAAGGTTGTTGACCTTGGATGTGGCAGAGGAGGCTGGAGTTATTACGCTGCCACCATCCGTAAAGTGCAGGAGGTCAGAGGATACACAAAGGGAGGTCCTGGCCATGAAGAGCCCATGCTGGTGCAAAGCTATGGGTGGAACATAGTTCGCCTCAAGAGTGGAGTGGACGTCTTCCACATGGCGGCTGAACCGTGTGACACTCTGCTGTGTGACATAGGCGAGTCATCATCCAGTCCTGAAGTGGAAGAGACGCGAACACTCAGAGTGCTCTCCATGGTGGGAGACTGGCTTGAAAAAAGACCAGGGGCCTTCTGCATAAAGGTGCTGTGCCCATACACCAGCACAATGATGGAGACCATGGAGCGACTGCAACGTAGGCATGGGGGAGGATTAGTCAGAGTGCCATTGTCCCGCAACTCTACACATGAGATGTATTGGGTCTCTGGAGCCAAAAGTAACATCATAAAGAGTGTGTCCACCACAAGTCAGCTCCTCTTGGGACGCATGGAAGGGCCTAGGAGGCCAGTGAAATATGAGGAGGATGTGAACCTCGGCTCAGGCACACGAGCTGTGGCAAGCTGCGCTGAGGCTCCCAACATGAAGATCATTGGTAGGCGCATTGAGAGAATCCGCAATGAACATGCAGAGACATGGTTCTTTGATGAAAACCACCCATACAGGACATGGGCCTACCATGGGAGCTACGAAGCCCCCACGCAGGGGTCAGCGTCATCCCTCGTGAACGGGGTTGTTAGACTCTTGTCAAAGCCCTGGGATGTGGTGACTGGAGTTACAGGAATAGCTATGACTGACACCACGCCATACGGCCAACAAAGAGTCTTCAAGGAAAAAGTGGACACCAGGGTGCCAGATCCCCAAGAAGGCACTCGCCAAGTGATGAACATGGTATCGTCTTGGCTATGGAAGGAGCTGGGAAAACGCAAGCGACCACGTGTCTGCACCAAAGAAGAGTTCATTAATAAGGTGCGCAGCAATGCAGCACTGGGAGCAATATTTGAAGAGGAAAAAGAATGGAAGACAGCTGTAGAAGCTGTGAATGATCCAAGATTTTGGGCTCTAGTGGACAAGGAAAGAGAACACCACCTGAGAGGAGAGTGTCATAGCTGTGTGTACAACATGATGGGCAAAAGAGAAAAGAAGCAAGGAGAATTCGGGAAAGCAAAAGGCAGCCGCGCAATCTGGTACATGTGGTTGGGAGCCAGATTTTTGGAGTTTGAAGCTCTTGGGTTCTTGAACGAGGACCACTGGATGGGGAGAGAAAACTCAGGAGGTGGCGTTGAAGGGCTAGGACTGCAAAGGCTTGGATATATCCTAGAAGAAATGAACCGGGCACCAGGAGGAAAGATGTATGCAGATGACACTGCTGGCTGGGACACCCGCATTAGCAAGTTTGATCTAGAGAATGAAGCCCTGATCACTAACCAGATGGAAGAAGGGCACAGAGCTCTGGCGTTGGCCGTGATTAAATACACATACCAAAACAAAGTGGTGAAGGTTCTCAGACCAGCTGAAGGAGGGAAAACAGTCATGGACATCATCTCAAGACAAGACCAGAGAGGGAGCGGACAAGTTGTTACTTATGCTCTCAACACATTCACCAACCTGGTGGTGCAGCTTATCCGGAACATGGAGGCTGAGGAAGTGCTAGAGATGCATGATCTGTGGTTGTTGAGGAAGCCAGAGAAAGTGACCAGATGGTTGCAGAGCAATGGATGGGACAGACTCAAACGGATGGCAGTTAGTGGAGATGACTGCGTTGTAAAGCCAATTGATGATAGGTTTGCACATGCCCTCAGGTTCTTGAATGACATGGGAAAGGTTAGGAAAGACACACAGGAATGGAAACCCTCGACTGGATGGAGCAATTGGGAAGAGGTCCCGTTCTGTTCTCACCACTTCAACAAGCTGCACCTCAAGGACGGGAGATCCATTGTGGTCCCCTGCCGCCACCAAGATGAACTGATTGGCCGAGCCCGTGTCTCACCAGGGGCAGGATGGAGCATCCGGGAGACTGCTTGTCTTGCAAAATCATATGCACAGATGTGGCAGCTTCTTTATTTCCACAGAAGAGACCTCCGACTGATGGCCAATGCCATTTGTTCGGCTGTGCCAATTGACTGGGTACCAACTGGGAGAACCACCTGGTCAATCCATGGAAAGGGAGAATGGATGACTACTGAGGACATGCTCATGGTGTGGAATAGAGTGTGGATTGAGGAGAACGACCACATGGAGGACAAGACCCCTGTAACAAAATGGACAGACATTCCCTATTTGGGAAAAAGGGAGGACTTATGGTGTGGATCCCTTATAGGGCATAGACCTCGCACCACTTGGGCTGAGAACATCAAAGACACAGTCAACATGGTGCGTAGGATCATAGGTGATGAAGAAAAGTTCATGGACTACCTATCCACCCAAGTACGCTACTTGGGTGAGGAAGGGTCCACACCTGGAGTGCTGTAA

>ENA|AMO03410|AMO03410.1 Zika virus polyprotein

ATGAAAAACCC-AAAAAAGAAATCCGGAGGATTCCGGATTGTCAATATGCTAAAACGCGGAGTAGCCCGTGTGAGCCCCTTTGGGGGCTTGAAGAGGCTGCCAGCCGGACTTCTGCTGGGTCATGGGCCCATCAGGATGGTCTTGGCGATTCTAGCCTTCTTGAGATTCACGGCAATCAAGCCATCACTGGGTCTCATCAATAGATGGGGTTCAGTGGGGAAAAAAGAGGCTATGGAAATAATAAAGAAGTTCAAGAAAGATCTGGCTGCCATGCTGAGAATAATCAATGCTAGGAAGGAGAAGAAGAGACGAGGCGCAGATACTAATGTCGGAATTGTTGGCCTCCTGCTGACCACAGCTATGGCAGCGGAGGTCACTAGACGTGGGAGTGCATACTATATGTACTTGGACAGAAACGATGCTGGGGAGGCCATATCTTTTCCAACCACATTGGGGATGAATAAGTGTTATATACAGATCATGGATCTTGGACACATGTGTGATGCCACCATGAGCTATGAATGCCCTATGCTGGATGAGGGGGTGGAACCAGATGACGTCGATTGTTGGTGCAACACGACGTCAACTTGGGTTGTGTACGGAACCTGCCATCACAAAAAAGGTGAAGCACGGAGATCTAGAAGAGCTGTGACGCTCCCCTCCCATTCCACTAGGAAGCTGCAAACGCGGTCGCAAACTTGGTTGGAATCAAGAGAATACACAAAGCACTTGATTAGAGTCGAAAATTGGATATTCAGGAACCCTGGCTTCGCGTTAGCAGCAGCTGCCATCGCTTGGCTTTTGGGAAGCTCAACGAGCCAAAAAGTCATATACTTGGTCATGATACTGCTGATTGCCCCGGCATACAGCATCAGGTGCATAGGAGTCAGCAATAGGGACTTTGTGGAAGGTATGTCAGGTGGGACTTGGGTTGATGTTGTCTTGGAACATGGAGGTTGTGTCACCGTAATGGCACAGGACAAACCGACTGTCGACATAGAGCTGGTTACAACAACAGTCAGCAACATGGCGGAGGTAAGATCCTACTGCTATGAGGCATCAATATCGGACATGGCTTCGGACAGCCGCTGCCCAACACAAGGTGAAGCCTACCTTGACAAGCAATCAGACACTCAATATGTCTGCAAAAGAACGTTAGTGGACAGAGGCTGGGGAAATGGATGTGGACTTTTTGGCAAAGGGAGCCTGGTGACATGCGCTAAGTTTGCATGCTCCAAGAAAATGACCGGGAAGAGCATCCAGCCAGAGAATCTGGAGTACCGGATAATGCTGTCAGTTCATGGCTCCCAGCACAGTGGGATGATCGTTAATGACACAGGACATGAAACTGATGAGAATAGAGCGAAGGTTGAGATAACGCCCAATTCACCAAGAGCCGAAGCCACCCTGGGGGGTTTTGGAAGCCTAGGACTTGATTGTGAACCGAGGACAGGCCTTGACTTTTCAGATTTGTATTACTTGACTATGAATAACAAGCACTGGTTGGTTCACAAGGAGTGGTTCCACGACATTCCATTACCTTGGCACGCTGGGGCAGACACCGGAACTCCACACTGGAACAACAAAGAAGCACTGGTAGAGTTCAAGGACGCACATGCCAAAAGGCAAACTGTCGTGGTTCTAGGGAGTCAAGAAGGAGCAGTTCACACGGCCCTTGCTGGAGCTCTGGAGGCTGAGATGGATGGTGCAAAGGGAAGGCTGTCCTCTGGCCACTTGAAATGTCGCCTGAAAATGGATAAACTTAGATTGAAGGGCGTGTCATACTCCTTGTGTACCGCAGCGTTCACATTCACCAAGATCCCGGCTGAAACACTGCACGGGACAGTCACAGTGGAGGTACAGTACGCAGGGACAGATGGACCTTGCAAGGTTCCAGCTCAGATGGCGGTGGACATGCAAACTCTGACCCCAGTTGGGAGGCTGATAACCGCTAACCCCGTAATCACTGAAAGCACTGAGAACTCCAAGATGATGCTGGAACTTGATCCACCATTTGGGGACTCTTACATTGTCATAGGAGTCGGGGAGAAGAAGATCACCCACCACTGGCACAGGAGTGGCAGCACCATTGGAAAAGCATTTGAAGCCACTGTGAGAGGTGCCAGGAGAATGGCAGTCTTGGGAGACACAGCCTGGGACTTTGGATCAGTTGGAGGCGCTCTCAACTCATTGGGCAAGGGCATCCATCAAATTTTTGGAGCAGCTTTCAAATCATTGTTTGGAGGAATGTCCTGGTTCTCACAAATTCTCATTGGAACGTTGCTGATGTGGTTGGGTCTGAACACAAAGAATGGATCTATTTCCCTTATGTGCTTGGCCTTAGGGGGAGTGTTGATCTTCTTATCCACAGCCGTCTCTGCTGATGTGGGGTGCTCGGTGGACTTCTCAAAGAAGGAGACGAGATGCGGTACAGGGGTGTTCGTCTATAACGACGTTGAAGCCTGGAGGGACAGGTACAAGTACCATCCTGACTCCCCCCGTAGATTGGCAGCAGCAGTCAAGCAAGCCTGGGAAGATGGTATCTGTGGGATCTCCTCTGTTTCAAGAATGGAAAACATCATGTGGAGATCAGTAGAAGGGGAGCTCAACGCAATCCTGGAAGAGAATGGAGTTCAACTGACGGTCGTTGTGGGATCTGTAAAAAACCCCATGTGGAGAGGTCCACAGAGATTGCCCGTGCCTGTGAACGAGCTGCCCCACGGCTGGAAGGCTTGGGGGAAATCGTACTTCGTCAGAGCAGCAAAGACAAATAACAGCTTTGTCGTGGATGGTGACACACTGAAGGAATGCCCACTCAAACATAGAGCATGGAACAGCTTTCTTGTGGAGGATCATGGGTTCGGGGTATTTCACACTAGTGTCTGGCTCAAGGTTAGAGAAGATTATTCATTAGAGTGTGATCCAGCCGTTATTGGAACAGCTGTTAAGGGAAAGGAGGCTGTACACAGTGATCTAGGCTACTGGATTGAGAGTGAGAAGAATGACACATGGAGGCTGAAGAGGGCCCATCTGATCGAGATGAAAACATGTGAATGGCCAAAGTCCCACACATTGTGGACAGATGGAATAGAAGAGAGTGATCTGATCATACCCAAGTCTTTAGCTGGGCCACTCAGCCATCACAATACCAGAGAGGGCTACAGGACCCAAATGAAAGGGCCATGGCACAGTGAAGAGCTTGAAATTCGGTTTGAGGAATGCCCAGGCACCAAGGTCCACGTGGAGGAAACATGTGGAACAAGAGGACCATCTCTGAGATCAACCACAGCAAGCGGAAGGGTGATCGAGGAATGGTGCTGCAGGGAGTGCACAATGCCCCCACTGTCGTTCCAGGCTAAAGATGGCTGTTGGTATGGAATGGAGATAAGGCCCAGGAAAGAACCAGAAAGTAACTTAGTAAGGTCAATGGTGACTGCAGGATCAACTGATCACATGGATCACTTCTCCCTTGGAGTGCTTGTGATTCTGCTCATGGTGCAGGAAGGGCTGAAGAAGAGAATGACCACAAAGATCATCATAAGCACATCAATGGCAGTGCTGGTAGCTATGATCCTGGGAGGATTTTCAATGAGTGACCTGGCTAAGCTTGCAATTTTGATGGGTGCCACCTTCGCGGAAATGAACACTGGAGGAGATGTAGCTCATCTGGCGCTGATAGCGGCATTCAAAGTCAGACCAGCGTTGCTGGTATCTTTCATCTTCAGAGCTAATTGGACACCCCGTGAAAGCATGCTGCTGGCCTTGGCCTCGTGTCTTTTGCAAACTGCGATCTCCGCCTTGGAAGGCGACCTGATGGTTCTCATCAATGGTTTTGCTTTGGCCTGGTTGGCAATACGAGCGATGGTTGTTCCACGCACTGATAACATCACCTTGGCAATCCTGGCTGCTCTGACACCACTGGCCCGGGGCACACTGCTTGTGGCGTGGAGAGCAGGCCTTGCTACTTGCGGGGGGTTTATGCTCCTCTCTCTGAAGGGAAAAGGCAGTGTGAAGAAGAACTTACCATTTGTCATGGCCCTGGGACTAACCGCTGTGAGGCTGGTCGACCCCATCAACGTGGTGGGACTGCTGTTGCTCACAAGGAGTGGGAAGCGGAGCTGGCCCCCTAGCGAAGTACTCACAGCTGTTGGCCTGATATGCGCATTGGCTGGAGGGTTCGCCAAGGCAGATATAGAGATGGCTGGGCCCATGGCCGCGGTCGGTCTGCTAATTGTCAGTTACGTGGTCTCAGGAAAGAGTGTGGACATGTACATTGAAAGAGCAGGTGACATCACATGGGAAAAAGATGCGGAAGTCACTGGAAACAGTCCCCGGCTTGATGTGGCGCTAGATGAGAGTGGTGATTTCTCCCTGGTGGAGGATGACGGTCCCCCCATGAGAGAGATCATACTCAAGGTGGTCCTGATGACCATCTGTGGCATGAACCCAATAGCCATACCCTTTGCAGCTGGAGCGTGGTACGTATACGTGAAGACTGGAAAAAGGAGTGGAGCTCTATGGGATGTGCCTGCTCCCAAGGAAGTAAAAAAGGGGGAGACCACAGATGGAGTGTACAGAGTGATGACTCGTAGACTGCTAGGTTCAACACAAGTTGGAGTGGGAGTTATGCAAGAGGGGGTCTTTCACACCATGTGGCACGTCACAAAAGGATCCGCGCTGAGAAGCGGTGAAGGGAGACTTGATCCATACTGGGGAGATGTCAAGCAGGATCTGGTGTCATACTGTGGTCCATGGAAGCTAGATGCCGCCTGGGACGGGCACAGCGAGGTGCAGCTCTTGGCCGTGCCCCCCGGAGAGAGAGCGAGGAACATCCAGACTCTGCCCGGAATATTTAAGACAAAGGATGGGGACATTGGAGCGGTTGCGCTGGATTACCCAGCAGGAACTTCAGGATCTCCAATCCTAGACAAGTGTGGGAGAGTGATAGGACTTTATGGCAATGGGGTCGTGATCAAAAATGGGAGTTATGTTAGTGCCATCACCCAAGGGAGGAGGGAGGAAGAGACTCCTGTTGAGTGCTTCGAGCCTTCGATGCTGAAGAAGAAGCAGCTAACTGTCTTAGACTTGCATCCTGGAGCTGGGAAAACCAGGAGAGTTCTTCCTGAAATAGTCCGTGAAGCCATAAAAACAAGACTCCGTACTGTGATCTTAGCTCCAACCAGGGTTGTCGCTGCCGAAATGGAGGAAGCCCTTAGAGGGCTTCCAGTGCGTTATATGACAACAGCAGTCAATGTCACCCACTCTGGAACAGAAATCGTCGACTTAATGTGCCATGCCACCTTCACTTCACGTCTACTACAGCCAATCAGAGTCCCCAACTATAATCTGTATATTATGGATGAGGCCCACTTCACAGATCCCTCAAGTATAGCAGCAAGAGGATACATTTCAACAAGGGTTGAGATGGGCGAGGCGGCTGCCATCTTCATGACCGCCACGCCACCAGGAACCCGTGACGCATTTCCGGACTCCAACTCACCAATTATGGACACCGAAGTGGAAGTCCCAGAGAGAGCCTGGAGCTCAGGCTTTGATTGGGTGACGGATCATTCTGGAAAAACAGTCTGGTTTGTTCCAAGCGTGAGGAACGGCAATGAGATCGCAGCTTGTCTGACAAAGGCTGGAAAACGGGTCATACAGCTCAGCAGAAAGACTTTTGAGACAGAGTTCCAGAAAACAAAACATCAAGAGTGGGACTTTGTCGTGACAACTGACATTTCAGAGATGGGCGCCAACTTTAAAGCTGACCGTGTCATAGATTCCAGGAGATGCCTAAAGCCGGTCATACTTGATGGCGAGAGAGTCATTCTGGCTGGACCCATGCCTGTCACACATGCCAGCGCTGCCCAGAGGAGGGGGCGCATAGGCAGGAATCCCAACAAACCTGGAGATGAGTATCTGTATGGAGGTGGGTGCGCAGAGACTGACGAAGACCATGCACACTGGCTTGAAGCAAGAATGCTCCTTGACAATATTTACCTCCAAGATGGCCTCATAGCCTCGCTCTATCGACCTGAGGCCGACAAAGTAGCAGCCATTGAGGGAGAGTTCAAGCTTAAGACGGAGCAAAGGAAGACCTTTGTGGAACTCATGAAAAGAGGAGATCTTCCTGTTTGGCTGGCCTATCAGGTTGCATCTGCCGGAATAACCTACACAGATAGAAGATGGTGCTTTGATGGCACGACCAACAACACCATAATGGAAGACAGTGTGCCGGCAGAAGTGTGGACCAGACACGGAGAGAAAAGAGTGCTCAAACCGAGGTGGATGGACGCCAGAGTTTGTTCAGATCATGCGGCCCTGAAGTCATTCAAAGAGTTTGCCGCTGGGAAAAGAGGAGCGGCCTTTGGAGTGATGGAAGCCCTGGGAACACTGCCAGGACATATGACAGAGAGATTCCAGGAGGCCATTGACAACCTCGCTGTGCTCATGCGGGCAGAGACTGGAAGCAGGCCCTACAAAGCCGCGGCGGCCCAATTACCGGAGACCCTAGAGACTATCATGCTTTTGGGGTTGCTGGGAACAGTCTCGCTGGGAATCTTTTTCGTCTTGATGCGGAACAAGGGCATAGGGAAGATGGGCTTTGGAATGGTGACTCTTGGGGCCAGCGCATGGCTTATGTGGCTCTCGGAAATTGAGCCAGCCAGAATTGCATGTGTCCTCATTGTTGTGTTCCTATTGCTGGTGGTGCTCATACCTGAGCCAGAAAAGCAAAGATCTCCCCAGGACAACCAAATGGCAATCATCATCATGGTAGCAGTGGGTCTTCTGGGCTTGATTACCGCCAATGAACTCGGATGGTTGGAGAGAACAAAGAGTGACCTAAGCCATCTAATGGGAAGGAGAGAGGAGGGGGCAACCATAGGATTCTCAATGGACATTGACCTGCGGCCAGCCTCAGCTTGGGCCATCTACGCTGCCTTGACAACTTTCATTACCCCAGCCGTCCAACATGCAGTGACCACTTCATACAACAACTACTCCTTAATGGCGATGGCCACGCAAGCTGGAGTGTTGTTTGGTATGGGCAAAGGGATGCCATTCTACGCATGGG-ACTTTGGAGTCCCGCTGCTAATGATAGGTTGCTACTCACAATTAACACCCCTGACCCTAATAGTAGCCATCATTTTGCTCGTGGCGCACTACATGTACTTGATCCCAGGGCTGCAGGCAGCAGCTGCGCGTGCTGCCCAGAAGAGAACGGCAGCTGGCATCATGAAGAACCCTGTTGTGGATGGAATAGTGGTGACTGACATTGACACAATGACAATTGACCCCCAAGTGGAGAAAAAGATGGGACAGGTGCTACTCATAGCAGTAGCCGTCTCCAGCGCCATACTGTCGCGGACCGCCTGGGGGTGGGGGGAGGCTGGGGCCCTGATCACAGCTGCAACTTCCACTTTGTGGGAAGGCTCTCCGAACAAGTACTGGAACTCCTCTACAGCCACTTCACTGTGTAACATTTTTAGGGGAAGTTACTTGGCTGGAGCTTCTCTAATCTACACAGTAACAAGAAACGCTGGCTTGGTCAAGAGACGTGGGGGTGGAACAGGAGAGACCCTGGGAGAGAAATGGAAGGCCCGCTTGAACCAGATGTCGGCCCTGGAGTTCTACTCCTACAAAAAGTCAGGCATCACCGAGGTGTGCAGAGAAGAGGCCCGCCGCGCCCTCAAGGACGGTGTGGCAACGGGAGGCCATGCTGTGTCCCGAGGAAGTGCAAAGCTGAGATGGTTGGTGGAGCGGGGATACCTGCAGCCCTATGGAAAGGTCATTGATCTTGGATGTGGCAGAGGGGGCTGGAGTTACTACGCCGCCACCATCCGCAAAGTTCAAGAAGTGAAAGGATACACAAAAGGAGGCCCTGGTCATGAAGAACCCATGTTGGTGCAAAGCTATGGGTGGAACATAATCCGTCTTAAGAGTGGGGTGGACGTCTTTCATATGGCGGCTGAGCCGTGTGACACGTTGCTGTGTGACATAGGTGAGTCATCATCTAGTCCTGAAGTGGAAGAAGCACGGACGCTCAGAGTCCTTTCCATGGTGGGGGATTGGCTTGAAAAAAGACCAGGAGCCTTTTGTATAAAAGTGTTGTGTCCATACACCAGCACTATGATGGAAACCCTGGAGCGACTGCAGCGTAGGTATGGGGGAGGACTGGTCAGAGTGCCACTCTCCCGCAACTCTACACATGAGATGTACTGGGTCTCTGGAGCGAAAAGCAACACCATAAAAAGTGTGTCCACCACGAGCCAGCTCCTCTTGGGGCGCATGGACGGGCCCAGGAGGCCAGTGAAATATGAGGAGGATGTGAATCTCGGCTCTGGCACGCGGGCTGTGGTAAGCTGCGCTGAAGCTCCCAACATGAAGATCATTGGTAACCGCATTGAAAGGATCCGCAGTGAGCACGCGGAAACGTGGTTCTTTGACGAGAACCACCCATATAGGACATGGGCTTACCATGGAAGCTATGAGGCCCCCACACAAGGGTCAGCGTCCTCTCTAATAAACGGGGTTGTCAGGCTCCTGTCAAAACCCTGGGATGTGGTGACTGGAGTCACAGGAATAGCCATGACCGACACCACACCGTATGGTCAGCAAAGAGTTTTCAAGGAAAAAGTGGACACTAGGGTGCCAGATCCCCAAGAAGGCACTCGTCAGGTTATGAGCATGGTCTCTTCCTGGTTGTGGAAAGAGCTAGGCAAACACAAACGGCCACGAGTCTGTACCAAAGAAGAGTTCATCAACAAGGTTCGTAGCAATGCAGCATTAGGGGCAATATTTGAAGAGGAAAAAGAGTGGAAGACTGCAGTGGAAGCTGTGAACGATCCAAGGTTCTGGGCTCTAGTGGACAAGGAAAGAGAGCACCACCTGAGAGGAGAGTGCCAGAGTTGTGTGTACAACATGATGGGAAAAAGAGAAAAGAAACAAGGGGAATTTGGAAAGGCCAAGGGCAGCCGCGCCATCTGGTATATGTGGCTAGGGGCTAGATTTCTAGAGTTCGAAGCCCTTGGATTCTTGAACGAGGATCACTGGATGGGGAGAGAGAACTCAGGAGGTGGTGTTGAAGGGCTGGGATTACAAAGACTCGGATATGTCCTAGAAGAGATGAGTCGCATACCAGGAGGAAGGATGTATGCAGATGACACTGCTGGCTGGGACACCCGCATCAGCAGGTTTGATCTGGAGAATGAAGCTCTAATCACCAACCAAATGGAGAAAGGGCACAGGGCCTTGGCATTGGCCATAATCAAGTACACATACCAAAACAAAGTGGTAAAGGTCCTTAGACCAGCTGAAAAAGGGAAGACAGTTATGGACATTATTTCGAGACAAGACCAAAGGGGGAGCGGACAAGTTGTCACTTACGCTCTTAACACATTTACCAACCTAGTGGTGCAACTCATTCGGAGTATGGAGGCTGAGGAAGTTCTAGAGATGCAAGACTTGTGGCTGCTGCGGAGGTCAGAGAAAGTGACCAACTGGCTGCAGAGCAACGGATGGGATAGGCTCAAACGAATGGCAGTCAGTGGAGATGATTGCGTTGTGAGGCCAATTGATGATAGGTTTGCACATGCCCTCAGGTTCTTGAATGATATGGGGAAAGTTAGGAAGGACACACAAGAGTGGAAACCCTCAACTGGATGGGACAACTGGGAGGAAGTTCCGTTTTGCTCCCACCACTTCAACAAGCTCCATCTCAAGGACGGGAGGTCCATTGTGGTTCCCTGCCGCCACCAAGATGAACTGATTGGCCGAGCTCGCGTCTCACCGGGGGCGGGATGGAGCATCCGGGAGACTGCTTGCCTAGCAAAATCATATGCGCAAATGTGGCAGCTCCTTTATTTCCACAGAAGGGACCTCCGACTGATGGCCAATGCCATTTGTTCATCTGTGCCAGTTGACTGGGTTCCAACTGGGAGAACTACCTGGTCAATCCATGGAAAGGGAGAATGGATGACCACTGAAGACATGCTTGTGGTGTGGAACAGAGTGTGGATTGAGGAGAACGACCACATGGAAGACAAGACCCCAGTTACGAAATGGACAGACATTCCCTATTTGGGAAAAAGGGAAGACTTGTGGTGTGGGTCTCTCATAGGGCACAGACCGCGCACCACCTGGGCTGAGAACATTAAAAACACAGTCAACATGATGCGTAGGATCATAGGTGATGAAGAAAAGTACGTGGACTACCTATCCACCCAAGTTCGCTACTTGGGCGAAGAAGGGTCCACACCTGGAGTGCTATAA

>ENA|AMQ48981|AMQ48981.1 Zika virus polyprotein

ATGAAAAACCC-AAAAAAGAAATCCGGAGGATTCCGGATTGTCAATATGCTAAAACGCGGAGTAGCCCGTGTGAGCCCCTTTGGGGGCTTGAAGAGGCTGCCAGCCGGACTTCTGCTGGGTCATGGGCCCATCAGGATGGTCTTGGCGATTCTAGCCTTTTTGAGATTCACGGCAATCAAGCCATCACTGGGTCTCATCAATAGATGGGGTTCAGTGGGGAAAAAAGAGGCTATGGAAATAATAAAGAAGTTCAAGAAAGATCTGGCTGCCATGCTGAGAATAATCAATGCTAGGAAGGAGAAGAAGAGACGAGGCGCAGATACTAGTGTCGGAATTGTTGGCCTCCTGCTGACCACAGCTATGGCAGCGGAGGTCACTAGACGTGGGAGTGCATACTACATGTACTTGGACAGAAACGATGCTGGGGAGGCCATATCTTTTCCAACCACATTGGGGATGAATAAGTGTTATATACAGATCATGGATCTTGGACACATGTGTGATGCCACCATGAGCTATGAATGCCCTATGCTGGATGAGGGGGTGGAACCAGATGACGTCGATTGTTGGTGCAACACGACGTCAACTTGGGTTGTGTACGGAACCTGCCATCACAAAAAAGGTGAAGCACGGAGATCTAGAAGAGCTGTGACGCTCCCCTCCCATTCCACTAGGAAGCTGCAAACGCGGTCGCAAACCTGGTTGGAATCGAGAGAATACACAAAGCACTTGATTAGAGTCGAAAATTGGATATTCAGGAACCCTGGCTTCGCGTTAGCAGCAGCTGCCATCGCTTGGCTTTTGGGAAGCTCAACGAGCCAAAAAGTCATATACTTGGTCATGATACTGCTGATTGCCCCGGCATACAGCATCAGGTGCATAGGAGTCAGCAATAGGGACTTTGTGGAAGGTATGTCAGGTGGGACTTGGGTTGATGTTGTCTTGGAACATGGAGGTTGTGTCACCGTAATGGCACAGGATAAACCGACTGTCGACATAGAGCTGGTTACAACAACAGTCAGCAACATGGCGGAGGTAAGATCCTACTGCTATGAGGCATCAATATCAGACATGGCTTCGGACAGCCGCTGCCCAACACAAGGTGAAGCCTACCTTGACAAGCAATCAGACACTCAATATGTCTGCAAAAGAACGTTAGTGGACAGAGGCTGGGGAAATGGATGTGGACTTTTTGGCAAAGGGAGCCTGGTGACATGCGCTAAGTTTGCATGCTCCAAGAAAATGACCGGGAAGAGCATCCAGCCAGAGAATCTGGAGTACCGGATAATGCTGTCAGTTCATGGCTCCCAGCACAGTGGGATGATCGTTAATGACACAGGACATGAAACTGATGAGAATAGAGCGAAGGTTGAGATAACGCCCAATTCACCAAGAGCCGAAGCCACCCTGGGGGGTTTTGGAAGCCTAGGACTTGATTGTGAACCGAGGACAGGCCTTGACTTTTCAGATTTGTATTACTTGACTATGAATAACAAGCACTGGTTGGTTCACAAGGAGTGGTTCCACGACATTCCATTGCCTTGGCACGCTGGGGCAGACACCGGAACTCCACACTGGAACAACAAAGAAGCACTGGTAGAGTTCAAGGACGCACATGCCAAAAGGCAAACTGTCGTGGTTCTAGGGAGTCAAGAAGGAGCAGTTCACACGGCCCTTGCTGGAGCTCTGGAGGCTGAGATGGATGGTGCAAAGGGAAGGCTGTCCTCTGGCCACTTGAAATGTCGCCTGAAAATGGATAAACTTAGATTGAAGGGCGTGTCATACTCCTTGTGTACCGCAGCGTTCACATTCACCAAGATCCCGGCTGAAACACTGCACGGGACAGTCACAGTGGAGGTACAGTACGCAGGGACAGATGGACCTTGCAAGGTTCCAGCTCAGATGGCGGTGGACATGCAAACTCTGACCCCAGTTGGGAGGTTGATAACCGCTAACCCCGTAATCACTGAAAGCACTGAGAACTCTAAGATGATGCTGGAACTTGATCCACCATTTGGGGACTCTTACATTGTCATAGGAGTCGGGGAGAAGAAGATCACCCACCACTGGCACAGGAGTGGCAGCACCATTGGAAAAGCATTTGAAGCCACTGTGAGAGGTGCCAAGAGAATGGCAGTCTTGGGAGACACAGCCTGGGACTTTGGATCAGTTGGAGGCGCTCTCAACTCATTGGGCAAGGGCATCCATCAAATTTTTGGAGCAGCTTTCAAATCATTGTTTGGAGGAATGTCCTGGTTCTCACAAATTCTCATTGGAACGTTGCTGATGTGGTTGGGCCTGAACACAAAGAATGGATCTATTTCCCTTATGTGCTTGGCCTTAGGGGGAGTGTTGATCTTCTTATCCACAGCCGTCTCTGCTGATGTGGGGTGCTCGGTGGACTTCTCAAAGAAGGAGACGAGATGCGGTACAGGGGTGTTCGTCTATAACGACGTTGAAGCCTGGAGGGACAGGTACAAGTACCATCCTGACTCCCCCCGTAGATTGGCAGCAGCAGTCAAGCAAGCCTGGGAAGATGGTATCTGCGGGATCTCCTCTGTTTCAAGAATGGAAAACATCATGTGGAGATCAGTAGAAGGGGAGCTCAACGCAATCCTGGAAGAGAATGGAGTTCAACTGACGGTCGTTGTGGGATCGGTAAAAAACCCCATGTGGAGAGGTCCACAGAGATTGCCCGTGCCTGTGAACGAGCTGCCCCACGGCTGGAAGGCTTGGGGGAAATCGTACTTCGTCAGAGCAGCAAAGACAAATAACAGCTTTGTCGTGGATGGTGACACACTGAAGGAATGCCCACTCAAACATAGAGCATGGAACAGCTTTCTTGTGGAGGATCATGGGTTCGGGGTATTTCACACTAGTGTCTGGCTCAAGGTTAGAGAAGATTATTCATTAGAGTGTGATCCAGCCGTTATTGGAACAGCTGTTAAGGGAAAGGAGGCTGTACACAGTGATCTAGGCTACTGGATTGAGAGTGAGAAGAATGACACATGGAGGCTGAAGAGGGCCCATCTGATCGAGATGAAAACATGTGAATGGCCAAAGTCCCACACATTGTGGACAGATGGAATAGAAGAGAGTGATCTGATCATACCCAAGTCTTTAGCTGGGCCACTCAGCCATCACAATACCAGAGAGGGCTACAGGACCCAAATGAAAGGGCCATGGCACAGTGAAGAGCTTGAAATTCGGTTTGAGGAATGCCCAGGCACTAAGGTCCACGTGGAGGAAACATGTGGAACAAGAGGACCATCTCTGAGATCAACCACTGCAAGCGGAAGGGTGATCGAGGAATGGTGCTGCAGGGAGTGCACAATGCCCCCACTGTCGTTCCGGGCTAAAGATGGCTGTTGGTATGGAATGGAGATAAGGCCCAGGAAAGAACCAGAAAGCAACTTAGTAAGGTCAATGGTGACTGCAGGATCAACTGATCACATGGATCACTTCTCCCTTGGAGTGCTTGTGATTCTGCTCATGGTGCAGGAAGGGCTGAAGAAGAGAATGACCACAAAGATCATCATAAGCACATCAATGGCAGTGCTGGTAGCTATGATCCTGGGAGGATTTTCAATGAGTGACCTGGCTAAGCTTGCAATTTTGATGGGTGCCACCTTCGCGGAAATGAACACTGGAGGAGATGTAGCTCATCTGGCGCTGATAGCGGCATTCAAAGTCAGACCAGCGTTGCTGGTATCTTTCATCTTCAGAGCTAATTGGACACCCCGTGAAAGCATGCTGCTGGCCTTGGCCTCGTGTCTTTTGCAAACTGCGATCTCCGCCTTGGAAGGCGACCTGATGGTTCTCATCAATGGTTTTGCTTTGGCCTGGTTGGCAATACGAGCGATGGTTGTTCCACGCACTGATAACATCACCTTGGCAATCCTGGCTGCTCTGACACCACTGGCCCGGGGCACACTGCTTGTGGCGTGGAGAGCAGGCCTTGCTACTTGCGGGGGGTTTATGCTCCTCTCTCTGAAGGGAAAAGGCAGTGTGAAGAAGAACTTACCATTTGTCATGGCCCTGGGACTAACCGCTGTGAGGCTGGTCGACCCCATCAACGTGGTGGGACTGCTGTTGCTCACAAGGAGTGGGAAGCGGAGCTGGCCCCCTAGCGAAGTACTCACAGCTGTTGGCCTGATATGCGCATTGGCTGGAGGGTTCGCCAAGGCAGATATAGAGATGGCTGGGCCCATAGCCGCGGTCGGTCTGCTAATTGTCAGTTACGTGGTCTCAGGAAAGAGTGTGGACATGTACATTGAAAGAGCAGGTGACATCACATGGGAAAAAGATGCGGAAGTCACTGGAAACAGTCCCCGGCTCGATGTGGCGCTAGATGAGAGTGGTGATTTCTCCCTGGTGGAGGATGACGGTCCCCCCATGAGAGAGATCATACTCAAGGTGGTCCTGATGACCATCTGTGGCATGAACCCAATAGCCATACCTTTTGCAGCTGGAGCGTGGTACGTATACGTGAAGACTGGAAAAAGGAGTGGTGCTCTATGGGATGTGCCTGCTCCCAAGGAAGTAAAAAAGGGGGAGACCACAGATGGAGTGTACAGAGTAATGACTCGTAGACTGCTGGGTTCAACACAAGTTGGAGTGGGAGTTATGCAAGAGGGGGTCTTTCACACTATGTGGCACGTCACAAAAGGATCCGCACTGAGAAGCGGTGAAGGGAGACTTGATCCATACTGGGGAGATGTCAAGCAGGATCTGGTGTCATACTGTGGTCCATGGAAGCTAGATGCCGCCTGGGACGGGCACAGCGAGGTGCAGCTCTTGGCCGTGCCCCCCGGAGAGAGAGCGAGGAACATCCAGACTCTGCCCGGAATATTTAAGACAAAGGATGGGGACATTGGAGCGGTTGCGCTGGATTACCCAGCAGGAACTTCAGGATCTCCAATCCTAGACAAGTGTGGGAGAGTGATAGGACTTTATGGCAATGGGGTCGTGATCAAAAATGGGAGTTATGTTAGTGCCATCACCCAAGGGAGGAGGGAGGAAGAGACTCCTGTTGAGTGCTTCGAGCCTTCGATGCTGAAGAAGAAGCAGCTAACTGTCTTAGACTTGCATCCTGGAGCTGGGAAAACCAGGAGAGTTCTTCCTGAAATAGTCCGTGAAGCCATAAAAACAAGACTCCGTACTGTGATCTTAGCTCCAACCAGGGTTGTCGCTGCTGAAATGGAGGAGGCCCTTAGAGGGCTTCCAGTGCGTTATATGACAACAGCAGTCAATGTCACCCACTCTGGAACAGAAATCGTCGACTTAATGTGCCATGCCACCTTCACTTCACGTCTACTACAGCCAATCAGAGTCCCCAACTATAATCTGTATATTATGGATGAGGCCCACTTCACAGATCCCTCAAGTATAGCAGCAAGAGGATACATTTCAACAAGGGTTGAGATGGGCGAGGCGGCTGCCATCTTCATGACCGCCACGCCACCAGGAACCCGTGACGCATTTCCGGACTCCAACTCACCAATTATGGACACCGAAGTGGAAGTCCCAGAGAGAGCCTGGAGCTCAGGCTTTGATTGGGTGACGGATCATTCTGGAAAAACAGTTTGGTTTGTTCCAAGCGTGAGGAACGGCAATGAGATCGCAGCTTGTCTGACAAAGGCTGGAAAACGGGTCATACAGCTCAGCAGAAAGACTTTTGAGACAGAGTTCCAGAAAACAAAACATCAAGAGTGGGACTTTGTCGTGACAACTGACATTTCAGAGATGGGCGCCAACTTTAAAGCTGACCGTGTCATAGATTCCAGGAGATGCCTAAAGCCGGTCATACTTGATGGCGAGAGAGTCATTCTGGCTGGACCCATGCCTGTCACACATGCCAGCGCTGCCCAGAGGAGGGGGCGCATAGGCAGGAATCCCAACAAACCTGGAGATGAGTATCTGTATGGAGGTGGGTGCGCAGAGACTGACGAAGACCATGCACACTGGCTTGAAGCAAGAATGCTCCTTGACAATATTTACCTCCAAGATGGCCTCATAGCCTCGCTCTATCGACCTGAGGCCGACAAAGTAGCAGCCATTGAGGGAGAGTTCAAGCTTAGGACGGAGCAAAGGAAGACCTTTGTGGAACTCATGAGAAGAGGAGATCTTCCTGTTTGGCTGGCCTATCAGGTTGCATCTGCCGGAATAACCTACACAGATAGAAGATGGTGCTTTGATGGCACGACCAACAACACCATAATGGAAGACAGTGTGCCGGCAGAGGTGTGGACCAGACACGGAGAGAAAAGAGTGCTCAAACCGAGGTGGATGGACGCCAGAGTTTGTTCAGATCATGCGGCCCTGAAGTCATTCAAGGAGTTTGCCGCTGGGAAAAGAGGAGCGGCTTTTGGAGTGATGGAAGCCCTGGGAACACTGCCAGGACACATGACAGAGAGATTCCAGGAAGCCATTGACAACCTCGCTGTGCTCATGCGGGCAGAGACTGGAAGCAGGCCTTACAAAGCCGCGGCGGCCCAATTGCCGGAGACCCTAGAGACCATTATGCTTTTGGGGTTGTTGGGAACAGTCTCGCTGGGAATCTTTTTCGTCTTGATGAGGAACAAGGGCATAGGGAAGATGGGCTTTGGAATGGTGACTCTTGGGGCCAGCGCATGGCTCATGTGGCTCTCGGAAATTGAGCCAGCCAGAATTGCATGTGTCCTCATTGTTGTGTTCCTATTGCTGGTGGTGCTCATACCTGAGCCAGAAAAGCAAAGATCTCCCCAGGACAACCAAATGGCAATCATCATCATGGTAGCAGTAGGTCTTCTGGGCTTGATTACCGCCAATGAACTCGGATGGTTGGAGAGAACAAAGAGTGACCTAAGCCATCTAATGGGAAGGAGAGAGGAGGGGGCAACCATAGGATTCTCAATGGACATTGACCTGCGGCCAGCCTCAGCTTGGGCCATCTATGCTGCCTTGACAACTTTCATTACCCCAGCCGTCCAACATGCAGTGACCACTTCATACAACAACTACTCCTTAATGGCGATGGCCACGCAAGCTGGAGTGTTGTTTGGTATGGGCAAAGGGATGCCATTCTACGCATGGG-ACTTTGGAGTCCCGCTGCTAATGATAGGTTGCTACTCACAATTAACACCCCTGACCCTAATAGTGGCCATCATTTTGCTCGTGGCGCACTACATGTACTTGATCCCAGGGCTGCAGGCAGCAGCTGCGCGTGCTGCCCAGAAGAGAACGGCAGCTGGCATCATGAAGAACCCTGTTGTGGATGGAATAGTGGTGACTGACATTGACACAATGACAATTGACCCCCAAGTGGAGAAAAAGATGGGACAGGTGCTACTCATAGCAGTAGCCGTCTCCAGCGCCATACTGTCGCGGACCGCCTGGGGGTGGGGGGAGGCTGGGGCCCTGATCACAGCCGCAACTTCCACTTTGTGGGAAGGCTCTCCGAACAAGTACTGGAACTCCTCTACAGCCACTTCACTGTGTAACATTTTTAGGGGAAGTTACTTGGCTGGAGCTTCTCTAATCTACACAGTAACAAGAAACGCTGGCTTGGTCAAGAGACGTGGGGGTGGAACAGGAGAGACCCTGGGAGAGAAATGGAAGGCCCGCTTGAACCAGATGTCGGCCCTGGAGTTCTACTCCTACAAAAAGTCAGGCATCACCGAGGTGTGCAGAGAAGAGGCCCGCCGCGCCCTCAAGGACGGTGTGGCAACGGGAGGCCATGCTGTGTCCCGAGGAAGTGCAAAGCTGAGATGGTTGGTGGAGCGGGGATACCTGCAGCCCTACGGAAAGGTCATTGATCTTGGATGTGGCAGAGGGGGCTGGAGTTACTACGCCGCCACCATCCGCAAAGTTCAAGAAGTGAAAGGATACACAAAAGGAGGCCCTGGTCATGAAGAACCCGTGTTGGTGCAAAGCTATGGGTGGAACATAGTCCGTCTTAAGAGTGGGGTGGACGTCTTTCATATGGCGGCTGAGCCGTGTGACACGTTGCTGTGTGACATAGGTGAGTCATCATCTAGTCCTGAAGTGGAAGAAGCACGGACGCTCAGAGTCCTCTCCATGGTGGGGGATTGGCTTGAAAAAAGACCAGGAGCCTTTTGTATAAAAGTGTTGTGCCCATACACCAGCACTATGATGGAAACCCTGGAGCGACTGCAGCGTAGGTATGGGGGAGGACTGGTCAGAGTGCCACTCTCCCGCAACTCTACACATGAGATGTACTGGGTCTCTGGAGCGAAAAGCAACACCATAAAAAGTGTGTCCACCACGAGCCAGCTCCTCTTGGGGCGCATGGACGGGCCTAGGAGGCCAGTGAAATATGAGGAGGATGTGAATCTCGGCTCTGGCACGCGGGCTGTGGTAAGCTGCGCTGAAGCTCCCAACATGAAGATCATTGGTAACCGCATTGAAAGGATCCGCAGTGAGCACGCGGAAACGTGGTTCTTTGACGAGAACCACCCATATAGGACATGGGCTTACCATGGAAGCTATGAGGCCCCCACACAAGGGTCAGCGTCCTCTCTAATAAACGGGGTTGTCAGGCTCCTGTCAAAACCCTGGGATGTGGTGACTGGAGTCACAGGAATAGCCATGACCGACACCACACCGTATGGTCAGCAAAGAGTTTTCAAGGAAAAAGTGGACACTAGGGTGCCAGACCCCCAAGAAGGCACTCGTCAGGTTATGAGCATGGTCTCTTCCTGGTTGTGGAAAGAGCTAGGCAAACACAAACGGCCACGAGTCTGTACCAAAGAAGAGTTCATCAACAAGGTTCGCAGCAATGCAGCATTAGGGGCAATATTTGAAGAGGAAAAAGAGTGGAAGACTGCAGTGGAAGCTGTGAACGATCCAAGGTTCTGGGCTCTAGTGGACAAGGAAAGAGAGCACCACCTGAGAGGAGAGTGCCAGAGTTGTGTGTACAACATGATGGGAAAAAGAGAAAAGAAACAAGGGGAATTTGGAAAGGCCAAGGGCAGCCGCGCCATCTGGTATATGTGGCTAGGGGCTAGATTTCTAGAGTTCGAAGCCCTTGGATTCTTGAACGAGGATCACTGGATGGGGAGAGAGAACTCAGGAGGTGGTGTTGAAGGGCTGGGATTACAAAGACTCGGATATGTCCTAGAAGAGATGAGTCGCATACCAGGAGGAAGGATGTATGCAGATGACACTGCTGGCTGGGACACCCGCATCAGCAGGTTTGATCTGGAGAATGAAGCTCTAATCACCAACCAAATGGAGAAAGGGCACAGGGCCTTGGCATTGGCCATAATCAAGTACACATACCAAAACAAAGTGGTAAAGGTCCTTAGACCAGCTGAAAAAGGGAAAACAGTTATGGACATTATTTCGAGACAAGACCAAAGGGGGAGCGGACAAGTTGTCACTTACGCTCTTAACACATTTACCAACCTAGTGGTGCAACTCATTCGGAATATGGAGGCTGAGGAAGTTCTAGAGATGCAAGACTTGTGGCTGCTGCGGAGGTCAGAGAAAGTGACCAACTGGTTGCAGAGCAACGGATGGGATAGGCTCAAACGAATGGCAGTCAGTGGAGATGATTGCGTTGTGAAGCCAATTGATGACAGGTTTGCACATGCCCTCAGGTTCTTGAATGATATGGGAAAAGTTAGGAAGGACACACAAGAGTGGAAACCCTCAACTGGATGGGACAACTGGGAAGAAGTTCCGTTTTGCTCCCACCACTTCAACAAGCTCCATCTCAAGGACGGGAGGTCCATTGTGGTTCCCTGCCGCCACCAAGATGAACTGATTGGCCGGGCCCGCGTCTCTCCAGGGGCGGGATGGAGCATCCGGGAGACTGCTTGCCTAGCAAAATCATATGCGCAAATGTGGCAGCTCCTTTATTTCCATAGAAGGGACCTCCGACTGATGGCCAATGCCATTTGTTCATCTGTGCCAGTTGACTGGGTTCCAACTGGGAGAACTACCTGGTCAATCCATGGAAAGGGAGAATGGATGACCACTGAAGACATGCTTGTGGTGTGGAACAGAGTGTGGATTGAGGAGAACGACCACATGGAAGACAAGACCCCAGTTACGAAATGGACAGACATTCCCTATTTGGGAAAAAGGGAAGACTTGTGGTGTGGATCTCTCATAGGGCACAGACCGCGCACCACCTGGGCTGAGAACATTAAAAACACAGTCAACATGGTGCGCAGGATCATAGGTGATGAAGAAAAGTACATGGACTACCTATCCACCCAAGTTCGCTACTTGGGTGAAGAAGGGTCTACACCTGGAGTGCTGTAA

>ENA|AMQ76465|AMQ76465.1 Zika virus polyprotein

ATGAAAAACCC-AAAAAAGAAATCCGGAGGATTCCGGATTGTCAATATGCTAAAACGCGGAGTAGCCCGTGTGAGCCCCTTTGGGGGCTTGAAGAGGCTGCCAGCCGGACTTCTGCTGGGTCATGGGCCCATCAGGATGGTCTTGGCGATTCTAGCCTTTTTGAGATTCACGGCAATCAAGCCATCACTGGGTCTCATCAATAGATGGGGTTCAGTGGGGAAAAAAGAGGCTATGGAAATAATAAAGAAGTTCAAGAAAGATCTGGCTGCCATGCTGAGAATAATCAATGCTAGGAAGGAGAAGAAGAGACGAGGCGCAGATACTAGTGTCGGAATTGTTGGCCTCCTGCTGACCACAGCTATGGCAGCGGAGGTCACTAGACGTGGGAGTGCATACTATATGTACTTGGACAGAAACGATGCTGGGGAGGCCATATCTTTTCCAACCACATTGGGGATGAATAAGTGTTATATACAGATCATGGATCTTGGACACATGTGTGATGCCACCATGAGCTATGAATGCCCTATGCTGGATGAGGGGGTGGAACCAGATGACGTCGATTGTTGGTGCAACACGACGTCAACTTGGGTTGTGTACGGAACCTGCCATCACAAAAAAGGTGAAGCACGGAGATCTAGAAGAGCTGTGACGCTCCCCTCCCATTCCACTAGGAAGCTGCAAACGCGGTCGCAAACCTGGTTGGAATCAAGAGAATACACAAAGCACTTGATTAGAGTCGAAAATTGGATATTCAGGAACCCTGGCTTCGCGTTAGCAGCAGCTGCCATCGCTTGGCTTTTGGGAAGCTCAACGAGCCAAAAAGTCATATACTTGGTCATGATACTGCTGATTGCCCCGGCATACAGCATCAGGTGCATAGGAGTCAGCAATAGGGACTTTGTGGAAGGTATGTCAGGTGGGACTTGGGTTGATGTTGTCTTGGAACATGGAGGTTGTGTCACCGTAATGGCACAGGACAAACCGACTGTCGACATAGAGCTGGTTACAACAACAGTCAGCAACATGGCGGAGGTAAGATCCTACTGCTATGAGGCATCAATATCAGACATGGCTTCGGACAGCCGCTGCCCAACACAAGGTGAAGCCTACCTTGACAAGCAATCAGACACTCAATATGTCTGCAAAAGAACGTTAGTGGACAGAGGCTGGGGAAATGGATGTGGACTTTTTGGCAAAGGGAGCCTGGTGACATGCGCTAAGTTTGCATGCTCCAAGAAAATGACCGGGAAGAGCATCCAGCCAGAGAATCTGGAGTACCGGATAATGCTGTCAGTTCATGGCTCCCAGCACAGTGGGATGATCGTTAATGACACAGGACATGAAACTGATGAGAATAGAGCGAAGGTTGAGATAACGCCCAATTCACCAAGAGCCGAAGCCACCCTGGGGGGTTTTGGAAGCCTAGGACTTGATTGTGAACCGAGGACAGGCCTTGACTTTTCAGATTTGTATTACTTGACTATGAATAACAAGCACTGGTTGGTTCACAAGGAGTGGTTCCACGACATTCCATTACCTTGGCACGCTGGGGCAGACACCGGAACTCCACACTGGAACAACAAAGAAGCACTGGTAGAGTTCAAGGACGCACATGCCAAAAGGCAAACTGTCGTGGTTCTAGGGAGTCAAGAAGGAGCAGTTCACACGGCCCTTGCTGGAGCTCTAGAGGCTGAGATGGATGGTGCAAAGGGAAGGCTGTCCTCTGGCCACTTGAAATGTCGCCTGAAAATGGATAAACTTAGATTGAAGGGCGTGTCATACTCCTTGTGTACCGCAGCGTTCACATTCACCAAGATCCCGGCTGAAACACTGCACGGGACAGTCACAGTGGAGGTACAGTACGCAGGGACTGATGGACCTTGCAAGGTTCCAGCTCAGATGGCGGTGGACATGCAAACTCTGACCCCAGTTGGGAGGTTGATAACCGCTAACCCCGTAATCACTGAAAGCACTGAGAACTCTAAGATGATGCTGGAACTTGATCCACCATTTGGGGACTCCTACATTGTCATAGGAGTCGGGGAGAAGAAGATCACCCACCACTGGCACAGGAGTGGCAGCACCATTGGAAAAGCATTTGAAGCCACTGTGAGAGGTGCCAAGAGAATGGCAGTCTTGGGAGACACAGCCTGGGACTTTGGATCAGTTGGAGGCGCTCTCAACTCATTGGGCAAGGGCATCCATCAAATTTTTGGAGCAGCTTTCAAATCATTGTTTGGAGGAATGTCCTGGTTCTCACAAATTCTCATTGGAACGTTGCTGATGTGGTTGGGTCTGAACACAAAGAATGGATCTATTTCCCTTATGTGCTTGGCCTTAGGGGGAGTGTTGATCTTCTTATCCACAGCCGTCTCTGCTGATGTGGGGTGCTCGGTGGACTTCTCAAAGAAGGAGACGAGATGCGGTACAGGGGTGTTCGTCTATAACGACGTTGAAGCCTGGAGGGACAGGTACAAGTACCATCCTGACTCCCCCCGTAGATTGGCAGCAGCAGTCAAGCAAGCCTGGGAAGATGGTATCTGCGGGATCTCCTCTGTTTCAAGAATGGAAAACATCATGTGGAGATCAGTAGAAGGGGAGCTCAACGCAATCCTGGAAGAGAATGGAGTTCAACTGACGGTCGTTGTGGGATCTGTAAAAAACCCCATGTGGAGAGGTCCACAGAGATTGCCCGTGCCTGTGAACGAGCTGCCCCACGGCTGGAAGGCTTGGGGGAAATCGTACTTCGTCAGAGCAGCAAAGACAAATAACAGCTTTGTCGTGGATGGTGACACACTGAAGGAATGCCCACTCAAACATAGAGCATGGAACAGCTTTCTTGTGGAGGATCATGGGTTCGGGGTATTTCACACTAGTGTCTGGCTCAAGGTTAGAGAAGATTATTCATTAGAGTGTGATCCAGCCGTTATTGGAACAGCTGTTAAGGGAAAGGAGGCTGTACACAGTGATCTAGGCTACTGGATTGAGAGTGAGAAGAATGACACATGGAGGCTGAAGAGGGCCCATCTGATCGAGATGAAAACATGTGAATGGCCAAAGTCCCACACATTGTGGACAGATGGAATAGAAGAGAGTGATCTGATCATACCCAAGTCTTTAGCTGGGCCACTCAGCCATCACAATACCAGAGAGGGCTACAGGACCCAAATGAAAGGGCCATGGCACAGTGAAGAGCTTGAAATTCGGTTTGAGGAATGCCCAGGCACTAAGGTCCACGTGGAGGAAACATGTGGAACAAGAGGACCATCTCTGAGATCAACCACTGCAAGCGGAAGGGTGATCGAGGAATGGTGCTGCAGGGAGTGCACAATGCCCCCACTGTCGTTCCGGGCTAAAGATGGCTGTTGGTATGGAATGGAGATAAGGCCCAGGAAAGAACCAGAAAGCAACTTAGTAAGGTCAATGGTGACTGCAGGATCAACTGATCACATGGATCACTTCTCCCTTGGAGTGCTTGTGATTCTGCTCATGGTGCAGGAAGGGCTGAAGAAGAGAATGACCACAAAGATCATCATAAGCACATCAATGGCAGTGCTGGTAGCTATGATCCTGGGAGGATTTTCAATGAGTGACCTGGCTAAGCTTGCAATTTTGATGGGTGCCACCTTCGCGGAAATGAACACTGGAGGAGATGTAGCTCATCTGGCGCTGATAGCGGCATTCAAAGTCAGACCAGCGTTGCTGGTATCTTTCATCTTCAGAGCTAATTGGACACCCCGTGAAAGCATGCTGCTGGCCTTGGCCTCGTGTCTTTTGCAAACTGCGATCTCCGCCTTGGAAGGCGACCTGATGGTTCTCATCAATGGTTTTGCTTTGGCCTGGTTGGCAATACGAGCGATGGTTGTTCCACGCACTGATAACATCACCTTGGCAATCTTGGCTGCTCTGACACCACTGGCCCGGGGCACACTGCTTGTGGCGTGGAGAGCAGGCCTTGCTACTTGCGGGGGGTTTATGCTCCTCTCTCTGAAGGGAAAAGGCAGTGTGAAGAAGAACTTACCATTTGTCATGGCCCTGGGACTAACCGCTGTGAGGCTGGTCGACCCCATCAACGTGGTGGGACTGCTGTTGCTCACAAGGAGTGGGAAGCGGAGCTGGCCCCCTAGCGAAGTACTCACAGCTGTTGGCCTGATATGCGCATTGGCTGGAGGGTTCGCCAAGGCAGATATAGAGATGGCTGGGCCCATGGCCGCGGTCGGTCTGCTAATTGTCAGTTACGTGGTCTCAGGAAAGAGTGTGGACATGTACATTGAAAGAGCAGGTGACATCACATGGGAAAAAGATGCGGAAGTCACTGGAAACAGTCCCCGGCTCGATGTGGCGCTAGATGAGAGTGGTGATTTCTCCCTGGTGGAGGATGACGGTCCCCCCATGAGAGAGATCATACTCAAGGTGGTCCTGATGACCATCTGTGGCATGAACCCAATAGCCATACCCTTTGCAGCTGGAGCGTGGTACGTATACGTGAAGACTGGAAAAAGGAGTGGTGCTCTATGGGATGTGCCTGCTCCCAAGGAAGTAAAAAAGGGGGAGACCACAGATGGAGTGTACAGAGTAATGACTCGTAGACTGCTAGGTTCAACACAAGTTGGAGTGGGAGTTATGCAAGAGGGGGTCTTTCACACTATGTGGCACGTCACAAAAGGATCCGCGCTGAGAAGCGGTGAAGGGAGACTTGATCCATACTGGGGAGATGTCAAGCAGGATCTGGTGTCATACTGTGGTCCATGGAAGCTAGATGCCGCCTGGGACGGGCACAGCGAGGTGCAGCTCTTGGCCGTGCCCCCCGGAGAGAGAGCGAGGAACATCCAGACTCTGCCCGGAATATTTAAGACAAAGGGTGGGGACATTGGAGCGGTTGCGCTGGATTACCCAGCAGGAACTTCAGGATCTCCAATCCTAGACAAGTGTGGGAGAGTGATAGGACTTTATGGCAATGGGGTCGTGATCAAAAATGGGAGTTATGTTAGTGCCATCACCCAAGGGAGGAGGGAGGAAGAGACTCCTGTTGAGTGCTTCGAGCCTTCGATGCTGAAGAAGAAGCAGCTAACTGTCTTAGACTTGCATCCTGGAGCTGGGAAAACCAGGAGAGTTCTTCCTGAAATAGTCCGTGAAGCCATAAAAACAAGACTCCGTACTGTGATCTTAGCTCCAACCAGGGTTGTCGCTGCTGAAATGGAGGAAGCCCTTAGAGGGCTTCCAGTGCGTTATATGACAACAGCAGTCAATGTCACCCACTCTGGAACAGAAATCGTCGACTTAATGTGCCATGCCACCTTCACTTCACGTCTACTACAGCCAATCAGAGTCCCCAATTATAATCTGTATATTATGGATGAGGCCCACTTCACAGATCCCTCAAGTATAGCAGCAAGAGGATACATTTCAACAAGGGTTGAGATGGGCGAGGCGGCTGCCATCTTCATGACCGCCACGCCACCAGGAACCCGTGACGCATTTCCGGACTCCAACTCACCAATTATGGACACCGAAGTGGAAGTCCCAGAGAGAGCCTGGAGCTCAGGCTTTGATTGGGTGACGGATCATTCTGGAAAAACAGTTTGGTTTGTTCCAAGCGTGAGGAACGGCAATGAGATCGCAGCTTGTCTGACAAAGGCTGGAAAACGGGTCATACAGCTCAGCAGAAAGACTTTTGAGACAGAGTTCCAGAAAACAAAACATCAAGAGTGGGACTTTGTCGTGACAACTGACATTTCAGAGATGGGCGCCAACTTTAAAGCTGACCGTGTCATAGATTCCAGGAGATGCCTAAAGCCGGTCATACTTGATGGCGAGAGAGTCATTCTGGCTGGACCCATGCCTGTCACACATGCCAGCGCTGCCCAGAGGAGGGGGCGCATAGGCAGGAATCCCAACAAACCTGGAGATGAGTATCTGTATGGAGGTGGGTGCGCAGAGACTGACGAAGACCATGCACACTGGCTTGAAGCAAGAATGCTCCTTGACAATATTTACCTCCAAGATGGCCTCATAGCCTCGCTCTATCGACCTGAGGCCGACAAAGTAGCAGCCATTGAGGGAGAGTTCAAGCTTAGGACGGAGCAAAGGAAGACCTTTGTGGAACTCATGAAAAGAGGAGATCTTCCTGTTTGGCTGGCCTATCAGGTTGCATCTGCCGGAATAACCTACACAGATAGAAGATGGTGCTTTGATGGCACGACCAACAACACCATAATGGAAGACAGTGTGCCGGCAGAGGTGTGGACCAGACACGGAGAGAAAAGAGTGCTCAAACCGAGGTGGATGGATGCCAGAGTTTGTTCAGATCATGCGGCCCTGAAGTCATTCAAGGAGTTTGCCGCTGGGAAAAGAGGAGCGGCTCTTGGAGTGATGGAAGCCCTGGGAACACTGCCAGGACACATGACAGAGAGATTCCAGGAAGCCATTGACAACCTCGCTGTGCTCATGCGGGCAGAGACTGGAAGCAGGCCTTACAAAGCCGCGGCGGCCCAAATGCCGGAGACCCTAGAGACCATTATGCTTTTGGGGTTGCTAGGAACAGTCTCGCTGGGAATCTTTTTCGTCTTGATGAGGAACAAGGGCATAGGGAAGATGGGCTTTGGAATGGTGACTCTTGGGGCCAGCGCATGGCTCATGTGGCTCTCGGAAATTGAGCCAGCCAGAATTGCATGTGTCCTCATTGTTGTGTTCCTATTGCTGGTGGTGCTCATACCTGAGCCAGAAAAGCAAAGATCTCCCCAGGACAACCAAATGGCAATCATCATCATGGTAGCAGTAGGTCTTCTGGGCTTGATTACCGCCAATGAACTCGGATGGTTGGAGAGAACAAAGAGTGACCTAAGCCATCTAATGGGAAGGAGAGAGGAGGGGGCAACCATAGGATTCTCAATGGACATTGACCTGCGGCCAGCCTCAGCTTGGGCCATCTATGCTGCCTTGACAACTTTCATTACCCCAGCCGTCCAACATGCAGTGACCACTTCATACAACAACTACTCCTTAATGGCGATGGCCACGCAAGCTGGAGTGTTGTTTGGTATGGGCAAAGGGATGCCATTCTACGCATGGG-ACTTTGGAGTCCCGCTGCTAATGATAGGTTGCTACTCACAATTAACACCCCTGACCCTAATAGTGGCCATCATTTTGCTCGTGGCGCACTACATGTACTTGATCCCAGGGCTGCAGGCAGCAGCTGCGCGTGCTGCCCAGAAGAGAACGGCAGCTGGCATCATGAAGAACCCTGTTGTGGATGGAATAGTGGTGACTGACATTGACACAATGACAATTGACCCCCAAGTGGAGAAAAAGATGGGACAGGTGCTACTCATAGCAGTAGCCGTCTCCAGCGCCATACTGTCGCGGACCGCCTGGGGGTGGGGGGAGGCTGGGGCTCTGATCACAGCCGCAACTTCCACTTTGTGGGAAGGCTCTCCGAACAAGTACTGGAACTCCTCTACAGCCACTTCACTGTGTAACATTTTTAGGGGAAGTTACTTGGCTGGAGCTTCTCTAATCTACACAGTAACAAGAAACGCTGGCTTGGTCAAGAGACGTGGGGGTGGAACAGGAGAGACCCTGGGAGAGAAATGGAAGGCCCGCTTGAACCAGATGTCGGCCCTGGAGTTCTACTCCTACAAAAAGTCAGGCATCACCGAGGTGTGCAGAGAAGAGGCCCGCCGCGCCCTCAAGGACGGTGTGGCAACGGGAGGCCATGCTGTGTCCCGAGGAAGTGCAAAGCTGAGATGGTTGGTGGAGCGGGGATACCTGCAGCCCCATGGAAAGGTCATTGATCTTGGATGTGGCAGAGGGGGCTGGAGTTACTACGCCGCCACCATCCGCAAAGTTCAAGAAGTGAAAGGATACACAAAAGGAGGCCCTGGTCATGAAGAACCCGTGTTGGTGCAAAGCTATGGGTGGAACATAGTCCGTCTTAAGAGTGGGGTGGACGTCTTTCATATGGCGGCTGAGCCGTGTGACACGTTGCTGTGTGACATAGGTGAGTCATCATCTAGTCCTGAAGTGGAAGAAGCACGGACGCTCAGAGTTCTCTCCATGGTGGGGGATTGGCTTGAAAAAAGACCAGGAGCCTTTTGTATAAAAGTGTTGTGCCCATACACCAGCACTATGATGGAAACCCTGGAGCGACTGCAGCGTAGGTATGGGGGAGGACTGGTCAGAGTGCCACTCTCCCGCAACTCTACACATGAGATGTACTGGGTCTCTGGAGCGAAAAGCAACACCATAAAAAGTGTGTCCACCACGAGCCAGCTCCTCTTGGGGCGCATGGACGGGCCTAGGAGGCCAGTGAAATATGAGGAGGATGTGAATCTCGGCTCTGGCACGCGGGCTGTGGTAAGCTGCGCTGAAGCTCCCAACATGAAGATCATTGGTAACCGCATTGAAAGGATCCGCAGTGAGCACGCGGAAACGTGGTTCTTTGACGAGAACCACCCATATAGGACATGGGCTTACCATGGAAGCTATGAGGCCCCCACACAAGGGTCAGCGTCCTCTCTAGTAAACGGGGTTGTCAGGCTCCTGTCAAAACCCTGGGATGTGGTGACTGGAGTCACAGGAATAGCCATGACCGACACCACACCGTATGGTCAGCAAAGAGTTTTCAAGGAAAAAGTGGACACTAGGGTGCCAGACCCCCAAGAAGGCACTCGTCAGGTTATGAGCATGGTCTCTTCCTGGTTGTGGAAAGAGCTAGGCAAACACAAACGGCCACGAGTCTGTACCAAAGAAGAGTTCATCAACAAGGTTCGTAGCAATGCAGCACTAGGGGCAATATTTGAAGAGGAAAAAGAGTGGAAGACTGCAGTGGAAGCTGTGAACGATCCAAGGTTCTGGGCTCTAGTGGACAAGGAAAGAGAGCACCACCTGAGAGGAGAGTGCCAGAGTTGTGTGTACAACATGATGGGAAAAAGAGAAAAGAAACAAGGGGAATTTGGAAAGGCCAAGGGCAGCCGCGCCATCTGGTATATGTGGCTAGGGGCTAGATTTCTAGAGTTCGAAGCCCTTGGATTCTTGAACGAGGATCACTGGATGGGGAGAGAGAACTCAGGAGGTGGTGTTGAAGGGCTGGGATTACAAAGACTCGGATATGTCCTAGAAGAGATGAGTCGCATACCAGGAGGAAGGATGTATGCAGATGACACTGCTGGCTGGGACACCCGCATCAGCAGGTTTGATCTGGAGAATGAAGCTCTAATCACCAACCAAATGGAGAAAGGGCACAGGGCCTTGGCATTGGCCATAATCAAGTACACATACCAAAACAAAGTGGTAAAGGTCCTTAGACCAGCTGAAAAAGGGAAAACAGTTATGGACATTATTTCGAGACAAGACCAAAGGGGGAGCGGACAAGTTGTCACTTACGCTCTTAACACATTTACCAACCTAGTGGTGCAACTCATTCGGAATATGGAGGCTGAGGAAGTTCTCGAGATGCAAGACTTGTGGCTGCTGCGGAGGCCAGAGAAAGTGACCAACTGGTTGCAGAGCAACGGATGGGATAGGCTCAAACGAATGGCAGTCAGTGGAGATGATTGCGTTGTGAAGCCAATTGATGATAGGTTTGCACATGCCCTCAGGTTCTTGAATGATATGGGAAAAGTTAGGAAGGACACACAAGAGTGGAAACCCTCAACTGGATGGGACAACTGGGAAGAAGTTCCGTTTTGCTCCCACCACTTCAACAAGCTCCATCTCAAGGACGGGAGGTCCATTGTGGTTCCCTGCCGCCACCAAGATGAACTGATTGGCCGGGCCCGCGTCTCTCCAGGGGCGGGATGGAGCATCCGGGAGACTGCTTGCCTAGCAAAATCATATGCGCAAATGTGGCAGCTCCTTTATTTCCACAGAAGGGACCTCCGACTGATGGCCAATGCCATTTGCTCATCTGTGCCAGTTGACTGGGTTCCAACTGGGAGAACTACCTGGTCAATCCATGGAAAGGGAGAATGGATGACCACTGAAGACATGCTTGTGGTGTGGAACAGAGTGTGGATTGAGGAGAACGACCACATGGAAGACAAGACCCCAGTTACGAAATGGACAGACATTCCCTATTTGGGAAAAAGGGAAGACTTGTGGTGTGGATCTCTCATAGGGCACAGACCGCGCACCACCTGGGCTGAGAACATTAAAAACACAGTCAACATGGTGCGCAGGATCATAGGTGATGAAGAAAAGTACATGGACTACCTATCCACCCAAGTTCGCTACTTGGGTGAAGAAGGGTCTACACCTGGAGTGCTGTAA

>ENA|AMQ48982|AMQ48982.1 Zika virus polyprotein

ATGAAAAACCC-AAAAAAGAAATCCGGAGGATTCCGGATTGTCAATATGCTAAAACGCGGAGTAGCCCGTGTGAGCCCCTTTGGGGGCTTGAAGAGGCTGCCAGCCGGACTTCTGCTGGGTCATGGGCCCATCAGGATGGTCTTGGCGATTCTAGCCTTTTTGAGATTCACGGCAATCAAGCCATCACTGGGTCTCATCAATAGATGGGGTTCAGTGGGGAAAAAAGAGGCTATGGAAATAATAAAGAAGTTCAAGAAAGATCTGGCTGCCATGCTGAGAATAATCAATGCTAGGAAGGAGAAGAAGAGACGAGGCGCAGATACTAGTGTCGGAATTGTTGGCCTCCTGCTGACCACAGCTATGGCAGCGGAGGTCACCAGACGTGGGAGTGCATACTATATGTACTTGGACAGAAACGATGCTGGGGAGGCCATATCTTTTCCAACCACATTGGGGATGAATAAGTGTTATATACAGATCATGGATCTTGGACACATGTGTGATGCCACCATGAGCTATGAATGCCCTATGCTGGATGAGGGGGTGGAACCAGATGACGTCGATTGTTGGTGCAACACGACGTCAACTTGGGTTGTGTACGGAACCTGCCATCACAAAAAAGGTGAAGCACGGAGATCTAGAAGAGCTGTGACGCTCCCCTCCCATTCCACTAGGAAGCTGCAAACGCGGTCGCAAACCTGGTTGGAATCAAGAGAATACACAAAGCACTTGATTAGAGTCGAAAATTGGATATTCAGGAACCCTGGCTTCGCGTTAGCAGCAGCTGCCATCGCTTGGCTTTTGGGAAGCTCAACGAGCCAAAAAGTCATATACTTGGTCATGATACTGCTGATTGCCCCGGCATACAGCATCAGGTGCATAGGAGTCAGCAATAGGGACTTTGTGGAAGGTATGTCAGGTGGGACTTGGGTTGATGTTGTCTTGGAACATGGAGGTTGTGTCACCGTAATGGCACAGGACAAACCGACTGTCGACATAGAGCTGGTTACAACAACAGTCAGCAACATGGCGGAGGTAAGATCCTACTGCTATGAGGCATCAATATCAGACATGGCTTCGGACAGCCGCTGCCCAACACAAGGTGAAGCCTACCTTGACAAGCAATCAGACACTCAATATGTCTGCAAAAGAACGTTGGTGGACAGAGGCTGGGGAAATGGATGTGGACTTTTTGGCAAAGGGAGCCTGGTGACATGCGCTAAGTTTGCATGCTCCAAGAAAATGACCGGGAAGAGCATCCAGCCAGAGAATCTGGAGTACCGGATAATGCTGTCAGTTCATGGCTCCCAGCACAGTGGGATGATCGTTAATGACACAGGACATGAAACTGATGAGAATAGAGCGAAGGTTGAGATAACGCCCAATTCACCAAGAGCCGAAGCCACCCTGGGGGGTTTTGGAAGCCTAGGACTTGATTGTGAACCGAGGACAGGCCTTGACTTTTCAGATTTGTATTACTTGACTATGAATAACAAGCATTGGTTGGTTCACAAGGAGTGGTTCCACGACATTCCATTACCTTGGCACGCTGGGGCAGACACCGGAACTCCACACTGGAACAACAAAGAAGCACTGGTAGAGTTCAAGGACGCACATGCCAAAAGGCAAACTGTCGTGGTTCTAGGGAGTCAAGAAGGAGCAGTTCACACGGCCCTTGCTGGAGCTCTGGAGGCTGAGATGGATGGTGCAAAGGGAAGGCTGTCCTCTGGCCACTTGAAATGTCGCCTGAAAATGGATAAACTTAGATTGAAGGGCGTGTCATACTCCTTGTGTACCGCAGCGTTCACATTCACCAAGATCCCGGCTGAAACACTGCACGGGACAGTCACAGTGGAGGTACAGTACGCAGGGGCAGATGGACCCTGCAAGGTTCCAGCTCAGATGGCGGTGGACATGCAAACTCTGACCCCAGTTGGGAGGTTGATAACCGCCAACCCCGTAATCACTGAAAGCACTGAGAACTCTAAGATGATGCTGGAACTTGATCCACCATTTGGGGACTCTTACATTGTCATAGGAGTCGGGGAGAAGAAGATCACCCACCACTGGCACAGGAGTGGCAGCACCATTGGAAAAGCATTTGAAGCCACTGTGAGAGGTGCCAAGAGAATGGCAGTCTTGGGAGACACAGCCTGGGACTTTGGATCAGTTGGAGGCGCTCTCAACTCATTGGGCAAGGGCATCCATCAAATTTTTGGAGCAGCTTTCAAATCATTGTTTGGAGGAATGTCCTGGTTCTCACAAATCCTCATTGGAACGTTGCTGATGTGGTTGGGTCTGAACACAAAGAATGGATCTATTTCCCTTATGTGCTTGGCCTTAGGGGGAGTGTTGATCTTCTTATCCACAGCCGTCTCTGCTGATGTGGGGTGCTCGGTGGACTTCTCAAAGAAGGAGACGAGATGCGGTACAGGGGTGTTCGTCTATAACGACGTTGAAGCCTGGAGGGATAGGTACAAGTACCATCCTGACTCCCCCCGTAGATTGGCAGCAGCAGTCAAGCAAGCCTGGGAAGATGGTATCTGCGGGATCTCCTCTGTTTCAAGAATGGAGAACATCATGTGGAGATCAGTAGAAGGGGAGCTCAACGCAATCCTGGAAGAGAATGGAGTTCAACTGACGGTCGTTGTGGGATCTGTAAAAAACCCCATGTGGAGAGGTCCACAGAGATTGCCCGTGCCTGTGAACGAGCTGCCCCACGGCTGGAAGGCTTGGGGGAAATCGTACTTCGTCAGAGCAGCAAAGACAAATAACAGCTTTGTCGTGGATGGTGACACACTGAAGGAATGCCCACTCAAACATAGAGCATGGAACAGCTTTCTTGTGGAGGATCATGGGTTCGGGGTATTTCACACTAGTGTCTGGCTCAAGGTTAGAGAAGATTATTCATTAGAGTGTGATCCAGCCGTTATTGGAACAGCTGTTAAGGGAAAGGAGGCTGTACACAGTGATCTAGGCTACTGGATTGAGAGTGAGAAGAATGACACATGGAGGCTGAAGAGGGCCCATCTGATCGAGATGAAAACATGTGAATGGCCAAAGTCCCACACATTGTGGACAGATGGAATAGAAGAGAGTGATCTGATCATACCCAAGTCTTTAGCTGGGCCACTCAGCCATCACAATACCAGAGAGGGCTACAGGACCCAAATGAAAGGGCCATGGCACAGTGAAGAGCTTGAAATTCGGTTTGAGGAATGCCCAGGCACTAAGGTCCACGTGGAGGAAACATGTGGAACAAGAGGACCATCTCTGAGATCAACCACTGCAAGCGGAAGGGTGATCGAGGAATGGTGCTGCAGGGAGTGCACAATGCCCCCACTGTCGTTCCGGGCTAAAGATGGCTGTTGGTATGGAATGGAGATAAGGCCCAGGAAAGAACCAGAAAGCAACTTAGTAAGGTCAGTGGTGACTGCAGGATCAACTGATCACATGGATCACTTCTCCCTTGGAGTGCTTGTGATTCTGCTCATGGTGCAGGAAGGGCTGAAGAAGAGAATGACCACAAAGATCATCATAAGCACATCAATGGCAGTGCTGGTAGCTATGATCCTGGGAGGATTTTCAATGAGTGACCTGGCTAAGCTTGCAATTTTGATGGGTGCCACCTTCGCGGAAATGAACACTGGAGGAGATGTAGCTCATCTGGCGCTGATAGCGGCATTCAAAGTCAGACCAGCGTTGCTGGTATCTTTCATCTTCAGAGCTAATTGGACACCCCGTGAAAGCATGCTGCTGGCCTTGGCCTCGTGTCTTTTGCAAACTGCGATCTCCGCCTTGGAAGGCGACCTGATGGTTCTCATCAATGGTTTTGCTTTGGCCTGGTTGGCAATACGAGCGATGGTTGTTCCACGCACTGATAACATCACCTTGGCAATCCTGGCTGCTCTGACACCACTGGCCCGGGGCACACTGCTTGTGGCGTGGAGAGCAGGCCTTGCTACTTGCGGGGGGTTTATGCTCCTCTCTCTGAAGGGAAAAGGCAGTGTGAAGAAGAACTTACCATTTGTCATGGCCCTGGGACTAACCGCTGTGAGGCTGGTCGACCCCATCAACGTGGTGGGACTGCTGTTGCTCACAAGGAGTGGGAAGCGGAGCTGGCCCCCTAGCGAAGTACTCACAGCTGTTGGCCTGATATGCGCATTGGCTGGAGGGTTCGCCAAGGCAGATATAGAGATGGCTGGGCCCATGGCCGCGGTCGGTCTGCTAATTGTCAGTTACGTGGTCTCAGGAAAGAGTGTGGACATGTACATTGAAAGAGCAGGTGACATCACATGGGAAAAAGATGCGGAAGTCACTGGAAACAGTCCCCGGCTCGATGTGGCGCTAGATGAGAGTGGTGATTTCTCCCTGGTGGAGGATGACGGTCCCCCCATGAGAGAGATCATACTCAAGGTGGTCCTGATGACCATCTGTGGCATGAACCCAATAGCCATACCCTTTGCAGCTGGAGCGTGGTACGTATACGTGAAGACTGGAAAAAGGAGTGGTGCTCTATGGGATGTGCCTGCTCCCAAGGAAGTAAAAAAGGGGGAGACCACAGATGGAGTGTACAGAGTAATGACTCGTAGACTGCTAGGTTCAACACAAGTTGGAGTGGGAGTTATGCAAGAGGGGGTCTTTCACACTATGTGGCACGTCACAAAAGGATCCGCGCTGAGAAGCGGTGAAGGGAGACTTGATCCATACTGGGGAGATGTCAAGCAGGATCTGGTGTCATACTGTGGTCCATGGAAGCTAGATGCCGCCTGGGACGGGCACAGCGAGGTGCAGCTCTTGGCCGTGCCCCCCGGAGAGAGAGCGAGGAACATCCAGACTCTGCCCGGAATATTTAAGACAAAGGATGGGGACATTGGAGCGGTTGCGCTGGATTACCCAGCAGGAACTTCAGGATCTCCAATCCTAGACAAGTGTGGGAGAGTGATAGGACTTTATGGCAATGGGGTCGTGATCAAAAATGGGAGTTATGTTAGTGCCATCACCCAAGGGAGGAGGGAGGAAGAGACTCCTGTTGAGTGCTTCGAGCCTTCGATGCTGAAGAAGAAGCAGCTAACTGTCTTAGACTTGCATCCTGGAGCCGGGAAAACCAGGAGAGTTCTTCCTGAAATAGTCCGTGAAGCCATAAAAACAAGACTCCGTACTGTGATCTTAGCTCCAACCAGGGTTGTCGCTGCTGAAATGGAGGAAGCCCTTAGAGGGCTTCCAGTGCGTTATATGACAACAGCAGTCAATGTCACCCACTCTGGAACAGAAATCGTCGACTTAATGTGCCATGCCACCTTCACTTCACGTCTACTACAGCCAATCAGAGTCCCCAACTATAATCTGTATATTATGGATGAGGCCCACTTCACAGATCCCTCAAGCATAGCAGCAAGAGGATACATTTCAACAAGGGTTGAGATGGGCGAGGCGGCTGCCATCTTCATGACCGCCACGCCACCAGGAACCCGTGACGCATTTCCGGACTCCAACTCACCAATTATGGACACCGAAGTGGAAGTCCCAGAGAGAGCCTGGAGCTCAGGCTTTGATTGGGTGACGGATCATTCTGGAAAAACAGTTTGGTTTGTTCCAAGCGTGAGGAACGGCAATGAGATCGCAGCTTGTCTGACAAAGGCTGGAAAACGGGTCATACAGCTCAGCAGAAAGACTTTTGAGACAGAGTTCCAGAAAACAAAACATCAAGAGTGGGACTTTGTCGTGACAACTGACATTTCAGAGATGGGCGCCAACTTTAAAGCTGACCGTGTCATAGATTCCAGGAGATGCCTAAAGCCGGTCATACTTGATGGCGAGAGAGTCATTCTGGCTGGACCCATGCCTGTCACACATGCCAGCGCTGCCCAGAGGAGGGGGCGCATAGGCAGGAATCCCAACAAACCTGGAGATGAGTATCTGTATGGAGGTGGGTGCGCAGAGACTGATGAAGACCATGCACACTGGCTTGAAGCAAGAATGCTCCTTGACAATATTTACCTCCAAGATGGCCTCATAGCCTCGCTCTATCGACCTGAGGCCGACAAAGTAGCAGCCATTGAGGGAGAGTTCAAGCTTAGGACGGAGCAAAGGAAGACCTTTGTGGAACTCATGAAAAGAGGAGATCTTCCTGTTTGGCTGGCCTATCAGGTTGCATCTGCCGGAATAACCTACACAGATAGAAGATGGTGCTTTGATGGCACGACCAACAACACCATAATGGAAGACAGTGTGCCGGCAGAGGTGTGGACCAGACACGGAGAGAAAAGAGTGCTCAAACCGAGGTGGATGGACGCCAGAGTTTGTTCAGATCATGCGGCCCTGAAGTCATTCAAGGAGTTTGCCGCTGGGAAAAGAGGAGCGACTTTTGGAGTGATGGAAGCCCTGGGAACACTGCCAGGACACATGACAGAGAGATTCCAGGAAGCCATTGACAACCTCGCTGTGCTCATGCGGGCAGAGACTGGAAGCAGGCCTTACAAAGCCGCGGCGGCCCAATTGCCGGAGACCTTAGAGACCATTATGCTTTTGGGATTGCTGGGAACAGTCTCGTTGGGAATCTTTTTCGTCTTGATGAGGAACAAGGGCATAGGGAAGATGGGCTTTGGAATGGTGACTCTTGGGGCCAGCGCATGGCTCATGTGGCTCTCGGAAATTGAGCCAGCCAGAATTGCATGTGTCCTCATTGTTGTGTTCCTATTGCTGGTGGTGCTCATACCTGAGCCAGAAAAGCAAAGATCTCCCCAGGACAACCAAATGGCAATCATCATCATGGTAGCAGTAGGTCTTCTGGGCTTGATTACCGCCAATGAACTCGGATGGTTGGAGAGAACAAAGAGTGACCTAAGCCATCTAATGGGAAGGAGAGAGGAGGGAGCAACCATAGGATTCTCAATGGACATTGACCTGCGGCCAGCCTCAGCTTGGGCCATCTATGCTGCCTTGACAACTTTCATTACTCCAGCCGTCCAACATGCAGTGACCACTTCATACAACAACTACTCCTTAATGGCGATGGCCACGCAAGCTGGAGTGTTGTTTGGTATGGGCAAAGGGATGCCATTCTACGCATGGG-ACTTTGGAGTCCCGCTGCTAATGATAGGTTGCTACTCACAATTAACACCCCTGACCCTAATAGTGGCCATCATTTTGCTCGTGGCGCACTACATGTACTTGATCCCAGGGCTGCAGGCAGCAGCTGCGCGTGCTGCCCAGAAGAGAACGGCAGCTGGCATCATGAAGAACCCTGTTGTGGATGGAATAGTGGTGACTGACATTGACACAATGACAATTGACCCCCAAGTGGAGAAAAAGATGGGACAGGTGCTACTCATAGCAGTAGCCGTCTCCAGCGCCATACTGTCGCGGACCGCCTGGGGGTGGGGGGAGGCTGGGGCCCTGATCACAGCCGCAACTTCCACTTTGTGGGAAGGCTCTCCGAACAAGTACTGGAACTCCTCTACAGCCACTTCACTGTGTAACATTTTTAGGGGAAGTTACTTGGCTGGAGCTTCTCTAATCTACACAGTAACAAGAAACGCTGGCTTGGTCAAGAGACGTGGGGGTGGAACAGGAGAGACCCTGGGAGAGAAATGGAAGGCCCGCTTGAACCAGATGTCGGCCCTGGAGTTCTACTCCTACAAAAAGTCAGGCATCACCGAGGTGTGCAGAGAAGAGGCCCGCCGCGCCCTCAAGGACGGTGTGGCAACGGGAGGCCATGCTGTGTCCCGAGGAAGTGCAAAGCTGAGATGGTTAGTGGAGCGGGGATACCTGCAGCCCTATGGAAAGGTCATTGATCTTGGATGTGGCAGAGGGGGCTGGAGTTACTACGCCGCCACCATCCGCAAAGTTCAAGAAGTGAAAGGATACACAAAAGGAGGCCCTGGTCATGAAGAACCCGTGTTGGTGCAAAGCTATGGGTGGAACATAGTCCGTCTTAAGAGTGGGGTGGACGTCTTTCATATGGCGGCTGAGCCGTGTGACACGTTGCTGTGTGACATAGGTGAGTCATCATCTAGTCCTGAAGTGGAAGAAGCACGGACGCTCAGAGTCCTCTCCATGGCGGGGGATTGGCTTGAAAAAAGACCAGGAGCCTTTTGTATAAAAGTGTTGTGCCCATACACCAGCACCATGATGGAAACCCTGGAGCGACTGCAGCGTAGGTATGGGGGAGGACTGGTCAGAGTGCCACTCTCCCGCAACTCTACACATGAGATGTACTGGGTCTCTGGAGCGAAAAGCAACACCATAAAAAGTGTGTCCACCACGAGCCAGCTCCTCTTGGGGCGCATGGACGGGCCTAGGAGGCCAGTGAAATATGAGGAGGATGTGAATCTCGGCTCTGGCACGCGGGCTGTGGTAAGCTGCGCTGAAGCTCCCAACATGAAGATCATTGGTAACCGCATTGAAAGGATCCGCAGTGAGCACGCGGAAACGTGGTTCTTTGACGAGAACCACCCATATAGGACATGGGCTTACCATGGAAGCTATGAGGCCCCCACACAAGGGTCAGCGTCCTCTCTAATAAACGGGGTTGTCAGGCTCCTGTCAAAACCCTGGGATGTGGTGACTGGAGTCACAGGAATAGCCATGACCGACACCACACCGTATGGTCAGCAAAGAGTTTTCAAGGAAAAAGTGGACACTAGGGTGCCAGACCCCCAAGAAGGCACTCGTCAGGTTATGAGCATGGTCTCTTCCTGGTTGTGGAAAGAGCTAGGCAAACACAAACGGCCACGAGTCTGTACCAAAGAAGAGTTCATCAACAAGGTTCGTAGCAATGCAGCATTAGGGGCAATATTTGAAGAGGAAAAAGAGTGGAAGACTGCAGTGGAAGCTGTGAACGATCCAAGGTTCTGGGCTCTAGTGGACAAGGAAAGAGAGCACCACCTGAGAGGAGAGTGCCAGAGTTGTGTGTACAACATGATGGGAAAAAGAGAAAAGAAACAAGGGGAATTTGGAAAGGCCAAGGGCAGCCGCGCCATCTGGTATATGTGGCTAGGGGCTAGATTTCTAGAGTTCGAAGCCCTTGGATTCTTGAACGAGGATCACTGGATGGGGAGAGAGAACTCAGGAGGTGGTGTTGAAGGGCTGGGATTACAAAGACTCGGATATGTCCTAGAAGAGATGAGTCGCATACCAGGAGGAAGGATGTATGCAGATGACACTGCTGGCTGGGACACCCGCATCAGCAGGTTTGATCTGGAGAATGAAGCTCTAATCACCAACCAAATGGAGAAAGGGCACAGGGCCTTGGCATTGGCCATAATCAAGTACACATACCAAAACAAAGTGGTAAAGGTCCTTAGACCAGCTGAAAAAGGGAAAACAGTTATGGACATTATTTCGAGACAAGACCAAAGGGGGAGCGGACAAGTTGTCACTTACGCTCTTAACACATTTACCAATCTAGTGGTGCAACTCATTCGGAATATGGAGGCTGAGGAAGTTCTAGAGATGCAAGACTTGTGGCTGCTGCGGAGGTCAGAGAAAGTGACCAACTGGTTGCAGAGCAACGGATGGGATAGGCTCAAACGAATGGCAGTCAGTGGAGATGATTGCGTTGTGAAGCCAATTGATGATAGGTTTGCACATGCCCTCAGGTTCTTGAATGATATGGGAAAAGTTAGGAAGGACACACAAGAGTGGAAACCCTCAACTGGATGGGACAACTGGGAAGAAGTTCCGTTTTGCTCCCACCACTTCAACAAGCTCCATCTCAAGGACGGGAGGTCCATTGTGGTTCCCTGCCGCCACCAAGATGAACTGATTGGCCGGGCCCGCGTCTCTCCAGGGGCGGGATGGAGCATCCGGGAGACTGCTTGCCTAGCAAAATCATATGCGCAAATGTGGCAGCTCCTTTATTTCCACAGAAGGGACCTCCGACTGATGGCCAATGCCATTTGTTCATCTGTGCCAGTTGACTGGGTTCCAACTGGGAGAACTACCTGGTCAATCCATGGAAAGGGAGAATGGATGACCACTGAAGACATGCTTGTGGTGTGGAACAGAGTGTGGATTGAGGAGAACGACCACATGGAAGACAAGACCCCAGTTACGAAATGGACAGACATTCCCTATTTGGGAAAAAGGGAAGACTTGTGGTGTGGATCTCTCATAGGGCACAGACCGCGCACCACCTGGGCTGAGAACATTAAAAACACAGTCAACATGGTGCGCAGGATCATAGGTGATGAAGAAAAGTACATGGACTACCTATCCACCCAAGTTCGCTACTTGGGTGAAGAAGGGTCTACACCTGGAGTGCTGTAA

>ENA|AMQ34003|AMQ34003.1 Zika virus polyprotein

ATGAAAAACCC-AAAAAAGAAATCCGGAGGATTCCGGATTGTCAATATGCTAAAACGCGGAGTAGCCCGTGTGAGCCCCTTTGGGGGCTTGAAGAGGCTGCCAGCCGGACTTCTGCTGGGTCATGGGCCCATCAGGATGGTCTTGGCGATTCTAGCCTTTTTGAGATTCACGGCAATCAAGCCATCACTGGGTCTCATCAATAGATGGGGTTCAGTGGGGAAAAAAGAGGCTATGGAAATAATAAAGAAGTTCAAGAAAGATCTGGCTGCCATGCTGAGAATAATCAATGCTAGGAAGGAGAAGAAGAGACGAGGCGCAGAAACTAGTGTCGGAATTGTTGGCCTCCTGCTGACCACAGCTATGGCAGCGGAGGTCACTAGACGTGGGAGTGCATACTATATGTACTTGGACAGAAACGATGCTGGGGAGGCCATATCTTTTCCAACCACATTGGGGATGAATAAGTGTTATATACAGATCATGGATCTTGGACACATGTGTGATGCCACCATGAGCTATGAATGCCCTATGCTGGATGAGGGGGTGGAACCAGATGACGTCGATTGTTGGTGCAACACGACGTCAACTTGGGTTGTGTACGGAACCTGCCATCACAAAAAAGGTGAAGCACGGAGATCTAGAAGAGCCGTGACGCTCCCCTCCCATTCCACTAGGAAGCTGCAAACGCGGTCGCAAACCTGGTTGGAATCAAGAGAATACACAAAGCACTTGATTAGAGTCGAAAATTGGATATTCAGGAACCCTGGTTTCGCTTTAGCAGCAGCTGCCATCGCGTGGCTTTTGGGAAGCTCAACGAGCCAAAAAGTCATATACTTGGTCATGATACTGCTGATTGCCCTGGCATACAGCATCAGGTGCATAGGAGTCAGCAATAGGGACTTTGTGGAAGGTATGTCAGGTGGGACTTGGGTTGATGTTGTCTTGGAACATGGAGGTTGTGTCACCGTAATGGCACAGGACAAACCGACTGTCGACATAGAGCTGGTTACAACAACAGTCAGCAACATGGCGGAGGTAAGATCCTACTGCTATGAGGCATCAATATCAGACATGGCTTCGGACAGCCGCTGCCCAACACAAGGTGAAGCCTACCTTGACAAGCAATCAGACACTCAATATGTCTGCAAAAGAACGTTAGTGGACAGAGGCTGGGGAAATGGATGTGGACTTTTTGGCAAAGGGAGCCTGGTGACATGCGCTAAGTTTGCATGCTCCAAGAAAATGACCGGGAAGAGCATCCAGCCAGAGAATCTGGAGTACCGGATAATGTTGTCAGTTCATGGCTCCCAGCACAGTGGGATGATCGTTAATGACACAGGACATGAAACTGATGAGAATAGAGCGAAGGTTGAGATAACGCCCAATTCACCAAGAGCCGAAGCCACCCTGGGGGGTTTTGGAAGCCTAGGACTTGATTGTGAACCGAGGACAGGCCTTGACTTTTCAGATTTGTATTACTTGACTATGAATAACAAGCACTGGTTGGTTCACAAGGAGTGGTTCCACGACATTCCATTACCTTGGCACGCTGGGGCAGACACCGGAACTCCACACTGGAACAACAAAGAAGCACTGGTAGAGTTCAAGGACGCACATGCCAAAAGGCAAACTGTCGTGGTTCTAGGGAGTCAAGAAGGAGCAGTTCACACGGCCCTTGCTGGAGCTCTGGAGGCTGAGATGGATGGTGCAAAGGGAAGGCTGTCCTCTGGCCACTTGAAATGTCGCCTGAAAATGGATAAACTTAGATTGAAGGGCGTGTCATACTCCTTGTGTACCGCAGCGTTCACATTCACCAAGATCCCGGCTGAAACACTGCACGGGACAGTCACAGTGGAGGTACAGTACGCAGGGACAGATGGACCTTGCAAGGTTCCAGCCCAGATGGCGGTGGACATGCAAACTCTGACCCCAGTTGGGAGGTTGATAACCGCTAACCCCGTAATCACTGAAAGCACTGAGAACTCTAAGATGATGCTGGAACTTGATCCACCATTTGGGGACTCTTACATTGTCATAGGAGTCGGGGAGAAGAAGATCACCCACCACTGGCACAGGAGTGGCAGCACCATTGGAAAAGCATTTGAAGCCACTGTGAGAGGTGCCAAGAGAATGGCAGTCTTGGGAGACACAGCCTGGGACTTTGGATCAGTTGGAGGCGCTCTCAACTCATTGGGCAAGGGCATCCATCAAATTTTTGGAGCAGCTTTCAAATCATTGTTTGGAGGAATGTCCTGGTTCTCACAAATTCTCATTGGAACGTTGCTGATGTGGTTGGGTCTGAACACAAAGAATGGATCTATTTCCCTTATGTGCTTGGCCTTAGGGGGAGTGTTGATCTTCTTATCCACAGCCGTCTCTGCTGATGTGGGGTGCTCGGTGGACTTCTCAAAGAAGGAGACGAGATGTGGTACAGGGGTGTTCGTCTATAACGACGTTGAAGCCTGGAGGGACAGGTACAAGTACCATCCTGACTCCCCCCGTAGATTGGCAGCAGCAGTCAAGCAAGCCTGGGAAGATGGTATCTGCGGGATCTCCTCTGTTTCAAGAATGGAAAACATCATGTGGAGATCAGTAGAAGGGGAGCTCAACGCAATCCTGGAAGAGAATGGAGTTCAACTGACGGTCGTTGTGGGATCTGTAAAAAACCCCATGTGGAGAGGTCCACAGAGATTGCCCGTGCCTGTGAACGAGCTGCCCCACGGCTGGAAGGCTTGGGGGAAATCGTACTTCGTCAGAGCAGCAAAGACAAATAACAGCTTTGTCGTGGATGGTGACACACTGAAGGAATGCCCACTCAAACATAGAGCATGGAACAGCTTTCTTGTGGAGGATCATGGGTTCGGGGTATTTCACACTAGTGTCTGGCTCAAGGTTAGAGAAGATTATTCATTAGAGTGTGATCCAGCCGTTATTGGAACAGCTGTTAAGGGAAAGGAGGCTGTACACAGTGATCTAGGCTACTGGATTGAGAGTGAGAAGAATGACACATGGAGGCTGAAGAGGGCCCATCTGATCGAGATGAAAACATGTGAATGGCCAAAATCCCACACATTGTGGACAGATGGAATAGAAGAGAGTGATCTGATCATACCCAAGTCTTTAGCTGGGCCACTCAGCCATCACAATACCAGAGAGGGCTACAGGACCCAAATGAAAGGGCCATGGCACAGTGAAGAGCTTGAAATTCGGTTTGAGGAATGCCCAGGCACTAAGGTCCACGTGGAGGAAACATGTGGAACAAGAGGACCATCTCTGAGATCAACCACTGCAAGCGGAAGGGTGATCGAGGAATGGTGCTGCAGGGAGTGCACAATGCCCCCACTGTCGTTCTGGGCTAAAGATGGCTGTTGGTATGGAATGGAGATAAGGCCCAGGAAAGAACCAGAAAGCAACTTAGTAAGGTCAATGGTGACTGCAGGATCAACTGATCACATGGATCACTTCTCCCTTGGAGTGCTTGTGATTCTGCTCATGGTGCAGGAAGGGCTGAAGAAGAGAATGACCACAAAGATCATCATAAGCACATCAATGGCAGTGCTGGTAGCTATGATCCTGGGAGGATTTTCAATGAGTGACCTGGCTAAGCTTGCAATTTTGATGGGTGCCACCTTCGCGGAAATGAACACTGGAGGAGATGTAGCTCATCTGGCGCTGACAGCGGCATTCAAAGTCAGACCAGCGTTGCTGGTATCTTTCATCTTCAGAGCTAATTGGACACCCCGTGAAAGCATGCTGCTGGCCTTGGCCTCGTGTCTTTTGCAAACTGCGATCTCTGCCTTGGAAGGCGACCTGATGGTTCTCATCAATGGTTTTGCTTTGGCCTGGTTGGCAATACGAGCGATGGTTGTTCCACGCACTGACAACATCACCTTGGCAATCCTGGCTGCTCTGACACCACTGGCCCGGGGCACACTGCTTGTGGCGTGGAGAGCAGGCCTTGCTACTTGCGGGGGGTTTATGCTCCTCTCTCTGAAGGGAAAAGGCAGTGTGAAGAAGAACTTACCATTTGTCATGGCCCTGGGACTAACCGCTGTGAGGCTGGTCGACCCCATCAACGTGGTGGGACTGCTGTTGCTCACAAGGAGTGGGAAGCGGAGCTGGCCCCCTAGCGAAGTACTCACAGCTGTTGGCCTGATATGTGCATTGGCTGGAGGGTTCGCCAAGGCAGATATAGAGATGGCTGGGCCCATGGCCGCGGTCGGTCTGCTAATTGTCAGTTACGTGGTCTCAGGAAAGAGTGTGGACATGTACATTGAAAGAGCAGGTGACATCACATGGGAAAAAGATGCGGAAGTCACTGGAAACAGTCCCCGGCTCGACGTGGCGCTAGATGAGAGTGGTGATTTCTCCCTGGTGGAGGATGACGGTCCCCCCATGAGAGAGATCATACTCAAGGTGGTCCTGATGACCATCTGTGGCATGAACCCAATAGCCATACCCTTTGCAGCTGGAGCGTGGTACGTATACGTGAAGACTGGAAAAAGGAGTGGTGCTCTATGGGATGTGCCTGCTCCCAAGGAAGTAAAAAAGGGGGAGACCACAGATGGAGTGTACAGAGTAATGACTCGTAGACTGCTAGGTTCAACACAAGTTGGAGTGGGAGTTATGCAAGAGGGGGTCTTTCACACTATGTGGCACGTCACAAAAGGATCCGCGCTGAGAAGCGGTGAAGGGAGACTTGATCCATACTGGGGAGATGTCAAGCAGGATCTGGTGTCATACTGTGGTCCATGGAAGCTAGATGCCGCCTGGGACGGGCACAGCGAGGTGCAGCTCTTGGCCGTGCCCCCCGGAGAGAGAGAGAGGAACATCCAGACTCTGCCCGGAATATTTAAGACAAAGGATGGGGACATTGGAGCGGTTGCGCTGGATTACCCAGCAGGAACTTCAGGATCTCCAATCCTAGACAAGTGTGGGAGAGTGATAGGACTTTATGGCAATGGGGTCGTGATCAAAAATGGGAGTTATGTTAGTGCCATCACCCAAGGGAGGAGGGAGGAAGAGACTCCTGTTGAGTGCTTCGAGCCTTCGATGCTGAAGAAGAAGCAGCTAACTGTCTTAGACTTGCATCCTGGAGCTGGGAAAACCAGGAGAGTTCTTCCTGAAATAGTCCGTGAAGCCATAAAAACAAGACTCCGTACTGTGATCTTAGCTCCAACCAGGGTTGTCGCTGCTGAAATGGAGGAAGCCCTTAGAGGGCTTCCAGTGCGTTATATGACAACAGCAGTCAATGTCACCCACTCTGGAACAGAAATCGTCGACTTAATGTGCCATGCCACCTTCACTTCACGTCTACTACAGCCAATCAGAGTCCCCAACTATAATCTGTATATTATGGATGAGGCCCACTTCACAGATCCCTCAAGTATAGCAGCAAGAGGATACATTTCAACAAGGGTTGAGATGGGCGAGGCGGCTGCCATCTTCATGACCGCCACGCCACCAGGAACCCGTGACGCATTTCCGGACTCCAACTCACCAATTATGGACACCGAAGTGGAAGTCCCAGAGAGAGCCTGGAGCTCAGGCTTTGATTGGGTGACGGATCATTCTGGAAAAACAGTTTGGTTTGTTCCAAGCGTGAGGAACGGCAATGAGATCGCAGCTTGTCTGACAAAGGCTGGAAAACGGGTCATACAGCTCAGCAGAAAGACTTTTGAGACAGAGTTCCAGAAAACAAAACATCAAGAGTGGGACTTTGTCGTGACAACTGACATTTCAGAGATGGGCGCCAACTTTAAAGCTGACCGTGTCATAGATTCCAGGAGATGCCTAAAGCCGGTCATACTTGATGGCGAGAGAGTCATTCTGGCTGGACCCATGCCTGTCACACATGCCAGCGCTGCCCAGAGGAGGGGGCGCATAGGCAGGAATCCCAATAAACCTGGAGATGAGTATCTGTATGGAGGTGGGTGCGCAGAGACTGACGAAGACCATGCACACTGGCTTGAAGCAAGAATGCTCCTTGACAATATTTACCTCCAAGATGGCCTCATAGCCTCGCTCTATCGACCTGAGGCCGACAAAGTAGCAGCCATTGAGGGAGAGTTCAAGCTTAGGACGGAGCAAAGGAAGACCTTTGTGGAACTCATGAAAAGAGGAGATCTTCCTGTTTGGCTGGCCTATCAGGTTGCATCTGCCGGAATAACCTACACAGATAGAAGATGGTGCTTTGATGGCACGACCAACAACACCATAATGGAAGACAGTGTGCCGGCAGAGGTGTGGACCAGACACGGAGAGAAAAGAGTGCTCAAACCGAGGTGGATGGACGCCAGAGTTTGTTCAGATCATGCGGCCCTGAAGTCATTCAAGGAGTTTGCCGCTGGGAAAAGAGGAGCGGCTTTTGGAGTGATGGAAGCCCTGGGAACACTGCCAGGACACATGACAGAGAGATTCCAGGAAGCCATTGACAACCTCGCTGTGCTCATGCGGGCAGAGACTGGAAGCAGGCCTTACAAAGCCGCGGCGGCCCAATTGCCGGAGACCCTAGAGACCATTATGCTTTTGGGGTTGCTGGGAACAGTCTCGCTGGGAATCTTTTTCGTCTTGATGAGGAACAAGGGCATAGGGAAGATGGGCTTTGGAATGGTGACTCTTGGGGCCAGCGCATGGCTCATGTGGCTCTCGGAAATTGAGCCAGCCAGAATTGCATGTGTCCTCATTGTTGTGTTCCTATTGCTGGTGGTGCTCATACCTGAGCCAGAAAAGCAAAGATCTCCCCAGGACAACCAAATGGCAATCATCATCATGGTAGCAGTAGGTCTTCTGGGCTTGATTACCGCCAATGAACTCGGATGGTTGGAGAGAACAAAGAGTGACCTAAGCCATCTAATGGGAAGGAGAGAGGAGGGGGCAACCATAGGATTCTCAATGGACATTGACCTGCGGCCAGCCTCAGCTTGGGCCATCTATGCTGCCTTGACAACTTTCATTACCCCAGCCGTCCAACATGCAGTGACCACTTCATACAACAACTACTCCTTAATGGCGATGGCCACGCAAGCTGGAGTGTTGTTTGGTATGGGCAAAGGGATGCCATTCTACGCATGGG-ACTTTGGAGTCCCACTGCTAATGATAGGTTGCTACTCACAATTAACACCCCTGACCCTAATAGTGGCCATCATTTTGCTCGTGGCGCACTACATGTACTTGATCCCAGGGCTGCAGGCAGCAGCTGCGCGTGCTGCCCAGAAGAGAACGGCAGCTGGCATCATGAAGAACCCTGTTGTGGATGGAATAGTGGTGACTGACATTGACACAATGACAATTGACCCCCAAGTGGAGAAAAAGATGGGACAGGTGCTACTCATAGCAGTAGCCGTCTCCAGCGCCATACTGTCGCGGACCGCCTGGGGGTGGGGGGAGGCTGGGGCCCTGATCACAGCCGCAACTTCCACTTTGTGGGAAGGCTCTCCGAACAAGTACTGGAACTCCTCTACAGCCACTTCACTGTGTAACATTTTTAGGGGAAGTTACTTGGCTGGAGCTTCTCTAATCTACACAGTAACAAGAAACGCTGGCTTGGTCAAGAGACGTGGGGGTGGAACAGGAGAGACCCTGGGAGAGAAATGGAAGGCCCGCTTGAACCAGATGTCGGCCCTGGAGTTCTACTCCTACAAAAAGTCAGGCATCACCGAGGTGTGCAGAGAAGAGGCCCGCCGCGCCCTCAAGGACGGTGTGGCAACGGGAGGCCATGCTGTGTCCCGAGGAAGTGCAAAGCTGAGATGGTTGGTGGAGCGGGGATACCTGCAGCCCTATGGAAAGGTCATTGATCTTGGATGTGGCAGAGGGGGCTGGAGTTACTACGCCGCCACCATCCGCAAAGTTCAAGAAGTGAAAGGATACACAAAAGGAGGCCCTGGTCATGAAGAACCCGTGTTGGTGCAAAGCTATGGGTGGAACATAGTCCGTCTTAAGAGTGGGGTGGACGTCTTTCATATGGCGGCTGAGCCGTGTGACACGTTGCTGTGTGACATAGGTGAGTCATCATCTAGTCCTGAAGTGGAAGAAGCACGGACGCTCAGAGTCCTCTCCATGGTGGGGGATTGGCTTGAAAAAAGACCAGGAGCCTTTTGTATAAAAGTGTTGTGCCCATACACCAGCACTATGATGGAAACCCTGGAGCGACTGCAGCGTAGGTATGGGGGAGGACTGGTCAGAGTGCCACTCTCCCGCAACTCTACACATGAGATGTACTGGGTCTCTGGAGCGAAAAGCAACACCATAAAAAGTGTGTCCACCACGAGCCAGCTCCTCTTGGGGCGCATGGACGGGCCTAGGAGGCCAGTGAAATATGAGGAGGATGTGAATCTCGGCTCTGGCACGCGGGCTGTGGTAAGCTGCGCTGAAGCTCCCAACATGAAGATCATTGGTAACCGCATTGAAAGGATCCGCAGTGAGCAAGCGGAAACGTGGTTCTTTGACGAGAACCACCCATATAGGACATGGGCTTACCATGGAAGCTATGAGGCCCCCACACAAGGGTCAGCGTCCTCTCTAATAAACGGGGTTGTCAGGCTCCTGTCAAAACCCTGGGATGTGGTGACTGGAGTCACAGGAATAGCCATGACCGACACCACACCGTATGGTCAGCAAAGAGTTTTCAAGGAAAAAGTGGACACTAGGGTGCCAGACCCCCAAGAAGGCACTCGTCAGGTTATGAGCATGGTCTCTTCCTGGTTGTGGAAAGAGCTAGGCAAACACAAACGGCCACGAGTCTGTACCAAAGAAGAGTTCATCAACAAGGTTCGTAGCAATGCAGCATTAGGGGCAATATTTGAAGAGGAAAAAGAGTGGAAGACTGCAGTGGAAGCTGTGAACGATCCAAGGTTCTGGGCTCTAGTGGACAAGGAAAGAGAGCACCACCTGAGAGGAGAGTGCCAGAGTTGTGTGTACAACATGATGGGAAAAAGAGAAAAGAAACAAGGGGAATTTGGAAAGGCCAAGGGCAGCCGCGCCATCTGGTATATGTGGCTAGGGGCTAGATTTCTAGAGTTCGAAGCCCTTGGATTCTTGAACGAGGATCACTGGATGGGGAGAGAGAACTCAGGAGGTGGTGTTGAAGTGCTGGGATTACAGAGACTCGGATATGTCCTAGAAGAGATGAGTCGCATACCAGGAGGAAGGATGTATGCAGATGACACTGCTGGCTGGGACACCCGCATCAGCAGGTTTGATCTGGAGAATGAAGCTCTAATCACCAACCAAATGGAGAAAGGGCACAGGGCCTTGGCATTGGCCATAATCAAGTACACATACCAAAACAAAGTGGTAAAGGTCCTTAGACCAGCTGAAAAAGGGAAAACAGTTATGGACATTATTTCGAGACAAGACCAAAGGGGGAGCGGACAAGTTGTCACTTACGCTCTTAACACATTTACCAACCTAGTGGTGCAACTCATTCGGAATATGGAGGCTGAGGAAGTTCTAGAGATGCAAGACTTGTGGCTGCTGCGGAGGTCAGAGAAAGTGACCAACTGGTTGCAGAGCAACGGATGGGATAGGCTCAAACGAATGGCAGTCAGTGGAGATGATTGCGTTGTGAAGCCAATTGATGATAGGTTTGCACATGCCCTCAGGTTCTTGAATGATATGGGAAAAGTTAGGAAGGACACACAAGAGTGGAAACCCTCAACTGGATGGGACAACTGGGAAGAAGTTCCGTTTTGCTCCCACCACTTCAACAAGCTCCATCTCAAGGACGGGAGGTCCATTGTGGTTCCCTGCCGCCACCAAGATGAACTGATTGGCCGGGCCCGCGTCTCTCCAGGGGCGGGATGGAGCATCCGGGAGACTGCTTGCCTAGCAAAATCATATGCGCAAATGTGGCAGCTCCTTTATTTCCACAGAAGGGACCTCCGACTGATGGCCAATGCCATTTGTTCATCTGTGCCAGTTGACTGGGTTCCAACTGGGAGAACTACCTGGTCAATCCATGGAAAGGGAGAATGGATGACCACTGAAGACATGCTTGTGGTGTGGAACAGAGTGTGGATTGAGGAGAACGACCACATGGAAGACAAGACCCCAGTTGCGAAATGGACAGACATTCCCTATTTGGGAAAAAGGGAAGACTTGTGGTGTGGATCTCTCATAGGGCACAGACCGCGCACCACCTGGGCTGAGAACATTAAAAACACAGTCAACATGGTGCGCAGGATCATAGGTGATGAAGAAAAGTACATGGACTACCTATCCACCCAAGTTCGCTACTTGGGTGAAGAAGGGTCTACACCTGGAGTGCTGTAA

>ENA|AMM39805|AMM39805.1 Zika virus polyprotein

ATGAAAAACCC-AAAAAAGAAATCCGGAGGATTCCGGATTGTCAATATGCTAAAACGCGGAGTAGCCCGTGTGAGCCCCTTTGGGGGCTTGAAGAGGCTGCCAGCCGGACTTCTGCTGGGTCATGGGCCCATCAGGATGGTCTTGGCGATTCTAGCCTTTTTGAGATTCACGGCAATCAAGCCATCACTGGGTCTCATCAATAGATGGGGTTCAGTGGGGAAAAAAGAGGCTATGGAAATAATAAAGAAGTTCAAGAAAGATCTGGCTGCCATGCTGAGAATAATCAATGCTAGGAAGGAGAAGAAGAGACGAGGCGCAGAAACTAGTGTCGGAATTGTTGGCCTCCTGCTGACCACAGCTATGGCAGCGGAGGTCACTAGACGTGGGAGTGCATACTATATGTACTTGGACAGAAACGATGCTGGGGAGGCCATATCTTTTCCAACCACATTGGGGATGAATAAGTGTTATATACAGATCATGGATCTTGGACACATGTGTGATGCCACCATGAGCTATGAATGCCCTATGCTGGATGAGGGGGTGGAACCAGATGACGTCGATTGTTGGTGCAACACGACGTCAACTTGGGTTGTGTACGGAACCTGCCATCACAAAAAAGGTGAAGCACGGAGATCTAGAAGAGCTGTGACGCTCCCTTCCCATTCCACTAGGAAGCTGCAAACGCGGTCGCAAACCTGGTTGGAATCAAGAGAATACACAAAGCACTTGATTAGAGTCGAAAATTGGATATTCAGGAACCCTGGCTTCGCGTTAGCAGCAGCTGCTATCGCTTGGCTTTTGGGAAGCTCAACGAGCCAAAAAGTCATATACTTGGTCATGATACTGCTGATTGCCCCGGCATACAGCATCAGGTGCATAGGAGTCAGCAATAGGGACTTTGTGGAAGGTATGTCAGGTGGGACTTGGGTTGATGTTGTCTTGGAACATGGAGGTTGTGTCACCGTAATGGCACAGGACAAACCGACTGTCGACATAGAGCTGGTTACAACAACAGTCAGCAACATGGCGGAGGTAAGATCCTACTGCTATGAGGCATCAATATCAGACATGGCTTCGGACAGCCGCTGCCCAACACAAGGTGAAGCCTACCTTGACAAGCAATCAGACACTCAATATGTCTGCAAAAGAACGTTAGTGGACAGAGGCTGGGGAAATGGATGTGGACTTTTTGGCAAAGGGAGCCTGGTGACATGCGCTAAGTTTGCATGCTCCAAGAAAATGACCGGGAAAAGCATCCAGCCAGAGAATCTGGAGTACCGGATAATGCTGTCAGTTCATGGCTCCCAGCACAGTGGGATGATCGTTAATGACACAGGACATGAAACTGATGAGAATAGAGCGAAAGTTGAGATAACGCCCAATTCACCAAGAGCCGAAGCCACCCTGGGGGGTTTTGGAAGCCTAGGACTTGATTGTGAACCGAGGACAGGCCTTGACTTTTCAGATTTGTATTACTTGACTATGAATAACAAGCACTGGTTGGTTCACAAGGAGTGGTTCCACGACATTCCATTACCTTGGCACGCTGGGGCAGACACCGGAACTCCACACTGGAACAACAAAGAAGCACTGGTAGAGTTCAAGGACGCACATGCCAAAAGGCAAACTGTCGTGGTTCTAGGGAGTCAAGAAGGAGCAGTTCACACGGCCCTTGCTGGAGCTCTGGAGGCTGAGATGGATGGAGCAAAGGGAAGGCTGTCCTCTGGCCACTTGAAATGTCGCCTGAAAATGGATAAACTTAGATTGAAGGGCGTGTCATACTCCTTGTGTACTGCAGCGTTCACATTCACCAAGATCCCGGCTGAAACACTGCACGGGACAGTCACAGTGGAGGTACAGTACGCAGGGACAGATGGACCTTGCAAGGTTCCAGCTCAGATGGCGGTGGACATGCAAACTCTGACCCCAGTTGGGAGGTTGATAACCGCTAACCCCGTAATCACTGAAAGCACTGAGAACTCTAAGATGATGCTGGAACTTGATCCACCATTTGGGGACTCTTACATTGTCATAGGAGTCGGGGAGAAGAAGATCACCCACCACTGGCACAGGAGTGGCAGCACCATTGGAAAAGCATTTGAAGCCACTGTGAGAGGTGCCAAGAGAATGGCAGTCTTGGGAGACACAGCCTGGGACTTTGGATCAGTTGGAGGCGCTCTCAACTCATTGGGCAAGGGCATCCATCAAATTTTTGGAGCAGCTTTCAAATCATTGTTTGGAGGAATGTCCTGGTTCTCACAAATTCTCATTGGAACGTTGCTGATGTGGTTGGGTCTGAACACAAAGAATGGATCTATTTCCCTTATGTGCTTGGCCTTAGGGGGAGTGTTGATCTTCTTATCCACAGCCGTCTCTGCTGATGTGGGGTGCTCGGTGGACTTCTCAAAGAAGGAGACGAGATGCGGTACAGGGGTGTTTGTCTATAACGACGTTGAAGCCTGGAGGGACAGGTACAAGTACCATCCTGACTCCCCCCGTAGATTGGCAGCAGCAGTCAAGCAAGCCTGGGAAGATGGTATCTGCGGGATCTCCTCTGTTTCAAGAATGGAAAACATCATGTGGAGATCAGTAGAAGGGGAGCTCAACGCAATCCTGGAAGAGAATGGAGTTCAACTGACGGTCGTTGTGGGATCTGTAAAAAACCCCATGTGGAGAGGTCCACAGAGATTGCCCGTGCCTGTGAACGAGCTGCCCCACGGCTGGAAGGCTTGGGGGAAATCGTACTTCGTCAGAGCAGCAAAGACAAATAACAGCTTTGTCGTGGATGGTGACACACTGAAGGAATGCCCACTCAAACATAGAGCATGGAACAGCTTTCTTGTGGAGGATCATGGGTTCGGGGTATTTCACACTAGTGTCTGGCTCAAGGTTAGAGAAGACTATTCATTAGAGTGTGATCCAGCCGTTATTGGAACAGCTGTTAAGGGAAAGGAGGCTGTACACAGTGATCTAGGCTACTGGATTGAGAGTGAGAAGAATGACACATGGAGGCTGAAGAGGGCCCATCTGATCGAGATGAAAACATGTGAATGGCCAAAGTCCCACACATTGTGGACAGATGGAATAGAAGAGAGTGATCTGATCATACCCAAGTCTTTAGCTGGGCCACTCAGCCATCACAATACCAGAGAGGGCTACAGGACCCAAATGAAAGGGCCATGGCACAGTGAGGAGCTTGAAATTCGGTTTGAGGAATGCCCAGGCACTAAGGTCCACGTGGAGGAAACATGTGGAACAAGAGGACCATCTCTGAGATCAACCACTGCAAGCGGAAGGGTGATCGAGGAATGGTGCTGCAGGGAGTGCACAATGCCCCCACTGTCGTTCCGGGCTAAAGATGGCTGTTGGTATGGAATGGAGATAAGGCCCAGGAAAGAACCAGAAAGCAACTTAGTAAGGTCAATGGTGACTGCAGGATCAACTGATCACATGGACCACTTCTCCCTTGGAGTGCTTGTGATCCTGCTCATGGTGCAGGAAGGGCTGAAGAAGAGAATGACCACAAAGATCATCATAAGCACATCAATGGCAGTGCTGGTAGCTATGATCCTGGGAGGATTTTCAATGAGTGACCTGGCTAAGCTTGCAATTTTGATGGGTGCCACCTTCGCGGAAATGAACACTGGAGGAGATGTAGCTCATCTGGCGCTGATAGCGGCATTCAAAGTCAGACCAGCGTTGCTGGTATCTTTCATCTTCAGAGCTAATTGGACACCCCGTGAAAGCATGCTGCTGGCCTTGGCCTCGTGTCTTTTGCAAACTGCGATCTCCGCCTTGGAAGGCGACCTGATGGTTCTCATCAATGGTTTTGCTTTGGCCTGGTTGGCAGTACGAGCGATGGTTGTTCCACGCACTGATAACATCACCTTAGCAATCCTGGCTGCTCTGACACCACTGGCCCGGGGCACACTGCTTGTGGCGTGGAGAGCAGGCCTTGCTACTTGCGGGGGGTTTATGCTCCTCTCTCTGAAGGGAAAAGGCAGTGTGAAGAAGAACTTACCATTTGTCATGGCCCTGGGACTAACCGCTGTGAGGCTGGTCGACCCCATCAACGTGGTGGGACTGCTGTTGCTCACAAGGAGTGGGAAGCGGAGCTGGCCCCCTAGCGAAGTACTCACAGCTGTTGGCCTGATATGCGCATTGGCTGGAGGGTTCGCCAAGGCAGATATAGAGATGGCTGGGCCCATGGCCGCGGTCGGTCTGCTAATTGTCAGTTACGTGGTCTCAGGAAAGAGTGTGGACATGTACATTGAAAGAGCAGGTGACATCACATGGGAAAAAGATGCGGAAGTCACTGGAAACAGTCCCCGGCTCGATGTGGCGCTAGATGAGAGTGGTGATTTCTCCCTGGTGGAGGATGACGGTCCCCCCATGAGAGAGATCATACTCAAGGTGGTCCTGATGACCATCTGTGGCATGAACCCAATAGCCATACCCTTTGCAGCTGGAGCGTGGTACGTATACGTGAAGACTGGAAAAAGGAGTGGTGCTCTATGGGATGTGCCTGCTCCCAAGGAAGTAAAAAAGGGGGAGACCACAGATGGAGTGTACAGAGTAATGACTCGCAGACTGCTAGGTTCAACACAAGTTGGAGTGGGAGTTATGCAAGAGGGGGTCTTTCACACTATGTGGCACGTCACAAAAGGATCCGCGCTGAGAAGCGGTGAAGGGAGACTTGATCCATACTGGGGAGATGTCAAGCAGGATCTGGTGTCATACTGTGGTCCATGGAAGCTAGATGCCGCCTGGGACGGGCACAGCGAGGTGCAGCTCTTGGCCGTGCCCCCCGGAGAGAGAGCGAGGAACATCCAGACTCTGCCCGGAATATTTAAGACAAAGGATGGGGACATTGGAGCGGTTGCGCTGGATTACCCAGCAGGAACTTCAGGATCTCCAATCCTAGACAAGTGTGGGAGAGTGATAGGACTTTATGGCAATGGGGTCGTGATCAAAAATGGGAGTTATGTTAGTGCCATCACCCAAGGGAGGAGGGAGGAAGAGACTCCTGTTGAGTGCTTCGAGCCTTCGATGCTGAAGAAGAAGCAGCTAACTGTCTTAGACTTGCATCCTGGAGCTGGGAAAACCAGGAGAGTTCTTCCTGAAATAGTCCGTGAAGCCATAAAAACAAGACTCCGTACTGTGATCTTAGCTCCAACCAGGGTTGTCGCTGCTGAAATGGAGGAGGCCCTTAGAGGGCTTCCAGTGCGTTATATGACAACAGCAGTCAATGTCACCCACTCTGGAACAGAAATCGTCGACTTAATGTGCCATGCCACCTTCACTTCACGTCTACTACAGCCAATTAGAGTCCCCAACTATAATCTGTATATTATGGATGAGGCCCACTTCACAGATCCCTCAAGTATAGCAGCAAGAGGATACATTTCAACAAGGGTTGAGATGGGCGAGGCGGCTGCCATCTTCATGACCGCCACGCCACCAGGAACCCGTGACGCATTTCCGGACTCCAACTCACCAATTATGGACACCGAAGTGGAAGTCCCAGAGAGAGCCTGGAGCTCAGGCTTTGATTGGGTGACGGATCATTCTGGAAAAACAGTTTGGTTTGTTCCAAGCGTGAGGAACGGCAATGAGATCGCAGCTTGTCTGACAAAGGCTGGAAAACGGGTCATACAGCTCAGCAGAAAGACTTTTGAGACAGAGTTCCAGAAAACAAAACATCAAGAGTGGGACTTTGTCGTGACAACTGACATTTCAGAGATGGGCGCCAACTTTAAAGCTGACCGTGTCATAGATTCCAGGAGATGCCTAAAGCCGGTCATACTTGATGGCGAGAGAGTCATTCTGGCTGGACCCATGCCTGTCACACATGCCAGCGCTGCCCAGAGGAGGGGGCGCATAGGCAGGAATCCCAACAAACCTGGAGATGAGTATCTGTATGGAGGTGGGTGCGCAGAGACTGACGAAGACCATGCACACTGGCTTGAAGCAAGAATGCTCCTTGACAATATTTACCTCCAAGATGGCCTCATAGCCTCGCTCTATCGACCTGAGGCCGACAAAGTAGCAGCCATTGAGGGAGAGTTCAAGCTTAGGACGGAGCAAAGGAAGACCTTTGTGGAACTCATGAAAAGAGGAGATCTTCCTGTTTGGCTGGCCTATCAGGTTGCATCTGCCGGAATAACCTACACAGATAGAAGATGGTGCTTTGATGGCACGACCAACAACACCATAATGGAAGACAGTGTGCCGGCAGAGGTGTGGACCAGACACGGAGAGAAAAGAGTGCTCAAACCGAGGTGGATGGACGCCAGAGTTTGTTCAGATCATGCGGCCCTGAAGTCATTCAAGGAGTTTGCCGCTGGGAAAAGAGGAGCGGCTTTTGGAGTGATGGAAGCCCTGGGAACACTGCCAGGACACATGACAGAGAGATTCCAGGAAGCCATTGACAACCTCGCTGTGCTCATGCGGGCAGAGACTGGAAGCAGGCCTTACAAAGCCGCGGCAGCCCAATTGCCGGAGACCCTAGAGACCATTATGCTTTTGGGGTTGCTGGGAACAGTCTCGCTGGGAATCTTCTTCGTCTTGATGAGGAACAAGGGCATAGGGAAGATGGGCTTTGGAATGGTGACTCTTGGGGCCAGCGCATGGCTCATGTGGCTCTCGGAAATTGAGCCAGCCAGAATTGCATGTGTCCTCATTGTTGTGTTCCTATTGCTGGTGGTGCTCATACCTGAGCCAGAAAAGCAAAGATCTCCCCAGGACAACCAAATGGCAATCATCATCATGGTAGCAGTAGGTCTTCTGGGCTTGATTACCGCCAATGAACTCGGATGGTTGGAGAGAACAAAGAGTGACCTAAGCCATCTAATGGGAAGGAGAGAGGAGGGGGCAACCATAGGATTCTCAATGGACATTGACCTGCGGCCAGCCTCAGCTTGGGCCATCTATGCTGCCTTGACAACTTTCATTACCCCAGCCGTCCAACATGCAGTGACCACCTCATACAACAACTACTCCTTAATGGCGATGGCCACGCAAGCTGGAGTGTTGTTTGGTATGGGCAAAGGGATGCCATTCTACGCATGGG-ACTTTGGAGTCCCGCTGCTAATGATAGGCTGCTACTCACAATTAACACCCCTGACCCTAATAGTGGCCATCATTTTGCTCGTGGCGCACTACATGTACTTGATCCCAGGGCTGCAGGCAGCAGCTGCGCGTGCTGCCCAGAAGAGAACGGCAGCTGGCATCATGAAGAACCCTGTTGTGGATGGAATAGTGGTGACTGACATTGACACAATGACAATTGACCCCCAAGTGGAGAAAAAGATGGGACAGGTGCTACTCATAGCAGTAGCCGTCTCCAGCGCCATACTGTCGCGGACCGCCTGGGGGTGGGGGGAGGCTGGGGCCCTGATCACAGCCGCAACTTCCACTTTGTGGGAAGGCTCTCCGAACAAGTACTGGAACTCCTCTACAGCCACTTCACTGTGTAACATTTTTAGGGGAAGTTACTTGGCTGGAGCTTCTCTAATCTACACAGTAACAAGAAACGCTGGCTTGGTCAAGAGACGTGGGGGTGGAACAGGAGAGACCCTGGGAGAGAAATGGAAGGCCCGCTTGAACCAGATGTCGGCCCTGGAGTTCTACTCCTACAAAAAGTCAGGCATCACCGAGGTGTGCAGAGAAGAGGCCCGCCGCGCCCTCAAGGACGGTGTGGCAACGGGAGGCCATGCTGTGTCCCGAGGAAGTGCAAAGCTGAGATGGTTGGTGGAGCGGGGATACCTGCAGCCCTATGGAAAGGTCATTGATCTTGGATGTGGCAGAGGGGGCTGGAGTTACTACGCCGCCACCATCCGCAAAGTTCAAGAAGTGAAAGGATACACAAAAGGAGGCCCTGGTCATGAAGAACCCGTGTTGGTGCAAAGCTATGGGTGGAACATAGTCCGTCTTAAGAGTGGGGTGGACGTCTTTCATATGGCGGCTGAGCCGTGTGACACGTTGCTGTGTGACATAGGTGAGTCATCATCTAGTCCTGAAGTGGAAGAAGCACGGACGCTCAGAGTCCTCTCCATGGTGGGGGATTGGCTTGAAAAAAGACCAGGAGCCTTTTGCATAAAAGTGTTGTGCCCATACACCAGCACTATGATGGAAACCCTGGAGCGACTGCAGCGTAGGTATGGGGGAGGACTGGTCAGAGTGCCACTCTCCCGCAACTCTACACATGAGATGTACTGGGTCTCTGGAGCGAAAAGCAACATCATAAAAAGTGTGTCCACCACGAGCCAGCTCCTCTTGGGGCGCATGGACGGGCCTAGGAGGCCAGTGAAATATGAGGAGGATGTGAATCTCGGCTCTGGCACGCGGGCTGTGGCAAGCTGCGCTGAAGCTCCCAACATGAAGATCATTGGTAACCGCATTGAAAGGATCCGCAGTGAGCACGCGGAAACGTGGTTCTTTGACGAGAACCACCCATATAGGACATGGGCTTACCATGGAAGCTATGAGGCCCCCACACAAGGGTCAGCGTCCTCTCTAATAAACGGGGTTGTCAGGCTCCTGTCAAAACCCTGGGATGTGGTGACTGGAGTCACAGGAATAGCCATGACCGACACCACACCGTATGGTCAGCAAAGAGTTTTCAAGGAAAAAGTGGACACTAGGGTGCCAGACCCCCAAGAAGGCACTCGTCAGGTTATGAGCATGGTCTCTTCCTGGTTGTGGAAAGAGCTAGGCAAACACAAACGGCCACGAGTCTGTACCAAAGAAGAGTTCATCAACAAGGTTCGTAGCAATGCAGCATTAGGGGCAATATTTGAAGAGGAAAAAGAGTGGAAGACTGCAGTGGAAGCTGTGAACGATCCAAGGTTCTGGGCTCTAGTGGACAAGGAAAGAGAGCACCACCTGAGAGGAGAGTGCCAGAGTTGTGTGTACAACATGATGGGAAAAAGAGAAAAGAAACAAGGGGAATTTGGAAAGGCCAAGGGCAGCCGCGCCATCTGGTATATGTGGCTAGGGGCTAGATTTCTAGAGTTCGAAGCCCTTGGATTCTTGAACGAGGATCACTGGATGGGGAGAGAGAACTCAGGAGGTGGTGTTGAAGGGCTGGGATTACAAAGACTCGGATATGTCCTAGAAGAGATGAGTCGTATACCAGGAGGAAGGATGTATGCAGATGACACTGCTGGCTGGGACACCCGCATTAGCAGGTTTGATCTGGAGAATGAAGCTCTAATCACCAACCAAATGGAGAAAGGGCACAGGGCCTTGGCATTGGCCATAATCAAGTACACATACCAAAACAAAGTGGTAAAGGTCCTTAGACCAGCTGAAAAAGGGAAAACAGTTATGGACATTATTTCGAGACAAGACCAAAGGGGGAGCGGACAAGTTGTCACTTACGCTCTTAACACATTTACCAACCTAGTGGTGCAACTCATTCGGAATATGGAGGCTGAGGAAGTTCTAGAGATGCAAGACTTGTGGCTGCTGCGGAGGTCAGAGAAAGTGACCAACTGGTTGCAGAGCAACGGATGGGATAGGCTCAAACGAATGGCAGTCAGTGGAGATGATTGCGTTGTGAAGCCAATTGATGATAGGTTTGCACATGCCCTCAGGTTCTTGAATGATATGGGAAAAGTTAGGAAGGACACACAAGAGTGGAAACCCTCAACTGGATGGGACAACTGGGAAGAAGTTCCGTTTTGCTCCCACCATTTCAACAAGCTCCATCTCAAGGACGGGAGGTCCATTGTGGTTCCCTGCCGCCACCAAGATGAACTGATTGGCCGGGCCCGCGTCTCTCCAGGGGCGGGATGGAGCATCCGGGAGACTGCTTGCCTAGCAAAATCATATGCGCAAATGTGGCAGCTCCTTTATTTCCACAGAAGGGACCTCCGACTGATGGCCAATGCCATTTGTTCATCTGTGCCAGTTGACTGGGTTCCAACTGGGAGAACTACCTGGTCAATCCATGGAAAGGGAGAATGGATGACCACTGAAGACATGCTTGTGGTGTGGAACAGAGTGTGGATTGAGGAGAACGACCACATGGAAGACAAGACCCCAGTTACGAAATGGACAGACATTCCCTATTTGGGAAAAAGGGAAGACTTGTGGTGTGGATCTCTCATAGGGCACAGACCGCGCACCACCTGGGCTGAGAACATTAAAAACACAGTCAACATGGTGCGCAGGATCATAGGTGATGAAGAAAAGTACATGGACTACCTATCCACCCAAGTTCGCTACTTGGGTGAAGAAGGGTCTACACCTGGAGTGCTGTAA

>ENA|AMS00611|AMS00611.1 Zika virus polyprotein

ATGAAAAACCC-AAAAAAGAAATCCGGAGGATTCCGGATTGTCAATATGCTAAAACGCGGAGTAGCCCGTGTGAGCCCCTTTGGGGGCTTGAAGAGGCTGCCAGCCGGACTTCTGCTGGGTCATGGGCCCATCAGGATGGTCTTGGCGATTCTAGCCTTTTTGAGATTCACGGCAATCAAGCCATCACTGGGTCTCATCAATAGATGGGGTTCAGTGGGGAAAAAAGAGGCTATGGAAATAATAAAGAAGTTCAAGAAAGATCTGGCTGCCATGCTGAGAATAATCAATGCTAGGAAGGAGAAGAAGAGACGAGGCGCAGATACTAGTGTCGGAATTGTTGGCCTCCTGCTGACCACAGCTATGGCAGCGGAGGTCACTAGACGTGGGAGTGCATACTATATGTACTTGGACAGAAACGATGCTGGGGAGGCCATATCTTTTCCAACCACATTGGGGATGAATAAGTGTTATATACAGATCATGGATCTTGGACACATGTGTGATGCCACCATGAGCTATGAATGCCCTATGCTGGATGAGGGGGTGGAACCAGATGACGTCGATTGTTGGTGCAACACGACGTCAACTTGGGTTGTGTACGGAACCTGCCATCACAAAAAAGGTGAAGCACGGAGATCTAGAAGAGCTGTGACGCTCCCCTCCCATTCCACTAGGAAGCTGCAAACGCGGTCGCAAACCTGGTTGGAATCAAGAGAATACACAAAGCACTTGATTAGAGTCGAAAATTGGATATTCAGGAACCCTGGCTTCGCGTTAGCAGCAGCTGCCATCGCTTGGCTTTTGGGAAGCTCAACGAGCCAAAAAGTCATATACTTGGTCATGATACTGCTGATTGCCCCGGCATACAGCATCAGGTGCATAGGAGTCAGCAATAGGGACTTTGTGGAAGGTATGTCAGGTGGGACTTGGGTTGATGTTGTCTTGGAACATGGGGGTTGTGTCACCGTAATGGCACAGGACAAACCGACTGTCGACATAGAGCTGGTTACAACAACAGTCAGCAACATGGCGGAGGTAAGATCCTACTGCTATGAGGCATCAATATCAGACATGGCTTCGGACAGCCGCTGCCCAACACAAGGTGAAGCCTACCTTGACAAGCAATCAGACACTCAATATGTCTGCAAAAGAACGTTAGTGGACAGAGGCTGGGGAAATGGATGTGGACTTTTTGGCAAAGGGAGCCTGGTGACATGCGCTAAGTTTGCATGCTCCAAGAAAATGACCGGGAAGAGCATCCAGCCAGAGAATCTGGAGTACCGGATAATGCTGTCAGTTCATGGCTCCCAGCACAGTGGGATGATCGTTAATGACACAGGACATGAAACTGATGAGAATAGAGCGAAGGTTGAGATAACGCCCAATTCACCAAGAGCCGAAGCCACCCTGGGGGGTTTTGGAAGCCTAGGACTTGATTGTGAACCGAGGACAGGCCTTGACTTTTCAGATTTGTATTACTTGACTATGAATAACAAGCACTGGTTGGTTCACAAGGAGTGGTTCCACGACATTCCATTACCTTGGCACGCTGGGGCAGACACCGGAACTCCACACTGGAACAACAAAGAAGCACTGGTAGAGTTCAAGGACGCACATGCCAAAAGGCAAACTGTCGTGGTTCTAGGGAGTCAAGAAGGAGCAGTTCACACGGCCCTTGCTGGAGCTCTGGAGGCTGAGATGGATGGTGCAAAGGGAAGGCTGTCCTCTGGCCACTTGAAATGTCGCCTGAAAATGGATAAACTTAGATTGAAGGGCGTGTCATACTCCTTGTGTACCGCAGCGTTCACATTCACCAAGATCCCGGCTGAAACACTGCACGGGACAGTCACAGTGGAGGTACAGTACGCAGGGACAGATGGACCTTGCAAGGTTCCAGCTCAGATGGCGGTGGACATGCAAACTCTGACCCCAGTTGGGAGGTTGATAACCGCTAACCCCGTAATCACTGAAAGCACTGAGAACTCTAAGATGATGCTGGAACTTGATCCACCATTTGGGGACTCTTACATTGTCATAGGAGTCGGGGAGAAGAAGATCACCCACCACTGGCACAGGAGTGGCAGCACCATTGGAAAAGCATTTGAAGCCACTGTGAGAGGTGCCAAGAGAATGGCAGTCTTGGGAGACACAGCCTGGGACTTTGGATCAGTTGGAGGCGCTCTCAACTCATTGGGCAAGGGCATCCATCAAATTTTTGGAGCAGCTTTCAAATCATTGTTTGGAGGAATGTCCTGGTTCTCACAAATTCTCATTGGAACGTTGCTGATGTGGTTGGGTCTGAACACAAAGAATGGATCTATTTCCCTTATGTGCTTGGCCTTAGGGGGAGTGTTGATCTTCTTATCCACAGCCGTCTCTGCTGATGTGGGGTGTTCGGTGGACTTCTCAAAGAAGGAGACGAGATGTGGTACAGGGGTGTTCGTCTATAACGACGTTGAAGCCTGGAGGGACAGGTACAAGTACCATCCTGACTCCCCCCGTAGATTGGCAGCAGCAGTCAAGCAAGCCTGGGAAGATGGTATCTGCGGGATCTCCTCTGTTTCAAGAATGGAAAATATCATGTGGAGATCAGTAGAAGGGGAGCTCAACGCAATCCTGGAAGAGAATGGAGTTCAACTGACGGTCGTTGTGGGATCTGTAAAAAACCCCATGTGGAGAGGTCCACAGAGATTGCCCGTGCCTGTGAACGAGCTGCCCCACGGCTGGAAGGCTTGGGGGAAATCGTACTTCGTCAGAGCAGCAAAGACAAATAACAGCTTTGTCGTGGATGGTGACACACTGAAGGAATGCCCACTCAAACATAGAGCATGGAACAGCTTTCTTGTGGAGGATCATGGGTTCGGGGTATTTCACACTAGTGTCTGGCTCAAGGTTAGAGAAGATTATTCATTAGAGTGTGATCCAGCCGTTATTGGAACAGCTGTTAAGGGAAAGGAGGCTGTACACAGTGATCTAGGCTACTGGATTGAGAGTGAGAAGAATGACACATGGAGGCTGAAGAGGGCCCATCTGATCGAGATGAAAACATGTGAATGGCCAAAGTCCCACACATTGTGGACAGATGGAATAGAAGAGAGTGATCTGATCATACCCAAGTCTTTAGCTGGGCCACTCAGCCATCACAATACCAGAGAGGGCTACAGGACCCAAATGGAAGGGCCATGGCACAGTGAAGAGCTTGAAATTCGGTTTGAGGAATGCCCAGGCACTAAGGTCCACGTGGAGGAAACATGTGGAACAAGAGGACCATCTCTGAGATCAACCACTGCAAGCGGAAGGGTGATCGAGGAATGGTGCTGCAGGGAGTGCACAATGCCCCCACTGTCGTTCCGGGCTAAAGATGGCTGTTGGTATGGAATGGAGATAAGGCCCAGGAAAGAACCAGAAAGCAACTTAGTAAGGTCAATGGTGACTGCAGGATCAACTGATCACATGGATCACTTCTCCCTTGGAGTGCTTGTGATTCTGCTCATGGTGCAGGAAGGGCTGAAGAAGAGAATGACCACAAAGATCATCATAAGCACATCAATGGCAGTGCTGGTAGCTATGATCCTGGGAGGATTTTCAATGAGTGACCTGGCTAAGCTTGCAATTTTGATGGGTGCCACCTTCGCGGAAATGAACACTGGAGGAGATGTAGCTCATCTGGCGCTGATAGCGGCATTCAAAGTCAGACCAGCGTTGCTGGTATCTTTCATCTTCAGAGCTAATTGGACACCCCGTGAAAGCATGCTGCTGGCCTTGGCCTCGTGTCTTTTGCAAACTGCGATCTCCGCCTTGGAAGGCGACCTGATGGTTCTCATCAATGGTTTTGCTTTGGCCTGGTTGGCAATACGAGCGATGGTTGTTCCACGCACTGACAACATCACCTTGGCAATCCTGGCTGCTCTGACACCACTGGCCCGGGGCACACTGCTTGTGGCGTGGAGAGCAGGCCTTGCTACTTGCGGGGGGTTTATGCTCCTCTCTCTGAAGGGAAAAGGCAGTGTGAAGAAGAACTTACCATTTGTCATGGCCCTGGGACTAACCGCTGTGAGGCTGGTCGACCCCATCAACGTGGTGGGACTGCTGTTGCTCACAAGGAGTGGGAAGCGGAGCTGGCCCCCTAGCGAAGTACTCACAGCTGTTGGCCTGATATGCGCATTGGCTGGAGGGTTCGCCAAGGCAGATATAGAGATGGCTGGGCCCATGGCCGCGGTCGGTCTGCTAATTGTCAGTTACGTGGTCTCAGGAAAGAGTGTGGACATGTACATTGAAAGAGCAGGTGACATCACATGGGAAAAAGATGCGGAAGTCACTGGAAACAGTCCCCGGCTCGATGTGGCGCTAGATGAGAGTGGTGATTTCTCCCTGGTGGAGGATGACGGTCCCCCCATGAGAGAGATCATACTCAAGGTGGTCCTGATGACCATCTGTGGCATGAACCCAATAGCCATACCCTTTGCAGCTGGAGCGTGGTACGTATACGTGAAGACTGGAAAAAGGAGTGGTGCTCTATGGGATGTGCCTGCTCCCAAGGAAGTAAAAAAGGGGGAGACCACAGATGGAGTGTACAGAGTAATGACTCGCAGACTGCTAGGTTCAACACAAGTTGGAGTGGGAGTTATGCAAGAGGGGGTCTTTCACACTATGTGGCACGTCACAAAAGGATCCGCGCTGAGAAGCGGTGAAGGGAGACTTGATCCATACTGGGGAGATGTCAAGCAGGATCTGGTGTCATACTGTGGTCCGTGGAAGCTAGATGCCGCCTGGGACGGGCACAGCGAGGTGCAGCTCTTGGCCGTGCCCCCCGGAGAGAGAGCGAGGAACATCCAGACTCTGCCCGGAATATTTAAGACAAAGGATGGGGACATTGGAGCGGTTGCGCTGGATTACCCAGCAGGAACTTCAGGATCTCCAATCCTAGACAAGTGTGGGAGAGTGATAGGACTTTATGGCAATGGGGTCGTGATCAAAAATGGGAGTTATGTTAGTGCCATCACCCAAGGGAGGAGGGAGGAAGAGACTCCTGTTGATTGCTTCGAGCCTTCGATGCTGAAGAAGAAGCAGCTAACTGTCTTAGACTTGCATCCTGGAGCTGGGAAAACCAGGAGAGTTCTTCCTGAAATAGTCCGTGAAGCCATAAAAACAAGACTCCGTACTGTGATCTTAGCTCCAACCAGGGTTGTCGCTGCTGAAATGGAGGAAGCCCTTAGAGGGCTTCCAGTGCGTTATATGACAACAGCAGTCAATGTCACCCACTCTGGAACAGAAATCGTCGACTTAATGTGCCATGCCACCTTCACTTCACGTCTACTACAGCCAATCAGAGTCCCCAACTATAATCTGTATATTATGGATGAGGCCCACTTCACAGATCCCTCAAGTATAGCAGCAAGAGGATACATTTCAACAAGGGTTGAGATGGGCGAGGCGGCTGCCATCTTCATGACCGCCACGCCACCAGGAACCCGTGACGCATTTCCGGACTCCAACTCACCAATTATGGACACCGAAGTGGAAGTCCCAGAGAGAGCCTGGAGCTCAGGCTTTGATTGGGTGACGGATCATTCTGGAAAAACAGTTTGGTTTGTTCCAAGCGTGAGGAACGGCAATGAGATCGCAGCTTGTCTGACAAAGGCTGGAAAACGGGTCATACAGCTCAGCAGAAAGACTTTTGAGACAGAGTTCCAGAAAACAAAACATCAAGAGTGGGACTTTGTCGTGACAACTGACATTTCAGAGATGGGCGCCAACTTTAAAGCTGACCGTGTCATAGATTCCAGGAGATGCCTAAAGCCGGTCATACTTGATGGCGAGAGAGTCATTCTGGCTGGACCCATGCCTGTCACACATGCCAGCGCTGCCCAGAGGAGGGGGCGCATAGGCAGGAATCCCAACAAACCTGGAGATGAGTATCTGTATGGAGGTGGGTGCGCAGAGACTGACGAAGACCATGCACACTGGCTTGAAGCAAGAATGCTCCTTGACAATATTTACCTCCAAGATGGCCTCATAGCCTCGCTCTATCGACCTGAGGCCGACAAAGTAGCAGCCATTGAGGGAGAGTTCAAGCTTAGGACGGAGCAAAGGAAGACCTTTGTGGAACTCATGAAAAGAGGAGATCTTCCTGTTTGGCTGGCCTATCAGGTTGCATCTGCCGGAATAACCTACACAGATAGAAGATGGTGCTTTGATGGCACGACCAACAACACCATAATGGAAGACAGTGTGCCGGCAGAGGTGTGGACCAGACACGGAGAGAAAAGAGTGCTCAAACCGAGGTGGATGGACGCCAGAGTTTGTTCAGATCATGCGGCCCTGAAGTCATTCAAGGAGTTTGCCGCTGGGAAAAGAGGAGCGGCTTTTGGAGTGATGGAAGCCCTGGGAACACTGCCAGGACACATGACAGAGAGATTCCAGGAAGCCATTGACAACCTCGCTGTGCTCATGCGGGCAGAGACTGGAAGCAGGCCTTACAAAGCCGCGGCGGCCCAATTGCCGGAGACCCTAGAGACCATTATGCTTTTGGGGTTGCTGGGAACAGTCTCGCTGGGAATCTTTTTCGTCTTGATGAGGAACAAGGGCATAGGGAAGATGGGCTTTGGAATGGTGACTCTTGGGGCCAGCGCATGGCTCATGTGGCTCTCGGAAATTGAGCCAGCCAGAATTGCATGTGTCCTCATTGTTGTGTTCCTATTGCTGGTGGTGCTCATACCTGAGCCAGAAAAGCAAAGATCTCCCCAGGACAACCAAATGGCAATCATCATCATGGTAGCAGTAGGTCTTCTGGGCTTGATTACCGCCAATGAACTCGGATGGTTGGAGAGAACAAAGAGTGACCTAAGCCATCTAATGGGAAGGAGAGAGGAGGGGGCAACCATAGGATTCTCAATGGACATTGACCTGCGGCCAGCCTCAGCTTGGGCCATCTATGCTGCCTTGACAACTTTCATTACCCCAGCCGTCCAACATGCAGTGACCACTTCATACAACAACTACTCCTTAATGGCGATGGCCACGCAAGCTGGAGTGTTGTTTGGTATGGGCAAAGGGATGCCATTCTACGCATGGG-ACTTTGGAGTCCCGCTGCTAATGATAGGTTGCTACTCACAATTAACACCCCTGACCCTAATAGTGGCCATCATTTTGCTCGTGGCGCACTACATGTACTTGATCCCAGGGCTGCAGGCAGCAGCTGCGCGTGCTGCCCAGAAGAGAACGGCAGCTGGCATCATGAAGAACCCTGTTGTGGATGGAATAGTGGTGACTGACATTGACACAATGACAATTGACCCCCAAGTGGAGAAAAAGATGGGACAGGTGCTACTCATAGCAGTAGCCGTCTCCAGCGCCATACTGTCGCGGACCGCCTGGGGGTGGGGGGAGGCTGGGGCCCTGATCACAGCCGCAACTTCCACTTTGTGGGAAGGCTCTCCGAACAAGTACTGGAACTCCTCTACAGCCACTTCACTGTGTAACATTTTTAGGGGAAGTTACTTGGCTGGAGCTTCTCTAATCTACACAGTAACAAGAAACGCTGGCTTGGTCAAGAGACGTGGGGGTGGAACAGGAGAGACCCTGGGAGAGAAATGGAAGGCCCGCTTGAACCAGATGTCGGCCCTGGAGTTCTACTCCTACAAAAAGTCAGGCATCACCGAGGTGTGCAGAGAAGAGGCCCGCCGCGCCCTCAAGGACGGTGTGGCAACGGGAGGCCATGCTGTGTCCCGAGGAAGTGCAAAGCTGAGATGGTTGGTGGAGCGGGGATACCTGCAGCCCTATGGAAAGGTCATTGATCTTGGATGTGGCAGAGGGGGCTGGAGTTACTACGCCGCCACCATCCGCAAAGTTCAAGAAGTGAAAGGATACACAAAAGGAGGCCCTGGTCATGAAGAACCCGTGTTGGTGCAAAGCTATGGGTGGAACATAGTCCGTCTTAAGAGTGGGGTGGACGTCTTTCATATGGCGGCTGAGCCGTGTGACACGTTGCTGTGTGACATAGGTGAGTCATCATCTAGTCCTGAAGTGGAAGAAGCACGGACGCTCAGAGTCCTCTCCATGGTGGGGGATTGGCTTGAAAAAAGACCAGGAGCCTTTTGTATAAAAGTGTTGTGCCCATACACCAGCACTATGATGGAAACCCTGGAGCGACTGCAGCGTAGGTATGGGGGAGGACTGGTCAGAGTGCCACTCTCCCGCAACTCTACACATGAGATGTACTGGGTCTCTGGAGCGAAAAGCAACACCATAAAAAGTGTGTCCACCACGAGCCAGCTCCTCTTGGGGCGCATGGACGGGCCTAGGAGGCCAGTGAAATATGAGGAGGATGTGAATCTCGGCTCTGGCACGCGGGCTGTGGTAAGCTGCGCTGAAGCTCCCAATATGAAGATCATTGGTAACCGCATTGAAAGGATCCGCAGTGAGCACGCGGAAACGTGGTTCTTTGACGAGAACCACCCATATAGGACATGGGCTTACCATGGAAGCTATGAGGCCCCCACACAAGGGTCAGCGTCCTCTCTAATAAACGGGGTTGTCAGGCTCCTGTCAAAACCCTGGGATGTGGTGACTGGAGTCACAGGAATAGCCATGACCGACACCACACCGTATGGTCAGCAAAGAGTTTTCAAGGAAAAAGTGGACACTAGGGTGCCAGACCCCCAAGAAGGCACTCGTCAGGTTATGAGCATGGTCTCTTCCTGGTTGTGGAAAGAGCTAGGCAAACACAAACGGCCACGAGTCTGTACCAAAGAAGAGTTCATCAACAAGGTTCGTAGCAATGCAGCATTAGGGGCAATATTTGAAGAGGAAAAAGAGTGGAAGACTGCAGTGGAAGCTGTGAACGATCCAAGGTTCTGGGCTCTAGTGGACAAGGAAAGAGAGCACCACCTGAGAGGAGAGTGCCAGAGTTGTGTGTACAACATGATGGGAAAAAGAGAAAAGAAACAAGGGGAATTTGGAAAGGCCAAGGGCAGCCGCGCCATCTGGTATATGTGGCTAGGGGCTAGATTTCTAGAGTTCGAAGCCCTTGGATTCTTGAACGAGGATCACTGGATGGGGAGAGAGAACTCAGGAGGTGGTGTTGAAGGGCTGGGATTACAAAGACTCGGATATGTCCTAGAAGAGATGAGTCGCATACCAGGAGGAAGGATGTATGCAGATGACACTGCTGGCTGGGATACCCGCATCAGCAGGTTTGATCTGGAGAATGAAGCTCTAATCACCAACCAAATGGAGAAAGGGCACAGGGCCTTGGCATTGGCCATAATCAAGTACACATACCAAAACAAAGTGGTAAAGGTCCTTAGACCAGCTGAAAAAGGGAAAACAGTTATGGACATTATTTCGAGACAAGACCAAAGGGGGAGCGGACAAGTTGTCACTTACGCTCTTAACACATTTACCAACCTAGTGGTGCAACTCATTCGGAATATGGAGGCTGAGGAAGTTCTAGAGATGCAAGACTTGTGGCTGCTGCGGAGGTCAGAGAAAGTGACCAACTGGTTGCAGAGCAACGGATGGGATAGGCTCAAACGAATGGCAGTCAGTGGAGATGATTGCGTTGTGAAGCCAATTGATGATAGGTTTGCACATGCCCTCAGGTTCTTGAATGATATGGGAAAAGTTAGGAAGGACACACAAGAGTGGAAACCCTCAACTGGATGGGACAACTGGGAAGAAGTTCCGTTTTGCTCCCACCACTTCAACAAGCTCCATCTCAAGGACGGGAGGTCCATTGTGGTTCCCTGCCGCCACCAAGATGAACTGATTGGCCGGGCCCGCGTCTCTCCAGGGGCGGGATGGAGCATCCGGGAGACTGCTTGCCTAGCAAAATCATATGCGCAGATGTGGCAGCTCCTTTATTTCCACAGAAGGGACCTCCGACTGATGGCCAATGCCATTTGTTCATCTGTGCCAGTTGACTGGGTTCCAACTGGGAGAACTACCTGGTCAATCCATGGAAAGGGAGAATGGATGACCACTGAAGACATGCTTGTGGTGTGGAACAGAGTGTGGATTGAGGAGAACGACCACATGGAAGACAAGACCCCAGTTACGAAATGGACAGACATTCCCTATTTGGGAAAAAGGGAAGACTTGTGGTGTGGATCTCTCATAGGGCACAGACCGCGCACCACCTGGGCTGAGAACATTAAAAACACAGTCAACATGGTGCGCAGGATCATAGGTGATGAAGAAAAGTACATGGACTACCTATCCACCCAAGTTCGCTACTTGGGTGAAGAAGGGTCTACACCTGGAGTGCTGTAA

>ENA|AMQ34004|AMQ34004.1 Zika virus polyprotein

ATGAAAAACCC-AAAAAAGAAATCCGGAGGATTCCGGATTGTCAATATGCTAAAACGCGGAGTAGCCCGTGTGAGCCCCTTTGGGGGCTTGAAGAGGCTGCCAGCCGGACTTCTGCTGGGTCATGGGCCCATCAGGATGGTCTTGGCGATTCTAGCCTTTTTGAGATTCACGGCAATCAAGCCATCACTGGGTCTCATCAATAGATGGGGTTCAGTGGGGAAAAAAGAGGCTATGGAAATAATAAAGAAGTTCAAGAAAGATCTGGCTGCCATGCTGAGAATAATCAATGCTAGGAAGGAGAAGAAGAGACGAGGCGCAGAAACTAGTGTCGGAATTGTTGGCCTCCTGCTGACCACAGCTATGGCAGCGGAGGTCACTAGACGTGGGAGTGCATACTATATGTACTTGGACAGAAACGATGCTGGGGAGGCCATATCTTTTCCAACCACATTGGGGATGAATAAGTGTTATATACAGATCATGGATCTTGGACACATGTGTGATGCCACCATGAGCTATGAATGCCCTATGCTGGATGAGGGGGTGGAACCAGATGACGTCGATTGTTGGTGCAACACGACGTCAACTTGGGTTGTGTACGGAACCTGCCATCACAAAAAAGGTGAAGCACGGAGATCTAGAAGAGCCGTGACGCTCCCCTCCCATTCCACTAGGAAGCTGCAAACGCGGTCGCAAACCTGGTTGGAATCAAGAGAATACACAAAGCACTTGATTAGAGTCGAAAATTGGATATTCAGGAACCCTGGTTTCGCTTTAGCAGCAGCTGCCATCGCGTGGCTTTTGGGAAGCTCAACGAGCCAAAAAGTCATATACTTGGTCATGATACTGCTGATTGCCCCGGCATACAGCATCAGGTGCATAGGAGTCAGCAATAGGGACTTTGTGGAAGGTATGTCAGGTGGGACTTGGGTTGATGTTGTCTTGGAACATGGAGGTTGTGTCACCGTAATGGCACAGGACAAACCGACTGTCGACATAGAGCTGGTTACAACAACAGTCAGCAACATGGCGGAGGTAAGATCCTACTGCTATGAGGCATCAATATCAGACATGGCTTCGGACAGCCGCTGCCCAACACAAGGTGAAGCCTACCTTGACAAGCAATCAGACACTCAATATGTCTGCAAAAGAACGTTAGTGGACAGAGGCTGGGGAAATGGATGTGGACTTTTTGGCAAAGGGAGCCTGGTGACATGCGCTAAGTTTGCATGCTCCAAGAAAATGACCGGGAAGAGCATCCAGCCAGAGAATCTGGAGTACCGGATAATGTTGTCAGTTCATGGCTCCCAGCACAGTGGGATGATCGTTAATGACACAGGACATGAAACTGATGAGAATAGAGCGAAGGTTGAGATAACGCCCAATTCACCAAGAGCCGAAGCCACCCTGGGGGGTTTTGGAAGCCTAGGACTTGATTGTGAACCGAGGACAGGCCTTGACTTTTCAGATTTGTATTACTTGACTATGAATAACAAGCACTGGTTGGTTCACAAGGAGTGGTTCCACGACATTCCATTACCTTGGCACGCTGGGGCAGACACCGGAACTCCACACTGGAACAACAAAGAAGCACTGGTAGAGTTCAAGGACGCACATGCCAAAAGGCAAACTGTCGTGGTTCTAGGGAGTCAAGAAGGAGCAGTTCACACGGCCCTTGCTGGAGCTCTGGAGGCTGAGATGGATGGTGCAAAGGGAAGGCTGTCCTCTGGCCACTTGAAATGTCGCCTGAAAATGGATAAACTTAGATTGAAGGGCGTGTCATACTCCTTGTGTACCGCAGCGTTCACATTCACCAAGATCCCGGCTGAAACACTGCACGGGACAGTCACAGTGGAGGTACAGTACGCAGGGACAGATGGACCTTGCAAGGTTCCAGCCCAGATGGCGGTGGACATGCAAACTCTGACCCCAGTTGGGAGGTTGATAACCGCTAACCCCGTAATCACTGAAAGCACTGAGAACTCTAAGATGATGCTGGAACTTGATCCACCATTTGGGGACTCTTACATTGTCATAGGAGTCGGGGAGAAGAAGATCACCCACCACTGGCACAGGAGTGGCAGCACCATTGGAAAAGCATTTGAAGCCACTGTGAGAGGTGCCAAGAGAATGGCAGTCTTGGGAGACACAGCCTGGGACTTTGGATCAGTTGGAGGCGCTCTCAACTCATTGGGCAAGGGCATCCATCAAATTTTTGGAGCAGCTTTCAAATCATTGTTTGGAGGAATGTCCTGGTTCTCACAAATTCTCATTGGAACGTTGCTGATGTGGTTGGGTCTGAACACAAAGAATGGATCTATTTCCCTTATGTGCTTGGCCTTAGGGGGAGTGTTGATCTTCTTATCCACAGCCGTCTCTGCTGATGTGGGGTGCTCGGTGGACTTCTCAAAGAAGGAGACGAGATGTGGTACAGGGGTGTTCGTCTATAACGACGTTGAAGCCTGGAGGGACAGGTACAAGTACCATCCTGACTCCCCCCGTAGATTGGCAGCAGCAGTCAAGCAAGCCTGGGAAGATGGTATCTGCGGGATCTCCTCTGTTTCAAGAATGGAAAACATCATGTGGAGATCAGTAGAAGGGGAGCTCAACGCAATCCTGGAAGAGAATGGAGTTCAACTGACGGTCGTTGTGGGATCTGTAAAAAACCCCATGTGGAGAGGTCCACAGAGATTGCCCGTGCCTGTGAACGAGCTGCCCCACGGCTGGAAGGCTTGGGGGAAATCGTACTTCGTCAGAGCAGCAAAGACAAATAACAGCTTTGTCGTGGATGGTGACACACTGAAGGAATGCCCACTCAAACATAGAGCATGGAACAGCTTTCTTGTGGAGGATCATGGGTTCGGGGTATTTCACACTAGTGTCTGGCTCAAGGTTAGAGAAGATTATTCATTAGAGTGTGATCCAGCCGTTATTGGAACAGCTGTTAAGGGAAAGGAGGCTGTACACAGTGATCTAGGCTACTGGATTGAGAGTGAGAAGAATGACACATGGAGGCTGAAGAGGGCCCATCTGATCGAGATGAAAACATGTGAATGGCCAAAATCCCACACATTGTGGACAGATGGAATAGAAGAGAGTGATCTGATCATACCCAAGTCTTTAGCTGGGCCACTCAGCCATCACAATACCAGAGAGGGCTACAGGACCCAAATGAAAGGGCCATGGCACAGTGAAGAGCTTGAAATTCGGTTTGAGGAATGCCCAGGCACTAAGGTCCACGTGGAGGAAACATGTGGAACAAGAGGACCATCTCTGAGATCAACCACTGCAAGCGGAAGGGTGATCGAGGAATGGTGCTGCAGGGAGTGCACAATGCCCCCACTGTCGTTCTGGGCTAAAGATGGCTGTTGGTATGGAATGGAGATAAGGCCCAGGAAAGAACCAGAAAGCAACTTAGTAAGGTCAATGGTGACTGCAGGATCAACTGATCACATGGATCACTTCTCCCTTGGAGTGCTTGTGATTCTGCTCATGGTGCAGGAAGGGCTGAAGAAGAGAATGACCACAAAGATCATCATAAGCACATCAATGGCAGTGCTGGTAGCTATGATCCTGGGAGGATTTTCAATGAGTGACCTGGCTAAGCTTGCAATTTTGATGGGTGCCACCTTCGCGGAAATGAACACTGGAGGAGATGTAGCTCATCTGGCGCTGACAGCGGCATTCAAAGTCAGACCAGCGTTGCTGGTATCTTTCATCTTCAGAGCTAATTGGACACCCCGTGAAAGCATGCTGCTGGCCTTGGCCTCGTGTCTTTTGCAAACTGCGATCTCTGCCTTGGAAGGCGACCTGATGGTTCTCATCAATGGTTTTGCTTTGGCCTGGTTGGCAATACGAGCGATGGTTGTTCCACGCACTGACAACATCACCTTGGCAATCCTGGCTGCTCTGACACCACTGGCCCGGGGCACACTGCTTGTGGCGTGGAGAGCAGGCCTTGCTACTTGCGGGGGGTTTATGCTCCTCTCTCTGAAGGGAAAAGGCAGTGTGAAGAAGAACTTACCATTTGTCATGGCCCTGGGACTAACCGCTGTGAGGCTGGTCGACCCCATCAACGTGGTGGGACTGCTGTTGCTCACAAGGAGTGGGAAGCGGAGCTGGCCCCCTAGCGAAGTACTCACAGCTGTTGGCCTGATATGTGCATTGGCTGGAGGGTTCGCCAAGGCAGATATAGAGATGGCTGGGCCCATGGCCGCGGTCGGTCTGCTAATTGTCAGTTACGTGGTCTCAGGAAAGAGTGTGGACATGTACATTGAAAGAGCAGGTGACATCACATGGGAAAAAGATGCGGAAGTCACTGGAAACAGTCCCCGGCTCGACGTGGCGCTAGATGAGAGTGGTGATTTCTCCCTGGTGGAGGATGACGGTCCCCCCATGAGAGAGATCATACTCAAGGTGGTCCTGATGACCATCTGTGGCATGAACCCAATAGCCATACCCTTTGCAGCTGGAGCGTGGTACGTATACGTGAAGACTGGAAAAAGGAGTGGTGCTCTATGGGATGTGCCTGCTCCCAAGGAAGTAAAAAAGGGGGAGACCACAGATGGAGTGTACAGAGTAATGACTCGTAGACTGCTAGGTTCAACACAAGTTGGAGTGGGAGTTATGCAAGAGGGGGTCTTTCACACTATGTGGCACGTCACAAAAGGATCCGCGCTGAGAAGCGGTGAAGGGAGACTTGATCCATACTGGGGAGATGTCAAGCAGGATCTGGTGTCATACTGTGGTCCATGGAAGCTAGATGCCGCCTGGGACGGGCACAGCGAGGTGCAGCTCTTGGCCGTGCCCCCCGGAGAGAGAGAGAGGAACATCCAGACTCTGCCCGGAATATTTAAGACAAAGGATGGGGACATTGGAGCGGTTGCGCTGGATTACCCAGCAGGAACTTCAGGATCTCCAATCCTAGACAAGTGTGGGAGAGTGATAGGACTTTATGGCAATGGGGTCGTGATCAAAAATGGGAGTTATGTTAGTGCCATCACCCAAGGGAGGAGGGAGGAAGAGACTCCTGTTGAGTGCTTCGAGCCTTCGATGCTGAAGAAGAAGCAGCTAACTGTCTTAGACTTGCATCCTGGAGCTGGGAAAACCAGGAGAGTTCTTCCTGAAATAGTCCGTGAAGCCATAAAAACAAGACTCCGTACTGTGATCTTAGCTCCAACCAGGGTTGTCGCTGCTGAAATGGAGGAAGCCCTTAGAGGGCTTCCAGTGCGTTATATGACAACAGCAGTCAATGTCACCCACTCTGGAACAGAAATCGTCGACTTAATGTGCCATGCCACCTTCACTTCACGTCTACTACAGCCAATCAGAGTCCCCAACTATAATCTGTATATTATGGATGAGGCCCACTTCACAGATCCCTCAAGTATAGCAGCAAGAGGATACATTTCAACAAGGGTTGAGATGGGCGAGGCGGCTGCCATCTTCATGACCGCCACGCCACCAGGAACCCGTGACGCATTTCCGGACTCCAACTCACCAATTATGGACACCGAAGTGGAAGTCCCAGAGAGAGCCTGGAGCTCAGGCTTTGATTGGGTGACGGATCATTCTGGAAAAACAGTTTGGTTTGTTCCAAGCGTGAGGAACGGCAATGAGATCGCAGCTTGTCTGACAAAGGCTGGAAAACGGGTCATACAGCTCAGCAGAAAGACTTTTGAGACAGAGTTCCAGAAAACAAAACATCAAGAGTGGGACTTTGTCGTGACAACTGACATTTCAGAGATGGGCGCCAACTTTAAAGCTGACCGTGTCATAGATTCCAGGAGATGCCTAAAGCCGGTCATACTTGATGGCGAGAGAGTCATTCTGGCTGGACCCATGCCTGTCACACATGCCAGCGCTGCCCAGAGGAGGGGGCGCATAGGCAGGAATCCCAATAAACCTGGAGATGAGTATCTGTATGGAGGTGGGTGCGCAGAGACTGACGAAGACCATGCACACTGGCTTGAAGCAAGAATGCTCCTTGACAATATTTACCTCCAAGATGGCCTCATAGCCTCGCTCTATCGACCTGAGGCCGACAAAGTAGCAGCCATTGAGGGAGAGTTCAAGCTTAGGACGGAGCAAAGGAAGACCTTTGTGGAACTCATGAAAAGAGGAGATCTTCCTGTTTGGCTGGCCTATCAGGTTGCATCTGCCGGAATAACCTACACAGATAGAAGATGGTGCTTTGATGGCACGACCAACAACACCATAATGGAAGACAGTGTGCCGGCAGAGGTGTGGACCAGACACGGAGAGAAAAGAGTGCTCAAACCGAGGTGGATGGACGCCAGAGTTTGTTCAGATCATGCGGCCCTGAAGTCATTCAAGGAGTTTGCCGCTGGGAAAAGAGGAGCGGCTTTTGGAGTGATGGAAGCCCTGGGAACACTGCCAGGACACATGACAGAGAGATTCCAGGAAGCCATTGACAACCTCGCTGTGCTCATGCGGGCAGAGACTGGAAGCAGGCCTTACAAAGCCGCGGCGGCCCAATTGCCGGAGACCCTAGAGACCATTATGCTTTTGGGGTTGCTGGGAACAGTCTCGCTGGGAATCTTTTTCGTCTTGATGAGGAACAAGGGCATAGGGAAGATGGGCTTTGGAATGGTGACTCTTGGGGCCAGCGCATGGCTCATGTGGCTCTCGGAAATTGAGCCAGCCAGAATTGCATGTGTCCTCATTGTTGTGTTCCTATTGCTGGTGGTGCTCATACCTGAGCCAGAAAAGCAAAGATCTCCCCAGGACAACCAAATGGCAATCATCATCATGGTAGCAGTAGGTCTTCTGGGCTTGATTACCGCCAATGAACTCGGATGGTTGGAGAGAACAAAGAGTGACCTAAGCCATCTAATGGGAAGGAGAGAGGAGGGGGCAACCATAGGATTCTCAATGGACATTGACCTGCGGCCAGCCTCAGCTTGGGCCATCTATGCTGCCTTGACAACTTTCATTACCCCAGCCGTCCAACATGCAGTGACCACTTCATACAACAACTACTCCTTAATGGCGATGGCCACGCAAGCTGGAGTGTTGTTTGGTATGGGCAAAGGGATGCCATTCTACGCATGGG-ACTTTGGAGTCCCACTGCTAATGATAGGTTGCTACTCACAATTAACACCCCTGACCCTAATAGTGGCCATCATTTTGCTCGTGGCGCACTACATGTACTTGATCCCAGGGCTGCAGGCAGCAGCTGCGCGTGCTGCCCAGAAGAGAACGGCAGCTGGCATCATGAAGAACCCTGTTGTGGATGGAATAGTGGTGACTGACATTGACACAATGACAATTGACCCCCAAGTGGAGAAAAAGATGGGACAGGTGCTACTCATAGCAGTAGCCGTCTCCAGCGCCATACTGTCGCGGACCGCCTGGGGGTGGGGGGAGGCTGGGGCCCTGATCACAGCCGCAACTTCCACTTTGTGGGAAGGCTCTCCGAACAAGTACTGGAACTCCTCTACAGCCACTTCACTGTGTAACATTTTTAGGGGAAGTTACTTGGCTGGAGCTTCTCTAATCTACACAGTAACAAGAAACGCTGGCTTGGTCAAGAGACGTGGGGGTGGAACAGGAGAGACCCTGGGAGAGAAATGGAAGGCCCGCTTGAACCAGATGTCGGCCCTGGAGTTCTACTCCTACAAAAAGTCAGGCATCACCGAGGTGTGCAGAGAAGAGGCCCGCCGCGCCCTCAAGGACGGTGTGGCAACGGGAGGCCATGCTGTGTCCCGAGGAAGTGCAAAGCTGAGATGGTTGGTGGAGCGGGGATACCTGCAGCCCTATGGAAAGGTCATTGATCTTGGATGTGGCAGAGGGGGCTGGAGTTACTACGCCGCCACCATCCGCAAAGTTCAAGAAGTGAAAGGATACACAAAAGGAGGCCCTGGTCATGAAGAACCCGTGTTGGTGCAAAGCTATGGGTGGAACATAGTCCGTCTTAAGAGTGGGGTGGACGTCTTTCATATGGCGGCTGAGCCGTGTGACACGTTGCTGTGTGACATAGGTGAGTCATCATCTAGTCCTGAAGTGGAAGAAGCACGGACGCTCAGAGTCCTCTCCATGGTGGGGGATTGGCTTGAAAAAAGACCAGGAGCCTTTTGTATAAAAGTGTTGTGCCCATACACCAGCACTATGATGGAAACCCTGGAGCGACTGCAGCGTAGGTATGGGGGAGGACTGGTCAGAGTGCCACTCTCCCGCAACTCTACACATGAGATGTACTGGGTCTCTGGAGCGAAAAGCAACACCATAAAAAGTGTGTCCACCACGAGCCAGCTCCTCTTGGGGCGCATGGACGGGCCTAGGAGGCCAGTGAAATATGAGGAGGATGTGAATCTCGGCTCTGGCACGCGGGCTGTGGTAAGCTGCGCTGAAGCTCCCAACATGAAGATCATTGGTAACCGCATTGAAAGGATCCGCAGTGAGCAAGCGGAAACGTGGTTCTTTGACGAGAACCACCCATATAGGACATGGGCTTACCATGGAAGCTATGAGGCCCCCACACAAGGGTCAGCGTCCTCTCTAATAAACGGGGTTGTCAGGCTCCTGTCAAAACCCTGGGATGTGGTGACTGGAGTCACAGGAATAGCCATGACCGACACCACACCGTATGGTCAGCAAAGAGTTTTCAAGGAAAAAGTGGACACTAGGGTGCCAGACCCCCAAGAAGGCACTCGTCAGGTTATGAGCATGGTCTCTTCCTGGTTGTGGAAAGAGCTAGGCAAACACAAACGGCCACGAGTCTGTACCAAAGAAGAGTTCATCAACAAGGTTCGTAGCAATGCAGCATTAGGGGCAATATTTGAAGAGGAAAAAGAGTGGAAGACTGCAGTGGAAGCTGTGAACGATCCAAGGTTCTGGGCTCTAGTGGACAAGGAAAGAGAGCACCACCTGAGAGGAGAGTGCCAGAGTTGTGTGTACAACATGATGGGAAAAAGAGAAAAGAAACAAGGGGAATTTGGAAAGGCCAAGGGCAGCCGCGCCATCTGGTATATGTGGCTAGGGGCTAGATTTCTAGAGTTCGAAGCCCTTGGATTCTTGAACGAGGATCACTGGATGGGGAGAGAGAACTCAGGAGGTGGTGTTGAAGTGCTGGGATTACAGAGACTCGGATATGTCCTAGAAGAGATGAGTCGCATACCAGGAGGAAGGATGTATGCAGATGACACTGCTGGCTGGGACACCCGCATCAGCAGGTTTGATCTGGAGAATGAAGCTCTAATCACCAACCAAATGGAGAAAGGGCACAGGGCCTTGGCATTGGCCATAATCAAGTACACATACCAAAACAAAGTGGTAAAGGTCCTTAGACCAGCTGAAAAAGGGAAAACAGTTATGGACATTATTTCGAGACAAGACCAAAGGGGGAGCGGACAAGTTGTCACTTACGCTCTTAACACATTTACCAACCTAGTGGTGCAACTCATTCGGAATATGGAGGCTGAGGAAGTTCTAGAGATGCAAGACTTGTGGCTGCTGCGGAGGTCAGAGAAAGTGACCAACTGGTTGCAGAGCAACGGATGGGATAGGCTCAAACGAATGGCAGTCAGTGGAGATGATTGCGTTGTGAAGCCAATTGATGATAGGTTTGCACATGCCCTCAGGTTCTTGAATGATATGGGAAAAGTTAGGAAGGACACACAAGAGTGGAAACCCTCAACTGGATGGGACAACTGGGAAGAAGTTCCGTTTTGCTCCCACCACTTCAACAAGCTCCATCTCAAGGACGGGAGGTCCATTGTGGTTCCCTGCCGCCACCAAGATGAACTGATTGGCCGGGCCCGCGTCTCTCCAGGGGCGGGATGGAGCATCCGGGAGACTGCTTGCCTAGCAAAATCATATGCGCAAATGTGGCAGCTCCTTTATTTCCACAGAAGGGACCTCCGACTGATGGCCAATGCCATTTGTTCATCTGTGCCAGTTGACTGGGTTCCAACTGGGAGAACTACCTGGTCAATCCATGGAAAGGGAGAATGGATGACCACTGAAGACATGCTTGTGGTGTGGAACAGAGTGTGGATTGAGGAGAACGACCACATGGAAGACAAGACCCCAGTTGCGAAATGGACAGACATTCCCTATTTGGGAAAAAGGGAAGACTTGTGGTGTGGATCTCTCATAGGGCACAGACCGCGCACCACCTGGGCTGAGAACATTAAAAACACAGTCAACATGGTGCGCAGGATCATAGGTGATGAAGAAAAGTACATGGACTACCTATCCACCCAAGTTCGCTACTTGGGTGAAGAAGGGTCTACACCTGGAGTGCTGTAA

>ENA|AMQ48986|AMQ48986.1 Zika virus polyprotein

ATGAAAAACCC-AAAAAAGAAATCCGGAGGATTCCGGATTGTCAATATGCTAAAACGCGGAGTAGCCCGTGTGAGCCCCTTTGGGGGCTTGAAGAGGCTGCCAGCCGGACTTCTGCTGGGTCATGGGCCCATCAGGATGGTCTTGGCGATTCTAGCCTTTTTGAGATTCACGGCAATCAAGCCATCACTGGGTCTCATCAATAGATGGGGTTCAGTGGGGAAAAAAGAGGCTATGGAAATAATAAAGAAGTTCAAGAAAGATCTGGCTGCCATGTTGAGAATAATCAATGCTAGGAAGGAGAAGAAGAGACGAGGCGCAGATACTAGTGTCGGAATTGTTGGCCTCCTGCTGACCACAGCTATGGCAGCGGAGGTCACTAGACGTGGGAGTGCATACTATATGTACTTGGACAGAAACGATGCTGGGGAGGCCATATCTTTTCCAACCACATTGGGGATGAATAAGTGTTATATACAGATCATGGATCTTGGACACATGTGTGATGCCACCATGAGCTATGAGTGCCCTATGCTGGATGAGGGGGTGGAACCAGATGACGTCGATTGTTGGTGCAACACGACGTCAACTTGGGTTGTGTACGGAACCTGCCATCACAAAAAAGGTGAAGCACGGAGATCTAGAAGAGCTGTGACGCTCCCCTCCCATTCCACTAGGAAGCTGCAAACGCGGTCGCAAACCTGGTTGGAATCAAGAGAATACACAAAGCACTTGATTAGAGTCGAAAATTGGATACTCAGGAACCCTGGCTTCGCGTTAGCAGCAGCTGCCATCGCTTGGCTTTTGGGAAGCTCAACGAGCCAAAAAGTCATATATTTGGTCATGATACTGCTGATTGCCCCGGCATACAGCATCAGGTGCATAGGAGTCAGCAATAGGGACTTTGTGGAAGGTATGTCAGGTGGGACTTGGGTTGATGTTGTCTTGGAACATGGAGGTTGTGTCACCGTAATGGCACAGGACAAACCGACTGTCGACATAGAGCTGGTTACAACAACAGTCAGCAACATGGCGGAGGTAAGATCCTACTGCTATGAGGCATCAATATCAGACATGGCTTCGGACAGCCGCTGCCCAACACAAGGTGAAGCCTACCTTGACAAGCAATCAGACACTCAATATGTCTGCAAAAGAACGTTAGTGGACAGAGGCTGGGGAAATGGATGTGGACTTTTTGGCAAAGGGAGCCTGGTGACATGCGCTAAGTTTGCATGCTCCAAGAAAATGACCGGGAAGAGCATCCAGCCAGAGAATCTGGAGTACCGGATAATGCTGTCAGTTCATGGCTCCCAGCACAGTGGGATGATCGTTAATGACACAGGACATGAAACTGATGAGAATAGAGCGAAGGTTGAGATAACGCCCAATTCACCAAGAGCCGAAGCCACCCTGGGGGGTTTTGGAAGCCTAGGACTTGATTGTGAACCGAGGACAGGCCTTGACTTTTCAGATTTGTATTACTTGACTATGAATAACAAGCACTGGTTGGTTCACAAGGAGTGGTTCCACGACATTCCATTACCTTGGCACGCTGGGGCAGACACCGGAACTCCACACTGGAACAACAAAGAAGCACTGGTAGAGTTCAAGGACGCACATGCCAAAAGGCAAACTGTCGTGGTTCTAGGGAGTCAAGAAGGAGCAGTTCACACGGCCCTTGCTGGAGCTCTGGAGGCTGAGATGGATGGTGCAAAGGGAAGGCTGTCCTCTGGCCACTTGAAATGTCGCCTGAAAATGGATAAACTCAGATTGAAGGGCGTGTCATACTCCTTGTGTACCGCAGCGTTCACATTCACCAAGATCCCGGCTGAAACACTGCACGGGACAGTCACAGTGGAGGTACAGTACGCAGGGACAGATGGACCTTGCAAGGTTCCAGCTCAGATGGCGGTGGACATGCAAACTCTGACCCCAGTTGGGAGGTTGATAACCGCTAACCCCGTAATCACTGAAAGCACTGAGAACTCTAAGATGATGCTGGAACTTGATCCACCATTTGGGGACTCTTACATTGTCATAGGAATCGGGGAGAAGAAGATCACCCACCACTGGCACAGGAGTGGCAGCACCATTGGAAAAGCATTTGAAGCCACTGTGAGAGGTGCCAAGAGAATGGCAGTCTTGGGAGACACAGCCTGGGACTTTGGATCAGTTGGAGGCGCTCTCAACTCATTGGGCAAGGGCATCCATCAAATTTTTGGAGCAGCTTTCAAATCATTGTTTGGAGGAATGTCCTGGTTCTCACAAATTCTCATTGGAACGTTGCTGATGTGGTTGGGTCTGAACACAAAGAATGGATCTATTTCCCTTATGTGCTTGGCCTTAGGGGGAGTGTTGATCTTCTTATCCACAGCCGTCTCTGCTGATGTGGGGTGCTCGGTGGACTTCTCAAAGAAGGAGACGAGATGCGGTACAGGGGTGTTCGTCTATAACGACGTTGAAGCCTGGAGGGACAGGTACAAGTACCATCCTGACTCCCCCCGTAGATTGGCAGCAGCAGTCAAGCAAGCCTGGGAAGATGGTATCTGCGGGATCTCCTCTGTTTCAAGAATGGAAAACATCATGTGGAGATCAGTAGAAGGGGAGCTCAACGCAATCCTGGAAGAGAATGGAGTTCAACTGACGGTCGTTGTGGGATCTGTAAAAAACCCCATGTGGAGAGCTCCACAGAGATTGCCCGTGCCTGTGAACGAGCTGCCCCACGGCTGGAAGGCTTGGGGGAAATCGTACTTCGTCAGAGCAGCAAAGACAAATAACAGCTTTGTCGTGGATGGTGACACACTGAAGGAATGCCCACTCAAACATAGAGCATGGAACAGCTTTCTTGTGGAGGATCATGGGTTCGGGGTATTTCACACTAGTGTCTGGCTCAAGGTTAGAGAAGATTATTCATTAGAGTGTGATCCAGCCGTTATTGGAACAGCTGTTAAGGGAAAGGAGGCTGTACACAGTGATCTAGGCTACTGGATTGAGAGTGAGAAGAATGACACATGGAGGCTGAAGAGGGCCCATCTGATCGAGATGAAAACATGTGAATGGCCACAGTCCCACACATTGTGGACAGATGGAATAGAAGAGAGTGATCTGATCATACCCAAGTCTTTAGCTGGGCCACTCAGCCATCACAATACCAGAGAGGGCTACAGGACCCAAATGAAAGGGCCATGGCACAGTGAAGAGCTTGAAATTCGGTTTGAGGAATGCCCAGGCACTAAGGTCCACGTGGAGGAAACATGTGGAACAAGAGGACCATCTCTGAGATCAACCACTGCAAGCGGAAGGGTGATCGAGGAATGGTGCTGCAGGGAGTGCACAATGCCCCCACTGTCGTTCCGGGCTAAAGATGGCTGTTGGTATGGAATGGAGATAAGGCCCAGGAAAGAACCAGAAAGCAACTTAGTAAGGTCAATGGTGACTGCAGGATCAACTGATCACATGGATCACTTCTCCCTTGGAGTGCTTGTGATTCTGCTCATGGTGCAGGAAGGGCTAAAGAAGAGAATGACCACAAAGATCATCATAAGCACATCAATGGCAGTGCTGGTAGCTATGATCCTGGGAGGATTTTCAATGAGTGACCTGGCTAAGCTTGCAATTTTGATGGGTGCCACCTTCGCGGAAATGAACACTGGAGGAGATGTAGCTCATCTGGCGCTGATAGCGGCATTCAAAGTCAGACCAGCGTTGCTGGTATCTTTCATCTTCAGAGCTAATTGGACACCCCGTGAAAGCATGCTACTGGCCTTGGCCTCGTGTCTTTTGCAAACTGCGATCTCCGCCTTGGAAGGCGACCTGATGGTTCTCATCAATGGTTTTGCTTTGGCCTGGTTGGCAATACGAGCGATGGTTGTTCCACGCACTGATAACATCACCTTGGCAATCCTGGCTGCTCTGACACCACTGGCCCGGGGCACACTGCTTGTGGCGTGGAGAGCAGGCCTTGCTACTTGCGGGGGGTTTATGCTCCTCTCTCTGAAGGGAAAAGGCAGTGTGAAGAAGAACTTACCATTTGTCATGGCCCTGGGACTAACCGCTGTGAGGCTGGTCGACCCCATCAACGTGGTGGGACTGCTGTTGCTCACAAGGAGTGGGAAGCGGAGCTGGCCCCCTAGCGAAGTACTCACAGCTGTTGGCCTGATATGCGCATTGGCTGGAGGGTTCGCCAAGGCAGATATAGAGATGGCTGGGCCCGTGGCCGCGGTCGGTCTGCTAATTGTCAGTTACGTGGTCTCAGGAAAGAGTGTGGACATGTACATTGAAAGAGCAGGTGACATCACATGGGAAAAAGATGCGGAAGTCACTGGAAACAGTCCCCGGCTCGATGTGGCGCTAGATGAGAGTGGTGATTTCTCCCTGGTGGAGGATGACGGTCCCCCCATGAGAGAGATCATACTCAAGGTGGTCCTGATGACCATCTGTGGCATGAACCCAATAGCCATACCCTTTGCAGCTGGAGCGTGGTACGTATACGTGAAGACTGGAAAAAGGAGTGGTGCTCTATGGGATGTGCCTGCTCCCAAGGAAGTAAAAAAGGGGGAGACCACAGATGGAGTGTACAGAGTAATGACTCGTAGACTGCTAGGTTCAACACAAGTTGGAGTGGGAGTTATGCAAGAGGGGGTCTTTCACACTATGTGGCACGTCACAAAAGGATCCGCACTGAGAAGCGGTGAAGGGAGACTTGATCCATACTGGGGAGATGTCAAGCAGGATCTGGTGTCATACTGTGGTCCATGGAAGCTAGATGCCGCCTGGGACGGGCACAGCGAGGTGCAGCTCTTGGCCGTGCCCCCCGGAGAGAGAGCGAGGAACATCCAGACTCTGCCCGGAATATTTAAGACAAAGGATGGGGACATTGGAGCGGTTGCGCTGGATTACCCAGCAGGAACTTCAGGATCTCCAATCCTAGACAAGTGTGGGAGAGTGATAGGACTTTATGGCAATGGGGTCGTGATCAAAAATGGGAGTTATGTTAGTGCCATCACCCAAGGGAGGAGGGAGGAAGAGACTCCTGTTGAGTGCTTCGAGCCTTCGATGCTGAAGAAGAAGCAGCTAACTGTCTTAGACTTACATCCTGGAGCTGGGAAAACCAGGAGAGTTCTTCCTGAAATAGTCCGTGAAGCCATAAAAACAAGACTCCGTACTGTGATCTTAGCTCCAACCAGGGTTGTCGCTGCTGAAATGGAGGAGGCCCTTAGAGGGCTTCCAGTGCGTTATATGACAACAGCAGTCAATGTCACCCACTCTGGAACAGAAATCGTCGACTTAATGTGCCATGCCACCTTCACTTCACGTCTACTACAGCCAATCAGAGTCCCCAACTATAATCTGTATATTATGGATGAGGCCCACTTCACAGATCCCTCAAGTATAGCAGCAAGAGGATACATTTCAACAAGGGTTGAGATGGGCGAGGCGGCTGCCATCTTCATGACCGCCACGCCACCAGGAACCCGTGACGCATTTCCGGACTCCAACTCACCAATTATGGACACCGAAGTGGAAGTCCCAGAGAGAGCCTGGAGCTCAGGCTTTGATTGGGTGACGGATCATTCTGGAAAAACAGTTTGGTTTGTTCCAAGCGTGAGGAACGGCAATGAGATCGCAGCTTGTCTGACAAAGGCTGGAAAACGGGTCATACAGCTCAGCAGAAAGACTTTTGAGACAGAGTTCCAGAAAACAAAACATCAAGAGTGGGACTTTGTCGTGACAACTGACATTTCAGAGATGGGCGCCAACTTTAAAGCTGACCGTGTCATAGATTCCAGGAGATGCCTAAAGCCGGTCATACTTGATGGCGAGAGAGTCATTCTGGCTGGACCCATGCCTGTCACACATGCCAGCGCTGCTCAGAGGAGGGGGCGCATAGGCAGGAATCCCAACAAACCTGGAGATGAGTATCTGTATGGAGGTGGGTGCGCAGAGACTGACGAAGACCATGCACACTGGCTTGAAGCAAGAATGCTCCTTGACAATATTTACCTCCAAGATGGCCTCATAGCCTCGCTCTATCGACCTGAGGCCGACAAAGTAGCAGCCATTGAGGGAGAGTTCAAGCTTAGGACGGAGCAAAGGAAGACCTTTGTGGAACTCATGAAAAGAGGAGATCTTCCTGTTTGGCTGGCCTATCAGGTTGCATCTGCCGGAATAACCTACACAGATAGAAGATGGTGCTTTGATGGCGCGACCAACAACACCATACTGGAAGACAGTGTGCCGGCAGAGGTGTGGACCAGACACGGAGAGAAAAGAGTGCTCAAACCGAGGTGGATGGACGCCAGAGTTTGTTCAGATCATGCGGCCCTGAAGTCATTCAAGGAGTTTGCCGCTGGGAAGAGAGGAGCGGCTTTTGGAGTGATGGAAGCCCTGGGAACACTGCCAGGACACATGACAGAGAGATTCCAGGAAGCCATTGACAACCTCGCTGTGCTCATGCGGGCAGAGACTGGAAGCAGGCCTTACAAAGCCGCGGCGGCCCAATTGCCGGAGACCCTAGAGACCATTATGCTTTTGGGGTTGCTGGGAACAGTCTCGCTGGGAATCTTTTTCGTCTTGATGAGGAACAAGGGCATAGGGAAGATGGGCTTTGGAATGGTGACCCTTGGGGCCAGTGCATGGCTCATGTGGCTCTCGGAAATTGAGCCAGCCAGAATTGCATGTGTCCTCATTGTTGTGTTCCTATTGCTGGTGGTGCTCATACCTGAGCCAGAAAAGCAAAGATCTCCCCAGGACAACCAAATGGCAATCATCATCATGGTAGCAGTAGGTCTTCTGGGCTTGATTACCGCCAATGAACTCGGATGGTTGGAGAGAACAAAGAGTGACCTAAGCCATCTAATGGGAAGGAGAGAGGAGGGGGCAACCATAGGATTCTCAATGGACATTGACCTGCGGCCAGCCTCAGCTTGGGCCATCTATGCTGCCTTGACAACTTTCATTACCCCAGCCGTCCAACATGCAGTGACCACTTCATACAACAACTACTCCTTAATGGCGATGGCCACGCAAGCTGGAGTGTTGTTTGGTATGGGCAAAGGGATGCCATTCTACGCATGGG-ACTTTGGAGTCCCGCTGCTAATGATAGGTTGCTACTCACAATTAACACCCCTGACCCTAATAGTGGCCATCATTTTGCTCGTGGCGCACTACATGTACTTGATCCCAGGGCTGCAGGCAGCAGCTGCGCGTGCTGCCCAGAAGAGAACGGCAGCTGGCATCATGAAGAACCCTGTTGTGGATGGAATAGTGGTGACTGACATTGACACAATGACAATTGACCCCCAAGTGGAGAAAAAGATGGGACAGGTGCTACTCATAGCAGTAGCCGTCTCCAGCGCCATACTGTCGCGGACCGCCTGGGGGTGGGGGGAGGCTGGGGCCCTGATCACAGCCGCAACTTCCACTTTGTGGGAAGGCTCTCCGAACAAGTACTGGAACTCCTCTACAGCCACTTCACTGTGTAACATTTTTAGGGGAAGTTACTTGGCTGGAGCTTCTCTAATCTACACAGTAACAAGAAACGCTGGCTTGGTCAAGAGACGTGGGGGTGGAACAGGAGAGACCCTGGGAGAGAAATGGAAGGCCCGCTTGAACCAGATGTCGGCCCTGGAGTTCTACTCCTACAAAAAGTCAGGCATCACCGAGGTGTGCAGAGAAGAGGCTCGCCGCGCCCTCAAGGACGGTGTGGCAACGGGAGGCCATGCTGTGTCCCGAGGAAGTGCAAAGCTGAGATGGTTGGTGGAGCGGGGATACCTGCAGCCCTATGGAAAGGTCATTGATCTTGGATGTGGCAGAGGGGGCTGGAGTTACTACGCCGCCACCATCCGCAAAGTTCAAGAAGTGAAAGGATACACAAAAGGAGGCCCTGGTCATGAAGAACCCGTGTTGGTGCAAAGCTATGGGTGGAACATAGTCCGTCTTAAGAGTGGGGTGGACGTCTTTCATATGGCGGCTGAGCCGTGTGACACGTTGCTGTGTGACATAGGTGAGTCATCATCTAGTCCTGAAGTGGAAGAAACACGGACGCTCAGAGTCCTCTCCATGGTGGGGGATTGGCTTGAAAAAAGACCAGGAGCCTTTTGTATAAAAGTGTTGTGCCCATACACCAGCACTATGATGGAAACCCTGGAGCGACTGCAGCGTAGGTATGGGGGAGGACTGGTCAGAGTGCCACTCTCCCGCAACTCTACACATGAGATGTACTGGGTCTCTGGAGCGAAAAGCAACACCATAAAAAGTGTGTCCACCACGAGCCAGCTCCTCTTGGGGCGCATGGACGGGCCTAGGAGGCCAGTGAAATATGAGGAGGATGTGAATCTCGGCTCTGGCACGCGGGCTGTGGTAAGCTGCGCTGAAGCTCCCAACATGAAGATCATTGGTAACCGCATTGAAAGGATCCGCAGTGAGCACGCGGAAACGTGGTTCTTTGACGAGAACCACCCATATAGGACATGGGCTTACCATGGAAGCTATGAGGCCCCCACACAAGGGTCAGCGTCCTCTCTAATAAACGGGGTTGTCAGGCTCCTGTCAAAACCCTGGGATGTGGTGACTGGAGTCACAGGAATAGCTATGACCGACACCACACCGTATGGTCAGCAAAGAGTTTTCAAGGAAAAAGTGGACACTAGGGTGCCAGACCCCCAAGAAGGCACTCGTCAGGTTATGAGCATGGTCTCTTCCTGGTTGTGGAAAGAGCTAGGCAAACACAAACGGCCACGAGTCTGTACCAAAGAAGAGTTCATCAACAAGGTTCGTAGCAATGCAGCATTAGGGGCAATATTTGAAGAGGAAAAAGAGTGGAAGACTGCAGTGGAAGCTGTGAACGATCCAAGGTTCTGGGCTCTAGTGGACAAGGAAAGAGAGCACCACCTGAGAGGAGAGTGCCAGAGTTGTGTGTACAACATGATGGGAAAAAGAGAAAAGAAACAAGGGGAATTTGGAAAGGCCAAGGGCAGCCGCGCCATCTGGTATATGTGGCTAGGGGCTAGATTTCTAGAGTTCGAAGCCCTTGGATTCTTGAACGAGGATCACTGGATGGGGAGAGAGAACTCAGGAGGTGGTGTTGAAGGGCTGGGATTACAAAGACTCGGATATGTCCTAGAAGAGATGAGTTGCATACCAGGAGGAAGGATGTATGCAGATGACACTGCTGGCTGGGACACCCGCATCAGCAGGTTTGATCTGGAGAATGAAGCTCTAATCACCAACCAAATGGAGAAAGGGCACAGGGCCTTGGCATTGGCCATAATCAAGTACACATACCAAAACAAAGTGGTAAAGGTCCTTAGACCAGCTGAAAAAGGGAAAACAGTTATGGACATTATTTCGAGACAAGACCAAAGGGGGAGCGGACAAGTTGTCACTTACGCTCTTAACACATTTACCAACCTAGTGGTGCAACTCATTCGGAATATGGAGGCTGAGGAAGTTCTAGAGATGCAAGACTTGTGGCTGCTGCGGAGGTCAGAGAAAGTGACCAACTGGTTGCAGAGCAACGGATGGGATAGGCTCAAACGAATGGCAGTCAGTGGAGATGATTGCGTTGTGAAGCCAATTGATGATAGGTTTGCACATGCCCTCAGGTTCTTGAATGATATGGGAAAAGTTAGGAAGGACACACAAGAGTGGAAACCCTCAACTGGATGGGACAACTGGGAAGAAGTTCCGTTTTGCTCCCACCACTTCAACAAGCTCCATCTCAAGGACGGGAGGTCCATTGTGGTTCCCTGCCGCCACCAAGATGAACTGATTGGCCGGGCCCGCGTCTCTCCAGGGGCGGGATGGAGCATCCGGGAGACTGCTTGCCTAGCAAAATCATATGCGCAAATGTGGCAGCTCCTTTATTTCCACAGAAGGGACCTCCGACTGATGGCCAATGCCATTTGTTCATCTGTGCCAGTTGACTGGGTTCCAACTGGGAGAACTACCTGGTCAATCCATGGAAAGGGAGAATGGATGACCACTGAAGACATGCTTGTGGTGTGGAACAGAGTGTGGATTGAGGAGAACGACCACATGGAAGACAAGACCCCAGTTACGAAATGGACAGACATTCCCTATTTGGGAAAAAGGGAAGACTTGTGGTGTGGATCTCTCATAGGGCACAGACCGCGCACCACCTGGGCTGAGAACATTAAAAACACAGTCAACATGGTGCGCAGGATCATAGGTGATGAAGAAAAGTACATGGACTACCTATCCACCCAAGTTCGCTACTTGGGTGAAGAAGGGTCTACACCTGGAGTGCTGTAA

>ENA|AMX81919|AMX81919.1 Zika virus polyprotein

ATGAAAAACCC-AAAAAAGAAATCCGGAGGATTCCGGATTGTCAATATGCTAAAACGCGGAGTAGCCCGTGTGAGCCCCTTTGGGGGCTTGAAGAGGCTGCCAGCCGGACTTCTGCTGGGTCATGGGCCCATCAGGATGGTCTTGGCGATTCTAGCCTTTTTGAGATTCACGGCAATCAAGCCATCACTGGGTCTCATCAATAGATGGGGTTCAGTGGGGAAAAAAGAGGCTATGGAAATAATAAAGAAGTTCAAGAAAGATCTGGCTGCCATGCTGAGAATAATCAATGCTAGGAAGGAGAAGAAGAGACGAGGCGCAGATACTAGTGTCGGAATTGTTGGCCTCCTGCTGACCACAGCTATGGCAGCGGAGGTCACTAGACGTGGGAGTGCATACTATATGTACTTGGACAGAAACGATGCTGGGGAGGCCATATCTTTTCCAACCACATTGGGGATGAATAAGTGTTATATACAGATCATGGATCTTGGACACATGTGTGATGCCACCATGAGCTATGAATGCCCTATGCTGGATGAGGGGGTGGAACCAGATGACGTCGATTGTTGGTGCAACACGACGTCAACTTGGGTTGTGTACGGAACCTGCCATCACAAAAAAGGTGAAGCACGGAGATCTAGAAGAGCTGTGACGCTCCCCTCCCATTCCACTAGGAAGCTGCAAACGCGGTCGCAAACCTGGTTGGAATCAAGAGAATACACAAAGCACTTGATTAGAGTCGAAAATTGGATATTCAGGAACCCTGGCTTCGCGTTAGCAGCAGCTGCCATCGCTTGGCTTTTGGGAAGCTCAACGAGCCAAAAAGTCATATACTTGGTCATGATACTGCTGATTGCCCCGGCATACAGCATCAGGTGCATAGGAGTCAGCAATAGGGACTTTGTGGAAGGTATGTCAGGTGGGACTTGGGTTGATGTTGTCTTGGAACATGGAGGTTGTGTCACCGTAATGGCACAGGACAAACCGACTGTCGACATAGAGCTGGTTACAACAACAGTCAGCAACATGGCGGAGGTAAGATCCTACTGCTATGAGGCATCAATATCAGACATGGCTTCGGACAGCCGCTGCCCAACACAAGGTGAAGCCTACCTTGACAAGCAATCAGACACTCAATATGTCTGCAAAAGAACGTTAGTGGACAGAGGCTGGGGAAATGGATGTGGACTTTTTGGCAAAGGGAGCCTGGTGACATGCGCTAAGTTTGCATGCTCCAAGAAAATGACCGGGAAGAGCATCCAGCCAGAGAATCTGGAGTACCGGATAATGCTGTCAGTTCATGGCTCCCAGCACAGTGGGATGATCGTTAATGACACAGGACATGAAACTGATGAGAATAGAGCGAAGGTTGAGATAACGCCCAATTCACCAAGAGCCGAAGCCACCCTGGGGGGTTTTGGAAGCCTAGGACTTGATTGTGAACCGAGGACAGGCCTTGACTTTTCAGATTTGTATTACTTGACTATGAATAACAAGCACTGGTTGGTCCACAAGGAGTGGTTCCACGACATTCCATTACCTTGGCACGCTGGGGCAGACACCGGAACTCCACACTGGAACAACAAAGAAGCACTGGTAGAGTTCAAGGACGCACATGCCAAAAGGCAAACTGTCGTGGTTCTAGGGAGTCAAGAAGGAGCAGTTCACACGGCCCTTGCTGGAGCTCTGGAGGCTGAGATGGATGGTGCAAAGGGAAGGCTGTCCTCTGGCCACTTGAAATGTCGCCTGAAAATGGATAAACTTAGATTGAAGGGCGTGTCATACTCCTTGTGTACCGCAGCGTTCACATTCACCAAGATCCCGGCTGAAACACTGCACGGGACAGTCACAGTGGAGGTACAGTACGCAGGGACAGATGGACCTTGCAAGGTTCCAGCTCAGATGGCGGTGGACATGCAAACTCTGACCCCAGTTGGGAGGTTGATAACCGCTAACCCCGTAATCACTGAAAGCACTGAGAACTCTAAGATGATGCTGGAACTTGATCCACCATTTGGGGACTCTTACATTGTCATAGGAGTCGGGGAGAAGAAGATCACCCACCACTGGCACAGGAGTGGCAGCACCATTGGAAAAGCATTTGAAGCCACTGTGAGAGGTGCCAAGAGAATGGCAGTCTTGGGAGACACAGCCTGGGACTTTGGATCAGTTGGAGGCGCTCTCAACTCATTGGGCAAGGGCATCCATCAAATTTTTGGAGCAGCTTTCAAATCATTGTTTGGAGGAATGTCCTGGTTCTCACAAATTCTCATTGGAACGTTGCTGATGTGGTTGGGTCTGAACACAAAGAATGGATCTATTTCCCTTATGTGCTTGGCCTTAGGGGGAGTGTTGATCTTCTTATCCACAGCCGTCTCTGCTGATGTGGGGTGCTCGGTGGACTTCTCAAAGAAGGAGACGAGATGCGGTACAGGGGTGTTCGTCTATAACGACGTTGAAGCCTGGAGGGACAGGTACAAGTACCATCCTGACTCCCCCCGTAGATTGGCAGCAGCAGTCAAGCAAGCCTGGGAAGATGGTATCTGCGGGATCTCCTCTGTTTCAAGAATGGAAAACATCATGTGGAGATCAGTAGAAGGGGAGCTCAACGCAATCTTGGAAGAGAATGGAGTTCAACTGACGGTCGTTGTGGGATCTGTAAAAAACCCCATGTGGAGAGGTCCACAGAGATTGCCCGTGCCTGTGAACGAGCTGCCCCACGGCTGGAAGGCTTGGGGGAAATCGCACTTCGTCAGAGCAGCAAAGACAAATAACAGCTTTGTCGTGGATGGTGACACACTGAAGGAATGCCCACTCAAACATAGAGCATGGAACAGCTTTCTTGTGGAGGATCATGGGTTCGGGGTATTTCACACTAGTGTCTGGCTCAAGGTTAGAGAAGATTATTCATTAGAGTGTGATCCAGCCGTTATTGGAACAGCTGTTAAGGGGAAGGAGGCTGTACACAGTGATCTAGGCTACTGGATTGAGAGTGAGAAGAATGACACATGGAGGCTGAAGAGGGCCCATCTGATCGAGATGAAAACATGTGAATGGCCAAAGTCCCACACATTGTGGACAGATGGAATAGAAGAGAGTGATCTGATCATACCCAAGTCTTTAGCTGGGCCACTCAGCCATCACAATACCAGAGAGGGCTACAGGACCCAAATGAAAGGGCCATGGCACAGTGAAGAGCTTGAAATTCGGTTTGAGGAATGCCCGGGCACTAAGGTCCACGTGGAGGAAACATGTGGAACAAGAGGACCATCTCTGAGATCAACCACTGCAAGCGGAAGGGTGATCGAGGAATGGTGCTGCAGGGAGTGCACAATGCCCCCACTGTCGTTCCGGGCTAAAGATGGCTGTTGGTATGGAATGGAGATAAGGCCCAGGAAAGAACCAGAAAGCAACTTAGTAAGGTCAATGGTGACTGCAGGATCAACTGATCACATGGATCACTTCTCCCTTGGAGTGCTTGTGATTCTGCTCATGGTGCAGGAAGGGCTGAAGAAGAGAATGACCACAAAGATCATCATAAGCACATCAATGGCAGTGCTGGTAGCTATGATCCTGGGAGGATTTTCAATGAGTGACCTGGCTAAGCTTGCAATTTTGATGGGCGCCACCTTCGCGGAAATGAACACTGGAGGAGATGTAGCTCATCTGGCGCTGATAGCGGCATTCAAAGTCAGACCAGCGTTGCTGGTATCTTTCATCTTCAGAGCTAATTGGACACCCCGTGAAAGCATGCTGCTGGCCTTGGCCTCGTGTCTTTTGCAAACTGCGATCTCCGCCTTGGAAGGCGACCTGATGGTTCTCATCAATGGTTTTGCTTTGGCCTGGTTGGCAATACGAGCGATGGTTGTTCCACGCACTGATAACATCACCTTGGCAATCCTGGCTGCTCTGACACCACTGGCCCGGGGCACACTGCTTGTGGCGTGGAGAGCAGGCCTTGCTACTTGCGGGGGGTTTATGCTCCTCTCTCTGAAGGGAAAAGGCAGTGTGAAGAAGAACTTACCATTTGTCATGGCCCTGGGACTAACCGCTGTGAGGCTGGTCGACCCCATCAACGTGGTGGGGCTGCTGTTGCTCACAAGGAGTGGGAAGCGGAGCTGGCCCCCTAGCGAAGTACTCACAGCTGTTGGCCTGATATGCGCATTGGCTGGAGGGTTCGCCAAGGCAGATATAGAGATGGCTGGGCCCATGGCCGCGGTCGGTCTGCTAATTGTCAGTTACGTGGTCTCAGGAAAGAGTGTGGACATGTACATTGAAAGAGCAGGTGACATCACATGGGAAAAAGATGCGGAAGTCACTGGAAACAGTCCCCGGCTCGATGTGGCGCTAGATGAGAGTGGTGATTTCTCCCTGGTGGAGGATGACGGTCCCCCCATGAGAGAGATCATACTCAAGGTGGTCCTGATGACCATCTGTGGCATGAACCCAATAGCCATACCCTTTGCAGCTGGAGCGTGGTACGTATACGTGAAGACTGGAAAAAGGAGTGGTGCTCTATGGGATGTGCCTGCTCCCAAGGAAGTAAAAAAGGGGGAGACCACAGATGGAGTGTACAGAGTAATGACTCGTAGACTGCTAGGTTCAACACAAGTTGGAGTGGGAGTTATGCAAGAGGGGGTCTTTCACACTATGTGGCACGTCACAAAAGGATCCGCGCTGAGAAGCGGTGAAGGGAGACTTGATCCATACTGGGGAGATGTCAAGCAGGATCTGGTGTCATACTGTGGTCCATGGAAGCTAGATGCCGCCTGGGACGGGCACAGCGAGGTGCAGCTCTTGGCCGTGCCCCCCGGAGAGAGAGCGAGGAACATCCAGACTCTGCCCGGAATATTTAAGACAAAGGATGGGGACATTGGAGCGGTTGCGCTGGATTACCCAGCAGGAACTTCAGGATCTCCAATCCTAGACAAGTGTGGGAGAGTGATAGGACTTTATGGCAATGGGGTCGTGATCAAAAATGGGAGTTATGTTAGTGCCATCACCCAAGGGAGGAGGGAGGAAGAGACTCCTGTTGAGTGCTTCGAGCCTTCGATGCTGAAGAAGAAGCAGCTAACTGTCTTAGACTTGCATCCTGGAGCTGGGAAAACCAGGAGAGTTCTTCCTGAAATAGTCCGTGAAGCCATAAAAACAAGACTCCGTACTGTGATCTTAGCTCCAACCAGGGTTGTCGCTGCTGAAATGGAGGAAGCCCTTAGAGGGCTTCCAGTGCGTTATATGACAACAGCAGTCAATGTCACCCACTCTGGAACAGAAATCGTCGACTTAATGTGCCATGCCACCTTCACTTCACGTCTACTACAGCCAATCAGAGTCCCCAACTATAATCTGTATATTATGGATGAGGCCCACTTCACAGATCCCTCAAGTATAGCAGCAAGAGGATACATTTCAACAAGGGTTGAGATGGGCGAGGCGGCTGCCATCTTCATGACCGCCACGCCACCAGGAACCCGTGACGCATTTCCGGACTCCAACTCACCAATTATGGACACCGAAGTGGAAGTCCCAGAGAGAGCCTGGAGCTCAGGCTTTGATTGGGTGACGGATTATTCTGGAAAAACAGTTTGGTTTGTTCCAAGCGTGAGGAACGGCAATGAGATCGCAGCTTGTCTGACAAAGGCTGGAAAACGGGTCATACAGCTCAGCAGAAAGACTTTTGAGACAGAGTTCCAGAAAACAAAACATCAAGAGTGGGACTTTGTCGTGACAACTGACATTTCAGAGATGGGCGCCAACTTTAAAGCTGACCGTGTCATAGATTCCAGGAGATGCCTAAAGCCGGTCATACTTGATGGCGAGAGAGTCATTTTGGCTGGACCCATGCCTGTCACACATGCCAGCGCTGCCCAGAGGAGGGGGCGCATAGGCAGGAATCCCAACAAACCTGGAGATGAGTATCTGTATGGAGGTGGGTGCGCAGAGACTGACGAAGACCATGCACACTGGCTTGAAGCAAGAATGCTCCTTGACAATATTTACCTCCAAGATGGCCTCATAGCCTCGCTCTATCGACCTGAGGCCGACAAAGTAGCAGCCATTGAGGGAGAGTTCAAGCTTAGGACGGAGCAAAGGAAGACCTTTGTGGAACTCATGAAAAGAGGAGATCTTCCTGTTTGGCTGGCCTATCAGGTTGCATCTGCCGGAATAACCTACACAGATAGAAGATGGTGCTTTGATGGCACGACCAACAACACCATAATGGAAGACAGTGTGCCGGCAGAGGTGTGGACCAGACACGGAGAGAAAAGAGTGCTCAAACCGAGGTGGATGGACGCCAGAGTTTGTTCAGATCATGCGGCCCTGAAGTCATTCAAGGAGTTTGCCGCTGGGAAAAGAGGAGCGGCTTTTGGAGTGATGGAAGCCCTGGGAACACTGCCAGGACACATGACAGAGAGATTCCAGGAAGCCATTGACAACCTCGCTGTGCTCATGCGGGCAGAGACTGGAAGCAGGCCTTACAAAGCCGCGGCGGCCCAATTGCCGGAGACCCTAGAGACCATTATGCTTTTGGGGTTGCTGGGAACAGTCTCGCTGGGAATCTTTTTCGTCTTGATGAGGAACAAGGGCATAGGGAAGATGGGCTTTGGAATGGTGACTCTTGGGGCCAGCGCATGGCTCATGTGGCTCTCGGAAATTGAGCCAGCCAGAATTGCATGTGTCCTCATTGTTGTGTTCCTATTGCTGGTGGTGCTCATACCTGAGCCAGAAAAGCAAAGATCTCCCCAGGACAACCAAATGGCAATCATCATCATGGTAGCAGTAGGTCTTCTGGGCTTGATTACCGCCAATGAACTCGGATGGTTGGAGAGAACAAAGAGTGACCTAAGCCATCTAATGGGAAGGAGAGAGGAGGGGGCAACCATGGGATTCTCAATGGACATTGACCTGCGGCCAGCCTCAGCTTGGGCCATCTATGCTGCCTTGACAACTTTCATTACCCCAGCCGTCCAACATGCAGTGACCACTTCATACAACAACTACTCCTTAATGGCGATGGCCACGCAAGCTGGAGTGTTGTTTGGTATGGGCAAAGGGATGCCATTCTACGCATGGG-ACTTTGGAGTCCCGCTGCTAATGATAGGTTGCTACTCACAATTAACGCCCCTGACCCTAATAGTGGCCATCATTTTGCTCGTGGCGCACTACATGTACTTGATCCCAGGGCTGCAGGCAGCAGCTGCGCGTGCTGCCCAGAAGAGAACGGCAGCTGGCATCATGAAGAACCCTGTTGTGGATGGAATAGTGGTGACTGACATTGACACAATGACAATTGACCCCCAAGTGGAGAAAAAGATGGGACAGGTGCTACTCATAGCAGTAGCAGTCTCCAGCGCCATACTGTCGCGGACCGCCTGGGGGTGGGGGGAGGCTGGGGCCCTGATCACAGCCGCAACTTCCACTTTGTGGGAAGGCTCTCCGAACAAGTACTGGAACTCCTCTACAGCCACTTCACTGTGTAACATTTTTAGGGGAAGTTACTTGGCTGGAGCTTCTCTAATCTACACAGTAACAAGAAACGCTGGCTTGGTCAAGAGACGTGGGGGTGGAACAGGAGAGACCCTGGGAGAGAAATGGAAGGCCCGCTTGAACCAGATGTCGGCCCTGGAGTTCTACTCCTACAAAAAGTCAGGCATCACCGAGGTGTGCAGAGAAGAGGCCCGCCGCGCCCTCAAGGACGGTGTGGCAACGGGAGGCCATGCTGTGTCCCGAGGAAGTGCAAAGCTGAGATGGTTGGTGGAGCGGGGATACCTGCAGCCCTATGGAAAGGTCATTGATCTTGGATGTGGCAGAGGGGGCTGGAGTTACTACGCCGCCACCATCCGCAAAGTTCAAGAAGTGAAAGGATACACAAAAGGAGGCCCTGGTCATGAAGAACCCGTGTTGGTGCAAAGCTATGGGTGGAACATAGTCCGTCTTAAGAGTGGGGTGGACGTCTTTCATATGGCGGCTGAGCCGTGTGACACGTTGCTGTGTGACATAGGTGAGTCATCATCTAGTCCTGAAGTGGAAGAAGCACGGACGCTCAGAGTCCTCTCCATGGTGGGGGATTGGCTTGAAAAAAGACCAGGAGCCTTTTGTATAAAAGTGTTGTGCCCATACACCAGCACTATGATGGAAACCCTGGAGCGACTGCAGCGTAGGTATGGGGGAGGACTGGTCAGAGTGCCACTCTCCCGCAACTCTACACATGAGATGTACTGGGTCTCTGGAGCGAAAAGCAACACCATAAAAAGTGTGTCCACCACGAGCCAGCTCCTCTTGGGGCGCATGGACGGGCCTAGGAGGCCAGTGAAATATGAGGAGGATGTGAATCTCGGCTCTGGCACGCGGGCTGTGGTAAGCTGCGCTGAAGCTCCCAACATGAAGATCATTGGTAACCGCATTGAAAGGATCCGCAGTGAGCACGCGGAAACGTGGTTCTTTGACGAGAACCACCCATATAGGACATGGGCTTACCATGGAAGCTATGAGGCCCCCACACAAGGGTCAGCGTCCTCTCTAATAAACGGGGTTGTCAGGCTCCTGTCAAAACCCTGGGATGTGGTGACTGGAGTCACAGGAATAGCCATGACCGACACCACACCGTATGGTCAGCAAAGAGTTTTCAAGGAAAAAGTGGACACTAGGGTGCCAGACCCCCAAGAAGGCACTCGTCAGGTTATGAGCATGGTCTCTTCCTGGTTGTGGAAAGAGCTAGGCAAACACAAACGGCCACGAGTCTGTACCAAAGAAGAGTTCATCAACAAGGTTCGTAGCAATGCAGCATTAGGGGCAATATTTGAAGAGGAAAAAGAGTGGAAGACTGCAGTGGAAGCTGTGAACGATCCAAGGTTCTGGGCTCTAGTGGACAAGGAAAGAGAGCACCACCTGAGAGGAGAGTGCCAGAGTTGTGTGTACAACATGATGGGAAAAAGAGAAAAGAAACAAGGGGAATTTGGAAAGGCCAAGGGCAGCCGCGCCATCTGGTATATGTGGCTAGGGGCTAGATTTCTAGAGTTCGAAGCCCTTGGATTCTTGAACGAGGATCACTGGATGGGGAGAGAGAACTCAGGAGGTGGTGTTGAAGGGCTGGGATTACAAAGACTCGGATATGTCCTAGAAGAGATGAGTCGCATACCAGGAGGAAGGATGTATGCAGATGACACTGCTGGCTGGGACACCCGCATCAGCAGGTTTGATCTGGAGAATGAAGCTCTAATCACCAACCAAATGGAGAAAGGGCACAGGGCCTTGGCATTGGCCATAATCAAGTACACATACCAAAACAAAGTGGTAAAGGTCCTTAGACCAGCTGAAAAAGGGAAGACAGTTATGGACATTATTTCGAGACAAGACCAAAGGGGGAGCGGACAAGTTGTCACTTACGCTCTTAACACATTTACCAACCTAGTGGTGCAACTCATTCGGAATATGGAGGCTGAGGAAGTTCTAGAGATGCAAGACTTGTGGCTGCTGCGGAGGTCAGAGAAAGTGACCAACTGGTTGCAGAGCAACGGATGGGATAGGCTCAAACGAATGGCAGTCAGTGGAGATGATTGCGTTGTGAAGCCAATTGATGATAGGTTTGCACATGCCCTCAGGTTCTTGAATGATATGGGAAAAGTTAGGAAGGACACACAAGAGTGGAAACCCTCAACTGGATGGGACAACTGGGAAGAAGTTCCGTTTTGCTCCCACCACTTCAACAAGCTCCATCTCAAGGACGGGAGGTCCATTGTGGTTCCCTGCCGCCACCAAGATGAACTGATTGGCCGGGCCCGCGTCTCTCCAGGGGCGGGATGGAGCATCCGGGAGACTGCTTGCCTAGCAAAATCATATGCGCAAATGTGGCAGCTCCTTTATTTCCACAGAAGGGACCTCCGACTGATGGCCAATGCCATTTGTTCATCTGTGCCAGTTGACTGGGTTCCAACTGGGAGAACTACCTGGTCAATCCATGGAAAGGGAGAATGGATGACCACTGAAGACATGCTTGTGGTGTGGAACAGAGTGTGGATTGAGGAGAACGACCACATGGAAGACAAGACCCCAGTTACGAAATGGACAGACATTCCCTATTTGGGAAAAAGGGAAGACTTGTGGTGTGGATCTCTCATAGGGCACAGACCGCGCACCACCTGGGCTGAGAACATTAAAAACACAGTCAACATGGTGCGCAGGATCATAGGTGATGAAGAAAAGTACATGGACTACCTATCCACCCAAGTTCGCTACTTGGGTGAAGAAGGGTCTACACCTGGAGTGCTGTAA
