## Supplementary file 2 for "Zika viral proteome analysis reveals an epitope cluster within NS3 helicase as a potential vaccine candidate: an *in silico* study"

| Uniprot ID | Collection Date | Country | Strain | EMBL Gene ID |
| --- | --- | --- | --- | --- |
| A0A125RTB5 | 19-Jul-2014 | Thailand | Zika virus/H.sapiens-tc/THA/2014/SV0127- 14 | AMD61710.3 |
| A0A125RTB6 | 09-May-2012 | Philippines | Zika virus/H.sapiens-tc/PHL/2012/CPC-0740 | AMD61711.3 |
| A0A024B7W1 | 28-Nov-2013 | French Polynesia | H/PF/2013 | AHZ13508.1 |
| A0A109PRQ3 | 2015 | Brazil | BeH819015 | AMA12085.1 |
| B3U3M3 | Jun-2007 | Micronesia |  | ACD75819.1 |
| A0A0X9QZM7 | 2015 | Brazil | BeH818995 | AMA12084.1 |
| A0A120IF34 | 01-Dec-2015 | Guatemala | 103344 | AMC13912.1 |
| A0A110ASP6 | 12-Dec-2014 | Haiti | Haiti/1225/2014 | AMB37295.3 |
| A0A0X8GJ44 | Dec-2015 | Martinique | MRS_OPY_Martinique_PaRi_2015 | AMC33116.1 |
| A0A0X1LH65 | 01-Dec-2015 | Puerto Rico | PRVABC59 | AMC13911.1 |
| W8Q6T3 |  |  | ArB1362 | AHL43500.1 |
| A0A0U4DG08 | 02-Oct-2015 | Suriname | Z1106033 | ALX35659.1 |
| W8Q7Q3 |  |  | ArD158084 | AHL43504.1 |
| A0A109ZYP8 | 2015 | Brazil:Rio Grande do Norte, Natal | Natal RGN | AMB18850.1 |
| W8QIT7 |  |  | ArD128000 | AHL43502.1 |
| A0A090BCI3 |  | Uganda | MR766-NIID | BAP47441.1 |
| W8R1T1 |  |  | ArD7117 | AHL43501.1 |
| A0A0U3FSM8 | Mar-2015 | Brazil | ZikaSPH2015 | ALU33341.1 |
| H9A910 | 2010 | Cambodia | FSS13025 | AFD30972.1 |
| H8XX10 | 1947 | Uganda | MR_766 | AEN75263.1 |
| H8XX13 | 1984 | Senegal | ArD_41519 | AEN75266.1 |
| H8XX11 | 1966 | Malaysia | P6-740 | AEN75264.1 |
| W8QFH2 |  |  | ArD157995 | AHL43503.1 |
| A0A140FB25 | 12-Feb-2016 | China | GDZ16001 | AML82110.1 |
| A0A127AMC3 | 17-Feb-2016 | China | ZJ03 | AMM39806.2 |
| A0A140GMI4 | 01-Feb-2016 | Italy:Padua | Dominican Republic/2016/PD2 | AMN14620.1 |
| A0A140CFV0 | 30-Nov-2015 | Brazil | Brazil-ZKV2015 | AMD16557.1 |
| A0A140DLX5 | 2015 | Brazil | BeH828305 | AMK49165.1 |
| A0A140GMI3 | 01-Feb-2016 | Italy:Padua | Dominican Republic/2016/PD1 | AMN14619.1 |
| A0A140D2T1 | 1947 | Uganda | MR 766 | AMK02027.1 |
| A0A140DLX4 | 2015 | Brazil | BeH823339 | AMK49164.2 |
| A0A140E7U5 | 06-Feb-2016 | China | VE_Ganxian | AMK79469.1 |
| A0A127AM58 | Dec-2015 | Colombia:Barranquilla | FLR | AMM39804.2 |
| A0A146CJG7 | 25-Feb-2016 | China | Zika virus/GZ02/2016 | AMX81918.1 |
| A0A142I5B9 | 2010 | Cambodia | Zika virus/H.sapiens-tc/KHM/2010/FSS13025 | AMR39834.1 |
| A0A142I5C1 | 14-Dec-1984 | Senegal | Zika virus/A.taylori-tc/SEN/1984/41671-DAK | AMR39836.1 |
| A0A142I5B8 | 06-Dec-1984 | Senegal | Zika virus/A.taylori-tc/SEN/1984/41662-DAK | AMR39833.1 |
| A0A142I5B7 | 20-Nov-1984 | Senegal | Zika virus/A.africanus-tc/SEN/1984/41525-DAK | AMR39832.1 |
| A0A141ZVF6 | 2016 | China:Shenzhen | Zika virus/SZ01/2016 | AMO03410.1 |
| A0A142EHG8 | 14-Jan-2016 | Brazil | Rio-U1 | AMQ48981.1 |
| A0A142FH79 | 01-Jul-2015 | Brazil | Bahia07 | AMQ76465.1 |
| A0A142EHG9 | 29-Jan-2016 | Brazil | Rio-S1 | AMQ48982.1 |
| A0A142DS37 | 25-Feb-2016 | Mexico:Chiapas | MEX/InDRE/Lm/2016 | AMQ34003.1 |
| A0A141VLG5 | 14-Feb-2016 | China | GZ01 | AMM39805.1 |
| A0A142K469 | 06-Mar-2016 | Italy:Rome | Brazil/2016/INMI1 | AMS00611.1 |
| A0A142DS38 | 25-Feb-2016 | Mexico:Chiapas | MEX/InDRE/Sm/2016 | AMQ34004.1 |
| A0A142EHH1 | 02-Feb-2016 | USA | FB-GWUH-2016 | AMQ48986.1 |
| A0A146CJH9 | 05-Feb-2016 | USA | Haiti/1/2016 | AMX81919.1 |

**Table 01:** Sequences with their date, region, strain and Gene ID

**Algorithm used for MHC I and II**

import csv

f = open("test.csv")

fw = open("result.csv",'w')

fieldnames = ['Start Point', 'Lowest IC50', "Unique Allele"]

writer = csv.DictWriter(fw, fieldnames=fieldnames)

writer.writeheader()

csvF = csv.DictReader(f)

rowList = []

for row in csvF:

t = (row['allele'],row['start'],row['end'],row['mhc_ic50'])

\# print(t)

rowList.append(t)

num1 = 0

print(num1)

num2 = rowList[len(rowList)-1]

num2=int(num2[2])

print(num2)

resultRange = list(range(num1,num2+1))

print(resultRange)

for i in resultRange:

allele = []

ic50 = []

empty = 0

for j in rowList:

start = int(j[1])

end = int(j[2])

if i>=start and i<=end:

ic50.append(int(j[3]))

if allele.count(j[0])==0:

allele.append(j[0])

else:

empty=1

if len(ic50) != 0:

print(str(i)+" "+str(min(ic50))+" " +str(len(allele))+"\n")

writer.writerow({'Start Point':i, "Lowest IC50":min(ic50), "Unique Allele":len(allele)})

else:

writer.writerow({'Start Point': i, "Lowest IC50": 0, "Unique Allele": 0})

f.close()

fw.close()

**Table 02:** *in-house* algorithm to calculate the position-specific unique allele count and IC_50_ score of each amino acid. The algorithm calculates the lowest IC_50_ score and the number of unique alleles only for a given amino acid at a specific position.

| **Epitope/Ligand** | **HLA/Receptor** | **Center** | | | **Size** | | |
| --- | --- | --- | --- | --- | --- | --- | --- |
|  |  | **X axis** | **Y axis** | **Z axis** | **X axis** | **Y axis** | **Z axis** |
| ARMLLDNIY | HLA-A*30:02 | 3.122 | -0.223 | 57.609 | 72 | 54 | 66 |
| RMLLDNIYL | HLA-A*02:06 | -6.03 | -9.999 | 3.295 | 76 | 88 | 88 |
| MLLDNIYLQ | HLA-A*02:06 | -3.547 | -9.28 | 1.427 | 62 | 92 | 90 |
| YLQDGLIAS | HLA-A*02:06 | -8.002 | -5.689 | 1.179 | 62 | 74 | 84 |
| LQDGLIASL | HLA-A*02:06 | -3.968 | -9.142 | 1.711 | 66 | 94 | 92 |
| QDGLIASLY | HLA-A*29:02 | -7.968 | -10.994 | 0.327 | 96 | 68 | 88 |

**Table 03:** Docking properties of MHC I epitopes with their corresponding HLAs.

| **Epitope/Ligand** | **HLA/Receptor** | **Center** | | | **Size** | | |
| --- | --- | --- | --- | --- | --- | --- | --- |
|  |  | **X axis** | **Y axis** | **Z axis** | **X axis** | **Y axis** | **Z axis** |
| WLEARMLLD | HLA-DRB1*04:05 | -32.214 | 19.861 | -20.822 | 42 | 68 | 66 |
| EARMLLDNI | HLA-DRB4*01:01 | -34.276 | 19.925 | -20.526 | 56 | 54 | 68 |
| ARMLLDNIY | HLA-DRB1*04:05 | -34.214 | 19.861 | -20.822 | 72 | 50 | 84 |
| RMLLDNIYL | HLA-DRB1*01:01 | 4.905 | 7.923 | 71.276 | 94 | 90 | 96 |
| MLLDNIYLQ | HLA-DRB1*03:01 | -34.089 | 19.924 | -20.754 | 72 | 76 | 82 |
| LLDNIYLQD | HLA-DRB1*04:05 | -32.797 | 20.664 | -20.415 | 62 | 56 | 72 |
| LDNIYLQDG | HLA-DRB4*01:01 | -33.355 | 19.086 | -21.389 | 86 | 58 | 68 |
| DNIYLQDGL | HLA-DRB4*01:01 | -34.276 | 19.925 | -20.526 | 78 | 52 | 82 |
| NIYLQDGLI | HLA-DRB1*01:01 | 0.691 | 11.175 | 75.56 | 106 | 106 | 94 |
| IYLQDGLIA | HLA-DRB1*04:05 | -34.214 | 19.861 | -20.822 | 78 | 54 | 74 |
| YLQDGLIAS | HLA-DRB1*04:04 | -34.201 | 19.822 | -20.809 | 74 | 66 | 56 |
| LQDGLIASL | HLA-DRB1*09:01 | -34.932 | 19.325 | -21.406 | 68 | 54 | 82 |
| QDGLIASLY | HLA-DRB1*01:01 | -1.816 | 10.676 | 76.476 | 70 | 122 | 110 |
| DGLIASLYR | HLA-DRB1*15:01 | -34.212 | 19.791 | -20.753 | 64 | 68 | 74 |

**Table 04:** Docking properties of MHC II epitopes with their corresponding HLAs.

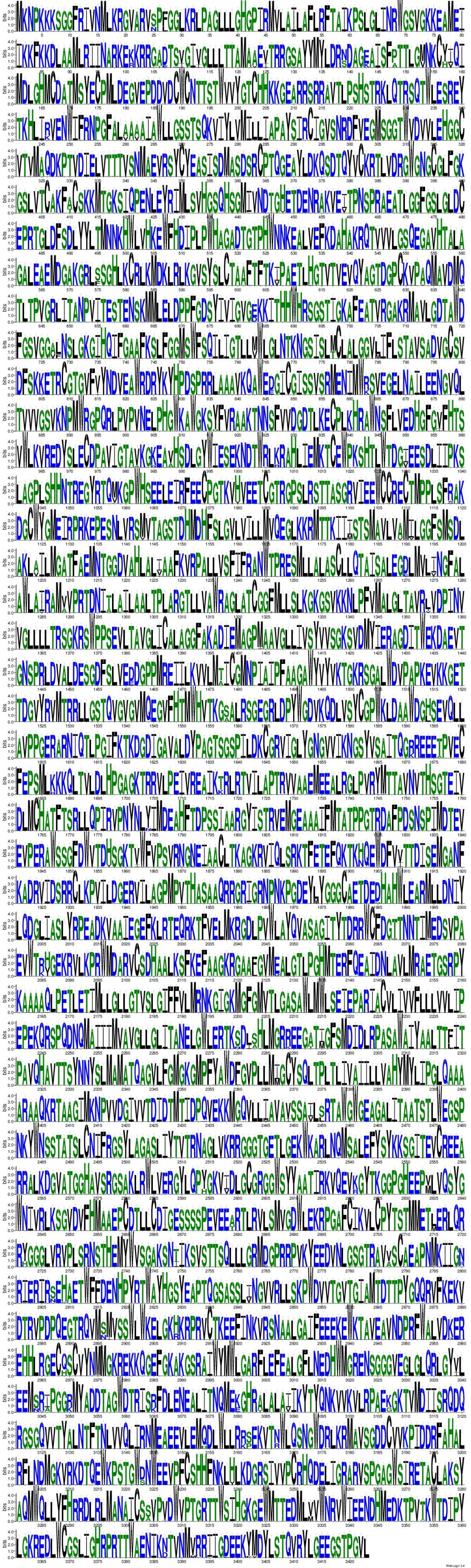

**Figure 01:** Weblogo
